# Modeling Heterogeneity of Triple-Negative Breast Cancer Uncovers a Novel Combinatorial Treatment Overcoming Primary Drug Resistance

**DOI:** 10.1101/2020.06.22.164418

**Authors:** Fabienne Lamballe, Fahmida Ahmad, Yaron Vinik, Olivier Castellanet, Fabrice Daian, Anna-Katharina Müller, Ulrike A. Köhler, Anne-Laure Bailly, Emmanuelle Josselin, Rémy Castellano, Christelle Cayrou, Emmanuelle Charafe-Jauffret, Gordon B. Mills, Vincent Géli, Jean-Paul Borg, Sima Lev, Flavio Maina

## Abstract

Triple-negative breast cancer (TNBC) is a highly aggressive breast cancer subtype characterized by a remarkable molecular heterogeneity. Currently, there are no effective druggable targets and advanced preclinical models of the human disease. Here, we generated a unique mouse model (*MMTV-R26*^*Met*^ mice) of mammary tumors driven by a subtle increase in the expression of the wild-type MET receptor. *MMTV-R26*^*Met*^ mice develop spontaneous, exclusive TNBC tumors, recapitulating primary resistance to treatment of patients. Proteomic profiling of *MMTV-R26*^*Met*^ tumors and machine learning approach showed that the model faithfully recapitulates inter-tumoral heterogeneity of human TNBC. Further signaling network analysis highlighted potential druggable targets, of which co-targeting of WEE1 and BCL-XL synergistically killed TNBC cells and efficiently induced tumor regression. Mechanistically, BCL-XL inhibition exacerbates the dependency of TNBC cells on WEE1 function, leading to Histone H3 and phosphoS_33_RPA32 upregulation, RRM2 downregulation, cell cycle perturbation, mitotic catastrophe and apoptosis. Our study introduces a unique, powerful mouse model for studying TNBC formation and evolution, its heterogeneity, and for identifying efficient therapeutic targets.

## 1. Introduction

Genetically engineered mouse models (GEMMs) of breast cancer have been proven as a powerful tool for gaining mechanistic insights into tumor initiation, progression, and metastasis as well as for developing innovative cancer therapy.^[1]^ GEMM models for breast cancer commonly use mammary-gland specific promoters, including MMTV (virus long terminal repeat), WAP (whey acidic protein), and C3 to ensure expression of transgenes in the mammary epithelium. More than 25 different murine GEMMs for breast cancer expressing different genes/oncogenes such as, *PyMT* (polyoma middle T antigen), SV40 T antigen, *ErbB2/Neu*, *cyclinD1*, *Ras*, *Myc*, *TGF-α*, and *Wnt1* have been established.^[2]^ The most widely used models are *MMTV-Neu* and *MMTV-PyMT*, which result in the development of multifocal adenocarcinoma and metastatic lesions in the lungs and/or lymph nodes. *MMTV-Neu* mice have been used for modelling epidermal growth factor receptor 2 (HER2)-positive breast cancer, whereas *MMTV-CyclinD1* for estrogen receptor (ER)-positive breast cancer. *MMTV-PyMT* mice lose the expression of ERα and progesterone receptor (PR) as they progress and concomitantly gain androgen receptor (AR) expression, therefore could be used for modelling luminal AR positive TNBC (LAR).^[3]^ However, only a small fraction of TNBC patients (~15%) are positive for AR, while the majority have been classified into different molecular subtypes, including basal-like (BL1 and BL2) and mesenchymal (M).^[4, 5]^

TNBC, which accounts for approximately 10-15% of all breast cancer patients, is defined by the lack of ER and PR expression, as well as HER2 amplification/overexpression. Compared to the other breast cancer subtypes, TNBC is characterized by the earliest age of onset, a high propensity for metastasis, and the worst prognosis in terms of relapse and survival rate.^[6–8]^ Over 80% of TNBC patients exhibit alterations in the *TP53* locus,^[9]^ whereas a smaller fraction has mutations in genes controlling the PI3K pathway and homologous recombination including BRCA1/2 negative. A molecular feature of TNBC is the dependency of cancer cells on signals that are rarely mutated, a phenomenon defined as “non-oncogene addiction”.^[10]^ Collectively, these traits are among the leading cause of limited efficacy of current TNBC therapies. Radiation therapy and chemotherapy, applied before and after surgery, are the mainstay of treatment, although frequently associated with drug resistance and recurrent disease.^[6–8]^

Extensive efforts have been made to search for molecular targeted therapies effective for TNBC treatment. Although some targeted therapies approved for treatment of other cancer types have been proposed in TNBC, they rarely turned out to be clinically relevant.^[11]^ These limited responses are associated with the high heterogeneity of the disease and the lack of suitable immunocompetent preclinical models that recapitulate the molecular diversity of TNBC. Among potential targets for TNBC subsets are PARP1, androgen receptor (AR), vascular endothelial growth factor receptor (VEGFR), epidermal growth factor receptor (EGFR), MET, PI3K/mTOR, MEK, Cyclin-dependent kinases (CDKs), heat shock protein 90 (HSP90), histone deacetylase (HDAC), hypoxia-inducible factor 1-α (HIF1-α), and integrins.^[11, 12]^ Inhibition of WEE1 kinase has been proposed as a promising treatment option for TNBC and several other types of solid cancer.^[13, 14]^ WEE1 plays central role in the G2/M checkpoint and controls DNA synthesis as part of the S phase checkpoint. Therefore, inhibition of WEE1 is associated with accumulation of DNA damage and aberrant mitosis. Co-inhibition of WEE1 with either radiotherapy or anticancer drugs such as cisplatin, gemcitabin, paclitaxel, or inhibitors of CDC25, ATR, or PARP causes death of breast cancer cells.^[15–22]^ The rational of these combined treatments is to associate DNA-damaging therapies together with perturbation of DNA damage checkpoint gatekeepers through WEE1 targeting. Nevertheless, the consequences of WEE1 targeting may be broader than cell cycle regulation, in view of recent studies showing that WEE1 inactivation increases CDK-dependent firing of dormant replication origins thereby leading to replication stress and increased dNTP demand.^[23, 24]^ Moreover, WEE1 was reported to modulate Histone H2B phosphorylation to inhibit transcription of several histone genes in yeast and humans and to function as a histone-sensing checkpoint in budding yeast.^[25, 26]^

Here, we report the generation of a unique mouse model (*MMTV-R26*^*Met*^ mice) in which a subtle increase in the expression levels of the wild-type MET receptor tyrosine kinase (RTK) leads to spontaneous TNBC formation. The tumorigenic switch correlated with a critical threshold of MET expression, whereas aggressiveness was associated with high MET levels and discrete signaling reprogramming. Proteomic profiling, signaling network analysis, and machine learning indicated that the *MMTV-R26*^*Met*^ mice not only model different tumorigenic stages of TNBC, but also largely recapitulates heterogeneity of the human disease as well as primary resistance to treatment. We used this unique model to identify potential therapeutic targets for TNBC through signaling reprogramming analysis and provide strong evidence that combination treatment with BCL-XL and WEE1 inhibitors could be a promising therapeutic approach with high clinical impact.

## 2. Results

### 2.1. Enhanced wild-type RTK MET expression levels in the mouse mammary gland induce spontaneous TNBC development

Previous studies showed that expression of oncogenic MET led to the development of diverse mammary tumors with basal characteristics.^[27]^ We assessed the sensitivity of the mammary gland to slighty increased wild-type MET levels by crossing the *MMTV-Cre* transgenic with *R26*^*stopMet*^ mice (referred to as *MMTV-R26*^*Met*^). The specificity of the LacZ-stop cassette deletion obtained by the *MMTV-Cre* mice was evaluated using the *R26*^*stopMet-Luc*^ mice,^[28]^ in which *Met*^*tg*^ is followed by an internal ribosome entry site-Luciferase reporter (Figure 1A). In vivo imaging of female *MMTV-R26*^*Met-Luc*^ mice revealed a strong luciferase signal in mammary glands only after the first lactation (Figure 1B), consistent with the expression of the Cre recombinase following MMTV promoter activation by prolactin.^[29, 30]^ This led to removal of the stop cassette, and thus *Met*^*tg*^ expression in the mammary gland of *MMTV-R26*^*Met*^ mice (Figure 1A). Consistently, the Luciferase-positive domains further increased after the second lactation and were significantly reduced in post-lactating females, in agreement with involution of the mammary gland occurring when the lactation phase is over (Figure 1B). This imaging analysis exemplifies the remodeling of the mammary gland overtime. In view of a dynamic regulation of the HGF/MET system in mammary gland morphogenesis previously reported,^[31, 32]^ we assessed *Met* and *Hgf* mRNA levels in *MMTV-R26*^*Met*^ and control mice from the virgin to the post-lactation state. RT-qPCR analysis revealed comparable dynamics of *Met* and *Hgf* transcript expression in both *MMTV-R26*^*Met*^ and control mice: high levels at virgin state, a progressive downregulation during pregnancy, reaching almost undetectable levels during lactation, and a restoration of *Met* and *Hgf* levels at the post-lactation stage (Figure 1C). Whereas *Met*^*tg*^ expression was undetectable in virgin animals, it became evident starting from the pregnancy stage, coherent with *MMTV* promoter activation by prolactin,^[29, 30]^ and remained expressed during subsequent phases. Western blot analysis confirmed Met^tg^ expression in the mammary gland of *MMTV-R26*^*Met*^ mice (Figure S1A).

**Figure 1.**
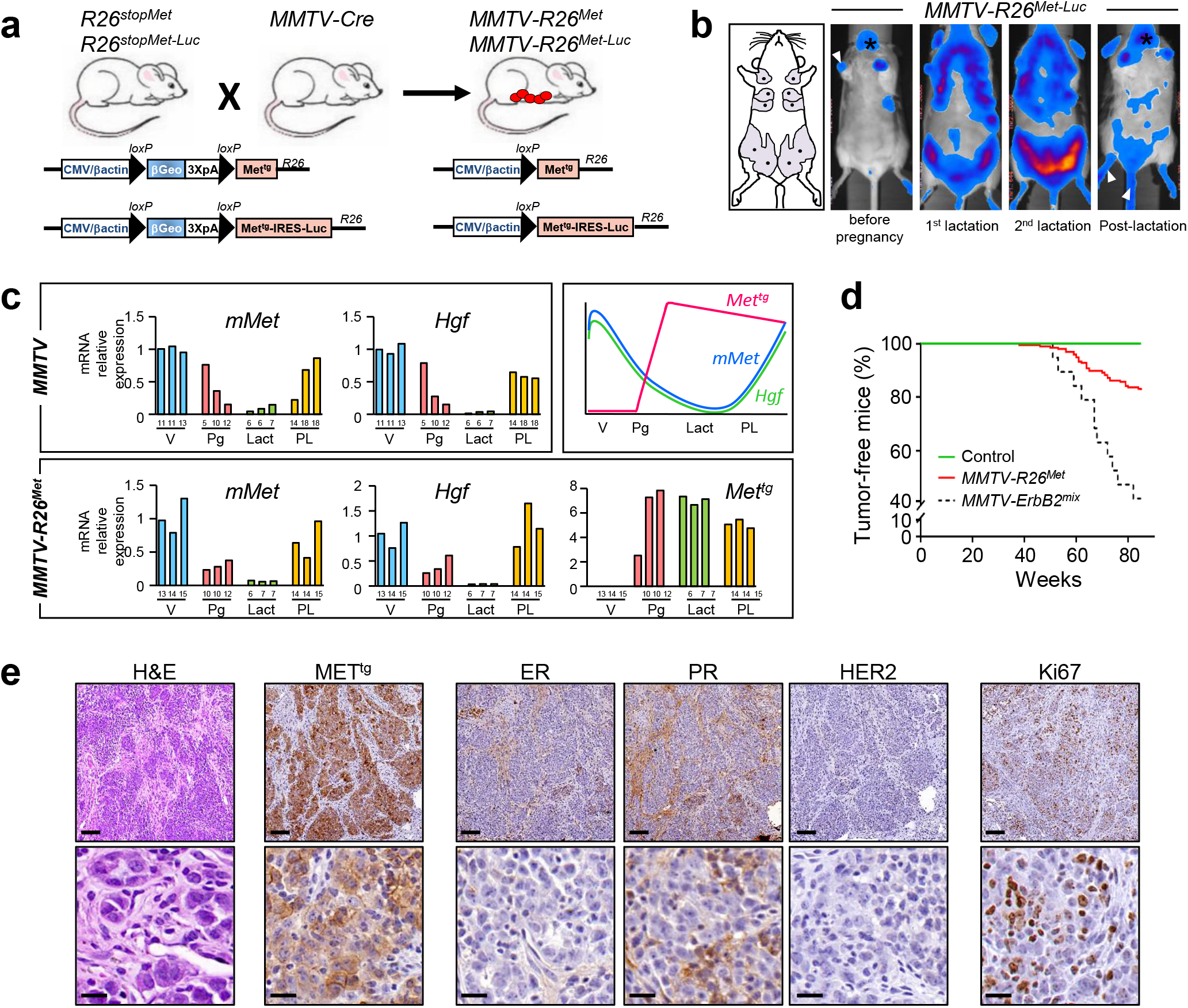
Increased expression of wild-type MET levels in the mouse mammary gland leads to Triple-Negative Breast Cancer formation. a) Strategy used to enhance wild-type MET in the mammary gland of mice. The *R26*^*stopMet*^ mouse line, carrying the LacZ-stop cassette followed by chimeric *Met*^*tg*^, was crossed with the *MMTV-Cre* mice, carrying the Cre recombinase under the control of the mouse mammary tumor virus *MMTV* promoter. After recombination, expression of the *Met*^*tg*^ is ensured by the removal of the LacZ-stop cassette (*MMTV-R26*^*Met*^ mice). The same strategy was used to generate transgenic mice carrying the LacZ-stop cassette followed by *Met*^*tg*^ and IRES-Luciferase before (*R26*^*stopMet-Luc*^) and after (*MMTV-R26*^*Met-Luc*^) Cre-mediated recombination. b) Non-invasive *in vivo* bioluminescence imaging of *MMTV-R26*^*Met-Luc*^ mice. Imaged mice were either not pregnant, under lactation (first or second lactation cycle), or in post-lactation phase. Although mainly detected in the mammary glands, low luciferase expression was also observed in the salivary gland (asterisk), in the skin of the paws and tail (white arrowhead), which is due to partial leakage of the *MMTV-Cre* line, as previously reported[63]. The five pairs of the mouse mammary glands are depicted on the scheme in the left. c) RT-qPCR analyses showing transcript levels of the endogenous mouse *Met* (*mMet*), *Hgf*, and the *Met*^*tg*^, in mammary glands of either *MMTV* (upper left panel) or *MMTV-R26*^*Met*^ (lower panel) mice. Mammary fat pads of three different mice were used for each stage. The age of each mouse is indicated (for virgin animals (V): in weeks; for the other stages: pregnancy (Pg), lactation (Lact), and post-lactation (PL): in days). The scheme on the top right illustrates the dynamic expression of the various transcripts. Note that during lactation, the expression levels of the endogenous *Met* and *Hgf* transcripts are very low, whereas expression of the *Met*^*tg*^ is maintained. d) Kaplan-Meier analysis of mammary gland tumor incidence in *MMTV-R26*^*Met*^, control (*R26*^*stopMet*^), and *MMTV-ErbB2* mice generated in the same mixed (C57/129, 50%/50%) genetic background (*MMTV-ErbB2*^*mix*^). e) Representative histopathological and immunohistological analysis of *MMTV-R26*^*Met*^ tumors using hematoxylin/eosin (H&E), anti-human MET staining to detect expression of the MET transgene (Met^tg^), anti-Ki67 to assess the proliferative index. Expression of the estrogen-(ER), progesterone-(PR), and ErbB2 receptors (HER2) were also analyzed. Scale bar: top panel: 100μm, bottom panel: 20μm.

We therefore hypothesized that the *MMTV-R26*^*Met*^ mice could be an appropriate genetic setting to assess the vulnerability of the mammary gland to subtle perturbation of wild-type MET levels overtime. In view of remarkable changes occurring during mammary gland morphogenesis, illustrated by our bioluminescence imaging and transcriptional analyses, and the susceptibility of parity-induced mammary epithelial subtypes to signaling perturbations,^[33]^ *MMTV-R26*^*Met*^ mice were kept under repeated cycles of pregnancy. Overtime, a proportion of *MMTV-R26*^*Met*^ mice spontaneously developed mammary gland tumors (Figure 1D). Remarkably, the kinetic of tumor formation was similar to that of *MMTV-ErbB2* mice generated in the same genetic background we used as reference (*MMTV-ErbB2*^*mix*^; Figures 1D, S1B). The percentage of mice with tumors correlated with the severity in RTK alteration: 16% of *MMTV-R26*^*Met*^ mice (with enhanced wild-type MET) developed tumors (32/196) compared to 58% of *MMTV-ErbB2*^*mix*^ mice (with oncogenic HERBB2 overexpression; 11/19; Figures 1D, S1B). A proportion of *MMTV-R26*^*Met*^ mice with mammary gland tumors also developed lung metastasis (19%; 6/32; Figure S1C, Table S1). Histological analyses of the *MMTV-R26*^*Met*^ tumors revealed highly aggressive and infiltrating breast carcinomas, which have been histologically identified as being exclusively TNBC (24 tumors analyzed; Figure 1E, Table S1).

### 2.2. The *MMTV-R26*^*Met*^ tumor model recapitulates heterogeneity and primary drug resistance of TNBC human patients

To further characterize the *MMTV-R26*^*Met*^ mammary tumors, we applied a semiquantitative proteomic profiling through reverse phase protein array (RPPA), a high-throughput antibody-based technique to analyze protein activities in signaling networks. Analysis of expression and/or phosphorylation levels (247 signals, Table S2) displayed that the *MMTV-R26*^*Met*^ tumors (n=24) clearly segregate from control mammary glands (n=3; Figure 2A). Interestingly, the *MMTV-R26*^*Met*^ tumors form 4 distinct clusters, highlighting heterogeneity in signaling levels, including the MET phosphorylation status (Figure 2A-B, S1D, Table S3). Heterogeneity was also observed at *Met* transcript levels, as revealed by RT-qPCR (Figure S1E), reflecting the heterogeneity of *MET* levels among TNBC patients.^[34–36]^ Thus, a slight increase in *Met* levels in the mouse mammary glands is sufficient to trigger the tumorigenic program of TNBC.

**Figure 2.**
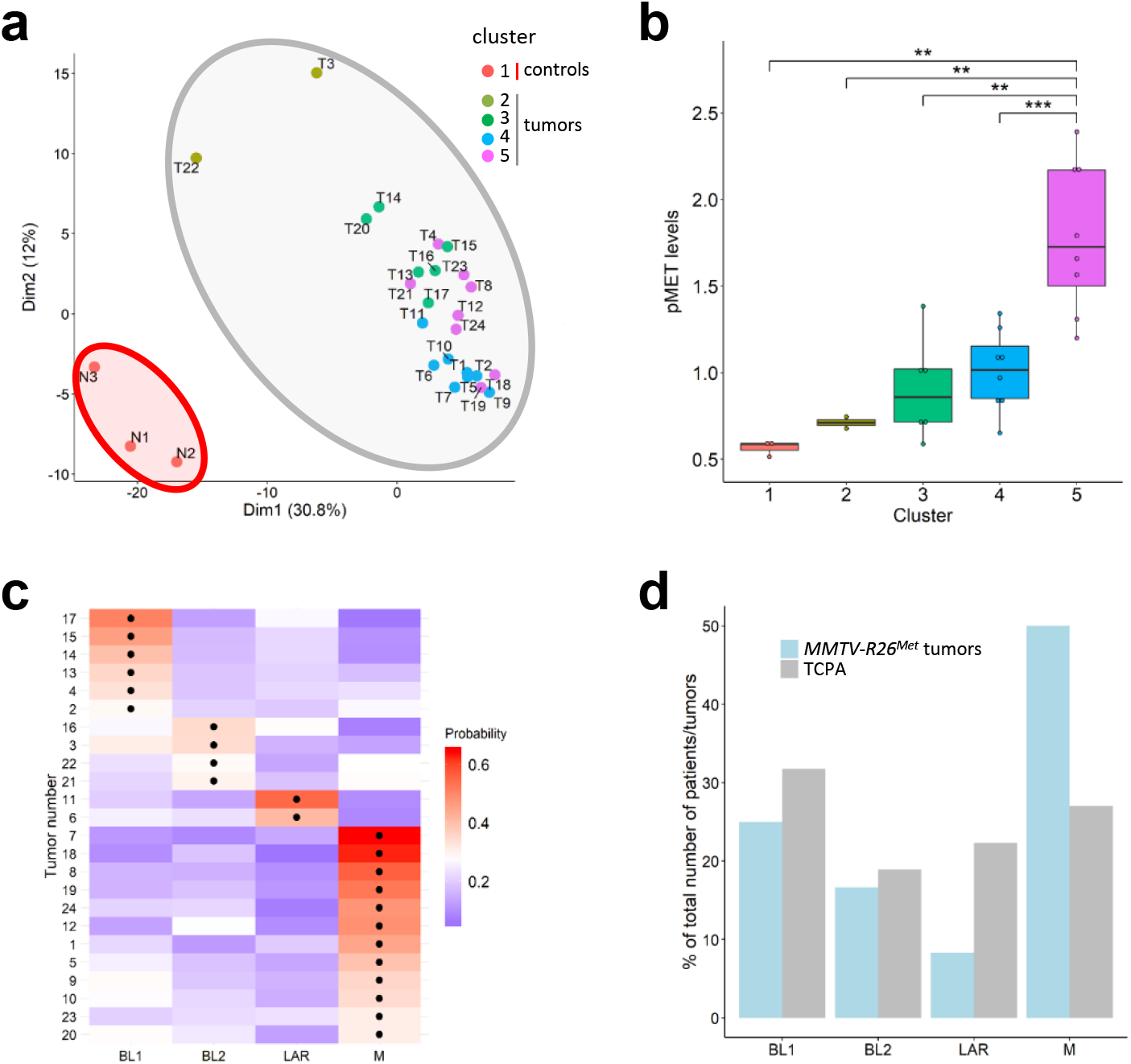
Machine learning processing of RPPA data from tumors and control illustrates that the *MMTV-R26*^*Met*^ model faithfully recapitulates inter-tumoral heterogeneity of human TNBC. a) K-means clustering of the RPPA data for tumor and control samples is depicted in the PCA plot. The red area includes the normal mammary tissue (cluster 1; n=3), whereas the grey area includes the tumors (n=24) separated into four clusters defined by the points color (cluster 2-5). b) Clusters defined in (a) are characterized by different MET phosphorylation status. Colors of the clusters in panels (a) and (b) are the same. c) Heatmap depicting the probability that each tumor belongs to a specific subtype. The black dots indicate the type with the highest probability for each tumor. BL1: basal-like-1; BL2: basal-like-2; LAR: luminal androgen receptor; M: mesenchymal. d) Histogram reporting enrichment of the tumors compared to the TCPA. Note that, even though all subtypes are represented, *MMTV-R26*^*Met*^ tumors are more enriched for the mesenchymal (M) subtype. Values are expressed as means ± s.e.m. ** P<0.01; *** P <0.001.

Next, we explored the possibility to classify the *MMTV-R26*^*Met*^ tumors to TNBC subtypes by analyzing the RPPA data applying the Random Forest machine learning algorithm previously used with transcriptomic data.^[37]^ As subtype classification is usually done on transcriptomic data, we first used the RPPA data of 152 TNBC patients from the TCGA dataset to build a model for subtyping prediction. We trained the model using 10-fold cross validation to optimize the method parameters (Figure S1F). The model was sensitive to the M class (balanced accuracy 0.89), and had lower sensitivity in distinguishing between BL1 and BL2 classes (balanced accuracy of 0.69 and 0.55, respectively). This was done to take into account that most of the patients are BL1, with a consequent 30% correct BL1 prediction called “no information rate”. Our Random Forest model had accuracy of 57% (or 71% without distinguishing BL1 and BL2), with a significance of p-value=0.002 compared to the “no-information rate”. We then applied the model on the RPPA data of *MMTV-R26*^*Met*^ tumors to predict their classification. Remarkably, we found that all TNBC subtypes are represented by the *MMTV-R26*^*Met*^ tumors with an enrichment of the mesenchymal subtype (Figure 2C-D). Collectively, these results showed that a moderate increase of MET levels in the mammary gland is sufficient to perturb tissue homeostasis, is able to initiate the TNBC program including the formation of lung metastasis, and that the resulting tumors recapitulate the heterogeneity characteristic of TNBC patients.

To further exploit the *MMTV-R26*^*Met*^ cancer model, we established and molecularly/biologically characterized six mammary gland tumor (MGT) cell lines from individual *MMTV-R26*^*Met*^ tumors (Figure 3A). Four cell lines, MGT4, MGT9, MGT11, and MGT13 exhibited tumorigenic properties in vivo, illustrated by the formation of tumors when injected heterotopically into the flank of nude mice, whereas the two other lines, MGT2 and MGT7 did not (Figure 3B). The four tumorigenic cell lines exhibited oncogenic features, whereas the MGT2 and MGT7 cell lines did not. In particular, we observed increased MET mRNA and protein levels (Figure S2A-C), with a heterogeneity similar to that observed among *MMTV-R26*^*Met*^ tumors and reported in TNBC patients.^[35, 36]^ MGT4, MGT11, and MGT13 (not MGT9) cell lines were capable of forming tumor spheroids when grown in self-renewal conditions (Figure 3D-E). Additionally, the tumorigenic cell lines were characterized by a high proliferation index, with a low proportion of cells in the G0 cell cycle phase (Figure 3C, S2D-F). Cells of the tumorigenic lines also exhibited increased motility, particularly for MGT13 cells that display a rather mesenchymal-like morphology compared with the other cell lines (Figure 3F, S2A). Furthermore, these cell lines also recapitulated the heterogeneity of p53 alterations observed in TNBC patients: p53 overexpression (likely oncogenic) in MGT4 and MGT9, decreased expression of p53 in MGT13, and comparable p53 levels in MGT11 (Figure S2G).

**Figure 3.**
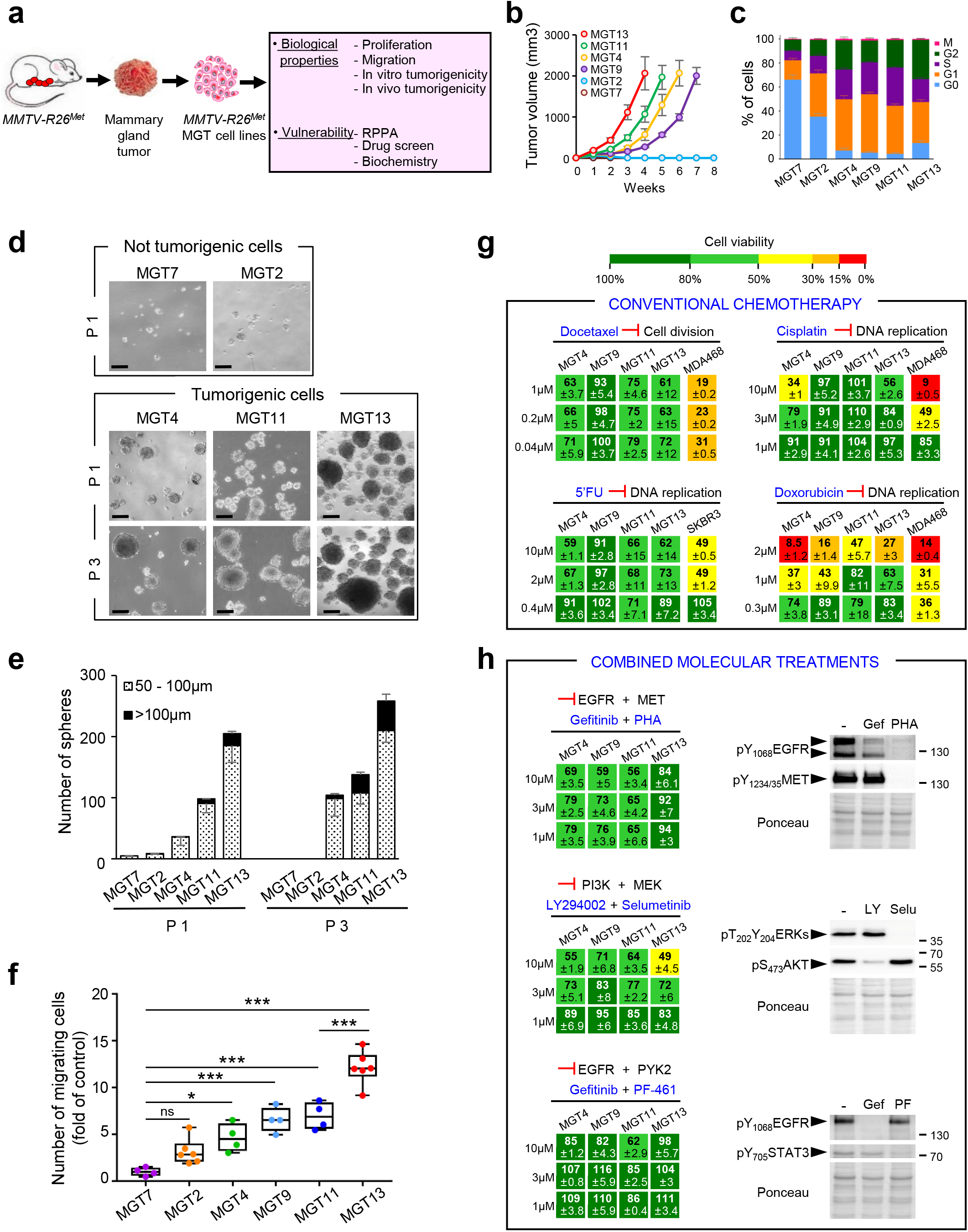
Cells derived from *MMTV-R26*^*Met*^ mammary gland tumors recapitulate primary resistance to drugs used in conventional chemotherapies and to combined molecular treatments. a) *MMTV-R26*^*Met*^ mammary gland tumors (MGT) were used to generate *MMTV-R26*^*Met*^ MGT cell lines, which were then utilized for assessing various biological properties and vulnerability to drugs. b) *In vivo* tumorigenic properties of the *MMTV-R26*^*Met*^ cell lines. Xenografts studies were performed by subcutaneous injection of cells in the flank of nude mice (n=4-5, injected bilaterally). Evolution of the tumor volume shows that MGT4, MGT9, MGT11, and MGT13 are highly tumorigenic cell lines, whereas the MGT2 and MGT7 cells do not form tumors. c) Histogram showing the percentage of cells from each cell line in each phase of the cell cycle as determined by flow cytometry using propidium iodide and Ki67 staining. Three independent experiments were performed. d-e) Tumor sphere formation assessing *in vitro* tumorigenicity of *MMTV-R26*^*Met*^ cells. d) Representative images of tumor spheres derived from MGT7, MGT2, MGT4, MGT11, and MGT13 cells, obtained after 1 (P1) or 3 (P3) passages. e) Histogram reporting the number of spheres, classified in 2 groups according to their size (dotted bars: 50-100μm; black bars >100μm), generated by the indicated *MMTV-R26*^*Met*^ cell lines. Note that: i) the very low capacity of MGT2 and MGT7 in forming spheres is totally abolished after 3 passages; ii) the number and size of spheres generated by MGT4, MGT11, and MGT13 increases from passage 1 to passage 3, reflecting their self-renewal capacity. Each experiment was done in triplicate. f) Quantification of the migration capacity of each *MMTV-R26*^*Met*^ cell line determined by the number of migrating cells compared to MGT7 (fold of control). g) Cell viability of *MMTV-R26*^*Met*^ MGT cells exposed to drugs conventionally used in chemotherapy. Human TNBC cell lines (MDA-MB-468 and SKBR3) were used as positive controls. Percentage of cell viability in presence of drugs compared to controls (untreated cells) is indicated. Percentages are reported using a color code (from green to red; the scale is shown on the top and is used as a reference in all studies). h) Dose-response effects of drug used in combined treatments on the viability of *MMTV-R26*^*Met*^ MGT cells. Western blots depict the effect of each drug on its specific target. Note loss of EGFR phosphorylation in cells treated with PHA-665752, the MET inhibitor. 5’FU: 5-fluorouracil; Gef: gefitinib; LY: LY294002; PF: PF-461; PHA: PHA-665752; Selu: selumetinib. Values are expressed as means ± s.e.m. not significant (ns) P>0.05; * P<0.05; *** P <0.001. Statistical analyses are reported in Table S8 (c), S9 (e), and S10 (f).

Interestingly, the tumorigenic MGT cell lines (MGT4, MGT9, MGT11, and MGT13) were resistant to conventional chemotherapeutic agents, such as Docetaxel, Cisplatin, 5-Fluorouracil (5’FU), and only partially sensitive to Doxorubicin, although only at high doses (Figure 3G). Furthermore, all these MGT cell lines were resistant to three drug combinations previously reported to be effective for TNBC treatment: combined inhibition of EGFR+MET, PI3K+MEK, and EGFR+PYK2^[38–40]^ (Figure 3H). Together, these results show that the *MMTV-R26*^*Met*^-derived cell lines are a relevant model to study as well drug resistance, an important feature of TNBC.

### 2.3. Signaling network analysis of *MMTV-R26*^*Met*^ tumor derived cells

To further characterize *MMTV-R26*^*Met*^ MGT cells, we examined their signaling status by RPPA and subsequent bioinformatics analysis (247 epitopes, listed in Table S2). The signaling profiles highlighted two major features, as illustrated by Principal Component Analysis (PCA). First, the tumorigenic MGT4, MGT9, MGT11, and MGT13 cells clearly segregate from the two types of non-tumorigenic cells: MGT7 and MGT2 (Figure 4A, S3A, Table S4). MGT2 cells, which express very low level of MET, can be considered as pre-tumorigenic. It is therefore tempting to speculate that critical levels of MET might establish a threshold for a tumorigenic switch, while higher MET levels are associated with aggressiveness. Second, the four tumorigenic *MMTV-R26*^*Met*^ MGT cell lines fall into two distinct TNBC subtypes that we named “subtype A” for MGT4, MGT9, MGT11, and “subtype B” for MGT13 (Figure 4A, S3A, Table S4). These two subtypes display distinct phenotypic features and MET levels (Figure S2A-C). Strikingly, by PCA analysis we could segregate into “subtype A” and “subtype B” both *MMTV-R26*^*Met*^ MGT cells and tumors (Figure 4B). Additionally, we identified ARID1A, Claudin-7, and E-Cadherin as hallmark of the “subtype A” (Figure 4C, Table S5).

**Figure 4.**
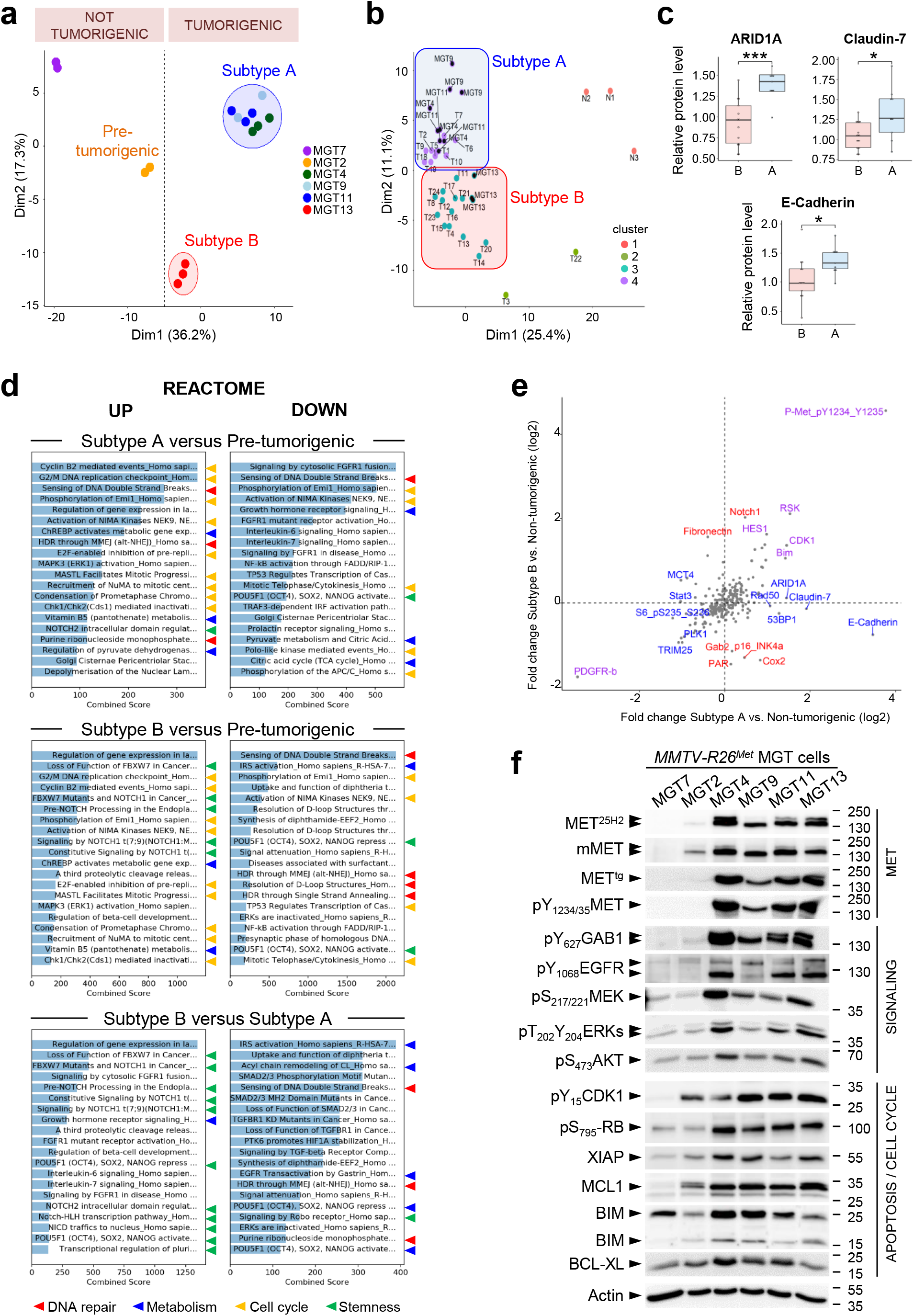
Proteomic analysis highlighted signaling changes in *MMTV-R26*^*Met*^ tumors, leading to the identification of a new potent drug combination for TNBC cells. a) Principal Component Analysis of the *MMTV-R26*^*Met*^ MGT cell lines, using Reverse Phase Protein Array (RPPA) data. The 2 non-tumorigenic cell lines are well separated from each other, with MGT2 that we named “pre-tumorigenic” cells. Both non-tumorigenic cells are distinct from the two tumorigenic cell clusters, designated as “subtype A” (MGT4, MGT9, MGT11) and “subtype B” (MGT13). b) Graph showing the combined PCA of *MMTV-R26*^*Met*^ MGT cell lines and tumors, according to k-mean clustering (k=4). Cluster 1: normal tissues; cluster 2: low phospho-MET tumors; cluster 3: “subtype B” cell line (MGT13) and tumors; cluster 4: “subtype A” cell lines (MGT4, MGT9, MGT11) and tumors. Cell lines are indicated by a black dot. c) Graphs reporting the expression levels in “subtype A” (A) and “subtype B” (B) *MMTV-R26*^*Met*^ tumors of the indicated proteins, considered as the hallmark of the “subtype A” cell lines (Table S5). d) Proteomic profiles of cells belonging to the different clusters (shown in a) were compared to identify enrichments by applying the Enrichr software. Histograms report the enriched cell signaling pathways, using the Reactome database, ordered according to the combined score. The 20 top ranked enrichments are highlighted. Note that the majority of the changes are related to signals (indicated by arrowheads) involved in DNA repair (red), metabolism (blue), cell cycle regulation (yellow), and stemness (green) (indicated by colored arrowheads). e) Graph showing the fold change (Log2) of protein phosphorylation or expression between “subtype A” (x-axis) or “subtype B” (y-axis) cell lines versus the non-tumorigenic cells (MGT2, MGT7). Proteins among the highest significant differentially expressed in “subtype A” (in blue), “subtype B” (in red), or both (in purple) are indicated (Table S6). f) Western blot analysis of total protein extracts from *MMTV-R26*^*Met*^ MGT cells. Actin and Ponceau S stainings were used as loading controls in all studies.

To obtain insights on molecular and cellular functions characterizing “subtype A” from “subtype B”, we performed a series of enrichment analyses by applying Enrichr, a web-based tool to highlight enrichments based on gene sets. Both subtypes showed an enrichment in pathways related to DNA repair, cell cycle regulation, and metabolism (Figure 4D, S3B). These enrichments are consistent with enhanced proliferation capacity of MGT4, MGT9, MGT11, and MGT13 cells versus non-tumorigenic cells. Moreover, “subtype B” is enriched in pathways related to stemness properties (Figure 4D, S3B), consistent with the enhanced capability of MGT13 to form tumor spheroids in vitro (Figure 3D-E). Further analysis of RPPA data using the limma package highlighted differences between the RPPA profiles of subtypes A and B versus the non-tumorigenic MGT cells. In particular, we detected upregulation of: a) Bim, which might sensitize the cells to anti-apoptotic drugs, b) CDK1 and RAD50, which are implicated in cell cycle regulation and DNA damage response (Figure 4E; Table S6). Biochemical studies supported the RPPA results and revealed consistent upregulation of oncogenic signals in MGT4, MGT9, MGT11, and MGT13 compared with control cells. This included phosphorylation of MET, EGFR, of their downstream adaptor GAB1, of MEK/ERKs, AKT, RB, and elevated anti-apoptotic signals such as MCL1, BCL-XL, and XIAP (Figure 4F).

### 2.4. Combined targeting of WEE1 and BCL-XL is deleterious for TNBC cells

Inspired by the signaling profiles of MGT cells and tumors, we designed a drug screen aiming at identifying combinatorial treatments effective for the two subtypes of TNBC cells modeled by the *MMTV-R26*^*Met*^ mice. Among all treatments tested in the MGT4 cell line (single or combined drugs), we uncovered that the simultaneous inhibition of BCL-XL and WEE1 drastically reduced tumor cell viability (Figure S4A-B). By further assessing the effects of this combined treatment on the six *MMTV-R26*^*Met*^ MGT cell lines, we found that BCL-XL+WEE1 inhibition was deleterious for all four tumorigenic *MMTV-R26*^*Met*^ MGT cells (MGT4, MGT9, MGT11, MGT13), but not for the non-tumorigenic cells (MGT2 and MGT7; Figure 5A-B). Importantly, this highlights lack of toxic effect of the newly identified drug combination. Combined inhibition of BCL-XL and WEE1 was synergistic (for 3 out of 4 *MMTV-R26*^*Met*^ MGT cell lines), as shown by the Bliss score and by the Chou-Talalay combination index score calculation (Figure 5A, S4C). Furthermore, BCL-XL+WEE1 targeting was detrimental for all six human TNBC cells tested (Figure 5C). Intriguingly, when this drug combination was tested on human non-TNBC cells, we found that inhibition of BCL-XL did not exacerbate the effects elicited by WEE1 targeting (Figure 5D), indicating that WEE1 inhibition is particularly detrimental in TNBC cells.

**Figure 5.**
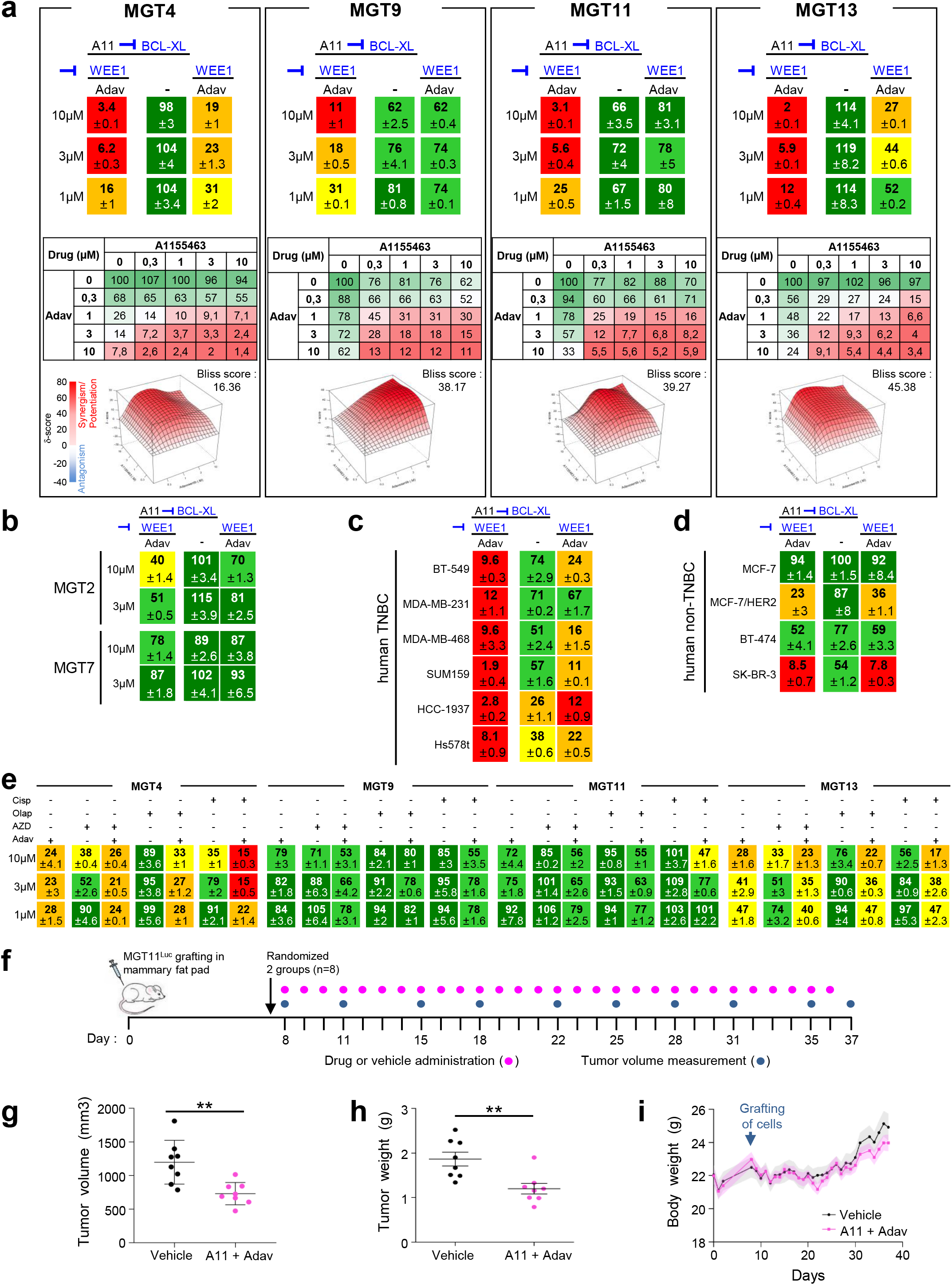
Combined inhibition of BCL-XL and WEE1 is deleterious for all *MMTV-R26*^*Met*^ MGT and human TNBC cell lines tested. a) Dose-response effects of A1155463 (A11, targeting BCL-XL) alone or in combination with Adavosertib (Adav, targeting WEE1) on the viability of the four tumorigenic *MMTV-R26*^*Met*^ MGT cell lines. Combined drug effects are reported on the left of the top panel. Detailed matrix (middle panel) and Loewe plots (lower panel) highlight the drug synergism. b) Cell viability assay performed on the non-tumorigenic MGT2 and MGT7 cell lines highlights the lack of in vitro toxic effect of A1155463+Adavosertib drug combination. c-d) Cell viability in response of A1155463 (1μM) and Adavosertib (3μM) in a panel of human TNBC (c) and human non-TNBC breast cancer (d) cell lines. In all figures, cell viability is presented as percentage of control (untreated cells) and labelled by the green (high) to-red (low) color code. In a-d, at least three independent experiments were performed. e) Dose-response effects on the viability of *MMTV-R26*^*Met*^ MGT cells treated with single or combined drugs as indicated. f-i) *In vivo* effects of A1155463+Adavosertib treatment in orthotopic tumors. f) Orthotopic injection of MGT11^Luc^ cells in the mammary fat pad of NSG mice, drug administration, and tumor volume measurement were performed as illustrated. Tumor volume (g) and tumor weight (h) measured at the end point of the experiment (day 37; n=8 mice per group). i) Graph reporting the evolution of the body weight of mice during the whole procedure. Body weight was measured every day, before drug administration. No significant changes were observed, indicating that the dose of drugs used *in vivo* were not toxic. A11: A1155463 (BCL-XL inhibitor); Adav: Adavosertib (WEE1 inhibitor); AZD: AZD6738 (ATR inhibitor); Cisp: cisplatin; Olap: Olaparib (PARP inhibitor). Values are expressed as the mean ± s.e.m. ** P <0.01.

Recent studies have reported the sensitivity of TNBC to WEE1 targeting in the presence of either PARP or ATR inhibitors, or Cisplatin.^[17, 19, 20, 22, 41]^ However, we found that *MMTV-R26*^*Met*^ MGT cells were either resistant or only partially sensitive to these drug combinations (Figure 5E), recapitulating other mechanisms of primary resistance beside those reported in Figure 3G-H. Thus, BCL-XL targeting is a preferable strategy to exacerbate WEE1 essentiality in TNBC.

Finally, we assessed in vivo the potency of BCL-XL+WEE1 co-targeting on tumor growth. We engineered a cell line for in vivo imaging by stably transfecting the MGT11 cells, characterized by strong tumorigenic properties, with a Luciferase reporter vector (defined as MGT11^Luc^). We confirmed that the MGT11^Luc^ cells have comparable biological properties as the parental cells, and maintain sensitivity to combined BCL-XL+WEE1 targeting (Figure S4D-F). Orthotopic studies showed that combinatorial BCL-XL+WEE1 inhibition reduced in vivo tumor growth of MGT11^Luc^ cells injected into the mammary fat pad of mice (Figure 5F-I, S4G). No obvious effects on mouse viability or murine weight indicated the lack of toxicity. Thus, the *MMTV-R26*^*Met*^ model recapitulating heterogeneity and resistance of TNBC led us to uncover a new potent drug combination based on BCL-XL and WEE1 inhibition, effective on human TNBC cell lines characterized by distinct features.

### 2.5. BCL-XL inhibition exacerbates WEE1 requirement in TNBC cells

While BCL-XL primarily has an anti-apoptotic function by inhibiting cytochrome C release, WEE1 acts on multiple regulatory circuits. Besides its well-established involvement in regulating the G2/M transition through phosphorylation of CDK1, recent studies have highlighted additional mechanistic roles of WEE1 in DNA replication stress and regulation of histone synthesis and levels. Therefore, we thoroughly examined the signaling changes occurring following BCL-XL and WEE1 inhibition in MGT cell lines by RPPA analysis (426 epitopes were analyzed, Table S2); some changes were validated by Western blot studies. As examples, the profile of proteins differentially expressed and/or phosphorylated in MGT4 is displayed in Figure 6A (Table S7). Interestingly, while BCL-XL inhibition alone had modest effects, WEE1 inhibition had marked effects on the cells, many of which were accentuated by treatment with the BCL-XL inhibitor. Among identified changes, some were consistently observed in all MGT cell lines, whereas others were specific to individual MGT lines. These changes covered a broad range of signaling/cellular functions, such as those associated with cell survival/death, cell cycle regulation, DNA damage/repair, histone levels, and oncogenic properties (Figure 6B, Table S7).

**Figure 6.**
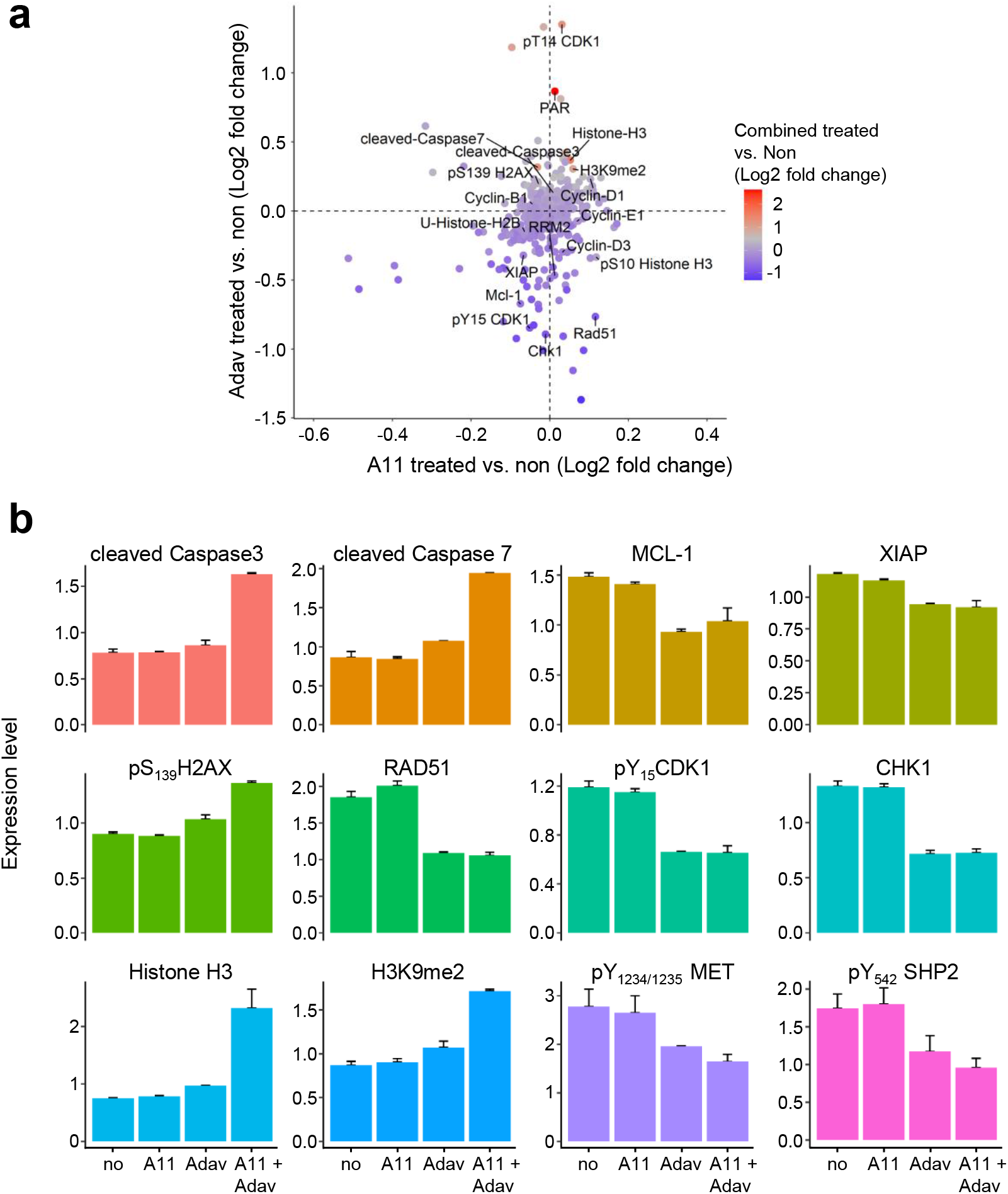
BCL-XL and WEE1 targeting leads to perturbation of several signals, including epigenetic, DNA damage/repair, apoptosis, and cell cycle regulators. a) Graph showing the fold change (Log2) of protein phosphorylation or expression in MGT4 cells between: A1155463 (A11) versus untreated (on the x-axis), Adavosertib (Adav) versus untreated (on the y-axis), and the combination versus untreated (colors of the dots). Changes related to epigenetics (HistoneH3, H3K9me2, phosphoS10HistoneH3, U-HistoneH2B, RRM2) DNA damage and repair (γH2AX, Rad51, PAR), apoptosis (cleaved Caspase7, cleaved Caspase3), and cell cycle (CHK1, CDK1, cyclins) are indicated. b) Changes in the expression/phosphorylation levels of the reported proteins in MGT4 cells untreated and treated with the indicated drugs, based on RPPA analysis (Table S7).

Consistent with the well-known regulatory activity of WEE1 in cell cycle progression, we observed decreased levels of phosphoY_15_CDK1 (the direct target of WEE1), phosphoS_795_RB, and CHK1 expression upon WEE1 inhibition (Figure 6B, 7A, Table S7). This was accompanied by an alteration in the distribution of cells in cycle phases as shown by FACS profiles (Figure 7B). Concerning cell survival signals, combined BCL-XL+WEE1 targeting in MGT cells led to a drastic down-regulation of MCL1 and XIAP anti-apoptotic signals associated with intense cleavage of Caspase3 and PARP (Figure 6A-B, 7A). Regarding DNA damage and repair, we observed an up-regulation of phosphoS_1987_ATM and phosphoS_139_H2AX (histone variant, γH2AX), reflecting increased levels of DNA damage in the MGT cells upon treatment (Figure 6A-B, 7A, S5A). This high proportion of DNA damage is associated to the downregulation of Rad51 (Figure 6A-B), which plays a major role in DSB repair by homologous recombination and in fork protection, and restart during replication stress,^[42]^ raising the possibility that this down-regulation is related to the increase of phosphoS_139_H2AX. Moreover, we observed a drastic downregulation of RRM2, a subunit of the ribonucleotide reductase required to maintain high levels of dNTPs,^[23]^ with an upregulation of Histone-H3 and H3K9me2 levels (Figure 6A-B, 7C-D, S5B, Table S7). This was further confirmed by cell fractionation studies, showing downregulation of RRM2, and increased pS_33_RPA32 in the chromatin fraction of cells co-treated with WEE1 and BCL-XL inhibitors (Figure 7E). Finally, concerning oncogenic signals, we found a significant downregulation of phosphorylation levels of MET and GAB1 (Figure 7A). Collectively, these results indicated that BCL-XL inhibition exacerbates the dependence of TNBC cells on the overall functions exerted by WEE1: an intact dNTP pool (by stabilizing RRM2 protein levels), appropriate histone levels, and proper cell cycle progression through G2/M.

**Figure 7.**
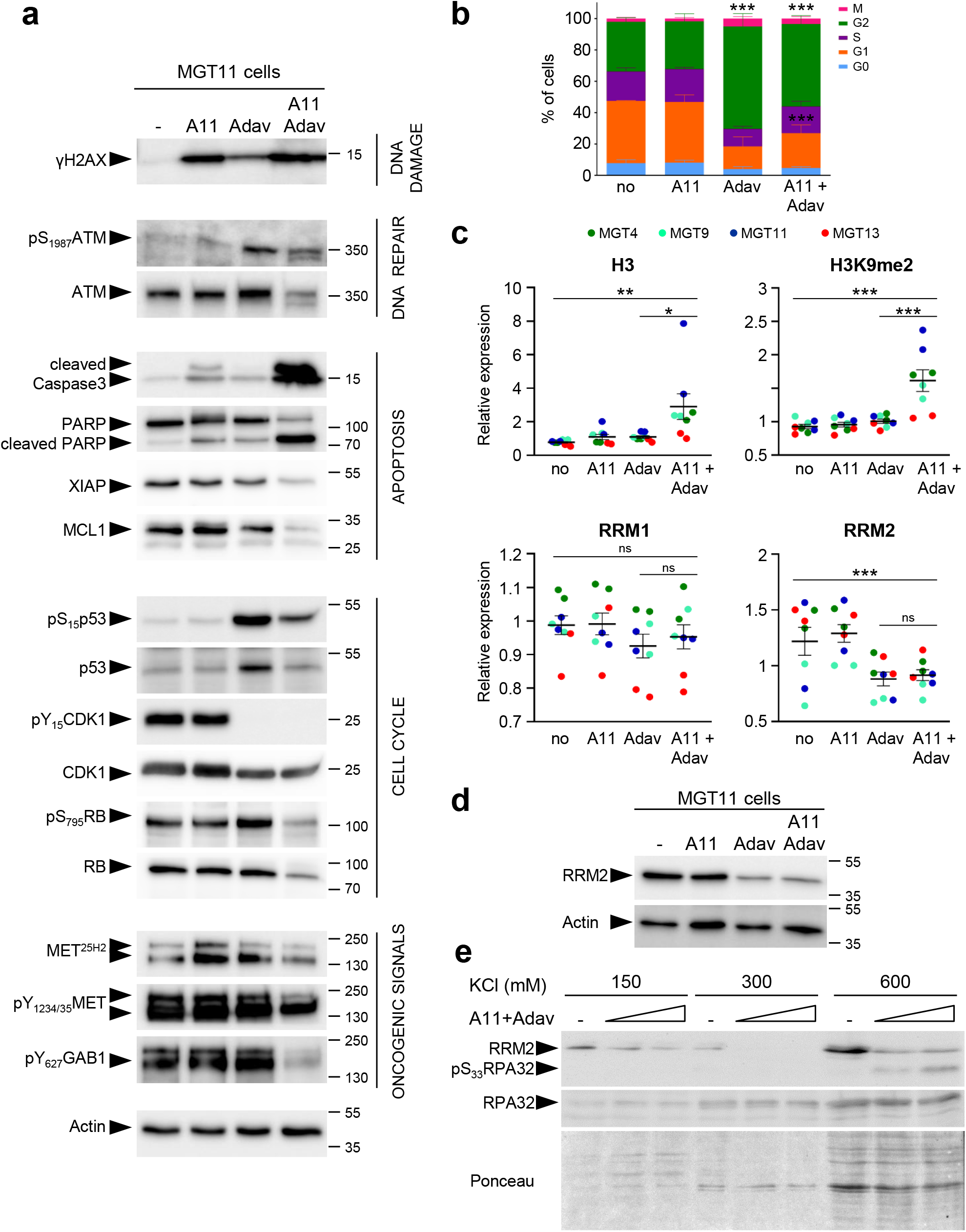
BCL-XL targeting exacerbates WEE1 requirement in TNBC cells. a) Western blots showing the effects of A1155463 (1μM), Adavosertib (3 μM), and combined treatment on the indicated signals in the MGT11 cells, 12hrs after treatment. b) Graph reporting the distribution of cells treated with the indicated drugs (for 12hrs), compared to untreated cells (no), in each phase of the cell cycle as determined by flow cytometry using PI and Ki67 staining. All statistical analyses are reported in Table S11. c) Graphs reporting changes by RPPA in levels of Histone H3, H3K9me2, RRM1, and RRM2 in MGT cells exposed to A1155463, Adavosertib, or in combination. Distinct MGT cell lines are indicated in colors. d) Western blot showing RRM2 downregulation in MGT1 cells treated with Adavosertib alone or in combination with A1155463. e) Western blot showing downregulation of RRM2 and increase of pS33RPA32 levels in the chromatin fraction (corresponding to the 600mM KCl) of cells co-treated with WEE1+BCL-XL (A11+Adav). Cells were treated with A1155463 (0.3μM) plus Adavosertib (1μM or 3μM). Three independent experiments were performed. A11: A1155463; Adav: Adavosertib. Values are expressed as the mean ± s.e.m. not significant (ns) P>0.05; * P<0.05; ** P <0.01; *** P <0.001.

### 2.6. Combined BCL-XL and WEE1 inhibition leads to mitotic catastrophe and apoptosis of TNBC cells

We explored at cellular levels the biological events associated with BCL-XL and WEE1 inhibition by immuno-cytochemistry. In cells experiencing the combined treatment, we found a significant increase in the number of γH2AX-positive cells as well as a raise in intensity of γH2AX staining per cell, reflecting an accumulation of DNA DSBs (Figure 8A). These findings are in agreement with the above results (Figure 7A, S5A) and reflect increased DNA damage in a high proportion of cells following BCL-XL+WEE1 targeting. In addition, we found a striking increase in phosphoS_10_Histone H3 (pH3)-positive cells when subjected to the combined treatment (Figure 8B), suggesting that a high proportion of cells are in G2/M.^[43]^ This could reflect a premature entry in mitosis due to WEE1 inhibition, but also an accumulation of unrepaired DNA damage in mitosis.^[13, 44]^ We investigated the consequences of this premature mitotic entry by performing a double immuno-staining with anti-α-Tubulin and anti-pH3 antibodies in cells treated (or not) with BCL-XL+WEE1 inhibitors. Interestingly, the staining highlighted a marked increase of cells harboring mitotic catastrophe revealed by monopolar, multipolar, or disorganized spindles, and even cytokinesis failure (Figure 8C). The results further showed that BCL-XL inhibition exacerbated the effects of WEE1 targeting by forcing cells to exit mitosis without undergoing complete chromosome segregation, a phenomenon called mitotic slippage. As a consequence, these excessive unscheduled and abnormal mitosis events led to an increased formation of micronuclei in treated cells (Figure 8D).

Finally, we assessed the terminal event associated with combined BCL-XL and WEE1 targeting on TNBC cells. We found that a pan-Caspase inhibitor (Z-VAD-FMK) significantly rescued cell death caused by the combined treatment of TNBC cells (Figures 8E-F). This result was consistent with Western blots analysis showing a strong increase in Caspase3 and PARP cleavage upon combined BCL-XL+WEE1 inhibition (Figure 7A). In contrast, inhibition of ferroptosis, another cell death mechanism to which TNBC cells are highly sensitive,^[45]^ did not prevent cell death (Figure 8E-F). Together, these findings indicate that the combined targeting of BCL-XL and WEE1 exacerbates the dependency of TNBC cells on WEE1 function in a context of low anti-apoptotic inputs, leading to mitotic catastrophe and apoptosis.

**Figure 8.**
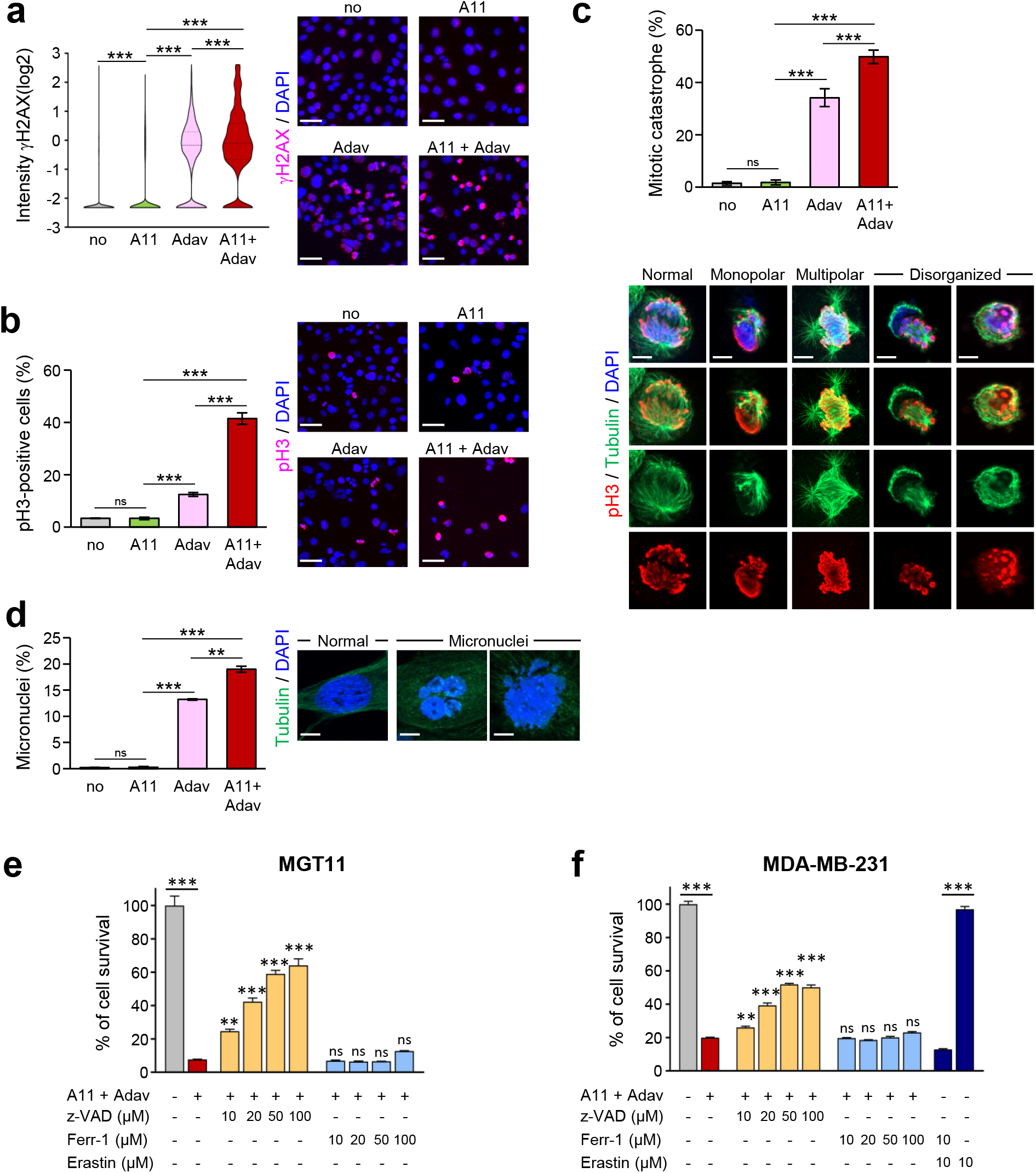
Combined BCL-XL and WEE1 inhibition leads to mitotic catastrophe and apoptosis. Cells untreated or treated for the indicated times with A1155463 (0.3μM), Adavosertib (3μM), or in combination, were immunostained with the indicated antibodies. DAPI was used to counterstain the nuclear DNA. a) MGT11 cells treated for 12hrs with the drugs were immunostained with anti-γH2AX antibodies. Left: the number of γH2AX-positive cells according to the intensity of staining is represented in the violin plot (Log2). Right: representative images of γH2AX immunostaining. Scale bar: 50μm. Four independent experiments were performed. b-c) MGT11 cells treated for 16hrs with the drugs were stained with anti-pH3 (red) and α-Tubulin (microtubules, green) antibodies. b) Left: percentage of cells in mitosis (pH3-positive cells) versus total number of cells. Right: representative images of pH3 immunostaining. Scale bar: 50μm. c) Quantifications (top) and examples (bottom) of cells harboring either typical mitotic phenotypes (normal) or mitotic catastrophe (monopolar, multipolar, or disorganized spindle) revealed by anti-pH3/α-Tubulin immunostaining. To calculate the percentage of cells harboring mitotic catastrophe, we considered cells in metaphase and anaphase among the pH3-positive cells. Scale bar: 5μm for multipolar spindle, 10μm for normal monopolar and disorganized spindle images. Three independent experiments were performed. d) MGT11 cells were treated for 24hrs with the indicated drugs. Nuclear DNA was counterstained with DAPI. Percentage of cells with micronuclei versus the total number of cells (left) and representative images (right) are shown. Scale bar: 10μm. Three independent experiments were performed. e-f) The pan-caspase inhibitor z-VAD-FMK rescues from cell death induced by the combination of A1155463 and Adavosertib. *MMTV-R26*^*Met*^ MGT11 (e) and MDA-MB-231 (f) cells were pretreated with either the apoptosis (z-VAD: z-VAD-FMK) or the ferroptosis (Ferr-1: Ferrostatin-1) inhibitor at the indicated doses, for 1hr and then for additional 24hrs in the absence or presence of A1155463 (A11, 0.3μM) + Adavosertib (Adav, 3μM). Histograms represent the percentage of cell viability in presence of drugs compared to controls (untreated cells). Statistics refer to cell viability obtained in presence of the apoptosis or ferroptosis inhibitors compared to A1155463+Adavosertib alone. The efficiency of Ferrostatin-1 was assessed in presence of Erastin, a ferroptosis inducer. Three independent experiments were performed. Data are expressed as means ± s.e.m. not significant (ns) P>0.05; ** P<0.01; *** P <0.001.

## 3. Discussion

In this study, we have developed a new TNBC mouse model that recapitulates primary drug resistance, we uncovered that the combined inhibition of WEE1 and BCL-XL selectively kills mouse and human TNBC cell lines, and provided mechanistic insights into inhibition of WEE1 and BCL-XL in TNBC cells.

A number of transgenic mice have been engineered to model TNBC, predominantly through drastic genetic manipulations (often combined), such as loss-of-tumor suppressors and overexpression of activated oncogenes.^[46]^ Although they have been instrumental to implicate candidate genes as oncogenes, each model generally recapitulates a fraction of disease features predominantly associated with the genetic manipulation employed. Patient-derived xenografts (PDXs) capture the heterogeneity that characterizes cancers like TNBC. However, transplantation-based models in immunocompromised mice do not report on the reciprocal crosstalk between cancer and immune cells, a limitation that can be overcome, in part, by laborious and expensive “humanized models”. However, most murine TNBC models do not recapitulate the formation of spontaneous cancers occurring in human patients. In this respect, the *MMTV-R26*^*Met*^ model is rather unique for a series of features.

(i) Tumors developed by *MMTV-R26*^*Met*^ mice are exclusively TNBC rather than covering a range of breast cancer types. This is not the case for other transgenic mice overexpressing MET oncogenic forms in the mammary gland,^[27, 47–49]^ generated in the past because of the implication of MET in breast cancer pathophysiology. Indeed, MET is overexpressed in about 40% of breast cancer patients (in luminal A: 36%; in Luminal B: 39%; in HER2+: 48%; in TNBC: 53%^[50]^), and its overexpression often correlates with poorly differentiated and aggressive forms of the disease.^[35]^ In TNBC, MET is particularly highly expressed, and implicated in malignancy progression, metastasis, and resistance to anticancer therapies.^[34, 35]^ Consequently, agents targeting MET are actively being explored for clinical purposes,^[51, 52]^ including in TNBC.^[53–55]^
(ii) In *MMTV-R26*^*Met*^ mice, the TNBC program is driven by a subtle increase of MET levels in the mammary gland, and by the wild-type form of MET rather than oncogenic versions of MET (mutated MET or TPR-MET).^[27, 47, 48]^ Consequently, *MMTV-R26*^*Met*^ TNBC cells are not addicted to MET, a feature predominantly characterizing cancers induced by driver oncogenes. This might explain why the *MMTV-R26*^*Met*^ model reported here recapitulates the tumor heterogeneity typical of TNBC patients.
(iii) TNBC heterogeneity is recapitulated by *MMTV-R26*^*Met*^ mice at different levels.
(iv) *MMTV-R26*^*Met*^ TNBC recapitulates primary resistance to conventional chemotherapy and to a set of targeted molecular treatments reported in previous studies.^[39]^

Histologically, the 28 *MMTV-R26*^*Met*^ tumors analyzed revealed differences in their grade (although the majority were high-grade), with a range of low to high mitotic index, and the absence or presence of necrotic areas. Heterogeneity is also evidenced by processing our RPPA analysis of 24 *MMTV-R26*^*Met*^ tumors through machine learning, indicating that the 4 TNBC subtypes (BL1, BL2, LAR, and mesenchymal) are indeed represented, with the most aggressive mesenchymal subclass enriched. Diversity was preserved in vitro by the four *MMTV-R26*^*Met*^ MGT cells we established from distinct tumors. Heterogeneity was maintained even concerning MET levels in TNBC, as shown for example by the different levels of MET expression and activation in *MMTV-R26*^*Met*^ tumors and cells (for example, phosphorylation levels of MET and of GAB-1, a MET downstream effector). It is tempting to speculate that although all tumors in *MMTV-R26*^*Met*^ mice originate from a common genetic setting characterized by a slight increase of MET levels, this context does not impose an oncogenic path in which MET would be systematically altered in all tumors to the same extent. Finally, *MMTV-R26*^*Met*^ MGT cells are also heterogeneous in terms of drug sensitivity: for example, while MGT9 and MGT11 are resistant to single WEE1 targeting, MGT4 and MGT13 are partially sensitive.

It is rather surprising that a subtle increase of MET levels, in its wild-type form, spontaneously initiates a destabilization process that fully recapitulates the whole TNBC program. Nevertheless, this sensitivity to MET levels is conditioned by a multiparous context, as females without multiple pregnancies did not develop tumors. Both MET and HGF are dynamically expressed during pregnancy/lactation, as we showed here consistent with previous reports.^[31, 32]^ Moreover, the HGF/MET system regulates mammary gland morphogenesis, especially ductal branching and proliferation of ductal end buds.^[56]^ These data illustrate how a tight regulation of the time and signal input levels required for the mammary gland remodeling is critical to prevent transformation. The vulnerability of the mammary gland to a slight increase in MET levels resembles the susceptibility of the liver we reported in previous studies.^[28, 57–59]^ The vulnerability of the mammary gland and the liver is contrasted by a remarkable resilience of other tissues, in which a tumorigenic event requires additional genetic alterations, as reported for malignant peripheral nerve sheath tumors.^[60]^ Whether such a mild MET perturbation in the mammary gland occurs in specific subgroups of women and/or physiological contexts and can increase susceptibility to tumor development remains an open issue. If this is not the case, such genetic manipulation nevertheless makes it possible to initiate a cascade of molecular events leading to a clinically relevant TNBC context.

The second major finding of this work is that combinatorial inhibition of WEE1 and BCL-XL kills a panel of heterogeneous *MMTV-R26*^*Met*^ and human TNBC cell lines. For decades, WEE1 has been considered primarily as a key regulator in cell cycle progression.^[13, 14]^ In particular, WEE1 regulates the G2/M checkpoint through phosphorylation and inactivation of CDK1, thus preventing entry of cells with unrepaired DNA damage into mitosis.^[13]^ Nevertheless, additional mechanistic functions of WEE1 have recently emerged. Indeed, WEE1 stabilizes RRM2 protein, a regulatory subunit of the ribonucleotide reductase required to maintain high dNTPs levels.^[23]^ In addition, WEE1 was reported in yeast and human to inhibit transcription of several histone genes by phosphorylating Histone H2B at Tyr_37_.^[25]^ In addition, the WEE1 yeast homolog Swe1^WEE1^ was recently reported to act as a histone-sensing checkpoint by sensing excess histone levels before cells enter mitosis, thus preventing aberrant chromosomal segregation and polyploidy.^[26]^ Thus, WEE1 targeting might affect several key cellular processes that are particularly relevant in cancer cells as they proliferate at high rates and are more prone to replication stress with higher demands in dNTP and histones. WEE1 inhibition may therefore particularly expose cancer cells to DNA damage. This is reflected by the marked increase of cells harboring mitotic catastrophe upon the combined inhibition of WEE1 and BCL-XL in TNBC cells.

WEE1 is an attractive target for cancer therapies including for TNBC, and strategies are being intensively explored in preclinical studies and clinical trials. It has been recently reported that WEE1 targeting, in combination with either cisplatin or inhibitors of ATR or PARP is effective in human TNBC cells lines.^[17, 19, 20, 22, 41]^ By testing them in *MMTV-R26*^*Met*^ MGT cells, mimicking primary resistant treatment contexts, we have shown that these three combinations are indeed effective, albeit to a varying degree and depending on the cell line. Nevertheless, the combinatorial targeting of WEE1 together with BCL-XL elicits superior effects, as shown by the loss of viability of all four very aggressive/highly tumorigenic *MMTV-R26*^*Met*^ MGT cell lines and of the six human TNBC cell lines tested. Interestingly, such vulnerability is specific to TNBC cells as three out of four non-TNBC cell lines were resilient to WEE1 plus BCL-XL inhibition. This resilience, as well as the absence of effects on two non-tumorigenic *MMTV-R26*^*Met*^ MGT cell lines (MGT2, MGT7) highlights two relevant points. First, the reduction of the stress support pathway by targeting BCL-XL exacerbates a specific requirement of WEE1 in TNBC. This effect resembles an “essentiality-induced” synthetic lethality, characterized by the essentiality of one gene following the targeting of a second gene.^[61]^ Nevertheless, we cannot exclude that BCL-XL targeting may also contribute to altered cell cycle progression.^[62]^ Second, in addition to the absence of in vivo side effects, BCL-XL plus WEE1 targeting appears to be a rather safe treatment for healthy cells.

## 4. Conclusion

We propose that the *MMTV-R26*^*Met*^ genetic setting we have generated is a relevant model for molecular and preclinical studies on TNBC in an immunocompetent context. The panel of *MMTV-R26*^*Met*^ MGT cells we established are particularly useful to screen TNBC therapeutic options. The usefulness of the *MMTV-R26*^*Met*^ model is further strengthened by its capability to recapitulate TNBC heterogeneity and primary resistance to treatments. We illustrated how the combination of this unique model with proteomic profiling, signaling network analysis, and machine learning can lead to the identification of a new, potent drug combination for TNBC treatment, based on WEE1 and BCL-XL targeting. Our findings may be particularly relevant from a translational perspective, considering that agents targeting WEE1 or BCL-XL are already in phase I/II clinical trials.

## 5. Experimental Section

### Reverse phase protein array (RPPA)

Protein lysates of dissected mammary gland tumors (n=24), control mammary glands (MMTV and *MMTV-R26*^*Met*^), and *MMTV-R26*^*Met*^ MGT cells either not treated or treated for 12hrs with A1155463 (1μM), Adavosertib (3μM), or A1155463 + Adavosertib (1μM, 3μM) were prepared according to the MD Anderson Cancer Center platform instructions. Samples were screened with 426 antibodies to identify signaling changes in protein expression and phosphorylation levels.

### Bioinformatic analysis

Random Forest was performed using the randomForest package in R. RPPA data for 152 TNBC patients in the TCGA dataset (TCPA: The Cancer Proteome Atlas) was split into training (80%) and test (20%) sets. The expression levels of 105 proteins (protein without missing data in the TCGA and whose expression was also evaluated in our RPPA) were scaled to have an average of 0 and standard deviation of 1, and were used to train a random forest model for TNBCtype-4 classification by optimizing the number of proteins randomly selected at each split, using 10-fold cross validation. The model with the highest accuracy was validated on the test set, and used to predict the classification of the mice tumors to the four TNBC subtypes. Hierarchical clustering of the RPPA data and partition clustering were performed and visualized using the gplots and Factoextra packages in R.

### Statistical analysis

Data are presented as the median or as the mean ± standard error of the mean (s.e.m.), according to sample distributions. For two sided comparisons, we used unpaired Student’s t test for data showing normal distributions and Wilcoxon test in other situations. For multiple comparisons, ANOVA test followed by Tukey test were used. ANOVA to analyze the RPPA outcomes was performed using the Partek Genomic Suite. P values are indicated in figures. The cumulative overall disease-free survival rates were calculated using the Kaplan-Meier method. P < 0.05 was considered significant.

## Supporting Information

Supporting information is available from the Wiley Online Library

## Acknowledgments

We thank: all members of our labs for helpful discussions and comments; R. Dono and F. Helmbacher for extremely valuable feedback on the study; A. Furlan for initial work with *MMTV-R26*^*Met*^ mice; S. Richelme for *in vivo* bioluminescence imaging reported in Fig. 1B and for work on a first cohort of mice; E. Marechal for her contribution to the *MMTV-R26*^*Met*^ characterization; M. Buferne for *in vivo* bioluminescence imaging; the animal house platform for excellent help with mouse husbandry. This research was supported by a grant from the Ministry of Foreign Affairs and International Development (MAEDI) and the Ministry of National Education, Higher Education and Research (MENESR) of France and by the Ministry of Science and Technology of Israel (Grant #3-14002) to F.M. and S.L. F.A. was supported by the Higher Education Commission (HEC) of Pakistan. U.A.K. was supported by the Rising Tide Foundation Cancer Research Postdoctoral Fellowship, the Dean Fellowship of the Faculty of Biology of the Weizmann Institute of Science (WIS), and the Postdoctoral Fellowship of the Swiss Society of Friends of the WIS. J-P.B. is a scholar of Institut Universitaire de France. V.G. is supported by the “Ligue Nationale Contre le Cancer” (Equipe Labellisée). The contribution of the Region Provence Alpes Côtes d’Azur and of the Aix-Marseille University to the IBDM animal facility and of the France-BioImaging/PICsL infrastructure (ANR-10-INBS-04-01) to the imaging facility are also acknowledged. The funders had no role in study design, data collection and analysis, decision to publish or preparation of the manuscript.

## Conflict of interest

The authors declare no conflict of interest.

## Notes

### Competing Interest Statement

The authors have declared no competing interest.

