## Supporting text for "Modeling Heterogeneity of Triple-Negative Breast Cancer Uncovers a Novel Combinatorial Treatment Overcoming Primary Drug Resistance"

**Supplementary Experimental Section**

*Ethics statement:* All procedures involving the use of animals were carried out in accordance with the European Community Council Directive of 22 September 2010 on the protection of animals used for experimental purposes (2010/63/EU). The experimental protocols were performed according to the institutional Ethical Committee guidelines for animal research (Comité d'éthique pour l'expérimentation animale – Comité d'éthique de Marseille) and in compliance with the French law under the agreement number D13-055-21, delivered by the "Préfecture de la Région Provence-Alpes-Côte-d'Azur et des Bouches-du-Rhône". Mice were housed under pathogen-free conditions in enriched cages, with a light/dark cycle, and fed ad libitum according to Safe Complete Care Competence (SAFE A04). The mouse project authorization of the Maina laboratory is: APAFIS #8214-2016121417291352.v5, delivered by the "Ministère de l'Enseignement Supérieur, de la Recherche et de l'Innovation".

*Mice: R26*$^{stopMet}$ mice (international nomenclature *Gt(ROSA)26Sor*$^{tm1(Actb\text{-}Met)Fmai}$), and *R26*$^{stopMet\text{-}Luc}$ mice (international nomenclature *Gt(ROSA)26Sor*$^{tm1(Actb\text{-}Met\text{-}IRES\text{-}Luc)Fmai}$) carrying a conditional mouse-human chimeric *Met* transgene in the *Rosa26* locus was previously reported[1,2]. In both lines, expression of the MET$^{tg}$ (with or without Luciferase) was conditioned by the Cre-mediated removal of a LoxP-stop-LoxP cassette. *MMTV-R26*$^{Met}$ and *MMTV-R26*$^{Met\text{-}Luc}$ transgenic mouse lines were generated by crossing the *R26*$^{stopMet}$ or *R26*$^{stopMet\text{-}Luc}$ mice, respectively, with the *MMTV-Cre* line (B6129-Tgn(MMTV-Cre)4Mam) obtained from the Jackson Laboratory. All animals were maintained in a mixed genetic background (50% 129/Sv, 50% C57/Bl6). Mice were genotyped by PCR analysis of genomic DNA as described elsewhere[1,2]. Since it has been well established that mammary gland tumor formation is accelerated in multiparous females [3], *MMTV-R26*$^{Met}$ mice were maintained in constant breeding. Eight week-old nude mice (Rj:NMRI-Foxn1 nu/nu; JanvierLabs) were used for xenograft studies. Healthy 8-week-old female NOD.Cg-Prkdc $^{scid}$/J (NSG) mice, used for orthotopic studies, were bred in the animal facility of the Cancer Research Center of Marseille (CRCM) and maintained under specific pathogen-free conditions with sterilized food, water provided ad libitum and on a 12-h light and 12-h dark cycle. Orthotopic experiments were approved by animal ethics committees (APAFIS#13349-2018013116278149 v2).

*In vivo bioluminescence imaging:* Imaging was conducted as previously described[4]. Briefly, anesthetized *MMTV-R26$^{Met\text{-}Luc}$* mice were injected intraperitonealy with luciferin (3 mg/mouse). Analysis of the luminescent images was performed using Berthold Technologies software.

*Immunohistochemistry:* For histopathological analysis, control mammary glands or dissected tumors were fixed for 4hrs in 4% paraformaldehyde (PFA, Sigma), processed for paraffin embedding, and sectioned. They were then either stained with hematoxylin, and counterstained with eosin (H&E) or subjected to immunostaining with antibodies directed against estrogen receptor (ER), progesterone receptor (PR), human EGFR2 (HER2), or Ki67 (antibodies, and dilutions used are listed in Table S12).

*Cell lines: MMTV-R26$^{Met}$* MGT cell lines were derived from independent *MMTV-R26$^{Met}$* tumors. To establish *MMTV-R26$^{Met}$* MGT cell lines, *MMTV-R26$^{Met}$* tumors were dissected, and chopped into 1 mm3 pieces. Cells were dissociated for 30-40 min at 37°C with type II collagenase (1mg/ml, ThermoFisher Scientific) and DNase I (20µg/ml, Roche) in DMEM/F12 (Dulbecco's modified Eagle's media/F12, 1/1, ThermoFisher Scientific) complemented 10% fetal bovine serum (FBS, ThermoFisher Scientific), penicillin-streptomycin (P/S, 100 U/ml/0.1 mg/ml, ThermoFisher Scientific), and fungizone (25µg/ml, Sigma). The cell suspension was then passed through a 40µm nylon cell strainer to remove aggregates, and cells were seeded in complete DMEM/F12 medium (DMEM/F12, supplemented with 10% FBS, P/S, glutamine (2mM, ThermoFisher Scientific), glucose (0.25%, Sigma), insulin (10 µg/ml, Sigma), transferrin (10µg/ml, Sigma), sodium selenite (5ng/ml, Sigma), hydrocortisone (0.5 µg/ml, Sigma), EGF (20ng/ml, Roche), and HGF (10 ng/ml, Peprotech), and cultured in this complete medium in a humidified incubator at 37C° in a 5% $CO_2$ atmosphere. The establishment of normal mammary epithelial cell culture was performed as described above. Mammary glands were dissected from

*MMTV-R26$^{Met}$* mice not carrying tumor.

Human TNBC (MDA-MB-231, MDA-MB-468, SUM-159, Hs578t, HCC-1937 and BT-549) and non-TNBC (MCF-7, SKBR-3, and BT-474) cell lines were obtained from the American Type Culture Collection (ATCC). All human breast cancer cells were grown in DMEM/F12 medium supplemented with P/S, glutamine (2mM), sodium pyruvate (1mM, ThermoFisher Scientific), non-essential amino acids (ThermoFisher Scientific), and insulin (10µg/ml; Sigma).

MGT11$^{Luc}$ cells were generated by lentivirus infection of the parental *MMTV-R26$^{Met}$* MGT11 cells. Briefly, the pHAGE PGK-GFP-IRES-LUC-W lentivirus construct, which allows expression of luciferase and GFP in the infected cells, was used to produce the lentivirus and perform infection. Infected cells were grown in complete DMEM/F12 medium. GFP-positive cells were then sorted by FACS, amplified, and tested for their bioluminescence.

All cells were tested by PCR-based assay to assess their mycoplasma-free status.

*Drugs:* Drug concentration and sources are reported in Table S13. Calculation of the Synergy maps and Bliss score has been done using online SynergyFinder tool v1.0[5] using "Viability" parameter as readout and "Bliss Method" with correction activated.

Synergy or additive effects of drug combinations were measured by employing the CompuSyn software v1.0 using the Chou-Talalay equation. Combination index CI<1 indicates synergism, CI<0.5 indicates strong synergism, CI=1 means additive effect and CI>1 stands for antagonism.

*Cell viability assay: MMTV-R26$^{Met}$* MGT and human breast cancer cells were plated in 96-well plates at 10,000 cells per well (150µl media/well). After 24hrs, cells were treated with single or combined drugs at the indicated concentrations (Table S13) and cell viability was assessed 48hrs later using the Cell Titer Glo Luminescent Assay (Promega) by detecting the luminescent signals with a luminometer microplate reader (Berthold). Concerning cell death rescue experiments, *MMTV-R26$^{Met}$* MGT and MDA-MB-231 cells were pre-treated for 1hr with inhibitors of apoptosis (Z-VAD-FMK) or ferroptosis (Ferrostatin-1) at the indicated concentrations, then for additional 24hrs in the absence or presence of A-1155463 (0.3µM) and Adavosertib (3µM). Effects of cell death inhibitors on specific death inducers were assessed as positive control. Data are mean values of at least three independent experiments done in triplicates.

*Cell cycle analysis by flow cytometry: MMTV-R26$^{Met}$* MGT cells were cultured in standard conditions before cell cycle analysis of non-treated cells. Regarding cell cycle analysis after drug treatment, cells were treated with vehicle, A1155463 (1µM), Adavosertib (3µM), alone or in combination for 12hrs. After trypsinisation, cells were resuspended at $10^6$ cells per condition. Single cell suspensions were first incubated with the antibodies indicated in Table S12 and Fixable Viability Dye eF506 (1:1000, eBioscience) in PBS-0.5% BSA for 20 min at 4°C. Cells were then fixed and permeabilized using eBioscience™ Foxp3 / Transcription Factor Staining Buffer Set, according to manufacturer's instructions. After centrifugation, cells were resuspended in permeabilization solution containing anti-Ki67antibody, propidium iodide (PI, 40µg/ml, ThermoFisher Scientific) supplemented with RNaseA (50µg/ml, ThermoFisher Scientific). Cells were incubated 20 minutes at room temperature, washed and rapidly analyzed by a BD LSR-II FACS (lasers 405, 488, 561, and 633nm). Cell cycle distribution was analyzed with the BD-Diva version 8.0.1 software. Three independent experiments were performed.

*Apoptosis assay by flow cytometry:* Quantification of apoptosis was assessed by Annexin V-PI staining, using the eBioscience™ Annexin V Apoptosis Detection Kit FITC, according to the manufacturer's instructions. Briefly, harvested cells were washed in cold PBS and incubated for 15 minutes at room temperature with Annexin V-FITC / PI (1µg/ml) solution prepared in binding buffer. Samples were rapidly analyzed by a BD LSR-II FACS (lasers 405, 488, 561, and 633nm). Results were analyzed with BD-Diva version 8.0.1 software. Four independent experiments were performed.

*Cell migration assay:* MGT cells ($10^4$) were seeded in the upper compartment of an 8µm-pore Boyden-like chamber (Transwell, Corning) in 200µl of DMEM/F12 complete medium supplemented with 0.5%FBS. To create a chemo-attractant gradient, the bottom chambers were filled with 700µl of complete DMEM/F12 medium containing 10%FBS, 20ng/ml EGF, 10ng/ml HGF. Media also contained Cytosine arabinoside (AraC, 10µM) to prevent cell proliferation. After 36hrs of incubation at 37°C, non-migrating cells present on the upper surface of the filter were removed with a cotton swab. The invading cells located on the underside were fixed with 4% PFA, and nuclei were stained with DAPI. The number of migrating cells was directly analyzed under a Zeiss Axiophot fluorescent microscope, and determined in at least five fields of each filter. Experiments were performed in triplicate in three independent experiments.

*Tumor spheroid forming assay:* Cells were seeded in 35mm low attachment plates at a density of 25,000 cells/dish in DMEM/F12(1:1) medium supplemented with P/S (50mg/ml), glutamine (2mM), bovine serum albumin (BSA, 0.01%, Sigma), B27 without vitamin A (2%, ThermoFisher Scientific), N2 supplement (1%, ThermoFisher Scientific), insulin (10µg/ml), hydrocortisone (0.5 µg/ml), EGF (20ng/ml), HGF (10 ng/ml), and bFGF (10ng/ml, ThermoFisher Scientific). Medium was changed every 2 days. After 10 days, primary spheres were dissociated into single cells and re-plated at the same density as previously described. Subspheroid forming assay (also called passage) was repeated every 10 days. Brightfield pictures of the whole dishes were acquired using an inverted Zeiss Axiophot microscope. Number and size of tumor spheroids were determined using the ImageJ software. Values are expressed as mean ± SEM of three independent experiments.

*β-Gal and immunofluorescence staining on MGT cells:* For β-Gal staining, cells cultured on glass coverslips were fixed for 2 min in PBS, 25% Gluteraldehyde, 37% formaldehyde, then washed with PBS, and incubated with the staining solution (1mg/ml X-Gal, 0.5mM K3K4, 2mM MgCl2 in PBS) for 4hrs at 37°C. Cells were then fixed with 4% PFA, counterstained with eosin, and mounted on glass microscopic slides with Eukitt hardening solution.

For immunofluorescence staining, cells cultured on coverslips and fixed with 4% PFA were permeabilized for 15 min in PBS/TritonX-100 (PBST) solution (see TritonX-100 concentrations in Table S12), incubated for 1hr in blocking solution (PBST, 10% BSA), then overnight at 4°C with primary antibodies (MET, $pY_{1234/1235}$MET, Ki67) diluted in the above blocking solution. The next day, cells were washed 4 times with PBST, subsequently incubated for 1hr with fluorescence-labelled secondary antibodies (Table S12), incubated for 15 min in presence of DAPI (1:1000), then mounted on glass microscopic slides with Prolong Gold antifade reagent (ThermoFischer Scientific). Images were taken with a Zeiss Axiophot fluorescent microscope.

*In vivo tumorigenesis assays:* Xenografts. To assess the in vivo tumorigenicity of the *MMTV-R26$^{Met}$* MGT cell lines, cells ($5x10^6$) were resuspended in 200µl of a PBS:Matrigel (1:1) solution (CorningBV), and injected subcutaneously into both flanks of nude mice (n=4 mice / cell line). Animals were sacrificed when the tumor volume reached a maximum of 2000 $mm^3$. Tumor volume, measured with a caliper, was determined by the formula: $(L \times W^2)/2$. L: length; W: width.

Orthotopic injections. To evaluate the in vivo efficacy of the drug combination, MGT11$^{Luc}$ ($1x10^5$) cells, suspended in 50% phenol red–free Matrigel (Becton Dickinson Bioscience), were grafted into the fourth mammary fat pad of NSG mice (TrGET Preclinical Platform). After seven days of graft, mice were randomly divided into two groups (n=8 per group). Starting from the eighth day post-graft, control vehicle (2% DMSO, 30% PEG300, 5% Tween-80) or drug combination were administered (A1155463, 5mg/Kg, intraperitoneally; Adavosertib, 60mg/Kg, *per os*) for a total of 29 days. Mice body weight was checked daily, before drug administration. Tumor volume was measured every three days and calculated with the formula: (3.14 x Length x Width x Height)/6. The experiment was terminated when tumor volume reached a maximum of 1500$mm^3$. After completion of the study, lung luminescence was assessed following addition of endotoxin-free luciferin (30 mg/kg) and autopsy of mice. Bioluminescence analysis was performed using Optima-PhotonIMAGER (Biospace Lab).

*Quantitative RT-PCR analysis:* Total RNA was extracted from tissues or cells using the RNeasy Mini Kit (Qiagen) as previously described[6]. DNase (RNase-free DNAse I Set, Qiagen) treatment was included to avoid possible genomic DNA contamination. cDNA was synthesized using a Reverse Transcription Kit (iScript Reverse Transcription Supermix, Bio-Rad). Real-time PCR reactions were performed in a qPCR

CFX 96 apparatus (Bio-Rad), using the SYBR® Green detection method (SYBR GreenER qPCR SuperMix, ThermoFisher Scientific), and specific primers (0.1µM; primer sequences are listed in Table S14). mRNA expression levels were normalized to the *Beta-2-microglobuline* (*B2M*) housekeeping gene, and analyzed using the $2^{-\Delta\Delta Ct}$ method. All reactions were run in triplicate and repeated in three independent experiments.

*Western blotting:* Protein extracts were biochemically analyzed as previously described[7]. Briefly, *MMTV-R26*$^{Met}$ MGT cells were lysed in EMB lysis buffer (1% Triton X-100, 50mM HEPES, 1mM EGTA, 150mM NaCl, 1.5mM MgCl2, 10% glycerol, 10mM NaF, 1mM NaPP, 1mM $Na_3VO_4$, 10mM β-glycerophosphate, 5µg/ml leupeptin, 5 µM pepstatin A, 2µg/ml aprotinin, 5mM benzamidin, 1mM phenylmethylsulfonyl fluoride (PMSF), and centrifuged at 14,000 rpm for 15 min at 4 °C. Protein concentration was measured by Bradford assay (Bio-Rad), and equal amounts of proteins were separated on 10% polyacrylamide gels or SDS-PAGE (for phospho-ATM).

For nuclear and chromatin extracts, cells were pelleted, washed in PBS, swollen in hypotonic buffer (20mM HEPES pH 7.9, 1.5mM MgCl2, 10mM KCl, 1mM DTT, 1mM PMSF, 1x protease Inhibitor cocktail (Sigma)) on ice, then centrifuged at 1500rpm for 5min. Nuclei were resuspended in nuclear extract low salt buffer (20mM HEPES pH 7.9, 350mM sucrose, 1.5mM MgCl2, 150mM KCl, 0.5mM EDTA, 0.2% NP-40, 1mM PMSF, 1x protease Inhibitor cocktail) for 30 min on ice. The nucleoplasmic extract was recovered in the supernatant after centrifugation at 3000rpm for 10min. Residual nuclei were then resuspended in nuclear extract buffer (20mM HEPES pH 7.9, 350mM sucrose, 1.5mM MgCl2, 300mM KCl, 0.5mM EDTA, 0.2% NP-40, 1mM PMSF, 1x protease Inhibitor cocktail). After centrifugation at 3000rpm for 10min, the nuclear extract (supernatant) was separated from the chromatin (pellet). DNA from pellet was digested with Universal Nuclease (PierceTM) in nuclear extract high salt buffer (20mM HEPES pH 7.9, 350mM sucrose, 1.5mM MgCl2, 600mM KCl, 0.5mM EDTA, 0.2% NP-40, 1mM PMSF, 1x protease Inhibitor cocktail) for 30 min on ice. Protein concentration was measured by Qubit Protein Assay Kit (ThermoFisher) and equal amounts of proteins were separated on 15% polyacrylamide gels.

Membranes were incubated in blocking buffer (5% milk, PBS, 0.1% Tween-20, 50mM NaF), then analyzed using standard procedures. PageRuler Plus Prestained protein ladder (ThermoFisher Scientific) was used as a molecular weight marker. Pictures of the membranes stained with Ponceau (Sigma) and immunoblotted were taken through BioRad ChemiDoc imager. Antibodies used are reported in Table S12.

**Author contributions**

F.L.: designed and supervised studies, performed the majority of the experiments, data analysis, and interpretation; contributed to write the manuscript.

F.A.: performed the majority of the experiments under supervision of F.L., data analysis, and interpretation.

Y.V.: performed computational work with mouse and human TNBC databases, machine learning analyses, and interpretation; provided inputs on studies and on the manuscript.

O.C.: contributed to cell viability and biochemical studies; data processing.

F.D.: contributed to computational work with drug response data on cells, data analysis, and interpretation.

A.K.M.: performed histological analysis on *MMTV-R26*$^{Met}$ samples, data analysis, and interpretation; provided input on the manuscript.

U.A.K.: performed histological analysis on *MMTV-R26*$^{Met}$ samples, data analysis, and interpretation; provided input on the manuscript.

A.L.B.: performed FACS studies, data analysis, and interpretation.

E.J.: performed orthotopic studies and data analysis.

R.C.: performed orthotopic studies and data analysis.

C.C.: performed cell fractionation studies and data analysis.

E.C.J.: supervised and performed histological analysis on *MMTV-R26*$^{Met}$ samples, data analysis, and interpretation; provided input on the manuscript.

G.B.M.: contributed to RPPA studies; provided input on the manuscript.

V.G.: provided input on studies and contributed to write the manuscript.

J.P.B.: provided input on studies and contributed to write the manuscript.

S.L.: designed and supervised studies on histological and RPPA analysis; provided input on studies; contributed to interpret data and to write the manuscript.

F.M.: designed and supervised the study, contributed to experimental work, analysed and interpreted data, ensured financial support, and wrote the manuscript.

**Supplementary Figure Legends**

**Figure S1.** Expression of the wild-type *Met* transgene in the mouse mammary gland leads to tumor formation. a) Western blot analysis of total protein extracts from either mammary gland or liver from adult wild-type (wt), *Alb-R26*$^{Met}$, *MMTV-R26*$^{Met}$, or *R26*$^{stopMet}$ mice. The MET$^{tg}$ is specifically expressed in the mammary gland of *MMTV-R26*$^{Met}$ mice. *Alb-R26*$^{Met}$ mice were used as control for expression of MET$^{tg}$ in the liver[4]. Expression levels of ERKs were used as a loading control. b) Kaplan-Meier analysis of mammary gland tumor incidence in *MMTV-ErbB2* mice kept in the same genetic background as the *MMTV-R26*$^{Met}$ (*MMTV-ErbB2*$^{mix}$; data already shown in Figure 1d), and in FVB/mix background (*MMTV-ErbB2*)[3]. c) Example of *MMTV-R26*$^{Met}$ mammary gland tumor and lung metastasis. Hematoxylin and eosin (H&E) staining of corresponding sections are shown on the bottom. d) Heatmap reporting upregulated (red) and downregulated (blue) signals in *MMTV-R26*$^{Met}$ tumors. e) Graph reporting total *Met* mRNA levels (endogenous plus exogenous, using primers in the *Met* extracellular domain (*Met EXT*)) in *MMTV-cre* (n=5) and *MMTV-R26*$^{Met}$ pre-tumorigenic (n=6) glands, and *MMTV-R26*$^{Met}$ tumors (n=24). f) Random Forest method, a very robust machine learning method for classification problems, was used to build the model predicting TNBC subtypes. TCPA was used as training (80%) and validation (20%) sets. The graph reports the accuracy (repeated cross-validation) of the number of proteins randomly selected at each split of the trees. Based on the high accuracy achieved, this model was used for analysis of the *MMTV-R26*$^{Met}$ tumors.

**Figure S2.** Biological properties of *MMTV-R26*$^{Met}$ MGT cell lines. a) Efficiency of Cre recombination (β-Gal staining, top images), and immunohistochemical analysis of MET$^{tg}$ (detected by human MET antibodies) and phosphoY$_{1234/1235}$-MET (pMET) in *MMTV-R26*$^{Met}$ cell lines. Lack of β-Gal expression, due to the excision of the LacZ-stop cassette, reveals that Cre recombination has occurred in cells. Percentages of cells with efficient recombination and positive for MET$^{tg}$ or phospho-MET are indicated on the corresponding images. Nuclei stained with DAPI are in blue. Scale bars: 100μm b) Graph reporting total *Met* mRNA levels (endogenous plus exogenous) in the 6 *MMTV-R26*$^{Met}$ MGT cell lines compared to normal mammary epithelial cells (control). c) Western blot analysis of total protein extracts from MGT2, MGT4, and MGT9 cell lines. The quantity of total protein extracts loaded in each lane is indicated. Ponceau staining was used as a loading control. The $MET^{25H2}$ antibodies recognize both the endogenous MET and the $MET^{tg}$. d-e) Quantification (d) and representative images (e) of cell proliferation capacity determined by analysis of the percentage of Ki67-positive cells compared to the total number of cells. The mean percentages are indicated on the corresponding images. Statistical analyses shown here were performed by using MGT2 as the control cell line. All statistical analyses performed by comparing one cell line to another are reported in Table S15. Scale bar: 100µm. f) Cell cycle distribution as measured by flow cytometry using PI and Ki67 staining. g) Western blot analysis of p53 expression levels in *MMTV-R26*$^{Met}$ MGT cell lines compared to normal mammary epithelial cells (control).

**Figure S3.** Multiple signaling networks are enriched in the *MMTV-R26*$^{Met}$ cell lines. a) Heatmap reporting expression or phosphorylation levels of proteins in the *MMTV-R26*$^{Met}$ cell lines as determined by RPPA. Note the hierarchical clustering of MGT4, MGT9, and MGT11 compared to MGT13, among the tumorigenic cell lines. Red: upregulated; blue: downregulated. b) Proteomic profiles of cells from the different clusters identified by PCA were compared one to another, as indicated. Enrichment analyses were done by using the Enrichr software. Histograms show: i) enriched pathways according to the Kyoto Encyclopedia of Genes and Genomes (KEGG) database; ii) cell signaling pathway enrichment using the WikiPathways database; iii) kinases identified based on their phosphorylation targets (Kinase Enrichment Analysis, KEA database); iv) enrichment according to the Jensen compartments database. In all panels, enrichments are ranked according to combined scores (values are indicated on the X axis). The 20 top ranked enrichments are presented. Note that the majority of the signaling changes are related to signals involved in DNA repair, cell cycle regulation, metabolism, and stemness.

**Figure S4.** A1155463 and Adavosertib exhibit synergistic effects on *MMTV-R26$^{Met}$* MGT cell lines. a-b) Representation of the drug screen outcomes performed on the MGT4 cell line, highlighting single and combined treatments tested. Drugs were used at a concentration of 3µM (a) and 10µM (b). Numbers indicate the percentage of viable cells in the presence of drugs compared to controls (untreated cells), and labelled by the green-to-red color code. Values are expressed as the mean ± s.e.m. Grey squares correspond to untested drug effects. Note that targeting BCL-XL (with Wehi) and WEE1 (with Adavosertib) is highly deleterious (red color) for *MMTV-R26$^{Met}$* MGT4 cells compared to other drugs exhibiting no or minimal effects (green, yellow, orange). c) Data from cell viability assays of combined drug effects were used by the Compusyn software to simulate combination index (Y-axis) for each affected fraction (X-axis, from 0 to 1). Each black dot corresponds to a tested dose. Based on the combination index scores, combinations of BCL-XL and WEE1 inhibition resulted in strong synergistic interactions for high doses on MGT4, and for all doses tested for MGT9, MGT11, and MGT13. (d-f) MGT11 and MGT11$^{Luc}$ cells exhibit similar biological properties. (d) Viability assay performed on MGT11 and MGT11$^{Luc}$ cells when exposed to A1155463 (A11, 0.3µM) alone or in combination with Adavosertib (Adav, 3µM). Percentage of cell viability in the presence of drugs compared to untreated cells is reported using the defined color code. Values are expressed as the mean ± s.e.m. (e) Graph representing the percentage of cells in each phase of the cell cycle as determined by flow cytometry using PI and Ki67 staining. (f) Tumor sphere assay performed with MGT11 and MGT11$^{Luc}$ cells. Values are expressed as means ± s.e.m. For all these *in vitro* assays, three independent experiments were performed. g) Effect of the combined A1155463+Adavosertib treatment (A11+Adav) on lung metastasis formation monitored using bioluminescence imaging in the treated mice. Quantification of the normalized photon flux in A11+Adav-treated mice compared to the control mice (vehicle). Results show a trend of reduced metastasis formation in the group of mice treated with the drug combination, although not significant. Wilcoxon test was used for statistical analysis. Data represent mean ± SD ($n$ = 8). ns: P>0.05.

**Figure S5.** Signaling changes in MGT4, MGT9, and MGT13 cells when exposed to drugs targeting BCL-XL and WEE1. a-b) Western blot analysis of total protein extracts from MGT4, MGT9, and MGT13 cells treated for 12hrs with either A1155463 (A11, 1µM) or Adavosertib (Adav, 3 µM) alone or in combination.

**Supplementary Table Legends**

**Table S1.** Mammary gland (tumors and controls) used for histopathological and RPPA analyses.

**Table S2.** Antibodies used for RPPA analysis of *MMTV-R26*$^{Met}$ tumors and cells. RPPA analysis on tumors and non-treated MGT cells was performed using the 247 rabbit antibodies. RPPA analysis done on treated MGT cells included mouse and rabbit antibodies (n= 426).

**Table S3.** Molecular/signaling characteristics of *MMTV-R26*$^{Met}$ tumors identified through RPPA analysis.

**Table S4.** Molecular/signaling characteristics of *MMTV-R26*$^{Met}$ MGT cells identified through RPPA analysis.

**Table S5.** Comparison of RPPA outcomes of *MMTV-R26*$^{Met}$ cells and tumors belonging to "subtype A" versus "subtype B".

**Table S6.** Comparisons of RPPA outcomes of *MMTV-R26*$^{Met}$ cells belonging to "subtype A" and "subtype B" versus those from the non-tumorigenic cells.

**Table S7.** Changes in protein expression and/or phosphorylation levels occurring in *MMTV-R26*$^{Met}$ MGT4 cells when exposed to single or combined treatment targeting BCL-XL (A1155463) and WEE1 (Adavosertib).

**Table S8.** Statistical analysis of the MMTV-R26Met MGT cell lines cell cycle distribution.

**Table S9.** Statistical analysis of the in vitro tumorigenic capacity of the MMTV-R26Met cell lines (determined by the tumor sphere assay).

**Table S10.** Statistical analysis of the migrating capacity of the MMTV-R26Met cell lines.

**Table S11.** Statistical analysis of cell cycle distribution of MGT11 cells treated with A1155463, or Adavosertib, alone or in combination.

**Table S12.** Antibodies used in the study.

**Table S13.** Drugs used for cell viability assays, with the indicated targets and the concentrations used.

**Table S14.** Oligonucleotides used for RT-qPCR experiments.

**Table S15.** Statistical analysis of the proliferation capacity (mitotic index) of the MMTV-R26Met cell lines.
