## Supplementary figures and images for "Modeling Heterogeneity of Triple-Negative Breast Cancer Uncovers a Novel Combinatorial Treatment Overcoming Primary Drug Resistance"

### Supporting Figures

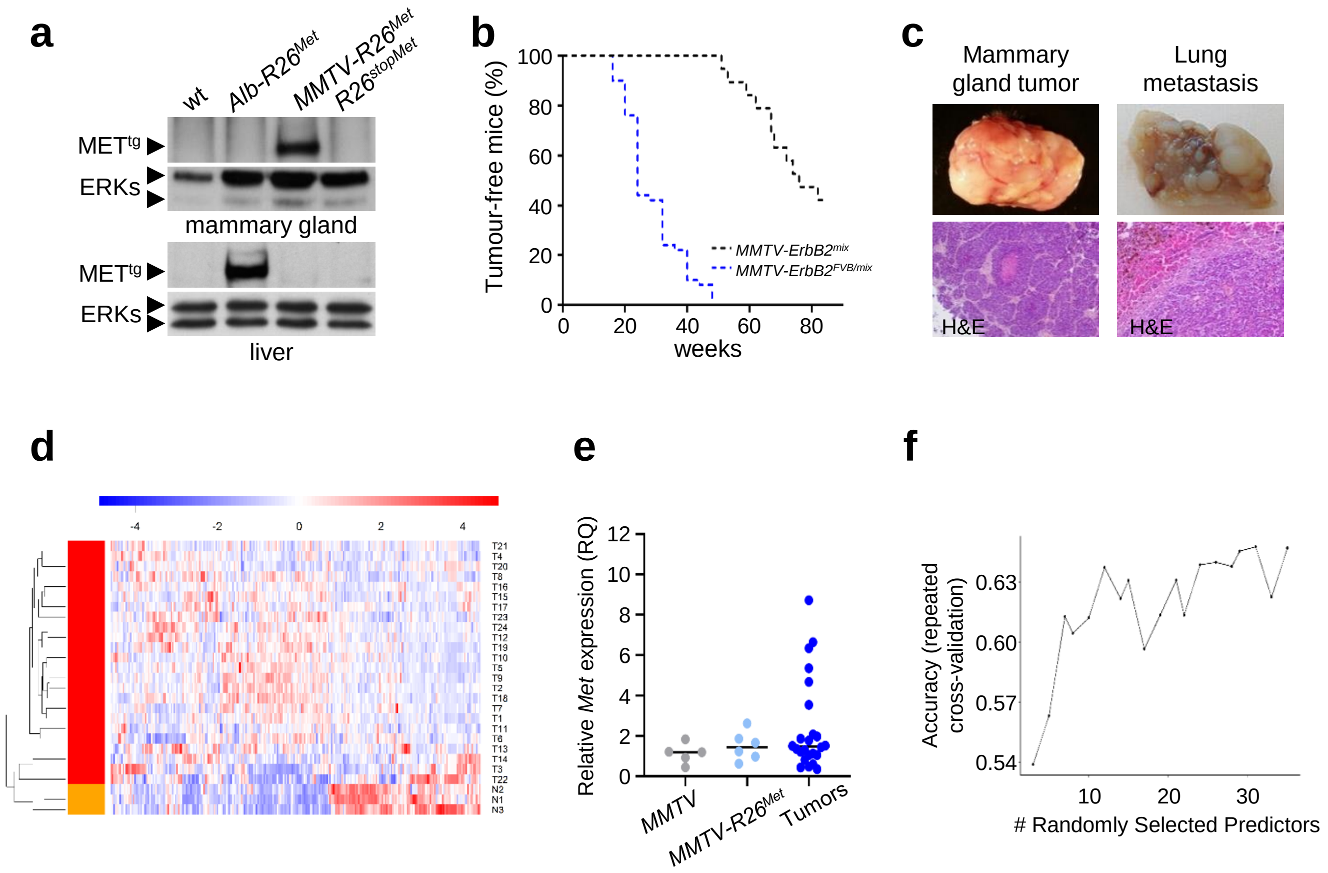

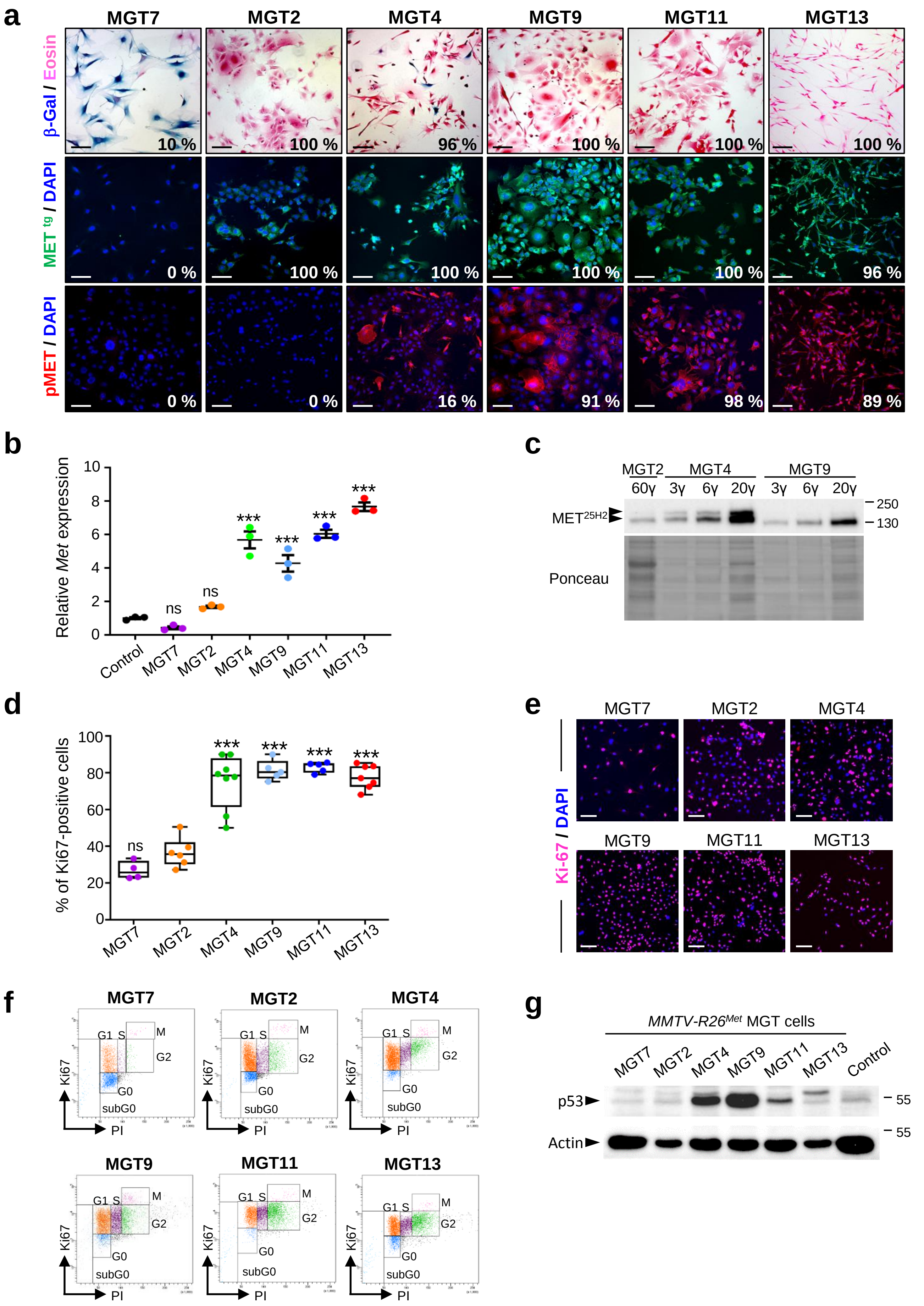

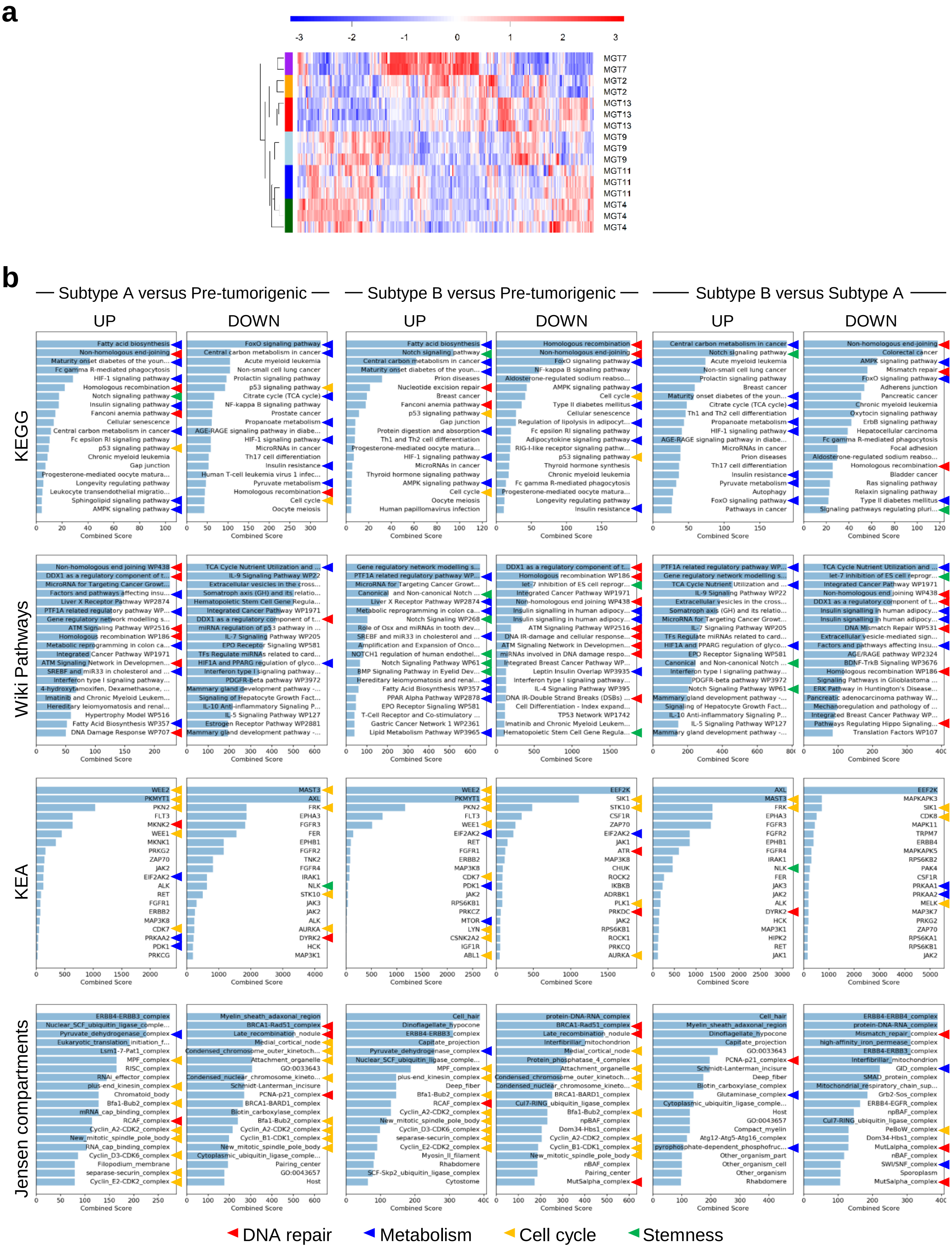

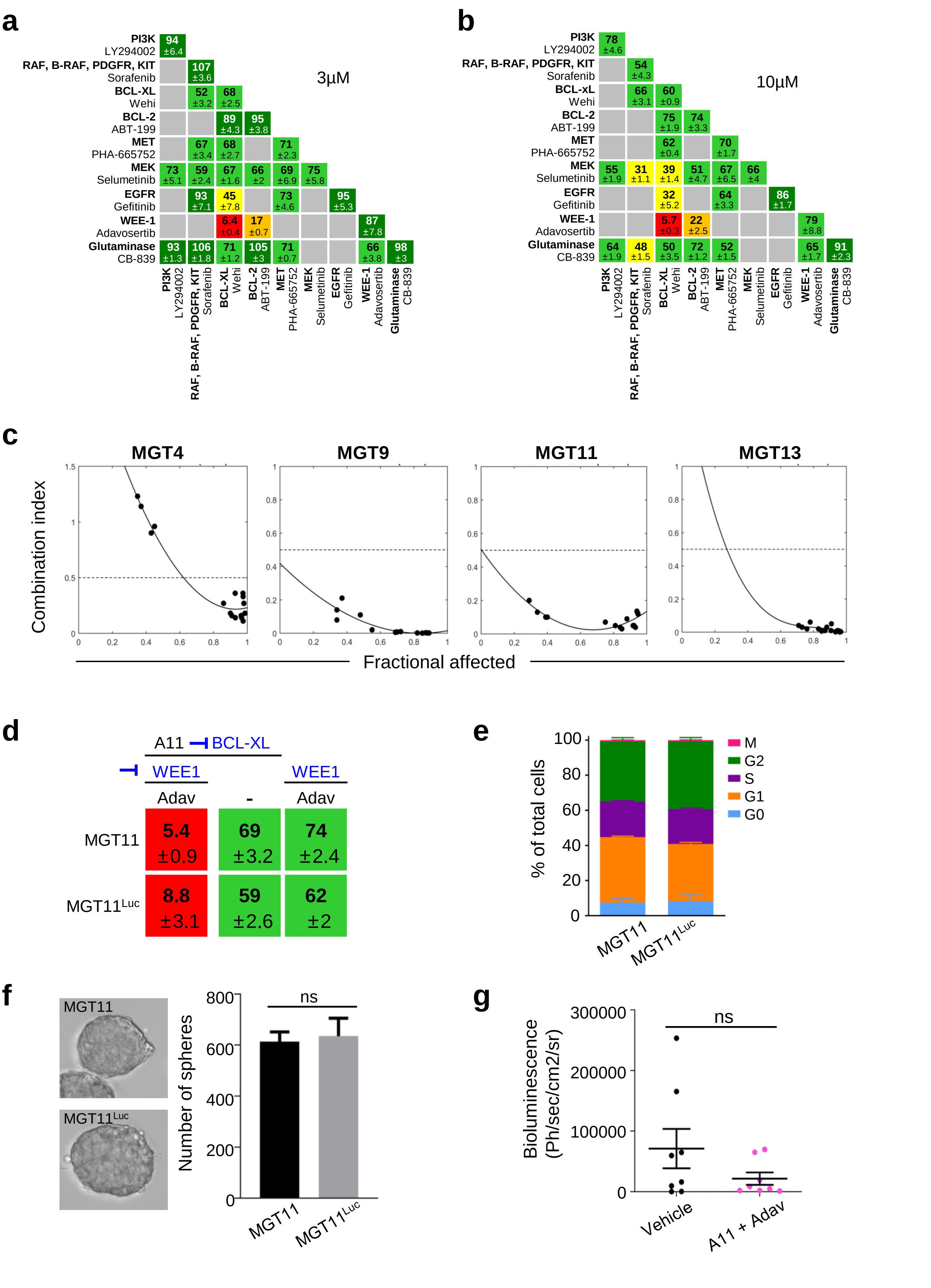

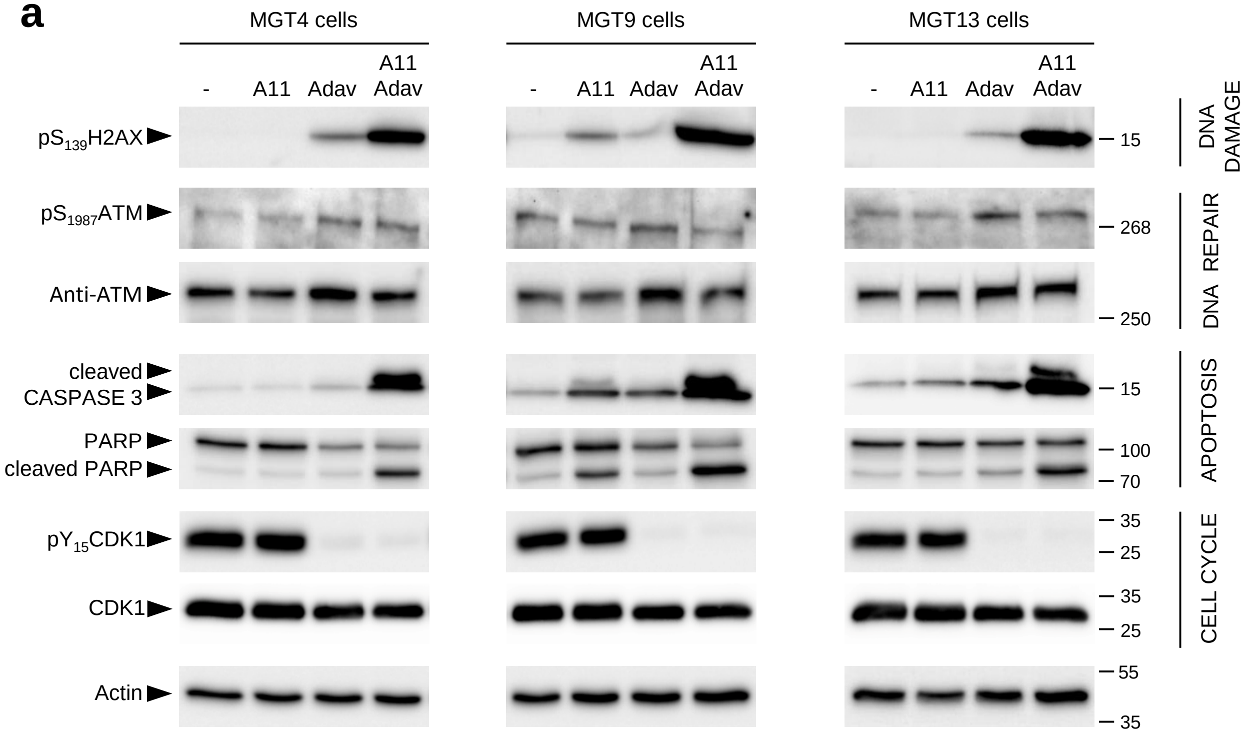

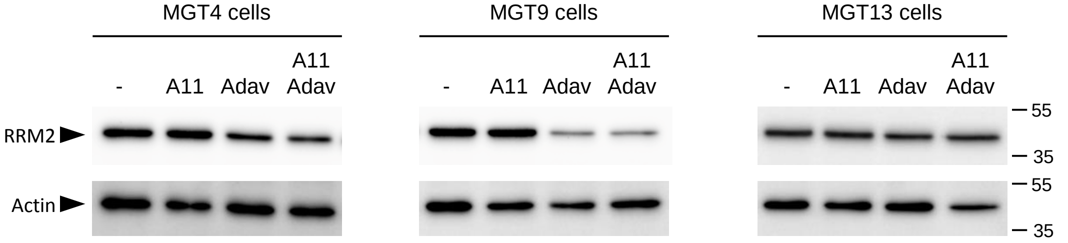
