## Supporting Tables for "Modeling Heterogeneity of Triple-Negative Breast Cancer Uncovers a Novel Combinatorial Treatment Overcoming Primary Drug Resistance"

**Table S1:** Mammary gland (tumors and controls) used for histopathological and RPPA analyses.

| Mouse # | Genotype | Gross aspect of gland | Histopathological analysis | | | | | | Lung metas-tases | RPPA (Tumor #) |
|---|---|---|---|---|---|---|---|---|---|---|
| | | | Tumor state | Grade | Necrosis | Prolif. status | ER/PR/ HER2 scoring | Met | | |
| 669 201 | *MMTV-R26*$^{Met}$ | Tumour | carcinoma | nd | nd | nd | 0 | nd | Yes | Yes (T1) |
| 669 352 | *MMTV-R26*$^{Met}$ | Tumour | carcinoma | nd | + | nd | 0 | nd | Yes | Yes (T2) |
| 669 713 | *MMTV-R26*$^{Met}$ | Tumour | carcinoma | nd | nd | nd | 0 | nd | _ | Yes (T3) |
| 669 163 | *MMTV-R26*$^{Met}$ | Tumour | carcinoma | nd | nd | nd | 0 | nd | _ | Yes (T4) |
| 669 789 | *MMTV-R26*$^{Met}$ | Tumour | carcinoma | nd | nd | nd | 0 | nd | Yes | Yes (T5) |
| 669 722 | *MMTV-R26*$^{Met}$ | Tumour | carcinoma | nd | nd | nd | 0 | nd | Yes | Yes (T6) |
| 669 175 | *MMTV-R26*$^{Met}$ | Tumour | carcinoma | nd | nd | nd | 0 | nd | _ | Yes (T7) |
| 669 715 | *MMTV-R26*$^{Met}$ | Tumour | carcinoma | nd | nd | nd | 0 | nd | _ | Yes (T8) |
| 671 802 (Tumor A) | *MMTV-R26*$^{Met}$ | Tumour | carcinoma | 3 | + | 3 | 0 | + | _ | Yes (T9) |
| 671 607 | *MMTV-R26*$^{Met}$ | Tumour | carcinoma | 3 | + | 3 | 0 | + | _ | Yes (T10) |
| 671 662 | *MMTV-R26*$^{Met}$ | Tumour | carcinoma | 3 | ++ | 2 | 0 | + | _ | Yes (T11) |
| 671 670 | *MMTV-R26*$^{Met}$ | Tumour | carcinoma | 3 | +(+) | 3 | 0 | + | Yes | Yes (T12) |
| 671 975 | *MMTV-R26*$^{Met}$ | Tumour | carcinoma | 2 | _ | 1 | 0 | + | _ | Yes (T13) |
| 672 961 | *MMTV-R26*$^{Met}$ | Tumour | carcinoma | 3 | ++ | 3 | 0 | + | _ | Yes (T14) |
| 671 711 | *MMTV-R26*$^{Met}$ | Tumour | carcinoma | 3 | ++ | 2 | 0 | + | _ | Yes (T15) |
| 672 046 | *MMTV-R26*$^{Met}$ | Tumour | carcinoma | 3 | _ | 2 | 0 | + | _ | Yes (T16) |
| 671 893 | *MMTV-R26*$^{Met}$ | Tumour | carcinoma | 3 | _ | 2 | 0 | + | Yes | Yes (T17) |
| 672 867 | *MMTV-R26*$^{Met}$ | Tumour | carcinoma | 3 | + | 2 | 0 | + | _ | Yes (T18) |
| 672 790 | *MMTV-R26*$^{Met}$ | Tumour | carcinoma | 2 | _ | 2 | 0 | + | _ | Yes (T19) |
| 672 955 | *MMTV-R26*$^{Met}$ | Tumour | carcinoma | 2-3 | + | 2 | 0 | + | _ | Yes (T20) |
| 672 941 | *MMTV-R26*$^{Met}$ | Tumour | carcinoma | 3 | _ | 2 | 0 | + | _ | Yes (T23) |

| | | | | | | | | | | |
|---|---|---|---|---|---|---|---|---|---|---|
| 672 876 | *MMTV-R26$^{Met}$* | Tumour | nd | nd | nd | nd | nd | nd | _ | Yes (T21) |
| 672 302 | *MMTV-R26$^{Met}$* | Tumour | nd | nd | nd | nd | nd | nd | _ | Yes (T22) |
| 672 870 | *MMTV-R26$^{Met}$* | Tumour | nd | nd | nd | nd | nd | nd | _ | Yes (T24) |
| 671 509 | *MMTV-R26$^{Met}$* | Tumour | carcinoma | 3 | + | 2 | 0 | + | _ | _ |
| 671 802 (Tumor B) | *MMTV-R26$^{Met}$* | Tumour | carcinoma | 3 | _ | 1 | 0 | + | _ | _ |
| 672 802 | *MMTV-R26$^{Met}$* | Tumour | carcinoma | 3 | + | 2 | 0 | + | _ | _ |
| 669 527 | *MMTV-R26$^{Met}$* | Tumour | carcinoma | nd | nd | nd | nd | nd | _ | _ |
| 669 355 | *MMTV-R26$^{Met}$* | Tumour | carcinoma | nd | nd | nd | nd | nd | _ | _ |
| 669 717 | *MMTV-R26$^{Met}$* | Tumour | carcinoma | nd | nd | nd | nd | nd | _ | _ |
| 669 854 | *MMTV-R26$^{Met}$* | Tumour | carcinoma | nd | nd | nd | nd | nd | _ | _ |
| 671 720 | *MMTV-R26$^{Met}$* | Tumour | nd | nd | nd | nd | nd | nd | _ | _ |
| 672 246 | *MMTV-R26$^{Met}$* | Tumour | nd | nd | nd | nd | nd | nd | _ | _ |
| 671 594 | *MMTV* | Normal | _ | nd | _ | nd | nd | - | _ | Yes (N1) |
| 671 600 | *MMTV* | Normal | _ | nd | _ | nd | nd | - | _ | Yes (N2) |
| 672 603 | *MMTV-R26$^{Met}$* | Normal | _ | nd | _ | nd | nd | nd | _ | Yes (N3) |

**Table S2:** Antibodies used for RPPA analysis of *MMTV-R26$^{Met}$* tumors and cells

| # | Official Ab Name | Ab Name Reported on Dataset | Gene Name | Company | Catalog # | Species | RPPA Dilution |
|---|---|---|---|---|---|---|---|
| 1 | 14-3-3 beta | 14-3-3-beta | YWHAB | Santa Cruz | sc-628 | Rabbit | 1:75 |
| 2 | 14-3-3 epsilon | 14-3-3-epsilon | YWHAE | Santa Cruz | SC-23957 | Mouse | 1:50 |
| 3 | 14-3-3 zeta | 14-3-3-zeta | YWHAZ | Santa Cruz | sc-1019 | Rabbit | 1:5000 |
| 4 | 4E-BP1 | 4E-BP1 | EIF4EBP1 | CST | 9452 | Rabbit | 1:100 |
| 5 | 4E-BP1 (phospho S65) | 4E-BP1_pS65 | EIF4EBP1 | CST | 9456 | Rabbit | 1:250 |
| 6 | 4E-BP1 (phospho T37/46) | 4E-BP1-pT37-T46 | EIF4EBP1 | CST | 9459 | Rabbit | 1:2000 |
| 7 | 53BP1 | 53BP1 | TP53BP1 | CST | 4937 | Rabbit | 1:300 |
| 8 | A1Up | UBQLN4 | UBQLN4 | Santa Cruz | sc-136145 | Mouse | 1:125 |
| 9 | Acetyl-CoA-Carboxylase | ACC1 | ACACA, B | Epitomics/ Abcam | 1768-1/ ab45174 | Rabbit | 1:1500 |
| 10 | Acetyl-CoA-Carboxylase (phospho S79) | ACC_pS79 | ACACA, B | CST | 3661 | Rabbit | 1:500 |
| 11 | ACSL1 (D2H5) | ACSL1 | ACSL1 | CST | 9189 | Rabbit | 1:500 |
| 12 | ACVRL1 | ACVRL1 | ACVRL1 | Epitomics/ Abcam | 2940-1/ ab108207 | Rabbit | 1:30 |
| 13 | ADAR1 | ADAR1 | ADAR | Abcam | ab88574 | Mouse | 1:100 |
| 14 | Akt | Akt | AKT1, 2, 3 | CST | 4691 | Rabbit | 1:7500 |
| 15 | Akt (phospho S473) | Akt_pS473 | AKT1, 2, 3 | CST | 9271 | Rabbit | 1:150 |
| 16 | Akt (phospho T308) | Akt_pT308 | AKT1, 2, 3 | CST | 2965 | Rabbit | 1:250 |
| 17 | Akt1 | Akt1 | AKT1 | CST | 2938 | Rabbit | 1:1000 |
| 18 | Akt1 (phospho S473) | Akt1_pS473 | AKT1 | CST | 9018 | Rabbit | 1:1000 |
| 19 | Akt2 | Akt2 | AKT2 | CST | 3063 | Rabbit | 1:3000 |
| 20 | Akt2 (phospho S474) | Akt2_pS474 | AKT2 | CST | 8599 | Rabbit | 1:1000 |
| 21 | Ambra1 (phospho S52) | Ambra1_pS52 | AMBRA1 | Millipore | ABC80 | Rabbit | 1:250 |
| 22 | AMPK alpha 2 (phospho S345) | AMPK-a2_pS345 | PRKAA1, 2 | Abcam | ab129081 | Rabbit | 1:200 |
| 23 | AMPKa | AMPKa | PRKAA1, 2 | CST | 2532 | Rabbit | 1:75 |

| | | | | | | | |
|---|---|---|---|---|---|---|---|
| **24** | AMPKa (phospho T172) | AMPKa_pT172 | PRKAA1, 2 | CST | 2535 | Rabbit | 1:100 |
| **25** | Androgen Receptor (D6F11) | AR | AR | CST | 5153 | Rabbit | 1:250 |
| **26** | Annexin I | Annexin-I | ANXA1 | BD Biosciences | 610066 | Mouse | 1:5000 |
| **27** | Annexin VII | Annexin-VII | ANXA7 | BD Biosciences | 610668 | Mouse | 1:20 |
| **28** | A-Raf | A-Raf | ARAF | CST | 4432 | Rabbit | 1:200 |
| **29** | A-Raf (phospho S299) | A-Raf_pS299 | ARAF | CST | 4431 | Rabbit | 1:25 |
| **30** | ARID1A | ARID1A | ARID1A | Sigma-Aldrich | HPA005456 | Rabbit | 1:1000 |
| **31** | ASNS | ASNS | ASNS | Sigma-Aldrich | HPA029318 | Rabbit | 1:500 |
| **32** | Atg3 | Atg3 | ATG3 | CST | 3415 | Rabbit | 1:72 |
| **33** | Atg4B | Atg4B | ATG4B | CST | 13507 | Rabbit | 1:200 |
| **34** | Atg5 | Atg5 | ATG5 | CST | 12994 | Rabbit | 1:1000 |
| **35** | Atg7 | Atg7 | ATG7 | CST | 8558 | Rabbit | 1:1000 |
| **36** | ATM | ATM | ATM | CST | 2873 | Rabbit | 1:250 |
| **37** | ATM (phospho S1981) | ATM_pS1981 | ATM | CST | 5883 | Rabbit | 1:20 |
| **38** | ATP5A | ATP5A | ATP5A | Abcam | ab14748 | Mouse | 1:500 |
| **39** | ATP5H | ATP5H | ATP5H | Abcam | ab110275 | Mouse | 1:30 |
| **40** | ATR | ATR | ATR | CST | 2790 | Rabbit | 1:30 |
| **41** | ATR (phospho S428) | ATR_pS428 | ATR | Abcam | ab178407 | Rabbit | 1:1000 |
| **42** | ATRX | ATRX | ATRX | Abcam | ab97508 | Rabbit | 1:300 |
| **43** | Aurora B/AIM1 | Aurora-B | AURKB | CST | 3094 | Rabbit | 1:38 |
| **44** | Axl | Axl | AXL | CST | 8661 | Rabbit | 1:500 |
| **45** | B7-H3 | B7-H3 | CD276 | CST | 14058 | Rabbit | 1:200 |
| **46** | B7-H4 | B7-H4 | VTCN1 | CST | 14572 | Rabbit | 1:50 |
| **47** | Bad (phospho S112) | Bad_pS112 | BAD | CST | 9291 | Rabbit | 1:50 |
| **48** | Bak | Bak | BAK1 | Epitomics/ Abcam | 1542-1/ ab32371 | Rabbit | 1:400 |
| **49** | BAP1 | BAP1 | BAP1 | Santa Cruz | sc-28383 | Mouse | 1:200 |
| **50** | Bax | Bax | BAX | CST | 2772 | Rabbit | 1:100 |
| **51** | b-Catenin | b-Catenin | CTNNB1 | CST | 9562 | Rabbit | 1:1500 |
| **52** | Bcl2 | Bcl2 | BCL2 | Dako | M0887 | Mouse | 1:50 |
| **53** | Bcl2A1 | Bcl2A1 | BCL2A1 | Abnova | PAB8528 | Rabbit | 1:250 |
| **54** | Bcl-xL | Bcl-xL | BCL2L1 | CST | 2762 | Rabbit | 1:100 |

| | | | | | | | |
|---|---|---|---|---|---|---|---|
| **55** | Beclin 1 | Beclin | BECN1 | ThermoFisher | PA1-16857 | Rabbit | 1:500 |
| **56** | beta Actin | b-Actin | ACTB | CST | 4970 | Rabbit | 1:50 |
| **57** | beta Catenin (phospho T41/S45) | b-Catenin_pT41_S45 | CTNNB1 | CST | 9565 | Rabbit | 1:30 |
| **58** | Bid | Bid | BID | CST | 2002 | Rabbit | 1:500 |
| **59** | Bim (C34C5) | Bim | BCL2L11 | Epitomics/ Abcam | 1036-1/ ab32158 | Rabbit | 1:400 |
| **60** | BiP/GRP78 | BiP-GRP78 | HSPA5 | BD Biosciences | 610978 | Mouse | 1:150 |
| **61** | BMK1/Erk5 (phospho T218/Y220) | BMK1-Erk5_pT218_Y220 | MAPK7 | Millipore | 07-507 | Rabbit | 1:500 |
| **62** | B-Raf | B-Raf | BRAF | CST | 14814 | Rabbit | 1:500 |
| **63** | B-Raf (phospho S445) | B-Raf_pS445 | BRAF | CST | 2696 | Rabbit | 1:75 |
| **64** | BRD4 | BRD4 | BRD4 | CST | 13440 | Rabbit | 1:1000 |
| **65** | CA9 (CAIX) | CA9 | CA9 | CST | 5649 | Rabbit | 1:200 |
| **66** | c-Abl | c-Abl | ABL1 | CST | 2862 | Rabbit | 1:100 |
| **67** | c-Abl (phospho Y412) | Abl_pY412 | ABL1 | CST | 2865 | Rabbit | 1:200 |
| **68** | Caspase 3 (cleaved asp175) | Caspase-3-cleaved | CASP3 | CST | 9661 | Rabbit | 1:500 |
| **69** | Caspase 7 (cleaved) | Caspase-7-cleaved | CASP7 | CST | 9491 | Rabbit | 1:60 |
| **70** | Caspase 8 | Caspase-8 | CASP8 | CST | 9746 | Mouse | 1:150 |
| **71** | Caspase 8 (cleaved asp391) | Caspase-8-cleaved | CASP8 | CST | 9496 | Rabbit | 1:500 |
| **72** | Caspase-3 | Caspase-3 | CASP3 | Epitomics/ Abcam | 1476-1/ ab32042 | Rabbit | 1:250 |
| **73** | Caveolin 1 | Caveolin-1 | CAV1 | CST | 3238 | Rabbit | 1:3000 |
| **74** | CD134/OX40 | CD134 | TNFRSF | Abcam | ab76000 | Rabbit | 1:100 |
| **75** | CD171 (L1) | CD171 | L1CAM | Biolegend | 826701 | Mouse | 1:1000 |
| **76** | CD20 | CD20 | MS4A1 | Epitomics/ Abcam | 1632-1/ ab78237 | Rabbit | 1:75 |
| **77** | CD26 | CD26 | DPP4 | Abcam | ab28340 | Rabbit | 1:1000 |
| **78** | CD29 | CD29 | ITGB1 | BD Biosciences | 610467 | Mouse | 1:30 |
| **79** | CD31 | CD31 | PECAM1 | Dako/ Fisher | M0823/ MS353S | Mouse | 1:25 |
| **80** | CD38 | CD38 | CD38 | Abcam | ab108403 | Rabbit | 1:250 |
| **81** | CD4 | CD4 | CD4 | Abcam | ab133616 | Rabbit | 1:500 |

| | | | | | | | |
|---|---|---|---|---|---|---|---|
| **82** | CD44 | CD44 | CD44 | CST | 3570 | Mouse | 1:20 |
| **83** | CD45 | CD45 | CD45 | DAKO/ ThermoFisher | M070129-2/ MS355P | Mouse | 1:1000 |
| **84** | CD49b | CD49b | ITGA2 | BD Biosciences | 611016 | Mouse | 1:50 |
| **85** | CD86 | CD86 | CD86 | Abcam | ab53004 | Rabbit | |
| **86** | Cdc2 (phospho Y15) | cdc2_pY15 | CDK | CST | 4539 | Rabbit | 1:38 |
| **87** | cdc25C | cdc25C | CDC25C | CST | 4688 | Rabbit | 1:250 |
| **88** | CDK1/2/3 (phospho T14) | CDK1_pT14 | CDK1, 2, 3 | Abcam | ab32384 | Rabbit | 1:1000 |
| **89** | CDKN2A/p16INK4a | p16INK4a | CDKN2A | Abcam | ab81278 | Rabbit | 1:500 |
| **90** | Chk1 | Chk1 | CHEK | CST | 2360 | Mouse | 1:100 |
| **91** | Chk1 (phospho S296) | Chk1_pS296 | CHEK1 | Abcam | ab79758 | Rabbit | 1:125 |
| **92** | Chk1 (phospho S345) | Chk1_pS345 | CHEK1 | CST | 2348 | Rabbit | 1:30 |
| **93** | Chk2 | Chk2 | CHEK2 | CST | 3440 | Mouse | 1:50 |
| **94** | Chk2 (phospho T68) | Chk2_pT68 | CHEK2 | CST | 2197 | Rabbit | 1:250 |
| **95** | c-IAP2 | c-IAP2 | BIRC3 | CST | 3130 | Rabbit | 1:50 |
| **96** | CIITA | CIITA | CIITA | CST | 3793 | Rabbit | 1:250 |
| **97** | c-Jun (phospho S73) | c-Jun_pS73 | JUN | CST | 9164 | Rabbit | 1:30 |
| **98** | c-Kit | c-Kit | KIT | Epitomics/ Abcam | 1522-1/ ab32363 | Rabbit | 1:250 |
| **99** | Claudin 7 | Claudin-7 | CLDN7 | Abcam | ab79481 | Rabbit | 1:250 |
| **100** | c-Myc | c-Myc | MYC | Santa Cruz | sc-764 | Rabbit | 1:250 |
| **101** | COG3 | COG3 | COG3 | ProteinTech | 11130-1-AP | Rabbit | 1:750 |
| **102** | Collagen-VI/COL6A1 | Collagen-VI | COL6A1 | Santa Cruz | sc-20649 | Rabbit | 1:6000 |
| **103** | Complex II Subunit | Complex-II-Subunit | SDHB | Life Technologies | 459230 | Mouse | 1:200 |
| **104** | Connexin 43 | Connexin-43 | GJA1 | CST | 3512 | Rabbit | 1:150 |
| **105** | Coup-TFII | Coup-TFII | NR2F2 | CST | 6434 | Rabbit | 1:50 |
| **106** | Cox2 | Cox2 | PTGS2 | CST | 4842 | Rabbit | 1:75 |
| **107** | Cox-IV | Cox-IV | COX4I1 | CST | 4850 | Rabbit | 1:5000 |
| **108** | C-Raf | C-Raf | RAF1 | Millipore | 04-739 | Rabbit | 1:100 |
| **109** | C-Raf (phospho S338) | C-Raf_pS338 | RAF1 | CST | 9427 | Rabbit | 1:200 |

| | | | | | | | |
|---|---|---|---|---|---|---|---|
| **110** | Creb | Creb | CREB1 | CST | 9197 | Rabbit | 1:75 |
| **111** | CSK | CSK | CSK | CST | 4980 | Rabbit | 1:300 |
| **112** | CtIP | CtIP | RBBP8 | CST | 9201 | Rabbit | 1:500 |
| **113** | Cyclin B1 | Cyclin B1 | CCNB1 | Epitomics/ Abcam | 1495-1/ ab32053 | Rabbit | 1:1500 |
| **114** | Cyclin D1 | Cyclin-D1 | CCND1 | Millipore Sigma | SAB4502603 | Rabbit | 1:200 |
| **115** | Cyclin D3 | Cyclin D3 | CCND3 | CST | 2936 | Mouse | 1:1000 |
| **116** | Cyclin E1 | Cyclin E1 | CCNE1 | Santa Cruz | sc-247 | Mouse | 1:25 |
| **117** | Cyclophilin-F | Cyclophilin-F | PPIF | Abcam | MSA04/ ab110324 | Mouse | 1:50000 |
| **118** | Cytokeratin 19 | Cytokeratin-19 | KRT19 | Dako | M0888 | Mouse | 1:50 |
| **119** | DAP Kinase 1 (phospho S308) | DAPK1_pS308 | DAPK1 | GeneTex | GTX10524 | Mouse | 1:200 |
| **120** | DAP Kinase 2 | DAPK2 | DAPK2 | Abcam | ab51601 | Rabbit | 1:250 |
| **121** | DDB-1 | DDB-1 | DDB1 | CST | 6998 | Rabbit | 1:5000 |
| **122** | Detyrosinated alpha-Tubulin | D-a-Tubulin | TUBA4A, TUBA3C | Abcam | ab48389 | Rabbit | 1:1500 |
| **123** | Di-Methyl-Histone H3 (Lys4/C64G9) | DM-Histone-H3 | HIST1H3A | CST | 9725 | Rabbit | 1:100 |
| **124** | Dimethyl-K9 Histone H3 | DM-K9-Histone-H3 | HIST3H3 | Abcam | ab1220 | Mouse | 1:250 |
| **125** | DNA Ligase IV | DNA-Ligase-IV | LIG4 | CST | 14649 | Rabbit | 1:1000 |
| **126** | DNA Polymerase gamma (D1Y6R) | POLG | POLG | CST | 13609 | Rabbit | 1:500 |
| **127** | DNMT1 (D63A6) | DNMT1 | DNMT1 | CST | 5032 | Rabbit | 1:500 |
| **128** | DRP1 (D8H5) | DRP1 | DNM1L | CST | 5391 | Rabbit | 1:1000 |
| **129** | DUSP4/MKP2 | DUSP4 | DUSP4 | CST | 5149 | Rabbit | 1:150 |
| **130** | DUSP6 | DUSP6 | DUSP6 | Abcam | ab76310 | Rabbit | 1:750 |
| **131** | Dvl3 | Dvl3 | DVL3 | CST | 3218 | Rabbit | 1:30 |
| **132** | E2F1 | E2F1 | E2F1 | Santa Cruz | sc-251 | Mouse | 1:20 |
| **133** | E-Cadherin | E-Cadherin | CDH1 | CST | 3195 | Rabbit | 1:150 |
| **134** | eEF2 | eEF2 | EEF2 | CST | 2332 | Rabbit | 1:50 |
| **135** | eEF2K | eEF2K | EEF2K | CST | 3692 | Rabbit | 1:50 |
| **136** | EGFR | EGFR | EGFR | CST | 2232 | Rabbit | 1:75 |
| **137** | EGFR (phospho Y1173) | EGFR_pY1173 | EGFR | Epitomics/ Abcam | 1124-1/ ab32578 | Rabbit | 1:300 |
| **138** | eIF4E | eIF4E | EIF4E | CST | 9742 | Rabbit | 1:75 |
| **139** | eIF4E (phospho S209) | eIF4E_pS209 | EIF4E | Abcam | ab76256 | Rabbit | 1:250 |

| | | | | | | | |
|---|---|---|---|---|---|---|---|
| **140** | eIF4G | eIF4G | EIF4G1 | CST | 2498 | Rabbit | 1:1000 |
| **141** | Elk1 (phospho S383) | Elk1_pS383 | ELK1 | CST | 9181 | Rabbit | 1:50 |
| **142** | Enolase-2 (D20H2) | Enolase-2 | ENO2 | CST | 8171 | Rabbit | 1:250 |
| **143** | ENY2 | ENY2 | ENY2 | GeneTex | GTX629542 | Mouse | 1:500 |
| **144** | Eph Receptor A2 | EPHA2 | EPHA2 | Abcam | ab133501 | Rabbit | 1:1000 |
| **145** | Epithelial Membrane Antigen | EMA | MUC1 | DAKO | M061329-2 | Mouse | 1:750 |
| **146** | ErbB3/HER3 | HER3 | ERBB3 | Santa Cruz | sc-285 | Rabbit | 1:300 |
| **147** | ErbB3/HER3 (phospho Y1289) | HER3_pY1289 | ERBB3 | CST | 4791 | Rabbit | 1:50 |
| **148** | ERCC1 | ERCC1 | ERCC1 | Santa Cruz | sc-17809 | Mouse | 1:38 |
| **149** | Erk5 | Erk5 | MAPK7 | CST | 3552 | Rabbit | 1:500 |
| **150** | ERRalpha (E1G1J) | ERRalpha | ESRRA | CST | 13826 | Rabbit | 1:500 |
| **151** | ERRFI1/MIG6 | MIG6 | ERRFI1 | Sigma-Aldrich | WH0054206 M1 | Mouse | 1:50 |
| **152** | Estrogen Receptor | ER | ESR1 | Lab Vision | RM-9101 | Rabbit | 1:40 |
| **153** | Estrogen Receptor alpha | ER-a | ERSA | CST | 13258 | Rabbit | 1:500 |
| **154** | Estrogen Receptor alpha (phospho S118) | ER-a_pS118 | ESR1 | Epitomics/ Abcam | 1091-1/ ab32396 | Rabbit | 1:500 |
| **155** | Ets-1 | Ets-1 | ETS1 | Bethyl | A303-501A | Rabbit | 1:100 |
| **156** | FAK | FAK | PTK2 | Epitomics/ Abcam | 1700-1/ ab40794 | Rabbit | 1:1000 |
| **157** | FAK (phospho Y397) | FAK_pY397 | PTK2 | CST | 3283 | Rabbit | 1:25 |
| **158** | Fatty Acid Synthase | FASN | FASN | CST | 3180 | Rabbit | 1:1000 |
| **159** | FGF-basic | FGF-basic | FGF2 | VWR | 10775-082 (500-P18) | Rabbit | 1:1000 |
| **160** | Fibronectin | Fibronectin | FN1 | Epitomics | 1574-1 | Rabbit | 1:10000 |
| **161** | FoxM1 | FOXM1 | FOXM1 | CST | 5436 | Rabbit | 1:30 |
| **162** | FoxO3a | FoxO3a | FOXO3 | CST | 2497 | Rabbit | 1:20 |
| **163** | FoxO3a (phospho S318/S321) | FoxO3a_pS318_S3 21 | FOXO3 | CST | 9465 | Rabbit | 1:30 |
| **164** | FRS2-a (phospho Y196) | FRS2-a_pY196 | FRS2 | CST | 3864 | Rabbit | 1:100 |
| **165** | G6PD | G6PD | G6PD | Santa Cruz | sc-373887 | Mouse | 1:75 |

| | | | | | | | |
|---|---|---|---|---|---|---|---|
| **166** | Gab2 | Gab2 | GAB2 | CST | 3239 | Rabbit | 1:300 |
| **167** | GAPDH | GAPDH | GAPDH | Ambion/ Invitrogen | AM4300 | Mouse | 1:75000 |
| **168** | GATA3 | GATA3 | GATA3 | BD Biosciences | 558686 | Mouse | 1:150 |
| **169** | GATA6 | GATA6 | GATA6 | CST | 5851 | Rabbit | 1:200 |
| **170** | GCLC | GCLC | GCLC | Proteintech Group | 12601-1-AP | Rabbit | 1:500 |
| **171** | GCLM | GCLM | GCLM | Abcam | ab124827 | Rabbit | 1:500 |
| **172** | GCN5L2 | GCN5L2 | KAT2A | CST | 3305 | Rabbit | 1:30 |
| **173** | Gli1 | Gli1 | GLI1 | CST | 3538 | Rabbit | 1:3000 |
| **174** | Gli3 | Gli3 | GLI3 | Abcam | ab69838 | Rabbit | 1:1000 |
| **175** | Glucose-6 Phosphate Dehydrogenase | G6PD | G6PD | CST | 8866 | Rabbit | 1:30 |
| **176** | Glutamate Dehydrogenase1/ 2 | Glutamate-D1-2 | GLUD1 | Novus | NBP2-16679 | Rabbit | 1:500 |
| **177** | Glutaminase | Glutaminase | GLS | Abcam | ab156876 | Rabbit | 1:150 |
| **178** | Glycogen Synthase | Gys | GYS1 | CST | 3886 | Rabbit | 1:2000 |
| **179** | Glycogen Synthase (phospho S641) | Gys_pS641 | GYS1 | CST | 3891 | Rabbit | 1:300 |
| **180** | GPBB | GPBB | PYGM | Novus | NBP1-32799 | Rabbit | 1:200 |
| **181** | Granzyme B | Granzyme-B | GZMB | CST | 4275 | Rabbit | 1:500 |
| **182** | GRB7 | GRB7 | GRB7 | Abcam | ab183737 | Rabbit | 1:500 |
| **183** | Grp75 (D13H4) | Grp75 | HSPA9 | CST | 3593 | Rabbit | 1:250 |
| **184** | GSK-3alpha/beta | GSK-3a-b | GSK3A, B | Santa Cruz | sc-7291 | Mouse | 1:750 |
| **185** | GSK-3alpha/beta (phospho S21/S9) | GSK-3a-b_pS21_S9 | GSK3A, B | CST | 9331 | Rabbit | 1:200 |
| **186** | GSK-3B | GSK-3B | GSK3B | CST | 9315 | Rabbit | 1:750 |
| **187** | GSK-3beta (phospho S9) | GSK-3b_pS9 | GSK3B | CST | 5558 | Rabbit | 1:250 |
| **188** | H2AX (phospho S140) | H2AX_pS140 | H2AFX | Pierce Biotechnolog y | MA12022 | Mouse | 1:100 |
| **189** | Hamartin/TSC1 | TSC1 | TSC1 | CST | 4906 | Rabbit | 1:200 |
| **190** | HER2 | HER2 | ERBB2 | Lab Vision | MS-325-P1 | Mouse | 1:300 |
| **191** | HER2 (phospho Y1248) | HER2_pY1248 | ERBB2 | R&D systems | AF1768 | Rabbit | 1:1500 |
| **192** | Heregulin | Heregulin | NRG1 | CST | 2573 | Rabbit | 01:30 |

| | | | | | | | |
|---|---|---|---|---|---|---|---|
| **193** | HES1 | HES1 | HES1 | CST | 11988 | Rabbit | 1:500 |
| **194** | Hexokinase II | Hexokinase II | HK2 | CST | 2106 | Rabbit | 1:100 |
| **195** | Hif-1-alpha | Hif-1-alpha | HIF1A | BD Biosciences | 610958 | Mouse | 1:20 |
| **196** | Histone H3 | Histone H3 | HIST3H3 | Abcam | ab1791 | Rabbit | 1:5000 |
| **197** | HLA-DQA1 | HLA-DQA1 | HLA-DQA1 | Abcam | ab128959 | Rabbit | 1:3000 |
| **198** | HLA-DR/DP/DQ/DX | HLA-DR-DP-DQ-DX | HLA-DRA | Santa Cruz | sc-53302 | Mouse | 1:250 |
| **199** | HMHA1 | HMHA1 | HMHA1 | ProteinTech | 14832-1-AP | Rabbit | 1:3000 |
| **200** | HSP27 | HSP27 | HSBP1 | CST | 2402 | Mouse | 1:75 |
| **201** | HSP27 (phospho S82) | HSP27_pS82 | HSBP1 | CST | 2401 | Rabbit | 1:75 |
| **202** | HSP60 | HSP60 | HSP60 | CST | 12165 | Rabbit | 1:1000 |
| **203** | HSP70 | HSP70 | HSPA1A | CST | 4872 | Rabbit | 1:50 |
| **204** | Hsp75/TRAP1 | TRAP1 | TRAP1 | BD Biosciences | 612344 | Mouse | 1:750 |
| **205** | IDO | IDO | IDO1 | CST | 86630 | Rabbit | 1:200 |
| **206** | IGF1R (phospho Y1135/Y1136 | IGF1R_pY1135_Y1136 | IGF1R, INSR | CST | 3024 | Rabbit | 1:30 |
| **207** | IGF-1Receptor beta | IGF1R-b | IGF1R | CST | 3018 | Rabbit | 1:50 |
| **208** | IGFBP2 | IGFBP2 | IGFBP2 | CST | 3922 | Rabbit | 1:50 |
| **209** | IGFBP3 | IGFBP3 | IGFBP3 | BD Biosciences | 611504 | Mouse | 1:1000 |
| **210** | IGFRb | IGFRb | IGF1R | CST | 3027 | Rabbit | 1:250 |
| **211** | IL-6 | IL-6 | IL6 | CST | 12153 | Rabbit | 1:250 |
| **212** | INPP4b | INPP4b | INPP4B | CST | 4039 | Rabbit | 01:30 |
| **213** | Insulin Receptor beta | IR-b | INSR | CST | 3025 | Rabbit | 1:100 |
| **214** | IRF-1 | IRF-1 | IRF1 | CST | 8478 | Rabbit | 1:250 |
| **215** | IRS1 | IRS1 | IRS1 | Millipore | 06-248 | Rabbit | 1:250 |
| **216** | IRS2 | IRS2 | IRS2 | CST | 4502 | Rabbit | 1:100 |
| **217** | JAB1 | JAB1 | COPS5 | Santa Cruz | sc-13157 | Mouse | 1:30 |
| **218** | Jagged1 | Jagged1 | JAG1 | Abcam | ab109536 | Rabbit | 01:50 |
| **219** | Jak2 | Jak2 | JAK2 | CST | 3230 | Rabbit | 1:750 |
| **220** | JNK (phospho T183/Y185) | JNK_pT183_Y185 | MAPK8 | CST | 4668 | Rabbit | 01:30 |
| **221** | JNK2 | JNK2 | MAPK9 | CST | 4672 | Rabbit | 1:25 |
| **222** | KAP1 | KAP1 | TRIM28 | Abcam | ab10484 | Rabbit | 1:2000 |
| **223** | KMT3A/HYPB/HIF-1 | SETD2 | SETD2 | abcam | ab184190 | Rabbit | 1:1000 |

| | | | | | | | |
|---|---|---|---|---|---|---|---|
| **224** | LAD1 | LAD1 | LAD1 | Atlas | HPA028732 | Rabbit | 1:500 |
| **225** | Lasu1/Ureb1 | Lasu1 | HUWE1 | Bethyl | IHC-00439 | Rabbit | 1:1000 |
| **226** | LC3A/B | LC3A-B | MAP1LC3A, B | CST | 4108 | Rabbit | 1:250 |
| **227** | Lck | Lck | LCK | CST | 2752 | Rabbit | 1:75 |
| **228** | LDHA | LDHA | LDHA | CST | 3582 | Rabbit | 1:250 |
| **229** | LRP6 (phospho S1490) | LRP6_pS1490 | LRP6 | CST | 2568 | Rabbit | 1:250 |
| **230** | MAPK (phospho T202/Y204) | MAPK_pT202/Y204 | MAPK1, 3 | CST | 4377 | Rabbit | 1:25 |
| **231** | Mcl-1 | Mcl-1 | MCL1 | CST | 5453 | Rabbit | 1:100 |
| **232** | MDM2 (phospho S166) | MDM2_pS166 | MDM2 | CST | 3521 | Rabbit | 1:60 |
| **233** | MEK1 | MEK1 | MAP2K1 | Epitomics/ Abcam | 1235-1/ ab32576 | Rabbit | 1:1500 |
| **234** | MEK1 (phospho S217/S221) | MEK1_p_S217/ S221 | MAP2K1, 2 | CST | 9154 | Rabbit | 1:50 |
| **235** | MEK2 | MEK2 | MAP2K2 | CST | 9125 | Rabbit | 1:50 |
| **236** | MelanA | MelanA | MLANA | Abcam | ab51061 | Rabbit | 1:500 |
| **237** | Melanoma gp100 | Melan-gp100 | PMEL | Abcam | ab137078 | Rabbit | 1:500 |
| **238** | MERIT40 | MERIT40 | MERIT40 | CST | 12711 | Rabbit | 1:3000 |
| **239** | MERIT40 (phospho S29) | MERIT40_pS29 | BABAM1 | CST | 12110 | Rabbit | 1:300 |
| **240** | Merlin/NF2 | Merlin | NF2 | Novus | 22710002 | Rabbit | 1:250 |
| **241** | MIF | MIF | MIF | Santa Cruz | sc-130329 | Rabbit | 1:100 |
| **242** | MITF (D5G7V) | MITF | MITF | CST | 12590 | Rabbit | 1:500 |
| **243** | Mitofusin-1 | Mitofusin-1 | MFN1 | CST | 14739 | Rabbit | 1:500 |
| **244** | Mitofusin-2 | Mitofusin-2 | MFN2 | CST | 11925 | Rabbit | 1:1000 |
| **245** | MLH1 (4C9C7) | MLH1 | MLH1 | CST | 3515 | Mouse | 1:500 |
| **246** | MLKL | MLKL | MLKL | CST | 14993 | Rabbit | 1:1000 |
| **247** | MMP2 | MMP2 | MMP2 | CST | 4022 | Rabbit | 1:75 |
| **248** | Mnk1 | Mnk1 | MKNK1 | CST | 2195 | Rabbit | 1:750 |
| **249** | Monocarboxylic Acid Transporter 4 | MCT4 | SLC16A4 | Millipore | AB3314P | Rabbit | 1:500 |
| **250** | MR1 | MR1 | MR1 | Santa Cruz | sc-377312 | Mouse | 1:500 |
| **251** | MRAP | MRAP | MRAP | Abcam | ab103319 | Rabbit | 1:500 |
| **252** | MSH2 (D24B5) | MSH2 | MSH2 | CST | 2017 | Rabbit | 1:750 |
| **253** | MSH6 | MSH6 | MSH6 | Novus | 22030002 | Rabbit | 1:1000 |
| **254** | MSI2 (EP1305Y) | MSI2 | MSI2 | Abcam | ab76148 | Rabbit | 1:1000 |

| | | | | | | | |
|---|---|---|---|---|---|---|---|
| **255** | MTCO1 | MTCO1 | MTCO1 | Abcam | ab14705 | Mouse | 1:500 |
| **256** | mTOR | mTOR | MTOR | CST | 2983 | Rabbit | 1:3000 |
| **257** | mTOR (phospho S2448) | mTOR_pS2448 | MTOR | CST | 2971 | Rabbit | 1:50 |
| **258** | MTSS1 | MTSS1 | MTSS1 | Novus | H00009788-M01A | Mouse | 1:250 |
| **259** | Myosin Heavy Chain 11 | Myosin-11 | MYH11 | Novus | 21370002 | Rabbit | 1:1000 |
| **260** | Myosin IIa | Myosin-IIa | MYH9 | CST | 3403 | Rabbit | 1:1000 |
| **261** | Myosin IIa (phospho S1943) | Myosin-IIa_pS1943 | MYH9 | CST | 5026 | Rabbit | 1:750 |
| **262** | Myt1 | Myt1 | PKMYT1 | CST | 4282 | Rabbit | 1:100 |
| **263** | NAPSIN-A | NAPSIN-A | NAPSA | Epitomics/ Abcam | 5795-1/ ab129189 | Rabbit | 1:150 |
| **264** | N-Cadherin | N-Cadherin | CDH2 | CST | 4061 | Rabbit | 1:25 |
| **265** | NDRG1 (phospho T346) | NDRG1_pT346 | NDRG1 | CST | 3217 | Rabbit | 01:50 |
| **266** | NDUFB4 | NDUFB4 | NDUFB4 | Abcam | ab110243 | Mouse | 1:25 |
| **267** | NF-kB p65 (phospho S536) | NF-kB-p65_pS536 | RELA | CST | 3033 | Rabbit | 1:30 |
| **268** | Notch1 | Notch1 | NOTCH1 | CST | 3268 | Rabbit | 01:30 |
| **299** | Notch1 (Cleaved) | Notch1-cleaved | NOTCH1 | CST | 4147 | Rabbit | 1:100 |
| **270** | Notch3 | Notch3 | NOTCH3 | Novus | H00004854-M01 | Mouse | 1:250 |
| **271** | NQO1 | NQO1 | NQO1 | CST | 3187 | Mouse | 1:15000 |
| **272** | N-Ras | N-Ras | NRAS | Santa Cruz | sc-31 | Mouse | 1:50 |
| **273** | NRF2 | NRF2 | NRF2 | CST | 12721 | Rabbit | 1:500 |
| **274** | Oct-4 | Oct-4 | POU5F1 | CST | 2750 | Rabbit | 1:40 |
| **275** | p16/INK4a | p16-INK4a | CDKN2A | Epitomics/ Abcam | 1712-1/ ab40803 | Rabbit | 1:500 |
| **276** | p21 | p21 | CDKN1A | Santa Cruz | sc-6246 | Rabbit | 1:150 |
| **277** | p27 (phospho T157) | p27_pT157 | CDKN1B | R&D Systems | AF1555 | Rabbit | 1:30 |
| **278** | p27 (phospho T198) | p27_pT198 | CDKN1B | Abcam | ab64949 | Rabbit | 01:50 |
| **279** | p27 KIP 1 | p27-Kip-1 | CDKN1B | Epitomics/ Abcam | 1591-1/ ab32034 | Rabbit | 01:40 |
| **280** | p38 (phospho T180/Y182) | p38_pT180_Y182 | MAPK11, 12, 13, 14 | CST | 9211 | Rabbit | 1:38 |
| **281** | p38 alpha MAPK | p38-a | MAPK1 | CST | 9228 | Mouse | 1:300 |
| **282** | p38 MAPK | p38-MAPK | MAPK11, 12, 14 | CST | 9212 | Rabbit | 1:1500 |

| | | | | | | | |
|---|---|---|---|---|---|---|---|
| **283** | p38/MAPK (phospho T180/Y182) | p38-MAPK_pT180_Y182 | MAPK14 | CST | 9215 | Rabbit | 1:250 |
| **284** | p44/42 MAPK | p44-42-MAPK | MAPK1, 3 | CST | 4695 | Rabbit | 1:2000 |
| **285** | p53 | p53 | TP53 | CST | 9282 | Rabbit | 1:2500 |
| **286** | p70 S6 Kinase (phospho T389) | p70-S6K_pT389 | RPS6KB1 | CST | 9205 | Rabbit | 1:50 |
| **287** | p70/S6K1 | p70-S6K1 | RPS6KB1 | Epitomics/ Abcam | 1494-1/ ab32529 | Rabbit | 1:300 |
| **288** | p90RSK (phospho T573) | p90RSK_pT573 | RPS6K | CST | 9346 | Rabbit | 1:25 |
| **289** | PAI-1 | PAI-1 | SERPINE1 | BD Biosciences | 612024 | Mouse | 1:50 |
| **290** | PAICS | PAICS | PAICS | Sigma-Aldrich | HPA035895 | Rabbit | 1:250 |
| **291** | PAK1 | PAK1 | PAK1 | CST | 2602 | Rabbit | 1:750 |
| **292** | PAK4 | PAK4 | PAK4 | CST | 3242 | Rabbit | 1:300 |
| **293** | PAR | PAR | PAR | Trevigen | 4336-BPC-100 | Rabbit | 1:30000 |
| **294** | PARG | PARG | PARG | CST | 66564 | Rabbit | 1:1000 |
| **295** | PARK7/DJ1 | DJ1 | PARK7 | Abcam | ab76008 | Rabbit | 1:5000 |
| **296** | PARP | PARP | PARP1 | CST | 9532 | Rabbit | 1:1000 |
| **297** | Patched | Patched | PTCH1 | Abcam | ab53715 | Rabbit | 1:1000 |
| **298** | Paxillin | Paxillin | PXN | CST | 2542 | Rabbit | 1:250 |
| **299** | P-Cadherin | P-Cadherin | CDH3 | CST | 2130 | Rabbit | 1:38 |
| **300** | PCNA | PCNA | PCNA | CST | 2586 | Mouse | 1:250 |
| **301** | PD-1 | PD-1 | PDCD1 | CST | 43248 | Mouse | 1:500 |
| **302** | Pdcd4 | Pdcd4 | PDCD4 | Rockland | 600-401-965 | Rabbit | 1:750 |
| **303** | PDGFRB | PDGFR-b | PDGFRB | Invitrogen | MA5-15143 | Rabbit | 1:500 |
| **304** | PDH | PDH | PDH | Abcam | ab110332 | Mouse | 1:100 |
| **305** | PDHK1 | PDHK1 | PDHK1 | CST | 3820 | Rabbit | 1:300 |
| **306** | PDK1 | PDK1 | PDPK1 | CST | 3062 | Rabbit | 01:50 |
| **307** | PDK1 (phospho S241) | PDK1_pS241 | PDPK1 | CST | 3061 | Rabbit | 01:50 |
| **308** | PD-L1 | PD-L1 | CD274 | CST | 13684 | Rabbit | 1:250 |
| **309** | PEA-15 | PEA-15 | PEA15 | CST | 2780S | Rabbit | 1:100 |
| **310** | PED/PEA-15 (phospho S116) | PEA-15_pS116 | PEA15 | Life Technologies | 44836G | Rabbit | 1:100 |
| **311** | PHGDH | PHGDH | PHGDH | CST | 13428 | Rabbit | 1:1000 |
| **312** | PI3 Kinase p110 alpha | PI3K-p110-a | PIK3CA | CST | 4255 | Rabbit | 1:50 |
| **313** | PI3K p110 beta | PI3K-p110-b | PIK3BC | Santa Cruz | sc-376412 | Mouse | 1:40 |

| | | | | | | | |
|---|---|---|---|---|---|---|---|
| **314** | PI3K p85 | PI3K-p85 | PIK3R1 | Millipore | 06-195 | Rabbit | 1:15000 |
| **315** | PKA RI alpha | PKA-a | PRKAR1A | CST | 5675 | Rabbit | 1:250 |
| **316** | PKC alpha/beta II (phospho T638/641) | PKC-a-b-II_pT638_T641 | PRKCA, B | CST | 9375 | Rabbit | 1:1000 |
| **317** | PKC(pan) beta II (phospho S660) | PKC-b-II_pS660 | PRKCA, B, D, E, H, Q | CST | 9371 | Rabbit | 1:200 |
| **318** | PKC delta (phospho S664) | PKC-delta_pS664 | PRKCD | Millipore | 07-875 | Rabbit | 1:75 |
| **319** | PKCalpha | PKCa | PRKCA | CST | 2056 | Rabbit | 1:200 |
| **320** | PKM2 | PKM2 | PKM | CST | 4053 | Rabbit | 1:300 |
| **321** | PLC gamma2 (phospho Y759) | PLC-gamma2_pY759 | PLCG2 | CST | 3874 | Rabbit | 01:25 |
| **322** | PLK1 | PLK1 | PLK1 | CST | 4513 | Rabbit | 1:125 |
| **323** | Met (phospho Y1234/Y1235) | c-Met_pY1234_Y1235 | MET | CST | 3129 | Rabbit | 1:100 |
| **324** | PMS2 | PMS2 | PMS2 | Novus Biologicals | 22510002 | Rabbit | 1:1500 |
| **325** | PRAS40 | PRAS40 | AKT1S1 | Life Technologies | AHO1031 | Mouse | 1:75 |
| **326** | PRAS40 (phospho T246) | PRAS40_pT246 | AKT1S1 | Life Technologies | 441100G | Rabbit | 1:500 |
| **327** | PREX1 | PREX1 | PREX1 | Abcam | ab102739 | Rabbit | 1:100 |
| **328** | Progesterone Receptor [YR85] | PR | PGR | abcam | 206926 | Rabbit | 1:500 |
| **329** | PTEN | PTEN | PTEN | CST | 9552 | Rabbit | 1:500 |
| **330** | PTPN12 | PTPN12 | PTPN12 | Abcam | ab76942 | Rabbit | 1:500 |
| **331** | Puma | Puma | BBC3 | CST | 4976 | Rabbit | 1:50 |
| **332** | PYGB | PYGB | PYGB | Sigma-Aldrich | SAB2900066 | Rabbit | 1:750 |
| **333** | PYGM | PYGM | PYGM | Novus | H00005837-M10 | Mouse | 1:500 |
| **334** | Pyk2 (phospho Y402) | Pyk2_pY402 | PYK2 | CST | 3291 | Rabbit | 1:500 |
| **335** | Pyruvate Dehydrogenase | PDHA1 | PDHA1 | CST | 3205 | Rabbit | 1:200 |
| **336** | Rab11 | Rab11 | RAB11A, B | CST | 3539 | Rabbit | 1:30 |
| **337** | Rab25 | Rab25 | RAB25 | CST | 4314 | Rabbit | 1:30 |
| **338** | Rac1/Cdc42 | Cdc42 | CDC42 | CST | 4651 | Rabbit | 1:100 |
| **339** | Rad23A | Rad23A | RAD23A | CST | 24555 | Rabbit | 1:1000 |
| **340** | Rad50 | Rad50 | RAD50 | CST | 3427 | Rabbit | 1:250 |
| **341** | Rad51 | Rad51 | RAD51 | Millipore | ABE257 | Rabbit | 1:1000 |

| | | | | | | | |
|---|---|---|---|---|---|---|---|
| **342** | Raptor | Raptor | RPTOR | CST | 2280 | Rabbit | 1:300 |
| **343** | Rb | Rb | RB1 | CST | 9309 | Mouse | 1:150 |
| **344** | Rb (phospho S807/811) | Rb_pS807_S811 | RB1 | CST | 9308 | Rabbit | 1:1000 |
| **345** | RBM15 | RBM15 | RBM15 | Novus | 21390002 | Rabbit | 1:5000 |
| **346** | Rheb | Rheb | RHEB | R&D Systems | MAB3426 | Mouse | 1:75 |
| **347** | Rictor | Rictor | RICTOR | CST | 2114 | Rabbit | 1:100 |
| **348** | Rictor (phospho T1135) | Rictor_pT1135 | RICTOR | CST | 3806 | Rabbit | 1:200 |
| **349** | RIP | RIP | RIP | CST | 4926 | Rabbit | 1:75 |
| **350** | RIP3 | RIP3 | RIP3 | CST | 13526 | Rabbit | 1:500 |
| **351** | RPA32 (phospho S4/S8) | RPA32_pS4/S8 | RPA2 | Bethyl | A300-245A | Rabbit | 1:250 |
| **352** | RPA32/RPA2 | RPA32 | RPA2 | CST | 2208 | Rat | 1:150 |
| **353** | RRM1 | RRM1 | RRM1 | CST | 3388 | Rabbit | 1:100 |
| **354** | RRM2 | RRM2 | RRM2 | Life Technologies | PA527856 | Rabbit | 1:250 |
| **355** | RSK | RSK | RPS6KA1, 2, 3 | CST | 9347 | Rabbit | 1:150 |
| **356** | S100A4 | S100A4 | S100A4 | CST | 13018 | Rabbit | 1:1000 |
| **357** | S6 (phospho S235/236) | S6_pS235_S236 | RPS6 | CST | 2211 | Rabbit | 1:2500 |
| **358** | S6 (phospho S240/244) | S6_pS240_S244 | RPS6 | CST | 2215 | Rabbit | 1:1000 |
| **359** | S6 Ribosomal Protein | S6 | RPS6 | CST | 2317 | Mouse | 1:750 |
| **360** | SCD | SCD | SCD | Santa Cruz | sc-58420 | Mouse | 1:20 |
| **361** | SDHA | SDHA | SDHA | CST | 11998 | Rabbit | 1:250 |
| **362** | SFRP1 | SFRP1 | SFRP1 | CST | 4690 | Rabbit | 1:500 |
| **363** | Shc_pY317 | Shc_pY317 | SHC1 | CST | 2431 | Rabbit | 01:25 |
| **364** | SHP-2 (phospho Y542) | SHP-2_pY542 | PTPN11 | CST | 3751 | Rabbit | 1:75 |
| **365** | SHP2 / PTPN11 | SHP2 | PTPN11 | CST | 3397 | Rabbit | 1:250 |
| **366** | SLC1A5 | SLC1A5 | SLC1A5 | Sigma-Aldrich | HPA035240 | Rabbit | 1:150000 |
| **367** | Slfn11 | Slfn11 | SLFN11 | Santa Cruz | sc-374339 | Mouse | 1:150 |
| **368** | Smac | Smac | DIABLO | CST | 2954 | Mouse | 1:150 |
| **369** | Smad1 | Smad1 | SMAD1 | Epitomics/ Abcam | 1649-1/ ab33902 | Rabbit | 1:500 |
| **370** | Smad3 | Smad3 | SMAD3 | Epitomics/ Abcam | 1735-1/ ab40854 | Rabbit | 1:150 |
| **371** | Smad4 | Smad4 | SMAD4 | Santa Cruz | sc-7966 | Mouse | 1:30 |

| | | | | | | | |
|---|---|---|---|---|---|---|---|
| **372** | Snail | Snail | SNAI1 | CST | 3895 | Mouse | 1:50 |
| **373** | SOD1 | SOD1 | SOD1 | CST | 4266 | Mouse | 1:500 |
| **374** | SOD2 (D9V9C) | SOD2 | SOD2 | CST | 13194 | Rabbit | 1:200 |
| **375** | Sox2 | Sox2 | SOX2 | CST | 2748 | Rabbit | 1:50 |
| **376** | Src | Src | SRC | Millipore | 05-184 | Mouse | 1:50 |
| **377** | Src (phospho Y416) | Src_pY419 | SRC | CST | 2101 | Rabbit | 1:25 |
| **378** | Src (phospho Y527) | Src_pY527 | SRC | CST | 2105 | Rabbit | 1:150 |
| **379** | SRSF1/SF2 | SF2 | SRSF1 | Invitrogen | 324500 | Mouse | 1:75 |
| **380** | Stat3 | Stat3 | STAT3 | CST | 4904 | Rabbit | 1:3000 |
| **381** | Stat3 (phospho Y705) | Stat3_pY705 | STAT3 | CST | 9145 | Rabbit | 1:100 |
| **382** | Stat5a | Stat5a | STAT5A | Epitomics/ Abcam | 1289-1/ ab32043 | Rabbit | 1:300 |
| **383** | Stathmin-1 | Stathmin-1 | STMN1 | Epitomics/ Abcam | 1972-1/ ab52630 | Rabbit | 1:75 |
| **384** | STING | STING | TMEM173 | CST | 13647 | Rabbit | 1:250 |
| **385** | Syk | Syk | SYK | Santa Cruz | sc-1240 | Mouse | 1:500 |
| **386** | Tau | Tau | MAPT | Millipore | 05-348 | Mouse | 1:100 |
| **387** | TAZ | TAZ | WWTR1 | CST | 4883 | Rabbit | 1:300 |
| **388** | TFAM | TFAM | TFAM | CST | 7495 | Rabbit | 1:300 |
| **389** | Transferrin R | TFRC | TFRC | Novus | 22500002 | Rabbit | 1:15000 |
| **390** | TIGAR | TIGAR | TIGAR | Epitomics/ Abcam | S1711/ ab137573 | Rabbit | 1:100 |
| **391** | Transglutaminase | Transglutaminase | TGM2 | Lab Vision | MS-224-P1 | Mouse | 1:150 |
| **392** | TRIM25 | TRIM25 | TRIM25 | Abcam | ab167154 | Rabbit | 1:3000 |
| **393** | TTF1 | TTF1 | NKX2-1 | Epitomics/ Abcam | 2044-1/ ab76013 | Rabbit | 1:150 |
| **394** | Tuberin | Tuberin | TSC2 | Epitomics/ Abcam | 1613-1/ ab32554 | Rabbit | 1:2500 |
| **395** | Tuberin/TSC2 (phospho T1462) | Tuberin_pT1462 | TSC2 | CST | 3617 | Rabbit | 1:38 |
| **396** | TUFM | TUFM | TUFM | Abcam | ab173300 | Rabbit | 1:38 |
| **397** | TWEAK Receptor/FN14 | FN14 | TNFRSF12A | CST | 4403 | Rabbit | 1:1000 |
| **398** | TWIST | TWIST | TWIST1 | Santa Cruz | sc-81417 | Mouse | 1:30 |
| **399** | Tyro3 | Tyro3 | TYRO3 | CST | 5585 | Rabbit | 1:30 |
| **400** | UBAC1 | UBAC1 | UBAC1 | Sigma-Aldrich | HPA005651 | Rabbit | 1:250 |
| **401** | Ubiquityl-Histone H2B | U-Histone-H2B | HIST1H2BB | CST | 5546 | Rabbit | 1:500 |

| | | | | | | | |
|---|---|---|---|---|---|---|---|
| **402** | UGT1A | UGT1A | UGT1A1, 3, 4, 5, 7, 8, 10 | Santa Cruz | sc-271268 | Mouse | 1:75 |
| **403** | ULK1 (phospho S757) | ULK1_pS757 | ULK1 | CST | 6888 | Rabbit | 1:300 |
| **404** | UQCRC2 | UQCRC2 | UQCRC2 | MitoSciences / Abcam | MS304/ ab14745 | Mouse | 01:50 |
| **405** | UVRAG | UVRAG | UVRAG | CST | 13115 | Rabbit | 1:100 |
| **406** | VASP | VASP | VASP | CST | 3112 | Rabbit | 1:100 |
| **407** | Vav1 | Vav1 | VAV1 | CST | 2502 | Rabbit | 1:500 |
| **408** | VDAC1/Porin | Porin | VDAC1 | Abcam | ab14734 | Mouse | 1:100 |
| **409** | VEGF Receptor 2 | VEGFR-2 | KDR | CST | 2479 | Rabbit | 1:3000 |
| **410** | VHL/EPPK1 | VHL-EPPK1 | EPPK1 | BD Biosciences | 556347 | Mouse | 1:1500 |
| **411** | Vimentin | Vimentin | VIM | Dako/Fisher | M0725/ MS-129-P | Mouse | 1:250 |
| **412** | Vinculin | Vinculin | VCL | Sigma-Aldrich | SAB4200080 | Mouse | 1:25000 |
| **413** | Wee1 | Wee1 | WEE1 | CST | 4936 | Rabbit | 1:250 |
| **414** | Wee1 (phospho S642) | Wee1_pS642 | WEE1 | CST | 4910 | Rabbit | 1:50 |
| **415** | WIPI1 | WIPI1 | WIPI1 | CST | 12124 | Rabbit | 1:150 |
| **416** | WIPI2 | WIPI2 | WIPI2 | CST | 8567 | Rabbit | 1:150 |
| **417** | XBP-1 | XBP-1 | XBP1 | Santa Cruz | sc-32136 | Goat | 1:200 |
| **418** | XIAP | XIAP | XIAP | CST | 2042 | Rabbit | 1:100 |
| **419** | XPA | XPA | XPA | Santa Cruz | sc-56813 | Mouse | 1:75 |
| **420** | XPF | XPF | ERCC4 | Abcam | ab73720 | Rabbit | 1:100 |
| **421** | XPG | ERCC5 | ERCC5 | Proteintech Group | 11331-1-AP | Rabbit | 1:250 |
| **422** | XRCC1 | XRCC1 | XRCC1 | CST | 2735 | Rabbit | 1:20 |
| **423** | YAP | YAP | YAP1 | Santa Cruz | sc-376830 | Mouse | 1:300 |
| **424** | YAP (phospho S127) | YAP_pS127 | YAP1 | CST | 4911 | Rabbit | 1:250 |
| **425** | YB1 (phospho S102) | YB1_pS102 | YBX1 | CST | 2900 | Rabbit | 1:50 |
| **426** | ZAP-70 | ZAP-70 | ZAP70 | CST | 2705 | Rabbit | 1:500 |

**Table S3.** Molecular/signaling characteristics of *MMTV-R26* $^{Met}$ tumors identified through RPPA analysis.

| Sample | 14-3-3-beta | 14-3-3-zeta | 4E-BP1 | 4E-BP1_pS65 | 53BP1 | A-Raf | ACC1 | ACC_pS79 | Akt | Akt_pS473 |
|---|---|---|---|---|---|---|---|---|---|---|
| T1 | 1,00030313 | 1,112909745 | 1,134492549 | 1,137233694 | 1,447094044 | 0,994553464 | 1,181854819 | 1,197543263 | 1,102463951 | 0,720638799 |
| T2 | 0,941037749 | 1,096944273 | 1,027642301 | 1,216365052 | 1,226844507 | 1,198216315 | 0,911067172 | 0,918659697 | 0,90597967 | 0,783979594 |
| T3 | 1,245118013 | 0,757216041 | 0,882534965 | 0,93677973 | 0,384268168 | 0,580713129 | 0,302714776 | 0,307450113 | 0,99411866 | 0,924505019 |
| T4 | 0,843563418 | 1,047775802 | 0,902704787 | 1,108629724 | 0,855440051 | 1,073170685 | 0,626047448 | 0,814158536 | 1,106024715 | 0,746302303 |
| T5 | 0,890252276 | 1,226663216 | 1,184383298 | 1,260307415 | 0,93378607 | 1,088196632 | 1,208043539 | 1,169484449 | 0,84737051 | 0,895688597 |
| T6 | 0,93645119 | 1,005972333 | 1,194694942 | 1,054710983 | 1,189703081 | 1,223393087 | 0,69113579 | 0,14513807 | 0,897245656 | 0,325770129 |
| T7 | 0,985372751 | 1,05955999 | 1,263612676 | 1,234111085 | 1,404953067 | 1,016783258 | 1,050783876 | 1,025081272 | 0,939280133 | 0,486231922 |
| T8 | 0,863746677 | 1,239080121 | 0,853463853 | 1,176759829 | 1,510982604 | 1,146751384 | 0,825043095 | 1,13633138 | 1,11054025 | 0,913958817 |
| T9 | 0,925172458 | 1,248622659 | 1,369888819 | 1,272146599 | 1,440378765 | 0,969565693 | 0,989826514 | 1,009446208 | 1,059704219 | 0,732273159 |
| T10 | 0,950840569 | 1,137862054 | 1,387515563 | 1,129187785 | 1,090606388 | 1,066259701 | 0,753114714 | 0,78318462 | 0,844195763 | 0,853441455 |
| T11 | 0,817212578 | 1,014257531 | 1,014197744 | 0,861902884 | 0,947313349 | 1,052522223 | 1,093881911 | 1,082935858 | 1,120470229 | 0,974542868 |
| T12 | 1,03998673 | 1,094035434 | 1,275213813 | 1,170383334 | 1,317040387 | 0,979142264 | 0,96358135 | 1,151345074 | 1,139495304 | 2,01446105 |
| T13 | 1,017625373 | 1,048012612 | 0,750049112 | 1,061406134 | 0,945045464 | 0,891689943 | 0,395264773 | 0,522335999 | 0,83651811 | 6,805906933 |
| T14 | 1,211802243 | 1,122590701 | 1,04265535 | 0,909412674 | 0,686463438 | 1,134026642 | 0,608433899 | 0,813731225 | 1,101593417 | 1,302656486 |
| T15 | 1,077863185 | 1,139889093 | 0,926640629 | 1,1019766 | 0,960782985 | 0,939882009 | 0,650313996 | 0,747168934 | 1,316225115 | 1,065531224 |
| T16 | 1,007403104 | 1,162129699 | 1,045959689 | 1,097375679 | 0,966304379 | 1,015226523 | 0,679954377 | 0,803804062 | 1,106787398 | 0,915525325 |
| T17 | 0,965560836 | 1,265364374 | 0,902678014 | 1,217621153 | 0,773950562 | 1,137843915 | 0,636906952 | 0,772715336 | 1,043402038 | 0,900258261 |
| T18 | 0,897611045 | 0,991430929 | 1,23006533 | 1,597264099 | 1,548351183 | 1,152894977 | 0,725471131 | 0,926021669 | 1,110552168 | 0,950555405 |
| T19 | 0,933238047 | 1,039550859 | 1,736692426 | 1,404388906 | 1,017718425 | 1,068116608 | 1 | 1,082159856 | 1,355815816 | 0,571355996 |
| T20 | 0,928451687 | 1,052360811 | 1,072346262 | 0,998826644 | 0,793427054 | 0,913167381 | 0,433369405 | 0,675786219 | 0,874258653 | 1,43691674 |
| T21 | 0,918862457 | 0,978238756 | 1,333393795 | 1,134903538 | 0,873341792 | 0,917925863 | 1,276197022 | 1,366606083 | 0,950787718 | 0,889009113 |
| T22 | 0,933876569 | 0,545456675 | 0,417475528 | 0,754254632 | 0,419997194 | 0,557967669 | 0,213328439 | 0,315396431 | 0,378923613 | 3,863099235 |
| T23 | 0,800170574 | 0,9657963 | 0,876890933 | 0,932487044 | 0,944547246 | 0,991183413 | 1,009073519 | 1,32439926 | 1,531700485 | 2,726033381 |
| T24 | 0,998593076 | 1,235547228 | 1,27248621 | 1,381722847 | 1,423449099 | 0,89640053 | 1,108624092 | 1,380400047 | 0,971670192 | 1,663675989 |
| N1 | 0,822850439 | 0,619543551 | 0,698145721 | 0,710435346 | 0,381816917 | 0,70722531 | 3,360065853 | 2,108678779 | 0,488063966 | 0,991773211 |
| N2 | 0,961319758 | 0,63981164 | 0,982891136 | 0,919567691 | 0,348989132 | 0,804085605 | 3,00322954 | 1,902650763 | 0,681699351 | 1,139334501 |
| N3 | 0,977415159 | 0,590343678 | 1,026865551 | 0,876695954 | 0,345709493 | 0,582900753 | 2,336784039 | 1,487356495 | 0,374327279 | 0,94939265 |

| Akt_pT308 | AMPK-a2_pS345 | AMPKa | AMPKa_pT172 | AR | ARID1A | Atg3 | Atg7 | ATM | ATM_pS1981 | ATR_pS428 |
|---|---|---|---|---|---|---|---|---|---|---|
| 0,906277073 | 1,039448488 | 0,983553321 | 1,005736217 | 1,034786543 | 0,99261002 | 0,952801101 | 0,897605057 | 1,250909885 | 0,954248231 | 1,069764814 |
| 0,919392833 | 0,962364296 | 1,137137669 | 1,023455848 | 0,928971023 | 1,608431904 | 0,950128565 | 1,016021747 | 1,746167849 | 0,898804229 | 0,947903557 |
| 1,234067532 | 1,027639557 | 0,717918343 | 0,517299149 | 0,48095966 | 0,305538572 | 1 | 1,134149856 | 0,574129574 | 1,409557465 | 0,841320619 |
| 1,038753193 | 1,031319897 | 1,10339874 | 1,190598039 | 0,800992049 | 0,876614626 | 0,877954285 | 1,180311361 | 0,761215968 | 1,014249365 | 0,748569997 |
| 0,902597672 | 0,906582726 | 1,151562842 | 1,081831828 | 0,897823961 | 1,304870796 | 0,882493758 | 0,848368391 | 1,453886617 | 0,986112008 | 1,00480032 |
| 0,634527581 | 0,968615001 | 1,207414439 | 1,090044705 | 1,104000705 | 1,422561781 | 1,040407658 | 0,977515608 | 0,999130409 | 0,947820407 | 0,998691021 |
| 0,852323375 | 0,996417132 | 0,934347428 | 0,991054466 | 0,888861114 | 1,32772081 | 1,03825765 | 0,882056647 | 1,003323421 | 1,062369295 | 0,930525225 |
| 1,137674538 | 1,088868073 | 1,294449187 | 1,39550735 | 0,554599806 | 0,656260779 | 0,82326045 | 1,106127266 | 0,684083925 | 0,943864558 | 0,87345653 |
| 0,900585396 | 1,049738913 | 1,086657044 | 1,095100742 | 0,967657026 | 1,505963851 | 0,876599424 | 0,942562061 | 1,419579567 | 1,021712492 | 1,074122299 |
| 0,950899754 | 1,134676365 | 1,15582576 | 1,095424885 | 0,998154866 | 1,303274437 | 0,774040907 | 0,906845548 | 1,388638709 | 1,043699853 | 1,094538709 |
| 1,053278972 | 1,010177859 | 1,03032509 | 1,005909801 | 1,164581939 | 0,686681104 | 0,779805722 | 1,068196832 | 1,05207154 | 0,947708578 | 0,977508318 |
| 1,32615107 | 0,993624302 | 0,976145435 | 0,976345945 | 0,96509194 | 1,215694809 | 0,774663563 | 0,921129611 | 1,04898057 | 0,976777129 | 0,967528786 |
| 2,393555921 | 0,862057129 | 1,084457112 | 0,659587568 | 0,881157428 | 1,222825587 | 0,845691189 | 0,830651548 | 0,917011594 | 1,063707892 | 0,976547973 |
| 1,063543863 | 0,95990482 | 1,170891812 | 0,901462475 | 0,856820149 | 0,542342364 | 1,292141354 | 1,3066609 | 0,885117722 | 0,864535343 | 0,713915378 |
| 1,038580954 | 0,913359358 | 0,923746717 | 0,893018102 | 0,656064662 | 0,975996094 | 1,004696031 | 1,200922363 | 1,011501996 | 0,954818725 | 0,97081817 |
| 1,012045259 | 0,936988553 | 1,155886853 | 1,01237763 | 1,0428335 | 1,135651751 | 0,936805474 | 0,938815886 | 0,944473673 | 1,037054605 | 0,890577823 |
| 0,907896729 | 0,859532338 | 1,079404996 | 0,938347597 | 1,048556388 | 0,751212882 | 0,994364598 | 1,112034836 | 1,490479498 | 0,960607868 | 0,976429598 |
| 0,863919693 | 0,964201089 | 1,125344876 | 1,050463834 | 0,909773111 | 1,475606851 | 0,929740052 | 1,002662306 | 1,749524112 | 0,929096619 | 0,936600607 |
| 0,801263831 | 1,094527698 | 1,127773688 | 1,182583672 | 0,847770527 | 1,504449406 | 0,807320289 | 0,905550182 | 1,274661418 | 0,924095499 | 0,933109343 |
| 1,266834357 | 0,821468171 | 1,120350589 | 0,82510951 | 0,710140773 | 0,565699336 | 0,884656242 | 1,097439935 | 0,70749545 | 1,07576532 | 0,899108485 |
| 1,090086204 | 1,046945894 | 0,813719584 | 0,755305277 | 0,696394513 | 0,964102876 | 0,792287944 | 0,874181241 | 0,763870683 | 1,218618686 | 0,935249645 |
| 2,025598422 | 1,071080068 | 0,567733495 | 0,425164667 | 0,744776102 | 0,85915432 | 0,502339187 | 0,592050323 | 1,002715565 | 1,369705949 | 0,915339048 |
| 1,558861966 | 1,037529874 | 0,981108579 | 1,001467012 | 0,93140676 | 1,038421038 | 0,703860751 | 1,01215664 | 1,54227391 | 1,025681767 | 0,992331284 |
| 1,213196916 | 1,023873253 | 1,170211407 | 1,216501015 | 0,815353401 | 1,439965449 | 0,911292024 | 0,969610934 | 1,068481297 | 0,932377422 | 0,965862452 |
| 1,038948409 | 1,016353635 | 0,955406936 | 0,423815167 | 2,056643929 | 0,664354985 | 1,115550242 | 0,636867421 | 0,969524908 | 1,19739591 | 1,249585999 |
| 0,996610783 | 1,02166068 | 1 | 0,696828063 | 1,918041384 | 0,514336123 | 1,055366827 | 0,736479036 | 0,926380569 | 1,134027187 | 1,072491495 |
| 0,810915732 | 1,025700295 | 1,582223106 | 0,659657529 | 2,313458017 | 0,465951245 | 1,07686836 | 0,849734189 | 1,102360025 | 1,26998829 | 0,82518238 |

| Aurora-B | Axl | b-Actin | b-Catenin | Catenin_pT41_S45 | B-Raf | B-Raf_pS445 | B7-H4 | Bad_pS112 | Bak | Bax |
|---|---|---|---|---|---|---|---|---|---|---|
| 1,18449035 | 1,087090696 | 1,414838545 | 1,056757564 | 1,139319141 | 1,223994396 | 1,061436479 | 0,890897194 | 0,948628259 | 0,874344503 | 1,003675359 |
| 1,007131724 | 0,925581062 | 0,887372578 | 1,531516363 | 1,237116735 | 1,292053137 | 1,175672764 | 0,901742913 | 1,017092646 | 1,033275584 | 1,032030568 |
| 0,991316655 | 0,954169469 | 1,151239261 | 0,415157814 | 1,321509279 | 0,570035416 | 0,809274063 | 1,135132098 | 1,088430419 | 1,178829844 | 0,62412262 |
| 1,013914341 | 0,745866166 | 1,162439661 | 0,699102862 | 1,032959535 | 0,95749834 | 1,017230248 | 0,924385515 | 1,162873541 | 0,693073514 | 0,883814664 |
| 1,07858742 | 0,956894829 | 0,836317598 | 1,417942978 | 1,176608926 | 1,035404321 | 0,87481818 | 0,973262461 | 1,068368231 | 0,897358066 | 0,879263858 |
| 0,954223754 | 1,204692915 | 1,05491283 | 1,109456768 | 0,947246857 | 1,086365493 | 0,53246517 | 1,078336519 | 0,68947732 | 1,007535349 | 1,104729135 |
| 0,957188586 | 0,935070648 | 0,966475676 | 0,760087919 | 1,150623757 | 0,880508094 | 0,779410705 | 0,937752587 | 0,870622717 | 0,964314864 | 1,176870188 |
| 1,031980975 | 0,841439381 | 1,045712538 | 0,86604425 | 1,188970298 | 1,083501823 | 1,308020672 | 0,961137001 | 1,222114906 | 0,821124106 | 0,909665389 |
| 1 | 0,69688318 | 0,833791744 | 1,068331143 | 1,252959539 | 1,349647405 | 1,271886779 | 0,958010684 | 1,02085139 | 0,930668195 | 1,310090331 |
| 0,907109041 | 1,056512193 | 1,062045188 | 1,202932251 | 1,105390737 | 1,179795334 | 1,156920715 | 0,964129424 | 1,074399405 | 1,135855839 | 1,041258404 |
| 1,028824741 | 1,094638832 | 1,097191901 | 1,064131705 | 0,951569777 | 1,231772081 | 1,060773619 | 1,053930163 | 0,968755511 | 0,928344324 | 0,870128269 |
| 0,97952723 | 0,977130346 | 0,963129348 | 0,842261193 | 1,067683553 | 1,034858574 | 1,038245784 | 0,949348223 | 1,00010892 | 0,923649979 | 0,995272542 |
| 0,869633496 | 1,165473776 | 1,024501989 | 1,430524229 | 1,155973698 | 1,154545064 | 1,177654184 | 1,112340422 | 1,197551203 | 1,186517395 | 0,860057172 |
| 1,01676825 | 0,973653862 | 1,23737939 | 1,034546851 | 1,198460601 | 0,893191744 | 1,296289679 | 1,044722596 | 1,30235415 | 1,141140537 | 1,007921252 |
| 0,915940562 | 0,962491778 | 1,5449353 | 1,237883177 | 1,199649718 | 0,946873852 | 1,078239539 | 0,913595313 | 1,206288219 | 1,06207767 | 1,207922839 |
| 0,962553983 | 0,839235759 | 1,256094867 | 1,03586576 | 0,911733989 | 0,880073965 | 0,944948797 | 1,013201595 | 1,001858327 | 0,939317157 | 0,932091887 |
| 1,096233911 | 1,083593244 | 1,470924543 | 0,87768101 | 1,019013065 | 1,172420033 | 1,282318686 | 0,981485639 | 1,100437513 | 1,14355804 | 1,036150536 |
| 0,947018765 | 0,937396078 | 0,993015439 | 1,246157485 | 0,972293942 | 1,34045197 | 1,365706925 | 0,960749696 | 1,034414238 | 0,982339164 | 1,382173991 |
| 1,065774866 | 1,066288334 | 1,1357293 | 1,180960895 | 1,070747818 | 1,235507501 | 1,291973377 | 0,933177395 | 1,093737168 | 0,918820678 | 1,272747135 |
| 1,195423101 | 0,866195733 | 1,10632144 | 0,715211258 | 0,728262956 | 0,918922268 | 1,147715957 | 1,246219241 | 1,047488516 | 0,897487425 | 0,803470748 |
| 1,109748069 | 0,912034056 | 1,057542813 | 0,479747648 | 1,117285584 | 0,898319133 | 0,923197351 | 1,014799263 | 0,952803794 | 1,039650434 | 1,310382267 |
| 1,431256454 | 1,311203329 | 0,991958809 | 0,485579231 | 1,10060945 | 0,781873804 | 0,660045144 | 1,345783071 | 0,734882944 | 1 | 0,5269289 |
| 1,071203057 | 0,963687336 | 1,484345303 | 0,980196064 | 0,940909402 | 1,043157092 | 0,985102592 | 0,961255447 | 1,04826267 | 0,846474543 | 1,030397297 |
| 0,947307727 | 0,967205113 | 0,923606977 | 1,151318933 | 0,981354633 | 1,307091798 | 1,307501638 | 0,907705767 | 1,17163516 | 0,93614206 | 0,928592653 |
| 1,570742641 | 1,703762778 | 0,808381267 | 0,520028735 | 0,935586377 | 0,667902531 | 0,425208977 | 1,100881429 | 0,708541616 | 0,892342891 | 0,508927683 |
| 1,016567707 | 1,728155595 | 0,827151506 | 1,199560769 | 0,973550477 | 0,736810905 | 0,643466345 | 1,152184302 | 0,860218649 | 1,019157262 | 0,575664963 |
| 1,492168044 | 2,101296125 | 0,705221341 | 0,444643066 | 1,012111385 | 0,699430949 | 0,512812891 | 1,401674197 | 0,973043762 | 1,090045698 | 0,571913786 |

| Bcl-xL | Beclin | Bid | Bim | BRD4 | c-Abl | c-IAP2 | c-Jun_pS73 | c-Kit | c-Myc | C-Raf |
|---|---|---|---|---|---|---|---|---|---|---|
| 1,02812525 | 0,942337878 | 0,867086961 | 1,088330566 | 1,031485972 | 1,035652119 | 1,030350751 | 1,129897396 | 1,010752369 | 1,065990853 | 0,897389178 |
| 1,049479618 | 0,996699251 | 1,008765878 | 1,057318013 | 1,001306126 | 0,958409887 | 1,024476555 | 1,531928896 | 1,052220677 | 1,299017692 | 1,088136514 |
| 0,709650836 | 1,295232334 | 0,878728627 | 0,653061586 | 1,079156784 | 0,820733097 | 1,364696468 | 1,183833471 | 1,000982091 | 1,147864342 | 0,778773298 |
| 0,774808429 | 0,998612148 | 0,844360562 | 0,919542513 | 1,070131651 | 0,856074545 | 0,964524365 | 1,082107816 | 1,005060253 | 1,165969073 | 0,907180267 |
| 1,002844274 | 0,903310071 | 0,936875551 | 0,961513103 | 1,018457063 | 0,857730878 | 1 | 1,287395119 | 1,012433073 | 1,116426133 | 1,017986523 |
| 1,169576179 | 0,976451837 | 1,026213329 | 1,510443173 | 0,985320707 | 1,162115946 | 1,002932635 | 0,745408287 | 1,011017669 | 0,839294358 | 0,984255747 |
| 1,388941037 | 1,01891683 | 1,037083184 | 1,30015777 | 1,110795888 | 0,9237513 | 1,088097092 | 1,137161874 | 1,050183739 | 1,079290679 | 0,972730721 |
| 0,982148759 | 0,966360133 | 0,826454436 | 1,403947784 | 1,089663486 | 0,816896168 | 0,961244037 | 1,270206362 | 0,984153951 | 1,062644675 | 1,000572219 |
| 0,961366513 | 0,913945405 | 0,950496335 | 1,072384157 | 1,006859012 | 1,021287842 | 1,055514055 | 1,098263047 | 1,036740704 | 1,089152633 | 1,032202142 |
| 0,96204955 | 0,963017775 | 1,156347774 | 1,58410474 | 1,016487478 | 0,934767227 | 0,810493521 | 1,322685114 | 1,001854448 | 1,271264311 | 1,049850659 |
| 1,042903124 | 0,89919724 | 0,888649406 | 0,973506738 | 0,903896155 | 0,927378367 | 0,836874959 | 0,761183577 | 0,841129909 | 0,829512094 | 1,057736083 |
| 1,028111046 | 0,953761915 | 1,027835641 | 1,244451012 | 0,911619918 | 1,106414618 | 0,980343069 | 0,913888045 | 0,972393649 | 1,168213481 | 1,019322995 |
| 0,963159274 | 1,034068169 | 0,905508399 | 0,947613892 | 0,920670803 | 1,182446114 | 0,993903451 | 1,019750728 | 0,899439672 | 1,022886854 | 0,97047247 |
| 0,809150092 | 1,158135454 | 0,96772053 | 1,070831529 | 1,028212103 | 1,081521573 | 0,970063076 | 0,934382083 | 0,818001065 | 0,977614878 | 0,813261655 |
| 1,019839124 | 1,031290179 | 0,901020239 | 1,048629397 | 1,073533708 | 0,931758096 | 1,096511572 | 1,45789205 | 0,962584025 | 1,121590387 | 0,969668697 |
| 1,194028993 | 1,142404416 | 1,103615663 | 0,891577392 | 0,957505241 | 0,949296343 | 1,053209743 | 1,082773054 | 0,95763726 | 0,906809263 | 0,943371259 |
| 0,920349426 | 0,948778337 | 1,051469718 | 1,290934703 | 0,923348174 | 0,837550428 | 1,012969497 | 1,056468866 | 0,933617766 | 0,911647579 | 1,029645422 |
| 1,017650891 | 0,907851307 | 1,168389286 | 1,069666864 | 0,962992132 | 1,005470806 | 1,050334743 | 1,569976703 | 1,035995554 | 1,486145285 | 1,117363905 |
| 1,153575149 | 0,890708141 | 1,04856261 | 1,155439679 | 1,025719333 | 1,039660708 | 1,071520265 | 1,305735811 | 0,982304013 | 1,02481481 | 1,126245699 |
| 0,797717794 | 1,044315547 | 1 | 0,823319837 | 1,137632313 | 0,846172205 | 0,964469849 | 1,310389106 | 0,964879176 | 1,018365263 | 0,810570357 |
| 1,006252656 | 0,961338323 | 1,240505506 | 1 | 1,031404503 | 0,889282632 | 1,042932212 | 1,132083415 | 0,911906068 | 1,136033269 | 0,915854414 |
| 0,675052583 | 1,406292641 | 0,67186617 | 0,725642062 | 1,613092877 | 0,575406234 | 0,685730028 | 0,948938437 | 1,160410471 | 0,943078712 | 0,666101539 |
| 1 | 0,807247744 | 0,85117856 | 0,801288626 | 0,91383452 | 0,940875593 | 0,803162154 | 0,878472301 | 1,043869389 | 1,04164572 | 1,067018148 |
| 0,854721764 | 0,964868303 | 0,887642712 | 1,083459665 | 1,018494823 | 1,008699993 | 1,027461922 | 1,133361907 | 1,037179429 | 1,392704517 | 1,050136418 |
| 0,7165335 | 0,835860413 | 1,118523593 | 0,631391532 | 1,545719658 | 1,079336223 | 0,869235926 | 1 | 0,764951697 | 0,965730945 | 0,669777722 |
| 0,82132354 | 1,042860458 | 1,388357397 | 0,715912341 | 1,463334493 | 1,03972651 | 0,823789435 | 1,282690361 | 0,783010752 | 0,877992605 | 0,661427409 |
| 0,874607689 | 1,269364038 | 1,467575094 | 0,693285152 | 1,773393339 | 0,996350895 | 0,93903938 | 1,272102378 | 0,83180102 | 0,814455707 | 0,673747483 |

| C-Raf_pS338 | Caspase-3 | Caspase-7-cleaved | Caveolin-1 | CD134 | CD20 | CD4 | cdc25C | Cdc2_pY15 | CDK1 | Chk1_pS296 |
|---|---|---|---|---|---|---|---|---|---|---|
| 0,867295438 | 1,000959699 | 0,925511774 | 0,479700257 | 0,871265319 | 0,775336088 | 0,921174923 | 0,945371715 | 0,914436008 | 1,330130543 | 1,035727144 |
| 1,082200845 | 1,055642329 | 1,471618901 | 0,577119438 | 0,922656268 | 1,025992994 | 0,991215887 | 0,917730545 | 1,068987286 | 1,533934756 | 0,805132758 |
| 0,760761809 | 1,020651627 | 1,656418837 | 0,913615018 | 1,175355304 | 1,426420718 | 0,896941767 | 0,832577116 | 1,007947697 | 0,621350221 | 1,401653969 |
| 1,245358426 | 1,042150336 | 1,42458202 | 0,496106717 | 0,819133732 | 0,895216074 | 0,774153016 | 0,987824652 | 0,981978466 | 1,088184206 | 1,268036046 |
| 0,987804014 | 1,121716925 | 0,964800664 | 0,401798549 | 0,85069715 | 0,777259699 | 0,887040712 | 1,006435906 | 1,081912856 | 1,292210402 | 0,783906993 |
| 0,60476035 | 1,208024556 | 1,135259287 | 0,808348742 | 0,926727113 | 1,10342551 | 1,083672538 | 1,019068139 | 0,824837575 | 1,650737269 | 1,336168974 |
| 0,81918803 | 1,105451397 | 0,881513895 | 0,527176076 | 0,944542981 | 0,920692748 | 0,971771899 | 1,244295796 | 0,841812414 | 1,406006635 | 0,929440861 |
| 1,353491634 | 1,071133213 | 0,895501141 | 1,29428858 | 0,851056776 | 0,912653434 | 0,894653217 | 0,982257452 | 1,087805217 | 1,249829473 | 0,992811591 |
| 0,964934684 | 0,999856339 | 1,069529537 | 0,4739244 | 0,930939972 | 0,797464757 | 0,938364634 | 0,924374332 | 0,895934448 | 1,373034655 | 0,898887702 |
| 0,884956688 | 1,397009276 | 1,342052156 | 0,846105771 | 0,877189442 | 0,894161711 | 0,84724859 | 0,783981702 | 1,15104014 | 1,190267401 | 0,999815562 |
| 0,998305431 | 0,788089429 | 1,594853655 | 0,543298329 | 1,055133054 | 0,9357062 | 0,775162183 | 0,778138271 | 0,876193576 | 1,024330131 | 1,20818014 |
| 0,971243396 | 1,030067002 | 0,859727914 | 0,566834893 | 0,961163576 | 0,917600511 | 0,80839878 | 0,82969613 | 1,088759348 | 1,362594587 | 0,918829458 |
| 1,265824216 | 0,941092439 | 1,228988608 | 7,583045684 | 0,910504853 | 1,057044541 | 0,907386727 | 1,073638504 | 0,803196327 | 0,627222038 | 1,011294869 |
| 1,181887288 | 0,916095557 | 1,042560376 | 5,192564468 | 0,86348801 | 1,438374133 | 0,791148521 | 1,140330711 | 0,872934898 | 0,620217611 | 1,649064774 |
| 0,900804745 | 1,023752558 | 1,353285357 | 0,562129699 | 0,93747421 | 1,101572723 | 0,922699262 | 0,732041258 | 0,932686296 | 1,15206987 | 2,393576918 |
| 0,880085552 | 1,029179469 | 1,185014134 | 1,088241416 | 0,94779079 | 1,14669623 | 0,920760915 | 0,906902156 | 0,992069432 | 0,973183086 | 1,307163094 |
| 1,271265267 | 0,990835302 | 1,236496459 | 0,944898342 | 0,885294343 | 1,096610194 | 0,877773094 | 1,229215983 | 1,110137571 | 1,172088445 | 1,492848628 |
| 1,082379719 | 1,163663349 | 0,857339649 | 0,80321579 | 0,918668782 | 0,919920257 | 0,955964239 | 0,991051272 | 1,030702958 | 1,371670732 | 0,92291082 |
| 1,358700295 | 1,148124041 | 0,711305922 | 0,797172169 | 0,844337092 | 0,869872351 | 0,88132508 | 1,315268248 | 0,89978305 | 1,222401633 | 1,03030341 |
| 1,023083781 | 0,97742663 | 0,917060382 | 1,415775029 | 1,01363621 | 1,330427002 | 1,061827022 | 1,034505738 | 0,988947286 | 0,838908303 | 1,348332517 |
| 1,144444169 | 1,147821445 | 1,122981971 | 0,3719211 | 0,861437391 | 0,998721011 | 0,980282838 | 0,929740945 | 0,936925023 | 1,166561163 | 0,986187429 |
| 1,075680646 | 1,364457488 | 0,695537312 | 2,42148905 | 0,59209477 | 1,715686572 | 1,962650888 | 1,454093415 | 0,974919174 | 1,085186283 | 1,226863681 |
| 1,070105172 | 0,840554838 | 0,896225802 | 0,939279714 | 0,712881511 | 0,983530707 | 0,719331316 | 0,759639113 | 1,047604692 | 1,5029133 | 1,384105564 |
| 1,472577509 | 0,993161813 | 0,820098726 | 1,034863972 | 0,988682913 | 0,952062335 | 0,955650914 | 0,903676673 | 1,025782305 | 1,502202511 | 0,935357221 |
| 1,072255545 | 0,907308002 | 0,865146641 | 2,216818839 | 1,234439539 | 1,350983279 | 1,208917135 | 2,837740832 | 1,120047929 | 0,730886764 | 0,81530021 |
| 1,192211116 | 0,967561694 | 1,066671931 | 7,233938264 | 1,142255671 | 1,320647366 | 1,150216031 | 2,966050295 | 1,260726922 | 0,792653889 | 0,859406067 |
| 1,252462698 | 0,995048436 | 0,940254744 | 20,75855182 | 1,2790254 | 2,199771741 | 1,293910141 | 2,78259476 | 1,046401161 | 1,193026237 | 0,980667937 |

| Chk2_pT68 | Claudin-7 | COG3 | Collagen-VI | Connexin-43 | Cox-IV | Cox2 | Creb | Cyclin-B1 | Cyclin-D1 | D-a-Tubulin |
|---|---|---|---|---|---|---|---|---|---|---|
| 0,856951624 | 1,089833776 | 1,124177148 | 0,808835596 | 1,106836373 | 0,941849188 | 0,97932357 | 0,978442357 | 0,869384814 | 0,981715786 | 0,938087843 |
| 1,002629451 | 1,509736017 | 0,852957157 | 0,891031502 | 0,948536741 | 0,915806141 | 0,995550269 | 1,024570722 | 0,915806141 | 0,962477144 | 1,009210002 |
| 1,111968503 | 0,908164 | 0,93483863 | 1,755729294 | 0,677509944 | 1,131397804 | 1,002453993 | 0,993968502 | 1,131397804 | 1,21863609 | 1,28252324 |
| 1,180429311 | 0,889878605 | 0,955582022 | 1,006786185 | 0,995920245 | 1,183189393 | 1,364693696 | 0,901656608 | 1,183189393 | 1,097666071 | 1,041964469 |
| 1,056126566 | 1,503343769 | 1,08258623 | 0,92222927 | 0,781972937 | 0,787643965 | 0,968077077 | 0,981872624 | 0,787643965 | 1,008012254 | 0,94209367 |
| 1,165901745 | 1,242272958 | 1,10136603 | 0,930498847 | 1,040417236 | 1 | 1,031721376 | 1,042446322 | 1 | 1,462593394 | 1,323577202 |
| 1,212816431 | 1,566390081 | 1,055773847 | 0,954448696 | 1,537724652 | 0,92930878 | 0,926266376 | 1,027377838 | 0,92930878 | 0,917632337 | 1,056515545 |
| 1,232645473 | 1,234427302 | 0,862636233 | 1,065954593 | 1,345336369 | 1,334799989 | 0,781790076 | 0,943838491 | 1,334799989 | 0,98731963 | 0,881044298 |
| 1,093771681 | 1,266547948 | 0,987702213 | 0,742729058 | 1,224230083 | 0,890984391 | 0,90419881 | 1,03569998 | 0,890984391 | 0,890852346 | 0,887201494 |
| 1,051903516 | 1,918802212 | 1,054638615 | 1,087574167 | 0,940341485 | 0,785632569 | 1,025667724 | 0,949100874 | 0,785632569 | 1,020541428 | 1,066657404 |
| 0,664719451 | 1,045652327 | 1,140437243 | 0,940499033 | 1,02503075 | 1,057420285 | 1,208436834 | 0,995817378 | 1,057420285 | 1,216293668 | 0,940640738 |
| 1,055730171 | 0,830178731 | 0,864524641 | 1,231259538 | 0,990305733 | 0,983226513 | 0,969679046 | 0,983840956 | 0,983226513 | 1,019815783 | 0,994720386 |
| 0,862321096 | 1,341299537 | 1,117482551 | 0,837601406 | 1,355625416 | 0,807683589 | 0,981432885 | 0,963077665 | 0,807683589 | 0,944659976 | 0,946266786 |
| 1,048541539 | 0,9745352 | 0,985755734 | 1,429696027 | 0,47900067 | 1,129335468 | 0,733697925 | 0,975560892 | 1,129335468 | 1,08162973 | 0,858747131 |
| 1,075802404 | 0,994695359 | 1,124952325 | 1,042784998 | 0,886944447 | 0,898645048 | 0,901587064 | 1,090832337 | 0,898645048 | 0,98145023 | 0,987495657 |
| 1,159810274 | 1,209315208 | 0,915075574 | 1,175183476 | 0,987355313 | 0,943653081 | 1,082969044 | 0,993784298 | 0,943653081 | 1,218609116 | 1,091432021 |
| 1,078250398 | 0,801810332 | 0,979854142 | 1,595750096 | 0,991577307 | 0,792985253 | 1,041451823 | 0,918882281 | 0,792985253 | 1,009470291 | 0,917361098 |
| 1,154008836 | 0,984884715 | 1,109028393 | 0,965498381 | 1,309507173 | 0,92304201 | 0,87854597 | 0,939084734 | 0,92304201 | 0,900719634 | 0,825394496 |
| 1,106712255 | 0,84686274 | 0,939831516 | 0,984321333 | 0,891894043 | 0,768032445 | 0,920841824 | 0,956485157 | 0,768032445 | 1,004358822 | 0,921979594 |
| 1,031524917 | 1,087900265 | 1,252374885 | 1,246551344 | 0,625525043 | 1,374071717 | 1,083370411 | 0,87634242 | 1,374071717 | 0,974613556 | 1,053544039 |
| 1,005322321 | 1,236897248 | 1,059581956 | 1,00896321 | 1,207959352 | 1,263434755 | 0,830633842 | 1,011825337 | 1,263434755 | 0,861888333 | 0,9087405 |
| 1,107296093 | 1,294522217 | 1,035721193 | 1,577995211 | 0,56418926 | 1,975982845 | 1,166217824 | 0,8039812 | 1,975982845 | 1,077293453 | 1,137387217 |
| 0,679801562 | 0,901030908 | 1,02508114 | 0,850421365 | 1,222489077 | 0,986457081 | 1,049217687 | 0,989055351 | 0,986457081 | 0,964578443 | 1,00319944 |
| 0,951460701 | 1,060401046 | 0,977993241 | 1,117617953 | 0,9613685 | 0,949246349 | 0,974307807 | 0,948653141 | 0,949246349 | 0,976983602 | 0,953276611 |
| 0,928430089 | 0,987240455 | 0,981538285 | 1,227351007 | 0,264894721 | 1,275737202 | 0,875563726 | 0,884070618 | 1,370364777 | 0,994589323 | 0,635117799 |
| 1,161562075 | 1,389829522 | 1,26372227 | 1,344971297 | 0,188643867 | 1,213257289 | 0,960815344 | 0,871941541 | 1,502541425 | 1,014646176 | 0,837628026 |
| 1,046064088 | 0,903366834 | 0,86278296 | 1,734807856 | 0,32674927 | 2,172345353 | 0,859555653 | 0,877532394 | 2,172345353 | 1,003039528 | 0,951453208 |

| DJ1 | DM-Histone-H3 | DM-K9-Histone-H3 | DUSP4 | E-Cadherin | eEF2 | eEF2K | EGFR | EGFR_pY1173 | eIF4E | eIF4E_pS209 |
|---|---|---|---|---|---|---|---|---|---|---|
| 1,158786545 | 0,978867184 | 1,124872792 | 1,134565628 | 1,1808459 | 1,194232448 | 0,857683052 | 1,201572786 | 0,95159528 | 1,072947362 | 0,959044531 |
| 1,198327219 | 1,24674163 | 1,341550787 | 0,790429848 | 1,801175199 | 1,077363279 | 1,022673739 | 1,136698087 | 0,98650542 | 0,943507446 | 0,956035749 |
| 0,87308216 | 0,859348889 | 1,052595139 | 0,570677023 | 0,092434432 | 0,723440718 | 0,811762264 | 1,023661592 | 1,151503808 | 0,605954477 | 0,944193822 |
| 0,808677842 | 1,043399199 | 1,010566332 | 0,986784734 | 0,761652238 | 0,864575423 | 0,848922012 | 0,955183024 | 0,989660337 | 0,787752059 | 0,958610867 |
| 0,943132865 | 1,038179371 | 1,090358702 | 1,131407192 | 1,329958384 | 0,99636624 | 1,006609374 | 1,160676215 | 0,968312919 | 0,925445599 | 0,997569902 |
| 0,868166191 | 0,978800311 | 1,00561982 | 1,411565467 | 0,974262653 | 0,791294984 | 0,8064532 | 1,273581292 | 0,996449798 | 0,711592366 | 0,901050179 |
| 0,937759193 | 1,229882697 | 1,298035438 | 1,131357371 | 1,508505265 | 1,061020457 | 1,02988088 | 1,111890951 | 1,066809816 | 1,079980081 | 0,971937903 |
| 0,85802021 | 0,923093424 | 1 | 0,6640873 | 0,382771605 | 1,037437814 | 0,957700516 | 0,994808894 | 1,140517092 | 1,021502107 | 1,073599126 |
| 1,313275279 | 1,250930154 | 1,302016257 | 0,919865829 | 1,254735772 | 1,183970927 | 0,930800861 | 1,118161304 | 1,071014854 | 1,525427502 | 0,96170535 |
| 1,233350171 | 1,277987103 | 1,253084749 | 1,35021207 | 1,228874164 | 1,006056502 | 0,875943936 | 1,103754937 | 0,954250371 | 1,015940657 | 1,057931529 |
| 1,28960445 | 1,096453764 | 1,051477773 | 1,298557958 | 0,961181881 | 1,077082755 | 0,78493774 | 0,956857195 | 0,905276688 | 1,189129776 | 1,057081636 |
| 0,981711664 | 1,061463312 | 1,073181293 | 0,937338428 | 1,008532155 | 1,012080788 | 0,903109697 | 0,997276586 | 0,986717165 | 1,139986926 | 0,961259521 |
| 1,270982153 | 0,783369934 | 0,877253135 | 1,007531213 | 1,905455396 | 0,792664911 | 2,053103477 | 1,352536015 | 0,964141036 | 0,826873836 | 0,895202354 |
| 1,364625996 | 0,889461945 | 0,991861714 | 0,743080386 | 0,768124783 | 0,760677067 | 1,213427416 | 0,94745712 | 0,962799011 | 0,754358537 | 0,934818361 |
| 1,212518742 | 0,847789747 | 0,943419519 | 0,849493569 | 1,218071128 | 1,026976188 | 1,034879119 | 1,230938952 | 1,045282665 | 0,825165791 | 0,898932849 |
| 0,742334088 | 0,993674652 | 1,146380569 | 0,931346261 | 1,338394499 | 0,964686943 | 0,99571023 | 1,244558631 | 1,061115433 | 0,971240509 | 0,836410578 |
| 1,178607983 | 0,881588416 | 0,882025439 | 1,056208321 | 1,226298395 | 0,850907713 | 1,051863815 | 1,114077002 | 0,954338256 | 0,887145853 | 0,837461628 |
| 1,206255841 | 0,942610213 | 1,042215613 | 0,935080026 | 1,50949452 | 1,011928725 | 0,890394325 | 1,040334284 | 1,023256396 | 1,15234593 | 0,981207932 |
| 1,098661217 | 0,97698689 | 1,016331836 | 0,87464069 | 1,529222172 | 0,817129519 | 0,854752015 | 1,055474498 | 1,040434166 | 0,965671722 | 0,997111608 |
| 0,912760796 | 0,902150393 | 0,973829524 | 0,854291159 | 0,980984919 | 0,791621303 | 0,985877437 | 1,009652404 | 1,049866485 | 0,859303746 | 0,884514508 |
| 0,689219241 | 1,093655086 | 1,080139716 | 0,700788623 | 0,968993754 | 1,028416901 | 0,906496949 | 0,960040701 | 1,003368391 | 1,233416037 | 0,965340216 |
| 1,279041327 | 0,778754238 | 0,816886547 | 1,092474827 | 0,25549001 | 0,614292997 | 1,41774034 | 0,910372655 | 0,946234515 | 0,502152986 | 0,949814838 |
| 0,974437355 | 0,776973889 | 0,87596045 | 1,133456376 | 0,850073525 | 1,828150787 | 0,999734615 | 0,966932709 | 0,97075509 | 1,11326491 | 1,028070973 |
| 0,930023773 | 1,025784289 | 1,034909685 | 1,067356774 | 1,349151774 | 1,069102306 | 0,906047827 | 1,096538851 | 1,042273732 | 1,203212725 | 0,990266875 |
| 1,2367623 | 1,224858699 | 0,971931065 | 1,721843353 | 0,888500872 | 0,701786993 | 1,608155822 | 0,93193392 | 0,918366682 | 0,433074134 | 1,256128208 |
| 0,775184714 | 1,248976985 | 0,88959639 | 1,714060662 | 1,217765089 | 0,693788313 | 1,845882574 | 0,999998924 | 0,898779416 | 0,561125829 | 1,246201557 |
| 2,175692268 | 1,255732627 | 0,718557078 | 1,535430576 | 0,351676173 | 0,45960962 | 1,558627721 | 0,715540739 | 0,977857936 | 0,391721541 | 1,204258998 |

| eIF4G | Elk1_pS383 | ER | ER-a_pS118 | ERCC5 | Ets-1 | FAK | FAK_pY397 | FASN | Fibronectin | FoxM1 |
|---|---|---|---|---|---|---|---|---|---|---|
| 1,315013646 | 0,928791664 | 0,919175582 | 1,012496168 | 0,924702287 | 0,897500837 | 0,849999366 | 0,911112828 | 1,360176881 | 0,638985218 | 0,956389904 |
| 1,431504612 | 0,90363113 | 1,066580732 | 1,071036268 | 0,912029905 | 1,001418681 | 0,981177295 | 0,893500385 | 1,061135918 | 0,510686146 | 0,918773117 |
| 0,481111903 | 1,435080806 | 1,432993438 | 0,972915058 | 1,803003043 | 1,265923565 | 1,167232966 | 1,012878956 | 0,336111395 | 3,14184878 | 1,0758361 |
| 0,701737018 | 1 | 0,842279874 | 0,958603573 | 1,096526263 | 0,894891944 | 0,592189025 | 1,06488503 | 0,757266569 | 4,023726701 | 1,009443613 |
| 1,23708784 | 1,009656335 | 0,826005727 | 1,07693309 | 1,201436495 | 0,91873935 | 0,896009694 | 1,033859784 | 0,857954345 | 0,610087608 | 0,989112629 |
| 1,06745528 | 0,878538737 | 0,98691606 | 1,060905892 | 1,078788353 | 1,046902972 | 1,099777003 | 0,789209671 | 0,691242972 | 0,984812536 | 0,894007474 |
| 1,332188708 | 0,981059606 | 0,89001465 | 1,111379608 | 0,926638896 | 1,03826658 | 0,959360201 | 0,971177957 | 1,043987445 | 2,24661883 | 1,012942802 |
| 1,242003361 | 1,000425105 | 0,881184811 | 1,109953601 | 0,912348101 | 0,800563464 | 0,94747494 | 1,134686946 | 0,927695963 | 2,023763514 | 0,749013376 |
| 1,079354609 | 0,925866824 | 0,870744695 | 1,081240609 | 0,866850284 | 0,881820409 | 1,021430464 | 1,128030369 | 1,002802969 | 0,803790761 | 0,920452437 |
| 1,015754048 | 0,901916493 | 0,949465159 | 1,033829801 | 0,985731004 | 0,952723122 | 0,865116865 | 0,989584838 | 0,758443443 | 1,1893404 | 1,252523204 |
| 0,893844666 | 0,920553181 | 1,050787538 | 0,828472596 | 0,821758286 | 1 | 0,838857793 | 0,922777908 | 1,221927868 | 1,092726455 | 1,062197781 |
| 1,070758025 | 1,073152631 | 0,989393874 | 0,97239421 | 0,880977828 | 0,956918684 | 0,976396295 | 1,311771699 | 0,917663922 | 2,054315063 | 0,989504245 |
| 0,913515991 | 1,004980034 | 0,986002608 | 0,970253317 | 0,895348989 | 0,961624318 | 0,742808953 | 1,060625921 | 0,559199902 | 0,567892936 | 1,223634694 |
| 0,586515788 | 1,014213514 | 1,39597075 | 0,811749309 | 2,039669873 | 1,165163655 | 1,171910758 | 1,055099531 | 0,600073137 | 0,86866115 | 0,968473072 |
| 1,049308134 | 0,910591915 | 1,088291486 | 0,908545509 | 1,626778971 | 1,085971585 | 0,904887701 | 0,768840475 | 0,583171847 | 0,926141112 | 0,953127235 |
| 0,964529409 | 1,141076938 | 1,115107235 | 0,980878313 | 1,062994268 | 1,077111957 | 1,033312789 | 0,966637089 | 0,843537342 | 3,082385077 | 0,922579843 |
| 0,775442132 | 0,911070767 | 1,021675499 | 0,969538787 | 0,974167561 | 0,975341607 | 0,877751763 | 0,984504773 | 0,74813353 | 2,08363982 | 1,19773172 |
| 1,107861159 | 1,022102191 | 0,869963023 | 1,135796472 | 0,9588583 | 0,880084106 | 0,824441948 | 1,165987317 | 0,979806119 | 0,937060677 | 0,962177777 |
| 1,198202529 | 1,075371223 | 0,82081471 | 1,091483729 | 0,908695733 | 0,892330032 | 0,810418515 | 1,755593074 | 0,824478588 | 3,074857993 | 1,302351482 |
| 0,748951433 | 1,234508518 | 1,008209983 | 0,980511458 | 0,997653816 | 1,008006538 | 1,111237372 | 1,292400372 | 0,53251362 | 1,619486766 | 1,189882783 |
| 1,060497804 | 1,187389535 | 0,873718443 | 0,925620616 | 0,991396898 | 0,860462281 | 0,791052713 | 1,222201535 | 1,215261159 | 2,425964719 | 0,817076861 |
| 0,441468568 | 1,078924065 | 1,385405254 | 0,87228443 | 1,284037723 | 1,174330375 | 1,14883508 | 1,10089117 | 0,450342168 | 1,043611817 | 1,099243877 |
| 0,876954142 | 0,904710302 | 0,952275344 | 0,867398504 | 0,989291151 | 1,086887459 | 0,890328885 | 1,134260787 | 1,020596331 | 1,557288478 | 0,926260438 |
| 1,091052344 | 0,997033984 | 1,0215505 | 0,984235713 | 0,935331683 | 0,959009569 | 1,086444692 | 1,359313882 | 0,875073151 | 0,776395861 | 0,946171035 |
| 0,771882222 | 1,266421447 | 1,178143079 | 1,018640234 | 1,856251941 | 1,016231296 | 1,349020309 | 1,382480254 | 4,092788879 | 0,168156724 | 1,621542176 |
| 1,179368564 | 0,906656204 | 1,186766968 | 1,15064117 | 1,473085515 | 0,954848358 | 1,093638483 | 1,326775825 | 4,474527914 | 0,189676912 | 1,518270734 |
| 0,511065174 | 0,975808903 | 1,280310171 | 1,259110064 | 1,281834569 | 1,164837493 | 1,247436959 | 1,815759712 | 4,007109025 | 0,397251237 | 1,169991856 |

| FoxO3a | (p_S318-S321) | G6PD | Gab2 | GATA6 | GCLM | GCN5L2 | Glutamate-D1-2 | Glutaminase | Granzyme-B | GSK-3a-b_pS21_S9 |
|---|---|---|---|---|---|---|---|---|---|---|
| 0,901597741 | 1,066827872 | 0,862699125 | 0,897094838 | 1,050870295 | 0,898350766 | 1,1937865 | 1,12985667 | 0,918771986 | 0,811095455 | 0,691021159 |
| 0,958584535 | 1,168062291 | 1,008756349 | 1,447456966 | 0,948374276 | 0,91644396 | 1,258503663 | 1,00001721 | 0,968517755 | 0,936931836 | 0,902155422 |
| 0,818162334 | 1,054353094 | 0,987818495 | 1,066693973 | 0,858195457 | 1,322310958 | 0,829126986 | 0,888883268 | 1,048129271 | 1,18080982 | 1,385143917 |
| 0,933632025 | 1,101101377 | 1,150403522 | 1,100954749 | 0,916842113 | 0,986074136 | 0,947809273 | 0,708719315 | 0,998690898 | 0,793881676 | 1,50702592 |
| 0,906799054 | 1,099650283 | 0,883506365 | 1,382565569 | 0,921090312 | 0,985499208 | 0,997213624 | 0,880065096 | 0,863894469 | 0,87848391 | 1,108181784 |
| 1,055114983 | 0,877271239 | 1,681225301 | 1,43174921 | 0,989914436 | 0,882319717 | 1,065266927 | 0,914925471 | 1,170732102 | 1,015226117 | 0,410957441 |
| 0,975838039 | 0,99259185 | 1,253963628 | 1,221446722 | 1,033404119 | 0,945422455 | 1,224326902 | 1,044474324 | 0,956249859 | 0,951662156 | 0,526581558 |
| 0,932026706 | 1,09818278 | 0,668416356 | 1,223675712 | 0,9286139 | 0,967358838 | 1,247681695 | 0,623187535 | 0,785281223 | 0,714220079 | 1,287444941 |
| 0,946216541 | 1,092511654 | 0,921212249 | 1,439950969 | 1,111246172 | 0,902600306 | 1,256146405 | 0,867030633 | 0,989479607 | 0,933395087 | 0,888799642 |
| 0,909596412 | 1,07413216 | 0,933749041 | 1,224913122 | 0,942323314 | 1,030651997 | 1,243890793 | 0,95360325 | 0,955508072 | 0,971399726 | 0,884698807 |
| 0,850032511 | 0,955947098 | 1,565894733 | 0,983676049 | 0,880669604 | 0,83855294 | 0,983755115 | 1,049600994 | 1,106631629 | 0,941383622 | 0,816755418 |
| 0,949908905 | 1,186122123 | 0,947147889 | 0,956511201 | 1,127800198 | 1,001886386 | 1,268325965 | 0,921341154 | 0,860333304 | 0,902823847 | 1,958136521 |
| 0,961061057 | 1,248565003 | 0,920703193 | 1,362716445 | 0,951449145 | 0,915666547 | 1,01945645 | 2,162940534 | 0,978541582 | 1,250460317 | 1,357101355 |
| 1,100010379 | 0,994256141 | 0,724254604 | 1,160020716 | 0,88407564 | 0,89774784 | 0,932631577 | 1,139202496 | 0,675765345 | 1,297127956 | 1,270951831 |
| 1,029912637 | 1,102051275 | 1,074838414 | 1,446878522 | 0,749885255 | 0,940204584 | 1,118437752 | 1,211839579 | 1,024608462 | 0,938852388 | 0,95277958 |
| 0,968156615 | 1,164011955 | 0,996253917 | 0,999852304 | 0,973447401 | 0,990839025 | 0,784586424 | 0,914287157 | 0,916296153 | 0,975513274 | 1,266448852 |
| 0,94778066 | 1,139643606 | 0,932636741 | 1,240236659 | 0,925582218 | 0,953922679 | 1,122368942 | 1,026021678 | 0,867336276 | 1,016276432 | 1,349874825 |
| 0,975482284 | 1,202325411 | 0,812790927 | 1,314537341 | 0,993192576 | 0,879637533 | 1,451514254 | 0,929867616 | 0,963157896 | 0,908987664 | 1,175853884 |
| 0,918333044 | 1,10938472 | 0,842979527 | 1,087249688 | 1,018403197 | 0,928626586 | 1,256626539 | 1,229956361 | 0,998948702 | 0,911892963 | 1,025274039 |
| 0,91134693 | 1,05863877 | 0,861723553 | 1,15607981 | 0,968961197 | 1,038574766 | 0,754682092 | 0,904914915 | 1,1259937 | 0,883702409 | 1,297753355 |
| 0,906312271 | 1,051127053 | 1,097989747 | 0,953414364 | 0,987925158 | 0,98879726 | 1,006578083 | 0,83827496 | 0,958541646 | 0,690056664 | 1,14512816 |
| 0,757416884 | 1,146116241 | 1,022005997 | 0,949632142 | 1,055934973 | 1,171734671 | 0,723000815 | 1,028268996 | 1,255968316 | 1,231467947 | 1,064061124 |
| 0,844499803 | 1,191855861 | 0,994043072 | 0,942512899 | 0,72364903 | 0,985770057 | 1,169243702 | 1,232068544 | 0,866810916 | 0,919459922 | 1,71652024 |
| 0,876530236 | 1,151499063 | 0,789705445 | 0,98288648 | 1,025803535 | 0,905126943 | 1,237446904 | 1,099329948 | 0,889537445 | 0,685651822 | 1,779251173 |
| 1,233995216 | 0,940445913 | 1,676687067 | 0,889995726 | 1,102013345 | 1,137606047 | 0,636658082 | 1,576716688 | 1,481445168 | 1,216564264 | 1,093685117 |
| 1,294522944 | 0,919634795 | 1,573606085 | 0,923422795 | 1,095621555 | 1,202187129 | 0,566606122 | 1,500861981 | 1,17254679 | 1,28969422 | 0,828930852 |
| 1,304979789 | 0,98342439 | 1,254560308 | 1,078957433 | 1,180977839 | 1,126641509 | 0,483814674 | 1,735537977 | 1,172077005 | 1,154635146 | 1,393437464 |

| Gys | Gys_pS641 | HER2_pY1248 | HER3 | HER3_pY1289 | Heregulin | HES1 | Hexokinase-II | Histone-H3 | HSP27_pS82 | HSP70 |
|---|---|---|---|---|---|---|---|---|---|---|
| 1,049553444 | 0,912156825 | 0,880815502 | 0,884828286 | 0,961465516 | 0,93640444 | 1,077403717 | 1,037911047 | 1,22001778 | 0,939253282 | 1,112679588 |
| 0,844165331 | 0,866946898 | 0,990160641 | 0,983648542 | 1,030672207 | 0,952395239 | 0,809917896 | 1,001226438 | 2,662414786 | 0,894575803 | 0,932270848 |
| 1,21987393 | 1,072448794 | 1,10876637 | 1,484306 | 0,767414859 | 1,434153907 | 0,419360607 | 0,718634537 | 1,496465702 | 0,844698135 | 0,907682163 |
| 1,007320755 | 0,975103388 | 1,050775444 | 1,280853226 | 1,11382633 | 1,18287408 | 1,244089979 | 0,85855306 | 1,169916162 | 0,970053719 | 0,952553888 |
| 0,664715364 | 0,77713048 | 0,933165559 | 0,900195261 | 1,035981783 | 0,923704098 | 0,76867998 | 0,989822396 | 1,338732214 | 0,652394744 | 1,368367939 |
| 1,111750411 | 0,481121078 | 0,883066606 | 0,838126108 | 0,938519546 | 0,881040265 | 1,067378181 | 1,02612275 | 0,976063789 | 0,99748292 | 1,132597454 |
| 1,135344858 | 0,962424844 | 0,860840157 | 1,067377639 | 1,006989346 | 0,969812391 | 1,069754361 | 1,035557024 | 2,341093702 | 0,839960816 | 1,124031043 |
| 0,698380774 | 0,797374758 | 1,018163452 | 1,407905109 | 0,932559287 | 1,012041851 | 1,366937767 | 0,879198959 | 1,025234274 | 1,301738925 | 1,036815655 |
| 1,146341684 | 0,966298915 | 0,96922283 | 0,976833932 | 1,017432209 | 0,873865846 | 1,0444836 | 1,048731176 | 2,384375199 | 0,867567823 | 1,003874303 |
| 0,892455423 | 0,759585658 | 0,908613967 | 0,968688977 | 0,968733251 | 1,125811622 | 0,805680283 | 0,865283519 | 2,806149312 | 1,078158241 | 1,393004257 |
| 1,400956904 | 1,059079115 | 1,043517303 | 0,823095885 | 0,971626627 | 1,058936126 | 0,892684904 | 0,971185871 | 1,801946445 | 1,029442695 | 1,089175622 |
| 1,124275213 | 1,055995129 | 0,966861816 | 1,133788507 | 1,10311801 | 0,958599037 | 1,55699793 | 1,008102205 | 1,253090979 | 1,188652082 | 1,037963847 |
| 1,277333889 | 1,302415552 | 0,964090472 | 0,883100806 | 0,878585465 | 1,040181211 | 1,617557388 | 0,923854649 | 0,759847755 | 1,14313598 | 1,405227004 |
| 0,823143347 | 0,905028826 | 1,011140482 | 0,827822295 | 0,734091245 | 0,913513276 | 0,705536722 | 0,911936696 | 0,961027066 | 1,132178095 | 1,044516333 |
| 0,767520394 | 0,797088908 | 0,986711084 | 0,853146173 | 0,957002476 | 0,938481006 | 0,709569531 | 0,910887351 | 0,868763684 | 0,905011353 | 1,396334472 |
| 1,113416322 | 0,99084064 | 0,974324246 | 1,128950934 | 1,097691236 | 0,975598209 | 1,159368665 | 0,93079763 | 1,595164904 | 1,000684715 | 1,195933591 |
| 0,712309505 | 0,796682039 | 0,920527336 | 0,856662947 | 0,857539165 | 0,883532215 | 0,894518249 | 1,029361004 | 0,86242617 | 0,729330624 | 1,043986359 |
| 0,837097866 | 0,84783119 | 1,024445437 | 1,08257963 | 1,098052884 | 0,970678368 | 1,28916537 | 1,009809612 | 1,24571197 | 1,09490544 | 0,88781933 |
| 0,976976142 | 0,937050161 | 0,972646824 | 1,167741083 | 1,065121575 | 0,853860466 | 1,422997855 | 1,070327135 | 1,110867521 | 0,909579342 | 0,982042285 |
| 1,072038642 | 0,97493016 | 1,090763259 | 0,969075738 | 0,918760183 | 1,155774217 | 0,86601182 | 0,790144439 | 0,835191611 | 1,090605189 | 0,924861029 |
| 1,565989929 | 1,257089509 | 1,05838967 | 1,177215174 | 0,934570495 | 1,103503965 | 0,784392362 | 0,872096652 | 1,277608193 | 1,128533906 | 0,994319884 |
| 1,492175916 | 1,305283752 | 1,192899607 | 0,86454857 | 0,840609718 | 1,631194536 | 0,930184677 | 0,980971489 | 0,514521982 | 1,575046179 | 1,142688074 |
| 1,235004077 | 1,32034671 | 1,056364255 | 0,883314458 | 1,107413546 | 0,956158688 | 1,145716187 | 0,857246369 | 0,8072434 | 1,03643598 | 1,288727412 |
| 0,875014599 | 0,959230458 | 1,044121807 | 0,932384243 | 1,004894781 | 0,949499318 | 1,021790231 | 0,922599835 | 1,304230865 | 1,132942633 | 0,932180442 |
| 2,259451543 | 2,335791098 | 1,271574984 | 0,739396843 | 0,622853087 | 1,175690431 | 0,569550895 | 1,368664499 | 0,664237015 | 1,184192211 | 0,905364527 |
| 0,449639446 | 1,012904335 | 1,185334111 | 0,787473723 | 0,653591857 | 0,851441618 | 0,546606881 | 1,434873682 | 0,54678543 | 1,444092626 | 0,884021139 |
| 4,680427801 | 3,198836313 | 1,034126057 | 0,771868563 | 0,586678519 | 0,970144416 | 0,560971008 | 1,373071068 | 0,359652797 | 2,593680618 | 0,92640376 |

| IGF1R_pY1135_Y1136 | IGFBP2 | IGFRb | INPP4b | IR-b | IRF-1 | IRS1 | Jagged1 | Jak2 | JNK2 | JNK_pT183_Y185 |
|---|---|---|---|---|---|---|---|---|---|---|
| 0,963931986 | 0,92379931 | 1,118917668 | 1,0448453 | 1,270201311 | 0,553958262 | 0,927650127 | 0,939114462 | 0,969747864 | 0,951161215 | 1,043682722 |
| 1,021453312 | 0,872290119 | 0,835360768 | 0,965240912 | 1,104968655 | 1,082058033 | 1,017348773 | 0,925626139 | 1,013758681 | 0,98557421 | 1 |
| 1,162504905 | 0,828774673 | 0,533480543 | 1,120922497 | 0,644759668 | 1,918551167 | 0,716596994 | 1,024424025 | 0,802834607 | 0,859621301 | 1,112814 |
| 1,080143575 | 0,868889 | 1,301601865 | 0,818984233 | 1,034532518 | 0,659931484 | 0,837298829 | 1,069097911 | 0,876487686 | 0,944514823 | 1,074612049 |
| 0,970188433 | 1,150275299 | 0,756279705 | 0,88901078 | 1,341806106 | 0,56595326 | 0,95242923 | 1,012765276 | 0,928450809 | 1,159680605 | 1,003904903 |
| 0,877354978 | 1,269866479 | 1,452017158 | 0,937184406 | 1,278371861 | 1,119535041 | 1,296444175 | 1,115800533 | 1,011326477 | 1,063537695 | 0,959421681 |
| 1,067667841 | 1,142819143 | 1,111790125 | 0,9839931 | 0,990486375 | 0,808119453 | 1,147630918 | 1,07240334 | 0,950401572 | 1,08720175 | 0,986865068 |
| 0,954165613 | 1,010408929 | 0,95248368 | 0,914036757 | 1,360853339 | 0,939810194 | 0,82629378 | 1,180976687 | 0,969078335 | 1,023857414 | 1,096938095 |
| 0,938719242 | 0,755582094 | 1,232725539 | 0,974572996 | 1,13754664 | 0,556858048 | 0,970806042 | 1,012934621 | 1,038561293 | 1,359068546 | 1,0215221 |
| 0,966056992 | 0,79750487 | 0,805272496 | 0,95819623 | 0,901102259 | 0,928225548 | 0,862076919 | 1,040696842 | 0,907634442 | 0,934841786 | 1,134407058 |
| 1,059504942 | 1,04144079 | 1,007749889 | 1,013784271 | 0,978239122 | 0,870678058 | 1,114802495 | 1,083470601 | 0,805547636 | 0,852227254 | 1,012905704 |
| 0,995843934 | 0,963002227 | 1,211403073 | 0,983085582 | 1,064259559 | 0,798710475 | 0,983491198 | 0,898328555 | 1,006107665 | 0,920754567 | 0,999376526 |
| 0,957423517 | 1,261185007 | 0,958211153 | 0,957106138 | 0,70954048 | 0,877429377 | 1,492749821 | 1,226763997 | 0,900631192 | 1,152904876 | 0,925821638 |
| 1,002288526 | 0,853580314 | 0,677575895 | 0,980781739 | 1,034168628 | 1,98927273 | 0,82836314 | 1,026830808 | 1,19411814 | 1,036337537 | 0,777725833 |
| 0,954133456 | 1,247558459 | 0,876179937 | 1,006795231 | 1,000973438 | 1,231129041 | 0,924115175 | 1,091773311 | 1,009398444 | 1,364150661 | 0,962196962 |
| 1,052543359 | 0,981464204 | 1,370088633 | 0,94895571 | 1,150889796 | 1,535115584 | 1,164225847 | 1,175353185 | 0,92811749 | 1,173717395 | 0,974234248 |
| 0,903655851 | 1,302509131 | 0,859421749 | 0,945713313 | 0,797275325 | 1,384146743 | 0,835483473 | 1,017524779 | 1,21955427 | 1,055037163 | 0,901517753 |
| 0,967541244 | 1,095456044 | 1,097375448 | 0,940743835 | 1,423881331 | 0,893387601 | 1 | 0,977888114 | 1,008598394 | 1,022155264 | 0,998915299 |
| 0,90141113 | 1,366950633 | 1,788587117 | 0,876742689 | 1,159226958 | 0,622912602 | 1,557932847 | 0,904098381 | 0,881008494 | 1,074386877 | 0,987007988 |
| 0,886185341 | 0,840436618 | 0,665484153 | 1,04589034 | 1,061192462 | 1,585358273 | 1,043955287 | 1,071097656 | 0,874384376 | 0,931888388 | 1,080535863 |
| 1,183872221 | 1,040942093 | 1,554702125 | 0,946109101 | 1,144815303 | 0,784516802 | 1,155937307 | 1,049671766 | 0,727778839 | 0,973458081 | 1,106853163 |
| 0,647322797 | 0,84751295 | 0,809757781 | 0,799420333 | 0,557675987 | 1,850275641 | 1,036500161 | 1,050319517 | 0,685915834 | 0,918428833 | 1,123888467 |
| 0,96118929 | 1,007587289 | 1,32437168 | 0,960691493 | 1,322298011 | 0,776419454 | 0,986191527 | 1,178575567 | 0,975283997 | 1,078187509 | 1,010337627 |
| 0,947374952 | 0,968348853 | 1,054096575 | 0,94938241 | 0,944110901 | 1,125254471 | 0,892711659 | 0,939067779 | 1,098573084 | 0,827778051 | 1,023730441 |
| 0,999978627 | 1,775840653 | 0,557664773 | 1,142153027 | 0,602214575 | 1,026728239 | 1,048016453 | 0,928902599 | 0,814247378 | 1,00724012 | 0,917856046 |
| 0,943185006 | 2,00300738 | 0,619963882 | 1,049725145 | 0,843976675 | 1,305874705 | 1,010253937 | 0,913703925 | 0,736708794 | 1,188324236 | 0,919718834 |
| 0,837474262 | 1,442790049 | 0,854735613 | 0,808369267 | 0,781304954 | 2,242135912 | 1,072428007 | 1,109107894 | 0,663609414 | 1,259740658 | 0,799734069 |

| LC3A-B | Lck | LDHA | LRP6_pS1490 | MAPK_pT202_Y204 | Mcl-1 | MCT4 | MDM2_pS166 | MEK1 | MEK1_pS217_S221 | MERIT40_pS29 |
|---|---|---|---|---|---|---|---|---|---|---|
| 1,016596909 | 0,885767516 | 0,869384814 | 1,048477911 | 0,83850575 | 1,151420529 | 0,82126549 | 1,108763364 | 0,869384814 | 1,044813711 | 1,304845059 |
| 1,005151876 | 0,902048348 | 0,915806141 | 1,184701776 | 0,979580984 | 1,16249644 | 0,819572802 | 1,198002682 | 0,915806141 | 1,034264492 | 0,896479757 |
| 1,042034595 | 1,151972966 | 1,131397804 | 0,915404526 | 1,216246678 | 0,851800991 | 1,032275686 | 1,147323453 | 1,131397804 | 0,989569788 | 0,742595883 |
| 0,966847506 | 1,005554761 | 1,183189393 | 1,00374454 | 0,880557902 | 1,033932272 | 1,250478339 | 1,058672487 | 1,183189393 | 0,887381809 | 0,954026931 |
| 1,027405586 | 0,839653052 | 0,848996338 | 1,112814821 | 0,955480323 | 1,145188583 | 0,905586643 | 1,232204826 | 0,862211 | 1,00446864 | 1,198959903 |
| 1,056348237 | 1,112785576 | 1,135767418 | 0,875504298 | 0,821955909 | 1,091971785 | 0,726444898 | 0,969303943 | 1,096790164 | 0,969348781 | 1,053487122 |
| 1,081734361 | 0,786156445 | 1,086739658 | 0,945014959 | 0,915902103 | 0,924418962 | 1,820403696 | 1,170418073 | 1,043445781 | 1,02749216 | 1,266963878 |
| 1,463200215 | 0,900908735 | 1,334799989 | 1,185050057 | 1,121479187 | 1,28081049 | 0,729754806 | 1,280730006 | 1,334799989 | 0,943429961 | 0,828436327 |
| 0,974136547 | 0,738884506 | 0,890984391 | 1,126405627 | 0,99516408 | 1,037871882 | 1,333516685 | 1,14382713 | 0,890984391 | 0,994462257 | 1,168420846 |
| 0,80752977 | 0,907238272 | 0,785632569 | 1,188061876 | 0,817520837 | 1,058113097 | 0,888123118 | 1,275200072 | 0,785632569 | 1,069930987 | 1,293506319 |
| 1,030438573 | 1,028336138 | 1,057420285 | 0,915345123 | 0,710592619 | 1,090110597 | 1,248438406 | 1,020564627 | 1,057420285 | 0,933839677 | 1,091115161 |
| 1,054678755 | 0,774833272 | 0,983226513 | 1,259468263 | 0,77281965 | 0,988400034 | 1,310090949 | 0,967594158 | 0,983226513 | 1,013363605 | 0,95744768 |
| 0,899527221 | 1,098748589 | 1,014984308 | 1,003209526 | 1,052165366 | 0,862560416 | 1,219696529 | 1,536644662 | 0,807683589 | 0,976462539 | 0,951256664 |
| 1,144694392 | 1,885135421 | 1,129335468 | 0,917461689 | 1,103159538 | 1,101256611 | 0,653760379 | 1,11343885 | 1,129335468 | 0,859267552 | 0,997598155 |
| 0,886423506 | 2,199748293 | 0,898645048 | 0,883457115 | 0,966349494 | 1,256514998 | 0,948383889 | 1,349374821 | 0,898645048 | 1,006786467 | 0,887397156 |
| 0,869098332 | 1,159105238 | 1,175731916 | 1,031610618 | 0,962005452 | 0,900458485 | 1,165852724 | 1,21881748 | 1,027307626 | 1,000894361 | 0,980712202 |
| 0,963081945 | 2,103376533 | 1,01918828 | 0,972094316 | 0,894959895 | 1,361053413 | 0,862770718 | 1,261251753 | 1,13269813 | 0,924878699 | 1,238479606 |
| 0,865646656 | 0,940189671 | 0,962329745 | 1,278523212 | 0,914833746 | 1,058402092 | 0,771056497 | 1,440844198 | 1,089089241 | 1,006411906 | 0,858158343 |
| 0,798177965 | 0,819176658 | 1,147273455 | 1,133317016 | 0,892235972 | 0,833849803 | 1,013930337 | 1,243769725 | 0,952476186 | 0,992651857 | 1,224457287 |
| 1,023229574 | 1,422399248 | 1,374071717 | 1,180736139 | 1,386290818 | 1,151854266 | 1,113049522 | 1,206570261 | 1,374071717 | 0,906465244 | 0,822086188 |
| 0,839048184 | 0,774319328 | 1,263434755 | 1,022540642 | 1,044477473 | 0,841086968 | 1,018098587 | 1,100169927 | 1,263434755 | 0,894184307 | 0,856249723 |
| 0,845222597 | 1,081525886 | 1,975982845 | 1,017053746 | 0,928588083 | 0,489001617 | 0,664428242 | 1,64166588 | 1,975982845 | 0,847015803 | 1,131124506 |
| 0,848287935 | 1,391947118 | 0,986457081 | 1,009421501 | 1,091100996 | 1,126334386 | 1,224691436 | 1,286112066 | 0,986457081 | 1,127873873 | 1,001764457 |
| 0,9625595 | 0,963506501 | 0,949246349 | 1,229314228 | 1,040483671 | 1,359925997 | 0,859221623 | 1,223987627 | 0,949246349 | 1,053270679 | 0,961127593 |
| 1,019820933 | 0,738949496 | 2,308844355 | 0,904300636 | 1,126056858 | 0,799888882 | 1,417064891 | 1,037066888 | 1,616932156 | 0,935498267 | 2,183517967 |
| 1,333954614 | 0,809330251 | 2,358333435 | 1,043090689 | 1,169290259 | 0,818378624 | 1,161325827 | 0,970583001 | 1,387126502 | 0,824946081 | 2,046556959 |
| 1,045217648 | 0,885887247 | 2,172345353 | 0,971344361 | 1 | 0,766228873 | 1,194233298 | 1,861534973 | 2,172345353 | 0,944579122 | 1,805303095 |

| Merlin | MIF | MMP2 | Mnk1 | MSH6 | MSI2 | mTOR | mTOR_pS2448 | Myosin-Heavy-Chain-11 | Myosin-IIa_pS1943 | Myt1 |
|---|---|---|---|---|---|---|---|---|---|---|
| 1,03751251 | 1,149713758 | 0,973275109 | 1,128837875 | 0,982119569 | 0,693055984 | 0,861570616 | 0,821734848 | 0,403735669 | 0,869384814 | 1,234192718 |
| 0,995490948 | 1,113588887 | 0,965066948 | 0,911186994 | 0,935187317 | 1,070011893 | 1,078506472 | 1,128140308 | 0,730576205 | 0,915806141 | 1,068370079 |
| 0,648548317 | 1,072290329 | 1,122300606 | 1,122813984 | 1,137537326 | 1,578551081 | 0,590048366 | 0,71980694 | 0,96472008 | 1,131397804 | 0,854119208 |
| 0,756151871 | 1,090551006 | 0,981613247 | 0,923406009 | 0,763634361 | 0,664184651 | 0,882477372 | 1,053144636 | 0,562265312 | 1,183189393 | 0,980670075 |
| 1,010056076 | 1,101487302 | 0,984294037 | 0,927597154 | 0,880109369 | 0,760307666 | 1,200339515 | 1,23571096 | 0,777282889 | 0,787643965 | 1,080601003 |
| 1,095315426 | 1,13719021 | 1,060617971 | 0,834105223 | 0,935286045 | 1,166767292 | 1,045391202 | 0,55068598 | 0,849224258 | 1 | 0,982141244 |
| 1,024839771 | 1,260168452 | 1,034600162 | 0,794347893 | 1,014484928 | 0,933539324 | 0,919161781 | 0,895221291 | 0,596763241 | 0,92930878 | 1,055481091 |
| 0,838804167 | 0,990723614 | 0,951788203 | 0,915705594 | 0,856345741 | 0,947212535 | 0,919727337 | 1,168166678 | 2,061934568 | 1,334799989 | 0,99005854 |
| 0,91433929 | 1,173729332 | 1,006507122 | 1,10545649 | 0,717910516 | 0,757392655 | 1,292729048 | 1,233071109 | 0,627770097 | 0,890984391 | 0,982947778 |
| 0,985547263 | 1,157995568 | 1,002287254 | 1,008128755 | 0,98186677 | 0,953592911 | 1,06019205 | 1,241291848 | 0,750225754 | 0,785632569 | 1,055727339 |
| 0,991616213 | 0,952495523 | 1,040752837 | 0,868726738 | 0,70049341 | 0,908338862 | 0,91703514 | 1,096206965 | 0,779137564 | 1,057420285 | 1,058619781 |
| 1,053807078 | 1,120425576 | 0,991132315 | 1,059341681 | 0,715004147 | 0,827346964 | 1,21683839 | 1,492230125 | 0,677870868 | 0,983226513 | 0,934189644 |
| 1,030283376 | 1,176565544 | 1,086355333 | 0,99356685 | 3,924871729 | 0,947910306 | 1,187289289 | 1,316581941 | 293,953477 | 0,807683589 | 0,958901751 |
| 0,756216424 | 0,962107244 | 0,984056791 | 1,537193431 | 0,935614489 | 1,675197774 | 1,139486153 | 1,206208147 | 44,2788609 | 1,129335468 | 0,833273967 |
| 1,00019248 | 1,188772743 | 1,022399616 | 1,366860214 | 2,585263415 | 1,094647926 | 1,031688935 | 1,103638782 | 0,91046629 | 0,898645048 | 0,873332338 |
| 0,973019207 | 1,062750423 | 0,928861834 | 1,040296935 | 2,69226294 | 1,459380535 | 1,032375478 | 1,134423417 | 1,541905634 | 0,943653081 | 0,942582638 |
| 1,045170696 | 1,022031413 | 0,896868186 | 1,337127623 | 0,967247994 | 1,263269725 | 1,146580675 | 1,209068259 | 1,400705465 | 0,792985253 | 1,082044535 |
| 1,060305964 | 1,13570824 | 0,840383812 | 0,829789602 | 0,931275709 | 0,97184603 | 1,183455308 | 1,300026684 | 0,897814869 | 0,92304201 | 0,975240498 |
| 0,998526719 | 1,038380144 | 0,92929718 | 0,979547986 | 1,05016363 | 0,743034861 | 1,373822306 | 1,225270929 | 0,736000694 | 0,768032445 | 1,071773289 |
| 0,692752814 | 1,051117456 | 1,067968165 | 1,122117228 | 1,434054949 | 1,322960741 | 0,898492659 | 1,135960792 | 2,259755987 | 1,374071717 | 0,840245285 |
| 0,787547699 | 1,110822385 | 0,971678011 | 0,869478482 | 1,096220722 | 0,701910056 | 0,776379959 | 0,831855449 | 0,568761838 | 1,263434755 | 0,843992393 |
| 0,8487584 | 1,125991923 | 1,29622275 | 0,817091091 | 1,267698912 | 1,834875694 | 0,551403819 | 1,112234555 | 48,93540834 | 1,975982845 | 0,973308101 |
| 1,259191414 | 0,889794665 | 1,159860933 | 1,222103834 | 0,74393369 | 0,840389308 | 1,184350194 | 1,482776579 | 0,893805317 | 0,986457081 | 1,058141566 |
| 0,92436426 | 0,998837708 | 1,064737325 | 1,092076723 | 0,763446037 | 0,944925041 | 1,028471065 | 1,284120262 | 3,434426202 | 0,949246349 | 1,016528998 |
| 0,793812301 | 0,90031492 | 1,310251797 | 1,096512735 | 1,321978612 | 0,959693404 | 0,628429658 | 0,596420497 | 53,01071797 | 1,091009894 | 1,265529892 |
| 0,775251859 | 0,93444739 | 0,757793186 | 1,09712442 | 1,075639824 | 1,290874647 | 0,682866547 | 0,63399926 | 95,82640676 | 1,014495827 | 1,193220269 |
| 0,765132672 | 0,912884651 | 1,042414284 | 1,112747295 | 1,393675986 | 2,187407395 | 0,702759816 | 0,802231146 | 16,46188768 | 2,172345353 | 1,242895045 |

| N-Cadherin | NAPSIN-A | NDRG1_pT346 | NF-kB-p65_pS536 | Notch1 | Notch3 | Oct-4 | P-Cadherin | Met_pY1234_Y1235 | p16INK4a | p21 |
|---|---|---|---|---|---|---|---|---|---|---|
| 0,907799347 | 0,973576468 | 0,79509645 | 0,948961674 | 1 | 1,017497131 | 1,024863341 | 0,942904386 | 0,86306001 | 0,961996733 | 1,017507133 |
| 0,973027481 | 1,007002379 | 1,037041584 | 1,146008476 | 1,046111106 | 1,095624577 | 0,969151423 | 0,925533572 | 1,259823078 | 0,889588549 | 0,839609133 |
| 1,128563764 | 1,102349088 | 1,853542962 | 1,088130609 | 0,675142002 | 0,930640199 | 1,165805125 | 0,972959818 | 0,745570797 | 0,839365216 | 1,565781587 |
| 1,067989265 | 1,094834181 | 1,276789683 | 0,971865969 | 0,782353788 | 0,943025734 | 1,006261273 | 0,826551652 | 2,178987282 | 1,022471585 | 1,064812813 |
| 1,011702018 | 1,103925844 | 1,965841759 | 1,120874005 | 1,1067517 | 1,010170594 | 0,919543957 | 1,018181294 | 1,119070998 | 0,858782276 | 0,928399324 |
| 1,059056908 | 0,963235048 | 0,223060666 | 0,769441403 | 1,182131487 | 1,143240774 | 0,946515225 | 1,363329143 | 0,651302121 | 1,219062202 | 0,718953697 |
| 1,033021531 | 0,956636148 | 0,437450177 | 0,953803798 | 1,055690299 | 1,062365413 | 1,091402215 | 4,906684496 | 0,81571637 | 1,161619179 | 0,904392176 |
| 0,829681267 | 1,046023833 | 1,016594839 | 0,890395453 | 1,064756082 | 1,079863962 | 0,982687357 | 0,830893411 | 1,200115414 | 1,751598668 | 0,778110096 |
| 1,000464373 | 0,980514317 | 0,858018913 | 1,191543113 | 1,288407824 | 1,003072351 | 1,012298263 | 1,048919823 | 1,058455889 | 1,045831542 | 1,003089591 |
| 0,909051369 | 1,221088642 | 0,813586932 | 1,012389533 | 1,105182806 | 1,081472695 | 0,900841132 | 0,930706282 | 0,970877698 | 0,834594324 | 1,119815175 |
| 0,773120387 | 1,038635719 | 1,07113835 | 0,679689717 | 0,95690224 | 0,996721122 | 0,892515956 | 0,98260267 | 1,341926824 | 0,707425305 | 0,970614531 |
| 0,95579699 | 1,013583929 | 4,263460592 | 1,162643647 | 1,519635807 | 1,028080198 | 1,065304767 | 0,828990683 | 1,309547091 | 0,990388862 | 1,094576667 |
| 1,064712874 | 0,832386951 | 3,092039389 | 1,176192395 | 1,419396838 | 1,012187557 | 1,154055967 | 0,984336702 | 0,58853928 | 1,223978673 | 1,103101172 |
| 1,015736434 | 0,988981791 | 1,824356781 | 0,95377778 | 0,678513889 | 0,859364357 | 0,870258076 | 1,117339989 | 0,716201694 | 1,003427054 | 1,082366171 |
| 1,125206266 | 0,940852936 | 0,806455932 | 1,0359667 | 1,021818085 | 0,954354645 | 0,949240901 | 1,009225813 | 0,714329755 | 1,301210833 | 1,031447067 |
| 1,104480377 | 0,978191195 | 0,992806635 | 1,016962123 | 0,974135976 | 1,038970594 | 0,997738332 | 0,997932073 | 1,02851646 | 0,955780344 | 1,065241189 |
| 0,955668465 | 0,885252719 | 1,239907315 | 1,003167472 | 0,821784747 | 0,968734446 | 0,848813703 | 0,987063282 | 1 | 1,010181649 | 1,002102525 |
| 1,021315252 | 0,974631284 | 0,911067254 | 1,13074306 | 1,352469788 | 1,066106916 | 0,938624442 | 1,002691509 | 2,169154008 | 1,286469942 | 0,915226086 |
| 0,937030539 | 0,873530636 | 2,907642321 | 1,045596523 | 1,149661115 | 1,064479338 | 0,915168472 | 0,924959992 | 1,565514032 | 0,96122835 | 0,983720813 |
| 0,89904873 | 0,936569616 | 1,233875876 | 1,066275166 | 0,776236804 | 0,881457611 | 0,989948672 | 0,958808888 | 1,384068135 | 0,901430696 | 1,201051312 |
| 1,058104217 | 0,916716498 | 1,700215217 | 0,958151361 | 0,717088013 | 1,071836673 | 1,242105947 | 0,854592288 | 1,791343887 | 1,032509594 | 1,165574036 |
| 1,067797812 | 1,112082481 | 0,738916296 | 0,863158239 | 0,927703565 | 0,998451823 | 0,824400179 | 1,06122827 | 0,677758136 | 0,651896838 | 1,319589666 |
| 0,903858192 | 1,00398102 | 2,48537846 | 1,03821956 | 0,954591485 | 0,941456022 | 0,840536698 | 0,811435509 | 1,659748136 | 1,428397803 | 0,994809977 |
| 0,951624118 | 0,987503795 | 2,865911183 | 1,402400334 | 1,049430406 | 1,007686779 | 0,970289301 | 0,760481005 | 2,391643494 | 0,854931408 | 1,031506238 |
| 1,071768342 | 0,939051762 | 0,597728308 | 0,953890966 | 0,870326248 | 0,713957167 | 0,909711289 | 1,369907476 | 0,515514337 | 0,925537199 | 1,242135556 |
| 1,172725615 | 0,975041807 | 0,943079948 | 0,987755422 | 0,926425268 | 0,701990652 | 0,820700543 | 1,372288938 | 0,587825193 | 1,007105696 | 1,325567322 |
| 1,140698638 | 1,035748683 | 0,871193254 | 0,929858271 | 0,679714688 | 0,793507415 | 0,976925792 | 1,377748187 | 0,595376807 | 0,766191185 | 1,345371701 |

| p27-Kip-1 | p27_pT198 | p38-MAPK | p38_pT180_Y 182 | p44-42-MAPK | p53 | p70-S6K1 | p70-S6K_pT389 | p90RSK_pT57 3 | PAICS | PAK1 |
|---|---|---|---|---|---|---|---|---|---|---|
| 1,190345081 | 1,002265522 | 1,077699534 | 0,833320185 | 1,292901327 | 0,865302841 | 1,197179028 | 0,934396381 | 0,772600576 | 0,952102266 | 1,234695402 |
| 1,091079005 | 0,985696935 | 0,888347247 | 0,832207757 | 0,973948692 | 0,928490451 | 1,26391758 | 1,004805496 | 1,189734178 | 0,886093897 | 1,230010481 |
| 1,158365499 | 1,04261697 | 1,140975333 | 1,201438516 | 1,034464335 | 0,943192027 | 0,772760315 | 0,99225317 | 1,250976373 | 0,643075627 | 0,410232093 |
| 1,053079568 | 0,815422732 | 1,124187285 | 1,306894909 | 1,079922135 | 1,019157169 | 1,055113804 | 0,938158931 | 1,153141893 | 1,167255232 | 0,893610747 |
| 1,033752677 | 1,054208295 | 0,765418108 | 1,235068809 | 0,871681366 | 0,980506143 | 1,142275115 | 0,866083708 | 1,189787695 | 0,839677295 | 1,402834747 |
| 0,977798364 | 0,999297093 | 1,160184448 | 0,622388247 | 1,081683503 | 0,996776382 | 0,907793265 | 0,845216009 | 0,656453291 | 1,060063034 | 1,080809485 |
| 1,075803191 | 0,951816539 | 0,944173265 | 0,659301593 | 1,025279428 | 1,004180217 | 1,217480884 | 0,830452832 | 0,727665148 | 1,019358437 | 1,009798919 |
| 0,964672675 | 0,863882333 | 1,323500617 | 1,62588548 | 1,045439333 | 0,873363501 | 1,385971798 | 0,850441212 | 0,952185465 | 0,773752029 | 0,979383042 |
| 1,034721643 | 0,925828571 | 0,905065204 | 0,876004019 | 0,987781236 | 0,945561891 | 1,599210891 | 0,952486271 | 0,862107253 | 0,942569872 | 1,305920003 |
| 0,993543645 | 1,089405585 | 0,901545114 | 0,91165953 | 0,775354686 | 1,029107524 | 1,105469577 | 0,857640748 | 1,041399905 | 1,231711414 | 0,91963244 |
| 0,98053013 | 0,945363281 | 0,954985921 | 0,734018303 | 1,030084202 | 1,054366375 | 1,059691858 | 0,876009136 | 0,810063586 | 1,446722616 | 1,068952783 |
| 0,997545257 | 0,994949834 | 0,918742506 | 0,89838299 | 1,044275794 | 1,068647162 | 0,881361461 | 0,971984724 | 1,655171137 | 0,996467749 | 0,968477876 |
| 1,209692285 | 1,001808189 | 1,0205517 | 1,394657467 | 0,868331536 | 0,955915983 | 0,981324799 | 1,190778117 | 1,915563815 | 0,83200736 | 1 |
| 1,097568087 | 0,875278987 | 1,535768128 | 1,379069371 | 1,214359835 | 0,984495828 | 1,041222959 | 1,131302756 | 1,301401162 | 0,831883895 | 0,904739487 |
| 0,962570393 | 0,987598496 | 1,431768424 | 1,358779443 | 1,04563897 | 0,901240404 | 1,013868091 | 1,061192529 | 0,887868979 | 0,846051552 | 1,102833857 |
| 1,113005285 | 0,944316381 | 1,066999107 | 1,210224245 | 1,032252817 | 1,034859911 | 0,957240643 | 0,98275118 | 0,832522467 | 1,244192116 | 1,035029733 |
| 1,038465501 | 0,942059389 | 1,273524052 | 1,216055752 | 0,906916481 | 1,05340187 | 1,324021585 | 1,199228823 | 1,038875808 | 1,19282651 | 0,833116892 |
| 1,094562739 | 0,991611446 | 0,998069986 | 1,096236719 | 0,913355671 | 1,024469336 | 1,536117087 | 0,921434083 | 1,140329788 | 1,279773485 | 1,036151865 |
| 1,097141178 | 0,977933568 | 0,998056545 | 0,822735812 | 0,881888125 | 0,91308238 | 0,965690224 | 0,96678597 | 0,97837733 | 1,391149104 | 0,882311698 |
| 0,940378634 | 0,790123076 | 1,206430882 | 1,397174372 | 0,999973953 | 0,910973263 | 0,957080164 | 0,955176258 | 1,387492697 | 0,996390458 | 0,80930932 |
| 1,027761835 | 0,817573969 | 0,762090541 | 0,784691504 | 0,983714136 | 0,944371054 | 1,159850771 | 0,916701051 | 0,934334455 | 1,134615185 | 1,379674925 |
| 0,787135488 | 1,050038394 | 0,670720232 | 1,532348151 | 0,375274943 | 1,040188475 | 0,516417739 | 0,941450088 | 1,956682813 | 0,838780161 | 0,437790981 |
| 0,972091291 | 0,99729365 | 1,258597383 | 1,820232192 | 1,126191685 | 0,891111148 | 1,022955741 | 1,022639112 | 1,991901058 | 1,395983007 | 0,975698527 |
| 1,00102756 | 0,962585282 | 0,968945351 | 1,100929859 | 0,959207674 | 1,040963917 | 0,947207158 | 1,024858901 | 1,440027058 | 1 | 0,920104829 |
| 0,902034538 | 1,043460912 | 0,513241217 | 0,971732513 | 0,528610894 | 1,4050874 | 0,640694934 | 1,139614728 | 1,473814651 | 0,604805199 | 0,486974371 |
| 0,934053617 | 0,826585289 | 0,403423996 | 0,855718787 | 0,607342885 | 1,364772155 | 0,660951351 | 1,324396067 | 1,359679186 | 0,577996697 | 0,588945995 |
| 0,824504616 | 0,945871767 | 0,953449985 | 1,230276979 | 0,504269947 | 1,161792538 | 0,61005565 | 1,122439535 | 1,879954794 | 0,922133605 | 0,405099397 |

| PAK4 | PAR | PARP | Paxillin | PD-L1 | Pdcd4 | PDGFR-b | PDHK1 | PDK1 | PDK1_pS241 | PEA-15 |
|---|---|---|---|---|---|---|---|---|---|---|
| 1,008511218 | 0,949483072 | 1,244225846 | 1,169322915 | 0,849401633 | 0,937328255 | 0,726457288 | 1,045458606 | 0,761037834 | 0,94075097 | 1,210846967 |
| 0,965789423 | 3,706514017 | 1,374407367 | 0,67769123 | 0,882143754 | 0,8848502 | 1,571059101 | 0,978248861 | 0,813346235 | 0,952677593 | 0,891847939 |
| 0,872571877 | 0,683035101 | 0,724717472 | 0,825573635 | 1,133927128 | 0,499522605 | 1,698652918 | 1,076629628 | 0,75191024 | 0,880247851 | 1,028494283 |
| 0,855261567 | 0,722774096 | 0,976044021 | 0,928864475 | 1,036703915 | 0,83165709 | 1,039117014 | 0,941280738 | 0,833256553 | 1,082257325 | 1,002283787 |
| 0,918024983 | 1,823386809 | 0,982309064 | 0,870602076 | 0,878147994 | 1,282186647 | 9,265792379 | 0,914575581 | 0,878234783 | 1,148775173 | 0,97411906 |
| 1,003082514 | 0,462575301 | 0,938507377 | 0,96009292 | 0,889691798 | 0,441509258 | 1,090747532 | 0,987548937 | 1,037747758 | 1,410806151 | 1,317277609 |
| 1,032653767 | 2,931661544 | 1,490635081 | 1,033473762 | 0,889035155 | 0,6553009 | 0,777672854 | 1,049398224 | 0,715731614 | 0,838459122 | 1,18874488 |
| 0,918892076 | 0,402605991 | 1,025633411 | 1,156116863 | 0,824165259 | 0,589326553 | 1,183812517 | 0,913055935 | 0,863728683 | 1,218710643 | 1,039877386 |
| 0,970840662 | 3,318713526 | 1,326271773 | 0,813304419 | 0,873319913 | 1,130988863 | 0,776550391 | 0,987164259 | 0,726843983 | 0,909152499 | 1,134701079 |
| 0,896802698 | 2,002969958 | 1,331369947 | 0,911486107 | 0,928547528 | 1,46553439 | 0,760814406 | 1,047477766 | 0,894606623 | 1,144621504 | 0,942978677 |
| 1,19356846 | 0,678830143 | 0,993011017 | 1,070110221 | 5,497039886 | 0,971335224 | 1,891477383 | 1,009613243 | 1,506797707 | 0,945984942 | 1,368007614 |
| 0,936419736 | 1,188740872 | 0,97743926 | 1,020483641 | 1,485707452 | 1,998147614 | 1,250282762 | 1,024469428 | 1 | 1,319576263 | 1,043764104 |
| 0,938886336 | 0,733269995 | 0,851663008 | 1,347637862 | 0,805877952 | 1,469133625 | 0,569399542 | 0,900843843 | 0,873838927 | 1,177096535 | 0,804922304 |
| 0,926435698 | 0,697254454 | 1,07448589 | 1,253805537 | 0,94029811 | 1,002909881 | 2,804433625 | 0,95585206 | 0,786350315 | 1,148269546 | 1,080301195 |
| 0,895874962 | 1,199803678 | 0,996182448 | 1,011268282 | 0,910016096 | 0,905061992 | 1,050272069 | 1,028253536 | 0,88331449 | 1,016984973 | 1,227421316 |
| 0,855931936 | 0,854606627 | 0,898078348 | 0,997702542 | 0,909184654 | 0,931917055 | 1,113767378 | 0,945923224 | 0,924392227 | 1,003565286 | 1,182178205 |
| 0,907141493 | 0,942113939 | 1,563988453 | 1,066695703 | 0,850153065 | 2,177939753 | 1,109370868 | 0,868736571 | 1,005211942 | 1,322229655 | 1,004118273 |
| 0,904157936 | 2,387238624 | 1,465535927 | 0,927076308 | 0,876205094 | 1,242974239 | 0,852018658 | 0,954229723 | 0,824068395 | 1,241440016 | 0,926197678 |
| 0,89265906 | 3,390547926 | 1,095006779 | 1,288317282 | 0,813156942 | 1,461665166 | 1,073442236 | 0,850825393 | 0,973987699 | 1,209371573 | 0,987884453 |
| 1,269614636 | 1,515394732 | 1,140995825 | 1,036182711 | 4,819660264 | 0,961681112 | 1,435912774 | 0,953598642 | 2,371226971 | 1,077495143 | 1,085285739 |
| 0,925669651 | 0,94576349 | 1,000258366 | 0,89599018 | 1,01006563 | 0,931638106 | 1,798063449 | 1,105165535 | 0,742629906 | 0,908208619 | 1,033227849 |
| 1,998670773 | 0,723439494 | 1,123570998 | 0,881972322 | 22,94153439 | 1,961872969 | 1,494478198 | 1,294514724 | 4,395941425 | 0,553196977 | 0,671469645 |
| 1,00898509 | 0,57528098 | 1,197863274 | 1,180114378 | 3,350507837 | 2,430732349 | 0,877941429 | 0,906311295 | 2,157889991 | 1,045868521 | 1,177264096 |
| 1,030417439 | 2,810951451 | 1,282743957 | 0,957952642 | 3,515948827 | 2,615564077 | 0,979233078 | 0,955019558 | 1,464811125 | 0,933331587 | 1,148629839 |
| 2,033828789 | 1,204290678 | 1,097431558 | 0,739853675 | 38,30942767 | 1,436991395 | 0,953770616 | 1,089825531 | 11,73716828 | 0,537868225 | 0,823366673 |
| 0,993005284 | 1,12222345 | 1,094565514 | 1,010061816 | 0,958722394 | 1,366271846 | 0,975952474 | 1,015597129 | 0,708394225 | 0,752944389 | 0,649386914 |
| 1,233962886 | 1,367253434 | 1,235225418 | 0,996057153 | 4,447379407 | 1,032068823 | 1,162460878 | 1,287499768 | 8,386683086 | 0,837620716 | 0,676131151 |

| PEA-15_pS116 | PI3K-p110-a | PI3K-p85 | PKA-a | PKC-b-II_pS660 | PKC-delta_pS664 | PKCa | PKM2 | gamma2_pY759 | PLK1 | PMS2 |
|---|---|---|---|---|---|---|---|---|---|---|
| 0,948732575 | 1,136447061 | 1,068873588 | 0,984248802 | 1,013517202 | 1,063355977 | 0,98775855 | 1,193374647 | 1,081900748 | 0,869384814 | 0,869384814 |
| 0,961183367 | 1,090306639 | 1,036097026 | 0,918375431 | 0,815233416 | 1,030515824 | 1,022078288 | 1,102529476 | 1,170640309 | 0,915806141 | 0,915806141 |
| 1,27477249 | 0,784813083 | 0,707399062 | 0,862231752 | 1,110485933 | 1,207939721 | 0,715698568 | 0,92515778 | 1,060132674 | 1,131397804 | 1,131397804 |
| 1,064515115 | 1,000462077 | 0,905676997 | 0,904312498 | 1,185794743 | 1,04823176 | 1,443737719 | 1,195651936 | 1,090700509 | 1,183189393 | 1,183189393 |
| 0,877641559 | 1,033741067 | 1,080331611 | 0,865328649 | 1,095334696 | 0,941791561 | 1,385700183 | 1,402931234 | 1,185238822 | 0,787643965 | 0,787643965 |
| 0,891854202 | 1,09542718 | 1,331209385 | 0,979534896 | 1,219888027 | 0,998041584 | 1,395086076 | 0,228054156 | 0,91714639 | 1 | 1 |
| 0,92940006 | 0,965052832 | 1,04944021 | 0,942483225 | 0,825534071 | 1,035460002 | 0,608122898 | 1,559513536 | 0,914799783 | 0,92930878 | 0,92930878 |
| 1,056727154 | 1,107038899 | 0,83029428 | 1,299646188 | 1,236174394 | 0,945949122 | 1,016145267 | 1,035191931 | 1,160381632 | 1,334799989 | 1,334799989 |
| 0,924034901 | 1,107459347 | 0,960699453 | 1,145378556 | 0,889022468 | 1,065807106 | 1,58614248 | 1,121955917 | 1,100741584 | 0,890984391 | 0,890984391 |
| 0,910405201 | 1,130310446 | 0,906655968 | 0,936367184 | 1,008393601 | 1,096167287 | 1 | 1,139811952 | 1,163707978 | 0,785632569 | 0,785632569 |
| 0,950864352 | 0,976032905 | 0,92618886 | 1,033007469 | 0,945210782 | 0,997056899 | 1,383412519 | 1,440437931 | 1,084432808 | 1,057420285 | 1,057420285 |
| 1,060975932 | 1,052025935 | 1,008984003 | 1,030006973 | 1,129662365 | 1,041340505 | 0,888710717 | 1,228223885 | 0,982560179 | 0,983226513 | 0,983226513 |
| 1,186423503 | 1,092847668 | 1,567419989 | 1,408155097 | 1,776066326 | 1,008680417 | 1,293045492 | 0,753240265 | 1,1276144 | 0,807683589 | 0,820931135 |
| 1,129224541 | 1 | 1,220700965 | 1,191270362 | 1,61359843 | 0,905187627 | 1,193051106 | 0,67807188 | 1,109716344 | 1,129335468 | 1,129335468 |
| 1,072966852 | 1,091055852 | 1,284467032 | 1,076337725 | 1,954480343 | 1,044439087 | 0,814551801 | 1,231315148 | 1,274513759 | 0,898645048 | 0,898645048 |
| 0,910173104 | 0,950635438 | 1,071544674 | 0,941005666 | 1,224078624 | 0,945501146 | 1,438554048 | 1,302665286 | 0,960068199 | 0,943653081 | 0,943653081 |
| 1,136237876 | 1,041927822 | 1,389362744 | 1,403938149 | 2,045459246 | 0,958822647 | 1,542692895 | 1,01054689 | 1,350011777 | 0,792985253 | 0,821716878 |
| 0,912695163 | 1,191524362 | 1,064746816 | 1,104042884 | 1,119780766 | 0,999797734 | 1,600846767 | 0,918639755 | 1,047353067 | 0,92304201 | 0,92304201 |
| 0,84038619 | 1,062573086 | 0,971369359 | 1,311438675 | 1,09675763 | 0,945443431 | 1,318065601 | 1,322462047 | 0,969584274 | 0,768032445 | 0,853180905 |
| 0,856116527 | 1,024391866 | 0,928026326 | 1,05736049 | 1,128069499 | 1,062995201 | 1,56283231 | 0,725970026 | 1,087429255 | 1,374071717 | 1,374071717 |
| 1,030372882 | 0,940121754 | 0,9949424 | 0,795501026 | 0,791406526 | 1,162332919 | 0,878487578 | 1,259673402 | 1,038015005 | 1,263434755 | 1,263434755 |
| 1,182712655 | 0,763361791 | 0,61835015 | 0,820076756 | 1,215395903 | 1,204161641 | 1,013699103 | 0,450631101 | 1,413339009 | 1,975982845 | 1,975982845 |
| 1,032035347 | 0,979158005 | 0,96061496 | 1,074905074 | 2,098133088 | 0,983970589 | 0,783546478 | 1,300878661 | 1,342033335 | 0,986457081 | 0,986457081 |
| 1,01732084 | 1,196631789 | 0,864070921 | 1,02589461 | 1,171154156 | 1,003116274 | 0,895691105 | 0,952072336 | 1,289283157 | 0,949246349 | 0,949246349 |
| 1,107706131 | 0,743959135 | 0,759704813 | 0,40195522 | 0,705040044 | 1,008786409 | 0,673514176 | 0,344863238 | 1,091749273 | 1,21104404 | 1,909150194 |
| 0,989134917 | 0,845617373 | 0,930669663 | 0,390212412 | 0,788473653 | 0,898606955 | 0,571519095 | 0,460984102 | 0,951953309 | 1,2774609 | 1,684543271 |
| 0,889585482 | 0,817523619 | 0,838561591 | 0,485458745 | 0,854188808 | 1,014178511 | 0,9798212 | 0,495412387 | 1,115854944 | 2,172345353 | 2,172345353 |

| PRAS40_pT246 | PREX1 | Progesterone-Receptor | PTEN | Rab11 | Rab25 | Rad50 | Rad51 | Raptor | RBM15 | Rb_pS807_S811 |
|---|---|---|---|---|---|---|---|---|---|---|
| 1,026829772 | 1,050184247 | 1,080115009 | 1,005460294 | 1,070154834 | 1,127594726 | 1,165325057 | 0,981281383 | 1,012687225 | 1,488230682 | 0,918489598 |
| 0,986150033 | 1,04358813 | 0,999886045 | 0,892706119 | 1,035260339 | 0,986315061 | 1,215743674 | 0,91630983 | 1,029916301 | 1,282269056 | 1,032926598 |
| 0,773792109 | 1,022678141 | 1,057004039 | 1,110147839 | 1,457344071 | 0,835036777 | 0,580377653 | 1,235290424 | 0,648200038 | 0,4902366 | 0,787991418 |
| 0,860289828 | 0,804579879 | 0,884643096 | 0,895121853 | 0,996161473 | 0,99663848 | 0,904440279 | 0,874094235 | 0,796082844 | 0,937684483 | 0,824063016 |
| 1,363534304 | 1,080110593 | 0,955782828 | 0,900742544 | 0,972319181 | 1,06870357 | 1,162248611 | 1,028782333 | 1,204213241 | 1,348858366 | 1,535870354 |
| 0,788871902 | 0,923314035 | 0,925424125 | 1,061041838 | 0,940475331 | 1,201081505 | 1,072965 | 0,931954254 | 0,840652333 | 1,057102439 | 0,696476001 |
| 0,997292474 | 0,936064196 | 1,060105237 | 0,995550771 | 0,967987433 | 1,152631938 | 1,228102445 | 0,928797665 | 1,076014378 | 1,236955634 | 0,647241262 |
| 1,294809336 | 0,858397482 | 0,891238305 | 1,033930113 | 0,855440521 | 0,800711887 | 1,090968375 | 1,082620128 | 0,947546035 | 1,131769518 | 0,929656158 |
| 0,984171735 | 0,940639601 | 1,199736482 | 1,181808728 | 0,954271988 | 1,150904668 | 0,939960033 | 0,944653257 | 1,275423262 | 1,150165192 | 0,975046479 |
| 0,978690739 | 0,92799341 | 1,001319727 | 0,898331632 | 1,084447099 | 1,246051762 | 1,041827192 | 0,94836289 | 1,182267654 | 1,126962255 | 0,866168858 |
| 0,770014012 | 1,013826434 | 0,965126377 | 1,020739127 | 1,142736405 | 1,00053879 | 0,934578586 | 0,919435176 | 1,058224528 | 0,877629712 | 0,736467851 |
| 1,236758131 | 0,953706816 | 0,877118611 | 1,001693519 | 0,953086312 | 0,859924968 | 1,020105663 | 1,050610078 | 1,059602105 | 1,319493921 | 1,161033123 |
| 1,657134768 | 1,2835323 | 1,051916115 | 0,587378969 | 1,147809814 | 1,0536683 | 1,151248525 | 0,852745478 | 1,070839334 | 1,004419939 | 1,587719833 |
| 0,818671945 | 1,159905581 | 0,930516846 | 1,174992124 | 1,045454254 | 1,036391072 | 0,924701456 | 0,615684829 | 1,040885162 | 0,678607715 | 0,812520281 |
| 0,925568484 | 1 | 0,959067596 | 1,049585391 | 0,947909371 | 0,906562264 | 1,005610508 | 1,064464488 | 1,260392863 | 0,959385007 | 0,931400565 |
| 1 | 1,128448256 | 1,072010728 | 1,036028609 | 0,961147203 | 0,935339336 | 0,872554809 | 0,802718133 | 1,01900945 | 0,889768188 | 1,138858904 |
| 1,043560951 | 1,191459323 | 1,071255091 | 0,98046051 | 0,843776674 | 1,182604402 | 1,20538135 | 0,977213531 | 1,098405625 | 0,864692105 | 1,495908301 |
| 1,159998938 | 0,936584288 | 0,951473527 | 1,107356236 | 0,882316542 | 1,166960015 | 1,235989872 | 0,97106787 | 1,142692684 | 1,021645954 | 1,232852253 |
| 1,479056289 | 1,120835538 | 1,099580134 | 1,28836633 | 0,986913996 | 1,03896793 | 1,30831411 | 0,941231002 | 1,200891332 | 1,263328099 | 1,082961111 |
| 1,006134848 | 1,018823222 | 0,960372873 | 0,96636917 | 1,139939307 | 0,85464162 | 0,726684762 | 0,921384287 | 0,876795057 | 0,720612429 | 0,957007974 |
| 0,908906517 | 1,002132369 | 1,10256127 | 0,866823302 | 1,108918104 | 1,065524689 | 0,729336331 | 0,932622122 | 0,902211083 | 0,735133746 | 0,890645101 |
| 0,960068687 | 1,132609829 | 0,976047824 | 0,382377744 | 1,069150666 | 0,98851124 | 0,764002473 | 1,511219908 | 1,105820392 | 0,469676465 | 1,376224629 |
| 1,231765821 | 0,87765853 | 0,956072255 | 1,29923238 | 1,103376036 | 0,972229276 | 1,139683835 | 0,964128813 | 1,32411987 | 1,098380617 | 1,036470882 |
| 1,725056124 | 0,942346001 | 1,084541388 | 1,074399583 | 0,957563734 | 1,180198468 | 0,954416626 | 1,079978915 | 0,94852438 | 1,1572205 | 1,727301584 |
| 0,979154843 | 1,685366443 | 1,523112976 | 0,606179838 | 1,061469605 | 1,791792467 | 1,054657298 | 1,342275887 | 0,688926573 | 1,013377674 | 1,068424358 |
| 0,77933938 | 1,690271015 | 1,642377051 | 0,595035898 | 1,00560672 | 1,880065593 | 1,008970155 | 1,374152726 | 0,723492948 | 0,913103967 | 1,035497407 |
| 0,960354811 | 1,277035593 | 1,178215713 | 0,570720295 | 0,828881401 | 1,659141028 | 0,942172003 | 1,429366241 | 0,614034358 | 0,652681034 | 1,512986403 |

| Rictor | Rictor_pT1135 | RIP | RPA32_pS4_S8 | RSK | S6_pS235_S236 | S6_pS240_S244 | SDHA | Shc_pY317 | SHP-2_pY542 | SLC1A5 |
|---|---|---|---|---|---|---|---|---|---|---|
| 1,327776901 | 0,904227673 | 0,926987351 | 1,097754534 | 1,02125946 | 1,067831132 | 1,118803103 | 0,955860022 | 1,170592463 | 0,849959485 | 0,819590002 |
| 1,688080549 | 0,912132664 | 0,916520742 | 1,19844302 | 1,003642265 | 0,737863774 | 0,733162707 | 1,064857191 | 0,992518138 | 1,037960153 | 0,855939523 |
| 0,510298407 | 1,548288011 | 0,857731662 | 0,73870385 | 0,556302306 | 1,357354049 | 1,092084604 | 0,750096129 | 0,953230837 | 0,954444615 | 1,057437872 |
| 1,000124845 | 1,064405757 | 0,567508594 | 1,01525174 | 1,313705779 | 1,673320198 | 1,411966752 | 0,894708279 | 1,204986895 | 1,209626298 | 1,105843824 |
| 1,592187833 | 1,021401171 | 0,99441702 | 1,083669551 | 0,880318531 | 1,38251747 | 1,325033732 | 1,027308369 | 1,043207173 | 1,045988352 | 1,409090101 |
| 1,233754787 | 0,730237821 | 1,060230684 | 0,780858461 | 1,056856119 | 0,48004743 | 0,469201614 | 1,05933113 | 0,697811601 | 0,609918018 | 1,010402526 |
| 1,415163637 | 0,891344406 | 0,928583643 | 1,105180729 | 0,91832939 | 0,691919947 | 0,798333744 | 1,196461819 | 0,821110019 | 0,7676151 | 0,935565186 |
| 1,279913158 | 1,041237397 | 0,800409028 | 0,981491389 | 0,770303246 | 2,918392201 | 2,420848242 | 0,920334084 | 1,10976255 | 1,021305711 | 1,247543573 |
| 1,847974823 | 0,918549642 | 0,952415819 | 1,03204694 | 0,891939199 | 0,614909257 | 0,660094033 | 0,969932776 | 0,911753512 | 0,961520256 | 0,832740381 |
| 1,440554872 | 1,050723311 | 1,302719603 | 0,929120143 | 0,962877355 | 0,713635426 | 0,831610274 | 0,92640485 | 1,282852487 | 0,951288219 | 1,363967859 |
| 1,043127954 | 0,913334661 | 1,144909157 | 0,732602814 | 0,906880081 | 1,418018895 | 1,303294893 | 1,085013965 | 0,955019738 | 0,870511646 | 0,988296293 |
| 1,801383083 | 1,384682002 | 1,101947 | 1,171700148 | 1,151029781 | 2,355104631 | 1,922537544 | 0,837036446 | 1,30715014 | 1,124939813 | 0,918952598 |
| 1,642179313 | 1,582005204 | 0,935481931 | 0,996863446 | 0,930246313 | 1,322681905 | 1,184627694 | 0,879168929 | 1,149749579 | 0,940780614 | 1,03487985 |
| 0,868924675 | 1,133865453 | 0,707058739 | 0,748297783 | 0,954753242 | 1,17286237 | 0,956578839 | 1,049471131 | 0,96054705 | 1,008432028 | 1,588098731 |
| 0,910791338 | 1,08955092 | 0,883074102 | 1,045812276 | 1,190605324 | 1,044868929 | 1,046405551 | 1,051308534 | 0,821319644 | 0,979805379 | 0,845997786 |
| 1,211323734 | 1,198071475 | 0,788601404 | 1,095927888 | 1,101453729 | 1,134295824 | 1,110167932 | 0,997640531 | 0,926268476 | 1,083368978 | 0,922187547 |
| 0,980737444 | 1,016458266 | 1,329401048 | 0,985362219 | 1,087354756 | 1,121891285 | 1,108718817 | 0,887515174 | 0,916956371 | 0,886021232 | 1,461399627 |
| 1,634175436 | 0,940225632 | 0,986959835 | 1,169444326 | 1,12896797 | 1,544463149 | 1,629991542 | 0,890938102 | 1,203906973 | 1,069943743 | 0,916827794 |
| 1,759431707 | 1,041662053 | 0,975632859 | 1,057331427 | 1,355010392 | 0,720879861 | 0,794719622 | 0,866844495 | 1,195081061 | 0,978639117 | 1,246342441 |
| 1,074991705 | 1,241982476 | 0,681790275 | 0,882161103 | 0,854113558 | 1,108281641 | 0,892394841 | 0,791930953 | 0,809928989 | 1,127359521 | 1,284248093 |
| 1,105569449 | 1,137993205 | 0,819203679 | 0,849378604 | 1,043783541 | 1,214739393 | 1,208601627 | 0,919437045 | 0,800506096 | 1,02066265 | 1,180843514 |
| 0,558139547 | 1,158967018 | 1,062723131 | 0,61276954 | 0,464377333 | 1,203513849 | 0,819974074 | 0,734907078 | 1,362822239 | 0,899856811 | 1,846812047 |
| 0,881098085 | 1,01860386 | 0,96231541 | 1,126471257 | 1,52103042 | 2,832885116 | 2,181120258 | 0,981703935 | 1,204791983 | 1,172400771 | 1,590464893 |
| 1,813284839 | 1,272769216 | 1,106965475 | 1,18100596 | 0,948229105 | 1,987524831 | 1,795996851 | 0,91245412 | 1,348869159 | 1,315143672 | 1,425543203 |
| 0,688723848 | 0,96547344 | 2,894107647 | 0,759624617 | 0,472578734 | 0,326636183 | 0,431963284 | 1,096266818 | 1,724182207 | 0,945613463 | 2,656884655 |
| 0,739365977 | 0,932059632 | 2,846876191 | 0,697825335 | 0,640315147 | 0,750295868 | 0,842388825 | 1,098850902 | 1,400538692 | 1,052519798 | 2,8025146 |
| 0,481973104 | 0,921115117 | 2,131561711 | 0,70541115 | 0,323815247 | 0,608082036 | 0,495015141 | 1,44815142 | 1,328309794 | 0,800926815 | 3,272926537 |

| Slfn11 | Smad1 | Smad3 | SOD2 | Sox2 | Src-(phospho-Y416) | Src_pY527 | Stat3 | Stat3_pY705 | Stat5a | Stathmin-1 |
|---|---|---|---|---|---|---|---|---|---|---|
| 0,918623857 | 1,071437428 | 1,04506218 | 0,920966593 | 0,883479796 | 0,97909468 | 0,522086572 | 1,401620256 | 1,118735842 | 0,92915578 | 0,962649318 |
| 0,969293414 | 1,088571895 | 1,166566219 | 1,000540148 | 0,818422895 | 1,100019054 | 0,869286752 | 1,113555951 | 1,114806437 | 1,069173503 | 1,003719447 |
| 1,128110889 | 0,828359213 | 1,049931935 | 0,921803986 | 2,417890041 | 0,74283989 | 1,154773187 | 0,448183649 | 1,072901932 | 0,379111403 | 1,137406246 |
| 0,765505826 | 0,950749534 | 1,006586347 | 1,041816692 | 2,711438096 | 1,534176497 | 1,558775287 | 0,8057947 | 1,106494211 | 0,951714253 | 0,910942049 |
| 0,849013609 | 1,090749813 | 1,056073485 | 1,05534269 | 0,744420736 | 1,052690465 | 0,843577989 | 1,389401072 | 1,075107979 | 1,128045935 | 0,946188827 |
| 1,037447952 | 1,027596894 | 1,103998723 | 1,135729725 | 0,90462786 | 0,627345694 | 0,12303244 | 0,94446899 | 0,954488922 | 0,643398362 | 0,982551811 |
| 0,922516929 | 1,123663905 | 1,063988667 | 1,037429508 | 1,471420964 | 0,849704501 | 0,674218668 | 1,210414875 | 0,871785889 | 1,038119625 | 1,068474778 |
| 0,942297556 | 1,402165436 | 1,08389842 | 0,994825865 | 1,910331621 | 0,904611188 | 2,117227404 | 1,343272791 | 1,138544106 | 1,376982323 | 0,846953027 |
| 0,895071351 | 1,034519403 | 1,110066741 | 1,080294105 | 1,087054519 | 1,017552476 | 0,92122884 | 1,720470398 | 1,174754167 | 1,128746809 | 1,015071534 |
| 0,877649154 | 1,02935771 | 1,082901337 | 1,120105202 | 0,901619884 | 1,02514403 | 0,728780633 | 1,277418314 | 0,994052359 | 0,867080235 | 1,041198402 |
| 0,921115408 | 1,055394761 | 0,91558109 | 0,966029005 | 1,040276761 | 1,09812281 | 1,035573324 | 0,985044063 | 1,054973843 | 0,784816344 | 1,043290614 |
| 1,017593112 | 0,773767016 | 0,947572342 | 0,940915032 | 1,540193632 | 1,274276923 | 2,714539243 | 1,108269531 | 1,019555099 | 1,49526987 | 1,074381957 |
| 1,20587405 | 1,010473068 | 0,84762259 | 0,832702674 | 0,782062264 | 0,818433984 | 1,524217807 | 0,883232903 | 0,87865077 | 1,001933637 | 1,246140145 |
| 0,967392838 | 0,862679171 | 0,947767985 | 1,019421399 | 0,874281142 | 0,726299209 | 1,704641155 | 0,930667114 | 1,131948293 | 0,702396489 | 0,878135272 |
| 0,901255775 | 1,011681283 | 0,931631556 | 0,958315561 | 1,205213434 | 0,862850078 | 1,045838492 | 1,062029151 | 1,00251678 | 0,63155085 | 1,022442637 |
| 1,03415231 | 0,976670583 | 0,995962595 | 1,007626232 | 2,1428006 | 0,93109291 | 1,491544761 | 0,93766313 | 0,985831082 | 1,223375595 | 1,03114005 |
| 0,82459492 | 0,9990809 | 0,858783284 | 1,051122163 | 1,110157516 | 1,098191636 | 1,08765768 | 0,807702189 | 0,970468906 | 0,729399509 | 0,964376058 |
| 0,937611179 | 1,135781994 | 0,956513746 | 0,984299896 | 1,134578865 | 2,031918569 | 1,477063349 | 1,134735675 | 1,13445417 | 0,902706372 | 0,985723234 |
| 0,856614975 | 0,968765291 | 0,921856558 | 1,083755044 | 1,923988284 | 1,870535682 | 1,582694529 | 0,96764987 | 1,002503705 | 2,414335791 | 0,997447994 |
| 1,0769374 | 0,957874515 | 0,844326461 | 0,947604463 | 1,267259539 | 1,082968276 | 1,196048599 | 0,782360363 | 1,133035868 | 0,984416555 | 0,962718541 |
| 1,144566733 | 0,920763358 | 0,838346956 | 0,954583957 | 1,761443238 | 1,665446856 | 1,206365592 | 0,820192907 | 1,065077834 | 1,263570824 | 1,246487119 |
| 0,993627972 | 0,967715386 | 1,099691205 | 0,931893227 | 0,926972881 | 1,43240083 | 0,916896053 | 0,460872647 | 0,693376018 | 1,262620628 | 1,160577565 |
| 0,932416323 | 0,894875504 | 1,02984355 | 0,786149341 | 1,21022701 | 1,635977221 | 2,04671902 | 1,176483565 | 1,116789117 | 0,986350843 | 0,981218367 |
| 0,972210701 | 0,951064662 | 1,028634329 | 0,919842167 | 0,815495585 | 1,775719049 | 1,780360775 | 1,167856473 | 1,10274435 | 1,395987609 | 1,011567366 |
| 1,03052794 | 0,816469372 | 0,707021271 | 1,212889262 | 0,914164587 | 0,971642775 | 0,79699733 | 0,332741439 | 0,844866312 | 1,454613005 | 0,809861042 |
| 1,025580911 | 0,988601559 | 0,809113041 | 1,422803958 | 0,874641687 | 0,924610379 | 0,815380601 | 0,438018643 | 0,927004317 | 0,968365826 | 0,969971192 |
| 1,119553701 | 0,852868294 | 0,722236612 | 1,56309088 | 1,007487475 | 0,921373628 | 0,794155712 | 0,331404756 | 0,802799441 | 1,161463217 | 0,667561634 |

| STING | TAZ | TFAM | TFRC | TIGAR | TRIM25 | TSC1 | TTF1 | Tuberin | Tuberin_pT1462 | TUFM |
|---|---|---|---|---|---|---|---|---|---|---|
| 1,635075879 | 1,359296317 | 1,056274663 | 0,869384814 | 1,13278611 | 0,869384814 | 1,151396659 | 0,97567229 | 0,958130252 | 0,886392259 | 1,025816648 |
| 0,982024628 | 1,268853401 | 0,877941422 | 0,915806141 | 1,089387752 | 0,915806141 | 1,032319405 | 1,046630843 | 0,677876027 | 0,981624513 | 0,999831436 |
| 1,011713456 | 0,465485208 | 1,08461928 | 1,131397804 | 1,038780413 | 1,131397804 | 1,054237927 | 1,2539141 | 0,837456111 | 1,196347832 | 0,638855985 |
| 1,47436741 | 0,661706943 | 1,134269505 | 1,183189393 | 1,099347582 | 1,183189393 | 1,084454886 | 0,86817791 | 0,875792036 | 0,840215124 | 0,821835383 |
| 0,974633629 | 1,443549472 | 0,859595053 | 0,787643965 | 0,983295709 | 0,787643965 | 1,038337911 | 0,845179666 | 0,809016846 | 0,857692439 | 0,974945574 |
| 1,549220626 | 1,293820664 | 0,95865422 | 1 | 1,101641102 | 1 | 1,014487422 | 0,924794775 | 0,828342624 | 0,968715614 | 1,006542842 |
| 1,085202003 | 1,038134105 | 0,935687132 | 0,92930878 | 1,064002354 | 0,92930878 | 0,959359291 | 0,91642433 | 0,966102312 | 0,98727856 | 1 |
| 1,149854394 | 0,920100313 | 1,279611642 | 1,334799989 | 1,343524488 | 1,334799989 | 1,056019052 | 0,907074862 | 0,988013591 | 0,841041122 | 0,989853423 |
| 1,225438787 | 1,034889397 | 0,854145946 | 0,890984391 | 0,98916527 | 0,890984391 | 1,051906323 | 0,953543135 | 1,179048813 | 0,945732605 | 1,008542807 |
| 1,510490854 | 1,135999112 | 0,978444106 | 0,785632569 | 0,817040062 | 0,785632569 | 0,993863384 | 0,842603089 | 1,087244222 | 0,807889763 | 0,987736515 |
| 2,36679459 | 1,112878272 | 1,069252323 | 1,057420285 | 1,041552141 | 1,057420285 | 1,105251422 | 0,804561457 | 0,925635213 | 0,732307511 | 0,857561105 |
| 0,670499848 | 0,901166488 | 1,022822516 | 0,983226513 | 0,976799963 | 0,983226513 | 0,812152372 | 0,889535907 | 0,811724532 | 0,883532327 | 1,012428624 |
| 0,626453603 | 0,961231481 | 1,034109378 | 0,807683589 | 0,966594455 | 0,807683589 | 1,264962877 | 1,014020763 | 1,099397318 | 1,028201064 | 0,869441687 |
| 1,047677481 | 1,043896526 | 1,082642212 | 1,129335468 | 0,913044894 | 1,129335468 | 1,38968228 | 1,260241775 | 1,104330493 | 1,206961548 | 0,705038147 |
| 1,799972184 | 1,27019522 | 0,950305768 | 0,898645048 | 1,157539192 | 0,898645048 | 1,273867975 | 1,082279943 | 0,965797341 | 1,062660548 | 0,714346992 |
| 1,058683538 | 1,008819507 | 0,937176276 | 0,943653081 | 0,971113718 | 0,943653081 | 0,937562142 | 1,050143136 | 1,05668004 | 1,16759096 | 0,754845891 |
| 1,168999812 | 0,751395957 | 0,933938866 | 0,875376447 | 0,986322509 | 0,792985253 | 1,282574912 | 0,939852895 | 1,400153234 | 1,058930622 | 0,921310875 |
| 0,595467692 | 0,998670346 | 1,009989097 | 0,92304201 | 0,963909042 | 0,92304201 | 0,927910107 | 0,89705044 | 1,090000033 | 0,969785599 | 1,014627116 |
| 0,921445889 | 0,859956485 | 0,960743888 | 0,768032445 | 0,898308691 | 0,768032445 | 1,281179134 | 0,871275878 | 1,281219669 | 0,992118232 | 0,96861751 |
| 0,724453776 | 0,83392434 | 1,31725965 | 1,374071717 | 0,953358635 | 1,374071717 | 0,940550313 | 1,078820434 | 1,088365505 | 1,174422038 | 0,8660515 |
| 1,087026237 | 0,828931551 | 1,21119706 | 1,263434755 | 0,857689371 | 1,263434755 | 0,933237417 | 0,976180559 | 1,098058722 | 0,820812322 | 0,789585385 |
| 1,041800232 | 0,463642866 | 1,894284293 | 1,975982845 | 0,529852256 | 1,975982845 | 0,826252983 | 0,875533946 | 1,462614565 | 1,030065855 | 0,742439498 |
| 0,863549627 | 0,660734004 | 1,069625508 | 0,986457081 | 1,022846746 | 0,986457081 | 0,981254534 | 0,980890911 | 1,158452475 | 0,891553127 | 0,980002848 |
| 0,850209903 | 1,003548566 | 1,112537908 | 0,949246349 | 0,932273339 | 0,949246349 | 1,023307365 | 0,954602554 | 0,970263913 | 0,956309053 | 0,97701058 |
| 1,26191856 | 0,741565158 | 1,575979809 | 1,976928763 | 0,631570154 | 1,091009894 | 0,591514295 | 1,197835408 | 1,798945443 | 1,082892423 | 1,178947784 |
| 1,290339987 | 1,43595783 | 1,580472488 | 2,051373506 | 0,684403222 | 1,014495827 | 0,682705156 | 1,234314003 | 1,973353602 | 0,989256968 | 1,080809105 |
| 1,145328715 | 0,545909435 | 2,08252804 | 2,172345353 | 0,700418485 | 2,172345353 | 0,714106342 | 1,367575367 | 1,607961305 | 1,527319217 | 0,96417767 |

| Tyro3 | U-Histone-H2B | UBAC1 | ULK1_pS757 | VASP | VEGFR-2 | Wee1 | Wee1_pS642 | WIPI1 | WIPI2 | XBP-1 |
|---|---|---|---|---|---|---|---|---|---|---|
| 1,07255918 | 1,042845431 | 0,748974666 | 0,837840723 | 1,113913268 | 1,097267533 | 1,246381379 | 0,956326435 | 1,000570206 | 0,96600351 | 0,647975726 |
| 0,987093223 | 0,995767973 | 0,939918031 | 0,969356501 | 1,251708511 | 1,066924804 | 1,114774033 | 0,997267426 | 1,338772815 | 0,589059808 | 1,127666012 |
| 1,0276038 | 1,079828853 | 0,878455668 | 0,977601121 | 0,402263485 | 0,774635256 | 0,732031697 | 1,141387393 | 0,611735766 | 0,677450451 | 2,185414307 |
| 0,890684749 | 1,234574781 | 0,766242051 | 1,139340495 | 1,061754069 | 1,186093778 | 0,765541648 | 1,0684342 | 1,013784043 | 0,695603564 | 0,897645663 |
| 1,171528576 | 1,301233784 | 0,8781879 | 1,096069067 | 0,779688523 | 1,199109565 | 0,782637558 | 1,053795907 | 1,329164896 | 0,9166671 | 0,657890848 |
| 1,086816485 | 0,438279513 | 1,075785876 | 0,33769572 | 1,595227486 | 1,163114465 | 0,975400546 | 0,801252 | 1,287007281 | 1,043631826 | 1,283286161 |
| 0,980602047 | 0,699858517 | 1,087788295 | 0,788528954 | 1,557212909 | 1,111861558 | 0,601277005 | 0,921103521 | 1,289842672 | 1,024495462 | 0,923734061 |
| 0,98996246 | 0,864018691 | 0,787776362 | 1,383230848 | 0,805585135 | 1,019638737 | 0,863636024 | 0,909490935 | 1,537678844 | 0,748530755 | 1,037537422 |
| 0,954152577 | 0,937122317 | 1,018027145 | 1,145412068 | 1,527626879 | 1,156473694 | 0,968577485 | 1,01982143 | 1,482083243 | 1,022185947 | 0,635833212 |
| 1,040056215 | 0,967246325 | 1,085508827 | 1,069853995 | 1,082498761 | 0,973414116 | 1,47896526 | 0,916754334 | 0,99484005 | 0,857196901 | 0,917594445 |
| 0,872302527 | 0,742372996 | 0,979218044 | 1,169581025 | 1,193049594 | 1,034374474 | 1,094257456 | 0,780005513 | 0,948267784 | 0,723169823 | 0,860061642 |
| 1,113742461 | 1,157995209 | 0,895513666 | 1,675307352 | 0,909161011 | 1,18418114 | 1,309701952 | 0,942468713 | 0,999942859 | 1,007899515 | 0,913504959 |
| 1,079588204 | 0,933082889 | 0,980298944 | 1,679892265 | 0,874063196 | 0,805700961 | 1,343772467 | 0,998576875 | 0,987391579 | 0,915045867 | 0,970659308 |
| 0,921933626 | 1,315111646 | 0,858064275 | 0,775102793 | 0,635192392 | 0,869970701 | 1,082038401 | 0,968648781 | 0,811596538 | 0,771267487 | 2,133887863 |
| 0,910728861 | 0,991612566 | 0,895131202 | 0,840758549 | 0,953851317 | 0,805423613 | 1,007596005 | 1,067631146 | 0,914513509 | 0,876094527 | 1,282765558 |
| 0,926752851 | 0,990979786 | 0,962494503 | 0,994047009 | 1,174873738 | 1,366587942 | 0,949651109 | 1,101018024 | 1,164990058 | 0,708377833 | 1,731916461 |
| 0,942999694 | 0,885897885 | 0,99631097 | 1,121433445 | 0,992313684 | 1,024308065 | 1,365739764 | 1,03644191 | 0,779503298 | 0,964154697 | 1,414963438 |
| 0,940280013 | 1,034802207 | 1,003397844 | 1,369615012 | 1,182558465 | 1,18918494 | 0,894701773 | 0,950805209 | 1,272217306 | 0,946564871 | 0,928579136 |
| 0,954726192 | 0,872833295 | 0,844165968 | 1,253740716 | 1,917276189 | 1,559713916 | 1,169221773 | 0,950086826 | 1,079591448 | 0,911636718 | 0,741307788 |
| 0,981110491 | 1,00611435 | 1,032103172 | 1,073665852 | 0,853862968 | 1,033352872 | 0,889045433 | 0,834632144 | 0,965141604 | 0,845954408 | 1,637130024 |
| 0,958235749 | 0,898971671 | 1,044608029 | 1,082354177 | 1,616228425 | 1,110001453 | 0,83009057 | 1,007127634 | 1,106082215 | 0,843180563 | 0,99397852 |
| 0,972801733 | 0,891973046 | 2,443872343 | 1,16518945 | 0,491010326 | 0,367253548 | 1,278491146 | 0,786621367 | 0,506944255 | 1,211172773 | 1,693683025 |
| 0,9340764 | 0,94361569 | 0,954686228 | 1,268530629 | 0,775113921 | 1,123773167 | 0,690896476 | 0,963318384 | 1,067478593 | 1,018214588 | 0,748305814 |
| 0,936482116 | 1,526407697 | 0,952958055 | 1,052863051 | 0,958834023 | 0,974122018 | 1,552666944 | 0,963265354 | 0,886467178 | 0,99070981 | 1,085860798 |
| 1,204939368 | 0,789415936 | 1,33411409 | 1,012248666 | 0,474485781 | 0,410074221 | 3,335943624 | 1,369396374 | 0,825688329 | 1,759317567 | 1,005801748 |
| 1,838268388 | 0,670531846 | 1,204980554 | 1,067323773 | 0,561232926 | 0,512731773 | 3,321566053 | 1,547048179 | 0,938021711 | 1,288950335 | 1,495129191 |
| 1,007024799 | 0,471283025 | 2,107449674 | 0,929508878 | 0,556443555 | 0,373792475 | 2,704586322 | 0,952218424 | 0,6676689 | 1,462912878 | 2,243137295 |

| XPF | XRCC1 | YAP | YAP_pS127 | YB1_pS102 | ZAP-70 |
|---|---|---|---|---|---|
| 0,94430059 | 1,420583345 | 1,027471429 | 1,160888573 | 0,841021121 | 1,181052484 |
| 0,992503616 | 1,514062263 | 1,058664854 | 1,058291758 | 0,897663993 | 0,889337827 |
| 1,305872272 | 0,473663941 | 1,072489255 | 0,922037345 | 1,587006077 | 1,291317269 |
| 1,020445963 | 1,012902573 | 1,100520198 | 0,909631286 | 1,235304985 | 1,149317352 |
| 1,004789204 | 1,379089484 | 1,04738806 | 1,090037108 | 0,913510783 | 0,810192433 |
| 0,951570426 | 1,193844085 | 1,015305174 | 0,655865742 | 0,759720163 | 1,22540778 |
| 1,025387585 | 1,545770007 | 0,916037027 | 1,160431599 | 0,940369368 | 0,881753492 |
| 0,84075884 | 1,064810892 | 0,850109364 | 0,886736164 | 1,207346976 | 0,84518364 |
| 0,926809343 | 1,587427783 | 0,903708821 | 0,882089647 | 0,948125224 | 0,750375619 |
| 1,129098156 | 1,425090928 | 1,199310253 | 0,950860182 | 1,027904779 | 1,102709727 |
| 0,951354293 | 1,042938652 | 0,943320451 | 0,839951866 | 0,799429065 | 1,364403967 |
| 0,99627765 | 0,981885298 | 1,044379974 | 1,940154246 | 1,003733585 | 0,836182304 |
| 1,083841174 | 0,94557109 | 0,953787478 | 2,225132126 | 0,968826396 | 1,564966525 |
| 1,065872168 | 0,764765666 | 0,865419057 | 1,129772849 | 0,941306611 | 2,575787897 |
| 1,037132838 | 0,791980956 | 0,974060989 | 0,985422409 | 1,002339102 | 2,569526438 |
| 1,009925372 | 0,944940313 | 0,930116551 | 1,240277521 | 1,006725237 | 1,114726261 |
| 0,93967275 | 0,969887142 | 1,050808613 | 0,739110542 | 0,873258438 | 1,469035233 |
| 0,918083676 | 1,402464819 | 0,951676849 | 1,168402662 | 1,018841449 | 0,875248997 |
| 0,983971639 | 1,109258988 | 1,114748738 | 1,159869022 | 0,974711615 | 0,980132555 |
| 0,954170986 | 0,75715604 | 1,036538306 | 0,893414949 | 1,324895429 | 1,151888094 |
| 0,869829936 | 1,221220442 | 1,169252304 | 1,26294544 | 1,272555998 | 0,980939402 |
| 1,110697398 | 0,546751359 | 1,196813111 | 1,016528504 | 1,21145404 | 1,566968074 |
| 0,991756064 | 1,015020336 | 0,796090361 | 1,102338051 | 1,031982895 | 1,418294842 |
| 0,891174062 | 0,924296106 | 0,986263809 | 1,147637031 | 0,942393359 | 0,903688487 |
| 1,003463752 | 1,226152486 | 1,589685338 | 0,627133314 | 0,795000518 | 1,434648467 |
| 1,031103821 | 1,019043386 | 1,565861008 | 1,09941583 | 0,905193199 | 1,488937757 |
| 0,924088243 | 0,804190519 | 1,357329751 | 0,855845806 | 0,894325112 | 1,67185964 |

**Table S4.** Molecular/signaling characteristics of *MMTV-R26* $^{Met}$ MGT cells identified through RPPA analysis.

| sample | 14-3-3-bet | 14-3-3-zet | 4E-BP1 | 4E-BP1_pS | 53BP1 | A-Raf | ACC1 | ACC_pS79 | ADAR1 | Akt | Akt_pS473 | Akt_pT308 | AMPK-a2_p | AMPKa |
|---|---|---|---|---|---|---|---|---|---|---|---|---|---|---|
| MGT2 - A | 1,061894 | 0,773692 | 0,567263 | 0,861403 | 0,787794 | 1,135853 | 0,817423 | 0,87167 | 0,942721 | 0,906558 | 0,603454 | 0,562641 | 0,870328 | 0,70348 |
| MGT2 - B | 1,112429 | 0,780973 | 0,61677 | 1,047868 | 0,669072 | 1,210138 | 0,954432 | 1,027931 | 0,978911 | 0,876482 | 0,646971 | 0,576709 | 0,894695 | 0,898233 |
| MGT4 - A | 0,944269 | 1,147524 | 0,939736 | 1,507674 | 1,244175 | 1,131135 | 1,381581 | 1,507208 | 1,096794 | 1,075582 | 0,829757 | 0,905952 | 1,011924 | 1,007448 |
| MGT4 - B | 0,927013 | 1,143091 | 0,890366 | 1,447299 | 1,364928 | 1,10769 | 1,568612 | 1,475278 | 1,168686 | 1,200284 | 0,879378 | 1,02913 | 0,956247 | 0,942412 |
| MGT4 - C | 0,957631 | 1,053998 | 0,92813 | 1,701487 | 3,502548 | 0,826835 | 1,537085 | 1,370714 | 1,135028 | 1,066843 | 1,267612 | 1,244567 | 1,006902 | 0,937592 |
| MGT7 - A | 0,983932 | 0,817087 | 1,118397 | 0,650066 | 0,870207 | 0,872195 | 0,531382 | 0,33931 | 0,992725 | 1,013167 | 0,508949 | 0,74041 | 0,857425 | 0,679711 |
| MGT7 - B | 0,99467 | 0,813931 | 1,060866 | 0,678065 | 0,847705 | 0,901238 | 0,549096 | 0,392927 | 0,979534 | 0,898701 | 0,576662 | 0,75749 | 0,907273 | 0,701624 |
| MGT8 - A | 0,994884 | 0,961993 | 1,399697 | 0,845108 | 0,42008 | 1,023148 | 1,028445 | 1,049691 | 1,044992 | 1,079825 | 0,604259 | 0,684113 | 1,049691 | 0,872814 |
| MGT8 - B | 1 | 1 | 1,356221 | 0,904133 | 0,456771 | 1 | 0,950028 | 1,157648 | 1,026418 | 1,135973 | 0,54491 | 0,656548 | 1,039683 | 0,932238 |
| MGT9 - A | 0,963546 | 0,947508 | 1,383455 | 1,170172 | 1,580132 | 0,975289 | 1,166519 | 1,015439 | 1,07095 | 0,757149 | 0,695144 | 0,783925 | 0,936875 | 0,828407 |
| MGT9 - B | 0,926588 | 0,904043 | 1,554455 | 1,398565 | 1,607038 | 1,034138 | 1,632789 | 1,158385 | 1,134757 | 0,827822 | 0,792719 | 0,696414 | 0,933259 | 1,016165 |
| MGT9 - C | 0,915921 | 0,867385 | 1,406538 | 1,451486 | 2,418755 | 0,817541 | 1,219583 | 2,049206 | 1,086857 | 0,675495 | 0,762742 | 0,909713 | 0,975745 | 0,776867 |
| MGT11 - A | 0,984726 | 1,086006 | 1,117997 | 1,450093 | 2,448217 | 1,04708 | 1,0308 | 0,87808 | 1,087955 | 0,997418 | 1,178316 | 1,234289 | 0,977591 | 1,018972 |
| MGT11 - B | 1,019739 | 1,125547 | 1,103159 | 1,169476 | 2,214632 | 1,071563 | 1 | 0,769391 | 0,933606 | 0,99925 | 1,206755 | 1,304766 | 0,753278 | 0,939625 |
| MGT11 - C | 1,049403 | 1,157388 | 1,019803 | 1,194263 | 1,652744 | 1,122258 | 0,991028 | 0,591985 | 0,909742 | 1,065215 | 1,130654 | 1,194263 | 1,089954 | 1,027391 |
| MGT13 - A | 1,002599 | 1,015075 | 0,824867 | 0,934115 | 0,556838 | 0,957682 | 1,300158 | 1,641682 | 0,940843 | 0,980086 | 1,458444 | 1,367861 | 0,785139 | 1,080881 |
| MGT13 - B | 0,98377 | 0,992169 | 0,758636 | 0,892288 | 0,682954 | 0,991863 | 1,195894 | 1,514237 | 0,992976 | 0,992976 | 1,511358 | 1,389082 | 0,755205 | 1,031048 |
| MGT13 - C | 0,909464 | 0,946696 | 0,774656 | 0,983326 | 0,774656 | 0,919941 | 1,248657 | 1,442013 | 0,980087 | 0,954359 | 1,241198 | 1,169117 | 0,904467 | 0,992274 |

| AMPKa_pT | Annexin-I | Annexin-VI | AR | ARID1A | Atg3 | Atg7 | ATM | ATM_pS19 | ATR_pS428 | Aurora-B | Axl | b-Actin | b-Catenin | b-Catenin_ |
|---|---|---|---|---|---|---|---|---|---|---|---|---|---|---|
| 1,185521 | 1,201896 | 0,971385 | 1,253836 | 0,953191 | 1,309958 | 0,948325 | 1,554367 | 0,997391 | 0,900957 | 0,877864 | 1,247242 | 0,981407 | 0,937762 | 0,829435 |
| 1,70488 | 1,291044 | 1,047486 | 1,255994 | 0,963976 | 1,199333 | 0,994087 | 1,351598 | 1,069209 | 0,950486 | 0,878911 | 1,194812 | 0,946358 | 1,179184 | 1,067835 |
| 0,905438 | 1,211442 | 1,025903 | 0,990556 | 1,196329 | 0,816748 | 0,843275 | 1,199663 | 0,967155 | 0,99612 | 0,972562 | 0,953189 | 1,181263 | 1,584491 | 1,044274 |
| 0,803473 | 0,929031 | 0,948913 | 0,910732 | 1,283746 | 0,879378 | 0,881532 | 1,227739 | 0,938498 | 1,103638 | 0,942057 | 0,883766 | 1,232506 | 1,60217 | 1,033801 |
| 0,779997 | 1,051156 | 1,106336 | 0,875048 | 3,111983 | 0,791852 | 0,745971 | 1,387719 | 0,923196 | 1,011579 | 1,011843 | 0,865188 | 1,005124 | 1,522636 | 1,010487 |
| 1,590474 | 1,61054 | 0,95383 | 1,00534 | 0,456977 | 1,022396 | 1,765628 | 0,610392 | 1,208691 | 0,888184 | 1,019878 | 1,30298 | 0,982012 | 0,721382 | 1,175808 |
| 1,672793 | 1,568894 | 0,907313 | 0,92069 | 0,445161 | 1,3327 | 1,74681 | 0,594609 | 1,214923 | 0,913988 | 0,857242 | 1,423572 | 1,049993 | 0,809721 | 0,937747 |
| 1,245132 | 1,049691 | 0,864323 | 0,978501 | 0,701769 | 1,082987 | 1,049691 | 0,947467 | 1,051689 | 0,737665 | 1,035074 | 1,138806 | 1,02921 | 0,873431 | 0,950814 |
| 1,254709 | 0,995527 | 0,922027 | 0,92067 | 0,643109 | 1,140186 | 1,202728 | 0,904733 | 1 | 1 | 1 | 1,188744 | 1 | 0,965574 | 0,863447 |
| 0,603922 | 1,190873 | 1,085405 | 1,025226 | 1,382848 | 0,927122 | 0,954719 | 1,085511 | 0,962209 | 1,034691 | 1,084444 | 0,930748 | 1,099959 | 1,097968 | 0,981921 |
| 0,740295 | 1,102447 | 1,136863 | 1,01899 | 1,346492 | 0,888263 | 0,963272 | 1,163242 | 0,972759 | 1,052382 | 1,07011 | 0,927136 | 1,126675 | 1,423028 | 1,077272 |
| 1,149256 | 1,207681 | 1,164739 | 1,059331 | 2,683182 | 0,860233 | 0,909617 | 1,140506 | 0,982103 | 1,151429 | 1,105967 | 0,866735 | 0,817431 | 1,218215 | 1,136989 |
| 0,915916 | 0,900549 | 1,014038 | 0,92359 | 2,154478 | 0,755984 | 1,074724 | 0,92612 | 0,96124 | 1,014636 | 1,007985 | 0,835638 | 1,400796 | 1,115006 | 1,034224 |
| 0,755229 | 0,953772 | 1 | 0,882089 | 2,041648 | 0,805057 | 1,080769 | 0,920186 | 0,948127 | 0,991537 | 0,921495 | 0,851309 | 1,345652 | 1,066787 | 1,036697 |
| 0,885351 | 1,033026 | 0,96827 | 1,114102 | 1,812168 | 0,892673 | 1,105737 | 1,021269 | 0,990947 | 1,045261 | 0,855745 | 0,732509 | 1,183896 | 1,095777 | 1,103492 |
| 2,561986 | 1,159995 | 1,005641 | 1,080881 | 0,66359 | 1,068775 | 0,938826 | 1,000923 | 0,998601 | 0,947494 | 0,96468 | 1,080881 | 0,704879 | 1,04387 | 0,972783 |
| 2,307006 | 1,130674 | 1,064119 | 1,093798 | 0,682323 | 1,153175 | 0,961801 | 1,277884 | 1,026769 | 1,006654 | 1,006273 | 1,058585 | 0,786105 | 0,992976 | 0,992976 |
| 2,02011 | 1,343256 | 1,146424 | 1,081794 | 0,797618 | 1,055499 | 0,957705 | 1,236715 | 0,978045 | 1,090488 | 0,89634 | 1,075952 | 0,800506 | 0,993869 | 0,915826 |

| B-Raf | B-Raf_pS4 | B7-H4 | Bad_pS112 | Bak | BAP1 | Bax | Bcl-xL | Bcl2 | Beclin | Bid | Bim | BiP-GRP78 | BRD4 | c-Abl |
|---|---|---|---|---|---|---|---|---|---|---|---|---|---|---|
| 1,223895 | 1,122653 | 1,093275 | 0,715953 | 0,788362 | 1,106712 | 0,776384 | 0,90773 | 1,228027 | 1,097514 | 0,940269 | 0,562243 | 1,0739 | 1,085903 | 1,054685 |
| 1,263187 | 1,299605 | 1,038922 | 0,863188 | 1,000426 | 1,080545 | 0,824511 | 0,952034 | 1,212509 | 1,212942 | 1,038453 | 0,663213 | 1,141144 | 0,988585 | 0,978247 |
| 0,911298 | 0,999674 | 0,986017 | 1,02196 | 1,132256 | 0,984244 | 1,100969 | 1,301399 | 0,994814 | 0,998904 | 1,294361 | 1,213675 | 0,970877 | 0,901208 | 0,968297 |
| 0,890547 | 0,94544 | 0,923383 | 0,879378 | 1,009808 | 1,029548 | 1,158225 | 1,272455 | 0,898573 | 0,932397 | 1,218634 | 1,018356 | 0,940586 | 0,945037 | 0,94461 |
| 0,806031 | 0,909242 | 0,949624 | 0,941913 | 0,993773 | 1,06161 | 2,030401 | 1,300983 | 0,904936 | 0,980654 | 1,229038 | 1,913522 | 0,956481 | 0,925352 | 0,990287 |
| 1,441049 | 1,094409 | 1,35341 | 0,751802 | 0,842679 | 1,100368 | 0,890537 | 0,885517 | 1,292679 | 1,241176 | 0,710707 | 0,438936 | 1,106307 | 1,302782 | 0,947473 |
| 1,462311 | 1,126586 | 1,305622 | 0,80263 | 0,849793 | 1,121779 | 0,912509 | 0,903069 | 1,197689 | 1,191126 | 0,933045 | 0,457017 | 1,021451 | 1,269095 | 0,976518 |
| 1,049691 | 0,949161 | 1,049691 | 0,857831 | 0,94476 | 1,022725 | 1,049691 | 0,972194 | 1,229907 | 1,002519 | 0,922555 | 0,761124 | 0,96961 | 1,077753 | 0,975049 |
| 1,102289 | 1 | 1,036146 | 0,897856 | 0,992839 | 0,960582 | 1,049764 | 0,992575 | 1,076737 | 0,985896 | 0,931273 | 0,660989 | 1 | 1 | 1,023613 |
| 0,888575 | 0,819395 | 0,932112 | 0,840306 | 0,986622 | 1,187014 | 0,556608 | 1,179691 | 1,104289 | 1,094486 | 1,34763 | 1,679256 | 1,033744 | 1,05257 | 0,935164 |
| 0,901169 | 0,931554 | 0,90369 | 0,915099 | 1,033364 | 0,924522 | 0,549547 | 1,296238 | 1,045 | 0,971396 | 1,23442 | 1,842925 | 1,029991 | 0,924708 | 0,91739 |
| 0,860714 | 0,777683 | 0,880319 | 0,975612 | 0,952845 | 1,131275 | 0,738631 | 1,011163 | 1,018456 | 0,971889 | 1,266608 | 2,927673 | 1,197485 | 0,96661 | 0,874671 |
| 1,107544 | 1,101552 | 0,960834 | 0,969369 | 1,072554 | 0,984216 | 1,146002 | 1,04407 | 0,915916 | 0,93268 | 1,099471 | 0,915916 | 0,980163 | 0,928742 | 0,93396 |
| 1,096921 | 1,110234 | 0,963313 | 0,987888 | 1,0837 | 1,029418 | 1,108654 | 1 | 0,898583 | 0,963226 | 0,939826 | 0,910475 | 0,999969 | 0,86236 | 0,970433 |
| 1,076435 | 1,082559 | 1,01679 | 0,832599 | 0,895569 | 0,978529 | 1,798961 | 1,447057 | 0,999688 | 1,006147 | 0,960022 | 1,126184 | 1,060115 | 0,940154 | 0,943392 |
| 0,934995 | 1,040925 | 1,037432 | 1,107431 | 1,080881 | 1,080881 | 1,005591 | 0,936309 | 1,024269 | 1,007626 | 1,080881 | 1,123235 | 1,154693 | 0,911476 | 1,080881 |
| 0,954367 | 0,989338 | 1,050373 | 1,038536 | 0,972574 | 1,029129 | 0,957975 | 0,903081 | 0,975935 | 0,992976 | 1,089684 | 1,062812 | 1,154889 | 0,999625 | 1,048668 |
| 0,951745 | 0,956248 | 1,024682 | 0,947335 | 0,970021 | 1,237369 | 0,980283 | 0,942852 | 1,067794 | 0,999439 | 1,256913 | 1,082469 | 1,197774 | 0,986549 | 1,005466 |

| c-IAP2 | c-Jun_pS73 | c-Kit | c-Myc | C-Raf | C-Raf_pS3 | Caspase-3 | Caspase-7- | Caspase-8 | Caveolin-1 | CD134 | CD171 | CD20 | CD29 | CD31 |
|---|---|---|---|---|---|---|---|---|---|---|---|---|---|---|
| 0,918496 | 0,915004 | 0,965431 | 0,981334 | 1,060973 | 0,755835 | 0,845005 | 0,762433 | 1,032591 | 0,725523 | 1,090526 | 0,549646 | 0,981009 | 1,016369 | 1,149756 |
| 0,985704 | 0,825093 | 1,026583 | 1,004596 | 1,108643 | 0,838515 | 0,888827 | 0,977796 | 1,030803 | 0,813294 | 1,192647 | 0,667473 | 1,052061 | 1,095131 | 1,221096 |
| 1,006856 | 1,414215 | 1,247367 | 1,383655 | 0,925823 | 1,140186 | 1,245658 | 1,065583 | 0,986292 | 0,56746 | 0,951092 | 2,660234 | 0,885279 | 0,986292 | 0,986292 |
| 0,969264 | 1,328411 | 1,228554 | 1,369265 | 0,935355 | 1,101159 | 1,189247 | 0,967348 | 1,01302 | 0,694315 | 0,95584 | 2,711138 | 0,910352 | 0,991814 | 0,932483 |
| 0,944587 | 1,3567 | 1,25374 | 1,191528 | 0,818779 | 0,811215 | 1,245679 | 0,930035 | 1,013997 | 1,220556 | 0,91933 | 1,540577 | 1,028159 | 0,921287 | 1,017927 |
| 0,993856 | 0,772572 | 0,821842 | 0,76926 | 0,816171 | 0,713753 | 0,884546 | 0,783433 | 1,108251 | 1,530223 | 1,198067 | 0,875274 | 1,133996 | 1,054797 | 1,226073 |
| 0,940632 | 0,736436 | 1,004123 | 0,780173 | 0,857776 | 0,752684 | 0,862104 | 0,766404 | 1,025622 | 1,795102 | 1,12859 | 0,717977 | 1,200966 | 1,119994 | 1,11274 |
| 0,733027 | 0,683021 | 1,049691 | 0,857537 | 1,014642 | 0,789679 | 1,047712 | 0,890101 | 0,775516 | 0,645213 | 1,049691 | 1,055586 | 0,834976 | 1,070672 | 1,164585 |
| 0,842101 | 0,74754 | 0,969934 | 0,81273 | 1,020161 | 0,727341 | 0,992429 | 1 | 0,896456 | 0,76045 | 1,050497 | 1 | 0,881183 | 1,014009 | 0,980222 |
| 0,948238 | 1,073543 | 1,136085 | 1,086487 | 0,961086 | 1,111256 | 1,101896 | 1,107287 | 1,044865 | 0,761327 | 1,064826 | 1,860883 | 1,002297 | 1,081222 | 1,057558 |
| 0,93285 | 1,113618 | 1,264318 | 1,163378 | 0,999824 | 1,139243 | 1,148335 | 1,07105 | 1,021301 | 0,749057 | 1,011653 | 1,548518 | 0,89484 | 1,086711 | 1,060554 |
| 0,958213 | 1,146458 | 1,243732 | 1,398914 | 0,882411 | 1,034489 | 1,240448 | 1,049541 | 1,010132 | 0,93624 | 0,967422 | 0,935968 | 0,906305 | 1,094001 | 1,071474 |
| 0,961363 | 1,442785 | 1,22678 | 1,392995 | 1,077933 | 1,259473 | 1,184421 | 0,887453 | 1,064392 | 1,555904 | 1,007012 | 1,495574 | 1,043424 | 0,945704 | 0,996692 |
| 1 | 1,416055 | 1,192656 | 1,230252 | 1,076432 | 1,308681 | 1 | 0,786909 | 1,05565 | 1,317761 | 0,958718 | 1,28488 | 1,007253 | 0,923796 | 0,966162 |
| 0,998445 | 1,526874 | 1,09416 | 1,271842 | 1,116935 | 1,328125 | 1,23835 | 0,842088 | 0,991184 | 1,425577 | 1,082599 | 1,097235 | 0,986934 | 0,976464 | 1,053898 |
| 0,972557 | 0,851241 | 0,977723 | 0,66138 | 0,975072 | 0,780106 | 1,033268 | 1,089097 | 0,99442 | 1,138974 | 1,053163 | 0,736265 | 1,080881 | 0,977647 | 1,038482 |
| 1,027664 | 0,812849 | 0,979864 | 0,800291 | 0,992976 | 0,783123 | 0,978493 | 1,017177 | 1,051798 | 1,114052 | 1,080679 | 0,741673 | 1,101653 | 0,970309 | 0,997811 |
| 0,932693 | 0,774656 | 1,073687 | 0,774656 | 0,966109 | 0,774656 | 1,081962 | 1,069246 | 1,064767 | 1,31035 | 1,125949 | 0,8858 | 1,12156 | 1,021588 | 1,081539 |

| CD4 | CD44 | CD45 | CD49b | cdc25C | Cdc2_pY15 | CDK1 | Chk1 | Chk1_pS29 | Chk2 | Chk2_pT68 | Claudin-7 | COG3 | Collagen-VI | Connexin-4 |
|---|---|---|---|---|---|---|---|---|---|---|---|---|---|---|
| 1,054569 | 1,156142 | 0,87791 | 1,094281 | 1,091421 | 1,119051 | 0,679159 | 1,013519 | 1,020439 | 1,208509 | 0,956695 | 0,856098 | 0,883873 | 1,108526 | 1,235065 |
| 1,24642 | 1,273873 | 0,949786 | 1,136502 | 1,305345 | 1,214552 | 0,723676 | 1,021771 | 0,999038 | 0,812163 | 0,898052 | 0,911448 | 0,909355 | 1,1315 | 1,270129 |
| 0,99876 | 0,972213 | 1,000326 | 0,986292 | 0,986292 | 1,22034 | 0,986292 | 1,18238 | 0,967123 | 1,026179 | 1,117705 | 2,315467 | 0,986292 | 0,975158 | 1,481903 |
| 0,96519 | 0,936561 | 0,979238 | 0,995671 | 1,034419 | 1,160656 | 0,987957 | 1,254243 | 0,990695 | 1,009955 | 1,029038 | 3,334419 | 1,032629 | 0,940488 | 1,720383 |
| 0,887411 | 0,899102 | 0,971245 | 0,98271 | 1,026712 | 0,882868 | 1,007773 | 0,965172 | 0,93115 | 0,863175 | 1,020861 | 2,344901 | 0,902623 | 0,899291 | 1,704662 |
| 1,217415 | 1,002184 | 1,028723 | 1,119491 | 1,583479 | 0,694923 | 0,306231 | 0,463791 | 0,898834 | 0,782855 | 0,872164 | 0,770352 | 0,910367 | 1,528199 | 0,456565 |
| 1,444363 | 1,037082 | 1,005342 | 1,091464 | 1,17282 | 0,800371 | 0,298313 | 0,48533 | 0,824335 | 1,047917 | 0,783591 | 0,853184 | 0,988663 | 1,488683 | 0,481463 |
| 0,936586 | 1,059205 | 0,893725 | 0,919568 | 0,8307 | 0,931704 | 0,504302 | 0,672942 | 1,093122 | 0,927269 | 0,964739 | 0,782536 | 1,29849 | 1,327711 | 0,87747 |
| 0,952029 | 1 | 0,973673 | 0,972169 | 0,724317 | 0,803935 | 0,520009 | 0,619492 | 1 | 0,947297 | 0,863914 | 0,781457 | 1,231626 | 1,230031 | 0,817406 |
| 1,048041 | 1,040862 | 0,919445 | 1,050734 | 1,400052 | 1,095157 | 1,871809 | 1,501789 | 0,978244 | 0,986051 | 1,0665 | 4,392841 | 0,821603 | 0,874927 | 0,983482 |
| 1,065106 | 0,910997 | 0,934149 | 0,99722 | 1,257571 | 1,14301 | 1,945243 | 1,643339 | 0,992789 | 0,877241 | 1,079381 | 4,899628 | 0,789282 | 0,897729 | 1,12155 |
| 1,00969 | 0,953311 | 0,9413 | 0,991803 | 1,250954 | 1,043432 | 2,010503 | 1,13687 | 0,976404 | 0,955154 | 1,125934 | 3,139896 | 0,873801 | 1,109596 | 1,094807 |
| 0,948852 | 0,938507 | 0,992838 | 1,001657 | 0,966655 | 1,134977 | 1,074393 | 1,342958 | 1,065175 | 0,936922 | 0,915916 | 3,203973 | 0,646992 | 0,888274 | 2,037159 |
| 1 | 0,960823 | 1,021155 | 1,016026 | 0,961259 | 1,077636 | 1,129241 | 1,285354 | 1,012707 | 1 | 0,851877 | 3,450831 | 0,682471 | 0,829465 | 1,775288 |
| 1,052103 | 0,979011 | 0,93914 | 1,05022 | 0,968211 | 0,978439 | 0,892918 | 1,194263 | 0,909112 | 0,819878 | 1,054151 | 2,478599 | 0,693404 | 0,867176 | 1,283 |
| 1,011019 | 1,046445 | 1,0309 | 0,990018 | 1,158557 | 1,055513 | 1,182665 | 1,016582 | 0,917577 | 0,773799 | 1,116169 | 0,747267 | 1,080728 | 1,080881 | 0,887749 |
| 1,060756 | 1,075204 | 0,992976 | 0,993172 | 1,096662 | 1,101007 | 1,221405 | 1,005941 | 0,940664 | 1,054793 | 1,052724 | 0,782416 | 1,133272 | 1,046374 | 0,910616 |
| 1,098075 | 1,01595 | 1,008754 | 0,964227 | 1,224113 | 1,052534 | 1,158359 | 0,96216 | 1,044675 | 1,064677 | 1,034637 | 0,774656 | 1,211025 | 1,350151 | 0,774656 |

| Cox-IV | Cox2 | Creb | Cyclin-B1 | Cyclin-D1 | Cyclin-D3 | Cyclin-E1 | Cyclophilin | D-a-Tubuli | DJ1 | DM-Histon | DM-K9-Hist | DUSP4 | E-Cadherin | E2F1 |
|---|---|---|---|---|---|---|---|---|---|---|---|---|---|---|
| 1,259658 | 1,988633 | 1,079824 | 1,259658 | 0,913282 | 1,012924 | 1,372787 | 0,772436 | 0,952351 | 0,985684 | 1,134366 | 0,954825 | 1,191387 | 1,259658 | 0,997629 |
| 1,423938 | 2,028096 | 1,03356 | 1,423938 | 0,94692 | 1,04469 | 1,481766 | 0,713809 | 1,126583 | 0,910307 | 1,203545 | 0,984739 | 1,237066 | 0,549113 | 1,071426 |
| 0,986292 | 3,114737 | 1,031419 | 0,986292 | 1,082751 | 0,838091 | 0,934767 | 1,283673 | 0,986138 | 1,209065 | 1,019721 | 1,039302 | 0,968439 | 7,083806 | 0,903782 |
| 0,879378 | 3,721234 | 0,999428 | 0,886247 | 1,016372 | 0,859479 | 1,019574 | 1,30328 | 0,997754 | 1,168636 | 0,968357 | 1,110661 | 0,962396 | 7,167034 | 0,966766 |
| 0,785413 | 4,855096 | 1,028977 | 0,945779 | 1,066268 | 1,018029 | 1,016799 | 1,176528 | 0,927183 | 1,060958 | 0,867055 | 1,005652 | 1,100587 | 5,058402 | 0,90793 |
| 2,462769 | 0,596399 | 1,126611 | 2,462769 | 1,025772 | 1,496313 | 0,751172 | 0,734717 | 0,91303 | 0,918757 | 1,267385 | 0,867975 | 0,937578 | 0,550883 | 1,1033 |
| 2,399087 | 0,578169 | 1,109842 | 2,399087 | 0,947781 | 1,523119 | 0,743736 | 0,768799 | 0,970575 | 0,933136 | 1,318301 | 0,774962 | 0,849922 | 0,23153 | 1,176469 |
| 1,049691 | 0,704906 | 0,970748 | 1,049691 | 1,019449 | 1,121982 | 1,045211 | 0,970245 | 1,150102 | 0,97709 | 0,898029 | 0,994254 | 1,169614 | 0,393104 | 1,070575 |
| 1 | 0,702912 | 1 | 1 | 1,029775 | 1,120704 | 0,990351 | 0,869159 | 1,096888 | 0,920665 | 0,92974 | 1 | 1,166206 | 0,381053 | 0,983255 |
| 0,924905 | 0,678607 | 1,071419 | 1,063756 | 0,975416 | 0,929494 | 0,906028 | 1,463037 | 0,927157 | 1,008225 | 1,054112 | 0,975784 | 1,238182 | 7,791182 | 0,929933 |
| 0,874874 | 0,704746 | 0,951422 | 1,005142 | 0,921767 | 0,965994 | 0,935639 | 1,470465 | 0,893199 | 1,0279 | 1,045079 | 1,004843 | 1,144523 | 7,934911 | 0,963552 |
| 0,913219 | 3,605096 | 1,02977 | 0,883853 | 1,147938 | 0,863879 | 0,908661 | 4,282414 | 0,957046 | 1,134866 | 1,098194 | 1,009627 | 1,329485 | 5,972277 | 0,901221 |
| 0,915916 | 1,934289 | 1,000329 | 0,915916 | 1,027073 | 0,809139 | 0,961172 | 1,301101 | 0,998349 | 1,216465 | 1,037937 | 1,08102 | 0,963477 | 4,756514 | 1,042183 |
| 1 | 1,736373 | 1,003905 | 1 | 1 | 0,778285 | 1,086849 | 1,291978 | 1 | 1,257844 | 0,978589 | 0,995876 | 1 | 4,99747 | 1,035708 |
| 1,194263 | 1,130383 | 1,048785 | 1,194263 | 0,938969 | 0,846362 | 0,992894 | 1,218944 | 1,017813 | 1,194263 | 1,00798 | 0,949856 | 0,973502 | 4,687067 | 0,955489 |
| 1,080881 | 0,437708 | 0,999077 | 1,080881 | 0,846108 | 1,143854 | 0,996177 | 0,642598 | 0,953935 | 0,91396 | 1,080881 | 0,902855 | 0,888867 | 0,322478 | 0,983665 |
| 1,061161 | 0,435655 | 0,972379 | 1,102001 | 0,938588 | 1,197092 | 0,992976 | 0,72565 | 0,976142 | 0,955822 | 1,07933 | 0,997234 | 0,87817 | 0,346751 | 0,992976 |
| 0,964792 | 0,397165 | 1,021848 | 1,0646 | 0,895951 | 1,132812 | 1,027922 | 0,824453 | 0,929883 | 1,031285 | 1,163898 | 0,991589 | 0,877525 | 0,300729 | 1,064934 |

| eEF2 | eEF2K | EGFR | EGFR_pY11 | eIF4E | eIF4E_pS2 | eIF4G | Elk1_pS38 | EMA | ENY2 | ER | ER-a_pS11 | ERCC1 | ERCC5 | Ets-1 |
|---|---|---|---|---|---|---|---|---|---|---|---|---|---|---|
| 1,240016 | 1,138301 | 0,905569 | 0,846762 | 0,787259 | 1,098051 | 0,888196 | 0,796644 | 1,259658 | 1,109659 | 1,137187 | 0,949187 | 0,886022 | 0,845669 | 1,069425 |
| 1,066032 | 0,967526 | 0,79218 | 0,942356 | 0,753642 | 1,122609 | 0,918481 | 0,93772 | 1,278522 | 1,166643 | 1,215817 | 0,979048 | 0,951081 | 0,768567 | 1,075157 |
| 1,174564 | 0,915655 | 1,009269 | 1,037564 | 0,972674 | 0,986292 | 1,086214 | 1,077293 | 1,031009 | 1,010436 | 1,021575 | 1,228792 | 1,125016 | 0,917006 | 1,072077 |
| 1,295604 | 1,090285 | 1,000572 | 1,01711 | 1,044555 | 0,980039 | 1,068552 | 1,027935 | 0,939262 | 0,962703 | 0,953965 | 1,193922 | 1,075943 | 0,879378 | 1,062635 |
| 1,12807 | 1,286046 | 0,972334 | 1,048398 | 0,998241 | 0,958576 | 1,079686 | 1,0471 | 0,878272 | 0,864917 | 0,954116 | 1,282801 | 0,866739 | 1,061614 | 1,000162 |
| 0,765886 | 1,269658 | 0,775098 | 0,856312 | 0,722217 | 1,013625 | 0,909117 | 0,879009 | 0,867827 | 0,727736 | 1,150272 | 1,04224 | 1,028134 | 1,277025 | 1,012491 |
| 0,770799 | 1,236827 | 1,022445 | 0,883854 | 0,715981 | 0,871494 | 1,040045 | 0,862913 | 0,980447 | 0,713497 | 1,166466 | 0,99283 | 1,046081 | 1,227056 | 0,974033 |
| 1,050182 | 1,696561 | 1,031656 | 0,934935 | 1,053007 | 1,04242 | 0,932422 | 0,874211 | 1,035197 | 1,022496 | 1,040964 | 0,893406 | 0,852761 | 1,141719 | 0,997978 |
| 1 | 1,722976 | 1 | 0,941435 | 1,037225 | 1,055293 | 0,944024 | 0,850709 | 1,086355 | 1 | 1 | 0,824356 | 1 | 1,137644 | 0,945865 |
| 1,1544 | 1,563629 | 1,000628 | 0,999372 | 1,151288 | 1,080773 | 0,959125 | 0,927984 | 1,225343 | 1,094041 | 1,121031 | 1,118101 | 0,973648 | 1,112793 | 1,020349 |
| 1,334161 | 1,375525 | 0,942029 | 1,01332 | 1,188548 | 0,990034 | 1,054065 | 1,010291 | 1,138718 | 1,152744 | 1,03436 | 1,182157 | 1,00265 | 0,839628 | 1,015104 |
| 0,913071 | 1,251005 | 1,094976 | 0,93948 | 1,179872 | 1,08105 | 1,038902 | 0,978365 | 1,139986 | 0,813274 | 1,035461 | 1,247337 | 0,924404 | 0,984333 | 0,994598 |
| 0,942961 | 1,112393 | 0,932877 | 1,031285 | 0,973771 | 1,02983 | 1,070264 | 1,016025 | 0,881356 | 0,937824 | 1,017144 | 1,167598 | 1,065507 | 0,921984 | 1,022081 |
| 0,927063 | 1 | 0,924729 | 1 | 1 | 1,018788 | 1,060021 | 1,035811 | 0,954738 | 1,040266 | 0,994351 | 1,069857 | 1,134333 | 0,927598 | 1,064196 |
| 0,87363 | 0,751671 | 1,005128 | 1,038947 | 0,99787 | 1,031804 | 1,194263 | 0,96829 | 0,781894 | 0,879845 | 0,98799 | 1,105752 | 0,925828 | 0,821037 | 1,081821 |
| 0,790432 | 0,853512 | 0,581879 | 1,004013 | 0,925466 | 0,971105 | 0,904357 | 0,98689 | 1,15452 | 0,937908 | 0,985812 | 0,89951 | 0,772486 | 1,07104 | 1,008247 |
| 0,978805 | 0,799856 | 0,828141 | 1,007065 | 0,940163 | 0,934215 | 0,945456 | 0,992976 | 1,209702 | 0,920567 | 1,00193 | 0,992976 | 0,964257 | 1,050227 | 0,992976 |
| 0,900654 | 1,127506 | 0,834724 | 0,96474 | 0,88562 | 0,920009 | 0,86132 | 0,948355 | 1,335755 | 0,793471 | 0,940001 | 1,051963 | 0,911074 | 1,336473 | 0,961262 |

| FAK | FAK_pY397 | FASN | Fibronectir | FoxM1 | FoxO3a | FoxO3a-(p_ | G6PD | Gab2 | GATA3 | GATA6 | GCLM | GCN5L2 | Glutamate- | Glutaminas |
|---|---|---|---|---|---|---|---|---|---|---|---|---|---|---|
| 0,762542 | 1,196141 | 0,810006 | 1,037631 | 1,137089 | 1,111002 | 0,871278 | 1,066818 | 0,643284 | 1,114531 | 0,843342 | 1,080801 | 0,859535 | 0,914201 | 1,081074 |
| 0,792806 | 1,275762 | 0,698052 | 1,304073 | 0,942205 | 1,177071 | 0,927761 | 0,984575 | 0,66633 | 1,153312 | 0,95854 | 0,979908 | 1,048417 | 0,918305 | 1,117758 |
| 1,03781 | 0,741775 | 1,202837 | 0,976896 | 0,96187 | 0,986501 | 0,986292 | 0,988248 | 1,410846 | 1,021905 | 0,981726 | 1,013243 | 1,104967 | 1,112667 | 0,997246 |
| 0,996049 | 0,830999 | 1,211248 | 0,879378 | 1,009688 | 1,017541 | 0,940346 | 0,900315 | 1,449647 | 1,042395 | 0,915931 | 0,982869 | 1,135906 | 1,075933 | 0,976334 |
| 1,141812 | 0,753227 | 1,003459 | 1,336248 | 0,937699 | 0,912997 | 0,986162 | 0,613331 | 1,202452 | 1,149242 | 0,77123 | 1,015702 | 1,348467 | 1,048482 | 0,817721 |
| 0,739792 | 1,521569 | 0,412473 | 1,27562 | 1,187182 | 1,080331 | 0,933468 | 0,974743 | 1,385018 | 0,83044 | 0,962548 | 0,888547 | 0,662072 | 1,195354 | 1,647547 |
| 0,888473 | 1,507928 | 0,470903 | 1,16704 | 0,975707 | 1,131682 | 0,880296 | 1,300572 | 1,384154 | 0,908661 | 0,967203 | 0,933011 | 0,581375 | 1,209507 | 1,705061 |
| 0,860803 | 2,057431 | 0,612523 | 1,14058 | 1,131857 | 0,984084 | 0,938987 | 1,120574 | 1,403426 | 1,049691 | 0,933549 | 1,081153 | 0,716654 | 1,049691 | 1,319355 |
| 0,929069 | 2,080686 | 0,769715 | 1,144729 | 1,039947 | 0,938808 | 0,952774 | 1,050601 | 1,471 | 0,981223 | 1 | 1 | 0,795436 | 1,049945 | 1,334861 |
| 0,909992 | 1,578856 | 1,081749 | 0,380105 | 0,959591 | 0,895472 | 0,912026 | 0,883376 | 1,050415 | 1,291561 | 0,781301 | 1,012958 | 1,366199 | 0,949509 | 0,678607 |
| 0,972936 | 1,515036 | 1,151841 | 0,360354 | 0,902927 | 0,87463 | 0,921141 | 0,951594 | 1,075868 | 1,227786 | 0,796235 | 0,991609 | 1,439041 | 0,979275 | 0,691577 |
| 1,129294 | 0,790125 | 0,978201 | 0,88207 | 0,918045 | 0,961057 | 0,965967 | 0,867297 | 1,315301 | 1,447523 | 0,829221 | 1,070633 | 1,489573 | 1,102235 | 0,621001 |
| 1,015796 | 2,094 | 1,075027 | 1,782758 | 0,882038 | 1,069413 | 0,996682 | 0,911175 | 0,98591 | 0,978533 | 0,873067 | 1,002312 | 1,451161 | 0,975904 | 0,951769 |
| 1,034434 | 2,055549 | 1,092571 | 1,502065 | 1 | 1,076498 | 0,962462 | 0,959675 | 0,971543 | 0,983761 | 0,866174 | 0,993383 | 1,430573 | 0,892893 | 0,984287 |
| 1,067356 | 1,194263 | 0,84012 | 1,07104 | 0,813077 | 0,957962 | 0,97451 | 0,921616 | 0,535427 | 0,990592 | 0,819824 | 1,016872 | 1,194263 | 0,899856 | 1,001292 |
| 0,923132 | 0,875055 | 0,968705 | 3,653135 | 1,044375 | 0,986787 | 0,99173 | 1,018443 | 0,407522 | 1,066194 | 1,10034 | 0,980343 | 0,827587 | 0,848751 | 1,008263 |
| 0,992976 | 0,886009 | 1,097244 | 2,848528 | 1,042575 | 0,992976 | 1,020374 | 0,998626 | 0,451903 | 1,097291 | 1,051942 | 0,970851 | 0,947641 | 0,873616 | 1,019516 |
| 0,95096 | 0,788544 | 1,025306 | 4,212246 | 0,973587 | 1,034699 | 0,986574 | 0,998194 | 0,435635 | 1,051219 | 1,062311 | 0,965957 | 0,86547 | 0,917381 | 0,908475 |

| Granzyme- | GSK-3a-b | GSK-3a-b_p | Gys | Gys_pS641 | H2AX_pS14 | HER2 | HER2_pY12 | HER3 | HER3_pY12 | Heregulin | HES1 | Hexokinase | Hif-1-alpha | Histone-H3 |
|---|---|---|---|---|---|---|---|---|---|---|---|---|---|---|
| 1,311703 | 1,012587 | 0,828409 | 1,066764 | 1,035103 | 0,899889 | 1,243608 | 0,770454 | 0,934409 | 0,870773 | 1,607701 | 0,483372 | 0,976377 | 1,140818 | 0,802812 |
| 1,279028 | 1,061349 | 0,901366 | 1,104995 | 0,990326 | 0,873729 | 1,153921 | 0,800021 | 0,941719 | 0,853943 | 1,67842 | 0,481514 | 1,009066 | 1,273751 | 0,867021 |
| 0,765757 | 1,026143 | 1,123733 | 0,826931 | 0,80756 | 0,986292 | 1,326283 | 1,251007 | 1,545014 | 1,035516 | 0,828693 | 2,176057 | 0,981651 | 0,986292 | 0,995125 |
| 0,814977 | 0,986318 | 1,083039 | 0,790138 | 0,879378 | 1,038741 | 1,291541 | 1,274349 | 1,534707 | 1,042704 | 0,890163 | 2,104699 | 0,9552 | 0,965294 | 0,984859 |
| 0,924733 | 0,967438 | 1,113762 | 0,88038 | 0,915345 | 0,981348 | 0,888933 | 1,156152 | 1,190292 | 1,055818 | 1,275605 | 2,703425 | 0,876117 | 0,940594 | 1,312241 |
| 1,466264 | 1,222176 | 1,298561 | 2,01477 | 1,738773 | 0,823398 | 0,695391 | 0,893566 | 1,145377 | 1,171408 | 0,887748 | 0,850235 | 1,37076 | 1,227999 | 0,856412 |
| 1,566259 | 0,999519 | 1,315297 | 1,991356 | 1,738359 | 0,810435 | 0,688566 | 0,978483 | 1,013578 | 1,194764 | 0,884112 | 0,844202 | 1,279147 | 1,176065 | 0,834267 |
| 1,485874 | 1,049691 | 0,868171 | 1,234322 | 1,109224 | 0,986032 | 0,950573 | 1,007623 | 0,841076 | 0,994528 | 1,109324 | 1,003608 | 0,964492 | 0,966512 | 0,68668 |
| 1,326697 | 0,967983 | 0,870835 | 1,291186 | 1,176551 | 1,003537 | 0,941285 | 1,032242 | 0,859457 | 1 | 1,039001 | 0,886914 | 0,962952 | 0,971495 | 0,709353 |
| 0,809318 | 0,836527 | 0,682319 | 0,792118 | 0,822421 | 0,901464 | 1,242881 | 1,373902 | 2,025411 | 0,951749 | 0,742183 | 0,792269 | 0,972488 | 1,134741 | 1,290188 |
| 0,756066 | 0,872785 | 0,720675 | 0,749057 | 0,833574 | 0,93051 | 1,346344 | 1,379385 | 1,904472 | 1,036636 | 0,749057 | 0,811372 | 0,953873 | 1,007983 | 1,104881 |
| 0,914552 | 0,783439 | 0,817611 | 0,803345 | 0,8398 | 0,988248 | 0,940895 | 0,789941 | 1,655316 | 0,877923 | 0,67886 | 1,033503 | 1,022391 | 1,069016 | 1,455678 |
| 0,801994 | 0,977255 | 1,112029 | 0,715559 | 0,757128 | 0,989766 | 1,053359 | 1,036553 | 1,402503 | 0,991732 | 0,934402 | 0,915916 | 1,005304 | 0,964894 | 0,947744 |
| 0,793574 | 0,974201 | 1,100623 | 0,74268 | 0,773634 | 1,033764 | 1,059629 | 0,994603 | 1,333759 | 0,968128 | 0,958712 | 0,940872 | 0,966237 | 1,02826 | 1 |
| 0,737215 | 1,018895 | 1,194263 | 0,786783 | 0,825191 | 1,048297 | 0,990721 | 0,925277 | 1,194263 | 0,989645 | 0,799143 | 1,150739 | 0,911964 | 1,027755 | 0,859499 |
| 1,080881 | 1,043228 | 1,120672 | 1,245169 | 0,995294 | 0,958844 | 1,080881 | 1,080881 | 0,854295 | 0,949158 | 0,793918 | 1,873081 | 1,080881 | 1,058965 | 0,831517 |
| 1,217441 | 1,008575 | 1,155399 | 1,272453 | 1,03852 | 0,969792 | 1,05558 | 1,216846 | 0,888925 | 1,038393 | 0,796336 | 2,043334 | 1,067453 | 1,064798 | 1,073433 |
| 1,2285 | 1,02552 | 1,07909 | 1,186726 | 1,049318 | 0,934899 | 1,039629 | 1,207428 | 0,906789 | 0,96971 | 0,832813 | 2,012099 | 1,032533 | 1,051862 | 0,993721 |

| HLA-DR-DP | HSP27 | HSP27_pS8 | HSP70 | IGF1R_pY1 | IGFBP2 | IGFRb | INPP4b | IR-b | IRF-1 | IRS1 | Jagged1 | Jak2 | JNK2 | JNK_pT183 |
|---|---|---|---|---|---|---|---|---|---|---|---|---|---|---|
| 0,896759 | 1,05974 | 1,101863 | 0,861499 | 0,864816 | 1,037606 | 0,707453 | 1,078832 | 0,719394 | 1,07267 | 1,160839 | 0,877804 | 1,021255 | 1,0087 | 0,934006 |
| 0,96042 | 1,110337 | 1,410483 | 0,941684 | 0,946026 | 1,236771 | 0,783126 | 1,112324 | 0,652075 | 1,127715 | 1,216245 | 0,857484 | 1,072637 | 1,041098 | 0,998093 |
| 0,986292 | 0,984472 | 1,018431 | 1,034085 | 0,969291 | 0,977155 | 0,781153 | 1,002853 | 0,986292 | 1,099876 | 1,474196 | 1,324285 | 0,948607 | 1,078275 | 1,03866 |
| 0,971678 | 0,945253 | 1,074139 | 1,005809 | 0,977078 | 0,997982 | 0,805757 | 0,961954 | 0,937932 | 1,11047 | 1,462553 | 1,309458 | 0,902246 | 1,202202 | 1,085179 |
| 0,989732 | 0,910932 | 0,76501 | 0,971697 | 1,020121 | 0,934737 | 0,665644 | 0,937318 | 1,062646 | 0,947043 | 1,999185 | 1,471895 | 0,998334 | 1,05457 | 0,919787 |
| 1,114195 | 1,292416 | 1,850525 | 0,8511 | 1,155188 | 1,039969 | 1,397669 | 1,05622 | 1,132482 | 1,10091 | 0,616654 | 1,145361 | 1,107786 | 0,957664 | 1,153113 |
| 1,105568 | 1,169192 | 1,578925 | 0,842047 | 1,06112 | 0,806968 | 1,376594 | 1,180301 | 1,333261 | 1,033303 | 0,700103 | 1,141185 | 1,141128 | 1,024102 | 1,216773 |
| 0,82059 | 1,037078 | 1,238832 | 1,035557 | 0,917146 | 0,642351 | 1,053945 | 0,964616 | 1,005702 | 0,980884 | 0,914997 | 0,94966 | 1,114348 | 0,911499 | 1,042238 |
| 0,891701 | 0,978892 | 1,070562 | 1 | 0,886021 | 0,776586 | 1,037007 | 1 | 1,023517 | 1 | 0,905795 | 1 | 1,162812 | 0,935971 | 1 |
| 0,976916 | 0,963118 | 0,923458 | 1,16126 | 1,06846 | 1,017639 | 0,842598 | 0,989761 | 1,494401 | 1,030356 | 0,745362 | 0,911424 | 0,988365 | 0,924897 | 1,130563 |
| 0,95837 | 0,991288 | 0,975102 | 1,201655 | 1,074157 | 1,006415 | 0,898003 | 0,968127 | 1,437409 | 1,034645 | 0,749057 | 0,902065 | 1,096815 | 1,011477 | 1,133961 |
| 0,94025 | 1,025949 | 0,933546 | 1,126558 | 1,016103 | 0,915439 | 0,708788 | 0,993329 | 1,43052 | 1,051343 | 0,901679 | 0,817691 | 1,046179 | 0,944939 | 0,869666 |
| 1,001413 | 0,941154 | 0,915916 | 1,123343 | 1,002632 | 1,16021 | 0,784093 | 0,963942 | 0,943696 | 1,036191 | 1,734534 | 1,249861 | 0,915916 | 0,956424 | 1,088826 |
| 1,013277 | 1,007324 | 0,987169 | 1,07797 | 1 | 1,007688 | 0,753999 | 0,91908 | 0,958699 | 0,969949 | 1,702839 | 1,210785 | 0,925651 | 0,91354 | 1,028002 |
| 0,911345 | 1,070272 | 1,177571 | 1,10313 | 0,90812 | 0,748446 | 0,695339 | 0,905544 | 0,915111 | 0,973192 | 1,519723 | 1,000312 | 1,042953 | 0,934592 | 1,008635 |
| 1,06495 | 1,080881 | 0,876699 | 0,76536 | 1,30274 | 1,126967 | 2,163686 | 0,92218 | 1,113609 | 0,953675 | 0,770392 | 0,805412 | 0,973971 | 0,943849 | 0,94903 |
| 1,058231 | 1,000264 | 0,901644 | 0,63489 | 1,309727 | 0,992976 | 2,257407 | 0,995766 | 1,181735 | 0,940633 | 0,801736 | 0,816817 | 1,008463 | 0,992976 | 0,955506 |
| 1,006531 | 1,000431 | 0,947274 | 0,809376 | 1,213346 | 1,082695 | 2,217586 | 0,982669 | 1,506156 | 1,031972 | 0,833949 | 0,834277 | 0,93048 | 1,004341 | 0,953183 |

| LC3A-B | Lck | LDHA | LRP6_pS14 | MAPK_pT2 | Mcl-1 | MCT4 | MDM2_pS: | MEK1 | MEK1_pS2: | MERIT40_p | Merlin | MIF | MIG6 | MMP2 |
|---|---|---|---|---|---|---|---|---|---|---|---|---|---|---|
| 0,914505 | 0,947561 | 1,097064 | 0,881259 | 0,813867 | 1,263939 | 0,636594 | 0,839038 | 1,29878 | 0,96309 | 0,900957 | 1,026067 | 1,055924 | 0,981191 | 1,032118 |
| 0,891237 | 0,972464 | 1,423938 | 0,918077 | 0,823666 | 1,335978 | 0,556306 | 0,908626 | 1,196825 | 0,904556 | 1,202905 | 0,888626 | 1,110797 | 0,981025 | 1,069618 |
| 0,886561 | 1,143672 | 1,057241 | 0,959064 | 1,173354 | 0,928047 | 0,685915 | 0,97418 | 0,971572 | 1,085309 | 0,955982 | 0,791671 | 1,027109 | 0,984935 | 1,073193 |
| 0,879378 | 1,18811 | 0,884431 | 0,938115 | 1,19477 | 0,879378 | 0,64045 | 0,969209 | 1,214176 | 1,074462 | 0,936733 | 0,94119 | 0,959185 | 0,995142 | 1,030937 |
| 1,116291 | 1,487838 | 0,870414 | 0,926187 | 1,036807 | 0,705679 | 0,897716 | 0,954171 | 1,275356 | 1,064064 | 0,862393 | 1,199554 | 0,853108 | 0,999838 | 1,053517 |
| 1,94076 | 0,919758 | 1,697672 | 0,944609 | 0,938551 | 0,993564 | 2,511424 | 0,876044 | 1,850695 | 0,91616 | 1,343773 | 0,921504 | 1,197281 | 0,922639 | 1,046428 |
| 1,978578 | 0,839831 | 1,653773 | 0,932661 | 1,050316 | 1,07514 | 2,473769 | 0,851767 | 1,802839 | 0,906297 | 0,954732 | 0,855076 | 1,170437 | 0,934643 | 1,085224 |
| 1,235996 | 1,00114 | 0,723588 | 1,0612 | 0,820385 | 1,045603 | 1,828918 | 1,037531 | 0,824009 | 0,982973 | 1,050247 | 0,890142 | 1,049691 | 0,99495 | 0,955835 |
| 1,391551 | 1,061959 | 0,739739 | 1 | 0,803335 | 1,02532 | 2,325304 | 1,017689 | 0,796443 | 0,967706 | 1 | 1,102457 | 1,060962 | 1 | 1 |
| 1,082416 | 1,042357 | 1,088081 | 0,859904 | 0,757636 | 1,28532 | 0,678607 | 0,776389 | 1,284103 | 0,97594 | 0,967376 | 1,2103 | 0,856395 | 1,026565 | 1,007691 |
| 1,086228 | 1,071496 | 0,929814 | 0,869119 | 0,749057 | 1,390475 | 0,631849 | 0,819903 | 1,236984 | 0,988924 | 1,014425 | 1,248579 | 0,866837 | 1,053377 | 1,032615 |
| 1,00872 | 1,042248 | 1,04609 | 0,796018 | 0,705109 | 0,970909 | 0,691808 | 0,782271 | 1,062206 | 1,042265 | 0,998754 | 1,296272 | 0,89282 | 1,109676 | 1,043149 |
| 0,921694 | 1,113745 | 0,952829 | 0,972482 | 0,963089 | 0,655787 | 0,454252 | 0,977404 | 0,937888 | 1,052953 | 0,93747 | 0,915916 | 1,028448 | 1,038142 | 1,036368 |
| 0,938523 | 0,983491 | 0,689334 | 0,90037 | 0,894234 | 0,647882 | 0,468354 | 0,999256 | 1 | 1,000481 | 0,898529 | 1,173957 | 0,995982 | 1,077273 | 0,972589 |
| 0,881747 | 0,974678 | 0,823247 | 0,829245 | 0,977089 | 0,903511 | 0,386696 | 0,843973 | 0,995266 | 1,11559 | 1,069017 | 1,250372 | 0,930804 | 1,137435 | 1,033977 |
| 0,975295 | 0,96567 | 1,270841 | 1,405657 | 1,119624 | 1,463317 | 1,651212 | 0,763097 | 1,28465 | 0,905754 | 1,057717 | 0,628192 | 0,925212 | 0,978291 | 0,980238 |
| 0,987455 | 0,992976 | 1,239993 | 1,485759 | 1,180339 | 1,536869 | 1,720575 | 0,992976 | 1,253552 | 0,992976 | 1,113013 | 0,624711 | 0,930508 | 1,019689 | 0,971601 |
| 0,906373 | 1,105087 | 1,176899 | 1,501659 | 1,023984 | 1,388217 | 1,93326 | 0,867747 | 1,389404 | 0,975128 | 1,060261 | 0,788788 | 0,911951 | 0,97334 | 0,931438 |

| Mnk1 | MSH6 | MSI2 | mTOR | mTOR_pS2 | Myosin-He | Myosin-IIa_ | Myt1 | N-Cadherin | N-Ras | NAPSIN-A | NDRG1_pT | NDUFB4 | NF-kB-p65_ | Notch1 |
|---|---|---|---|---|---|---|---|---|---|---|---|---|---|---|
| 1,110739 | 1,04767 | 1,143311 | 0,92796 | 0,943175 | 1,07319 | 1,259658 | 0,90844 | 0,97101 | 1,046718 | 0,811153 | 0,450859 | 1,111708 | 1,088148 | 0,819963 |
| 0,964196 | 0,877233 | 1,220312 | 0,916326 | 0,977379 | 1,051917 | 1,423938 | 0,863568 | 0,998619 | 1,037752 | 0,874816 | 0,444728 | 1,16164 | 1,056013 | 0,809647 |
| 0,960173 | 1,181663 | 0,959081 | 0,99589 | 1,033089 | 1,052429 | 0,986292 | 0,986292 | 1,113882 | 1,026597 | 1,013411 | 1,221704 | 1,004114 | 1,11878 | 1,111945 |
| 0,947585 | 1,237729 | 0,914411 | 0,950432 | 1,021723 | 1,162808 | 0,879378 | 0,974612 | 1,094278 | 1,01097 | 0,949358 | 1,311155 | 0,931246 | 1,136327 | 1,083881 |
| 1,002355 | 1,207447 | 0,926345 | 1,015662 | 0,951308 | 0,928747 | 1,108681 | 0,949385 | 1,055535 | 0,913274 | 0,950624 | 1,143231 | 0,974773 | 0,871418 | 1,284239 |
| 1,329371 | 0,997198 | 0,946785 | 0,80739 | 0,911712 | 2,124396 | 2,462769 | 0,864005 | 1,134655 | 0,998131 | 0,855509 | 2,319897 | 0,955211 | 0,875263 | 0,515025 |
| 1,364377 | 1,102031 | 0,984487 | 0,818084 | 0,915238 | 2,038653 | 2,399087 | 0,772712 | 1,097919 | 1,000481 | 0,884431 | 2,218576 | 1,0007 | 0,966122 | 0,533368 |
| 1,077366 | 1,049691 | 0,985946 | 0,931217 | 0,79211 | 1,380158 | 1,049691 | 0,859924 | 1,049691 | 1,038003 | 1,097087 | 0,710741 | 0,945695 | 0,915963 | 0,958098 |
| 1 | 1,008999 | 1 | 0,942116 | 0,869397 | 1,091155 | 1 | 0,860362 | 1,073071 | 0,965365 | 1 | 0,752319 | 0,940461 | 0,972572 | 0,940667 |
| 1,051641 | 1,352641 | 0,884569 | 0,819757 | 0,702299 | 0,740417 | 0,724602 | 1,126654 | 0,747959 | 0,968858 | 0,84942 | 0,782599 | 1,007305 | 0,863791 | 0,716396 |
| 1,104656 | 1,328653 | 0,871735 | 0,845685 | 0,739029 | 0,690343 | 0,749057 | 1,153767 | 0,798436 | 0,928805 | 0,884915 | 0,796349 | 0,991527 | 0,918225 | 0,699052 |
| 1,024815 | 1,203324 | 0,977406 | 0,857807 | 0,758408 | 0,749908 | 0,703929 | 1,113355 | 0,792461 | 0,931472 | 1,021706 | 1,776848 | 1,063236 | 0,832103 | 0,70633 |
| 1,338719 | 1,046094 | 0,975996 | 1,005462 | 0,953829 | 1,051725 | 0,915916 | 0,992256 | 0,976166 | 1,03622 | 0,998272 | 2,025421 | 1,031877 | 1,114146 | 0,969763 |
| 1,209747 | 0,914 | 0,961456 | 1 | 0,982291 | 1 | 1 | 0,910119 | 0,99119 | 1,069486 | 0,987569 | 1,91767 | 1 | 1,138945 | 1,011614 |
| 1,135564 | 0,872961 | 1,014437 | 1,112106 | 1,098148 | 1,048049 | 1,194263 | 0,939898 | 0,945746 | 0,951561 | 1,029808 | 1,891997 | 1,036377 | 1,200654 | 0,783945 |
| 0,97431 | 0,916501 | 1,083088 | 1,065699 | 1,080881 | 0,93433 | 1,080881 | 0,995881 | 0,938657 | 1,012586 | 0,757738 | 0,929716 | 0,973045 | 0,716876 | 2,672888 |
| 0,986131 | 0,858202 | 1,14104 | 1,08239 | 1,082585 | 0,916485 | 0,992976 | 1,033418 | 0,956343 | 0,992976 | 0,959994 | 0,992976 | 0,99831 | 0,755466 | 2,72818 |
| 1,127816 | 0,993854 | 1,118498 | 1,041854 | 0,959881 | 0,970005 | 0,774656 | 1,043663 | 0,955736 | 0,929298 | 0,911255 | 0,977825 | 1,020037 | 0,774656 | 2,602653 |

| Notch3 | oct-04 | P-Cadherin | P-Met_pY1 | p16INK4a | p21 | p27-Kip-1 | p27_pT198 | p38-MAPK | p38_pT180 | p44-42-MA | p53 | p70-S6K1 | p70-S6K_p | p90RSK_pT |
|---|---|---|---|---|---|---|---|---|---|---|---|---|---|---|
| 0,863041 | 1,103883 | 1,075566 | 0,084234 | 0,698135 | 0,79412 | 0,917514 | 1,097539 | 0,94479 | 0,819106 | 0,977588 | 0,914048 | 1,067787 | 1,048461 | 0,788604 |
| 1,009729 | 1,09051 | 1,035056 | 0,094166 | 0,711997 | 0,82359 | 0,913999 | 0,941683 | 1,030003 | 0,823262 | 0,999574 | 0,806096 | 1,126393 | 1,145075 | 0,845638 |
| 1,081849 | 0,979321 | 1,01104 | 0,986292 | 1,558868 | 0,897469 | 0,98836 | 0,975158 | 0,933252 | 1,285209 | 0,902727 | 1,067182 | 1,289397 | 1,13788 | 1,129377 |
| 1,06292 | 1,006867 | 1,018846 | 1,114634 | 1,745272 | 0,879378 | 0,967136 | 1,002682 | 0,916942 | 1,231771 | 0,923677 | 1,001642 | 1,316478 | 1,138281 | 1,191809 |
| 1,044908 | 1,023402 | 0,996735 | 1,784451 | 2,024797 | 0,774441 | 0,983165 | 1,175819 | 0,868151 | 1,184975 | 0,877686 | 1,023084 | 1,076615 | 0,704307 | 1,038109 |
| 0,994702 | 1,166548 | 1,168682 | 0,074825 | 1,155974 | 0,914867 | 0,946758 | 0,913729 | 1,184852 | 0,964611 | 1,181568 | 0,991562 | 1,17002 | 1,900563 | 0,540575 |
| 0,947347 | 1,11147 | 1,412444 | 0,073108 | 1,169048 | 0,890207 | 0,882859 | 0,929997 | 1,24017 | 0,971148 | 1,237145 | 0,945685 | 1,176312 | 2,021467 | 0,563025 |
| 1,00557 | 0,932013 | 1,039683 | 0,770386 | 1,78203 | 0,999559 | 1,019551 | 0,949258 | 0,984844 | 0,904842 | 1,297959 | 0,946465 | 1,049691 | 1,00262 | 0,711899 |
| 1,023948 | 1 | 1 | 0,755592 | 1,674917 | 1,113509 | 1 | 0,997304 | 1,001487 | 0,89603 | 1,144891 | 0,857037 | 1,115049 | 0,954797 | 0,698215 |
| 1,025898 | 1,006048 | 1,113545 | 1,282598 | 1,08649 | 0,821372 | 0,945984 | 1,014086 | 1,208296 | 1,055021 | 0,728582 | 1,101162 | 1,024335 | 0,754359 | 0,951753 |
| 1,029546 | 1,045591 | 1,119174 | 1,429457 | 1,05396 | 0,820063 | 0,929048 | 1,036666 | 1,342873 | 1,078659 | 0,720221 | 1,051569 | 1,105879 | 0,808552 | 0,95733 |
| 1,118091 | 0,984603 | 1,121748 | 0,317081 | 1,165569 | 0,851564 | 0,968926 | 1,218161 | 0,933056 | 1,105284 | 0,704125 | 1,150461 | 1,038137 | 0,73706 | 1,150384 |
| 1,059402 | 1,038081 | 0,959042 | 1,493735 | 0,998882 | 0,919929 | 0,979979 | 1,109273 | 1,020949 | 1,120617 | 1,003503 | 1,019117 | 0,994312 | 0,724977 | 1,029505 |
| 1 | 0,968069 | 0,936242 | 1,212844 | 1 | 0,787326 | 1,006803 | 1,120244 | 1 | 1,085248 | 1 | 1,017184 | 0,996949 | 0,737939 | 0,93254 |
| 1,050156 | 1,015303 | 1,083471 | 1,223094 | 0,98531 | 0,69866 | 1,009572 | 1,194263 | 1,092962 | 1,194263 | 1,005653 | 0,725031 | 0,914426 | 0,878778 | 1,214363 |
| 0,891613 | 1,044472 | 0,891577 | 2,04571 | 0,347246 | 1,00753 | 0,824371 | 0,968216 | 0,952038 | 0,904827 | 0,994374 | 0,884883 | 0,880907 | 1,080881 | 1,076377 |
| 0,941664 | 1,019627 | 0,951473 | 1,86761 | 0,373882 | 1,069552 | 0,896777 | 1,034655 | 1,03336 | 0,925596 | 1,062354 | 0,992976 | 0,965911 | 1,132224 | 1,086228 |
| 0,918719 | 1,040605 | 0,928164 | 1,848636 | 0,385501 | 1,025307 | 0,86841 | 1,005849 | 0,946718 | 1,038849 | 1,04144 | 0,938744 | 0,924055 | 0,856342 | 0,914855 |

| PAI-1 | PAICS | PAK1 | PAK4 | PAR | PARP | Paxillin | PCNA | PD-1 | PD-L1 | Pdcd4 | PDGFR-b | PDHK1 | PDK1 | PDK1_pS24 |
|---|---|---|---|---|---|---|---|---|---|---|---|---|---|---|
| 0,898206 | 0,899554 | 0,977166 | 0,905988 | 3,52169 | 1,014518 | 0,745916 | 1,047238 | 0,939747 | 1,084449 | 1,110518 | 2,132013 | 1,038907 | 0,725539 | 0,685787 |
| 0,946851 | 0,968838 | 1,032286 | 0,926292 | 3,168963 | 0,888372 | 0,812777 | 0,938238 | 0,907776 | 1,125584 | 1,023391 | 2,154569 | 1,05005 | 0,758166 | 0,754924 |
| 0,996996 | 0,986292 | 1,49457 | 1,137789 | 1,271324 | 1,111637 | 0,986292 | 0,986292 | 1,010366 | 0,965586 | 0,947221 | 0,292474 | 0,894261 | 1,059967 | 1,02363 |
| 0,95498 | 0,954782 | 1,455662 | 1,14418 | 1,252301 | 1,252256 | 0,997498 | 1,010882 | 0,97341 | 0,960295 | 0,965982 | 0,325128 | 0,879378 | 0,994958 | 1,09082 |
| 0,860225 | 1,132277 | 1,285362 | 0,903726 | 0,634258 | 1,506098 | 1,101156 | 0,939375 | 0,891963 | 0,888868 | 1,077382 | 0,336405 | 0,86028 | 0,938042 | 1,044656 |
| 0,947997 | 1,10535 | 0,653418 | 1,383926 | 0,733318 | 0,756448 | 0,798164 | 0,922082 | 1,346829 | 1,15629 | 0,814151 | 5,552944 | 1,225257 | 0,893287 | 1,196531 |
| 0,883629 | 1,054578 | 0,668512 | 1,469062 | 0,838122 | 0,726995 | 0,789633 | 0,980529 | 1,267889 | 1,166716 | 0,745634 | 5,358982 | 1,313244 | 0,95132 | 1,215594 |
| 0,955129 | 1,021966 | 1,43772 | 0,961848 | 0,938854 | 0,847564 | 0,93454 | 0,974502 | 0,9088 | 1,223371 | 1,688251 | 2,002783 | 1,123006 | 0,946861 | 1,020485 |
| 0,962453 | 0,936545 | 1,61388 | 1 | 0,873599 | 0,628519 | 0,955589 | 0,995165 | 0,958437 | 1,060939 | 1,678876 | 2,135528 | 1,001002 | 1 | 1 |
| 0,900852 | 1,682711 | 0,694692 | 1,069138 | 2,492672 | 1,235254 | 1,018913 | 1,155265 | 0,962998 | 0,99789 | 0,902372 | 0,331679 | 0,871916 | 1,027362 | 1,147451 |
| 0,913508 | 1,613203 | 0,729471 | 1,029141 | 1,63618 | 1,288657 | 1,046976 | 1,11368 | 0,942904 | 0,941878 | 0,749057 | 0,333568 | 0,823667 | 1,063667 | 1,26348 |
| 0,897457 | 1,282533 | 0,65461 | 0,899877 | 5,561362 | 1,264099 | 1,035885 | 0,959608 | 0,97637 | 0,969296 | 0,860719 | 0,344078 | 0,865263 | 0,991715 | 1,146981 |
| 0,997508 | 1,775151 | 1,081841 | 1,02332 | 1,892851 | 1,472783 | 1,083211 | 1,01934 | 0,976054 | 0,955445 | 0,687464 | 0,277468 | 0,916044 | 1,008656 | 1,103926 |
| 1,006409 | 1,694393 | 1,018422 | 1,025801 | 1,572881 | 1,498993 | 1,084063 | 1,042377 | 0,979124 | 0,941523 | 0,741277 | 0,277066 | 0,923225 | 1,07099 | 1,102947 |
| 0,861673 | 1,093352 | 0,934637 | 1,104929 | 2,727331 | 1,194263 | 0,98977 | 0,999101 | 0,992601 | 0,920481 | 0,648838 | 0,305694 | 0,876347 | 0,944978 | 1,061658 |
| 1,011665 | 0,967408 | 0,960372 | 0,832853 | 0,53854 | 0,845681 | 1,029585 | 0,99872 | 1,080881 | 0,958723 | 1,277697 | 0,993901 | 0,989227 | 0,840384 | 0,93099 |
| 0,992976 | 1,007387 | 0,992976 | 0,841185 | 0,574823 | 0,962993 | 1,070555 | 1,000626 | 1,062276 | 0,992976 | 1,412184 | 0,992976 | 1,046585 | 0,943649 | 0,927254 |
| 0,913241 | 1,224047 | 1,023523 | 0,831612 | 0,662971 | 1,088861 | 1,026606 | 0,949289 | 1,050629 | 1,034087 | 1,415768 | 1,046861 | 1,082944 | 0,889433 | 0,842514 |

| PEA-15 | PEA-15_pS | PI3K-p110- | PI3K-p110- | PI3K-p85 | PKA-a | PKC-b-II_p | PKC-delta_ | PKCa | PKM2 | PLC-gamm | PLK1 | PMS2 | Porin | PRAS40 |
|---|---|---|---|---|---|---|---|---|---|---|---|---|---|---|
| 0,976802 | 1,046995 | 1,054171 | 0,880419 | 1,428987 | 0,902652 | 0,607773 | 1,031526 | 1,324588 | 0,903764 | 0,93152 | 1,259658 | 1,152988 | 1,001692 | 1,142735 |
| 1,093581 | 0,84125 | 1,097873 | 0,922782 | 1,515364 | 0,952908 | 0,546593 | 1,082015 | 1,335694 | 0,905369 | 0,775467 | 1,423938 | 1,153188 | 1,204022 | 1,196842 |
| 1,14046 | 0,918047 | 0,975627 | 0,941413 | 1,007006 | 0,801776 | 0,986292 | 0,970673 | 1,002463 | 0,833253 | 1,081073 | 0,986292 | 1,042513 | 1,007718 | 0,977942 |
| 1,118576 | 0,91692 | 0,966788 | 0,912976 | 0,979056 | 0,861551 | 1,0589 | 0,944551 | 1,04481 | 1,064896 | 1,062875 | 0,879378 | 1,082366 | 1,009418 | 1,025759 |
| 1,110059 | 1,113443 | 0,942971 | 0,785663 | 1,019701 | 0,730758 | 0,860507 | 0,95065 | 0,722393 | 1,114624 | 1,051733 | 0,727944 | 1,122476 | 0,962666 | 0,953268 |
| 1,274792 | 1,077846 | 0,9203 | 0,852661 | 0,615112 | 0,908381 | 0,653587 | 0,930886 | 1,314891 | 1,370196 | 1,03794 | 2,462769 | 1,984845 | 1,03798 | 1,30746 |
| 1,273922 | 0,853542 | 0,891007 | 0,865341 | 0,65354 | 0,896583 | 0,728707 | 1,021641 | 1,32434 | 1,297676 | 1,0294 | 2,399087 | 1,93352 | 1,010901 | 1,157014 |
| 0,987162 | 0,989523 | 1,049691 | 1,106701 | 0,939246 | 1,436631 | 1,034382 | 1,066736 | 1,232597 | 1,038408 | 1,01445 | 1,049691 | 0,950864 | 1,041064 | 1,035254 |
| 0,967184 | 1 | 1,003802 | 1,048916 | 1 | 1,377012 | 1,06541 | 1,029349 | 1,221836 | 1,008262 | 1 | 1 | 1,116078 | 1,023568 | 1,023313 |
| 0,932827 | 0,893957 | 0,973993 | 0,848465 | 0,941515 | 0,981068 | 0,909642 | 0,956721 | 0,639098 | 0,733996 | 1,058238 | 0,83733 | 1,302833 | 0,943646 | 1,062461 |
| 1,002702 | 0,891957 | 0,988706 | 0,882149 | 0,949127 | 1,008176 | 0,929906 | 0,926407 | 0,655327 | 0,817686 | 0,969465 | 0,780588 | 1,199742 | 0,968531 | 0,956968 |
| 0,982227 | 1,193647 | 0,977315 | 0,888005 | 0,902108 | 0,818894 | 0,783816 | 0,914259 | 0,741639 | 0,703929 | 0,928072 | 0,784114 | 1,27348 | 0,983572 | 0,973293 |
| 1,310397 | 0,906001 | 0,953254 | 0,937614 | 0,855493 | 0,915916 | 1,027191 | 0,966757 | 0,915925 | 1,1292 | 0,996704 | 0,915916 | 0,960355 | 0,98049 | 0,968615 |
| 1,395458 | 0,899868 | 0,959016 | 0,958459 | 0,916988 | 0,890778 | 1,064473 | 0,948354 | 0,914129 | 1,15473 | 0,97896 | 1 | 0,990358 | 1 | 1,005535 |
| 1,140258 | 1,464637 | 0,992413 | 0,871614 | 0,964311 | 1,048716 | 0,784235 | 0,938908 | 0,687841 | 0,993798 | 1,049061 | 1,194263 | 0,962505 | 1,116794 | 1,053413 |
| 1,031055 | 0,696158 | 1,02767 | 0,960553 | 1,04227 | 1,773768 | 0,909855 | 0,97001 | 0,822355 | 1,076318 | 1,084376 | 1,080881 | 1,042686 | 0,979486 | 1,024282 |
| 0,992976 | 0,925313 | 1,044659 | 0,992976 | 1,118699 | 1,844069 | 0,984381 | 0,992976 | 0,828453 | 1,053712 | 1,130979 | 0,992976 | 0,992976 | 0,942149 | 0,992976 |
| 0,715045 | 0,842654 | 1,063596 | 0,939521 | 1,164416 | 1,74345 | 0,904724 | 0,981662 | 0,774656 | 0,968784 | 1,09432 | 0,9679 | 1,185635 | 0,875676 | 0,990293 |

| PRAS40_pT | PREX1 | Progestero | PTEN | Rab11 | Rab25 | Rad50 | Rad51 | Raptor | Rb | RBM15 | Rb_pS807_ | Rheb | Rictor | Rictor_pT1 |
|---|---|---|---|---|---|---|---|---|---|---|---|---|---|---|
| 0,783757 | 1,073379 | 1,174212 | 1,336228 | 1,020127 | 1,003993 | 1,012127 | 1,121063 | 1,070231 | 1,027466 | 1,231582 | 1,162312 | 0,778012 | 0,912333 | 0,906903 |
| 0,764754 | 1,129822 | 1,188086 | 1,423938 | 0,954738 | 0,940699 | 0,900009 | 1,191678 | 1,128669 | 0,939827 | 1,144504 | 1,158007 | 0,734241 | 0,957358 | 0,967295 |
| 0,60011 | 1,069112 | 0,981959 | 0,965334 | 0,98977 | 0,989371 | 1,162593 | 1,096145 | 1,112089 | 1,001873 | 0,986292 | 1,393446 | 1,07106 | 1,098694 | 0,916777 |
| 1,292289 | 1,140074 | 0,972801 | 0,937297 | 1,00058 | 0,952765 | 1,280059 | 1,111387 | 1,18112 | 1,029062 | 1,029465 | 1,475538 | 1,189663 | 1,093224 | 0,909495 |
| 1,09101 | 1,137543 | 0,948076 | 0,749324 | 0,909056 | 0,832323 | 1,425593 | 1,124988 | 1,067114 | 1,00624 | 3,78744 | 1,003546 | 1,135085 | 1,015395 | 0,899912 |
| 0,774025 | 1,25608 | 1,241883 | 0,930775 | 0,88355 | 1,102194 | 0,538179 | 1,379008 | 0,914127 | 0,953005 | 1,069425 | 0,336628 | 0,827067 | 0,863217 | 1,264299 |
| 0,75428 | 1,354762 | 1,292414 | 0,928534 | 0,953 | 1,174134 | 0,524263 | 1,14427 | 0,850631 | 0,94241 | 1,041461 | 0,348682 | 0,859554 | 0,895435 | 1,296224 |
| 0,945084 | 0,976633 | 1,014381 | 1,317207 | 0,918381 | 1,061478 | 0,593171 | 0,952739 | 0,999043 | 1,023819 | 0,455679 | 0,841791 | 1,002801 | 0,937996 | 0,953124 |
| 0,903163 | 0,958168 | 1 | 1,253493 | 1,062546 | 1,024762 | 0,627811 | 0,916004 | 1 | 1,019329 | 0,696113 | 0,810632 | 0,752078 | 1 | 0,953164 |
| 0,559187 | 1,256472 | 1,055534 | 0,679171 | 1,122393 | 0,804382 | 1,871582 | 1,338084 | 0,952034 | 1,005181 | 1,307094 | 1,622494 | 1,095746 | 0,890037 | 0,875883 |
| 1,130864 | 1,23638 | 1,048138 | 0,714014 | 1,121154 | 0,936372 | 1,808515 | 1,430562 | 0,921165 | 1,010791 | 1,337885 | 1,743532 | 1,140396 | 0,926827 | 0,886213 |
| 1,094882 | 1,188966 | 1,016674 | 0,658882 | 1,01341 | 0,961149 | 1,828486 | 1,293817 | 1,016772 | 1,007513 | 3,939999 | 1,130987 | 1,112371 | 0,921975 | 0,769547 |
| 1,004489 | 1,140382 | 0,964185 | 0,877598 | 0,915916 | 0,915916 | 1,561186 | 1,044909 | 0,979831 | 1,006165 | 1,062864 | 0,848774 | 0,9852 | 1,129026 | 0,952193 |
| 1,058061 | 1 | 0,968968 | 0,948905 | 0,951064 | 0,933334 | 1,516273 | 1 | 0,977127 | 1 | 0,984865 | 0,875929 | 1,019216 | 1,129124 | 0,960566 |
| 1,210965 | 1,07844 | 0,955618 | 1,048221 | 0,784721 | 0,913909 | 1,136368 | 1,030089 | 0,990088 | 1,072304 | 1,358911 | 0,732814 | 0,864039 | 1,123118 | 1,366657 |
| 1,080881 | 0,862464 | 1,176043 | 1,253849 | 0,732141 | 0,94207 | 0,644533 | 1,062883 | 1,011068 | 1,019936 | 1,019114 | 1,495186 | 1,003743 | 0,856162 | 1,080881 |
| 1,024481 | 0,94472 | 1,219129 | 1,288876 | 0,923808 | 0,988196 | 0,621569 | 1,039385 | 1,075762 | 0,98768 | 1,015032 | 1,584516 | 0,992976 | 0,90795 | 1,094828 |
| 1,092973 | 1,058045 | 1,17501 | 1,263834 | 0,891319 | 0,929279 | 0,774656 | 1,03984 | 1,054841 | 0,997396 | 0,954631 | 1,592242 | 0,920337 | 0,850301 | 1,06254 |

| RIP | RPA32 | RPA32_pS4 | RSK | S6 | S6_pS235_ | S6_pS240_ | SCD | SDHA | SF2 | Shc_pY317 | SHP-2_pY5 | SLC1A5 | Slfn11 | Smac |
|---|---|---|---|---|---|---|---|---|---|---|---|---|---|---|
| 1,181462 | 0,902496 | 0,834532 | 0,465701 | 0,746581 | 1,281591 | 1,259658 | 1,146944 | 1,041615 | 0,941627 | 0,805473 | 1,032299 | 1,042276 | 0,973074 | 0,938562 |
| 0,977379 | 0,848516 | 0,884014 | 0,468922 | 0,816224 | 1,341056 | 1,313391 | 1,228249 | 1,144441 | 0,964809 | 0,637371 | 1,057344 | 0,856969 | 1,016487 | 0,990142 |
| 0,986292 | 1,106276 | 0,988174 | 0,986292 | 1,03895 | 1,717998 | 1,354109 | 0,98964 | 0,904659 | 1,032003 | 0,99054 | 0,9417 | 0,815319 | 0,789243 | 0,985701 |
| 0,856519 | 1,050118 | 0,961886 | 0,95735 | 1,031898 | 1,701257 | 1,363355 | 0,974749 | 0,914631 | 1,014511 | 0,946344 | 0,916077 | 0,966327 | 0,998141 | 1,023628 |
| 0,946453 | 1,011263 | 1,012369 | 0,939953 | 1,305651 | 0,853301 | 1,16766 | 0,925124 | 0,766047 | 1,028592 | 0,864704 | 0,915577 | 0,961345 | 0,924048 | 1,075798 |
| 1,195042 | 0,767256 | 0,700419 | 0,215158 | 0,629747 | 2,27565 | 1,260633 | 1,149747 | 1,001869 | 0,89197 | 0,700647 | 1,289157 | 1,395135 | 0,961575 | 1,073861 |
| 0,660471 | 0,71784 | 0,674217 | 0,209594 | 0,652722 | 2,373635 | 1,327655 | 1,067152 | 0,963634 | 0,89968 | 0,705503 | 1,343456 | 1,282507 | 0,941553 | 1,043602 |
| 1,226192 | 1,064484 | 0,701889 | 0,558934 | 0,978109 | 0,635405 | 0,696846 | 0,887881 | 1,049691 | 0,93067 | 1,048791 | 1,221494 | 1,535829 | 0,810122 | 0,960266 |
| 1,147143 | 0,831412 | 0,681842 | 0,557407 | 0,900187 | 0,639939 | 0,629177 | 0,851971 | 1,015112 | 0,972333 | 0,971891 | 1,264247 | 0,985279 | 0,832196 | 1,01648 |
| 1,204609 | 1,255403 | 0,938543 | 0,802354 | 0,905193 | 0,678607 | 0,709937 | 1,009378 | 0,789176 | 0,981654 | 0,986828 | 1,00644 | 1,348535 | 0,949338 | 1,154516 |
| 1,132935 | 1,209103 | 0,948976 | 0,796276 | 1,000704 | 0,773581 | 0,813484 | 1,013187 | 0,841349 | 0,962714 | 0,934543 | 1,022754 | 1,253565 | 0,969546 | 1,140416 |
| 1,161952 | 1,167104 | 1,146793 | 0,751148 | 1,237398 | 0,428371 | 0,580502 | 0,995239 | 0,836571 | 1,011899 | 0,735213 | 0,7595 | 1,711417 | 0,946611 | 1,189539 |
| 0,783258 | 1,293438 | 1,040753 | 1,008705 | 1,07154 | 0,344742 | 0,417637 | 0,961793 | 0,877276 | 1,02072 | 0,915916 | 0,915916 | 0,892876 | 0,964031 | 0,999671 |
| 0,904736 | 1,165429 | 1,065931 | 1,044497 | 1 | 0,331246 | 0,409092 | 1,008705 | 0,938382 | 1,020975 | 0,939939 | 0,904562 | 1,026959 | 1,047598 | 1 |
| 0,717186 | 0,965577 | 1,194263 | 0,918296 | 1,322706 | 1,165454 | 1,080735 | 1,038724 | 0,86035 | 1,124701 | 0,87002 | 0,872038 | 1,139195 | 0,867117 | 0,891845 |
| 1,03134 | 0,954596 | 0,711159 | 1,376112 | 0,957385 | 1,469344 | 1,447141 | 1,000954 | 1,089926 | 0,9761 | 1,120447 | 0,971196 | 1,517346 | 0,82489 | 1,011563 |
| 1,366179 | 0,992976 | 0,662134 | 1,378299 | 0,959057 | 1,579205 | 1,579105 | 0,992976 | 1,155491 | 0,992976 | 1,100488 | 1,010714 | 1,441858 | 0,992976 | 1,009156 |
| 1,301459 | 1,126663 | 0,815662 | 1,338494 | 0,934666 | 1,080032 | 1,225084 | 1,006028 | 1,255046 | 0,902937 | 0,996162 | 0,917054 | 1,530496 | 0,981443 | 1,169876 |

| Smad1 | Smad3 | Smad4 | Snail | SOD1 | SOD2 | Sox2 | Src | Src-(phospl | Src_pY527 | Stat3 | Stat3_pY70 | Stat5a | Stathmin-1 | STING |
|---|---|---|---|---|---|---|---|---|---|---|---|---|---|---|
| 1,082396 | 0,963851 | 1,107341 | 0,904425 | 0,838599 | 1,221219 | 1,38889 | 0,601047 | 0,512646 | 0,510417 | 1,259658 | 1,090569 | 1,576881 | 0,863542 | 0,813923 |
| 1,049896 | 1,065299 | 1,183335 | 0,963041 | 0,885144 | 1,271683 | 1,423938 | 0,684988 | 0,523765 | 0,401414 | 1,202713 | 1,12751 | 1,609319 | 0,939941 | 0,846419 |
| 1,037288 | 1,122139 | 0,986292 | 0,950724 | 1,002961 | 0,946824 | 1,095054 | 1,136968 | 1,080051 | 1,175358 | 0,669456 | 0,914568 | 0,885516 | 1,012527 | 0,806323 |
| 1,041788 | 1,217699 | 0,984753 | 0,939587 | 1,10311 | 0,896522 | 1,061933 | 1,00123 | 1,119247 | 1,262509 | 0,714927 | 0,879378 | 0,879378 | 0,947004 | 0,83733 |
| 0,962545 | 1,172112 | 0,893888 | 1,159437 | 1,001231 | 0,863877 | 1,172984 | 1,031802 | 0,975759 | 1,490128 | 1,119321 | 1,040828 | 0,89371 | 0,839185 | 4,009113 |
| 1,06995 | 0,780971 | 1,085551 | 0,950322 | 1,2161 | 1,316446 | 0,932964 | 0,911917 | 0,508993 | 0,760177 | 1,507206 | 1,45068 | 1,051604 | 0,82839 | 1,219278 |
| 1,014657 | 0,956947 | 1,334128 | 0,934397 | 1,054741 | 1,28038 | 0,912672 | 0,993729 | 0,509645 | 0,790757 | 1,478969 | 1,452552 | 1,115838 | 0,793561 | 1,18775 |
| 1,028638 | 1,049691 | 1,052074 | 0,922043 | 1,122959 | 1,036134 | 0,75066 | 0,850825 | 1,103428 | 0,969073 | 1,288514 | 1,070253 | 1,141989 | 1,031499 | 1,060777 |
| 1 | 0,987907 | 0,957665 | 0,927141 | 1 | 0,976759 | 0,78346 | 0,81054 | 1 | 1,011001 | 1,316595 | 1,132736 | 1,12437 | 0,961004 | 1 |
| 0,8977 | 1,107282 | 1,112234 | 0,882002 | 1,034168 | 0,95237 | 1,364227 | 0,777335 | 1,16542 | 0,729647 | 0,716158 | 0,805673 | 0,634698 | 0,916496 | 0,780969 |
| 0,896368 | 1,166307 | 1,097544 | 0,876115 | 1,055071 | 0,912918 | 1,47808 | 0,865483 | 1,209812 | 0,779335 | 0,758824 | 0,78038 | 0,648879 | 0,911128 | 0,659505 |
| 0,959011 | 1,247945 | 1,030162 | 0,900939 | 1,04714 | 1,010153 | 1,763691 | 0,985463 | 0,622069 | 1,185747 | 0,456359 | 0,808032 | 0,770242 | 0,852639 | 1,531909 |
| 1,209829 | 1,231951 | 1,021224 | 0,989996 | 0,916334 | 0,965742 | 0,971682 | 1,40293 | 1,04105 | 1,177925 | 0,546442 | 0,848521 | 0,871091 | 1,021188 | 0,775639 |
| 1,183958 | 1,133744 | 0,999862 | 1 | 0,94548 | 1,002203 | 0,954323 | 1,321462 | 1,069283 | 1,139062 | 0,562763 | 0,850637 | 0,941419 | 1,088611 | 0,668304 |
| 0,98145 | 1,144056 | 0,979113 | 1,032098 | 0,989807 | 1,016056 | 0,858994 | 1,286863 | 0,856594 | 1,110187 | 0,591812 | 0,927176 | 0,929465 | 0,91177 | 3,138413 |
| 0,880075 | 0,837783 | 0,989218 | 1,681781 | 0,989958 | 1,080881 | 0,868808 | 0,935915 | 1,054052 | 0,774316 | 1,350186 | 1,241614 | 1,736671 | 1,006443 | 1,291906 |
| 0,951255 | 0,933189 | 0,999298 | 1,705629 | 0,9059 | 1,04897 | 1,024981 | 0,992976 | 1,073301 | 0,813692 | 1,423168 | 1,239971 | 1,80706 | 0,992976 | 1,468064 |
| 0,94258 | 1,056667 | 0,970758 | 1,858618 | 1,128111 | 1,012296 | 0,936067 | 0,912849 | 0,993566 | 0,774656 | 1,390717 | 1,215533 | 1,847587 | 0,930463 | 1,634673 |

| Syk | Tau | TAZ | TFAM | TFRC | TIGAR | Transglutar | TRIM25 | TSC1 | TTF1 | Tuberin | Tuberin_p1 | TUFM | TWIST | Tyro3 |
|---|---|---|---|---|---|---|---|---|---|---|---|---|---|---|
| 1,074244 | 0,959291 | 0,990624 | 1,20484 | 1,379739 | 0,939645 | 1,033771 | 1,259658 | 0,747865 | 1,169061 | 0,927574 | 1,052066 | 1,010892 | 1,046581 | 0,981486 |
| 1,102293 | 1,023852 | 1,100224 | 1,142537 | 1,421291 | 0,976072 | 1,059209 | 1,423938 | 0,759268 | 1,273517 | 0,977036 | 1,059944 | 0,831692 | 1,095127 | 1,0183 |
| 0,986292 | 0,986292 | 1,224967 | 0,881934 | 0,984458 | 1,075635 | 0,939404 | 0,986292 | 0,893141 | 1,039432 | 1,000745 | 0,986292 | 0,986292 | 1,009453 | 0,986292 |
| 1,056112 | 0,981408 | 1,258005 | 0,962908 | 0,967946 | 1,072875 | 1,057714 | 0,879378 | 0,893214 | 0,97635 | 1,102116 | 0,973655 | 1,1635 | 1,045235 | 1,019718 |
| 1,063328 | 0,964459 | 1,224135 | 1,169587 | 0,97674 | 1,258528 | 0,922339 | 0,634258 | 0,810645 | 0,93856 | 1,20703 | 0,948248 | 1,115084 | 1,013226 | 1,009674 |
| 1,190794 | 1,327599 | 0,973302 | 1,678335 | 2,458191 | 1,038322 | 1,396761 | 2,462769 | 1,610557 | 1,131865 | 1,053345 | 1,302918 | 0,982739 | 1,174665 | 0,925463 |
| 1,177116 | 1,196987 | 1,112522 | 1,634937 | 2,394627 | 1,000822 | 1,239524 | 2,399087 | 1,608897 | 1,125417 | 0,917884 | 1,24131 | 0,969138 | 1,138804 | 0,875287 |
| 0,839293 | 1,032911 | 0,867753 | 1,482198 | 1,158962 | 0,821894 | 1,120102 | 1,049691 | 0,998306 | 1,041181 | 0,880775 | 0,965632 | 0,944583 | 0,98312 | 0,957661 |
| 0,84849 | 1,053653 | 0,893387 | 1,147018 | 1 | 0,891323 | 1,226213 | 1 | 1,071241 | 1,026241 | 0,971407 | 0,983699 | 1,089152 | 0,986286 | 0,971463 |
| 1,05805 | 1,014498 | 1,020113 | 1,323555 | 1,45203 | 0,91122 | 0,821903 | 0,749394 | 0,73616 | 1,071473 | 1,181354 | 0,959244 | 1,117791 | 1,010339 | 1,025931 |
| 0,999853 | 0,976049 | 1,055618 | 1,242711 | 1,490942 | 0,911167 | 0,874578 | 0,749057 | 0,763629 | 1,039137 | 1,073047 | 0,915718 | 1,107489 | 1,000147 | 1,007447 |
| 1,017571 | 1,001246 | 1,116552 | 1,243276 | 1,489813 | 1,234703 | 0,885556 | 0,703929 | 0,729089 | 1,030157 | 1,121181 | 0,918775 | 1,299448 | 1,011748 | 1,019717 |
| 1,021438 | 0,948098 | 1,177182 | 0,877693 | 0,914214 | 1,001338 | 1,05069 | 0,915916 | 1,048317 | 1,019502 | 0,915916 | 0,996666 | 1,182885 | 1,037334 | 1,059461 |
| 1,090665 | 0,960511 | 1,20603 | 0,899211 | 0,998141 | 1 | 1 | 1 | 1,01268 | 0,980498 | 0,964328 | 1,0037 | 1,110885 | 1 | 1,049124 |
| 1,045612 | 0,983868 | 1,174754 | 1,11783 | 1,192043 | 1,02003 | 1,060762 | 1,194263 | 1,027419 | 0,97134 | 0,915512 | 1,161093 | 1,103812 | 1,01903 | 0,930678 |
| 0,992542 | 1,029271 | 0,984202 | 1,080881 | 1,078872 | 0,919597 | 0,908978 | 1,080881 | 0,747589 | 0,972761 | 1,173056 | 1,011499 | 0,896285 | 1,058337 | 1,035127 |
| 1,007876 | 1,05616 | 0,992976 | 1,101499 | 1,218849 | 0,979227 | 0,887558 | 0,992976 | 0,796463 | 0,992976 | 1,342246 | 1,006145 | 0,899027 | 1,054572 | 1,043953 |
| 0,993236 | 1,04099 | 0,886637 | 1,286304 | 1,683917 | 0,958202 | 0,883189 | 0,774656 | 0,781883 | 0,975501 | 1,413788 | 0,978797 | 1,033829 | 1,110529 | 1,059052 |

| U-Histone- | UBAC1 | UGT1A | ULK1_pS75 | VASP | VEGFR-2 | VHL-EPPK1 | Vimentin | Wee1 | Wee1_pS6 | WIPI1 | WIPI2 | XBP-1 | XPA | XPF |
|---|---|---|---|---|---|---|---|---|---|---|---|---|---|---|
| 0,932433 | 0,898303 | 1,060004 | 1,245882 | 0,598098 | 0,879417 | 1,210167 | 1,036088 | 1,259658 | 1,123111 | 1,02301 | 1,139143 | 1,114943 | 1,050436 | 0,998308 |
| 1,037185 | 0,835064 | 1,105376 | 1,284125 | 0,631731 | 0,900164 | 1,182996 | 1,118407 | 1,176998 | 1,127334 | 0,97011 | 0,795944 | 1,215033 | 1,11419 | 1,028056 |
| 1,036492 | 1,050215 | 0,795553 | 0,935606 | 1,190166 | 1,178648 | 1,015777 | 1,014584 | 0,953329 | 1,222099 | 1,217283 | 0,844962 | 0,985623 | 0,932375 | 0,986292 |
| 0,839695 | 1,054645 | 0,76337 | 0,808469 | 1,159558 | 1,247721 | 0,879378 | 0,959078 | 0,944934 | 1,183285 | 1,301786 | 0,933654 | 0,981335 | 1,008925 | 0,939108 |
| 0,984641 | 0,895345 | 1,004169 | 0,977794 | 1,155481 | 1,236111 | 0,855087 | 0,925779 | 0,995263 | 1,046773 | 1,899271 | 1,068879 | 0,968069 | 0,985571 | 0,870396 |
| 0,916765 | 0,911688 | 2,462769 | 1,429545 | 0,79692 | 1,325867 | 1,524105 | 1,215629 | 0,936991 | 0,976631 | 0,933282 | 2,074235 | 1,131528 | 1,196846 | 0,859216 |
| 1,122818 | 0,877129 | 2,468861 | 1,447459 | 0,816582 | 1,351333 | 1,25214 | 1,24864 | 0,849265 | 0,790907 | 1,07879 | 1,308433 | 1,115546 | 1,092217 | 0,877089 |
| 0,081435 | 1,049691 | 1,788952 | 1,33562 | 1,049691 | 1,364159 | 0,822632 | 1,000441 | 1,032001 | 0,927317 | 0,948974 | 0,975437 | 0,92972 | 1,049691 | 0,957794 |
| 0,99782 | 0,968053 | 1,650543 | 1,393383 | 1,054479 | 1,340881 | 0,755416 | 0,960072 | 0,68125 | 0,98528 | 1 | 1,055161 | 0,998378 | 1,208721 | 0,922925 |
| 0,946214 | 1,072158 | 0,639407 | 0,898969 | 0,883596 | 0,866494 | 1,361868 | 0,878781 | 1,326334 | 1,222926 | 1,371579 | 1,23088 | 0,985686 | 1,310597 | 0,935231 |
| 0,935863 | 1,096475 | 0,660479 | 0,878981 | 0,896601 | 0,920926 | 1,194448 | 0,962162 | 1,346947 | 1,163324 | 1,442467 | 1,204506 | 0,986511 | 1,060791 | 0,914426 |
| 1,025952 | 1,015073 | 0,719598 | 1,061122 | 0,807071 | 1,054887 | 1,234776 | 0,941561 | 1,362145 | 1,236206 | 1,155141 | 1,189526 | 0,925046 | 1,059591 | 0,865821 |
| 1,004319 | 0,954146 | 0,737858 | 1,006484 | 1,230342 | 0,878723 | 1,100234 | 0,98415 | 1,025474 | 1,190371 | 0,905985 | 0,817811 | 0,980568 | 0,871255 | 0,924479 |
| 1,046263 | 0,939226 | 0,711903 | 1 | 1,24579 | 0,861742 | 0,820683 | 1 | 0,953305 | 1,13304 | 0,927986 | 0,888417 | 1 | 1,225553 | 0,925501 |
| 1,125591 | 0,845798 | 0,875309 | 0,962292 | 0,773446 | 0,826634 | 0,850939 | 0,99149 | 0,7043 | 1,066693 | 1,114688 | 0,959914 | 1,010789 | 0,914011 | 0,955512 |
| 1,080881 | 1,027974 | 1,210541 | 1,059406 | 0,526837 | 0,79279 | 1,384301 | 1,016531 | 1,25686 | 0,971692 | 1,172463 | 1,080881 | 1,026143 | 0,984104 | 1,019593 |
| 1,052176 | 1,023191 | 1,138754 | 1,149461 | 0,517509 | 0,839287 | 1,315587 | 1,035149 | 1,239583 | 0,992976 | 1,20629 | 1,210376 | 1,057007 | 1,037756 | 0,999736 |
| 0,992426 | 1,013035 | 1,384123 | 1,007291 | 0,617872 | 0,774656 | 1,1213 | 0,999569 | 1,306187 | 0,988184 | 1,31791 | 1,147144 | 1,012588 | 1,100193 | 0,986878 |

| XRCC1 | YAP | YAP_pS127 | YB1_pS102 | ZAP-70 |
|---|---|---|---|---|
| 1,071328 | 0,994923 | 0,777948 | 0,841969 | 1,096174 |
| 0,98798 | 0,985166 | 0,822088 | 0,915462 | 1,105441 |
| 0,967433 | 1,003541 | 0,945719 | 1,083507 | 0,986292 |
| 1,027219 | 0,955034 | 0,879378 | 1,094627 | 0,948043 |
| 1,07879 | 0,932544 | 1,2831 | 0,919685 | 0,918945 |
| 0,795401 | 0,925105 | 1,748823 | 0,880809 | 1,238507 |
| 0,855545 | 0,910758 | 1,795218 | 0,87392 | 1,262238 |
| 0,811682 | 1,027301 | 1,485821 | 0,93884 | 1,011524 |
| 0,718895 | 0,992591 | 1,514294 | 0,896192 | 1,091368 |
| 1,164403 | 1,015211 | 0,725 | 0,849462 | 1,198034 |
| 1,035781 | 0,91377 | 0,734288 | 0,911244 | 1,142111 |
| 1,188473 | 1,058353 | 0,785413 | 0,859468 | 1,068986 |
| 1,074997 | 1,018775 | 0,796259 | 1,025891 | 1,040723 |
| 0,948487 | 1 | 0,79319 | 1 | 0,953993 |
| 0,9029 | 0,858556 | 0,75535 | 0,934128 | 0,959888 |
| 1,080881 | 0,96763 | 1,188918 | 1,067591 | 0,97983 |
| 1,219603 | 1,028987 | 1,190246 | 1,088315 | 1,111305 |
| 1,27131 | 1,002983 | 1,135393 | 0,974743 | 1,091465 |

### ANOVA - Subtype A (MGT4, 9, 11) vs. MGT2

| # | Protein | p-value | Ratio (MGT4, 9, 11 vs. 2) | Fold change (MGT4, 9, 11 vs. 2) | F(cell lines) | SS(cell lines) | SS(Error) | F(Error) |
|---|---|---|---|---|---|---|---|---|
| 173 | PDGFR-b | 1,08E-13 | 0,146377 | -6,83167 | 2680,48 | 45,9792 | 0,0314478 | 1 |
| 220 | Stat5a | 2,64E-10 | 0,519908 | -1,92342 | 212,947 | 2,68086 | 0,0230804 | 1 |
| 106 | Heregulin | 2,92E-06 | 0,531313 | -1,88213 | 15,8753 | 1,22152 | 0,141065 | 1 |
| 149 | Notch1 | 0,0748195 | 1,14099 | 1,14099 | 261,231 | 8,7418 | 0,0613504 | 1 |
| 218 | Stat3 | 0,000198521 | 0,553762 | -1,80583 | 22,5764 | 2,24621 | 0,182405 | 1 |
| 114 | IGFRb | 0,610093 | 1,03396 | 1,03396 | 224,45 | 5,12321 | 0,041847 | 1 |
| 16 | AMPKa_pT172 | 0,00315167 | 0,57961 | -1,7253 | 20,1532 | 5,16458 | 0,469821 | 1 |
| 192 | PTEN | 6,24E-07 | 0,610168 | -1,63889 | 36,5922 | 1,00525 | 0,0503651 | 1 |
| 184 | PKCa | 2,95E-05 | 0,611766 | -1,63461 | 18,1378 | 1,02674 | 0,103781 | 1 |
| 99 | Granzyme-B | 4,49E-06 | 0,627723 | -1,59306 | 41,2121 | 1,35911 | 0,0604605 | 1 |
| 180 | PI3K-p85 | 1,16E-08 | 0,644194 | -1,55233 | 65,5844 | 0,787884 | 0,0220243 | 1 |
| 227 | TRIM25 | 0,00053431 | 0,646933 | -1,54576 | 45,9436 | 4,35433 | 0,173755 | 1 |
| 187 | PLK1 | 0,000101582 | 0,671224 | -1,48982 | 74,7899 | 4,09094 | 0,100282 | 1 |
| 20 | Atg7 | 0,60119 | 0,979264 | -1,02117 | 90,159 | 1,23961 | 0,0252068 | 1 |
| 129 | MCT4 | 0,867443 | 1,03122 | 1,03122 | 84,0128 | 9,79881 | 0,21383 | 1 |
| 19 | Atg3 | 6,48E-05 | 0,674586 | -1,48239 | 11,0157 | 0,46522 | 0,0774264 | 1 |
| 206 | S6_pS240_S24 | 0,0212966 | 0,681984 | -1,46631 | 8,71822 | 1,99056 | 0,41859 | 1 |
| 143 | Myosin-IIa_pS | 0,000470945 | 0,684166 | -1,46163 | 56,6744 | 4,16063 | 0,13459 | 1 |
| 62 | Cox-IV | 0,00012595 | 0,701733 | -1,42504 | 81,2113 | 3,84541 | 0,0868097 | 1 |
| 181 | PKA-a | 0,55918 | 0,964983 | -1,03629 | 75,668 | 2,16153 | 0,0523711 | 1 |
| 25 | Axl | 1,52E-06 | 0,71399 | -1,40058 | 41,9318 | 0,580668 | 0,0253878 | 1 |
| 128 | Mcl-1 | 0,00361495 | 0,715149 | -1,39831 | 11,3896 | 1,13117 | 0,18208 | 1 |
| 126 | LRP6_pS1490 | 0,88489 | 0,994255 | -1,00578 | 66,1155 | 0,789773 | 0,0218998 | 1 |
| 101 | Gys | 4,96E-07 | 0,725166 | -1,379 | 324,947 | 2,62375 | 0,014803 | 1 |
| 145 | N-Cadherin | 0,0559853 | 0,960772 | -1,04083 | 64,9493 | 0,208575 | 0,00588748 | 1 |
| 125 | LDHA | 0,00230132 | 0,735287 | -1,36001 | 19,2759 | 1,35548 | 0,12892 | 1 |
| 65 | Cyclin-B1 | 0,00024449 | 0,735434 | -1,35974 | 82,2768 | 3,5958 | 0,0801235 | 1 |
| 188 | PMS2 | 0,344717 | 0,957489 | -1,0444 | 60,6591 | 1,46813 | 0,0443722 | 1 |
| 123 | LC3A-B | 0,215517 | 1,08316 | 1,08316 | 58,4615 | 1,87329 | 0,0587456 | 1 |
| 142 | Myosin-Heavy- | 0,0813902 | 0,880941 | -1,13515 | 58,0716 | 2,47971 | 0,078285 | 1 |
| 236 | ULK1_pS757 | 8,34E-05 | 0,749204 | -1,33475 | 23,8444 | 0,644312 | 0,0495395 | 1 |
| 29 | B-Raf | 8,33E-08 | 0,762985 | -1,31064 | 111,25 | 0,617595 | 0,0101776 | 1 |
| 214 | SOD2 | 7,50E-07 | 0,763649 | -1,3095 | 35,5291 | 0,303646 | 0,0156684 | 1 |
| 21 | ATM | 0,000815902 | 0,770212 | -1,29834 | 18,4361 | 0,970152 | 0,0964744 | 1 |
| 163 | p70-S6K_pT38 | 0,0367157 | 0,77218 | -1,29503 | 19,7553 | 2,1438 | 0,198949 | 1 |
| 166 | PAK1 | 0,559137 | 1,03392 | 1,03392 | 49,2351 | 1,5473 | 0,0576156 | 1 |
| 110 | HSP27_pS82 | 0,0200727 | 0,775754 | -1,28907 | 10,1753 | 1,07461 | 0,193617 | 1 |
| 17 | AR | 0,000157581 | 0,779128 | -1,28349 | 7,24728 | 0,173602 | 0,0439159 | 1 |
| 98 | Glutaminase | 0,000155622 | 0,780194 | -1,28173 | 80,6271 | 1,4637 | 0,0332823 | 1 |
| 207 | SDHA | 0,000392208 | 0,785629 | -1,27286 | 12,7619 | 0,273363 | 0,0392706 | 1 |
| 219 | Stat3_pY705 | 3,10E-05 | 0,786986 | -1,27067 | 62,5686 | 0,749615 | 0,0219646 | 1 |
| 172 | Pdcd4 | 0,000937141 | 0,789402 | -1,26678 | 71,9431 | 1,77986 | 0,0453565 | 1 |
| 59 | COG3 | 0,0971174 | 0,920635 | -1,08621 | 38,9353 | 0,588651 | 0,0277176 | 1 |
| 247 | YAP_pS127 | 0,476629 | 1,0691 | 1,0691 | 37,918 | 2,09533 | 0,101309 | 1 |
| 30 | B-Raf_pS445 | 0,000232716 | 0,796073 | -1,25617 | 10,5878 | 0,221288 | 0,0383174 | 1 |
| 139 | MSI2 | 4,27E-06 | 0,79778 | -1,25348 | 18,0605 | 0,144807 | 0,0146995 | 1 |
| 102 | Gys_pS641 | 3,33E-05 | 0,817827 | -1,22275 | 189,262 | 1,40427 | 0,0136028 | 1 |
| 60 | Collagen-VI | 0,0210347 | 0,821627 | -1,2171 | 13,1673 | 0,713145 | 0,0992937 | 1 |
| 229 | TTF1 | 7,57E-06 | 0,824851 | -1,21234 | 15,007 | 0,108667 | 0,0132754 | 1 |
| 191 | Progesterone-I | 8,65E-08 | 0,83835 | -1,19282 | 94,235 | 0,221118 | 0,00430183 | 1 |
| 174 | PDHK1 | 0,000351092 | 0,841501 | -1,18835 | 28,3303 | 0,29458 | 0,019063 | 1 |
| 243 | XBP-1 | 1,63E-05 | 0,841555 | -1,18828 | 13,4597 | 0,085545 | 0,011652 | 1 |
| 215 | Sox2 | 0,0197289 | 0,846989 | -1,18065 | 19,6576 | 1,20302 | 0,112197 | 1 |
| 90 | FoxO3a | 0,000710501 | 0,850018 | -1,17645 | 7,09168 | 0,0949975 | 0,0245586 | 1 |
| 131 | MEK1 | 0,0862033 | 0,888455 | -1,12555 | 23,975 | 1,28431 | 0,098209 | 1 |
| 36 | Beclin | 0,000579213 | 0,851373 | -1,17457 | 10,1533 | 0,129077 | 0,0233067 | 1 |
| 225 | TFRC | 0,0726572 | 0,830355 | -1,2043 | 23,0032 | 3,23592 | 0,2579 | 1 |
| 171 | PD-L1 | 0,00109392 | 0,858836 | -1,16437 | 10,7281 | 0,133421 | 0,0228005 | 1 |
| 108 | Hexokinase-II | 0,342714 | 0,967622 | -1,03346 | 22,6331 | 0,23349 | 0,0189132 | 1 |
| 81 | ER | 0,000131643 | 0,861309 | -1,16102 | 12,1807 | 0,0968804 | 0,0145816 | 1 |
| 118 | IRS1 | 0,152006 | 1,14885 | 1,14885 | 21,6986 | 2,81443 | 0,237794 | 1 |
| 135 | MIF | 0,0016136 | 0,862612 | -1,15927 | 12,857 | 0,162293 | 0,023142 | 1 |
| 140 | mTOR | 0,218133 | 1,03657 | 1,03657 | 21,6487 | 0,141672 | 0,0119976 | 1 |
| 69 | DM-Histone-H | 0,00112926 | 0,862785 | -1,15904 | 16,9499 | 0,224844 | 0,0243195 | 1 |
| 8 | A-Raf | 0,0484134 | 0,865165 | -1,15585 | 2,42788 | 0,121051 | 0,0914079 | 1 |
| 52 | CD4 | 0,0237995 | 0,866791 | -1,15368 | 7,06838 | 0,237332 | 0,0615571 | 1 |

| | | | | | | | | |
|---|---|---|---|---|---|---|---|---|
| 50 | CD134 | 0,00160553 | 0,86804 | -1,15202 | 6,86224 | 0,0886066 | 0,0236724 | 1 |
| 88 | Fibronectin | 0,612654 | 0,870297 | -1,14903 | 20,1859 | 16,8349 | 1,52898 | 1 |
| 115 | INPP4b | 0,000924067 | 0,876443 | -1,14098 | 7,69294 | 0,0688118 | 0,0163988 | 1 |
| 209 | SHP-2_pY542 | 0,0370133 | 0,877828 | -1,13918 | 13,4274 | 0,381753 | 0,0521231 | 1 |
| 160 | p44-42-MAPK | 0,00353761 | 0,884115 | -1,13108 | 48,2315 | 0,455429 | 0,0173114 | 1 |
| 71 | DUSP4 | 0,00934556 | 0,885848 | -1,12886 | 17,8602 | 0,340866 | 0,0349897 | 1 |
| 31 | B7-H4 | 0,000169984 | 0,887563 | -1,12668 | 55,7304 | 0,254248 | 0,00836387 | 1 |
| 3 | 14-3-3-beta | 0,000312794 | 0,888024 | -1,1261 | 7,01879 | 0,0383397 | 0,0100145 | 1 |
| 183 | PKC-delta_pS6 | 0,000290416 | 0,895524 | -1,11666 | 7,28739 | 0,0321434 | 0,00808651 | 1 |
| 45 | C-Raf | 0,00690107 | 0,900781 | -1,11015 | 12,349 | 0,127875 | 0,0189843 | 1 |
| 179 | PI3K-p110-a | 3,95E-05 | 0,901476 | -1,10929 | 17,5983 | 0,0447948 | 0,00466657 | 1 |
| 39 | BRD4 | 0,0244602 | 0,904827 | -1,10518 | 15,2571 | 0,215063 | 0,0258425 | 1 |
| 244 | XPF | 0,00502473 | 0,912061 | -1,09642 | 5,6978 | 0,036381 | 0,011706 | 1 |
| 78 | eIF4E_pS209 | 0,0133372 | 0,916362 | -1,09127 | 5,70269 | 0,0556886 | 0,0179031 | 1 |
| 151 | 04-oct | 0,00265902 | 0,918227 | -1,08906 | 8,60402 | 0,0456702 | 0,00973133 | 1 |
| 40 | c-Abl | 0,0120691 | 0,926762 | -1,07903 | 5,76258 | 0,0348276 | 0,0110802 | 1 |
| 197 | Raptor | 0,029677 | 0,929489 | -1,07586 | 10,2826 | 0,0973311 | 0,0173536 | 1 |
| 22 | ATM_pS1981 | 0,00410807 | 0,929732 | -1,07558 | 26,8204 | 0,106612 | 0,00728759 | 1 |
| 156 | p27-Kip-1 | 0,00932017 | 1,06517 | 1,06517 | 12,5046 | 0,0442074 | 0,00648135 | 1 |
| 91 | FoxO3a-(p_S31 | 0,011967 | 1,06793 | 1,06793 | 5,07456 | 0,0205944 | 0,00744033 | 1 |
| 97 | Glutamate-D1- | 0,028718 | 1,09586 | 1,09586 | 15,1267 | 0,181095 | 0,0219485 | 1 |
| 200 | Rictor | 0,00154301 | 1,10861 | 1,10861 | 27,7896 | 0,161173 | 0,0106329 | 1 |
| 61 | Connexin-43 | 0,15408 | 1,1711 | 1,1711 | 15,6721 | 3,01638 | 0,352858 | 1 |
| 112 | IGF1R_pY1135 | 0,019286 | 1,10887 | 1,10887 | 21,4428 | 0,272795 | 0,0233237 | 1 |
| 223 | TAZ | 0,0145134 | 1,11144 | 1,11144 | 14,9438 | 0,237227 | 0,0291034 | 1 |
| 132 | MEK1_pS217_ | 0,00307844 | 1,11846 | 1,11846 | 6,74308 | 0,0568754 | 0,0154635 | 1 |
| 150 | Notch3 | 0,0063406 | 1,12403 | 1,12403 | 4,82015 | 0,0564768 | 0,0214808 | 1 |
| 238 | VEGFR-2 | 0,00990612 | 1,13284 | 1,13284 | 57,4204 | 0,813718 | 0,0259806 | 1 |
| 76 | EGFR_pY1173 | 0,000443573 | 1,13345 | 1,13345 | 9,80828 | 0,056272 | 0,0105182 | 1 |
| 24 | Aurora-B | 0,0207669 | 1,13493 | 1,13493 | 3,69143 | 0,0699945 | 0,0347624 | 1 |
| 235 | UBAC1 | 0,0139717 | 1,14396 | 1,14396 | 4,508 | 0,0808853 | 0,0328948 | 1 |
| 146 | NAPSIN-A | 0,0403182 | 1,14475 | 1,14475 | 2,99184 | 0,0810062 | 0,0496389 | 1 |
| 199 | Rb_pS807_S81 | 0,766182 | 1,03693 | 1,03693 | 14,168 | 2,7481 | 0,355602 | 1 |
| 231 | Tuberin_pT146 | 0,0661435 | 0,93259 | -1,07228 | 14,1568 | 0,169301 | 0,0219248 | 1 |
| 196 | Rad51 | 0,888414 | 1,00602 | 1,00602 | 13,7869 | 0,317955 | 0,0422807 | 1 |
| 167 | PAK4 | 0,0631488 | 1,13252 | 1,13252 | 13,7548 | 0,466078 | 0,0621221 | 1 |
| 11 | Akt | 0,13488 | 1,07993 | 1,07993 | 13,6563 | 0,261477 | 0,0351029 | 1 |
| 127 | MAPK_pT202_ | 0,0255476 | 1,14687 | 1,14687 | 18,2811 | 0,389619 | 0,0390732 | 1 |
| 144 | Myt1 | 0,00031202 | 1,14701 | 1,14701 | 27,5113 | 0,17194 | 0,0114579 | 1 |
| 100 | GSK-3a-b_pS2: | 0,00328648 | 1,14954 | 1,14954 | 49,6985 | 0,584682 | 0,0215684 | 1 |
| 105 | HER3_pY1289 | 0,00153558 | 1,15315 | 1,15315 | 12,0771 | 0,118349 | 0,0179656 | 1 |
| 213 | Smad3 | 0,0259735 | 1,15464 | 1,15464 | 6,38273 | 0,233252 | 0,0669977 | 1 |
| 141 | mTOR_pS2448 | 0,302264 | 0,953442 | -1,04883 | 12,5741 | 0,210657 | 0,0307143 | 1 |
| 80 | Elk1_pS383 | 0,000999443 | 1,16458 | 1,16458 | 8,30934 | 0,0843929 | 0,01862 | 1 |
| 228 | TSC1 | 0,000222615 | 1,16694 | 1,16694 | 227,87 | 1,22223 | 0,00983348 | 1 |
| 32 | Bad_pS112 | 0,0334893 | 1,17703 | 1,17703 | 3,55901 | 0,115742 | 0,0596216 | 1 |
| 15 | AMPKa | 0,0357664 | 1,17858 | 1,17858 | 5,72029 | 0,200858 | 0,0643744 | 1 |
| 79 | eIF4G | 0,00179545 | 1,18217 | 1,18217 | 6,89226 | 0,109649 | 0,0291666 | 1 |
| 161 | p53 | 0,0389507 | 1,1829 | 1,1829 | 2,40202 | 0,106289 | 0,0811244 | 1 |
| 37 | Bid | 0,0141726 | 1,18932 | 1,18932 | 10,6969 | 0,434923 | 0,074541 | 1 |
| 138 | MSH6 | 0,00825374 | 1,19423 | 1,19423 | 10,5037 | 0,348983 | 0,0609123 | 1 |
| 186 | PLC-gamma2_ | 0,00103584 | 1,19459 | 1,19459 | 6,24204 | 0,0866953 | 0,025463 | 1 |
| 185 | PKM2 | 0,504394 | 1,04975 | 1,04975 | 10,8879 | 0,454477 | 0,0765257 | 1 |
| 26 | b-Actin | 0,0401365 | 1,19808 | 1,19808 | 7,4584 | 0,493307 | 0,121259 | 1 |
| 203 | RPA32_pS4_S8 | 0,0100323 | 1,20227 | 1,20227 | 14,4553 | 0,444965 | 0,0564341 | 1 |
| 68 | DJ1 | 0,00103 | 1,20467 | 1,20467 | 11,0921 | 0,209944 | 0,0347 | 1 |
| 111 | HSP70 | 0,000450799 | 1,20842 | 1,20842 | 26,2536 | 0,374713 | 0,0261668 | 1 |
| 43 | c-Kit | 0,00152899 | 1,21456 | 1,21456 | 9,67881 | 0,248033 | 0,0469817 | 1 |
| 190 | PREX1 | 0,457057 | 1,03358 | 1,03358 | 10,3415 | 0,233884 | 0,041463 | 1 |
| 51 | CD20 | 0,191735 | 0,947099 | -1,05586 | 10,3099 | 0,151296 | 0,0269041 | 1 |
| 82 | ER-a_pS118 | 0,000421124 | 1,22119 | 1,22119 | 14,7376 | 0,266244 | 0,0331203 | 1 |
| 232 | TUFM | 0,0100379 | 1,22861 | 1,22861 | 3,36369 | 0,152081 | 0,0828898 | 1 |
| 27 | b-Catenin | 0,00685493 | 1,23092 | 1,23092 | 19,371 | 1,03206 | 0,0976768 | 1 |
| 208 | Shc_pY317 | 0,0126903 | 1,26048 | 1,26048 | 6,59602 | 0,258832 | 0,0719412 | 1 |
| 44 | c-Myc | 0,00316716 | 1,28552 | 1,28552 | 18,0174 | 1,00698 | 0,102464 | 1 |
| 49 | Caveolin-1 | 0,0971409 | 1,33265 | 1,33265 | 10,0118 | 1,9585 | 0,358633 | 1 |
| 35 | Bcl-xL | 0,0149623 | 1,29682 | 1,29682 | 4,17741 | 0,376623 | 0,165288 | 1 |
| 148 | NF-kB-p65_pS! | 0,399194 | 0,952911 | -1,04942 | 9,70749 | 0,315754 | 0,0596325 | 1 |
| 164 | p90RSK_pT573 | 0,00639564 | 1,30474 | 1,30474 | 11,5145 | 0,621942 | 0,0990253 | 1 |
| 119 | Jagged1 | 0,000739508 | 1,30593 | 1,30593 | 20,9169 | 0,677713 | 0,0594006 | 1 |

| | | | | | | | | |
|---|---|---|---|---|---|---|---|---|
| 245 | XRCC1 | 0,825676 | 1,01312 | 1,01312 | 9,47025 | 0,333549 | 0,0645713 | 1 |
| 239 | Wee1 | 0,104662 | 0,876613 | -1,14075 | 9,40644 | 0,667283 | 0,130055 | 1 |
| 249 | ZAP-70 | 0,0820368 | 0,930328 | -1,07489 | 9,2011 | 0,145113 | 0,0289141 | 1 |
| 109 | Histone-H3 | 0,0187727 | 1,32418 | 1,32418 | 5,92002 | 0,561613 | 0,173922 | 1 |
| 85 | FAK | 0,000694245 | 1,32953 | 1,32953 | 5,73598 | 0,170349 | 0,054447 | 1 |
| 170 | Paxillin | 5,24E-06 | 1,33213 | 1,33213 | 19,796 | 0,194305 | 0,0179948 | 1 |
| 205 | S6_pS235_S23 | 0,122847 | 0,677394 | -1,47625 | 8,58233 | 5,39909 | 1,15334 | 1 |
| 137 | Mnk1 | 0,397508 | 1,04691 | 1,04691 | 8,53919 | 0,256266 | 0,0550193 | 1 |
| 4 | 14-3-3-zeta | 1,19E-06 | 1,34827 | 1,34827 | 33,826 | 0,267897 | 0,0145197 | 1 |
| 240 | Wee1_pS642 | 0,477296 | 1,03335 | 1,03335 | 8,22266 | 0,209994 | 0,0468206 | 1 |
| 224 | TFAM | 0,352684 | 0,920053 | -1,08689 | 8,10877 | 0,744333 | 0,168288 | 1 |
| 230 | Tuberin | 0,142468 | 1,10623 | 1,10623 | 8,06143 | 0,324571 | 0,0738141 | 1 |
| 53 | cdc25C | 0,258835 | 0,913464 | -1,09473 | 7,90478 | 0,588719 | 0,13654 | 1 |
| 152 | P-Cadherin | 0,767333 | 0,985474 | -1,01474 | 7,73996 | 0,194149 | 0,0459873 | 1 |
| 74 | eEF2K | 0,441313 | 1,09181 | 1,09181 | 7,71876 | 1,10977 | 0,263588 | 1 |
| 47 | Caspase-3 | 0,000118548 | 1,35782 | 1,35782 | 10,1404 | 0,28436 | 0,0514109 | 1 |
| 175 | PDK1 | 2,39E-05 | 1,363 | 1,363 | 9,15835 | 0,134561 | 0,0269365 | 1 |
| 48 | Caspase-7-clea | 0,0880836 | 1,1119 | 1,1119 | 7,46312 | 0,198301 | 0,048713 | 1 |
| 77 | eIF4E | 8,20E-09 | 1,37103 | 1,37103 | 105,413 | 0,35314 | 0,0061418 | 1 |
| 169 | PARP | 0,00558074 | 1,38083 | 1,38083 | 9,62553 | 1,05198 | 0,200366 | 1 |
| 96 | GCN5L2 | 0,000667096 | 1,39302 | 1,39302 | 22,6414 | 1,42422 | 0,115323 | 1 |
| 155 | p21 | 0,707418 | 1,02342 | 1,02342 | 7,13402 | 0,169398 | 0,0435327 | 1 |
| 159 | p38_pT180_Y1 | 1,73E-06 | 1,3992 | 1,3992 | 24,6677 | 0,308891 | 0,0229572 | 1 |
| 87 | FASN | 0,00198794 | 1,42008 | 1,42008 | 14,6017 | 0,886435 | 0,111297 | 1 |
| 162 | p70-S6K1 | 0,809641 | 0,988121 | -1,01202 | 7,05292 | 0,193159 | 0,0502098 | 1 |
| 46 | C-Raf_pS338 | 0,000330295 | 1,4264 | 1,4264 | 17,2098 | 0,742969 | 0,0791471 | 1 |
| 103 | HER2_pY1248 | 0,0149297 | 1,44063 | 1,44063 | 2,57293 | 0,36422 | 0,259524 | 1 |
| 10 | ACC_pS79 | 0,233658 | 1,26526 | 1,26526 | 6,88379 | 2,70112 | 0,719379 | 1 |
| 94 | GATA6 | 0,265404 | 0,941572 | -1,06205 | 6,86886 | 0,135719 | 0,036224 | 1 |
| 9 | ACC1 | 0,00184344 | 1,44582 | 1,44582 | 15,9361 | 1,4716 | 0,169296 | 1 |
| 86 | FAK_pY397 | 0,835995 | 1,03868 | 1,03868 | 6,81998 | 3,40543 | 0,915442 | 1 |
| 165 | PAICS | 0,0196074 | 1,45279 | 1,45279 | 4,40808 | 1,03963 | 0,432387 | 1 |
| 6 | 4E-BP1_pS65 | 0,000634097 | 1,45378 | 1,45378 | 17,6814 | 1,46513 | 0,151915 | 1 |
| 42 | c-Jun_pS73 | 8,11E-08 | 1,50932 | 1,50932 | 116,358 | 1,45267 | 0,0228882 | 1 |
| 41 | c-IAP2 | 0,600646 | 1,01761 | 1,01761 | 6,4537 | 0,0613278 | 0,0174217 | 1 |
| 176 | PDK1_pS241 | 1,68E-07 | 1,54022 | 1,54022 | 35,1936 | 0,390214 | 0,0203273 | 1 |
| 12 | Akt_pS473 | 0,00436292 | 1,55379 | 1,55379 | 18,3478 | 1,69049 | 0,168916 | 1 |
| 177 | PEA-15 | 0,256578 | 1,08761 | 1,08761 | 6,15327 | 0,346978 | 0,10338 | 1 |
| 57 | Chk2_pT68 | 0,0629573 | 1,10963 | 1,10963 | 5,93181 | 0,140716 | 0,0434908 | 1 |
| 195 | Rad50 | 0,000125957 | 1,57946 | 1,57946 | 40,8683 | 3,70815 | 0,166346 | 1 |
| 120 | Jak2 | 0,135485 | 0,940837 | -1,06288 | 5,87721 | 0,0853043 | 0,0266098 | 1 |
| 182 | PKC-b-II_pS66( | 0,000345515 | 1,618 | 1,618 | 7,55104 | 0,363053 | 0,0881463 | 1 |
| 63 | Cox2 | 0,571433 | 1,18839 | 1,18839 | 5,80238 | 23,967 | 7,57267 | 1 |
| 104 | HER3 | 0,000111356 | 1,63288 | 1,63288 | 21,6693 | 2,19332 | 0,185566 | 1 |
| 116 | IR-b | 0,000218778 | 1,64733 | 1,64733 | 14,4859 | 0,963358 | 0,121922 | 1 |
| 237 | VASP | 0,000844433 | 1,68805 | 1,68805 | 10,586 | 0,902805 | 0,156352 | 1 |
| 93 | Gab2 | 0,00149893 | 1,69641 | 1,69641 | 20,7642 | 2,40937 | 0,21273 | 1 |
| 13 | Akt_pT308 | 0,000132405 | 1,81449 | 1,81449 | 19,8553 | 1,27872 | 0,11807 | 1 |
| 121 | JNK2 | 0,537938 | 0,977972 | -1,02252 | 5,46307 | 0,0676374 | 0,0226982 | 1 |
| 136 | MMP2 | 0,378155 | 0,981625 | -1,01872 | 5,4171 | 0,0235161 | 0,00795865 | 1 |
| 83 | ERCC5 | 0,127808 | 1,16538 | 1,16538 | 5,40905 | 0,348821 | 0,118229 | 1 |
| 134 | Merlin | 0,166822 | 1,16384 | 1,16384 | 5,0951 | 0,56156 | 0,202062 | 1 |
| 210 | SLC1A5 | 0,231834 | 1,18357 | 1,18357 | 5,08099 | 0,946591 | 0,341551 | 1 |
| 154 | p16INK4a | 2,05E-05 | 1,83106 | 1,83106 | 61,1803 | 4,11802 | 0,123401 | 1 |
| 70 | DM-K9-Histon | 0,217551 | 1,05094 | 1,05094 | 5,04787 | 0,0706856 | 0,0256723 | 1 |
| 54 | Cdc2_pY15 | 0,283329 | 0,927179 | -1,07854 | 5,02374 | 0,279873 | 0,102135 | 1 |
| 194 | Rab25 | 0,224555 | 0,941539 | -1,06209 | 4,99539 | 0,0956965 | 0,0351211 | 1 |
| 73 | eEF2 | 0,443447 | 0,938933 | -1,06504 | 4,95311 | 0,381503 | 0,141209 | 1 |
| 55 | CDK1 | 6,94E-08 | 1,88604 | 1,88604 | 184,164 | 4,38771 | 0,0436792 | 1 |
| 67 | D-a-Tubulin | 0,0756837 | 0,930459 | -1,07474 | 4,74738 | 0,0633336 | 0,024458 | 1 |
| 201 | Rictor_pT1135 | 0,892292 | 1,01225 | 1,01225 | 4,62353 | 0,3117 | 0,123596 | 1 |
| 242 | WIPI2 | 0,754261 | 1,04946 | 1,04946 | 4,54417 | 0,991671 | 0,400087 | 1 |
| 248 | YB1_pS102 | 0,0852765 | 1,09731 | 1,09731 | 4,53269 | 0,0910193 | 0,0368145 | 1 |
| 5 | 4E-BP1 | 3,26E-08 | 1,94132 | 1,94132 | 83,065 | 1,3759 | 0,0303675 | 1 |
| 84 | Ets-1 | 0,134148 | 0,96709 | -1,03403 | 4,43743 | 0,020748 | 0,00857209 | 1 |
| 204 | RSK | 2,84E-09 | 1,95084 | 1,95084 | 326,695 | 2,16415 | 0,0121447 | 1 |
| 124 | Lck | 0,0587615 | 1,1629 | 1,1629 | 4,23264 | 0,228688 | 0,0990542 | 1 |
| 216 | Src-(phospho-\ | 0,00166818 | 1,9596 | 1,9596 | 5,81883 | 0,827672 | 0,260774 | 1 |
| 233 | Tyro3 | 0,66492 | 1,01211 | 1,01211 | 4,15696 | 0,0302252 | 0,0133301 | 1 |
| 212 | Smad1 | 0,355836 | 0,955667 | -1,04639 | 4,05 | 0,0956214 | 0,0432855 | 1 |

| | | | | | | | | |
|---|---|---|---|---|---|---|---|---|
| 168 | PAR | 0,119847 | 0,632429 | -1,5812 | 3,96584 | 20,7026 | 9,57041 | 1 |
| 217 | Src_pY527 | 4,72E-05 | 2,44926 | 2,44926 | 11,0253 | 1,13465 | 0,188674 | 1 |
| 64 | Creb | 0,172066 | 0,963746 | -1,03762 | 3,71864 | 0,0251136 | 0,0123813 | 1 |
| 130 | MDM2_pS166 | 0,61547 | 1,02953 | 1,02953 | 3,7033 | 0,0906338 | 0,0448687 | 1 |
| 38 | Bim | 0,00855248 | 2,45677 | 2,45677 | 6,77479 | 5,19634 | 1,40619 | 1 |
| 7 | 53BP1 | 0,0210697 | 2,75068 | 2,75068 | 3,72702 | 8,23053 | 4,04863 | 1 |
| 18 | ARID1A | 0,0605014 | 1,97199 | 1,97199 | 3,43223 | 6,68877 | 3,57282 | 1 |
| 95 | GCLM | 0,499991 | 0,981277 | -1,01908 | 3,36925 | 0,0253068 | 0,0137703 | 1 |
| 107 | HES1 | 2,11E-05 | 2,90854 | 2,90854 | 43,8796 | 7,33319 | 0,306388 | 1 |
| 241 | WIPI1 | 0,0946799 | 1,26392 | 1,26392 | 3,21009 | 0,652047 | 0,372394 | 1 |
| 158 | p38-MAPK | 0,543201 | 1,04838 | 1,04838 | 3,14191 | 0,178812 | 0,104338 | 1 |
| 75 | EGFR | 0,0847649 | 1,16266 | 1,16266 | 3,12587 | 0,163053 | 0,0956314 | 1 |
| 202 | RIP | 0,36507 | 0,894917 | -1,11742 | 3,09244 | 0,437652 | 0,25946 | 1 |
| 193 | Rab11 | 0,886034 | 0,991129 | -1,00895 | 3,07432 | 0,107638 | 0,0641887 | 1 |
| 34 | Bax | 0,159603 | 1,41421 | 1,41421 | 3,06105 | 1,45184 | 0,869541 | 1 |
| 147 | NDRG1_pT346 | 0,000376259 | 3,19268 | 3,19268 | 15,1644 | 5,64423 | 0,682373 | 1 |
| 221 | Stathmin-1 | 0,400626 | 1,04742 | 1,04742 | 2,83115 | 0,0664936 | 0,0430585 | 1 |
| 23 | ATR_pS428 | 0,0685999 | 1,1284 | 1,1284 | 2,5898 | 0,0881786 | 0,0624221 | 1 |
| 58 | Claudin-7 | 8,26E-05 | 3,71646 | 3,71646 | 20,2027 | 31,189 | 2,83031 | 1 |
| 113 | IGFBP2 | 0,111664 | 0,856469 | -1,16758 | 2,56926 | 0,224706 | 0,160342 | 1 |
| 72 | E-Cadherin | 1,34E-06 | 6,81231 | 6,81231 | 50,6271 | 154,73 | 5,60315 | 1 |
| 153 | P-Met_pY1234 | 0,00111418 | 13,5079 | 13,5079 | 10,2702 | 6,56734 | 1,17234 | 1 |
| 56 | Chk1_pS296 | 0,494521 | 0,970922 | -1,02995 | 2,38404 | 0,0404177 | 0,0310814 | 1 |
| 66 | Cyclin-D1 | 0,0920015 | 1,09624 | 1,09624 | 2,24121 | 0,0517592 | 0,0423395 | 1 |
| 92 | G6PD | 0,149912 | 0,866254 | -1,1544 | 2,1683 | 0,167204 | 0,141374 | 1 |
| 157 | p27_pT198 | 0,253347 | 1,073 | 1,073 | 2,1435 | 0,0802349 | 0,0686247 | 1 |
| 14 | AMPK-a2_pS34 | 0,256513 | 1,08803 | 1,08803 | 2,12234 | 0,0877857 | 0,0758317 | 1 |
| 89 | FoxM1 | 0,113757 | 0,89613 | -1,11591 | 2,09024 | 0,0810734 | 0,0711088 | 1 |
| 226 | TIGAR | 0,220047 | 1,10031 | 1,10031 | 2,02815 | 0,108713 | 0,0982706 | 1 |
| 122 | JNK_pT183_Y1 | 0,297994 | 1,07002 | 1,07002 | 2,02018 | 0,0760564 | 0,0690219 | 1 |
| 117 | IRF-1 | 0,0981009 | 0,934488 | -1,0701 | 1,94612 | 0,0303863 | 0,0286253 | 1 |
| 33 | Bak | 0,0710451 | 1,13801 | 1,13801 | 1,87467 | 0,0702506 | 0,0687017 | 1 |
| 234 | U-Histone-H2B | 0,95682 | 1,00922 | 1,00922 | 1,55594 | 0,410705 | 0,483925 | 1 |
| 28 | b-Catenin_pT4 | 0,152139 | 1,10792 | 1,10792 | 1,32746 | 0,0577034 | 0,0796934 | 1 |
| 211 | Slfn11 | 0,351537 | 0,944448 | -1,05882 | 1,24644 | 0,0394885 | 0,0580819 | 1 |
| 133 | MERIT40_pS29 | 0,330414 | 0,912678 | -1,09568 | 1,0799 | 0,0862763 | 0,14647 | 1 |
| 198 | RBM15 | 0,4592 | 1,4772 | 1,4772 | 0,951534 | 5,1036 | 9,83319 | 1 |
| 189 | PRAS40_pT246 | 0,188228 | 1,29757 | 1,29757 | 0,904605 | 0,239531 | 0,48545 | 1 |
| 246 | YAP | 0,689856 | 0,982647 | -1,01766 | 0,788196 | 0,0136047 | 0,0316444 | 1 |
| 178 | PEA-15_pS116 | 0,597375 | 1,08254 | 1,08254 | 0,56699 | 0,114289 | 0,369549 | 1 |
| 222 | STING | 0,435835 | 1,76771 | 1,76771 | 0,35209 | 2,14706 | 11,1798 | 1 |

### ANOVA - Subtype B (MGT13) vs. MGT2

| # | Protein | p-value | Ratio (MGT13 vs. 2) | Fold change (MGT13 vs. 2) | F(cell lines) | SS(cell lines) | SS(Error) | F(Error) |
|---|---|---|---|---|---|---|---|---|
| 168 | PAR | 0,00796666 | 0,176996 | -5,64983 | 3,96584 | 20,7026 | 9,57041 | 1 |
| 173 | PDGFR-b | 1,09E-10 | 0,471819 | -2,11946 | 2680,48 | 45,9792 | 0,0314478 | 1 |
| 106 | Heregulin | 5,94E-06 | 0,491576 | -2,03427 | 15,8753 | 1,22152 | 0,141065 | 1 |
| 154 | p16INK4a | 0,0051752 | 0,52318 | -1,91139 | 61,1803 | 4,11802 | 0,123401 | 1 |
| 228 | TSC1 | 0,442482 | 1,02886 | 1,02886 | 227,87 | 1,22223 | 0,00983348 | 1 |
| 184 | PKCa | 0,000105671 | 0,607821 | -1,64522 | 18,1378 | 1,02674 | 0,103781 | 1 |
| 215 | Sox2 | 0,000387963 | 0,670703 | -1,49097 | 19,6576 | 1,20302 | 0,112197 | 1 |
| 102 | Gys_pS641 | 0,649504 | 1,01481 | 1,01481 | 189,262 | 1,40427 | 0,0136028 | 1 |
| 118 | IRS1 | 0,014978 | 0,674798 | -1,48193 | 21,6986 | 2,81443 | 0,237794 | 1 |
| 42 | c-Jun_pS73 | 0,197381 | 0,934333 | -1,07028 | 116,358 | 1,45267 | 0,0228882 | 1 |
| 61 | Connexin-43 | 0,0342814 | 0,684716 | -1,46046 | 15,6721 | 3,01638 | 0,352858 | 1 |
| 148 | NF-kB-p65_pS5 | 0,00054733 | 0,698641 | -1,43135 | 9,70749 | 0,315754 | 0,0596325 | 1 |
| 191 | Progesterone-I | 0,631253 | 1,00755 | 1,00755 | 94,235 | 0,221118 | 0,00430183 | 1 |
| 20 | Atg7 | 0,681356 | 0,981025 | -1,01934 | 90,159 | 1,23961 | 0,0252068 | 1 |
| 143 | Myosin-IIa_pS1 | 0,00254135 | 0,707636 | -1,41316 | 56,6744 | 4,16063 | 0,13459 | 1 |
| 227 | TRIM25 | 0,00573092 | 0,707636 | -1,41316 | 45,9436 | 4,35433 | 0,173755 | 1 |
| 134 | Merlin | 0,0469397 | 0,710885 | -1,4067 | 5,0951 | 0,56156 | 0,202062 | 1 |
| 195 | Rad50 | 0,0318499 | 0,711511 | -1,40546 | 40,8683 | 3,70815 | 0,166346 | 1 |
| 110 | HSP27_pS82 | 0,0152316 | 0,72326 | -1,38263 | 10,1753 | 1,07461 | 0,193617 | 1 |
| 71 | DUSP4 | 4,66E-05 | 0,725994 | -1,37742 | 17,8602 | 0,340866 | 0,0349897 | 1 |
| 44 | c-Myc | 0,016986 | 0,750724 | -1,33205 | 18,0174 | 1,00698 | 0,102464 | 1 |
| 180 | PI3K-p85 | 2,33E-06 | 0,752941 | -1,32813 | 65,5844 | 0,787884 | 0,0220243 | 1 |
| 187 | PLK1 | 0,0031451 | 0,755642 | -1,32338 | 74,7899 | 4,09094 | 0,100282 | 1 |
| 29 | B-Raf | 3,82E-07 | 0,761564 | -1,31309 | 111,25 | 0,617595 | 0,0101776 | 1 |
| 145 | N-Cadherin | 0,129919 | 0,964898 | -1,03638 | 64,9493 | 0,208575 | 0,00588748 | 1 |
| 62 | Cox-IV | 0,00307026 | 0,771809 | -1,29566 | 81,2113 | 3,84541 | 0,0868097 | 1 |
| 73 | eEF2 | 0,027315 | 0,771852 | -1,29559 | 4,95311 | 0,381503 | 0,141209 | 1 |
| 188 | PMS2 | 0,19857 | 0,931209 | -1,07387 | 60,6591 | 1,46813 | 0,0443722 | 1 |
| 123 | LC3A-B | 0,439528 | 1,05926 | 1,05926 | 58,4615 | 1,87329 | 0,0587456 | 1 |
| 142 | Myosin-Heavy- | 0,140632 | 0,884918 | -1,13005 | 58,0716 | 2,47971 | 0,078285 | 1 |
| 238 | VEGFR-2 | 0,0741087 | 0,90161 | -1,10913 | 57,4204 | 0,813718 | 0,0259806 | 1 |
| 229 | TTF1 | 1,07E-05 | 0,802768 | -1,24569 | 15,007 | 0,108667 | 0,0132754 | 1 |
| 31 | B7-H4 | 0,279971 | 0,97317 | -1,02757 | 55,7304 | 0,254248 | 0,00836387 | 1 |
| 72 | E-Cadherin | 0,39156 | 0,357502 | -2,79719 | 50,6271 | 154,73 | 5,60315 | 1 |
| 21 | ATM | 0,00722314 | 0,806507 | -1,23991 | 18,4361 | 0,970152 | 0,0964744 | 1 |
| 166 | PAK1 | 0,854121 | 0,987623 | -1,01253 | 49,2351 | 1,5473 | 0,0576156 | 1 |
| 160 | p44-42-MAPK | 0,248371 | 1,04465 | 1,04465 | 48,2315 | 0,455429 | 0,0173114 | 1 |
| 65 | Cyclin-B1 | 0,00673208 | 0,806749 | -1,23954 | 82,2768 | 3,5958 | 0,0801235 | 1 |
| 8 | A-Raf | 0,0246201 | 0,815429 | -1,22635 | 2,42788 | 0,121051 | 0,0914079 | 1 |
| 111 | HSP70 | 0,00345903 | 0,816935 | -1,22409 | 26,2536 | 0,374713 | 0,0261668 | 1 |
| 99 | Granzyme-B | 0,104485 | 0,907549 | -1,10187 | 41,2121 | 1,35911 | 0,0604605 | 1 |
| 30 | B-Raf_pS445 | 0,00207868 | 0,821963 | -1,2166 | 10,5878 | 0,221288 | 0,0383174 | 1 |
| 81 | ER | 8,49E-05 | 0,829505 | -1,20554 | 12,1807 | 0,0968804 | 0,0145816 | 1 |
| 214 | SOD2 | 0,000123197 | 0,840291 | -1,19006 | 35,5291 | 0,303646 | 0,0156684 | 1 |
| 192 | PTEN | 0,099198 | 0,919403 | -1,08766 | 36,5922 | 1,00525 | 0,0503651 | 1 |
| 162 | p70-S6K1 | 0,0168898 | 0,841885 | -1,18781 | 7,05292 | 0,193159 | 0,0502098 | 1 |
| 236 | ULK1_pS757 | 0,00924834 | 0,84747 | -1,17998 | 23,8444 | 0,644312 | 0,0495395 | 1 |
| 78 | eIF4E_pS209 | 0,000794488 | 0,848195 | -1,17897 | 5,70269 | 0,0556886 | 0,0179031 | 1 |
| 174 | PDHK1 | 0,899871 | 0,995315 | -1,00471 | 28,3303 | 0,29458 | 0,019063 | 1 |
| 135 | MIF | 0,00274436 | 0,851569 | -1,1743 | 12,857 | 0,162293 | 0,023142 | 1 |
| 17 | AR | 0,0135122 | 0,864991 | -1,15608 | 7,24728 | 0,173602 | 0,0439159 | 1 |
| 22 | ATM_pS1981 | 0,19837 | 0,968875 | -1,03213 | 26,8204 | 0,106612 | 0,00728759 | 1 |
| 36 | Beclin | 0,00353933 | 0,865642 | -1,15521 | 10,1533 | 0,129077 | 0,0233067 | 1 |
| 190 | PREX1 | 0,0240693 | 0,86699 | -1,15342 | 10,3415 | 0,233884 | 0,041463 | 1 |
| 131 | MEK1 | 0,491393 | 1,04921 | 1,04921 | 23,975 | 1,28431 | 0,098209 | 1 |
| 212 | Smad1 | 0,0310573 | 0,86727 | -1,15304 | 4,05 | 0,0956214 | 0,0432855 | 1 |
| 225 | TFRC | 0,610392 | 0,947661 | -1,05523 | 23,0032 | 3,23592 | 0,2579 | 1 |
| 96 | GCN5L2 | 0,446808 | 0,922699 | -1,08378 | 22,6414 | 1,42422 | 0,115323 | 1 |
| 108 | Hexokinase-II | 0,101829 | 1,06806 | 1,06806 | 22,6331 | 0,23349 | 0,0189132 | 1 |
| 218 | Stat3 | 0,209103 | 1,12739 | 1,12739 | 22,5764 | 2,24621 | 0,182405 | 1 |
| 240 | Wee1_pS642 | 0,0373854 | 0,874746 | -1,14319 | 8,22266 | 0,209994 | 0,0468206 | 1 |

| | | | | | | | | |
|---|---|---|---|---|---|---|---|---|
| 104 | HER3 | 0,653377 | 0,941659 | -1,06196 | 21,6693 | 2,19332 | 0,185566 | 1 |
| 152 | P-Cadherin | 0,0475988 | 0,875323 | -1,14244 | 7,73996 | 0,194149 | 0,0459873 | 1 |
| 25 | Axl | 0,00590256 | 0,877791 | -1,13922 | 41,9318 | 0,580668 | 0,0253878 | 1 |
| 119 | Jagged1 | 0,482063 | 0,943746 | -1,05961 | 20,9169 | 0,677713 | 0,0594006 | 1 |
| 93 | Gab2 | 0,106578 | 0,659258 | -1,51686 | 20,7642 | 2,40937 | 0,21273 | 1 |
| 58 | Claudin-7 | 0,807363 | 0,869129 | -1,15058 | 20,2027 | 31,189 | 2,83031 | 1 |
| 90 | FoxO3a | 0,00805024 | 0,878312 | -1,13855 | 7,09168 | 0,0949975 | 0,0245586 | 1 |
| 115 | INPP4b | 0,0038102 | 0,882522 | -1,13312 | 7,69294 | 0,0688118 | 0,0163988 | 1 |
| 243 | XBP-1 | 0,000932885 | 0,885771 | -1,12896 | 13,4597 | 0,085545 | 0,011652 | 1 |
| 117 | IRF-1 | 0,0214383 | 0,886596 | -1,12791 | 1,94612 | 0,0303863 | 0,0286253 | 1 |
| 163 | p70-S6K_pT389 | 0,560881 | 0,932876 | -1,07195 | 19,7553 | 2,1438 | 0,198949 | 1 |
| 3 | 14-3-3-beta | 0,00101994 | 0,887888 | -1,12627 | 7,01879 | 0,0383397 | 0,0100145 | 1 |
| 27 | b-Catenin | 0,586225 | 0,95443 | -1,04775 | 19,371 | 1,03206 | 0,0976768 | 1 |
| 125 | LDHA | 0,757713 | 0,975203 | -1,02543 | 19,2759 | 1,35548 | 0,12892 | 1 |
| 98 | Glutaminase | 0,0350438 | 0,890247 | -1,12328 | 80,6271 | 1,4637 | 0,0332823 | 1 |
| 171 | PD-L1 | 0,0229596 | 0,900676 | -1,11028 | 10,7281 | 0,133421 | 0,0228005 | 1 |
| 45 | C-Raf | 0,0168169 | 0,90159 | -1,10915 | 12,349 | 0,127875 | 0,0189843 | 1 |
| 136 | MMP2 | 0,00377979 | 0,91457 | -1,09341 | 5,4171 | 0,0235161 | 0,00795865 | 1 |
| 139 | MSI2 | 0,0677338 | 0,942797 | -1,06067 | 18,0605 | 0,144807 | 0,0146995 | 1 |
| 84 | Ets-1 | 0,00674098 | 0,920921 | -1,08587 | 4,43743 | 0,020748 | 0,00857209 | 1 |
| 183 | PKC-delta_pS6 | 0,0112644 | 0,92882 | -1,07664 | 7,28739 | 0,0321434 | 0,00808651 | 1 |
| 6 | 4E-BP1_pS65 | 0,869373 | 0,981083 | -1,01928 | 17,6814 | 1,46513 | 0,151915 | 1 |
| 179 | PI3K-p110-a | 0,130633 | 0,971456 | -1,02938 | 17,5983 | 0,0447948 | 0,00466657 | 1 |
| 46 | C-Raf_pS338 | 0,821624 | 0,977571 | -1,02294 | 17,2098 | 0,742969 | 0,0791471 | 1 |
| 69 | DM-Histone-H3 | 0,183532 | 0,947886 | -1,05498 | 16,9499 | 0,224844 | 0,0243195 | 1 |
| 200 | Rictor | 0,0472823 | 0,932208 | -1,07272 | 27,7896 | 0,161173 | 0,0106329 | 1 |
| 156 | p27-Kip-1 | 0,0369904 | 0,942593 | -1,0609 | 12,5046 | 0,0442074 | 0,00648135 | 1 |
| 151 | 04-oct | 0,0424524 | 0,943223 | -1,06019 | 8,60402 | 0,0456702 | 0,00973133 | 1 |
| 39 | BRD4 | 0,135088 | 0,931202 | -1,07388 | 15,2571 | 0,215063 | 0,0258425 | 1 |
| 76 | EGFR_pY1173 | 0,00543045 | 1,10886 | 1,10886 | 9,80828 | 0,056272 | 0,0105182 | 1 |
| 97 | Glutamate-D1- | 0,391946 | 0,960342 | -1,0413 | 15,1267 | 0,181095 | 0,0219485 | 1 |
| 219 | Stat3_pY705 | 0,0115838 | 1,11121 | 1,11121 | 62,5686 | 0,749615 | 0,0219646 | 1 |
| 223 | TAZ | 0,079226 | 0,913127 | -1,09514 | 14,9438 | 0,237227 | 0,0291034 | 1 |
| 82 | ER-a_pS118 | 0,735367 | 1,01801 | 1,01801 | 14,7376 | 0,266244 | 0,0331203 | 1 |
| 91 | FoxO3a-(p_S31 | 0,00145173 | 1,11121 | 1,11121 | 5,07456 | 0,0205944 | 0,00744033 | 1 |
| 80 | Elk1_pS383 | 0,0144634 | 1,12557 | 1,12557 | 8,30934 | 0,0843929 | 0,01862 | 1 |
| 203 | RPA32_pS4_S8 | 0,0729682 | 0,84915 | -1,17765 | 14,4553 | 0,444965 | 0,0564341 | 1 |
| 220 | Stat5a | 0,000487859 | 1,12806 | 1,12806 | 212,947 | 2,68086 | 0,0230804 | 1 |
| 231 | Tuberin_pT146 | 0,188121 | 0,945842 | -1,05726 | 14,1568 | 0,169301 | 0,0219248 | 1 |
| 196 | Rad51 | 0,0803365 | 0,905739 | -1,10407 | 13,7869 | 0,317955 | 0,0422807 | 1 |
| 167 | PAK4 | 0,263038 | 0,911669 | -1,09689 | 13,7548 | 0,466078 | 0,0621221 | 1 |
| 11 | Akt | 0,130426 | 1,09454 | 1,09454 | 13,6563 | 0,261477 | 0,0351029 | 1 |
| 101 | Gys | 0,000984417 | 1,13713 | 1,13713 | 324,947 | 2,62375 | 0,014803 | 1 |
| 209 | SHP-2_pY542 | 0,237513 | 0,924868 | -1,08124 | 13,4274 | 0,381753 | 0,0521231 | 1 |
| 60 | Collagen-VI | 0,660698 | 1,03493 | 1,03493 | 13,1673 | 0,713145 | 0,0992937 | 1 |
| 105 | HER3_pY1289 | 0,00653797 | 1,14309 | 1,14309 | 12,0771 | 0,118349 | 0,0179656 | 1 |
| 207 | SDHA | 0,203236 | 1,06751 | 1,06751 | 12,7619 | 0,273363 | 0,0392706 | 1 |
| 141 | mTOR_pS2448 | 0,121926 | 1,08418 | 1,08418 | 12,5741 | 0,210657 | 0,0307143 | 1 |
| 57 | Chk2_pT68 | 0,0324054 | 1,15147 | 1,15147 | 5,93181 | 0,140716 | 0,0434908 | 1 |
| 140 | mTOR | 0,000668745 | 1,15309 | 1,15309 | 21,6487 | 0,141672 | 0,0119976 | 1 |
| 144 | Myt1 | 0,00065574 | 1,15611 | 1,15611 | 27,5113 | 0,17194 | 0,0114579 | 1 |
| 245 | XRCC1 | 0,0419477 | 1,15631 | 1,15631 | 9,47025 | 0,333549 | 0,0645713 | 1 |
| 159 | p38_pT180_Y1 | 0,00783357 | 1,16469 | 1,16469 | 24,6677 | 0,308891 | 0,0229572 | 1 |
| 128 | Mcl-1 | 0,193048 | 1,12527 | 1,12527 | 11,3896 | 1,13117 | 0,18208 | 1 |
| 68 | DJ1 | 0,717612 | 1,02007 | 1,02007 | 11,0921 | 0,209944 | 0,0347 | 1 |
| 235 | UBAC1 | 0,0101172 | 1,17852 | 1,17852 | 4,508 | 0,0808853 | 0,0328948 | 1 |
| 19 | Atg3 | 0,0578352 | 0,87075 | -1,14843 | 11,0157 | 0,46522 | 0,0774264 | 1 |
| 185 | PKM2 | 0,119922 | 1,14191 | 1,14191 | 10,8879 | 0,454477 | 0,0765257 | 1 |
| 248 | YB1_pS102 | 0,00972866 | 1,18759 | 1,18759 | 4,53269 | 0,0910193 | 0,0368145 | 1 |
| 37 | Bid | 0,0663586 | 1,15478 | 1,15478 | 10,6969 | 0,434923 | 0,074541 | 1 |
| 94 | GATA6 | 0,00764717 | 1,18935 | 1,18935 | 6,86886 | 0,135719 | 0,036224 | 1 |
| 237 | VASP | 0,587338 | 0,901056 | -1,10981 | 10,586 | 0,902805 | 0,156352 | 1 |
| 138 | MSH6 | 0,571692 | 0,958856 | -1,04291 | 10,5037 | 0,348983 | 0,0609123 | 1 |
| 47 | Caspase-3 | 0,0232792 | 1,18955 | 1,18955 | 10,1404 | 0,28436 | 0,0514109 | 1 |
| 51 | CD20 | 0,0869841 | 1,08345 | 1,08345 | 10,3099 | 0,151296 | 0,0269041 | 1 |

| | | | | | | | | |
|---|---|---|---|---|---|---|---|---|
| 197 | Raptor | 0,177609 | 0,952498 | -1,04987 | 10,2826 | 0,0973311 | 0,0173536 | 1 |
| 77 | eIF4E | 2,96E-05 | 1,19032 | 1,19032 | 105,413 | 0,35314 | 0,0061418 | 1 |
| 175 | PDK1 | 0,00701433 | 1,20126 | 1,20126 | 9,15835 | 0,134561 | 0,0269365 | 1 |
| 48 | Caspase-7-clea | 0,0100828 | 1,21651 | 1,21651 | 7,46312 | 0,198301 | 0,048713 | 1 |
| 85 | FAK | 0,0181695 | 1,22891 | 1,22891 | 5,73598 | 0,170349 | 0,054447 | 1 |
| 176 | PDK1_pS241 | 0,000784845 | 1,24973 | 1,24973 | 35,1936 | 0,390214 | 0,0203273 | 1 |
| 164 | p90RSK_pT573 | 0,0346426 | 1,25541 | 1,25541 | 11,5145 | 0,621942 | 0,0990253 | 1 |
| 4 | 14-3-3-zeta | 6,25E-05 | 1,2667 | 1,2667 | 33,826 | 0,267897 | 0,0145197 | 1 |
| 43 | c-Kit | 0,813488 | 1,01448 | 1,01448 | 9,67881 | 0,248033 | 0,0469817 | 1 |
| 169 | PARP | 0,909064 | 1,01513 | 1,01513 | 9,62553 | 1,05198 | 0,200366 | 1 |
| 59 | COG3 | 0,000234566 | 1,27332 | 1,27332 | 38,9353 | 0,588651 | 0,0277176 | 1 |
| 239 | Wee1 | 0,629776 | 1,0404 | 1,0404 | 9,40644 | 0,667283 | 0,130055 | 1 |
| 249 | ZAP-70 | 0,411638 | 0,963717 | -1,03765 | 9,2011 | 0,145113 | 0,0289141 | 1 |
| 155 | p21 | 0,00238152 | 1,27851 | 1,27851 | 7,13402 | 0,169398 | 0,0435327 | 1 |
| 206 | S6_pS240_S24 | 0,478707 | 1,1015 | 1,1015 | 8,71822 | 1,99056 | 0,41859 | 1 |
| 172 | Pdcd4 | 0,000320693 | 1,28267 | 1,28267 | 71,9431 | 1,77986 | 0,0453565 | 1 |
| 205 | S6_pS235_S23 | 0,830312 | 1,04947 | 1,04947 | 8,58233 | 5,39909 | 1,15334 | 1 |
| 137 | Mnk1 | 0,903037 | 0,992242 | -1,00782 | 8,53919 | 0,256266 | 0,0550193 | 1 |
| 15 | AMPKa | 0,00648871 | 1,29203 | 1,29203 | 5,72029 | 0,200858 | 0,0643744 | 1 |
| 186 | PLC-gamma2_ | 0,000141087 | 1,2926 | 1,2926 | 6,24204 | 0,0866953 | 0,025463 | 1 |
| 224 | TFAM | 0,879908 | 0,985123 | -1,0151 | 8,10877 | 0,744333 | 0,168288 | 1 |
| 100 | GSK-3a-b_pS21 | 6,08E-05 | 1,2931 | 1,2931 | 49,6985 | 0,584682 | 0,0215684 | 1 |
| 53 | cdc25C | 0,71148 | 0,967785 | -1,03329 | 7,90478 | 0,588719 | 0,13654 | 1 |
| 32 | Bad_pS112 | 0,0042153 | 1,3059 | 1,3059 | 3,55901 | 0,115742 | 0,0596216 | 1 |
| 74 | eEF2K | 0,391829 | 0,880374 | -1,13588 | 7,71876 | 1,10977 | 0,263588 | 1 |
| 5 | 4E-BP1 | 0,00193054 | 1,32775 | 1,32775 | 83,065 | 1,3759 | 0,0303675 | 1 |
| 170 | Paxillin | 1,94E-05 | 1,33734 | 1,33734 | 19,796 | 0,194305 | 0,0179948 | 1 |
| 199 | Rb_pS807_S81 | 0,0340238 | 1,34233 | 1,34233 | 14,168 | 2,7481 | 0,355602 | 1 |
| 26 | b-Actin | 0,0609298 | 0,792451 | -1,26191 | 7,4584 | 0,493307 | 0,121259 | 1 |
| 127 | MAPK_pT202_ | 0,000246362 | 1,35323 | 1,35323 | 18,2811 | 0,389619 | 0,0390732 | 1 |
| 87 | FASN | 0,0118659 | 1,36655 | 1,36655 | 14,6017 | 0,886435 | 0,111297 | 1 |
| 230 | Tuberin | 0,000572089 | 1,37529 | 1,37529 | 8,06143 | 0,324571 | 0,0738141 | 1 |
| 112 | IGF1R_pY1135 | 2,61E-06 | 1,40848 | 1,40848 | 21,4428 | 0,272795 | 0,0233237 | 1 |
| 52 | CD4 | 0,196579 | 0,918402 | -1,08885 | 7,06838 | 0,237332 | 0,0615571 | 1 |
| 9 | ACC1 | 0,00846659 | 1,40896 | 1,40896 | 15,9361 | 1,4716 | 0,169296 | 1 |
| 83 | ERCC5 | 0,00381892 | 1,42802 | 1,42802 | 5,40905 | 0,348821 | 0,118229 | 1 |
| 79 | eIF4G | 0,993821 | 1,00041 | 1,00041 | 6,89226 | 0,109649 | 0,0291666 | 1 |
| 247 | YAP_pS127 | 0,00138756 | 1,46437 | 1,46437 | 37,918 | 2,09533 | 0,101309 | 1 |
| 208 | Shc_pY317 | 0,000596084 | 1,48646 | 1,48646 | 6,59602 | 0,258832 | 0,0719412 | 1 |
| 50 | CD134 | 0,220672 | 0,95183 | -1,05061 | 6,86224 | 0,0886066 | 0,0236724 | 1 |
| 86 | FAK_pY397 | 0,170627 | 0,687624 | -1,45428 | 6,81998 | 3,40543 | 0,915442 | 1 |
| 38 | Bim | 0,172047 | 1,77812 | 1,77812 | 6,77479 | 5,19634 | 1,40619 | 1 |
| 132 | MEK1_pS217_ | 0,495467 | 1,02584 | 1,02584 | 6,74308 | 0,0568754 | 0,0154635 | 1 |
| 103 | HER2_pY1248 | 0,0194902 | 1,48794 | 1,48794 | 2,57293 | 0,36422 | 0,259524 | 1 |
| 41 | c-IAP2 | 0,496687 | 1,02682 | 1,02682 | 6,4537 | 0,0613278 | 0,0174217 | 1 |
| 213 | Smad3 | 0,333724 | 0,929006 | -1,07642 | 6,38273 | 0,233252 | 0,0669977 | 1 |
| 49 | Caveolin-1 | 0,027565 | 1,54377 | 1,54377 | 10,0118 | 1,9585 | 0,358633 | 1 |
| 177 | PEA-15 | 0,194855 | 0,881987 | -1,1338 | 6,15327 | 0,346978 | 0,10338 | 1 |
| 210 | SLC1A5 | 0,00592718 | 1,57596 | 1,57596 | 5,08099 | 0,946591 | 0,341551 | 1 |
| 109 | Histone-H3 | 0,276939 | 1,15727 | 1,15727 | 5,92002 | 0,561613 | 0,173922 | 1 |
| 120 | Jak2 | 0,118718 | 0,927432 | -1,07825 | 5,87721 | 0,0853043 | 0,0266098 | 1 |
| 16 | AMPKa_pT172 | 0,000884095 | 1,58896 | 1,58896 | 20,1532 | 5,16458 | 0,469821 | 1 |
| 63 | Cox2 | 0,0603857 | 0,210873 | -4,7422 | 5,80238 | 23,967 | 7,57267 | 1 |
| 40 | c-Abl | 0,34578 | 1,02808 | 1,02808 | 5,76258 | 0,0348276 | 0,0110802 | 1 |
| 10 | ACC_pS79 | 0,0296819 | 1,61365 | 1,61365 | 6,88379 | 2,70112 | 0,719379 | 1 |
| 182 | PKC-b-II_pS66( | 0,00114756 | 1,61645 | 1,61645 | 7,55104 | 0,363053 | 0,0881463 | 1 |
| 126 | LRP6_pS1490 | 2,60E-08 | 1,62767 | 1,62767 | 66,1155 | 0,789773 | 0,0218998 | 1 |
| 244 | XPF | 0,716109 | 0,989032 | -1,01109 | 5,6978 | 0,036381 | 0,011706 | 1 |
| 121 | JNK2 | 0,306082 | 0,956571 | -1,0454 | 5,46307 | 0,0676374 | 0,0226982 | 1 |
| 55 | CDK1 | 3,87E-06 | 1,69297 | 1,69297 | 184,164 | 4,38771 | 0,0436792 | 1 |
| 217 | Src_pY527 | 0,018099 | 1,72741 | 1,72741 | 11,0253 | 1,13465 | 0,188674 | 1 |
| 116 | IR-b | 8,31E-05 | 1,8479 | 1,8479 | 14,4859 | 0,963358 | 0,121922 | 1 |
| 181 | PKA-a | 3,08E-08 | 1,92621 | 1,92621 | 75,668 | 2,16153 | 0,0523711 | 1 |
| 216 | Src-(phospho-Y | 0,00341385 | 2,00752 | 2,00752 | 5,81883 | 0,827672 | 0,260774 | 1 |
| 70 | DM-K9-Histone | 0,896171 | 0,993927 | -1,00611 | 5,04787 | 0,0706856 | 0,0256723 | 1 |

| | | | | | | | | |
|---|---|---|---|---|---|---|---|---|
| 54 | Cdc2_pY15 | 0,29313 | 0,916766 | -1,09079 | 5,02374 | 0,279873 | 0,102135 | 1 |
| 194 | Rab25 | 0,717302 | 0,980291 | -1,02011 | 4,99539 | 0,0956965 | 0,0351211 | 1 |
| 147 | NDRG1_pT346 | 0,0433171 | 2,15912 | 2,15912 | 15,1644 | 5,64423 | 0,682373 | 1 |
| 150 | Notch3 | 0,645938 | 0,979653 | -1,02077 | 4,82015 | 0,0564768 | 0,0214808 | 1 |
| 67 | D-a-Tubulin | 0,0706427 | 0,917123 | -1,09037 | 4,74738 | 0,0633336 | 0,024458 | 1 |
| 201 | Rictor_pT1135 | 0,169374 | 1,15187 | 1,15187 | 4,62353 | 0,3117 | 0,123596 | 1 |
| 242 | WIPI2 | 0,32701 | 1,18458 | 1,18458 | 4,54417 | 0,991671 | 0,400087 | 1 |
| 12 | Akt_pS473 | 2,65E-05 | 2,2451 | 2,2451 | 18,3478 | 1,69049 | 0,168916 | 1 |
| 13 | Akt_pT308 | 8,16E-06 | 2,29725 | 2,29725 | 19,8553 | 1,27872 | 0,11807 | 1 |
| 204 | RSK | 7,79E-12 | 2,91947 | 2,91947 | 326,695 | 2,16415 | 0,0121447 | 1 |
| 165 | PAICS | 0,480766 | 1,14139 | 1,14139 | 4,40808 | 1,03963 | 0,432387 | 1 |
| 124 | Lck | 0,494363 | 1,06378 | 1,06378 | 4,23264 | 0,228688 | 0,0990542 | 1 |
| 35 | Bcl-xL | 0,982799 | 0,997346 | -1,00266 | 4,17741 | 0,376623 | 0,165288 | 1 |
| 233 | Tyro3 | 0,17434 | 1,04616 | 1,04616 | 4,15696 | 0,0302252 | 0,0133301 | 1 |
| 129 | MCT4 | 1,68E-06 | 2,96479 | 2,96479 | 84,0128 | 9,79881 | 0,21383 | 1 |
| 114 | IGFRb | 3,07E-11 | 2,96917 | 2,96917 | 224,45 | 5,12321 | 0,041847 | 1 |
| 7 | 53BP1 | 0,919947 | 0,921818 | -1,08481 | 3,72702 | 8,23053 | 4,04863 | 1 |
| 64 | Creb | 0,0806068 | 0,944237 | -1,05906 | 3,71864 | 0,0251136 | 0,0123813 | 1 |
| 130 | MDM2_pS166 | 0,989642 | 1,00089 | 1,00089 | 3,7033 | 0,0906338 | 0,0448687 | 1 |
| 24 | Aurora-B | 0,15978 | 1,08809 | 1,08809 | 3,69143 | 0,0699945 | 0,0347624 | 1 |
| 88 | Fibronectin | 2,12E-05 | 3,05017 | 3,05017 | 20,1859 | 16,8349 | 1,52898 | 1 |
| 18 | ARID1A | 0,648131 | 0,745381 | -1,3416 | 3,43223 | 6,68877 | 3,57282 | 1 |
| 95 | GCLM | 0,100177 | 0,943737 | -1,05962 | 3,36925 | 0,0253068 | 0,0137703 | 1 |
| 232 | TUFM | 0,788764 | 1,02361 | 1,02361 | 3,36369 | 0,152081 | 0,0828898 | 1 |
| 241 | WIPI1 | 0,188192 | 1,23647 | 1,23647 | 3,21009 | 0,652047 | 0,372394 | 1 |
| 158 | p38-MAPK | 0,912259 | 0,989848 | -1,01026 | 3,14191 | 0,178812 | 0,104338 | 1 |
| 75 | EGFR | 0,262043 | 0,881459 | -1,13448 | 3,12587 | 0,163053 | 0,0956314 | 1 |
| 202 | RIP | 0,296755 | 1,14227 | 1,14227 | 3,09244 | 0,437652 | 0,25946 | 1 |
| 193 | Rab11 | 0,0727852 | 0,859896 | -1,16293 | 3,07432 | 0,107638 | 0,0641887 | 1 |
| 34 | Bax | 0,495722 | 1,22592 | 1,22592 | 3,06105 | 1,45184 | 0,869541 | 1 |
| 146 | NAPSIN-A | 0,597461 | 1,03956 | 1,03956 | 2,99184 | 0,0810062 | 0,0496389 | 1 |
| 221 | Stathmin-1 | 0,216511 | 1,08305 | 1,08305 | 2,83115 | 0,0664936 | 0,0430585 | 1 |
| 23 | ATR_pS428 | 0,221339 | 1,09631 | 1,09631 | 2,5898 | 0,0881786 | 0,0624221 | 1 |
| 149 | Notch1 | 1,95E-11 | 3,27429 | 3,27429 | 261,231 | 8,7418 | 0,0613504 | 1 |
| 113 | IGFBP2 | 0,540373 | 0,938759 | -1,06524 | 2,56926 | 0,224706 | 0,160342 | 1 |
| 107 | HES1 | 9,00E-07 | 4,09618 | 4,09618 | 43,8796 | 7,33319 | 0,306388 | 1 |
| 161 | p53 | 0,336443 | 1,09161 | 1,09161 | 2,40202 | 0,106289 | 0,0811244 | 1 |
| 56 | Chk1_pS296 | 0,40415 | 0,958307 | -1,04351 | 2,38404 | 0,0404177 | 0,0310814 | 1 |
| 66 | Cyclin-D1 | 0,531901 | 0,960701 | -1,04091 | 2,24121 | 0,0517592 | 0,0423395 | 1 |
| 92 | G6PD | 0,845788 | 0,979908 | -1,0205 | 2,1683 | 0,167204 | 0,141374 | 1 |
| 157 | p27_pT198 | 0,82104 | 0,983616 | -1,01666 | 2,1435 | 0,0802349 | 0,0686247 | 1 |
| 14 | AMPK-a2_pS34 | 0,391724 | 0,92343 | -1,08292 | 2,12234 | 0,0877857 | 0,0758317 | 1 |
| 89 | FoxM1 | 0,795726 | 0,981274 | -1,01908 | 2,09024 | 0,0810734 | 0,0711088 | 1 |
| 226 | TIGAR | 0,950174 | 0,994241 | -1,00579 | 2,02815 | 0,108713 | 0,0982706 | 1 |
| 122 | JNK_pT183_Y1 | 0,85555 | 0,98605 | -1,01415 | 2,02018 | 0,0760564 | 0,0690219 | 1 |
| 153 | P-Met_pY1234 | 7,25E-05 | 21,532 | 21,532 | 10,2702 | 6,56734 | 1,17234 | 1 |
| 33 | Bak | 0,144181 | 1,12682 | 1,12682 | 1,87467 | 0,0702506 | 0,0687017 | 1 |
| 234 | U-Histone-H2B | 0,771409 | 1,0579 | 1,0579 | 1,55594 | 0,410705 | 0,483925 | 1 |
| 28 | b-Catenin_pT4 | 0,881122 | 1,01254 | 1,01254 | 1,32746 | 0,0577034 | 0,0796934 | 1 |
| 211 | Slfn11 | 0,372425 | 0,937999 | -1,0661 | 1,24644 | 0,0394885 | 0,0580819 | 1 |
| 133 | MERIT40_pS29 | 0,816288 | 1,02383 | 1,02383 | 1,0799 | 0,0862763 | 0,14647 | 1 |
| 198 | RBM15 | 0,828227 | 0,838571 | -1,1925 | 0,951534 | 5,1036 | 9,83319 | 1 |
| 189 | PRAS40_pT246 | 0,15625 | 1,37695 | 1,37695 | 0,904605 | 0,239531 | 0,48545 | 1 |
| 246 | YAP | 0,844661 | 1,00992 | 1,00992 | 0,788196 | 0,0136047 | 0,0316444 | 1 |
| 178 | PEA-15_pS116 | 0,478533 | 0,869988 | -1,14944 | 0,56699 | 0,114289 | 0,369549 | 1 |
| 222 | STING | 0,504687 | 1,76455 | 1,76455 | 0,35209 | 2,14706 | 11,1798 | 1 |

### ANOVA - Subtype B (MGT13) vs. subtype A (MGT4, 9, 11)

| # | Protein | p-value | Ratio (Subtype B vs. A) | Fold change (Subtype B vs. A) | F(cell lines) | SS(cell lines) | SS(Error) | F(Error) |
|---|---|---|---|---|---|---|---|---|
| 72 | E-Cadherin | 9,26E-08 | 0,0524788 | -19,0553 | 50,6271 | 154,73 | 5,60315 | 1 |
| 63 | Cox2 | 0,00455839 | 0,177443 | -5,6356 | 5,80238 | 23,967 | 7,57267 | 1 |
| 58 | Claudin-7 | 1,29E-05 | 0,23386 | -4,27607 | 20,2027 | 31,189 | 2,83031 | 1 |
| 168 | PAR | 0,0322396 | 0,279868 | -3,57312 | 3,96584 | 20,7026 | 9,57041 | 1 |
| 154 | p16INK4a | 4,85E-08 | 0,285726 | -3,49986 | 61,1803 | 4,11802 | 0,123401 | 1 |
| 7 | 53BP1 | 0,00715569 | 0,335124 | -2,98397 | 3,72702 | 8,23053 | 4,04863 | 1 |
| 18 | ARID1A | 0,0102002 | 0,377984 | -2,64561 | 3,43223 | 6,68877 | 3,57282 | 1 |
| 93 | Gab2 | 1,49E-05 | 0,388619 | -2,57321 | 20,7642 | 2,40937 | 0,21273 | 1 |
| 195 | Rad50 | 6,55E-07 | 0,450477 | -2,21987 | 40,8683 | 3,70815 | 0,166346 | 1 |
| 237 | VASP | 7,86E-05 | 0,533786 | -1,87341 | 10,586 | 0,902805 | 0,156352 | 1 |
| 29 | B-Raf | 0,932125 | 0,998138 | -1,00187 | 111,25 | 0,617595 | 0,0101776 | 1 |
| 104 | HER3 | 1,22E-05 | 0,576685 | -1,73405 | 21,6693 | 2,19332 | 0,185566 | 1 |
| 44 | c-Myc | 4,85E-06 | 0,583984 | -1,71238 | 18,0174 | 1,00698 | 0,102464 | 1 |
| 20 | Atg7 | 0,958228 | 1,0018 | 1,0018 | 90,159 | 1,23961 | 0,0252068 | 1 |
| 61 | Connexin-43 | 0,00034275 | 0,584679 | -1,71034 | 15,6721 | 3,01638 | 0,352858 | 1 |
| 118 | IRS1 | 0,000128719 | 0,587368 | -1,70251 | 21,6986 | 2,81443 | 0,237794 | 1 |
| 65 | Cyclin-B1 | 0,120747 | 1,09697 | 1,09697 | 82,2768 | 3,5958 | 0,0801235 | 1 |
| 62 | Cox-IV | 0,140669 | 1,09986 | 1,09986 | 81,2113 | 3,84541 | 0,0868097 | 1 |
| 134 | Merlin | 0,000554037 | 0,610809 | -1,63717 | 5,0951 | 0,56156 | 0,202062 | 1 |
| 42 | c-Jun_pS73 | 4,30E-09 | 0,619042 | -1,6154 | 116,358 | 1,45267 | 0,0228882 | 1 |
| 187 | PLK1 | 0,102764 | 1,12577 | 1,12577 | 74,7899 | 4,09094 | 0,100282 | 1 |
| 26 | b-Actin | 0,000163746 | 0,661434 | -1,51187 | 7,4584 | 0,493307 | 0,121259 | 1 |
| 86 | FAK_pY397 | 0,045405 | 0,662017 | -1,51054 | 6,81998 | 3,40543 | 0,915442 | 1 |
| 96 | GCN5L2 | 4,01E-05 | 0,662375 | -1,50972 | 22,6414 | 1,42422 | 0,115323 | 1 |
| 145 | N-Cadherin | 0,797114 | 1,00429 | 1,00429 | 64,9493 | 0,208575 | 0,00588748 | 1 |
| 6 | 4E-BP1_pS65 | 0,000126524 | 0,674848 | -1,48182 | 17,6814 | 1,46513 | 0,151915 | 1 |
| 111 | HSP70 | 3,24E-07 | 0,676037 | -1,47921 | 26,2536 | 0,374713 | 0,0261668 | 1 |
| 188 | PMS2 | 0,489107 | 0,972554 | -1,02822 | 60,6591 | 1,46813 | 0,0443722 | 1 |
| 123 | LC3A-B | 0,666452 | 0,977938 | -1,02256 | 58,4615 | 1,87329 | 0,0587456 | 1 |
| 142 | Myosin-Heavy-Chain-11 | 0,941442 | 1,00452 | 1,00452 | 58,0716 | 2,47971 | 0,078285 | 1 |
| 147 | NDRG1_pT346 | 0,0176701 | 0,676271 | -1,4787 | 15,1644 | 5,64423 | 0,682373 | 1 |
| 143 | Myosin-IIa_pS1943 | 0,677581 | 1,0343 | 1,0343 | 56,6744 | 4,16063 | 0,13459 | 1 |
| 5 | 4E-BP1 | 5,14E-07 | 0,683945 | -1,46211 | 83,065 | 1,3759 | 0,0303675 | 1 |
| 46 | C-Raf_pS338 | 5,62E-05 | 0,685341 | -1,45913 | 17,2098 | 0,742969 | 0,0791471 | 1 |
| 217 | Src_pY527 | 0,00310425 | 0,705281 | -1,41787 | 11,0253 | 1,13465 | 0,188674 | 1 |
| 166 | PAK1 | 0,355714 | 0,955222 | -1,04688 | 49,2351 | 1,5473 | 0,0576156 | 1 |
| 203 | RPA32_pS4_S8 | 5,41E-05 | 0,706291 | -1,41585 | 14,4553 | 0,444965 | 0,0564341 | 1 |
| 227 | TRIM25 | 0,351887 | 1,09383 | 1,09383 | 45,9436 | 4,35433 | 0,173755 | 1 |
| 119 | Jagged1 | 4,98E-05 | 0,72266 | -1,38378 | 20,9169 | 0,677713 | 0,0594006 | 1 |
| 148 | NF-kB-p65_pS536 | 0,000171846 | 0,733165 | -1,36395 | 9,70749 | 0,315754 | 0,0596325 | 1 |
| 169 | PARP | 0,00262143 | 0,735164 | -1,36024 | 9,62553 | 1,05198 | 0,200366 | 1 |
| 75 | EGFR | 0,00274615 | 0,758143 | -1,31901 | 3,12587 | 0,163053 | 0,0956314 | 1 |
| 35 | Bcl-xL | 0,00584901 | 0,769068 | -1,30028 | 4,17741 | 0,376623 | 0,165288 | 1 |
| 27 | b-Catenin | 0,000695335 | 0,775379 | -1,28969 | 19,371 | 1,03206 | 0,0976768 | 1 |
| 215 | Sox2 | 0,00361076 | 0,791867 | -1,26284 | 19,6576 | 1,20302 | 0,112197 | 1 |
| 238 | VEGFR-2 | 5,44E-05 | 0,795888 | -1,25646 | 57,4204 | 0,813718 | 0,0259806 | 1 |
| 138 | MSH6 | 0,000808125 | 0,802907 | -1,24547 | 10,5037 | 0,348983 | 0,0609123 | 1 |
| 213 | Smad3 | 0,00106329 | 0,804585 | -1,24288 | 6,38273 | 0,233252 | 0,0669977 | 1 |
| 167 | PAK4 | 0,00195353 | 0,804994 | -1,24224 | 13,7548 | 0,466078 | 0,0621221 | 1 |
| 177 | PEA-15 | 0,0071599 | 0,810941 | -1,23314 | 6,15327 | 0,346978 | 0,10338 | 1 |
| 144 | Myt1 | 0,715015 | 1,00793 | 1,00793 | 27,5113 | 0,17194 | 0,0114579 | 1 |
| 176 | PDK1_pS241 | 1,54E-05 | 0,811401 | -1,23244 | 35,1936 | 0,390214 | 0,0203273 | 1 |
| 71 | DUSP4 | 0,000312223 | 0,819547 | -1,22019 | 17,8602 | 0,340866 | 0,0349897 | 1 |
| 223 | TAZ | 8,36E-05 | 0,821571 | -1,21718 | 14,9438 | 0,237227 | 0,0291034 | 1 |
| 73 | eEF2 | 0,0269716 | 0,822052 | -1,21647 | 4,95311 | 0,381503 | 0,141209 | 1 |
| 159 | p38_pT180_Y182 | 5,65E-05 | 0,832394 | -1,20135 | 24,6677 | 0,308891 | 0,0229572 | 1 |
| 225 | TFRC | 0,135824 | 1,14127 | 1,14127 | 23,0032 | 3,23592 | 0,2579 | 1 |
| 232 | TUFM | 0,0075522 | 0,833148 | -1,20027 | 3,36369 | 0,152081 | 0,0828898 | 1 |
| 82 | ER-a_pS118 | 0,000232118 | 0,833625 | -1,19958 | 14,7376 | 0,266244 | 0,0331203 | 1 |
| 43 | c-Kit | 0,00079813 | 0,835262 | -1,19723 | 9,67881 | 0,248033 | 0,0469817 | 1 |
| 190 | PREX1 | 0,000925702 | 0,83882 | -1,19215 | 10,3415 | 0,233884 | 0,041463 | 1 |
| 200 | Rictor | 6,88E-06 | 0,840881 | -1,18923 | 27,7896 | 0,161173 | 0,0106329 | 1 |
| 79 | eIF4G | 0,000568829 | 0,846252 | -1,18168 | 6,89226 | 0,109649 | 0,0291666 | 1 |
| 240 | Wee1_pS642 | 0,00175013 | 0,846517 | -1,18131 | 8,22266 | 0,209994 | 0,0468206 | 1 |
| 68 | DJ1 | 0,000678414 | 0,846762 | -1,18097 | 11,0921 | 0,209944 | 0,0347 | 1 |
| 14 | AMPK-a2_pS345 | 0,0236485 | 0,848719 | -1,17825 | 2,12234 | 0,0877857 | 0,0758317 | 1 |
| 162 | p70-S6K1 | 0,00445748 | 0,852006 | -1,1737 | 7,05292 | 0,193159 | 0,0502098 | 1 |
| 193 | Rab11 | 0,0272591 | 0,867593 | -1,15261 | 3,07432 | 0,107638 | 0,0641887 | 1 |
| 77 | eIF4E | 2,50E-06 | 0,868193 | -1,15182 | 105,413 | 0,35314 | 0,0061418 | 1 |

| | | | | | | | | |
|---|---|---|---|---|---|---|---|---|
| 150 | Notch3 | 0,000778652 | 0,87155 | -1,14738 | 4,82015 | 0,0564768 | 0,0214808 | 1 |
| 170 | Paxillin | 0,883219 | 1,0039 | 1,0039 | 19,796 | 0,194305 | 0,0179948 | 1 |
| 163 | p70-S6K_pT389 | 0,0750696 | 1,20811 | 1,20811 | 19,7553 | 2,1438 | 0,198949 | 1 |
| 47 | Caspase-3 | 0,00844546 | 0,876076 | -1,14145 | 10,1404 | 0,28436 | 0,0514109 | 1 |
| 97 | Glutamate-D1-2 | 0,00156385 | 0,876338 | -1,14111 | 15,1267 | 0,181095 | 0,0219485 | 1 |
| 66 | Cyclin-D1 | 0,0110868 | 0,876358 | -1,14109 | 2,24121 | 0,0517592 | 0,0423395 | 1 |
| 21 | ATM | 0,416287 | 1,04712 | 1,04712 | 18,4361 | 0,970152 | 0,0964744 | 1 |
| 175 | PDK1 | 0,00390704 | 0,881331 | -1,13465 | 9,15835 | 0,134561 | 0,0269365 | 1 |
| 228 | TSC1 | 0,000285401 | 0,881672 | -1,13421 | 227,87 | 1,22223 | 0,00983348 | 1 |
| 184 | PKCa | 0,93687 | 0,993551 | -1,00649 | 18,1378 | 1,02674 | 0,103781 | 1 |
| 156 | p27-Kip-1 | 2,47E-05 | 0,884919 | -1,13005 | 12,5046 | 0,0442074 | 0,00648135 | 1 |
| 152 | P-Cadherin | 0,0207777 | 0,888225 | -1,12584 | 7,73996 | 0,194149 | 0,0459873 | 1 |
| 55 | CDK1 | 0,00810602 | 0,89763 | -1,11404 | 184,164 | 4,38771 | 0,0436792 | 1 |
| 196 | Rad51 | 0,0171023 | 0,90032 | -1,11072 | 13,7869 | 0,317955 | 0,0422807 | 1 |
| 212 | Smad1 | 0,0456033 | 0,907502 | -1,10193 | 4,05 | 0,0956214 | 0,0432855 | 1 |
| 132 | MEK1_pS217_S221 | 0,00533167 | 0,917191 | -1,09029 | 6,74308 | 0,0568754 | 0,0154635 | 1 |
| 78 | eIF4E_pS209 | 0,0168423 | 0,925611 | -1,08037 | 5,70269 | 0,0556886 | 0,0179031 | 1 |
| 9 | ACC1 | 0,700552 | 0,974509 | -1,02616 | 15,9361 | 1,4716 | 0,169296 | 1 |
| 106 | Heregulin | 0,405603 | 0,92521 | -1,08084 | 15,8753 | 1,22152 | 0,141065 | 1 |
| 136 | MMP2 | 0,00235366 | 0,93169 | -1,07332 | 5,4171 | 0,0235161 | 0,00795865 | 1 |
| 39 | BRD4 | 0,41524 | 1,02915 | 1,02915 | 15,2571 | 0,215063 | 0,0258425 | 1 |
| 4 | 14-3-3-zeta | 0,0239199 | 0,9395 | -1,0644 | 33,826 | 0,267897 | 0,0145197 | 1 |
| 84 | Ets-1 | 0,0221805 | 0,95226 | -1,05013 | 4,43743 | 0,020748 | 0,00857209 | 1 |
| 229 | TTF1 | 0,268852 | 0,973228 | -1,02751 | 15,007 | 0,108667 | 0,0132754 | 1 |
| 91 | FoxO3a-(p_S318-S321) | 0,0462245 | 1,04054 | 1,04054 | 5,07456 | 0,0205944 | 0,00744033 | 1 |
| 22 | ATM_pS1981 | 0,0380081 | 1,0421 | 1,0421 | 26,8204 | 0,106612 | 0,00728759 | 1 |
| 87 | FASN | 0,559426 | 0,962304 | -1,03917 | 14,6017 | 0,886435 | 0,111297 | 1 |
| 116 | IR-b | 0,0758736 | 1,12175 | 1,12175 | 14,4859 | 0,963358 | 0,121922 | 1 |
| 243 | XBP-1 | 0,0368888 | 1,05254 | 1,05254 | 13,4597 | 0,085545 | 0,011652 | 1 |
| 179 | PI3K-p110-a | 0,00019083 | 1,07763 | 1,07763 | 17,5983 | 0,0447948 | 0,00466657 | 1 |
| 231 | Tuberin_pT1462 | 0,647424 | 1,01421 | 1,01421 | 14,1568 | 0,169301 | 0,0219248 | 1 |
| 186 | PLC-gamma2_pY759 | 0,0243459 | 1,08204 | 1,08204 | 6,24204 | 0,0866953 | 0,025463 | 1 |
| 244 | XPF | 0,00427448 | 1,08439 | 1,08439 | 5,6978 | 0,036381 | 0,011706 | 1 |
| 11 | Akt | 0,735998 | 1,01353 | 1,01353 | 13,6563 | 0,261477 | 0,0351029 | 1 |
| 31 | B7-H4 | 0,000425646 | 1,09645 | 1,09645 | 55,7304 | 0,254248 | 0,00836387 | 1 |
| 209 | SHP-2_pY542 | 0,307113 | 1,05359 | 1,05359 | 13,4274 | 0,381753 | 0,0521231 | 1 |
| 50 | CD134 | 0,0102322 | 1,09653 | 1,09653 | 6,86224 | 0,0886066 | 0,0236724 | 1 |
| 135 | MIF | 0,703097 | 0,987199 | -1,01297 | 12,857 | 0,162293 | 0,023142 | 1 |
| 69 | DM-Histone-H3 | 0,00886289 | 1,09863 | 1,09863 | 16,9499 | 0,224844 | 0,0243195 | 1 |
| 214 | SOD2 | 0,00295949 | 1,10036 | 1,10036 | 35,5291 | 0,303646 | 0,0156684 | 1 |
| 108 | Hexokinase-II | 0,00411948 | 1,1038 | 1,1038 | 22,6331 | 0,23349 | 0,0189132 | 1 |
| 45 | C-Raf | 0,975277 | 1,0009 | 1,0009 | 12,349 | 0,127875 | 0,0189843 | 1 |
| 81 | ER | 0,151447 | 0,963075 | -1,03834 | 12,1807 | 0,0968804 | 0,0145816 | 1 |
| 105 | HER3_pY1289 | 0,753547 | 0,991278 | -1,0088 | 12,0771 | 0,118349 | 0,0179656 | 1 |
| 164 | p90RSK_pT573 | 0,536997 | 0,962192 | -1,03929 | 11,5145 | 0,621942 | 0,0990253 | 1 |
| 40 | c-Abl | 0,00049692 | 1,10932 | 1,10932 | 5,76258 | 0,0348276 | 0,0110802 | 1 |
| 17 | AR | 0,0266135 | 1,1102 | 1,1102 | 7,24728 | 0,173602 | 0,0439159 | 1 |
| 140 | mTOR | 0,000486706 | 1,11241 | 1,11241 | 21,6487 | 0,141672 | 0,0119976 | 1 |
| 100 | GSK-3a-b_pS21_S9 | 0,00147089 | 1,12488 | 1,12488 | 49,6985 | 0,584682 | 0,0215684 | 1 |
| 185 | PKM2 | 0,161928 | 1,0878 | 1,0878 | 10,8879 | 0,454477 | 0,0765257 | 1 |
| 171 | PD-L1 | 0,155913 | 1,04872 | 1,04872 | 10,7281 | 0,133421 | 0,0228005 | 1 |
| 37 | Bid | 0,546167 | 0,970957 | -1,02991 | 10,6969 | 0,434923 | 0,074541 | 1 |
| 30 | B-Raf_pS445 | 0,442357 | 1,03252 | 1,03252 | 10,5878 | 0,221288 | 0,0383174 | 1 |
| 236 | ULK1_pS757 | 0,0179527 | 1,13116 | 1,13116 | 23,8444 | 0,644312 | 0,0495395 | 1 |
| 141 | mTOR_pS2448 | 0,00444242 | 1,13712 | 1,13712 | 12,5741 | 0,210657 | 0,0307143 | 1 |
| 98 | Glutaminase | 0,00708487 | 1,14106 | 1,14106 | 80,6271 | 1,4637 | 0,0332823 | 1 |
| 245 | XRCC1 | 0,014799 | 1,14133 | 1,14133 | 9,47025 | 0,333549 | 0,0645713 | 1 |
| 197 | Raptor | 0,35991 | 1,02476 | 1,02476 | 10,2826 | 0,0973311 | 0,0173536 | 1 |
| 51 | CD20 | 0,00147571 | 1,14397 | 1,14397 | 10,3099 | 0,151296 | 0,0269041 | 1 |
| 110 | HSP27_pS82 | 0,471567 | 0,932331 | -1,07258 | 10,1753 | 1,07461 | 0,193617 | 1 |
| 36 | Beclin | 0,601858 | 1,01676 | 1,01676 | 10,1533 | 0,129077 | 0,0233067 | 1 |
| 180 | PI3K-p85 | 0,000227896 | 1,16881 | 1,16881 | 65,5844 | 0,787884 | 0,0220243 | 1 |
| 49 | Caveolin-1 | 0,204318 | 1,15842 | 1,15842 | 10,0118 | 1,9585 | 0,358633 | 1 |
| 76 | EGFR_pY1173 | 0,308719 | 0,978301 | -1,02218 | 9,80828 | 0,056272 | 0,0105182 | 1 |
| 208 | Shc_pY317 | 0,0115756 | 1,17928 | 1,17928 | 6,59602 | 0,258832 | 0,0719412 | 1 |
| 127 | MAPK_pT202_Y204 | 0,00135986 | 1,17994 | 1,17994 | 18,2811 | 0,389619 | 0,0390732 | 1 |
| 131 | MEK1 | 0,00869448 | 1,18093 | 1,18093 | 23,975 | 1,28431 | 0,098209 | 1 |
| 160 | p44-42-MAPK | 8,92E-05 | 1,18158 | 1,18158 | 48,2315 | 0,455429 | 0,0173114 | 1 |
| 139 | MSI2 | 2,18E-05 | 1,18178 | 1,18178 | 18,0605 | 0,144807 | 0,0146995 | 1 |
| 249 | ZAP-70 | 0,30524 | 1,03589 | 1,03589 | 9,2011 | 0,145113 | 0,0289141 | 1 |
| 174 | PDHK1 | 0,000121269 | 1,18278 | 1,18278 | 28,3303 | 0,29458 | 0,019063 | 1 |
| 239 | Wee1 | 0,0188 | 1,18684 | 1,18684 | 9,40644 | 0,667283 | 0,130055 | 1 |
| 151 | 04-oct | 0,194053 | 1,02722 | 1,02722 | 8,60402 | 0,0456702 | 0,00973133 | 1 |

| | | | | | | | | |
|---|---|---|---|---|---|---|---|---|
| 191 | Progesterone-Receptor | 1,02E-08 | 1,20182 | 1,20182 | 94,235 | 0,221118 | 0,00430183 | 1 |
| 137 | Mnk1 | 0,254237 | 0,947779 | -1,0551 | 8,53919 | 0,256266 | 0,0550193 | 1 |
| 80 | Elk1_pS383 | 0,243206 | 0,966507 | -1,03465 | 8,30934 | 0,0843929 | 0,01862 | 1 |
| 83 | ERCC5 | 0,010715 | 1,22537 | 1,22537 | 5,40905 | 0,348821 | 0,118229 | 1 |
| 224 | TFAM | 0,374221 | 1,07073 | 1,07073 | 8,10877 | 0,744333 | 0,168288 | 1 |
| 25 | Axl | 6,31E-05 | 1,22942 | 1,22942 | 41,9318 | 0,580668 | 0,0253878 | 1 |
| 53 | cdc25C | 0,399524 | 1,05947 | 1,05947 | 7,90478 | 0,588719 | 0,13654 | 1 |
| 102 | Gys_pS641 | 3,62E-06 | 1,24086 | 1,24086 | 189,262 | 1,40427 | 0,0136028 | 1 |
| 74 | eEF2K | 0,0539679 | 0,806347 | -1,24016 | 7,71876 | 1,10977 | 0,263588 | 1 |
| 115 | INPP4b | 0,800602 | 1,00694 | 1,00694 | 7,69294 | 0,0688118 | 0,0163988 | 1 |
| 182 | PKC-b-II_pS660 | 0,988281 | 0,99904 | -1,00096 | 7,55104 | 0,363053 | 0,0881463 | 1 |
| 48 | Caspase-7-cleaved | 0,064765 | 1,09409 | 1,09409 | 7,46312 | 0,198301 | 0,048713 | 1 |
| 230 | Tuberin | 0,000658781 | 1,24322 | 1,24322 | 8,06143 | 0,324571 | 0,0738141 | 1 |
| 183 | PKC-delta_pS664 | 0,0775669 | 1,03718 | 1,03718 | 7,28739 | 0,0321434 | 0,00808651 | 1 |
| 155 | p21 | 0,000457045 | 1,24925 | 1,24925 | 7,13402 | 0,169398 | 0,0435327 | 1 |
| 60 | Collagen-VI | 0,00309077 | 1,25961 | 1,25961 | 13,1673 | 0,713145 | 0,0992937 | 1 |
| 90 | FoxO3a | 0,326212 | 1,03329 | 1,03329 | 7,09168 | 0,0949975 | 0,0245586 | 1 |
| 52 | CD4 | 0,258857 | 1,05954 | 1,05954 | 7,06838 | 0,237332 | 0,0615571 | 1 |
| 94 | GATA6 | 0,000113335 | 1,26315 | 1,26315 | 6,86886 | 0,135719 | 0,036224 | 1 |
| 3 | 14-3-3-beta | 0,994291 | 0,999848 | -1,00015 | 7,01879 | 0,0383397 | 0,0100145 | 1 |
| 13 | Akt_pT308 | 0,00215215 | 1,26606 | 1,26606 | 19,8553 | 1,27872 | 0,11807 | 1 |
| 10 | ACC_pS79 | 0,0783184 | 1,27535 | 1,27535 | 6,88379 | 2,70112 | 0,719379 | 1 |
| 112 | IGF1R_pY1135_Y1136 | 2,51E-06 | 1,2702 | 1,2702 | 21,4428 | 0,272795 | 0,0233237 | 1 |
| 202 | RIP | 0,0243554 | 1,2764 | 1,2764 | 3,09244 | 0,437652 | 0,25946 | 1 |
| 19 | Atg3 | 0,00106264 | 1,29079 | 1,29079 | 11,0157 | 0,46522 | 0,0774264 | 1 |
| 38 | Bim | 0,108906 | 0,723765 | -1,38166 | 6,77479 | 5,19634 | 1,40619 | 1 |
| 199 | Rb_pS807_S811 | 0,0130711 | 1,29452 | 1,29452 | 14,168 | 2,7481 | 0,355602 | 1 |
| 125 | LDHA | 0,00151046 | 1,32629 | 1,32629 | 19,2759 | 1,35548 | 0,12892 | 1 |
| 41 | c-IAP2 | 0,747176 | 1,00905 | 1,00905 | 6,4537 | 0,0613278 | 0,0174217 | 1 |
| 210 | SLC1A5 | 0,00888782 | 1,33153 | 1,33153 | 5,08099 | 0,946591 | 0,341551 | 1 |
| 207 | SDHA | 8,99E-06 | 1,3588 | 1,3588 | 12,7619 | 0,273363 | 0,0392706 | 1 |
| 247 | YAP_pS127 | 0,00044078 | 1,36972 | 1,36972 | 37,918 | 2,09533 | 0,101309 | 1 |
| 57 | Chk2_pT68 | 0,374469 | 1,03771 | 1,03771 | 5,93181 | 0,140716 | 0,0434908 | 1 |
| 109 | Histone-H3 | 0,124636 | 0,873953 | -1,14423 | 5,92002 | 0,561613 | 0,173922 | 1 |
| 120 | Jak2 | 0,676892 | 0,985752 | -1,01445 | 5,87721 | 0,0853043 | 0,0266098 | 1 |
| 216 | Src-(phospho-Y416) | 0,813303 | 1,02445 | 1,02445 | 5,81883 | 0,827672 | 0,260774 | 1 |
| 59 | COG3 | 1,30E-06 | 1,38309 | 1,38309 | 38,9353 | 0,588651 | 0,0277176 | 1 |
| 107 | HES1 | 0,000318396 | 1,40833 | 1,40833 | 43,8796 | 7,33319 | 0,306388 | 1 |
| 85 | FAK | 0,123423 | 0,924317 | -1,08188 | 5,73598 | 0,170349 | 0,054447 | 1 |
| 15 | AMPKa | 0,102413 | 1,09626 | 1,09626 | 5,72029 | 0,200858 | 0,0643744 | 1 |
| 219 | Stat3_pY705 | 1,10E-07 | 1,41198 | 1,41198 | 62,5686 | 0,749615 | 0,0219646 | 1 |
| 12 | Akt_pS473 | 0,000280348 | 1,44492 | 1,44492 | 18,3478 | 1,69049 | 0,168916 | 1 |
| 121 | JNK2 | 0,484001 | 0,978116 | -1,02237 | 5,46307 | 0,0676374 | 0,0226982 | 1 |
| 99 | Granzyme-B | 1,48E-05 | 1,44578 | 1,44578 | 41,2121 | 1,35911 | 0,0604605 | 1 |
| 204 | RSK | 4,24E-10 | 1,49652 | 1,49652 | 326,695 | 2,16415 | 0,0121447 | 1 |
| 192 | PTEN | 1,28E-06 | 1,5068 | 1,5068 | 36,5922 | 1,00525 | 0,0503651 | 1 |
| 205 | S6_pS235_S236 | 0,0450751 | 1,54927 | 1,54927 | 8,58233 | 5,39909 | 1,15334 | 1 |
| 101 | Gys | 1,39E-09 | 1,56809 | 1,56809 | 324,947 | 2,62375 | 0,014803 | 1 |
| 70 | DM-K9-Histone-H3 | 0,114037 | 0,945754 | -1,05736 | 5,04787 | 0,0706856 | 0,0256723 | 1 |
| 54 | Cdc2_pY15 | 0,853437 | 0,98877 | -1,01136 | 5,02374 | 0,279873 | 0,102135 | 1 |
| 194 | Rab25 | 0,338673 | 1,04116 | 1,04116 | 4,99539 | 0,0956965 | 0,0351211 | 1 |
| 128 | Mcl-1 | 6,57E-05 | 1,57347 | 1,57347 | 11,3896 | 1,13117 | 0,18208 | 1 |
| 153 | P-Met_pY1234_Y1235 | 0,00722186 | 1,59402 | 1,59402 | 10,2702 | 6,56734 | 1,17234 | 1 |
| 67 | D-a-Tubulin | 0,667794 | 0,985668 | -1,01454 | 4,74738 | 0,0633336 | 0,024458 | 1 |
| 201 | Rictor_pT1135 | 0,0911178 | 1,13793 | 1,13793 | 4,62353 | 0,3117 | 0,123596 | 1 |
| 242 | WIPI2 | 0,32589 | 1,12876 | 1,12876 | 4,54417 | 0,991671 | 0,400087 | 1 |
| 248 | YB1_pS102 | 0,0642147 | 1,08227 | 1,08227 | 4,53269 | 0,0910193 | 0,0368145 | 1 |
| 235 | UBAC1 | 0,428827 | 1,03021 | 1,03021 | 4,508 | 0,0808853 | 0,0328948 | 1 |
| 206 | S6_pS240_S244 | 0,00161609 | 1,61514 | 1,61514 | 8,71822 | 1,99056 | 0,41859 | 1 |
| 165 | PAICS | 0,0500058 | 0,785655 | -1,27282 | 4,40808 | 1,03963 | 0,432387 | 1 |
| 124 | Lck | 0,160695 | 0,914763 | -1,09318 | 4,23264 | 0,228688 | 0,0990542 | 1 |
| 172 | Pdcd4 | 9,07E-08 | 1,62486 | 1,62486 | 71,9431 | 1,77986 | 0,0453565 | 1 |
| 233 | Tyro3 | 0,170435 | 1,03364 | 1,03364 | 4,15696 | 0,0302252 | 0,0133301 | 1 |
| 126 | LRP6_pS1490 | 8,48E-10 | 1,63707 | 1,63707 | 66,1155 | 0,789773 | 0,0218998 | 1 |
| 181 | PKA-a | 7,46E-10 | 1,9961 | 1,9961 | 75,668 | 2,16153 | 0,0523711 | 1 |
| 218 | Stat3 | 5,00E-06 | 2,03587 | 2,03587 | 22,5764 | 2,24621 | 0,182405 | 1 |
| 64 | Creb | 0,376458 | 0,979757 | -1,02066 | 3,71864 | 0,0251136 | 0,0123813 | 1 |
| 130 | MDM2_pS166 | 0,568461 | 0,972175 | -1,02862 | 3,7033 | 0,0906338 | 0,0448687 | 1 |
| 24 | Aurora-B | 0,295692 | 0,958726 | -1,04305 | 3,69143 | 0,0699945 | 0,0347624 | 1 |
| 32 | Bad_pS112 | 0,0624282 | 1,10949 | 1,10949 | 3,55901 | 0,115742 | 0,0596216 | 1 |
| 220 | Stat5a | 3,63E-12 | 2,16972 | 2,16972 | 212,947 | 2,68086 | 0,0230804 | 1 |
| 95 | GCLM | 0,129298 | 0,961744 | -1,03978 | 3,36925 | 0,0253068 | 0,0137703 | 1 |
| 16 | AMPKa_pT172 | 4,17E-07 | 2,74143 | 2,74143 | 20,1532 | 5,16458 | 0,469821 | 1 |

| | | | | | | | | |
|---|---|---|---|---|---|---|---|---|
| 241 | WIPI1 | 0,827622 | 0,978283 | -1,0222 | 3,21009 | 0,652047 | 0,372394 | 1 |
| 158 | p38-MAPK | 0,392472 | 0,944171 | -1,05913 | 3,14191 | 0,178812 | 0,104338 | 1 |
| 149 | Notch1 | 1,28E-12 | 2,86969 | 2,86969 | 261,231 | 8,7418 | 0,0613504 | 1 |
| 114 | IGFRb | 1,21E-12 | 2,87166 | 2,87166 | 224,45 | 5,12321 | 0,041847 | 1 |
| 129 | MCT4 | 8,25E-08 | 2,87503 | 2,87503 | 84,0128 | 9,79881 | 0,21383 | 1 |
| 34 | Bax | 0,438382 | 0,866859 | -1,15359 | 3,06105 | 1,45184 | 0,869541 | 1 |
| 146 | NAPSIN-A | 0,0732502 | 0,908105 | -1,10119 | 2,99184 | 0,0810062 | 0,0496389 | 1 |
| 221 | Stathmin-1 | 0,457431 | 1,03401 | 1,03401 | 2,83115 | 0,0664936 | 0,0430585 | 1 |
| 23 | ATR_pS428 | 0,566134 | 0,971562 | -1,02927 | 2,5898 | 0,0881786 | 0,0624221 | 1 |
| 103 | HER2_pY1248 | 0,723671 | 1,03284 | 1,03284 | 2,57293 | 0,36422 | 0,259524 | 1 |
| 113 | IGFBP2 | 0,269581 | 1,09608 | 1,09608 | 2,56926 | 0,224706 | 0,160342 | 1 |
| 8 | A-Raf | 0,357691 | 0,942513 | -1,06099 | 2,42788 | 0,121051 | 0,0914079 | 1 |
| 161 | p53 | 0,197587 | 0,922828 | -1,08363 | 2,40202 | 0,106289 | 0,0811244 | 1 |
| 56 | Chk1_pS296 | 0,726063 | 0,987007 | -1,01316 | 2,38404 | 0,0404177 | 0,0310814 | 1 |
| 173 | PDGFR-b | 6,75E-10 | 3,22331 | 3,22331 | 2680,48 | 45,9792 | 0,0314478 | 1 |
| 92 | G6PD | 0,151232 | 1,1312 | 1,1312 | 2,1683 | 0,167204 | 0,141374 | 1 |
| 157 | p27_pT198 | 0,111426 | 0,916702 | -1,09087 | 2,1435 | 0,0802349 | 0,0686247 | 1 |
| 88 | Fibronectin | 5,67E-07 | 3,50475 | 3,50475 | 20,1859 | 16,8349 | 1,52898 | 1 |
| 89 | FoxM1 | 0,126873 | 1,09501 | 1,09501 | 2,09024 | 0,0810734 | 0,0711088 | 1 |
| 226 | TIGAR | 0,135172 | 0,9036 | -1,10668 | 2,02815 | 0,108713 | 0,0982706 | 1 |
| 122 | JNK_pT183_Y185 | 0,152745 | 0,921521 | -1,08516 | 2,02018 | 0,0760564 | 0,0690219 | 1 |
| 117 | IRF-1 | 0,149577 | 0,948751 | -1,05402 | 1,94612 | 0,0303863 | 0,0286253 | 1 |
| 33 | Bak | 0,852836 | 0,990169 | -1,00993 | 1,87467 | 0,0702506 | 0,0687017 | 1 |
| 234 | U-Histone-H2B | 0,738196 | 1,04823 | 1,04823 | 1,55594 | 0,410705 | 0,483925 | 1 |
| 28 | b-Catenin_pT41_S45 | 0,139094 | 0,913904 | -1,09421 | 1,32746 | 0,0577034 | 0,0796934 | 1 |
| 211 | Slfn11 | 0,897035 | 0,993172 | -1,00688 | 1,24644 | 0,0394885 | 0,0580819 | 1 |
| 133 | MERIT40_pS29 | 0,156747 | 1,12178 | 1,12178 | 1,0799 | 0,0862763 | 0,14647 | 1 |
| 198 | RBM15 | 0,253967 | 0,567676 | -1,76157 | 0,951534 | 5,1036 | 9,83319 | 1 |
| 189 | PRAS40_pT246 | 0,66926 | 1,06118 | 1,06118 | 0,904605 | 0,239531 | 0,48545 | 1 |
| 246 | YAP | 0,466005 | 1,02776 | 1,02776 | 0,788196 | 0,0136047 | 0,0316444 | 1 |
| 178 | PEA-15_pS116 | 0,128782 | 0,803653 | -1,24432 | 0,56699 | 0,114289 | 0,369549 | 1 |
| 222 | STING | 0,996961 | 0,998216 | -1,00179 | 0,35209 | 2,14706 | 11,1798 | 1 |

**Table S5.** Comparison of RPPA outcomes of *MMTV-R26* $^{Met}$ cells and tumors belonging to “subtype A” versus “subtype B”.

Cluster 3 (cluster with subtype A) vs. cluster 4 (cluster with subtype B)

| Protein | logFC | AveExpr | t | P.Value | adj.P.Val |
|---|---|---|---|---|---|
| PAR | 1,131472 | 0,3170698 | 3,330956 | 0,0029563 | 0,04152945 |
| ARID1A | 0,625451 | 0,0859351 | 4,062781 | 4,96E-04 | 0,02858153 |
| Histone-H3 | 0,616463 | 0,3605703 | 3,054654 | 0,0056961 | 0,05733171 |
| XRCC1 | 0,550487 | 0,1489197 | 6,796958 | 6,87E-07 | 1,70E-04 |
| ATM | 0,481253 | 0,1386044 | 3,422094 | 0,0023745 | 0,04152945 |
| E-Cadherin | 0,443419 | 0,1698936 | 2,34252 | 0,0283516 | 0,1024357 |
| VASP | 0,430123 | 0,1169322 | 2,906673 | 0,008042 | 0,06207435 |
| Rictor | 0,402752 | 0,3815294 | 3,060335 | 0,0056207 | 0,05733171 |
| P-Cadherin | 0,400019 | 0,0334093 | 1,909318 | 0,0690093 | 0,17902479 |
| eIF4G | 0,398827 | 0,0164296 | 3,67933 | 0,0012718 | 0,03925799 |
| RBM15 | 0,374286 | 0,0558702 | 3,587849 | 0,0015894 | 0,03925799 |
| CDK1 | 0,367319 | 0,2349106 | 2,646612 | 0,0145488 | 0,07645874 |
| 53BP1 | 0,348198 | 0,1029189 | 2,691726 | 0,0131446 | 0,07189393 |
| TAZ | 0,340197 | -0,0059708 | 3,046745 | 0,0058028 | 0,05733171 |
| 4E-BP1 | 0,333041 | 0,1455759 | 3,325777 | 0,0029933 | 0,04152945 |
| Stat3 | 0,328124 | 0,0928596 | 3,007286 | 0,0063644 | 0,05822247 |
| beta-Catenin | 0,324496 | 0,0162367 | 2,143181 | 0,043113 | 0,12986481 |
| Claudin-7 | 0,316824 | 0,1779344 | 2,562707 | 0,0175428 | 0,08405637 |
| ACC1 | 0,31187 | -0,2919491 | 1,689445 | 0,104897 | 0,23990341 |
| WIPI1 | 0,287525 | 0,1169281 | 2,937817 | 0,0074821 | 0,06063795 |
| Notch1 | 0,256416 | 0,0348844 | 2,150548 | 0,0424624 | 0,12948413 |
| Rad50 | 0,252598 | 0,0485918 | 2,930486 | 0,0076104 | 0,06063795 |
| MERIT40_pS29 | 0,236574 | 0,0348573 | 2,887791 | 0,0084007 | 0,06287832 |
| GCN5L2 | 0,228901 | 0,1400084 | 2,546477 | 0,0181848 | 0,08405637 |
| FASN | 0,223324 | -0,2280732 | 1,474832 | 0,1540659 | 0,31192027 |
| DM-K9-Histone-H3 | 0,221614 | 0,0792837 | 3,479287 | 0,0020682 | 0,04152945 |
| PARP | 0,21735 | 0,1754915 | 2,170756 | 0,040723 | 0,12573225 |
| DM-Histone-H3 | 0,21678 | 0,0015729 | 2,793482 | 0,0104327 | 0,06750283 |
| TUFM | 0,215179 | -0,1295122 | 3,427262 | 0,0023451 | 0,04152945 |
| Bim | 0,208428 | 0,120133 | 2,008825 | 0,0566601 | 0,15550059 |
| 4E-BP1_pS65 | 0,207393 | 0,1963715 | 2,611425 | 0,0157408 | 0,07934629 |
| Bax | 0,205523 | 0,0459626 | 2,362578 | 0,0271577 | 0,10011854 |
| Caspase-3 | 0,203513 | 0,053363 | 3,115907 | 0,0049315 | 0,05733171 |
| Rab25 | 0,19528 | 0,053111 | 2,942803 | 0,007396 | 0,06063795 |
| DUSP4 | 0,192819 | -0,034058 | 1,63052 | 0,1168633 | 0,25608136 |
| ER-alpha_pS118 | 0,191709 | -0,0097066 | 4,014505 | 5,59E-04 | 0,02858153 |
| Gab2 | 0,187292 | 0,2209397 | 2,027495 | 0,0545749 | 0,15146063 |
| c-Jun_pS73 | 0,187168 | 0,1662713 | 1,583037 | 0,1273203 | 0,2750364 |
| p70-S6K1 | 0,185401 | 0,1487478 | 1,941728 | 0,0647494 | 0,17383816 |
| Bcl-xL | 0,18301 | -0,0045359 | 2,377068 | 0,0263241 | 0,09851602 |
| B-Raf | 0,182396 | 0,119589 | 2,176267 | 0,0402599 | 0,12573225 |
| AR | 0,179985 | -0,1801834 | 1,721602 | 0,0988159 | 0,23025968 |
| IR-b | 0,178742 | 0,1179838 | 1,763467 | 0,0913511 | 0,22057594 |
| PAK1 | 0,177609 | 0,0452871 | 1,969404 | 0,0612968 | 0,16637706 |

| | | | | | |
|---|---|---|---|---|---|
| RIP | 0,172828 | -0,1031336 | 1,449313 | 0,1609979 | 0,31813179 |
| VEGFR-2 | 0,170852 | 0,112644 | 1,91996 | 0,0675841 | 0,17902479 |
| Smad3 | 0,163193 | -0,0207871 | 2,986563 | 0,0066799 | 0,05892596 |
| Myt1 | 0,147849 | -0,0118133 | 2,625587 | 0,0152504 | 0,07847609 |
| Hexokinase-II | 0,146195 | -0,0752403 | 3,065559 | 0,0055521 | 0,05733171 |
| AMPK_pT172 | 0,145741 | 0,0074493 | 1,501703 | 0,1470302 | 0,30013613 |
| DJ-1 | 0,141801 | 0,0489347 | 1,203647 | 0,2411937 | 0,42553468 |
| Merlin | 0,13962 | -0,0647313 | 1,65167 | 0,1124426 | 0,25248474 |
| Connexin-43 | 0,133795 | 0,0121025 | 0,85322 | 0,4024979 | 0,59110886 |
| SOD2 | 0,132995 | -0,0158534 | 2,529887 | 0,0188637 | 0,08471496 |
| ATR_pS428 | 0,131701 | -0,0822292 | 2,189481 | 0,0391689 | 0,124035 |
| MIF | 0,126377 | 0,1137327 | 2,483055 | 0,0209097 | 0,08904635 |
| Smad1 | 0,126119 | 0,0125196 | 1,803546 | 0,0846632 | 0,20704768 |
| p27_pT198 | 0,12259 | -0,0775066 | 2,440183 | 0,0229618 | 0,09452611 |
| Chk2_pT68 | 0,121915 | 0,0296161 | 1,224318 | 0,2334535 | 0,41784797 |
| GATA6 | 0,11837 | -0,0708088 | 1,849068 | 0,0775839 | 0,19554318 |
| mTOR | 0,112892 | 0,0728351 | 1,263187 | 0,2194103 | 0,40443542 |
| b-Catenin_pT41_S45 | 0,112507 | 0,0870738 | 1,433354 | 0,1654593 | 0,32435275 |
| A-Raf | 0,111316 | 0,0495721 | 1,906472 | 0,0693948 | 0,17902479 |
| Notch3 | 0,11089 | 0,0179976 | 2,520372 | 0,0192636 | 0,08496624 |
| c-Kit | 0,109967 | -0,0346328 | 2,541729 | 0,0183767 | 0,08405637 |
| CD4 | 0,109381 | -0,1543716 | 1,723178 | 0,0985256 | 0,23025968 |
| Raptor | 0,106089 | 0,0743287 | 1,270748 | 0,2167555 | 0,40254593 |
| cdc25C | 0,105267 | -0,0550074 | 1,087078 | 0,288471 | 0,47820368 |
| c-Myc | 0,104214 | 0,1120016 | 1,154996 | 0,2601691 | 0,4462622 |
| IRS1 | 0,101347 | 0,0292407 | 0,925277 | 0,3646062 | 0,55497351 |
| WIPI2 | 0,1013 | -0,1982764 | 1,070823 | 0,2955621 | 0,48028836 |
| EGFR | 0,099325 | 0,1196557 | 1,526609 | 0,1407455 | 0,29213552 |
| C-Raf | 0,097531 | -0,0158396 | 1,731508 | 0,0970041 | 0,23025968 |
| RPA32_pS4_S8 | 0,097246 | 0,0030646 | 1,077405 | 0,2926762 | 0,48028836 |
| Bid | 0,092652 | -0,0289827 | 1,315894 | 0,2014139 | 0,37976522 |
| Tyro3 | 0,090933 | -0,0275295 | 1,825937 | 0,0811143 | 0,20125619 |
| IGFRb | 0,085766 | 0,0774075 | 0,529761 | 0,6014485 | 0,77315301 |
| SDHA | 0,083141 | -0,0625461 | 1,325743 | 0,1981819 | 0,37654562 |
| Glutaminase | 0,082261 | -0,0878491 | 1,096358 | 0,2844782 | 0,47477109 |
| MEK1_pS217_S221 | 0,079621 | -0,0247615 | 1,759823 | 0,0919811 | 0,22057594 |
| eIF4E | 0,076387 | -0,0104446 | 0,6724 | 0,5081521 | 0,69729755 |
| PI3K-p110-a | 0,076263 | 0,0773417 | 1,630262 | 0,1169183 | 0,25608136 |
| YAP | 0,072962 | -0,0114551 | 1,115955 | 0,2761775 | 0,46723176 |
| HER3_pY1289 | 0,071337 | -0,0242126 | 1,110572 | 0,2784398 | 0,46785463 |
| LRP6_pS1490 | 0,070404 | 0,0860639 | 0,949516 | 0,3524114 | 0,54673438 |
| STAT5-alpha | 0,070151 | 0,0335454 | 0,374359 | 0,7116303 | 0,84501285 |
| AMPK-a2_pS345 | 0,067148 | -0,0243644 | 1,224603 | 0,2333481 | 0,41784797 |
| STING | 0,066374 | 0,1183359 | 0,310674 | 0,7588984 | 0,87185075 |
| JNK2 | 0,066189 | 0,0509504 | 0,825831 | 0,4175405 | 0,60311405 |
| c-Abl | 0,06544 | -0,0534208 | 0,9741 | 0,340328 | 0,53542056 |
| PR | 0,064823 | 3,60E-04 | 1,171619 | 0,2535651 | 0,43810296 |
| PTEN | 0,064072 | 0,0033979 | 0,632158 | 0,5336332 | 0,72025901 |
| G6PD | 0,056212 | -0,0405422 | 0,41671 | 0,6808257 | 0,83048272 |
| cIAP | 0,054272 | -0,0154717 | 0,94158 | 0,3563734 | 0,54673438 |

| | | | | | |
|---|---|---|---|---|---|
| p27-Kip-1 | 0,053339 | 0,0575981 | 1,127133 | 0,2715221 | 0,46252386 |
| AMPK_alpha | 0,047113 | 0,106793 | 0,689381 | 0,4976073 | 0,69440119 |
| Atg3 | 0,046415 | -0,159205 | 0,553954 | 0,5850549 | 0,76306001 |
| UBAC1 | 0,046399 | -0,0899037 | 0,658319 | 0,5169899 | 0,70550553 |
| Rad51 | 0,046157 | -0,0949432 | 0,605841 | 0,5506624 | 0,73920439 |
| PKC-alpha | 0,045954 | 0,1939478 | 0,274739 | 0,7860152 | 0,8806955 |
| eIF4E_pS209 | 0,045378 | -0,0631162 | 0,945002 | 0,3546613 | 0,54673438 |
| NAPSIN-A | 0,044088 | -0,0234169 | 0,791903 | 0,436658 | 0,62343651 |
| Axl | 0,042812 | -0,0614344 | 0,507958 | 0,6164094 | 0,77680158 |
| FoxM1 | 0,039265 | -0,0026116 | 0,451163 | 0,656175 | 0,81037612 |
| Creb | 0,036445 | -0,0330347 | 0,922075 | 0,3662376 | 0,55497351 |
| eEF2 | 0,035399 | -0,0115659 | 0,298964 | 0,767702 | 0,87383596 |
| D-a-Tubulin | 0,033737 | -0,0371452 | 0,511837 | 0,6137346 | 0,77680158 |
| JNK_pT183_pY185 | 0,031993 | 0,0011722 | 0,588901 | 0,5617736 | 0,74601111 |
| BRD4 | 0,030861 | 0,0060905 | 0,651368 | 0,5213838 | 0,70759235 |
| PDHK1 | 0,025525 | -0,0481166 | 0,536905 | 0,5965844 | 0,77149923 |
| Granzyme-B | 0,025318 | -0,1310343 | 0,258625 | 0,7982674 | 0,88349877 |
| COG3 | 0,022993 | 0,0297644 | 0,344984 | 0,7332995 | 0,86249986 |
| XPF | 0,02051 | -0,0346966 | 0,4101 | 0,685598 | 0,83048272 |
| Bak | 0,01909 | -0,0570662 | 0,23899 | 0,8132694 | 0,88349877 |
| FAK | 0,019023 | -0,1338132 | 0,196817 | 0,8457312 | 0,90820757 |
| NF-kB-p65_pS536 | 0,018266 | 0,0282627 | 0,192106 | 0,8493763 | 0,90820757 |
| FoxO3a | 0,017794 | -0,0865903 | 0,384539 | 0,7041769 | 0,84432861 |
| PKC-delta_pS664 | 0,017134 | 0,0161086 | 0,389313 | 0,7006921 | 0,84424857 |
| N-Cadherin | 0,015604 | -0,0350948 | 0,253208 | 0,8023984 | 0,88349877 |
| Jak2 | 0,011693 | -0,0562983 | 0,156208 | 0,877257 | 0,91592889 |
| PAICS | 0,009199 | 0,068113 | 0,078792 | 0,9378923 | 0,95993902 |
| IGFBP2 | 0,002766 | 0,0294213 | 0,02627 | 0,9792726 | 0,98726669 |
| EGFR_pY1173 | 0,001998 | 0,0068537 | 0,04823 | 0,9619566 | 0,97571776 |
| HSP70 | -9,36E-04 | 0,127176 | -0,01018 | 0,9919661 | 0,99196612 |
| MDM2_pS166 | -0,00152 | 0,2541291 | -0,02032 | 0,9839646 | 0,98796441 |
| 14-3-3_zeta | -0,00262 | 0,1376385 | -0,04581 | 0,9638669 | 0,97571776 |
| Shc_pY317 | -0,00873 | 0,0301483 | -0,07547 | 0,9405071 | 0,95993902 |
| Stat3_pY705 | -0,00942 | 0,0686024 | -0,16531 | 0,8701717 | 0,91592889 |
| PI3K-p85 | -0,00967 | 0,0713216 | -0,09531 | 0,924907 | 0,95188347 |
| Aurora-B | -0,0097 | 0,01747 | -0,1729 | 0,8642672 | 0,91592889 |
| CD134 | -0,00978 | -0,1496793 | -0,1745 | 0,8630245 | 0,91592889 |
| INPP4b | -0,01017 | -0,0663281 | -0,23944 | 0,8129208 | 0,88349877 |
| PDK1_pS241 | -0,01383 | 0,111296 | -0,15417 | 0,8788467 | 0,91592889 |
| Stathmin | -0,01555 | 0,0087648 | -0,25689 | 0,7995869 | 0,88349877 |
| TIGAR | -0,01592 | 0,0086581 | -0,22454 | 0,8243579 | 0,88915461 |
| PRAS40_pT246 | -0,01648 | 0,1035279 | -0,1188 | 0,9064814 | 0,94076011 |
| 04-oct | -0,01819 | -0,0377163 | -0,28753 | 0,7763327 | 0,87558987 |
| p53 | -0,01975 | -0,041495 | -0,4161 | 0,6812657 | 0,83048272 |
| PAK4 | -0,02093 | -0,0564756 | -0,32651 | 0,7470436 | 0,87181917 |
| Cdc2_pY15 | -0,02183 | -0,0437417 | -0,3196 | 0,7522109 | 0,87185075 |
| Cyclin_D1 | -0,02217 | 0,0211284 | -0,29165 | 0,7732166 | 0,87558987 |
| 14-3-3_beta | -0,02295 | -0,0765912 | -0,36975 | 0,7150109 | 0,84501285 |
| HER3 | -0,02299 | -0,0146603 | -0,24109 | 0,8116594 | 0,88349877 |
| PREX1 | -0,02378 | 0,0037087 | -0,32485 | 0,748282 | 0,87181917 |

| | | | | | |
|---|---|---|---|---|---|
| HES1 | -0,02381 | 0,040922 | -0,15602 | 0,8774025 | 0,91592889 |
| Wee1_pS642 | -0,02829 | -0,0582548 | -0,47399 | 0,6400616 | 0,79444828 |
| Rab11 | -0,0347 | 6,85E-04 | -0,57397 | 0,5716632 | 0,75508456 |
| TSC1 | -0,03475 | 0,084765 | -0,40968 | 0,6859048 | 0,83048272 |
| PDGFR-b | -0,03484 | 0,2735999 | -0,09909 | 0,9219449 | 0,95188347 |
| ATM_pS1981 | -0,03672 | -0,0181914 | -0,71053 | 0,4846512 | 0,68405048 |
| GCLM | -0,03747 | -0,0844496 | -0,89226 | 0,3816656 | 0,56958898 |
| RSK | -0,03772 | 0,0447656 | -0,37143 | 0,7137828 | 0,84501285 |
| PEA15 | -0,03997 | 0,100227 | -0,48655 | 0,6312653 | 0,78748748 |
| Glutamate-D1-2 | -0,04005 | 0,006518 | -0,27214 | 0,7879907 | 0,8806955 |
| Cox2 | -0,04352 | -0,0368758 | -0,51248 | 0,6132918 | 0,77680158 |
| IGF1R_pY1135_Y1136 | -0,0443 | -0,0292372 | -0,8901 | 0,3828007 | 0,56958898 |
| Src_pY416 | -0,04576 | 0,1671187 | -0,23603 | 0,8155373 | 0,88349877 |
| Wee1 | -0,04659 | 0,0202702 | -0,30498 | 0,7631726 | 0,872702 |
| LC3A-B | -0,04699 | -0,0407871 | -0,55111 | 0,5869692 | 0,76306001 |
| FOXO3a_pS318_S321 | -0,04802 | 0,1285134 | -0,86926 | 0,3938533 | 0,58252548 |
| MMP2 | -0,04894 | -0,0069953 | -1,00785 | 0,3242056 | 0,52219947 |
| Tuberin_pT1462 | -0,04902 | -0,0744172 | -0,60081 | 0,5539491 | 0,73959697 |
| FAK_pY397 | -0,05659 | 0,09564 | -0,49238 | 0,6272014 | 0,78638962 |
| Ets-1 | -0,05815 | -0,0496285 | -0,96877 | 0,3429218 | 0,53608663 |
| Mcl-1 | -0,0605 | 0,1007123 | -0,68002 | 0,5034044 | 0,69648258 |
| Beclin | -0,06576 | -0,0428242 | -1,22691 | 0,2324949 | 0,41784797 |
| p16_INK4a | -0,06889 | 0,0593482 | -0,55142 | 0,5867618 | 0,76306001 |
| B7-H4 | -0,07178 | -0,0254777 | -1,36892 | 0,1844887 | 0,3588088 |
| PKM2 | -0,07668 | 0,0484725 | -0,3143 | 0,7561821 | 0,87185075 |
| p44-42-MAPK | -0,07773 | 2,21E-05 | -1,07351 | 0,2943824 | 0,48028836 |
| MCT4 | -0,08051 | 6,78E-04 | -0,52584 | 0,6041236 | 0,77315301 |
| Slfn11 | -0,08247 | -0,0769721 | -1,23643 | 0,2290046 | 0,41784797 |
| Elk1_pS383 | -0,08984 | -0,0110248 | -1,51092 | 0,1446778 | 0,29779513 |
| Tuberin | -0,09163 | 0,0086241 | -0,90071 | 0,3772518 | 0,56817809 |
| TTF1 | -0,09406 | -0,0691308 | -1,46065 | 0,1578879 | 0,31703801 |
| PKA-a | -0,09458 | 0,0742455 | -1,00492 | 0,3255819 | 0,52219947 |
| Heregulin | -0,0977 | -0,0312672 | -1,69577 | 0,1036767 | 0,2393285 |
| Gys | -0,10409 | -0,0146164 | -0,74778 | 0,4623131 | 0,65627208 |
| Jagged1 | -0,1053 | 0,0592771 | -1,89412 | 0,0710898 | 0,1810225 |
| PLC-gamma2_pY759 | -0,10893 | 0,1434968 | -1,57983 | 0,1280534 | 0,2750364 |
| Caspase-7_cleavedD198 | -0,11207 | 0,0887793 | -0,84964 | 0,4044429 | 0,59110886 |
| HER2_pY1248 | -0,11683 | -0,028885 | -2,84922 | 0,0091814 | 0,06670045 |
| Akt | -0,11736 | 0,0629356 | -1,20863 | 0,2393095 | 0,42524774 |
| ERCC5 | -0,12431 | 0,0282191 | -0,98398 | 0,3355524 | 0,53471894 |
| B-Raf_pS445 | -0,12751 | 0,1013743 | -0,97938 | 0,3377704 | 0,53480307 |
| Rb_pS807_S811 | -0,12758 | 0,0272653 | -0,79409 | 0,4354084 | 0,62343651 |
| MAPK_pT202_Y204 | -0,13335 | -0,0711768 | -1,52798 | 0,140405 | 0,29213552 |
| YAP_pS127 | -0,13727 | 0,1047023 | -0,83629 | 0,4117556 | 0,59825671 |
| p70S6K_pT389 | -0,1463 | -0,0543965 | -2,38207 | 0,0260422 | 0,09851602 |
| eEF2K | -0,15185 | -0,0364881 | -1,31039 | 0,203239 | 0,38030336 |
| ER-alpha | -0,1525 | -0,0489758 | -2,12593 | 0,0446718 | 0,13293895 |
| p21 | -0,16075 | -0,0093602 | -2,21492 | 0,0371438 | 0,12071739 |
| Atg7 | -0,16082 | -0,0104942 | -2,30048 | 0,0310121 | 0,107887 |
| P-Met_pY1234_Y1235 | -0,16251 | 0,2264404 | -0,67788 | 0,5047384 | 0,69648258 |

| | | | | | |
|---|---|---|---|---|---|
| YB1_pS102 | -0,16926 | -0,0186932 | -2,05423 | 0,0517075 | 0,14513365 |
| Paxillin | -0,16968 | 0,0199992 | -1,82359 | 0,0814802 | 0,20125619 |
| Bad_pS112 | -0,1814 | 0,0648426 | -2,38673 | 0,0257817 | 0,09851602 |
| PEA15_pS116 | -0,18554 | -0,0263777 | -3,62304 | 0,0014589 | 0,03925799 |
| mTOR_pS2448 | -0,18887 | 0,1628695 | -1,45599 | 0,1591608 | 0,31703801 |
| MEK1 | -0,19369 | 0,0244747 | -2,21758 | 0,0369377 | 0,12071739 |
| MSI2 | -0,19481 | -0,0673892 | -1,33243 | 0,1960093 | 0,37530469 |
| ACC_pS79 | -0,19619 | -0,1864269 | -0,69342 | 0,4951176 | 0,69440119 |
| U-Histone-H2B | -0,20031 | -0,0548845 | -1,34821 | 0,1909613 | 0,36849573 |
| SLC1A5 | -0,20084 | 0,1496055 | -1,55452 | 0,1339673 | 0,28281975 |
| TFAM | -0,20188 | 0,0319333 | -3,1974 | 0,0040662 | 0,05285995 |
| LDHA | -0,20267 | 0,0494123 | -2,4196 | 0,0240121 | 0,09566098 |
| HSP27_pS82 | -0,20785 | -0,0132763 | -2,27218 | 0,0329295 | 0,11296646 |
| Mnk1 | -0,20914 | 0,034214 | -2,11814 | 0,0453926 | 0,13347587 |
| b-Actin | -0,21224 | 0,1361256 | -2,08337 | 0,0487362 | 0,13997493 |
| SHP-2_pY542 | -0,21356 | -0,0152545 | -2,33117 | 0,0290485 | 0,10249987 |
| CD20 | -0,23204 | -0,0273627 | -2,72529 | 0,012184 | 0,06839641 |
| p38_MAPK | -0,23446 | 0,0700919 | -2,25274 | 0,0343083 | 0,11485535 |
| Cox-IV | -0,24076 | -0,043776 | -2,5984 | 0,0162049 | 0,08005199 |
| PMS2 | -0,24249 | -0,0387305 | -2,75431 | 0,0114071 | 0,06750283 |
| Cyclin_B1 | -0,25359 | -0,0490261 | -2,75183 | 0,0114717 | 0,06750283 |
| Myosin-IIa_pS1943 | -0,25359 | -0,0490261 | -2,75183 | 0,0114717 | 0,06750283 |
| PLK1 | -0,25359 | -0,0490261 | -2,75183 | 0,0114717 | 0,06750283 |
| TRIM25 | -0,25359 | -0,0490261 | -2,75183 | 0,0114717 | 0,06750283 |
| C-Raf_pS338 | -0,25626 | 0,0632267 | -2,25134 | 0,0344101 | 0,11485535 |
| TFRC | -0,26456 | -0,0425438 | -3,0103 | 0,0063198 | 0,05822247 |
| Collagen_VI | -0,26524 | 0,0298493 | -2,68355 | 0,0133892 | 0,07189393 |
| Gys_pS641 | -0,28804 | -0,1177172 | -2,45877 | 0,0220503 | 0,09231214 |
| Rictor_pT1135 | -0,30625 | 0,0756234 | -3,70501 | 0,0011945 | 0,03925799 |
| ULK1_pS757 | -0,30751 | 0,0702128 | -1,62915 | 0,1171546 | 0,25608136 |
| Pdcd4 | -0,31026 | 0,1711035 | -1,17143 | 0,2536386 | 0,43810296 |
| MSH6 | -0,32168 | 0,0879335 | -1,18801 | 0,2471765 | 0,43299717 |
| Sox2 | -0,32872 | 0,2627177 | -1,56069 | 0,1325045 | 0,28214312 |
| p90RSK_pT573 | -0,36455 | 0,1115986 | -2,19125 | 0,0390248 | 0,124035 |
| Chk1_pS296 | -0,38385 | 0,1682085 | -2,75111 | 0,0114903 | 0,06750283 |
| PDK1 | -0,38557 | -0,0217785 | -2,0993 | 0,0471778 | 0,13709323 |
| PKC-b-II_pS660 | -0,42941 | 0,2542991 | -2,81445 | 0,009944 | 0,06750283 |
| ZAP-70 | -0,43507 | 0,188693 | -2,39244 | 0,0254655 | 0,09851602 |
| XBP1 | -0,44064 | 0,0239927 | -2,5454 | 0,0182283 | 0,08405637 |
| Lck | -0,46571 | 0,0837977 | -2,74122 | 0,0117515 | 0,06750283 |
| IRF-1 | -0,4735 | -0,109279 | -2,49305 | 0,0204565 | 0,08864496 |
| Akt_pT308 | -0,48868 | 0,0607433 | -3,89334 | 7,53E-04 | 0,03098879 |
| p38_pT180_Y182 | -0,49449 | 0,0719863 | -3,32118 | 0,0030264 | 0,04152945 |
| Fibronectin | -0,58387 | 0,376004 | -1,68088 | 0,1065695 | 0,24149247 |
| S6_pS240_S244 | -0,63893 | 0,1788402 | -3,11939 | 0,0048911 | 0,05733171 |
| GSK-3a-b_pS21_S9 | -0,72138 | 0,1130619 | -4,17276 | 3,78E-04 | 0,02858153 |
| PD-L1 | -0,78775 | 0,2728692 | -2,33819 | 0,0286156 | 0,1024357 |
| Caveolin-1 | -0,79932 | -0,2343902 | -1,90511 | 0,0695805 | 0,17902479 |
| NDRG1_pT346 | -0,88585 | 0,3263951 | -2,42061 | 0,0239595 | 0,09566098 |
| S6_pS235_S236 | -0,89273 | 0,2497408 | -4,00027 | 5,79E-04 | 0,02858153 |

| | | | | | |
|---|---|---|---|---|---|
| Akt_pS473 | -1,03289 | 0,0325736 | -3,35669 | 0,0027793 | 0,04152945 |
| Src_pY527 | -1,07674 | 0,1555428 | -3,45293 | 0,0022043 | 0,04152945 |
| MYH11 | -1,75648 | 0,5057535 | -2,06601 | 0,0504883 | 0,14334031 |

**Table S6.** Comparisons of RPPA outcomes of *MMTV-R26*$^{Met}$ cells belonging to “subtype A” and “subtype B” versus those from the non-tumorigenic cells.

### Subtype A (MGT4+9+11) vs. non-tumorigenic (MGT2 & 7)

| Protein | logFC | AveExpr | t | P.Value | adj.P.Val |
|---|---|---|---|---|---|
| P-Met_pY1234_Y1235 | 3,771668 | -1,0119078 | 10,90563 | 6,84E-08 | 5,77E-06 |
| E-Cadherin | 3,468199 | 1,5253849 | 10,61925 | 9,35E-08 | 5,77E-06 |
| Claudin-7 | 1,911199 | 1,0822876 | 10,33591 | 1,28E-07 | 6,33E-06 |
| RSK | 1,524384 | -0,6111489 | 7,299833 | 6,18E-06 | 1,53E-04 |
| CDK1 | 1,448314 | -0,1166139 | 4,725375 | 4,02E-04 | 0,00183884 |
| ARID1A | 1,446252 | 0,3963761 | 4,852469 | 3,20E-04 | 0,00155111 |
| Bim | 1,416402 | 0,0453894 | 4,942895 | 2,73E-04 | 0,00142332 |
| 53BP1 | 1,269014 | 0,5378448 | 5,401039 | 1,23E-04 | 7,89E-04 |
| Rad50 | 1,061401 | 0,2447965 | 5,410073 | 1,21E-04 | 7,89E-04 |
| HES1 | 0,981841 | 0,0343482 | 2,795423 | 0,0152226 | 0,03357122 |
| Src_pY416 | 0,958743 | -0,2971786 | 6,696275 | 1,52E-05 | 2,50E-04 |
| ACC_pS79 | 0,939118 | -0,11619 | 2,665268 | 0,0195073 | 0,04054706 |
| Connexin-43 | 0,895784 | 0,2361452 | 3,028428 | 0,0097416 | 0,02382356 |
| FASN | 0,887145 | -0,1825017 | 5,620712 | 8,49E-05 | 7,23E-04 |
| Src_pY527 | 0,883245 | -0,1438668 | 4,22756 | 9,98E-04 | 0,00347248 |
| Rb_pS807_S811 | 0,870722 | -0,0627291 | 2,38951 | 0,0328075 | 0,06157929 |
| ACC1 | 0,865206 | 0,0651942 | 4,657714 | 4,54E-04 | 0,00196783 |
| Cox2 | 0,828262 | 0,6923661 | 1,422211 | 0,1786289 | 0,24648788 |
| 4E-BP1_pS65 | 0,794498 | 0,2176293 | 6,006903 | 4,49E-05 | 5,28E-04 |
| GCN5L2 | 0,78403 | 0,1609002 | 5,658902 | 7,96E-05 | 7,23E-04 |
| Akt_pS473 | 0,706044 | -0,29258 | 4,279576 | 9,06E-04 | 0,00334177 |
| c-Jun_pS73 | 0,687669 | 0,1712913 | 6,835796 | 1,23E-05 | 2,17E-04 |
| p90RSK_pT573 | 0,660934 | -0,1175991 | 5,401963 | 1,23E-04 | 7,89E-04 |
| PARP | 0,639323 | 0,189449 | 6,210382 | 3,24E-05 | 4,44E-04 |
| Akt_pT308 | 0,631021 | -0,1776652 | 3,715662 | 0,0026124 | 0,00750295 |
| HER3 | 0,584665 | 0,4124621 | 4,245167 | 9,66E-04 | 0,00340895 |
| C-Raf_pS338 | 0,560357 | -6,44E-04 | 5,077137 | 2,15E-04 | 0,0011679 |
| PKC-b-II_pS660 | 0,557357 | -0,279136 | 5,524983 | 9,97E-05 | 7,73E-04 |
| IRS1 | 0,543443 | 0,197673 | 1,828737 | 0,0905815 | 0,1399653 |
| VASP | 0,536384 | -0,1346535 | 3,548721 | 0,0035909 | 0,00996573 |
| c-Myc | 0,536074 | 0,1818384 | 5,782582 | 6,48E-05 | 6,96E-04 |
| beta-Catenin | 0,520031 | 0,2023753 | 3,444927 | 0,0043796 | 0,01163174 |
| eIF4E | 0,500608 | -0,0795851 | 7,337273 | 5,85E-06 | 1,53E-04 |
| 4E-BP1 | 0,49076 | 0,0226499 | 2,387859 | 0,0329088 | 0,06157929 |
| Notch1 | 0,475117 | -0,2848443 | 2,481032 | 0,027637 | 0,0541772 |
| p16_INK4a | 0,460764 | 0,1754814 | 2,047757 | 0,061451 | 0,10328628 |
| Caspase-3 | 0,432888 | 0,0986794 | 7,15535 | 7,63E-06 | 1,63E-04 |
| RPA32_pS4_S8 | 0,422635 | -0,0877175 | 4,922764 | 2,83E-04 | 0,00142496 |
| NDRG1_pT346 | 0,421693 | 0,3033009 | 0,904163 | 0,3824397 | 0,47708386 |
| RBM15 | 0,419121 | 0,4528309 | 1,104033 | 0,2896937 | 0,37859436 |
| Paxillin | 0,399942 | -0,0701099 | 8,728997 | 8,82E-07 | 3,63E-05 |
| PAICS | 0,395387 | 0,2794241 | 2,228145 | 0,0442483 | 0,07751302 |
| Bcl-xL | 0,392312 | 0,1383302 | 4,062771 | 0,001357 | 0,00424273 |
| 14-3-3_zeta | 0,388742 | -0,0596922 | 4,658633 | 4,53E-04 | 0,00196783 |
| Bid | 0,381066 | 0,1070465 | 3,283919 | 0,0059639 | 0,01534459 |
| HER2_pY1248 | 0,3781 | 0,0388001 | 2,664526 | 0,0195348 | 0,04054706 |
| FAK | 0,377805 | -0,0713128 | 5,6286 | 8,38E-05 | 7,23E-04 |
| Histone-H3 | 0,376417 | 0,0086542 | 3,004335 | 0,0102027 | 0,02470651 |
| p38_pT180_Y182 | 0,363108 | 0,0857146 | 5,278316 | 1,52E-04 | 9,11E-04 |
| Shc_pY317 | 0,352321 | -0,2506858 | 4,352649 | 7,92E-04 | 0,00305704 |
| c-Kit | 0,345628 | 0,1667719 | 5,504168 | 1,03E-04 | 7,73E-04 |
| AMPK_alpha | 0,343258 | -0,1949831 | 3,730602 | 0,0025393 | 0,00737877 |
| PRAS40_pT246 | 0,333433 | -0,1478686 | 1,602546 | 0,1331629 | 0,19578124 |
| Smad3 | 0,32264 | 0,1278642 | 4,631279 | 4,76E-04 | 0,00202851 |
| HSP70 | 0,316225 | 0,0233271 | 5,187279 | 1,78E-04 | 0,00102115 |
| WIPI1 | 0,299004 | 0,2067124 | 1,857936 | 0,0860821 | 0,13542849 |
| PDK1 | 0,288736 | -0,0743783 | 4,052317 | 0,0013838 | 0,00427252 |
| PAK1 | 0,288257 | -0,0960324 | 1,206203 | 0,2493386 | 0,3329007 |
| Bax | 0,282279 | -0,0402982 | 0,891662 | 0,3888601 | 0,48265551 |
| DJ-1 | 0,28216 | 0,1007253 | 4,336833 | 8,16E-04 | 0,00309901 |
| IR-b | 0,271336 | 0,0634504 | 1,320554 | 0,2095391 | 0,28437448 |
| Lck | 0,270305 | 0,0645107 | 2,784147 | 0,0155539 | 0,0339984 |
| Merlin | 0,25649 | 0,0583102 | 2,001088 | 0,0668193 | 0,11002916 |
| TUFM | 0,255489 | 0,0967286 | 3,668656 | 0,0028567 | 0,00811037 |
| Myt1 | 0,251577 | -0,0590917 | 3,421506 | 0,0045805 | 0,01203606 |
| PDK1_pS241 | 0,24966 | 0,0705303 | 1,828203 | 0,0906658 | 0,1399653 |
| ATM | 0,248594 | 0,0743071 | 1,061881 | 0,3077141 | 0,40002836 |
| ER-alpha_pS118 | 0,247404 | 0,1571599 | 4,543131 | 5,59E-04 | 0,00222667 |
| Bad_pS112 | 0,246678 | -0,1849946 | 3,61229 | 0,0031806 | 0,00892728 |
| Caspase-7_cleavedD19 | 0,233133 | -0,1285137 | 2,305651 | 0,0383474 | 0,06814244 |
| Bak | 0,22832 | -0,0479864 | 3,170981 | 0,0074085 | 0,01848373 |
| Chk2_pT68 | 0,22803 | -0,034172 | 2,891423 | 0,0126689 | 0,02897413 |
| Wee1_pS642 | 0,223293 | 0,1465578 | 2,56915 | 0,0234061 | 0,04700253 |
| Elk1_pS383 | 0,217788 | -0,0541057 | 4,129958 | 0,0011969 | 0,00394175 |
| Cdc2_pY15 | 0,208341 | 0,0433277 | 1,556484 | 0,1437134 | 0,20758604 |
| b-Actin | 0,207393 | 0,1279735 | 1,793327 | 0,0963213 | 0,14686021 |
| EGFR_pY1173 | 0,201168 | -0,0426149 | 4,551783 | 5,50E-04 | 0,00222667 |
| STING | 0,200475 | 0,1379571 | 0,41888 | 0,6821762 | 0,75222104 |
| ATR_pS428 | 0,192485 | 0,00212 | 3,993173 | 0,0015462 | 0,00465732 |

### Subtype B (MGT13) vs. non-tumorigenic (MGT2 & 7)

| Protein | logFC | AveExpr | t | P.Value | adj.P.Val |
|---|---|---|---|---|---|
| P-Met_pY1234_Y1235 | 4,563148 | -1,6674278 | 44,05051 | 1,05E-09 | 2,60E-07 |
| RSK | 2,114524 | -0,7602672 | 6,313678 | 4,25E-04 | 0,00656047 |
| Notch1 | 2,029212 | 0,2558908 | 10,54554 | 1,68E-05 | 5,19E-04 |
| HES1 | 1,627067 | 0,0519271 | 6,696406 | 2,98E-04 | 0,0056618 |
| Fibronectin | 1,565388 | 0,9234731 | 10,57193 | 1,65E-05 | 5,19E-04 |
| ACC_pS79 | 1,380298 | -0,174792 | 3,366794 | 0,0122547 | 0,05132152 |
| CDK1 | 1,366848 | -0,5335007 | 3,823602 | 0,0067068 | 0,03524618 |
| Rb_pS807_S811 | 1,304012 | -0,1066745 | 2,532854 | 0,0395758 | 0,11217854 |
| Akt_pS473 | 1,265382 | -0,2390729 | 11,70825 | 8,45E-06 | 3,64E-04 |
| IGFRb | 1,122679 | 0,5041849 | 4,190776 | 0,004226 | 0,03070037 |
| Bim | 1,058487 | -0,4815593 | 7,109478 | 2,07E-04 | 0,00451158 |
| Src_pY416 | 1,017153 | -0,5250009 | 18,32814 | 4,21E-07 | 5,08E-05 |
| Akt_pT308 | 0,99835 | -0,1866618 | 7,260346 | 1,82E-04 | 0,00448387 |
| PKA-a | 0,965586 | 0,2854551 | 17,31854 | 6,17E-07 | 5,08E-05 |
| ACC1 | 0,852846 | -0,1682892 | 3,899131 | 0,0060888 | 0,03379328 |
| FASN | 0,838035 | -0,4375213 | 3,49291 | 0,0103442 | 0,0464548 |
| LRP6_pS1490 | 0,671783 | 0,1657914 | 11,62867 | 8,84E-06 | 3,64E-04 |
| p90RSK_pT573 | 0,607544 | -0,3147928 | 3,325557 | 0,0129593 | 0,05334913 |
| Shc_pY317 | 0,593491 | -0,2402474 | 6,355699 | 4,08E-04 | 0,00656047 |
| AMPK_pT172 | 0,585846 | 0,8578037 | 3,89801 | 0,0060975 | 0,03379328 |
| Pdcd4 | 0,584895 | 0,116794 | 3,924916 | 0,0058923 | 0,03379328 |
| PKC-b-II_pS660 | 0,563848 | -0,4233496 | 5,389217 | 0,0010738 | 0,01386302 |
| STING | 0,544972 | 0,2327264 | 3,055622 | 0,0187941 | 0,06981802 |
| MCT4 | 0,534748 | 0,5135805 | 0,88172 | 0,4076264 | 0,52833899 |
| AMPK_alpha | 0,481 | -0,2264803 | 4,409915 | 0,003239 | 0,02424311 |
| PRAS40_pT246 | 0,470494 | -0,177067 | 8,623494 | 6,16E-05 | 0,00169047 |
| IR-b | 0,453404 | 0,0699186 | 1,657671 | 0,1420237 | 0,25420179 |
| STAT5-alpha | 0,451643 | 0,587122 | 2,638098 | 0,0339911 | 0,10494747 |
| HER2_pY1248 | 0,445391 | -0,0320792 | 4,41123 | 0,0032339 | 0,02424311 |
| Tuberin | 0,432484 | 0,1376954 | 4,905013 | 0,0018226 | 0,01911688 |
| SLC1A5 | 0,412054 | 0,3457002 | 2,460839 | 0,0439391 | 0,11545689 |
| Src_pY527 | 0,410403 | -0,5794568 | 1,690426 | 0,1354647 | 0,24602776 |
| Paxillin | 0,406428 | -0,1728096 | 7,041667 | 2,19E-04 | 0,00451158 |
| Bad_pS112 | 0,397009 | -0,1856245 | 4,420566 | 0,0031979 | 0,02424311 |
| Caspase-7_cleavedD198 | 0,371346 | -0,130765 | 3,639518 | 0,0085192 | 0,04125948 |
| XRCC1 | 0,366656 | 0,0387943 | 3,039023 | 0,0192347 | 0,06986708 |
| IGF1R_pY1135_Y1136 | 0,348902 | 0,1505626 | 3,239624 | 0,0145717 | 0,05900362 |
| Bid | 0,345436 | -0,0087244 | 2,510036 | 0,0409076 | 0,11226863 |
| Mcl-1 | 0,334732 | 0,3562144 | 2,893978 | 0,0235855 | 0,08205113 |
| RIP | 0,325051 | 0,1058554 | 1,440921 | 0,193453 | 0,30269376 |
| ATM | 0,31833 | 0,0386307 | 0,832604 | 0,4329695 | 0,55411127 |
| COG3 | 0,306351 | 0,0145178 | 4,422202 | 0,0031917 | 0,02424311 |
| 14-3-3_zeta | 0,305893 | -0,197724 | 5,347493 | 0,0011225 | 0,01386302 |
| eIF4E | 0,300825 | -0,2972351 | 4,907438 | 0,0018176 | 0,01911688 |
| WIPI1 | 0,299748 | 0,1281728 | 3,925854 | 0,0058853 | 0,03379328 |
| mTOR | 0,29646 | -0,0810146 | 4,04117 | 0,0050885 | 0,03307519 |
| MAPK_pT202_Y204 | 0,294888 | -0,0230378 | 2,702394 | 0,0309894 | 0,09801214 |
| Chk2_pT68 | 0,285975 | -0,0694781 | 3,509896 | 0,0101125 | 0,04625544 |
| PAK1 | 0,283941 | -0,1739056 | 1,541461 | 0,1677734 | 0,28383578 |
| Wee1 | 0,282046 | 0,180511 | 1,947076 | 0,0932163 | 0,18604001 |
| p21 | 0,275143 | -0,1092629 | 3,873219 | 0,0062935 | 0,03379328 |
| Myt1 | 0,267636 | -0,1185596 | 3,748261 | 0,007392 | 0,03651662 |
| FAK | 0,266835 | -0,2185122 | 3,366532 | 0,012259 | 0,05132152 |
| YB1_pS102 | 0,248129 | -0,0819382 | 3,803376 | 0,0068837 | 0,03529816 |
| Caspase-3 | 0,244179 | -0,0963645 | 4,070668 | 0,0049042 | 0,03273887 |
| 4E-BP1_pS65 | 0,236739 | -0,2309479 | 1,380441 | 0,2105612 | 0,32505382 |
| PLC-gamma2_pY759 | 0,23491 | 0,0072565 | 2,118022 | 0,0725491 | 0,1585807 |
| Bak | 0,215446 | -0,11372 | 2,249705 | 0,0598096 | 0,14342683 |
| UBAC1 | 0,214845 | -0,092247 | 3,790107 | 0,0070025 | 0,03529816 |
| Bax | 0,208181 | -0,146502 | 2,8076 | 0,026662 | 0,08780673 |
| GATA6 | 0,201924 | -0,0159812 | 2,874659 | 0,0242395 | 0,08315488 |
| Cdc2_pY15 | 0,197787 | -0,0161427 | 0,987127 | 0,3569608 | 0,49533325 |
| PI3K-p85 | 0,197071 | 0,0344405 | 0,548609 | 0,6005859 | 0,70571753 |
| GCN5L2 | 0,19554 | -0,298087 | 0,966581 | 0,3664395 | 0,50283648 |
| PARP | 0,19536 | -0,1694338 | 1,365077 | 0,2151167 | 0,32798665 |
| Histone-H3 | 0,194263 | -0,1686865 | 2,039951 | 0,081353 | 0,16967817 |
| Stathmin | 0,191596 | -0,1444441 | 2,518617 | 0,0404014 | 0,11217854 |
| ERCC5 | 0,189449 | 0,0877916 | 0,905446 | 0,3957795 | 0,52557809 |
| beta-Catenin | 0,171947 | -0,083955 | 1,057484 | 0,3259429 | 0,46268912 |
| EGFR_pY1173 | 0,169934 | -0,1090561 | 2,744401 | 0,0291791 | 0,09360055 |
| Elk1_pS383 | 0,169645 | -0,1321767 | 2,395789 | 0,0483065 | 0,12300738 |
| SDHA | 0,16948 | 0,1233133 | 2,017755 | 0,0840447 | 0,17156231 |
| PTEN | 0,164032 | 0,2496951 | 0,94282 | 0,3776406 | 0,5125122 |
| Connexin-43 | 0,158935 | -0,3158979 | 0,379397 | 0,7158123 | 0,78931092 |
| mTOR_pS2448 | 0,150508 | -0,0301328 | 2,20943 | 0,0634461 | 0,14779565 |
| Lck | 0,150488 | -0,0581287 | 1,887569 | 0,1016952 | 0,19935492 |
| ATR_pS428 | 0,150049 | -0,066832 | 2,208924 | 0,0634932 | 0,14779565 |
| FOXO3a_pS318_S321 | 0,146765 | -0,08466 | 2,59818 | 0,0360061 | 0,10715061 |
| Tyro3 | 0,14106 | -0,0157264 | 2,036651 | 0,0817478 | 0,16967817 |

| | | | | | |
|---|---|---|---|---|---|
| DM-K9-Histone-H3 | 0,191106 | -0,032935 | 2,955169 | 0,0112119 | 0,02688678 |
| eEF2 | 0,187505 | 0,0403709 | 1,234271 | 0,2390563 | 0,32090713 |
| Rictor | 0,187175 | -0,0121124 | 2,449399 | 0,0293286 | 0,05615625 |
| eIF4G | 0,186366 | 0,0353867 | 3,198586 | 0,0070258 | 0,017708 |
| EGFR | 0,182966 | -0,0770142 | 2,536964 | 0,0248733 | 0,04954598 |
| MSH6 | 0,182251 | 0,1297574 | 1,5399 | 0,1476837 | 0,21208063 |
| MEK1_pS217_S221 | 0,17823 | 0,0065781 | 3,843738 | 0,0020492 | 0,00602553 |
| XRCC1 | 0,173953 | 0,002085 | 1,931898 | 0,0755753 | 0,12362311 |
| p27_pT198 | 0,17211 | 0,0723343 | 2,328813 | 0,0367341 | 0,06622864 |
| Gab2 | 0,171561 | 0,0479602 | 0,624194 | 0,5433525 | 0,62422358 |
| NAPSIN-A | 0,170103 | -0,1065433 | 2,9256 | 0,0118658 | 0,02764966 |
| UBAC1 | 0,166895 | -0,0687808 | 2,357465 | 0,0348277 | 0,06419738 |
| Jagged1 | 0,158894 | 0,1041255 | 0,993534 | 0,3386726 | 0,43119655 |
| TAZ | 0,153525 | 0,1660362 | 2,347866 | 0,0354557 | 0,06439373 |
| p53 | 0,147901 | -0,0309641 | 1,44288 | 0,172831 | 0,24118219 |
| Notch3 | 0,144246 | 0,0288049 | 2,942955 | 0,0114776 | 0,02699964 |
| PAR | 0,143159 | 0,7936152 | 0,264154 | 0,7958228 | 0,84364046 |
| Stathmin | 0,139431 | -0,1300271 | 1,902231 | 0,079641 | 0,12857082 |
| mTOR | 0,136343 | -0,1136771 | 1,652966 | 0,1223894 | 0,18321324 |
| Aurora-B | 0,133051 | -0,0498797 | 1,806192 | 0,0941993 | 0,14451695 |
| YB1_pS102 | 0,130054 | -0,0982415 | 1,779569 | 0,0986378 | 0,14946961 |
| PLC-gamma2_pY759 | 0,119652 | -0,0105832 | 1,610908 | 0,131321 | 0,19539936 |
| AMPK-a2_pS345 | 0,115932 | -0,1005424 | 1,507819 | 0,1556308 | 0,22220123 |
| Tuberin | 0,115824 | 0,0325309 | 1,403352 | 0,1840597 | 0,25117539 |
| Tyro3 | 0,092494 | -0,0121462 | 1,858878 | 0,0859403 | 0,13542849 |
| p27-Kip-1 | 0,091811 | -0,0645913 | 2,350207 | 0,0353015 | 0,06439373 |
| FOXO3a_pS318_S321 | 0,088969 | -0,0859653 | 2,14559 | 0,0514697 | 0,08828487 |
| Cyclin_D1 | 0,087443 | -0,0020169 | 1,432363 | 0,175761 | 0,24389313 |
| PEA15_pS116 | 0,085991 | -0,0163314 | 0,631631 | 0,5386309 | 0,62422358 |
| TIGAR | 0,084525 | 0,0411889 | 0,976319 | 0,3468127 | 0,43929614 |
| b-Catenin_pT41_S45 | 0,079037 | 0,0461708 | 1,01903 | 0,3268706 | 0,41832662 |
| Chk1_pS296 | 0,071257 | -0,0518607 | 1,16746 | 0,264095 | 0,34883135 |
| GCLM | 0,062402 | -0,0037227 | 1,236505 | 0,2382525 | 0,32090713 |
| Raptor | 0,049775 | 0,0118633 | 0,593726 | 0,5629346 | 0,64075962 |
| MDM2_pS166 | 0,04399 | -0,1730021 | 0,553736 | 0,5892041 | 0,66758449 |
| Akt | 0,039874 | -0,0891319 | 0,293054 | 0,7741296 | 0,82774897 |
| DUSP4 | 0,037353 | 0,0838537 | 0,318001 | 0,7555598 | 0,81494878 |
| MAPK_pT202_Y204 | 0,034884 | -0,1252682 | 0,241066 | 0,8132833 | 0,85481267 |
| Rab11 | 0,032688 | -0,048905 | 0,381195 | 0,7092465 | 0,77514998 |
| C-Raf | 0,030052 | -0,0492449 | 0,301115 | 0,7681125 | 0,82488607 |
| Sox2 | 0,028709 | 0,2078454 | 0,150345 | 0,8828119 | 0,90329557 |
| NF-kB-p65_pS536 | 0,027659 | 0,0087904 | 0,245802 | 0,8096926 | 0,85467557 |
| YAP | 0,026607 | -0,0505988 | 0,45738 | 0,654985 | 0,72874462 |
| cIAP | 0,014036 | -0,0504367 | 0,346067 | 0,7348567 | 0,79609472 |
| Ets-1 | 0,006433 | 0,0497873 | 0,147502 | 0,8850102 | 0,90329557 |
| Wee1 | 0,005595 | 0,0635073 | 0,032864 | 0,9742851 | 0,97428513 |
| IGF1R_pY1135_Y1136 | 0,003058 | 0,0031501 | 0,042925 | 0,9664169 | 0,9703454 |
| eIF4E_pS209 | -0,0069 | 0,0259484 | -0,10684 | 0,9165562 | 0,92403831 |
| U-Histone-H2B | -0,01159 | -0,0096519 | -0,15162 | 0,8818259 | 0,90329557 |
| JNK2 | -0,01277 | 0,0018004 | -0,1781 | 0,8614012 | 0,8939752 |
| A-Raf | -0,01623 | 0,0166796 | -0,14277 | 0,8886742 | 0,9033026 |
| XPF | -0,02234 | -0,1081235 | -0,39114 | 0,7020631 | 0,7707093 |
| HER3_pY1289 | -0,0246 | -0,0026466 | -0,26741 | 0,7933662 | 0,84364046 |
| PI3K-p110-a | -0,02523 | -0,036344 | -0,46261 | 0,6513318 | 0,72874462 |
| eEF2K | -0,02755 | 0,178513 | -0,16833 | 0,8689247 | 0,89801002 |
| PI3K-p85 | -0,02841 | -0,069686 | -0,13565 | 0,8941876 | 0,90518167 |
| D-a-Tubulin | -0,03121 | -0,0398156 | -0,53242 | 0,6034589 | 0,68061341 |
| SLC1A5 | -0,03636 | 0,143932 | -0,19651 | 0,8472655 | 0,88301506 |
| RIP | -0,03699 | -0,0590641 | -0,21562 | 0,8326489 | 0,87145884 |
| MMP2 | -0,03721 | 0,055784 | -0,96445 | 0,3525077 | 0,44423161 |
| PEA15 | -0,03872 | 0,1718814 | -0,35493 | 0,7283627 | 0,79253564 |
| PKA-a | -0,04 | -0,1560579 | -0,45884 | 0,6539663 | 0,72874462 |
| LRP6_pS1490 | -0,04168 | -0,1509694 | -0,71962 | 0,4845572 | 0,58099818 |
| mTOR_pS2448 | -0,04959 | -0,1289684 | -0,42059 | 0,6809608 | 0,75222104 |
| p21 | -0,05003 | -0,2618167 | -0,69608 | 0,4986958 | 0,58824698 |
| JNK_pT183_pY185 | -0,05417 | 0,0593339 | -0,62554 | 0,5424947 | 0,62422358 |
| Slfn11 | -0,05446 | -0,0775038 | -0,82271 | 0,425579 | 0,52038616 |
| Smad1 | -0,05639 | 0,0367162 | -0,69372 | 0,500129 | 0,58824698 |
| Rad51 | -0,06129 | 0,2265507 | -0,6108 | 0,5519147 | 0,6311247 |
| Glutamate-D1-2 | -0,06848 | 0,0224619 | -0,73836 | 0,4734825 | 0,57048871 |
| c-Abl | -0,07024 | -0,0653876 | -1,56494 | 0,1417235 | 0,20591584 |
| IGFBP2 | -0,07332 | -0,0240207 | -0,64764 | 0,528544 | 0,61580364 |
| p70-S6K1 | -0,07489 | 0,1299386 | -0,82936 | 0,4219433 | 0,51850749 |
| IRF-1 | -0,07708 | 0,0617817 | -1,4848 | 0,161554 | 0,22802199 |
| PREX1 | -0,0774 | 0,2077197 | -1,05361 | 0,3113465 | 0,40263137 |
| p38_MAPK | -0,09257 | 0,0649005 | -0,80506 | 0,4353339 | 0,52969197 |
| Creb | -0,09465 | 0,0546613 | -2,20983 | 0,0457627 | 0,07960132 |
| PKC-delta_pS664 | -0,10128 | -0,0486037 | -2,32125 | 0,0372539 | 0,06667902 |
| ERCC5 | -0,10161 | -0,0637429 | -0,77106 | 0,4545252 | 0,55033195 |
| 14-3-3_beta | -0,10446 | -0,0199976 | -2,04436 | 0,0618276 | 0,10328628 |
| GSK-3a-b_pS21_S9 | -0,12418 | 0,0017529 | -0,68567 | 0,505028 | 0,59119387 |
| Mnk1 | -0,12863 | 0,1504279 | -1,1942 | 0,2538398 | 0,33708831 |
| Pdcd4 | -0,13265 | -0,2257078 | -0,91752 | 0,3756589 | 0,47100381 |
| VEGFR-2 | -0,13295 | 0,0340055 | -0,86012 | 0,4053808 | 0,50064526 |

| | | | | | |
|---|---|---|---|---|---|
| S6_pS240_S244 | 0,127725 | 0,4220785 | 1,371766 | 0,2131229 | 0,32696494 |
| ARID1A | 0,115095 | -0,5555488 | 0,353378 | 0,7343676 | 0,80617243 |
| DM-K9-Histone-H3 | 0,1107 | -0,1177962 | 1,125894 | 0,2978848 | 0,43027804 |
| PDK1 | 0,106406 | -0,2286699 | 0,953741 | 0,3724603 | 0,50827461 |
| p38_pT180_Y182 | 0,09872 | -0,1233594 | 1,03424 | 0,3359452 | 0,46880484 |
| XPF | 0,095511 | -0,0517258 | 1,156349 | 0,2860453 | 0,41560697 |
| Raptor | 0,088684 | 0,0154113 | 0,808989 | 0,4455471 | 0,56435964 |
| c-Kit | 0,086084 | -0,0356159 | 0,919586 | 0,3888417 | 0,51915619 |
| PI3K-p110-a | 0,082661 | 0,016552 | 0,925104 | 0,3861591 | 0,51837655 |
| Akt | 0,081209 | -0,0819331 | 1,179535 | 0,2772956 | 0,40527812 |
| c-Abl | 0,079637 | 0,0173695 | 1,262439 | 0,2478299 | 0,37010532 |
| PAICS | 0,07913 | 0,0396074 | 0,710801 | 0,5005479 | 0,61942666 |
| Aurora-B | 0,075079 | -0,1098153 | 0,899516 | 0,3987164 | 0,52664673 |
| GSK-3a-b_pS21_S9 | 0,07313 | 0,1190676 | 0,398842 | 0,7020784 | 0,77763844 |
| Caveolin-1 | 0,070314 | 0,2043218 | 0,209433 | 0,840166 | 0,88684184 |
| YAP | 0,068369 | -0,039718 | 1,111025 | 0,3038096 | 0,4362847 |
| IGFBP2 | 0,065561 | 0,0548377 | 0,461112 | 0,658917 | 0,75348374 |
| MSI2 | 0,060932 | 0,120869 | 0,599495 | 0,5680198 | 0,67129616 |
| U-Histone-H2B | 0,059832 | 0,0240155 | 0,674241 | 0,522123 | 0,63473657 |
| MEK1_pS217_S221 | 0,053707 | -0,0937947 | 0,882184 | 0,4073923 | 0,52833899 |
| Chk1_pS296 | 0,051428 | -0,079152 | 0,534241 | 0,6099673 | 0,71066945 |
| p27_pT198 | 0,050474 | -0,0251875 | 0,621983 | 0,5539641 | 0,66101032 |
| DJ-1 | 0,044436 | -0,0755722 | 0,670191 | 0,5245494 | 0,63473657 |
| p53 | 0,040754 | -0,1158909 | 0,463467 | 0,6573101 | 0,75348374 |
| C-Raf | 0,03794 | -0,0537902 | 0,312643 | 0,7637999 | 0,82744992 |
| Stat3 | 0,033749 | 0,4534121 | 0,347239 | 0,7387735 | 0,80742064 |
| C-Raf_pS338 | 0,028811 | -0,3762361 | 0,414808 | 0,6908904 | 0,7721716 |
| NAPSIN-A | 0,026572 | -0,2129188 | 0,285877 | 0,7833742 | 0,84494947 |
| cIAP | 0,026387 | -0,0488455 | 0,417351 | 0,6891158 | 0,7721716 |
| Bcl-xL | 0,024299 | -0,1228564 | 0,436409 | 0,6758845 | 0,76229901 |
| CD20 | 0,016336 | 0,1297929 | 0,200478 | 0,8468943 | 0,89013995 |
| GCLM | 0,006495 | -0,0441405 | 0,082951 | 0,9362486 | 0,95240706 |
| c-Jun_pS73 | 0,004896 | -0,302689 | 0,055213 | 0,9575349 | 0,96535153 |
| Smad3 | 0,003638 | -0,0939425 | 0,028646 | 0,9779587 | 0,98193417 |
| MDM2_pS166 | 0,001828 | -0,2026732 | 0,01909 | 0,9853102 | 0,98531024 |
| TUFM | -0,0077 | -0,0834473 | -0,08199 | 0,9369835 | 0,95240706 |
| MERIT40_pS29 | -0,01226 | 0,1136221 | -0,08342 | 0,935886 | 0,95240706 |
| Wee1_pS642 | -0,01489 | -0,0144094 | -0,11372 | 0,9127006 | 0,94325128 |
| ER-alpha_pS118 | -0,01584 | -0,0209075 | -0,21508 | 0,835933 | 0,88616077 |
| AR | -0,01759 | 0,1283755 | -0,14103 | 0,8918789 | 0,92951097 |
| Rictor_pT1135 | -0,02078 | 0,1220186 | -0,14653 | 0,8876933 | 0,92906879 |
| YAP_pS127 | -0,02338 | 0,2413937 | -0,0691 | 0,946876 | 0,95851789 |
| 4E-BP1 | -0,03111 | -0,3304412 | -0,11791 | 0,9094996 | 0,94325128 |
| HER3_pY1289 | -0,03615 | -0,0011075 | -0,25026 | 0,8096846 | 0,86576661 |
| JNK2 | -0,03974 | -0,0063912 | -0,6728 | 0,5229878 | 0,63473657 |
| PR | -0,03991 | 0,2736279 | -0,66643 | 0,5268057 | 0,63473657 |
| Stat3_pY705 | -0,04201 | 0,325379 | -0,33924 | 0,7445334 | 0,81013109 |
| PKC-delta_pS664 | -0,04845 | 7,52E-04 | -0,71805 | 0,4963418 | 0,6191738 |
| FoxM1 | -0,04982 | 0,0565277 | -0,50459 | 0,6295739 | 0,72665769 |
| b-Catenin_pT41_S45 | -0,0504 | -0,0301463 | -0,40772 | 0,6958495 | 0,77421098 |
| PDK1_pS241 | -0,05081 | -0,1240855 | -0,22416 | 0,8291295 | 0,88273697 |
| D-a-Tubulin | -0,05104 | -0,0400841 | -0,60807 | 0,5626368 | 0,66813119 |
| eIF4G | -0,05348 | -0,1165538 | -0,70028 | 0,5066953 | 0,62265539 |
| TIGAR | -0,05361 | -0,0403032 | -0,8807 | 0,4081404 | 0,52833899 |
| Notch3 | -0,05379 | -0,0941086 | -0,73702 | 0,4854356 | 0,60864262 |
| Rictor | -0,05741 | -0,1662992 | -0,92512 | 0,3861535 | 0,51837655 |
| NDRG1_pT346 | -0,06058 | -0,0146028 | -0,08866 | 0,9318744 | 0,95240706 |
| Ets-1 | -0,06377 | 0,0180023 | -1,03536 | 0,3354585 | 0,46880484 |
| Slfn11 | -0,06517 | -0,0677315 | -0,79237 | 0,454551 | 0,572827 |
| Rad50 | -0,07267 | -0,5211625 | -0,27644 | 0,7903156 | 0,84873025 |
| RPA32_pS4_S8 | -0,07986 | -0,4145376 | -0,64032 | 0,5426597 | 0,65066474 |
| p44-42-MAPK | -0,0827 | 0,0931581 | -0,81917 | 0,4400946 | 0,56032659 |
| eEF2 | -0,08428 | -0,1255586 | -0,4373 | 0,6752698 | 0,76229901 |
| CD134 | -0,08434 | 0,1674613 | -1,35055 | 0,2195032 | 0,33262135 |
| p27-Kip-1 | -0,08497 | -0,1645682 | -1,44029 | 0,193626 | 0,30269376 |
| PKM2 | -0,08995 | 0,0966772 | -0,51679 | 0,6214707 | 0,72067255 |
| A-Raf | -0,09276 | -0,0118433 | -0,70906 | 0,50156 | 0,61942666 |
| G6PD | -0,09602 | 0,0621284 | -0,87988 | 0,4085536 | 0,52833899 |
| Cyclin_D1 | -0,10112 | -0,1058925 | -1,48559 | 0,1816317 | 0,2990869 |
| 14-3-3_beta | -0,10458 | 0,0075003 | -1,46237 | 0,1876933 | 0,30269376 |
| OCT4 | -0,11116 | 0,1129347 | -2,05236 | 0,0798856 | 0,16864745 |
| Hexokinase-II | -0,11298 | 0,1487511 | -0,84776 | 0,4250306 | 0,5467841 |
| eIF4E_pS209 | -0,11764 | -0,0196953 | -1,19881 | 0,2701927 | 0,39872105 |
| AMPK-a2_pS345 | -0,11883 | -0,23173 | -1,54239 | 0,1675505 | 0,28383578 |
| MSH6 | -0,12202 | -0,0487089 | -1,25977 | 0,2487348 | 0,37010532 |
| Creb | -0,12371 | 0,0671715 | -2,12557 | 0,0717497 | 0,15823379 |
| TAZ | -0,12866 | 0,0046104 | -1,61369 | 0,1513017 | 0,26693938 |
| WIPI2 | -0,12954 | 0,2692593 | -0,44044 | 0,6731039 | 0,76229901 |
| cdc25C | -0,13853 | 0,2915575 | -1,05224 | 0,328177 | 0,46319843 |
| MMP2 | -0,13915 | 0,0219057 | -2,56676 | 0,0376792 | 0,11079485 |
| Claudin-7 | -0,14004 | -0,2087461 | -1,94576 | 0,0933966 | 0,18604001 |
| IRS1 | -0,14048 | -0,2387615 | -0,54511 | 0,6028599 | 0,70571753 |
| N-Cadherin | -0,14183 | 0,0073635 | -1,93337 | 0,0951062 | 0,1879298 |
| Rab25 | -0,14206 | 0,0114982 | -1,60086 | 0,154112 | 0,26996927 |

| | | | | | |
|---|---|---|---|---|---|
| GATA6 | -0,1389 | -0,1986784 | -2,02354 | 0,0641846 | 0,10639995 |
| Jak2 | -0,14217 | 0,0189833 | -2,3765 | 0,0336141 | 0,06242619 |
| PAK4 | -0,14445 | 0,0926862 | -1,13604 | 0,2765507 | 0,36334052 |
| OCT4 | -0,15027 | 0,0565422 | -3,86126 | 0,0019824 | 0,0058994 |
| N-Cadherin | -0,1619 | -0,0439391 | -1,45835 | 0,1685958 | 0,23660887 |
| P-Cadherin | -0,16595 | 0,1045818 | -2,04382 | 0,0618881 | 0,10328628 |
| AR | -0,1731 | 0,0160716 | -1,84732 | 0,0876941 | 0,13709139 |
| COG3 | -0,1775 | -0,2396591 | -1,48881 | 0,1605088 | 0,2278487 |
| CD20 | -0,18028 | -0,0020163 | -2,72244 | 0,0174965 | 0,03757937 |
| MERIT40_pS29 | -0,18033 | -0,0059637 | -1,87899 | 0,0829643 | 0,13306605 |
| MSI2 | -0,18129 | -0,0307509 | -2,63475 | 0,0206712 | 0,04240027 |
| FoxM1 | -0,18286 | -0,0487133 | -2,49443 | 0,026949 | 0,05325125 |
| ZAP-70 | -0,20149 | 0,0909259 | -2,62787 | 0,0209426 | 0,04240027 |
| Rab25 | -0,20263 | -0,0678976 | -2,90888 | 0,0122521 | 0,02828296 |
| INPP4b | -0,20465 | 0,0035981 | -4,54366 | 5,58E-04 | 0,00222667 |
| ER-alpha | -0,20535 | 0,0807164 | -4,26333 | 9,34E-04 | 0,00334396 |
| FAK_pY397 | -0,21048 | 0,3062533 | -0,70205 | 0,4950855 | 0,58791402 |
| FoxO3a | -0,21308 | 0,0217331 | -3,44668 | 0,0043649 | 0,01163174 |
| CD134 | -0,21867 | 0,0522217 | -4,13803 | 0,001179 | 0,00393535 |
| ATM_pS1981 | -0,21994 | 0,0095047 | -4,08413 | 0,0013038 | 0,00412884 |
| Caveolin-1 | -0,2202 | 0,0217395 | -0,70632 | 0,4925168 | 0,58768911 |
| TTF1 | -0,22137 | 0,0775741 | -4,47425 | 6,34E-04 | 0,00248458 |
| XBP1 | -0,22244 | 0,0394977 | -5,52061 | 1,00E-04 | 7,73E-04 |
| Rictor_pT1135 | -0,22379 | -0,0240085 | -1,66556 | 0,1198209 | 0,18046192 |
| PKM2 | -0,23357 | -0,026473 | -1,4102 | 0,1820713 | 0,24984234 |
| cdc25C | -0,23387 | 0,1890208 | -1,90632 | 0,0790684 | 0,12848621 |
| Tuberin_pT1462 | -0,23808 | 0,0478807 | -3,2277 | 0,0066437 | 0,01691757 |
| PD-L1 | -0,25607 | 0,0026159 | -6,13692 | 3,64E-05 | 4,69E-04 |
| Hexokinase-II | -0,25661 | 0,0195186 | -3,05028 | 0,0093414 | 0,02307335 |
| Beclin | -0,26967 | 0,0575019 | -5,26686 | 1,55E-04 | 9,11E-04 |
| SDHA | -0,27331 | -0,1385362 | -4,27001 | 9,23E-04 | 0,00334396 |
| B-Raf_pS445 | -0,27469 | 0,0216048 | -2,79904 | 0,0151178 | 0,03357122 |
| MIF | -0,28066 | -0,0150759 | -4,2845 | 8,98E-04 | 0,00334177 |
| G6PD | -0,28562 | -0,094455 | -2,45428 | 0,0290612 | 0,056079 |
| DM-Histone-H3 | -0,28801 | 0,0980327 | -4,60032 | 5,04E-04 | 0,0021091 |
| BRD4 | -0,30036 | -0,0010115 | -4,09225 | 0,0012842 | 0,00411945 |
| PR | -0,30594 | 0,0789249 | -6,64866 | 1,63E-05 | 2,52E-04 |
| CD4 | -0,3083 | 0,0887386 | -4,15699 | 0,0011381 | 0,00385073 |
| WIPI2 | -0,31893 | 0,1039822 | -1,59248 | 0,1354101 | 0,19790704 |
| B7-H4 | -0,33225 | 0,0212483 | -4,74741 | 3,86E-04 | 0,00180088 |
| p44-42-MAPK | -0,33678 | -0,104557 | -2,86825 | 0,0132437 | 0,02973803 |
| TSC1 | -0,33822 | -0,0948599 | -1,60718 | 0,13214 | 0,19544057 |
| SHP-2_pY542 | -0,35906 | -0,0189077 | -4,10303 | 0,0012586 | 0,0040904 |
| Mcl-1 | -0,36634 | -0,0408638 | -1,86243 | 0,0854074 | 0,13542849 |
| TFAM | -0,38584 | 0,2122181 | -2,6807 | 0,0189432 | 0,03999125 |
| PDHK1 | -0,38987 | -0,0669717 | -6,22144 | 3,18E-05 | 4,44E-04 |
| Fibronectin | -0,40037 | -0,0245861 | -1,03917 | 0,3177623 | 0,40878793 |
| IGFRb | -0,40486 | -0,2572525 | -2,41451 | 0,03131 | 0,05948891 |
| SOD2 | -0,42029 | 0,0561037 | -8,10914 | 1,99E-06 | 6,15E-05 |
| PTEN | -0,44637 | -0,1296311 | -2,87312 | 0,0131207 | 0,02973227 |
| LC3A-B | -0,44985 | 0,1000738 | -2,23768 | 0,0434784 | 0,07670833 |
| PMS2 | -0,45375 | 0,2736831 | -2,94846 | 0,0113571 | 0,02697308 |
| MEK1 | -0,45539 | 0,2784054 | -3,46083 | 0,0042482 | 0,01153095 |
| Atg7 | -0,46796 | 0,0609753 | -2,73123 | 0,0172061 | 0,03727985 |
| Heregulin | -0,49019 | -0,0687039 | -2,46804 | 0,0283203 | 0,05507965 |
| YAP_pS127 | -0,49942 | -0,0943395 | -2,17194 | 0,049053 | 0,0847279 |
| Collagen_VI | -0,50292 | 0,030011 | -5,13438 | 1,95E-04 | 0,00109412 |
| B-Raf | -0,51067 | 0,0724391 | -5,4379 | 1,16E-04 | 7,89E-04 |
| ULK1_pS757 | -0,51347 | 0,0762755 | -7,12813 | 7,94E-06 | 1,63E-04 |
| Atg3 | -0,51804 | -0,0841261 | -6,85944 | 1,18E-05 | 2,17E-04 |
| Stat3_pY705 | -0,54483 | -0,0338044 | -5,75694 | 6,76E-05 | 6,96E-04 |
| Axl | -0,56846 | -0,0268375 | -8,15267 | 1,88E-06 | 6,15E-05 |
| HSP27_pS82 | -0,59208 | 0,1358536 | -4,68489 | 4,32E-04 | 0,00194184 |
| LDHA | -0,65569 | 0,079271 | -4,93506 | 2,77E-04 | 0,00142332 |
| STAT5-alpha | -0,67924 | -0,0766854 | -4,76584 | 3,74E-04 | 0,00177584 |
| Gys_pS641 | -0,68198 | -0,0642531 | -4,86307 | 3,14E-04 | 0,00155111 |
| Glutaminase | -0,68524 | -0,0335895 | -4,03746 | 0,0014229 | 0,00433884 |
| MYH11 | -0,68998 | 0,094718 | -3,28948 | 0,0059006 | 0,01534159 |
| S6_pS240_S244 | -0,6902 | -0,1104895 | -2,13149 | 0,0528086 | 0,08995675 |
| TFRC | -0,69256 | 0,4027866 | -3,52715 | 0,003742 | 0,01026971 |
| PKC-alpha | -0,72642 | -0,0970691 | -5,45213 | 1,13E-04 | 7,89E-04 |
| Granzyme-B | -0,78897 | -0,0597221 | -10,6371 | 9,17E-08 | 5,77E-06 |
| p70S6K_pT389 | -0,81525 | -0,0130179 | -4,20323 | 0,0010443 | 0,00358243 |
| Cyclin_B1 | -0,87661 | 0,244537 | -5,39404 | 1,25E-04 | 7,89E-04 |
| AMPK_pT172 | -0,8829 | -0,0045108 | -6,11099 | 3,80E-05 | 4,69E-04 |
| Gys | -0,90761 | -0,0679323 | -5,7821 | 6,49E-05 | 6,96E-04 |
| Cox-IV | -0,94707 | 0,1957551 | -5,62421 | 8,44E-05 | 7,23E-04 |
| Myosin-IIa_pS1943 | -0,99755 | 0,160807 | -5,07158 | 2,18E-04 | 0,0011679 |
| PLK1 | -1,01791 | 0,1467133 | -5,59318 | 8,89E-05 | 7,32E-04 |
| MCT4 | -1,02875 | -0,4278113 | -2,6302 | 0,0208504 | 0,04240027 |
| Stat3 | -1,03562 | -0,278021 | -5,65456 | 8,02E-05 | 7,23E-04 |
| TRIM25 | -1,08168 | 0,1025633 | -5,34429 | 1,36E-04 | 8,37E-04 |
| S6_pS235_S236 | -1,21966 | -0,0407079 | -2,6995 | 0,0182779 | 0,03891927 |
| PDGFR-b | -3,45067 | -0,6152202 | -14,695 | 1,90E-09 | 4,69E-07 |

| | | | | | |
|---|---|---|---|---|---|
| ZAP-70 | -0,14738 | 0,1672586 | -1,76888 | 0,1209039 | 0,22568701 |
| PDHK1 | -0,14793 | 0,1395381 | -1,46221 | 0,1877337 | 0,30269376 |
| Atg3 | -0,14802 | 0,2110792 | -1,41202 | 0,2014673 | 0,31297125 |
| XBP1 | -0,14841 | 0,1298931 | -2,51885 | 0,0403879 | 0,11217854 |
| DM-Histone-H3 | -0,15033 | 0,2330026 | -2,06291 | 0,0786587 | 0,16835293 |
| IRF-1 | -0,15225 | 0,0498924 | -2,39699 | 0,0482217 | 0,12300738 |
| ATM_pS1981 | -0,16042 | 0,0930213 | -1,84521 | 0,1081839 | 0,20876115 |
| Jak2 | -0,16069 | 0,0485434 | -2,48551 | 0,0423908 | 0,11506074 |
| FoxO3a | -0,16263 | 0,0995509 | -2,81837 | 0,0262562 | 0,08763892 |
| p38_MAPK | -0,16318 | 0,0590571 | -1,51641 | 0,173862 | 0,29016159 |
| JNK_pT183_pY185 | -0,16694 | 0,0252905 | -1,63531 | 0,1466713 | 0,26063165 |
| RBM15 | -0,16872 | 0,0903623 | -2,20324 | 0,0640248 | 0,14779565 |
| Rab11 | -0,17184 | -0,1451814 | -1,73941 | 0,1261882 | 0,23260071 |
| Collagen_VI | -0,17461 | 0,3033535 | -1,10641 | 0,3056669 | 0,43641459 |
| HER3 | -0,1871 | -0,0724925 | -2,16897 | 0,0673238 | 0,14981071 |
| PD-L1 | -0,18744 | 0,0995665 | -3,17387 | 0,0159508 | 0,06155995 |
| Smad1 | -0,18966 | -0,0055285 | -3,21912 | 0,0149875 | 0,05970839 |
| INPP4b | -0,19469 | 0,0618413 | -2,99018 | 0,0205961 | 0,07372812 |
| B7-H4 | -0,19824 | 0,1663108 | -1,85572 | 0,1065382 | 0,20720414 |
| Mnk1 | -0,20086 | 0,1533918 | -1,46557 | 0,1868452 | 0,30269376 |
| Rad51 | -0,20229 | 0,1822849 | -2,40296 | 0,0478038 | 0,12300738 |
| MEK1 | -0,20639 | 0,5052242 | -1,19607 | 0,2711949 | 0,39872105 |
| Tuberin_pT1462 | -0,21457 | 0,1207483 | -2,28442 | 0,0568454 | 0,13765495 |
| PEA15_pS116 | -0,21797 | -0,1692776 | -1,58466 | 0,1577332 | 0,27352611 |
| B-Raf_pS445 | -0,21915 | 0,1178505 | -2,76894 | 0,0281732 | 0,09156284 |
| CD4 | -0,22355 | 0,2063683 | -2,0594 | 0,0790645 | 0,16835293 |
| EGFR | -0,23428 | -0,3040888 | -1,44886 | 0,1913027 | 0,30269376 |
| LDHA | -0,23618 | 0,4319941 | -1,51999 | 0,1729796 | 0,29016159 |
| c-Myc | -0,23945 | -0,2919121 | -1,83696 | 0,1094939 | 0,20931118 |
| DUSP4 | -0,23995 | -0,0448429 | -1,69424 | 0,1347189 | 0,24602776 |
| Beclin | -0,2442 | 0,139538 | -3,88753 | 0,0061795 | 0,03379328 |
| 53BP1 | -0,24702 | -0,4465708 | -1,76536 | 0,1215238 | 0,22568701 |
| HSP70 | -0,25318 | -0,3041033 | -2,59833 | 0,0359985 | 0,10715061 |
| Glutamate-D1-2 | -0,25518 | -0,0394912 | -2,01831 | 0,0839767 | 0,17156231 |
| Granzyme-B | -0,2555 | 0,3769877 | -2,69438 | 0,031348 | 0,09801214 |
| Gys | -0,25676 | 0,4503697 | -0,97707 | 0,3615775 | 0,49893661 |
| BRD4 | -0,2582 | 0,0962749 | -2,32405 | 0,0536424 | 0,13249685 |
| ER-alpha | -0,25858 | 0,1120595 | -4,53533 | 0,0027905 | 0,02424311 |
| TTF1 | -0,25943 | 0,1196475 | -4,01472 | 0,0052602 | 0,0333148 |
| Axl | -0,26673 | 0,252402 | -3,6237 | 0,0086982 | 0,04131634 |
| TFAM | -0,27437 | 0,3617507 | -1,67315 | 0,1388871 | 0,25040237 |
| Sox2 | -0,27551 | 0,0698959 | -1,448 | 0,1915353 | 0,30269376 |
| SHP-2_pY542 | -0,28024 | 0,1095672 | -2,47284 | 0,0431787 | 0,11545689 |
| SOD2 | -0,2808 | 0,2267275 | -4,88818 | 0,0018575 | 0,01911688 |
| Jagged1 | -0,28263 | -0,1270044 | -2,2353 | 0,0610854 | 0,14507791 |
| MIF | -0,29557 | 0,0525548 | -4,60683 | 0,0025659 | 0,02424311 |
| p70-S6K1 | -0,29743 | 0,0543159 | -4,58138 | 0,0026435 | 0,02424311 |
| eEF2K | -0,32344 | 0,0589671 | -2,17741 | 0,0664964 | 0,14931476 |
| PREX1 | -0,33267 | 0,1187313 | -3,05014 | 0,0189385 | 0,06981802 |
| ULK1_pS757 | -0,33353 | 0,2888177 | -3,99062 | 0,0054222 | 0,03348217 |
| P-Cadherin | -0,33443 | 0,0761427 | -2,94261 | 0,0220207 | 0,07770164 |
| PEA15 | -0,34867 | 0,0492564 | -2,18148 | 0,0661002 | 0,14931476 |
| VASP | -0,35052 | -0,6562173 | -2,5361 | 0,03939 | 0,11217854 |
| S6_pS235_S236 | -0,36186 | 0,6485896 | -1,32262 | 0,2281608 | 0,34363247 |
| Gys_pS641 | -0,36883 | 0,2498172 | -1,5819 | 0,1583572 | 0,27352611 |
| b-Actin | -0,37532 | -0,1764572 | -5,26931 | 0,0012205 | 0,01435583 |
| NF-kB-p65_pS536 | -0,40735 | -0,1849384 | -4,54176 | 0,0027694 | 0,02424311 |
| Merlin | -0,44466 | -0,3098276 | -4,07534 | 0,0048757 | 0,03273887 |
| VEGFR-2 | -0,44476 | -0,0645609 | -2,46791 | 0,0434894 | 0,11545689 |
| PAK4 | -0,45249 | -0,0012319 | -2,34393 | 0,0521064 | 0,13000281 |
| Atg7 | -0,45482 | 0,1900279 | -1,79011 | 0,1172291 | 0,22103502 |
| Glutaminase | -0,47371 | 0,2377846 | -2,51829 | 0,0404206 | 0,11217854 |
| LC3A-B | -0,47688 | 0,2071291 | -1,44397 | 0,1926244 | 0,30269376 |
| PMS2 | -0,48922 | 0,378151 | -2,08762 | 0,0758579 | 0,16435883 |
| TFRC | -0,49996 | 0,6679799 | -1,83278 | 0,1101638 | 0,20931118 |
| B-Raf | -0,50455 | 0,2097444 | -6,12404 | 5,10E-04 | 0,00740602 |
| TSC1 | -0,50695 | -0,0779737 | -1,56969 | 0,1611479 | 0,27641343 |
| p70S6K_pT389 | -0,52884 | 0,3247387 | -1,98594 | 0,0880579 | 0,17828111 |
| Heregulin | -0,57913 | 0,0224571 | -2,18899 | 0,0653759 | 0,14931476 |
| MYH11 | -0,66165 | 0,2888351 | -2,30342 | 0,055286 | 0,13520441 |
| HSP27_pS82 | -0,68487 | 0,2522355 | -4,0859 | 0,0048119 | 0,03273887 |
| FAK_pY397 | -0,6886 | 0,1568537 | -6,40614 | 3,89E-04 | 0,00656047 |
| PKC-alpha | -0,71319 | 0,1001862 | -13,2164 | 3,78E-06 | 2,34E-04 |
| Cyclin_B1 | -0,73721 | 0,5354757 | -2,85891 | 0,0247867 | 0,0838673 |
| E-Cadherin | -0,75573 | -1,1995603 | -1,49073 | 0,180314 | 0,29890978 |
| Cox-IV | -0,80271 | 0,5074037 | -3,0851 | 0,0180379 | 0,06854393 |
| PLK1 | -0,8331 | 0,494379 | -3,20507 | 0,0152797 | 0,05990612 |
| Myosin-IIa_pS1943 | -0,9402 | 0,4484787 | -3,38942 | 0,0118857 | 0,05132152 |
| TRIM25 | -0,9402 | 0,4484787 | -3,38942 | 0,0118857 | 0,05132152 |
| Gab2 | -1,14244 | -0,560432 | -3,57162 | 0,0093168 | 0,04341993 |
| p16_INK4a | -1,29666 | -0,6992205 | -5,94748 | 6,06E-04 | 0,00831592 |
| Cox2 | -1,35992 | -0,4638683 | -2,61733 | 0,0350245 | 0,10680296 |
| PAR | -1,45606 | 0,0704788 | -2,36477 | 0,0505441 | 0,12739176 |
| PDGFR-b | -1,75802 | 1,0202718 | -4,44228 | 0,0031161 | 0,02424311 |

**Subtype A vs. MGT2**

| Protein | logFC | AveExpr | t | P.Value | adj.P.Val |
|---|---|---|---|---|---|
| P-Met_pY1 | 3,637657 | -0,512787 | 7,504304 | 1,24E-05 | 4,37E-04 |
| E-Cadherin | 2,858419 | 2,07281 | 9,734716 | 1,02E-06 | 1,22E-04 |
| Claudin-7 | 1,849309 | 1,334111 | 7,172082 | 1,88E-05 | 4,90E-04 |
| NDRG1_pT | 1,59218 | 0,143565 | 4,439983 | 0,001011 | 0,0078003 |
| 53BP1 | 1,390249 | 0,675538 | 4,255926 | 0,001372 | 0,0089284 |
| HES1 | 1,388026 | 0,084085 | 3,031517 | 0,011495 | 0,0359412 |
| Src_pY527 | 1,271446 | -0,103271 | 5,51767 | 1,86E-04 | 0,0025465 |
| Bim | 1,1928 | 0,264333 | 3,052958 | 0,011065 | 0,0350382 |
| RSK | 0,955445 | -0,315825 | 7,968723 | 7,06E-06 | 3,26E-04 |
| Src_pY416 | 0,946306 | -0,174236 | 4,702663 | 6,59E-04 | 0,0060253 |
| 4E-BP1 | 0,931205 | 0,004343 | 4,626002 | 7,46E-04 | 0,0063553 |
| ARID1A | 0,902423 | 0,6773 | 2,706231 | 0,020532 | 0,0533842 |
| CDK1 | 0,841402 | 0,176039 | 2,594209 | 0,025065 | 0,0612974 |
| Akt_pT308 | 0,828394 | -0,134122 | 3,761853 | 0,003179 | 0,0153978 |
| p16_INK4a | 0,821494 | 0,167892 | 3,066624 | 0,010799 | 0,0350382 |
| VASP | 0,732469 | -0,102788 | 3,829014 | 0,002831 | 0,0145695 |
| Gab2 | 0,711829 | -0,028675 | 2,434552 | 0,033271 | 0,0760914 |
| IR-b | 0,692957 | 0,020946 | 3,352691 | 0,006502 | 0,024048 |
| PKC-b-II_p | 0,687287 | -0,232603 | 5,540261 | 1,79E-04 | 0,0025465 |
| HER3 | 0,684612 | 0,467884 | 3,693476 | 0,003579 | 0,0164203 |
| Rad50 | 0,638681 | 0,455258 | 3,340458 | 0,006644 | 0,0241119 |
| PDK1_pS2 | 0,62223 | 0,034216 | 8,196864 | 5,40E-06 | 3,26E-04 |
| Akt_pS473 | 0,603119 | -0,184994 | 2,678357 | 0,021578 | 0,0549461 |
| c-Jun_pS7 | 0,585643 | 0,276401 | 4,455335 | 9,85E-04 | 0,0078003 |
| 4E-BP1_pS | 0,535982 | 0,364639 | 3,84237 | 0,002767 | 0,0145413 |
| ACC1 | 0,510477 | 0,238597 | 2,540506 | 0,027575 | 0,0648665 |
| HER2_pY1 | 0,504194 | 0,063467 | 2,666076 | 0,022055 | 0,0555883 |
| FASN | 0,501761 | -7,63E-04 | 4,079585 | 0,001845 | 0,0109818 |
| C-Raf_pS3 | 0,500747 | 0,080728 | 3,308351 | 0,007032 | 0,0248118 |
| PAICS | 0,500277 | 0,310122 | 2,043257 | 0,065897 | 0,136744 |
| p38_pT18 | 0,481667 | 0,109864 | 6,322686 | 5,82E-05 | 0,0010269 |
| GCN5L2 | 0,47722 | 0,315373 | 3,78688 | 0,003045 | 0,0153479 |
| STING | 0,468437 | 0,114471 | 0,704499 | 0,495859 | 0,612386 |
| PARP | 0,461149 | 0,302319 | 3,968254 | 0,00223 | 0,0122401 |
| eIF4E | 0,451018 | -0,007555 | 4,968384 | 4,31E-04 | 0,004632 |
| PDK1 | 0,445608 | -0,066556 | 7,13043 | 1,98E-05 | 4,90E-04 |
| Caspase-3 | 0,438372 | 0,152172 | 5,332925 | 2,45E-04 | 0,0031884 |
| 14-3-3_zet | 0,423332 | -0,017049 | 3,691773 | 0,00359 | 0,0164203 |
| Paxillin | 0,41394 | -0,022313 | 6,874556 | 2,76E-05 | 6,21E-04 |
| FAK | 0,407971 | -0,029242 | 5,123623 | 3,38E-04 | 0,004157 |
| Histone-H3 | 0,385838 | 0,054322 | 2,196682 | 0,050528 | 0,1114322 |
| p90RSK_pT | 0,378022 | 0,017034 | 3,531836 | 0,004745 | 0,01947 |
| Bax | 0,368331 | -0,020412 | 0,826717 | 0,426078 | 0,5396982 |
| Bcl-xL | 0,364332 | 0,192799 | 2,719429 | 0,020055 | 0,0526977 |
| Jagged1 | 0,35794 | 0,087936 | 1,749995 | 0,1081 | 0,1964232 |
| c-Myc | 0,357068 | 0,281863 | 3,662234 | 0,003779 | 0,016971 |
| Shc_pY317 | 0,338664 | -0,203854 | 3,023125 | 0,011668 | 0,0360265 |
| Caveolin-1 | 0,334515 | -0,106836 | 0,950612 | 0,362349 | 0,4775705 |
| RBM15 | 0,334174 | 0,521032 | 0,624377 | 0,545193 | 0,6474162 |
| PRAS40_pT | 0,323953 | -0,104174 | 1,103932 | 0,293333 | 0,4188231 |
| WIPI1 | 0,304193 | 0,243406 | 1,358826 | 0,201582 | 0,317138 |
| TUFM | 0,300465 | 0,120711 | 3,217371 | 0,008261 | 0,0283398 |
| beta-Caten | 0,289844 | 0,309686 | 1,608168 | 0,136277 | 0,2404317 |
| ER-alpha_p | 0,286177 | 0,181252 | 4,07261 | 0,001867 | 0,0109818 |
| c-Kit | 0,279571 | 0,222288 | 4,388641 | 0,0011 | 0,0079908 |
| HSP70 | 0,27151 | 0,071262 | 3,377812 | 0,00622 | 0,0240053 |
| DJ-1 | 0,265733 | 0,139229 | 2,994441 | 0,01228 | 0,0369909 |
| RPA32_pS | 0,261734 | -0,005264 | 2,862701 | 0,015532 | 0,044098 |
| PLC-gamm | 0,260835 | -0,021192 | 3,350843 | 0,006523 | 0,024048 |
| ACC_pS79 | 0,251971 | 0,126956 | 0,670382 | 0,516529 | 0,6254047 |
| MSH6 | 0,246726 | 0,140975 | 1,52971 | 0,154501 | 0,2651031 |
| b-Actin | 0,245096 | 0,147224 | 1,529492 | 0,154554 | 0,2651031 |
| AMPK_alph | 0,241763 | -0,133322 | 2,038531 | 0,066434 | 0,136744 |
| Bid | 0,241507 | 0,180387 | 1,811547 | 0,097594 | 0,1824368 |
| eIF4G | 0,239594 | 0,049167 | 3,538797 | 0,004687 | 0,01947 |
| Bad_pS112 | 0,238065 | -0,152378 | 2,575245 | 0,025924 | 0,0627778 |

**Subtype B vs. MGT 2**

| Protein | logFC | AveExpr | t | P.Value | adj.P.Val |
|---|---|---|---|---|---|
| P-Met_pY1 | 4,429137 | -0,83157 | 57,94388 | 2,68E-10 | 6,62E-08 |
| HES1 | 2,033252 | 0,168379 | 33,24106 | 1,09E-08 | 9,01E-07 |
| Notch1 | 1,710942 | 0,731063 | 31,59293 | 1,54E-08 | 9,49E-07 |
| Fibronectir | 1,599821 | 1,178052 | 9,284537 | 4,58E-05 | 4,52E-04 |
| IGFRb | 1,571705 | 0,517034 | 24,26581 | 8,87E-08 | 3,13E-06 |
| MCT4 | 1,567953 | 0,191969 | 17,06514 | 9,06E-07 | 1,97E-05 |
| RSK | 1,545585 | -0,170202 | 29,36662 | 2,50E-08 | 1,23E-06 |
| Akt_pT308 | 1,195723 | -0,094465 | 13,88509 | 3,49E-06 | 5,74E-05 |
| Akt_pS473 | 1,162458 | 0,01902 | 12,2847 | 7,70E-06 | 1,06E-04 |
| NDRG1_pT | 1,109908 | -0,493183 | 19,43433 | 3,85E-07 | 1,06E-05 |
| Src_pY416 | 1,004716 | -0,345657 | 16,91783 | 9,59E-07 | 1,97E-05 |
| PKA-a | 0,945893 | 0,458861 | 16,1482 | 1,30E-06 | 2,47E-05 |
| IR-b | 0,875025 | -0,021004 | 6,686059 | 3,40E-04 | 0,0020021 |
| Bim | 0,834885 | -0,210663 | 10,30311 | 2,37E-05 | 2,93E-04 |
| STING | 0,812935 | 0,218965 | 8,125268 | 1,05E-04 | 9,25E-04 |
| Src_pY527 | 0,798604 | -0,664382 | 7,787181 | 1,36E-04 | 0,0010837 |
| CDK1 | 0,759937 | -0,056419 | 12,81882 | 5,85E-06 | 9,03E-05 |
| LRP6_pS14 | 0,702505 | 0,268665 | 11,88844 | 9,51E-06 | 1,24E-04 |
| PKC-b-II_p | 0,693778 | -0,378662 | 9,59041 | 3,73E-05 | 4,39E-04 |
| ACC_pS79 | 0,693151 | 0,336689 | 7,578502 | 1,60E-04 | 0,0012387 |
| AMPK_pT1 | 0,684977 | 0,918582 | 4,163988 | 0,004651 | 0,0149183 |
| SLC1A5 | 0,662634 | 0,316106 | 7,351803 | 1,93E-04 | 0,0013138 |
| Caveolin-1 | 0,625031 | -0,005511 | 6,850821 | 2,95E-04 | 0,0018659 |
| Shc_pY317 | 0,579833 | -0,133043 | 5,338815 | 0,001242 | 0,0061373 |
| HER2_pY1 | 0,571485 | -0,006164 | 8,016517 | 1,14E-04 | 9,50E-04 |
| YAP_pS127 | 0,550479 | 0,007843 | 9,450377 | 4,10E-05 | 4,52E-04 |
| ERCC5 | 0,506844 | -0,006689 | 4,430638 | 0,003387 | 0,0124122 |
| ACC1 | 0,498117 | 0,119805 | 6,114026 | 5,75E-04 | 0,0032184 |
| IGF1R_pY1 | 0,494727 | 0,152044 | 7,286968 | 2,04E-04 | 0,0013228 |
| PRAS40_pT | 0,461013 | -0,092619 | 7,99944 | 1,15E-04 | 9,50E-04 |
| Tuberin | 0,455819 | 0,202501 | 5,167735 | 0,001488 | 0,0068051 |
| FASN | 0,452651 | -0,139704 | 5,22551 | 0,001399 | 0,0065509 |
| MAPK_pT2 | 0,433972 | -0,028119 | 5,953306 | 6,71E-04 | 0,0035067 |
| Rb_pS807_ | 0,424155 | 0,468813 | 7,429753 | 1,81E-04 | 0,0013065 |
| Paxillin | 0,420426 | -0,108736 | 6,765916 | 0,000317 | 0,0019426 |
| 4E-BP1 | 0,409332 | -0,511953 | 6,093394 | 5,86E-04 | 0,0032184 |
| Bad_pS112 | 0,388397 | -0,114121 | 3,819178 | 0,007119 | 0,0204478 |
| AMPK_alph | 0,379505 | -0,103425 | 3,568891 | 0,009808 | 0,0259636 |
| PLC-gamm | 0,376092 | -0,008946 | 4,423101 | 0,003417 | 0,0124122 |
| GSK-3a-b_ | 0,371556 | 0,012234 | 5,765962 | 8,05E-04 | 0,0040593 |
| Pdcd4 | 0,358724 | 0,30753 | 4,945374 | 0,001892 | 0,0080555 |
| p21 | 0,354248 | -0,093737 | 6,182004 | 5,39E-04 | 0,0030976 |
| COG3 | 0,347162 | 0,050711 | 5,221219 | 0,001406 | 0,0065509 |
| 14-3-3_zet | 0,340484 | -0,159122 | 5,936645 | 6,81E-04 | 0,0035067 |
| p90RSK_pT | 0,324632 | -0,097478 | 3,609566 | 0,009304 | 0,0255765 |
| PDK1_pS2 | 0,321764 | -0,281823 | 4,349438 | 0,003726 | 0,0129631 |
| WIPI1 | 0,304937 | 0,177482 | 4,349768 | 0,003725 | 0,0129631 |
| FAK | 0,297001 | -0,184835 | 5,008786 | 0,001765 | 0,007649 |
| Bax | 0,294233 | -0,145235 | 5,048647 | 0,001691 | 0,0074564 |
| Caspase-7_ | 0,293288 | -0,035883 | 2,760869 | 0,029328 | 0,0619152 |
| WIPI2 | 0,265893 | 0,088879 | 1,838137 | 0,110552 | 0,1870289 |
| PDK1 | 0,263278 | -0,273178 | 3,876257 | 0,006627 | 0,0197203 |
| GATA6 | 0,252854 | -0,001738 | 3,563194 | 0,009881 | 0,0259636 |
| eIF4E | 0,251234 | -0,225829 | 4,329857 | 0,003814 | 0,0130825 |
| Caspase-3 | 0,249663 | -0,056699 | 3,822671 | 0,007088 | 0,0204478 |
| YB1_pS102 | 0,247645 | -0,039208 | 3,393642 | 0,012343 | 0,030293 |
| UBAC1 | 0,237905 | -0,064641 | 4,115195 | 0,004934 | 0,0154269 |
| p38_pT18 | 0,217278 | -0,15386 | 2,938324 | 0,022889 | 0,0504783 |
| Myt1 | 0,209453 | -0,049406 | 3,66189 | 0,008698 | 0,024413 |
| XRCC1 | 0,207335 | 0,165378 | 2,451464 | 0,045538 | 0,086523 |
| Bid | 0,205877 | 0,106317 | 2,356773 | 0,052183 | 0,0969118 |
| mTOR | 0,205357 | 0,006248 | 3,857133 | 0,006787 | 0,0199584 |
| Rictor_pT1 | 0,204619 | 0,028295 | 3,573809 | 0,009746 | 0,0259636 |
| Histone-H3 | 0,203684 | -0,139153 | 1,838476 | 0,110497 | 0,1870289 |
| Chk2_pT68 | 0,203435 | 0,012562 | 3,233 | 0,015298 | 0,0363328 |
| HER3_pY1 | 0,191942 | -0,098544 | 3,108271 | 0,018117 | 0,0414349 |

| | | | | | |
|---|---|---|---|---|---|
| p53 | 0,234861 | -0,028158 | 1,717783 | 0,113997 | 0,2055273 |
| Elk1_pS383 | 0,223288 | -0,027692 | 3,153167 | 0,009258 | 0,0304905 |
| EGFR | 0,218885 | -0,060514 | 3,053962 | 0,011045 | 0,0350382 |
| ERCC5 | 0,21579 | -0,13424 | 2,001351 | 0,070806 | 0,1432118 |
| SLC1A5 | 0,214218 | 0,093795 | 0,925357 | 0,37476 | 0,4872195 |
| TSC1 | 0,209308 | -0,236983 | 1,440225 | 0,177827 | 0,2889694 |
| Smad3 | 0,208068 | 0,189307 | 3,351871 | 0,006511 | 0,024048 |
| Lck | 0,206677 | 0,110104 | 1,586843 | 0,141034 | 0,2470589 |
| Merlin | 0,203843 | 0,100168 | 1,149809 | 0,27476 | 0,4057239 |
| HER3_pY12 | 0,203492 | -0,047216 | 2,946901 | 0,013366 | 0,0393039 |
| Bak | 0,193494 | -0,012915 | 1,976923 | 0,073823 | 0,1461908 |
| Myt1 | 0,193394 | -0,016846 | 2,059608 | 0,06407 | 0,1341125 |
| NAPSIN-A | 0,193247 | -0,08934 | 2,482779 | 0,030548 | 0,0705178 |
| UBAC1 | 0,189955 | -0,051966 | 1,969361 | 0,074781 | 0,1465416 |
| Connexin-4 | 0,18699 | 0,477773 | 0,749889 | 0,469156 | 0,5852598 |
| EGFR_pY11 | 0,182097 | -0,013826 | 3,197102 | 0,008563 | 0,028975 |
| Aurora-B | 0,17813 | -0,041328 | 1,975649 | 0,073983 | 0,1461908 |
| PAK4 | 0,174688 | 0,016478 | 1,855111 | 0,09073 | 0,1723861 |
| GSK-3a-b_ | 0,174242 | -0,068138 | 0,819201 | 0,430172 | 0,5421047 |
| MAPK_pT2 | 0,173967 | -0,146165 | 0,912501 | 0,381192 | 0,4929548 |
| ATR_pS428 | 0,173213 | 0,029853 | 2,740563 | 0,019313 | 0,0515209 |
| Notch3 | 0,172453 | 0,041833 | 2,738095 | 0,019399 | 0,0515209 |
| VEGFR-2 | 0,161657 | -0,036295 | 0,967953 | 0,353999 | 0,4751501 |
| MEK1_pS2 | 0,160906 | 0,032162 | 2,643838 | 0,022946 | 0,0572498 |
| b-Catenin_ | 0,158045 | 0,041754 | 1,869614 | 0,088544 | 0,1695382 |
| Notch1 | 0,156847 | -0,167172 | 0,700098 | 0,498497 | 0,6125812 |
| Caspase-7_ | 0,155075 | -0,084976 | 1,138371 | 0,279303 | 0,4057239 |
| TAZ | 0,151172 | 0,185789 | 1,807623 | 0,098235 | 0,1824368 |
| IGF1R_pY1 | 0,148883 | -0,022979 | 2,229583 | 0,047709 | 0,106625 |
| Chk2_pT68 | 0,14549 | 0,009538 | 1,474399 | 0,168586 | 0,2794674 |
| Rictor | 0,143098 | 0,019462 | 1,386335 | 0,193273 | 0,3079892 |
| Tuberin | 0,139159 | 0,042867 | 1,276209 | 0,228333 | 0,3481377 |
| Cyclin_D1 | 0,129665 | 0,001313 | 1,650296 | 0,127294 | 0,2261986 |
| TIGAR | 0,129574 | 0,043638 | 1,09703 | 0,296209 | 0,4188231 |
| YB1_pS102 | 0,12957 | -0,081783 | 1,285143 | 0,225307 | 0,3461398 |
| Glutamate- | 0,127577 | -0,021805 | 1,417977 | 0,184073 | 0,2971628 |
| IRS1 | 0,116085 | 0,34378 | 0,316469 | 0,757609 | 0,8075234 |
| AMPK-a2_ | 0,11558 | -0,085886 | 1,095769 | 0,296737 | 0,4188231 |
| PEA15 | 0,11237 | 0,139538 | 0,833019 | 0,422664 | 0,5396982 |
| LC3A-B | 0,109187 | -0,058193 | 1,065715 | 0,309528 | 0,4319408 |
| PEA15_pS1 | 0,101697 | -0,008363 | 0,556717 | 0,588938 | 0,686168 |
| p27_pT198 | 0,101498 | 0,106837 | 1,053753 | 0,314734 | 0,4355815 |
| eEF2K | 0,100409 | 0,15178 | 0,451722 | 0,660302 | 0,7379841 |
| FOXO3a_p | 0,094894 | -0,075844 | 1,823714 | 0,095631 | 0,1803112 |
| JNK_pT183 | 0,093291 | 0,025704 | 0,959186 | 0,358203 | 0,4756777 |
| p27-Kip-1 | 0,090625 | -0,052819 | 1,913175 | 0,082268 | 0,158752 |
| Akt | 0,089004 | -0,093046 | 0,474568 | 0,644435 | 0,7235247 |
| WIPI2 | 0,076508 | -0,008059 | 0,424551 | 0,6794 | 0,7491602 |
| YAP_pS127 | 0,074437 | -0,261542 | 0,418235 | 0,683875 | 0,7507427 |
| DM-K9-His | 0,070306 | 0,013084 | 1,135106 | 0,28061 | 0,4057239 |
| Stathmin | 0,063385 | -0,09865 | 0,67448 | 0,514019 | 0,6254047 |
| Mnk1 | 0,061391 | 0,099689 | 0,508216 | 0,621402 | 0,7124215 |
| p38_MAPK | 0,056059 | 0,026224 | 0,387848 | 0,705576 | 0,7685687 |
| PKM2 | 0,046362 | -0,106769 | 0,241959 | 0,813292 | 0,8548214 |
| mTOR | 0,045241 | -0,079951 | 0,426884 | 0,677751 | 0,7491602 |
| Wee1_pS6 | 0,045056 | 0,207071 | 0,643198 | 0,533358 | 0,6395114 |
| PREX1 | 0,044776 | 0,175763 | 0,55076 | 0,592877 | 0,6875148 |
| IGFRb | 0,044164 | -0,389855 | 0,427417 | 0,677375 | 0,7491602 |
| MDM2_pS | 0,03625 | -0,166058 | 0,330196 | 0,747494 | 0,8062492 |
| p21 | 0,029076 | -0,282497 | 0,317732 | 0,756676 | 0,8075234 |
| cIAP | 0,025597 | -0,050772 | 0,50585 | 0,623008 | 0,7124215 |
| Tyro3 | 0,016713 | 0,013275 | 0,306917 | 0,764675 | 0,8075234 |
| XRCC1 | 0,014632 | 0,052949 | 0,148387 | 0,884739 | 0,9143541 |
| U-Histone- | 0,010909 | -0,015202 | 0,117134 | 0,908879 | 0,9315062 |
| MCT4 | 0,004452 | -0,74516 | 0,017185 | 0,986599 | 0,992555 |
| Rictor_pT1 | 0,001608 | -0,09316 | 0,01033 | 0,991944 | 0,992555 |
| Rad51 | -0,00124 | 0,207918 | -0,00955 | 0,992555 | 0,992555 |
| Rb_pS807_ | -0,00913 | 0,206846 | -0,03008 | 0,976542 | 0,9885487 |
| LRP6_pS14 | -0,01096 | -0,161801 | -0,1418 | 0,889819 | 0,915772 |

| | | | | | |
|---|---|---|---|---|---|
| PKM2 | 0,189979 | -0,030715 | 2,921563 | 0,023427 | 0,0512087 |
| RIP | 0,187817 | 0,216472 | 1,363352 | 0,216825 | 0,3113704 |
| Gys | 0,185007 | 0,229644 | 3,15264 | 0,017055 | 0,0393704 |
| PAICS | 0,18402 | 0,011217 | 1,71356 | 0,132265 | 0,21214 |
| Bak | 0,18062 | -0,062856 | 1,652588 | 0,144328 | 0,2270633 |
| Elk1_pS383 | 0,175146 | -0,105295 | 2,191381 | 0,066274 | 0,1186205 |
| STAT5-alph | 0,173442 | 0,775827 | 3,072549 | 0,019024 | 0,0431092 |
| Stat3 | 0,173035 | 0,403483 | 3,030139 | 0,020164 | 0,0452767 |
| Mcl-1 | 0,169579 | 0,479658 | 2,567735 | 0,038559 | 0,0767051 |
| FOXO3a_p | 0,15269 | -0,06187 | 2,656238 | 0,033999 | 0,0694039 |
| Stat3_pY70 | 0,15226 | 0,240467 | 2,857044 | 0,025629 | 0,0550477 |
| EGFR_pY11 | 0,150863 | -0,072296 | 2,273822 | 0,058821 | 0,1068297 |
| S6_pS240_ | 0,1319 | 0,442304 | 1,231873 | 0,259488 | 0,3580641 |
| ATR_pS428 | 0,130777 | -0,033401 | 1,752501 | 0,125071 | 0,2059507 |
| Akt | 0,130338 | -0,087664 | 2,386733 | 0,049979 | 0,0935216 |
| p53 | 0,127715 | -0,143689 | 1,597836 | 0,156039 | 0,2423994 |
| Aurora-B | 0,120157 | -0,114976 | 1,811734 | 0,114845 | 0,1916666 |
| CD20 | 0,11635 | 0,092588 | 1,985751 | 0,089289 | 0,1553125 |
| Stathmin | 0,115549 | -0,081181 | 1,728994 | 0,129368 | 0,2102227 |
| mTOR_pS2 | 0,114578 | 0,01004 | 1,580779 | 0,159865 | 0,2467916 |
| Hexokinase | 0,094923 | 0,04622 | 1,717849 | 0,131454 | 0,21214 |
| SDHA | 0,093427 | 0,182789 | 1,170325 | 0,281823 | 0,3762714 |
| Lck | 0,08686 | -0,00688 | 1,189175 | 0,274816 | 0,370927 |
| U-Histone- | 0,082333 | 0,025272 | 1,155404 | 0,287475 | 0,3817538 |
| LC3A-B | 0,082153 | -0,098236 | 1,329239 | 0,227257 | 0,3226006 |
| MEK1 | 0,069103 | 0,359646 | 0,978753 | 0,361722 | 0,4600599 |
| Tyro3 | 0,065279 | 0,038768 | 1,218976 | 0,264039 | 0,3603457 |
| p44-42-MA | 0,06256 | 0,020878 | 1,091774 | 0,312642 | 0,4085853 |
| Wee1 | 0,057608 | 0,318637 | 0,960062 | 0,370391 | 0,4667683 |
| S6_pS235_ | 0,051162 | 0,42135 | 0,330397 | 0,751193 | 0,8246435 |
| NAPSIN-A | 0,049716 | -0,217621 | 0,466318 | 0,655773 | 0,7430087 |
| MERIT40_ | 0,048588 | 0,087179 | 0,412583 | 0,692798 | 0,7813746 |
| TSC1 | 0,040575 | -0,38389 | 0,717029 | 0,497592 | 0,5995378 |
| c-Abl | 0,040335 | 0,046742 | 0,63463 | 0,546726 | 0,6523741 |
| Collagen_V | 0,040136 | 0,187521 | 0,356881 | 0,732159 | 0,8120777 |
| cIAP | 0,037947 | -0,048946 | 0,566057 | 0,589807 | 0,6904381 |
| TUFM | 0,037279 | -0,102757 | 0,355465 | 0,733171 | 0,8120777 |
| MEK1_pS2 | 0,036383 | -0,077658 | 0,55098 | 0,599535 | 0,6985145 |
| b-Catenin_ | 0,02861 | -0,070389 | 0,263338 | 0,800207 | 0,8519445 |
| DJ-1 | 0,028009 | -0,061384 | 0,382096 | 0,714224 | 0,8018785 |
| ER-alpha_ | 0,022935 | -0,039132 | 0,296226 | 0,776031 | 0,8336778 |
| Gys_pS641 | 0,021184 | 0,030585 | 0,369732 | 0,722994 | 0,8080525 |
| c-Kit | 0,020027 | 0,005564 | 0,295583 | 0,776501 | 0,8336778 |
| PARP | 0,017185 | -0,064674 | 0,149695 | 0,885413 | 0,904904 |
| YAP | 0,013803 | -0,006171 | 0,246721 | 0,812515 | 0,8576546 |
| PR | 0,010656 | 0,24656 | 0,198898 | 0,848245 | 0,876638 |
| eIF4G | -2,47E-04 | -0,147012 | -0,00395 | 0,99696 | 0,99696 |
| Bcl-xL | -0,00368 | -0,107499 | -0,06513 | 0,949972 | 0,9577273 |
| MDM2_pS | -0,00591 | -0,199264 | -0,05304 | 0,959248 | 0,9631474 |
| PDHK1 | -0,00776 | 0,058109 | -0,12715 | 0,902555 | 0,9174122 |
| TIGAR | -0,00856 | -0,067512 | -0,14815 | 0,886586 | 0,904904 |
| DM-K9-His | -0,0101 | -0,050499 | -0,15294 | 0,882953 | 0,904904 |
| Mnk1 | -0,01085 | 0,042951 | -0,11676 | 0,910477 | 0,9216714 |
| p38_MAPK | -0,01454 | -0,028369 | -0,20919 | 0,840521 | 0,8723052 |
| XPF | -0,01589 | 0,009206 | -0,2963 | 0,775978 | 0,8336778 |
| PAK1 | -0,01791 | -0,004488 | -0,30113 | 0,77245 | 0,8336778 |
| JNK_pT183 | -0,01948 | -0,062312 | -0,34436 | 0,74113 | 0,8172287 |
| p27_pT198 | -0,02014 | 0,01171 | -0,25504 | 0,806344 | 0,8547941 |
| FoxM1 | -0,02167 | 0,036726 | -0,24184 | 0,816141 | 0,8578158 |
| 4E-BP1_pS | -0,02178 | -0,086957 | -0,2319 | 0,823542 | 0,8619269 |
| LDHA | -0,02473 | 0,306928 | -0,22367 | 0,829685 | 0,8646926 |
| TFAM | -0,02557 | 0,215198 | -0,29125 | 0,779674 | 0,8336778 |
| Notch3 | -0,02558 | -0,114612 | -0,32481 | 0,755233 | 0,8254095 |
| Atg7 | -0,02732 | -0,058942 | -0,499 | 0,633749 | 0,734911 |
| G6PD | -0,02819 | 0,018532 | -0,47475 | 0,650055 | 0,7430087 |
| Rab25 | -0,02846 | -0,058297 | -0,46709 | 0,655251 | 0,7430087 |
| C-Raf_pS33 | -0,0308 | -0,347453 | -0,4849 | 0,643199 | 0,7423844 |
| B7-H4 | -0,03884 | 0,068568 | -0,70454 | 0,504849 | 0,6053288 |
| PI3K-p110- | -0,04162 | 0,080436 | -0,76123 | 0,472466 | 0,5748723 |

| | | | | | |
|---|---|---|---|---|---|
| PAK1 | -0,0136 | -0,004866 | -0,04448 | 0,965322 | 0,9812126 |
| Rab11 | -0,01981 | -0,035245 | -0,1721 | 0,866507 | 0,9030688 |
| P-Cadherin | -0,02388 | 0,057863 | -0,30645 | 0,765022 | 0,8075234 |
| GCLM | -0,02593 | 0,020193 | -0,50009 | 0,626924 | 0,713596 |
| p70-S6K1 | -0,02627 | 0,111673 | -0,21259 | 0,835559 | 0,8745042 |
| MMP2 | -0,02702 | 0,049248 | -0,55704 | 0,588724 | 0,686168 |
| YAP | -0,02796 | -0,037328 | -0,37465 | 0,715085 | 0,7746759 |
| JNK2 | -0,03743 | 0,004678 | -0,38679 | 0,706336 | 0,7685687 |
| Atg7 | -0,04046 | -0,075657 | -0,30686 | 0,764719 | 0,8075234 |
| Chk1_pS29 | -0,04384 | -0,021965 | -0,73728 | 0,476481 | 0,5914106 |
| Hexokinase | -0,0487 | -0,050583 | -0,80062 | 0,440407 | 0,5521855 |
| Ets-1 | -0,04892 | 0,060662 | -0,97684 | 0,349774 | 0,4746931 |
| Creb | -0,05367 | 0,035296 | -1,02208 | 0,328831 | 0,4487363 |
| Cox2 | -0,05874 | 0,957895 | -0,08284 | 0,935477 | 0,9548053 |
| PKA-a | -0,05969 | -0,157512 | -0,4941 | 0,631013 | 0,7149549 |
| FAK_pY397 | -0,06338 | 0,253015 | -0,15167 | 0,882213 | 0,9143541 |
| PMS2 | -0,07143 | 0,147057 | -0,59999 | 0,560745 | 0,6626989 |
| N-Cadherin | -0,07154 | -0,080749 | -0,47941 | 0,641096 | 0,7230625 |
| Smad1 | -0,07287 | 0,032616 | -0,65233 | 0,527671 | 0,6357796 |
| CD20 | -0,08026 | -0,042893 | -1,05163 | 0,315664 | 0,4355815 |
| NF-kB-p65_ | -0,08295 | 0,032383 | -0,56537 | 0,583242 | 0,6860037 |
| mTOR_pS2 | -0,08552 | -0,128678 | -0,52021 | 0,61329 | 0,7078633 |
| Slfn11 | -0,08637 | -0,078558 | -0,96595 | 0,354957 | 0,4751501 |
| GATA6 | -0,08797 | -0,225422 | -0,96402 | 0,355882 | 0,4751501 |
| Rab25 | -0,08902 | -0,114058 | -1,14255 | 0,277638 | 0,4057239 |
| Jak2 | -0,09052 | -0,008305 | -1,16934 | 0,267136 | 0,3974848 |
| IRF-1 | -0,09924 | 0,056109 | -1,47526 | 0,168358 | 0,2794674 |
| D-a-Tubulin | -0,10018 | -0,031204 | -1,40656 | 0,187348 | 0,3004864 |
| eEF2 | -0,10324 | 0,116836 | -0,63082 | 0,541127 | 0,6456925 |
| Tuberin_pT | -0,10398 | -0,006469 | -1,31947 | 0,213982 | 0,3324126 |
| ATM_pS19 | -0,10456 | -0,039159 | -2,17966 | 0,052048 | 0,1137689 |
| Raptor | -0,10909 | 0,047014 | -1,25124 | 0,236964 | 0,3590808 |
| c-Abl | -0,10954 | -0,067083 | -1,96493 | 0,075347 | 0,1465416 |
| ZAP-70 | -0,10962 | 0,048859 | -1,13442 | 0,280886 | 0,4057239 |
| Cdc2_pY15 | -0,11392 | 0,128145 | -1,08775 | 0,300112 | 0,4211795 |
| MERIT40_p | -0,11948 | -0,039726 | -1,17769 | 0,263926 | 0,3950895 |
| OCT4 | -0,12349 | 0,032758 | -2,62763 | 0,023618 | 0,0583365 |
| eIF4E_pS2 | -0,12707 | 0,046931 | -2,2271 | 0,047917 | 0,106625 |
| XPF | -0,13373 | -0,090681 | -2,36272 | 0,037769 | 0,0855867 |
| COG3 | -0,13669 | -0,269422 | -0,82924 | 0,424707 | 0,5396982 |
| TFAM | -0,13704 | 0,118415 | -0,8328 | 0,422781 | 0,5396982 |
| cdc25C | -0,13826 | 0,142199 | -0,92531 | 0,374784 | 0,4872195 |
| BRD4 | -0,1446 | -0,06714 | -2,08815 | 0,060995 | 0,1289559 |
| PI3K-p110- | -0,14952 | -0,016923 | -3,41611 | 0,005814 | 0,0231627 |
| FoxM1 | -0,15471 | -0,076849 | -1,74971 | 0,108152 | 0,1964232 |
| C-Raf | -0,15709 | -0,011436 | -1,46843 | 0,170171 | 0,2802145 |
| PKC-delta_ | -0,15902 | -0,050856 | -3,52424 | 0,004808 | 0,01947 |
| 14-3-3_bet | -0,17231 | -0,020808 | -2,84679 | 0,01598 | 0,0448519 |
| B7-H4 | -0,17286 | -0,049555 | -2,91179 | 0,01423 | 0,0413515 |
| RIP | -0,17423 | -0,038769 | -0,94825 | 0,363495 | 0,4775705 |
| MEK1 | -0,1799 | 0,170994 | -1,37431 | 0,196868 | 0,3117081 |
| DUSP4 | -0,18444 | 0,128881 | -1,47938 | 0,167273 | 0,2794674 |
| INPP4b | -0,19089 | -0,024663 | -3,62402 | 0,004039 | 0,0178147 |
| p44-42-MA | -0,19152 | -0,17336 | -1,28421 | 0,225621 | 0,3461398 |
| SHP-2_pY5 | -0,19254 | -0,09438 | -2,0873 | 0,061084 | 0,1289559 |
| CD4 | -0,20334 | 0,03085 | -2,55706 | 0,026776 | 0,063593 |
| CD134 | -0,20466 | 0,02215 | -2,95002 | 0,013292 | 0,0393039 |
| MYH11 | -0,20505 | -0,080303 | -1,09766 | 0,295948 | 0,4188231 |
| DM-Histon | -0,21518 | 0,048536 | -2,76747 | 0,018408 | 0,0505211 |
| ER-alpha | -0,21616 | 0,056836 | -3,38394 | 0,006153 | 0,0240053 |
| MIF | -0,21648 | -0,062072 | -2,56651 | 0,02633 | 0,0631408 |
| G6PD | -0,21778 | -0,142741 | -1,51267 | 0,158728 | 0,2703856 |
| A-Raf | -0,21779 | 0,051285 | -1,71144 | 0,115192 | 0,2061771 |
| Wee1 | -0,21884 | 0,105018 | -1,02382 | 0,328047 | 0,4487363 |
| PD-L1 | -0,21999 | -0,036177 | -4,263 | 0,001355 | 0,0089284 |
| IGFBP2 | -0,2265 | -0,005399 | -1,78229 | 0,102466 | 0,1888741 |
| Beclin | -0,23185 | 0,016681 | -3,53411 | 0,004726 | 0,01947 |
| FoxO3a | -0,23736 | -6,74E-04 | -2,86391 | 0,015499 | 0,044098 |
| XBP1 | -0,24793 | 0,016132 | -4,894 | 4,85E-04 | 0,0049924 |

| | | | | | |
|---|---|---|---|---|---|
| cdc25C | -0,04292 | 0,229565 | -0,47012 | 0,65319 | 0,7430087 |
| ATM_pS19 | -0,04503 | 0,019368 | -0,75503 | 0,475937 | 0,5762568 |
| N-Cadherin | -0,05146 | -0,053096 | -0,97553 | 0,363205 | 0,4600599 |
| ZAP-70 | -0,05551 | 0,105245 | -0,78236 | 0,460763 | 0,5634078 |
| MSH6 | -0,05754 | -0,095416 | -0,59024 | 0,574397 | 0,6779627 |
| beta-Caten | -0,05824 | 0,037597 | -0,58706 | 0,576405 | 0,6779627 |
| Cyclin_D1 | -0,0589 | -0,140117 | -0,90985 | 0,394464 | 0,4920839 |
| Glutamate- | -0,05912 | -0,161659 | -1,01008 | 0,347548 | 0,444789 |
| Chk1_pS29 | -0,06366 | -0,024299 | -0,88828 | 0,405158 | 0,5018846 |
| JNK2 | -0,0644 | -0,003338 | -1,11781 | 0,302133 | 0,3990735 |
| Raptor | -0,07018 | 0,094162 | -1,1847 | 0,276466 | 0,371125 |
| CD134 | -0,07033 | 0,1474 | -1,07888 | 0,317957 | 0,413344 |
| DM-Histon | -0,07749 | 0,178097 | -1,21892 | 0,264059 | 0,3603457 |
| Tuberin_pT | -0,08047 | 0,030326 | -1,523 | 0,173486 | 0,2628894 |
| GCLM | -0,08184 | -0,007694 | -1,31115 | 0,232966 | 0,3288146 |
| Creb | -0,08273 | 0,029571 | -1,46449 | 0,188359 | 0,2819681 |
| Jagged1 | -0,08358 | -0,255073 | -1,56374 | 0,163774 | 0,2498952 |
| OCT4 | -0,08438 | 0,083166 | -1,61517 | 0,152237 | 0,2379901 |
| MSI2 | -0,08455 | 0,189506 | -1,42272 | 0,199681 | 0,2918411 |
| p27-Kip-1 | -0,08616 | -0,17866 | -1,44399 | 0,19384 | 0,2864007 |
| HER3 | -0,08715 | -0,144546 | -1,5632 | 0,163899 | 0,2498952 |
| IGFBP2 | -0,08762 | 0,127348 | -0,93491 | 0,382307 | 0,4793395 |
| Slfn11 | -0,09708 | -0,066141 | -1,05563 | 0,327722 | 0,4216005 |
| c-Jun_pS73 | -0,09713 | -0,261039 | -1,35224 | 0,220177 | 0,3143565 |
| Rictor | -0,10149 | -0,15851 | -1,69216 | 0,136386 | 0,2173371 |
| BRD4 | -0,10243 | -0,010294 | -1,44056 | 0,194772 | 0,2864007 |
| TFRC | -0,10351 | 0,423692 | -0,59613 | 0,570677 | 0,6776791 |
| PKC-delta_ | -0,10618 | 0,01554 | -1,94196 | 0,095138 | 0,1631877 |
| PMS2 | -0,1069 | 0,14136 | -1,28062 | 0,242886 | 0,3389432 |
| Jak2 | -0,10903 | 3,33E-04 | -1,77367 | 0,12132 | 0,2011138 |
| p70S6K_pT | -0,10932 | 0,066268 | -0,88584 | 0,406384 | 0,5018846 |
| Smad3 | -0,11093 | -0,04749 | -1,06217 | 0,324952 | 0,4202255 |
| GCN5L2 | -0,11127 | -0,141842 | -1,10175 | 0,30858 | 0,4054211 |
| SHP-2_pY5 | -0,11372 | -0,005081 | -1,81284 | 0,114661 | 0,1916666 |
| CD4 | -0,11859 | 0,126065 | -1,40431 | 0,204861 | 0,2976512 |
| Ets-1 | -0,11913 | 0,029213 | -2,19915 | 0,065532 | 0,118149 |
| AMPK-a2_p | -0,11918 | -0,25196 | -1,38655 | 0,209974 | 0,3032958 |
| D-a-Tubulin | -0,12001 | -0,021247 | -1,47906 | 0,18455 | 0,2779508 |
| PTEN | -0,1206 | 0,391667 | -2,1152 | 0,07401 | 0,1315135 |
| Cdc2_pY15 | -0,12447 | 0,146667 | -2,01365 | 0,08575 | 0,1502142 |
| 53BP1 | -0,12579 | -0,537412 | -0,87951 | 0,409567 | 0,503299 |
| MMP2 | -0,12896 | -0,006025 | -2,28913 | 0,057534 | 0,1052666 |
| TAZ | -0,13101 | -0,016504 | -1,67688 | 0,139403 | 0,2207208 |
| PAK4 | -0,13335 | -0,206457 | -2,56304 | 0,038818 | 0,0767051 |
| Rad51 | -0,14224 | 0,123587 | -2,51801 | 0,041397 | 0,0798835 |
| Granzyme- | -0,14226 | 0,287889 | -1,94506 | 0,094712 | 0,1631877 |
| C-Raf | -0,1492 | 0,027571 | -2,73584 | 0,03038 | 0,0635929 |
| VEGFR-2 | -0,15015 | -0,258649 | -2,51138 | 0,041792 | 0,0800201 |
| PD-L1 | -0,15136 | 0,053002 | -2,54202 | 0,04 | 0,0777962 |
| VASP | -0,15443 | -0,79474 | -1,73271 | 0,128679 | 0,2102227 |
| Glutaminas | -0,16944 | 0,03487 | -2,42971 | 0,046984 | 0,0885875 |
| 14-3-3_bet | -0,17243 | 0,016716 | -2,63356 | 0,035111 | 0,0705078 |
| XBP1 | -0,17389 | 0,114646 | -2,80976 | 0,027382 | 0,0583048 |
| IRF-1 | -0,17441 | 0,032658 | -2,67768 | 0,032982 | 0,0678885 |
| MYH11 | -0,17671 | -0,018565 | -3,17057 | 0,016645 | 0,0387865 |
| INPP4b | -0,18093 | 0,022965 | -3,00863 | 0,02077 | 0,046217 |
| FoxO3a | -0,18691 | 0,081383 | -3,20605 | 0,015864 | 0,0373192 |
| Axl | -0,18778 | 0,175097 | -3,47248 | 0,011124 | 0,0283261 |
| P-Cadherin | -0,19236 | -0,038015 | -3,31283 | 0,013744 | 0,0332818 |
| Wee1_pS6 | -0,19312 | 0,054334 | -3,72218 | 0,008052 | 0,0228591 |
| eEF2K | -0,19548 | -0,047663 | -1,28135 | 0,242646 | 0,3389432 |
| PEA15 | -0,19758 | -0,070949 | -1,24298 | 0,255622 | 0,354712 |
| EGFR | -0,19836 | -0,358619 | -1,20859 | 0,267753 | 0,363379 |
| Atg3 | -0,19938 | 0,20625 | -2,87195 | 0,025102 | 0,0543879 |
| Claudin-7 | -0,20193 | -0,300118 | -3,44577 | 0,011522 | 0,0290396 |
| PEA15_pS1 | -0,20226 | -0,212926 | -1,44046 | 0,194799 | 0,2864007 |
| Smad1 | -0,20614 | -0,031447 | -3,39845 | 0,012265 | 0,030293 |
| Beclin | -0,20638 | 0,082546 | -3,28545 | 0,014257 | 0,0341886 |
| AR | -0,20926 | 0,202032 | -4,06792 | 0,005227 | 0,0161384 |

| | | | | | |
|---|---|---|---|---|---|
| PDHK1 | -0,24969 | -0,141532 | -4,83918 | 5,29E-04 | 0,0052289 |
| PR | -0,25538 | 0,031221 | -4,62549 | 7,46E-04 | 0,0063553 |
| Sox2 | -0,27523 | 0,266721 | -1,20916 | 0,252109 | 0,3797007 |
| TTF1 | -0,27763 | 0,059937 | -4,58585 | 7,96E-04 | 0,0065523 |
| Collagen_V | -0,28817 | -0,072339 | -3,15669 | 0,009201 | 0,0304905 |
| Gys_pS641 | -0,29197 | -0,221008 | -4,19661 | 0,001515 | 0,0093546 |
| TFRC | -0,29611 | 0,24353 | -1,45362 | 0,174154 | 0,2848743 |
| MSI2 | -0,32677 | -0,02712 | -5,08369 | 3,60E-04 | 0,004157 |
| B-Raf_pS44 | -0,33541 | -0,001932 | -2,50787 | 0,029219 | 0,0680857 |
| SDHA | -0,34936 | -0,159111 | -4,4032 | 0,001074 | 0,0079908 |
| Stat3_pY70 | -0,35056 | -0,137707 | -3,70164 | 0,003529 | 0,0164203 |
| Pdcd4 | -0,35882 | -0,201283 | -2,09119 | 0,060675 | 0,1289559 |
| HSP27_pS8 | -0,36439 | 0,019929 | -2,68632 | 0,021274 | 0,0547365 |
| AR | -0,36478 | 0,029133 | -3,98232 | 0,002177 | 0,0122401 |
| Fibronectir | -0,36594 | -0,081243 | -0,67256 | 0,515194 | 0,6254047 |
| Glutaminas | -0,38097 | -0,175164 | -1,99715 | 0,071316 | 0,1432118 |
| ATM | -0,3847 | 0,220743 | -2,75482 | 0,018829 | 0,0511061 |
| SOD2 | -0,39075 | -0,00217 | -5,75394 | 1,31E-04 | 0,0021542 |
| p70S6K_pT | -0,39572 | -0,191915 | -2,10826 | 0,058912 | 0,1276423 |
| B-Raf | -0,39896 | -0,012151 | -3,33268 | 0,006736 | 0,0241119 |
| ULK1_pS75 | -0,42069 | -0,005227 | -4,72419 | 6,36E-04 | 0,0060253 |
| LDHA | -0,44425 | -0,041709 | -2,82167 | 0,016712 | 0,0463804 |
| Cyclin_B1 | -0,44665 | 0,056019 | -4,26203 | 0,001358 | 0,0089284 |
| Gys | -0,46584 | -0,262499 | -6,48289 | 4,67E-05 | 9,61E-04 |
| Axl | -0,48951 | -0,112747 | -5,70622 | 1,40E-04 | 0,0021659 |
| Cox-IV | -0,51711 | -0,001632 | -4,23952 | 0,00141 | 0,0089284 |
| Mcl-1 | -0,5315 | -0,056949 | -2,00174 | 0,070759 | 0,1432118 |
| Myosin-IIa_ | -0,56759 | -0,042934 | -3,00941 | 0,011957 | 0,0364621 |
| Atg3 | -0,5694 | -0,139995 | -7,22159 | 1,77E-05 | 4,90E-04 |
| PLK1 | -0,58795 | -0,05959 | -3,76235 | 0,003177 | 0,0153978 |
| PI3K-p85 | -0,63576 | 0,037165 | -9,3692 | 1,48E-06 | 1,22E-04 |
| TRIM25 | -0,65172 | -0,111768 | -3,25323 | 0,007752 | 0,0269689 |
| Granzyme- | -0,67573 | -0,179623 | -7,87182 | 7,92E-06 | 3,26E-04 |
| S6_pS240_ | -0,68602 | -0,198127 | -1,49949 | 0,162064 | 0,2741772 |
| PTEN | -0,731 | -0,134067 | -4,24212 | 0,001404 | 0,0089284 |
| PKC-alpha | -0,73215 | -0,187465 | -3,91269 | 0,002452 | 0,0131676 |
| AMPK_pT1 | -0,78377 | -0,133669 | -3,97227 | 0,002215 | 0,0122401 |
| S6_pS235_ | -0,80664 | -0,269325 | -1,32602 | 0,211876 | 0,3312243 |
| Stat3 | -0,89634 | -0,433704 | -3,59133 | 0,004276 | 0,0185294 |
| PAR | -0,90248 | 1,00175 | -1,54287 | 0,151303 | 0,2631822 |
| Heregulin | -0,93558 | -0,049425 | -5,06575 | 3,70E-04 | 0,004157 |
| STAT5-alph | -0,95745 | -0,111603 | -6,36315 | 5,50E-05 | 0,0010269 |
| PDGFR-b | -2,77677 | -1,172098 | -30,5446 | 6,21E-12 | 1,53E-09 |

| | | | | | |
|---|---|---|---|---|---|
| PREX1 | -0,2105 | 0,01283 | -2,29758 | 0,056836 | 0,1047657 |
| Rab11 | -0,22434 | -0,153641 | -2,10942 | 0,074633 | 0,1316734 |
| MIF | -0,23139 | -0,023785 | -4,21668 | 0,004365 | 0,0141847 |
| eIF4E_pS2( | -0,23782 | 0,008209 | -4,27932 | 0,004049 | 0,0133362 |
| ULK1_pS75 | -0,24075 | 0,194529 | -3,39095 | 0,012387 | 0,030293 |
| RPA32_pS4 | -0,24076 | -0,363868 | -2,55644 | 0,039186 | 0,0768163 |
| p70-S6K1 | -0,24881 | -0,016118 | -3,88946 | 0,006518 | 0,0196337 |
| SOD2 | -0,25126 | 0,166774 | -4,31333 | 0,003889 | 0,0131586 |
| RBM15 | -0,25367 | 0,095418 | -4,00354 | 0,005657 | 0,0172507 |
| ER-alpha | -0,2694 | 0,072059 | -4,38667 | 0,003566 | 0,0127663 |
| B-Raf_pS44 | -0,27987 | 0,104569 | -3,51451 | 0,010528 | 0,0273731 |
| A-Raf | -0,29433 | 0,052879 | -4,72652 | 0,002412 | 0,0099295 |
| HSP70 | -0,29789 | -0,329619 | -2,71675 | 0,03121 | 0,0647802 |
| Cyclin_B1 | -0,30724 | 0,237113 | -4,47017 | 0,003235 | 0,0122912 |
| ATM | -0,31496 | 0,346519 | -2,63756 | 0,034912 | 0,0705078 |
| TTF1 | -0,31568 | 0,097674 | -5,23531 | 0,001385 | 0,0065509 |
| b-Actin | -0,33762 | -0,255878 | -4,66094 | 0,002598 | 0,0104183 |
| Cox-IV | -0,37275 | 0,197812 | -4,62509 | 0,002706 | 0,0106087 |
| eEF2 | -0,37502 | -0,023708 | -3,48264 | 0,010977 | 0,028242 |
| B-Raf | -0,39284 | 0,078568 | -7,40208 | 1,85E-04 | 0,0013065 |
| PLK1 | -0,40314 | 0,179577 | -5,07954 | 0,001635 | 0,0073431 |
| PI3K-p85 | -0,41028 | 0,311163 | -5,98445 | 6,51E-04 | 0,0034941 |
| c-Myc | -0,41846 | -0,261359 | -4,65506 | 0,002615 | 0,0104183 |
| ARID1A | -0,42873 | -0,318287 | -4,87571 | 0,002042 | 0,0085497 |
| HSP27_pS8 | -0,45719 | 0,043754 | -4,29815 | 0,00396 | 0,0132169 |
| DUSP4 | -0,46174 | 0,002738 | -8,67448 | 7,00E-05 | 6,40E-04 |
| Rad50 | -0,49539 | -0,364531 | -4,59453 | 0,002802 | 0,0108141 |
| Merlin | -0,4973 | -0,364996 | -4,13384 | 0,004824 | 0,0152748 |
| Myosin-IIa_ | -0,51024 | 0,115317 | -3,59979 | 0,009423 | 0,0255765 |
| TRIM25 | -0,51024 | 0,115317 | -3,59979 | 0,009423 | 0,0255765 |
| NF-kB-p65_ | -0,51796 | -0,210527 | -8,6935 | 6,90E-05 | 6,40E-04 |
| FAK_pY397 | -0,54151 | -0,02003 | -7,45072 | 1,78E-04 | 0,0013065 |
| Connexin-4 | -0,54986 | -0,005134 | -6,74757 | 3,22E-04 | 0,0019426 |
| IRS1 | -0,56784 | -0,091901 | -9,31209 | 4,49E-05 | 4,52E-04 |
| Sox2 | -0,57945 | 0,144242 | -7,32754 | 1,97E-04 | 0,0013138 |
| Gab2 | -0,60218 | -0,972387 | -9,2849 | 4,58E-05 | 4,52E-04 |
| PKC-alpha | -0,71892 | -0,019783 | -12,4502 | 7,07E-06 | 1,03E-04 |
| p16_INK4a | -0,93593 | -1,065797 | -14,4963 | 2,63E-06 | 4,65E-05 |
| Heregulin | -1,02453 | 0,101335 | -18,0143 | 6,35E-07 | 1,57E-05 |
| PDGFR-b | -1,08412 | 0,449338 | -19,4359 | 3,85E-07 | 1,06E-05 |
| E-Cadherin | -1,36551 | -1,085202 | -4,44167 | 0,003344 | 0,0124122 |
| Cox2 | -2,24692 | -0,342198 | -34,5506 | 8,46E-09 | 9,01E-07 |
| PAR | -2,5017 | 0,239122 | -25,1183 | 7,06E-08 | 2,90E-06 |

**Subtype A vs. MGT7**

| Protein | logFC | AveExpr | t | P.Value | adj.P.Val |
|---|---|---|---|---|---|
| E-Cadherin | 4,077979 | 1,851072 | 13,59682 | 3,49E-08 | 1,73E-06 |
| P-Met_pY1 | 3,905679 | -0,561518 | 8,062376 | 6,39E-06 | 8,49E-05 |
| RSK | 2,093323 | -0,522712 | 17,44287 | 2,58E-09 | 2,12E-07 |
| CDK1 | 2,055225 | -0,044656 | 6,337862 | 5,75E-05 | 3,23E-04 |
| ARID1A | 1,990081 | 0,479544 | 5,962624 | 9,74E-05 | 4,63E-04 |
| Claudin-7 | 1,973089 | 1,311605 | 7,625586 | 1,08E-05 | 1,03E-04 |
| Rb_pS807_ | 1,75058 | -0,113102 | 5,758942 | 1,31E-04 | 5,87E-04 |
| Cox2 | 1,715259 | 0,635351 | 2,416961 | 0,034359 | 0,0606194 |
| Bim | 1,640005 | 0,183023 | 4,214912 | 0,001475 | 0,0044977 |
| ACC_pS79 | 1,626265 | -0,122915 | 4,328442 | 0,001221 | 0,0039119 |
| Connexin-4 | 1,604578 | 0,22003 | 6,424789 | 5,10E-05 | 3,02E-04 |
| Rad50 | 1,484122 | 0,301541 | 7,84141 | 8,30E-06 | 8,92E-05 |
| FASN | 1,27253 | -0,140903 | 10,44204 | 5,14E-07 | 1,50E-05 |
| ACC1 | 1,219935 | 0,109605 | 6,171473 | 7,24E-05 | 0,000389 |
| PAR | 1,188793 | 0,62152 | 2,029457 | 0,067524 | 0,1030536 |
| 53BP1 | 1,147778 | 0,719624 | 3,536219 | 0,004722 | 0,01178 |
| GCN5L2 | 1,090841 | 0,203806 | 9,041021 | 2,13E-06 | 4,04E-05 |
| 4E-BP1_pS | 1,053014 | 0,270633 | 8,000785 | 6,87E-06 | 8,49E-05 |
| Src_pY416 | 0,971179 | -0,178759 | 4,825654 | 5,44E-04 | 0,0019745 |
| IRS1 | 0,9708 | 0,188377 | 2,63659 | 0,023276 | 0,0438869 |
| p90RSK_pT | 0,943846 | -0,085843 | 8,897744 | 2,49E-06 | 4,39E-05 |
| ATM | 0,881887 | -0,009546 | 6,501237 | 4,59E-05 | 2,91E-04 |
| PARP | 0,817498 | 0,237528 | 7,295409 | 1,62E-05 | 1,29E-04 |
| Akt_pS473 | 0,808969 | -0,222421 | 3,568563 | 0,004462 | 0,0114804 |
| Notch1 | 0,793386 | -0,282907 | 3,537211 | 0,004713 | 0,01178 |
| c-Jun_pS73 | 0,789696 | 0,239301 | 6,09568 | 8,06E-05 | 4,06E-04 |
| beta-Caten | 0,750219 | 0,225981 | 4,315244 | 0,001248 | 0,0039119 |
| c-Myc | 0,71508 | 0,21677 | 7,349606 | 1,52E-05 | 1,25E-04 |
| C-Raf_pS33 | 0,619967 | 0,059052 | 4,13766 | 0,001679 | 0,0050585 |
| PAK1 | 0,590111 | -0,114631 | 1,930564 | 0,079928 | 0,1189295 |
| RPA32_pS4 | 0,583537 | -0,063773 | 6,43274 | 5,04E-05 | 3,02E-04 |
| PI3K-p85 | 0,578939 | -0,183689 | 8,546006 | 3,67E-06 | 5,90E-05 |
| HES1 | 0,575656 | 0,231788 | 1,256218 | 0,23527 | 0,3026657 |
| eIF4E | 0,550199 | -0,025588 | 6,108774 | 7,91E-05 | 4,06E-04 |
| Cdc2_pY15 | 0,5306 | 0,01096 | 4,900449 | 4,83E-04 | 0,0018066 |
| Bid | 0,520625 | 0,129638 | 3,549615 | 0,004612 | 0,0117445 |
| RBM15 | 0,504069 | 0,490142 | 0,941461 | 0,366848 | 0,4441738 |
| Src_pY527 | 0,495044 | 0,037893 | 2,216448 | 0,048859 | 0,0807804 |
| HER3 | 0,484719 | 0,504228 | 2,581701 | 0,025662 | 0,0473706 |
| eEF2 | 0,478251 | 0,01111 | 2,996859 | 0,01225 | 0,0254274 |
| AMPK_alph | 0,444753 | -0,170229 | 4,313517 | 0,001251 | 0,0039119 |
| Smad3 | 0,437212 | 0,147645 | 5,818215 | 1,20E-04 | 5,59E-04 |
| Akt_pT308 | 0,433648 | -0,06235 | 1,968314 | 0,074962 | 0,1129002 |
| PKC-b-II_pS | 0,427428 | -0,185356 | 3,442722 | 0,005562 | 0,0134699 |
| Caspase-3 | 0,427404 | 0,154166 | 5,252264 | 2,79E-04 | 0,0011333 |
| Bcl-xL | 0,420292 | 0,182625 | 3,147138 | 0,009378 | 0,020498 |
| c-Kit | 0,411685 | 0,198267 | 5,25049 | 2,80E-04 | 0,0011333 |
| Wee1_pS6 | 0,40153 | 0,142258 | 4,645249 | 7,26E-04 | 0,0023914 |
| Paxillin | 0,385944 | -0,017222 | 6,852062 | 2,87E-05 | 2,07E-04 |
| Histone-H3 | 0,366995 | 0,057748 | 2,099147 | 0,059892 | 0,0954415 |
| Shc_pY317 | 0,365979 | -0,208821 | 3,786027 | 0,003059 | 0,0084894 |
| HSP70 | 0,360941 | 0,055002 | 4,667178 | 7,01E-04 | 0,0023714 |
| 14-3-3_zet | 0,354151 | -0,00447 | 3,090472 | 0,010371 | 0,0222748 |
| FAK | 0,34764 | -0,018272 | 3,843346 | 0,002771 | 0,0077789 |
| PRAS40_pT | 0,342914 | -0,107622 | 1,167702 | 0,267813 | 0,3392298 |
| VASP | 0,3403 | -0,031485 | 1,781609 | 0,102633 | 0,1491199 |
| Lck | 0,333933 | 0,086966 | 2,531525 | 0,028053 | 0,0504822 |
| XRCC1 | 0,333274 | -0,004986 | 3,395825 | 0,006041 | 0,0142101 |
| Sox2 | 0,332648 | 0,156197 | 1,460706 | 0,172288 | 0,2325423 |
| DM-K9-His | 0,311906 | -0,030843 | 4,64969 | 7,21E-04 | 0,0023914 |
| Caspase-7_ | 0,31119 | -0,113361 | 2,543016 | 0,027486 | 0,0499203 |
| Chk2_pT68 | 0,310569 | -0,020476 | 3,082131 | 0,010526 | 0,0224127 |
| Myt1 | 0,30976 | -0,038003 | 3,199444 | 0,008547 | 0,019191 |
| Merlin | 0,309137 | 0,081024 | 1,768057 | 0,104967 | 0,1516185 |
| DJ-1 | 0,298587 | 0,133255 | 3,448291 | 0,005508 | 0,0134699 |
| WIPI1 | 0,293815 | 0,245293 | 1,298115 | 0,221019 | 0,288845 |

**subtype B vs. MGT 7**

| Protein | logFC | AveExpr | t | P.Value | adj.P.Val |
|---|---|---|---|---|---|
| P-Met_pY1 | 4,697158 | -0,938778 | 74,92326 | 1,46E-10 | 3,62E-08 |
| RSK | 2,683463 | -0,625353 | 54,86156 | 1,05E-09 | 1,29E-07 |
| Notch1 | 2,347481 | 0,476448 | 45,8311 | 3,26E-09 | 2,03E-07 |
| Rb_pS807_ | 2,18387 | -0,235072 | 39,93065 | 7,76E-09 | 3,83E-07 |
| ACC_pS79 | 2,067445 | -0,213028 | 23,94125 | 1,93E-07 | 4,51E-06 |
| CDK1 | 1,97376 | -0,541948 | 38,45157 | 9,84E-09 | 4,05E-07 |
| Fibronectir | 1,530955 | 1,205599 | 9,730992 | 4,91E-05 | 2,53E-04 |
| Akt_pS473 | 1,368307 | -0,06332 | 13,46006 | 6,88E-06 | 6,30E-05 |
| Bim | 1,28209 | -0,389545 | 24,19337 | 1,81E-07 | 4,51E-06 |
| FASN | 1,22342 | -0,448012 | 14,95374 | 3,60E-06 | 3,58E-05 |
| HES1 | 1,220882 | 0,493327 | 21,1277 | 4,21E-07 | 6,94E-06 |
| ACC1 | 1,207576 | -0,163979 | 21,16275 | 4,17E-07 | 6,94E-06 |
| Src_pY416 | 1,029589 | -0,355606 | 18,66111 | 9,13E-07 | 1,11E-05 |
| PKA-a | 0,985279 | 0,443107 | 19,2266 | 7,59E-07 | 1,04E-05 |
| ATM | 0,951623 | -0,160116 | 8,725685 | 9,39E-05 | 4,37E-04 |
| p90RSK_pT | 0,890456 | -0,323807 | 10,28256 | 3,53E-05 | 2,03E-04 |
| Connexin-4 | 0,867729 | -0,57217 | 10,61205 | 2,92E-05 | 1,76E-04 |
| Pdcd4 | 0,811065 | 0,126594 | 11,36853 | 1,93E-05 | 1,22E-04 |
| PI3K-p85 | 0,804418 | -0,174715 | 12,15262 | 1,29E-05 | 9,34E-05 |
| Akt_pT308 | 0,800977 | 0,063433 | 9,42914 | 5,93E-05 | 2,93E-04 |
| STAT5-alph | 0,729843 | 0,553266 | 12,95944 | 8,68E-06 | 6,70E-05 |
| IGFRb | 0,673653 | 0,876255 | 13,66633 | 6,27E-06 | 5,96E-05 |
| ARID1A | 0,658924 | -0,75335 | 7,53846 | 2,20E-04 | 8,26E-04 |
| LRP6_pS14 | 0,641061 | 0,293243 | 11,98267 | 1,40E-05 | 9,88E-05 |
| Shc_pY317 | 0,607148 | -0,143969 | 9,197737 | 6,87E-05 | 3,33E-04 |
| PAK1 | 0,585795 | -0,245972 | 11,11462 | 2,21E-05 | 1,36E-04 |
| AMPK_alph | 0,582495 | -0,184621 | 10,14781 | 3,82E-05 | 2,15E-04 |
| XRCC1 | 0,525977 | 0,037921 | 6,388376 | 5,63E-04 | 0,0018298 |
| Cdc2_pY15 | 0,520045 | -0,11114 | 7,186166 | 2,90E-04 | 0,0010225 |
| Wee1 | 0,506484 | 0,139087 | 8,132213 | 1,42E-04 | 6,15E-04 |
| GCN5L2 | 0,502351 | -0,38729 | 5,969369 | 8,19E-04 | 0,0023802 |
| Mcl-1 | 0,499886 | 0,347536 | 7,515356 | 2,24E-04 | 8,26E-04 |
| 4E-BP1_pS | 0,495256 | -0,29377 | 8,153183 | 1,40E-04 | 6,15E-04 |
| AMPK_pT1 | 0,486715 | 0,997887 | 4,799688 | 0,002606 | 0,0060716 |
| Bid | 0,484994 | -0,00533 | 3,802381 | 0,00811 | 0,0147343 |
| PRAS40_pT | 0,479974 | -0,100203 | 8,915874 | 8,26E-05 | 3,93E-04 |
| RIP | 0,462285 | 0,106685 | 1,844754 | 0,112134 | 0,1465453 |
| Caspase-7_ | 0,449403 | -0,098329 | 8,343125 | 1,22E-04 | 5,59E-04 |
| PTEN | 0,448663 | 0,163963 | 9,436692 | 5,90E-05 | 2,93E-04 |
| PKC-b-II_pS | 0,433918 | -0,274718 | 6,139734 | 7,02E-04 | 0,0021397 |
| Tuberin | 0,409148 | 0,22117 | 4,123747 | 0,005535 | 0,0110677 |
| Bad_pS112 | 0,405622 | -0,121011 | 5,186137 | 0,001744 | 0,0045816 |
| beta-Caten | 0,402134 | -0,146554 | 6,042147 | 7,66E-04 | 0,0023086 |
| Paxillin | 0,39243 | -0,097537 | 7,902622 | 1,68E-04 | 6,86E-04 |
| mTOR | 0,387562 | -0,066634 | 7,952201 | 1,62E-04 | 6,77E-04 |
| PARP | 0,373534 | -0,207213 | 3,540071 | 0,011208 | 0,0197741 |
| Chk2_pT68 | 0,368514 | -0,05347 | 5,439334 | 0,001355 | 0,0035975 |
| eIF4E | 0,350416 | -0,265501 | 6,760994 | 4,10E-04 | 0,0013834 |
| Rad50 | 0,350054 | -0,702708 | 3,519606 | 0,0115 | 0,0201447 |
| Myt1 | 0,325818 | -0,095952 | 5,030162 | 0,002046 | 0,0050525 |
| HER2_pY12 | 0,319297 | 0,094711 | 4,232412 | 0,004881 | 0,0101314 |
| WIPI1 | 0,294558 | 0,181634 | 3,502838 | 0,011745 | 0,0204291 |
| IRS1 | 0,286879 | -0,433787 | 3,969903 | 0,006632 | 0,0130014 |
| STING | 0,27701 | 0,433335 | 2,789512 | 0,029918 | 0,0461539 |
| 14-3-3_zet | 0,271302 | -0,131449 | 5,086183 | 0,001931 | 0,0049116 |
| Stathmin | 0,267642 | -0,142018 | 4,610754 | 0,003195 | 0,0072399 |
| COG3 | 0,26554 | 0,083359 | 3,802082 | 0,008113 | 0,0147343 |
| Bak | 0,250272 | -0,090717 | 3,83869 | 0,007761 | 0,0143458 |
| YB1_pS102 | 0,248613 | -0,039595 | 3,901744 | 0,007194 | 0,0135278 |
| Raptor | 0,247552 | -0,032933 | 4,238046 | 0,00485 | 0,0101314 |
| SDHA | 0,245534 | 0,121946 | 3,433324 | 0,012824 | 0,0216948 |
| Caspase-3 | 0,238695 | -0,052312 | 3,961999 | 0,006695 | 0,0130208 |
| FAK | 0,23667 | -0,160703 | 2,734696 | 0,032237 | 0,0485526 |
| DM-K9-His | 0,231501 | -0,147139 | 3,087097 | 0,020079 | 0,0324157 |
| C-Raf | 0,225082 | -0,122143 | 4,418151 | 0,003954 | 0,0085667 |
| IGFBP2 | 0,218739 | 0,004805 | 1,89843 | 0,103935 | 0,1372836 |

| | | | | | |
|---|---|---|---|---|---|
| PAICS | 0,290497 | 0,348264 | 1,187647 | 0,260186 | 0,3312678 |
| Bak | 0,263146 | -0,025579 | 3,326898 | 0,006821 | 0,0157461 |
| DUSP4 | 0,259141 | 0,04823 | 2,047837 | 0,065427 | 0,1010029 |
| Bad_pS112 | 0,25529 | -0,15551 | 3,110719 | 0,010004 | 0,021676 |
| HER2_pY1 | 0,252006 | 0,109319 | 1,324687 | 0,212352 | 0,2813903 |
| p38_pT18 | 0,244549 | 0,152977 | 3,217093 | 0,008283 | 0,0187706 |
| p27_pT19 | 0,242723 | 0,08116 | 2,729391 | 0,019731 | 0,0386796 |
| Rictor | 0,231252 | 0,003434 | 2,248235 | 0,04622 | 0,077663 |
| Wee1 | 0,230033 | 0,023405 | 1,07217 | 0,306787 | 0,3827096 |
| mTOR | 0,227446 | -0,113079 | 2,147609 | 0,055074 | 0,0883326 |
| EGFR_pY11 | 0,220239 | -0,020761 | 4,315428 | 0,001247 | 0,0039119 |
| C-Raf | 0,217194 | -0,079488 | 2,02889 | 0,06759 | 0,1030536 |
| Stathmin | 0,215478 | -0,126303 | 2,337273 | 0,039539 | 0,0682952 |
| Elk1_pS38 | 0,212288 | -0,025692 | 3,610465 | 0,004147 | 0,0107835 |
| ATR_pS42 | 0,211757 | 0,022845 | 3,414579 | 0,005845 | 0,0138809 |
| TUFM | 0,210513 | 0,137065 | 2,61265 | 0,024288 | 0,0454489 |
| Raptor | 0,208643 | -0,010756 | 2,375235 | 0,036984 | 0,0647868 |
| ER-alpha_ | 0,208631 | 0,195351 | 2,953036 | 0,013245 | 0,0270372 |
| Bax | 0,196227 | 0,010879 | 0,440265 | 0,66834 | 0,7219861 |
| MEK1_pS2 | 0,195554 | 0,025863 | 3,328584 | 0,006801 | 0,0157461 |
| Chk1_pS29 | 0,18635 | -0,063817 | 2,978386 | 0,01266 | 0,0260589 |
| A-Raf | 0,185335 | -0,02201 | 1,464524 | 0,171263 | 0,2324288 |
| b-Actin | 0,169691 | 0,160934 | 1,054955 | 0,314248 | 0,3900466 |
| Tyro3 | 0,168276 | -0,014282 | 3,054048 | 0,011065 | 0,0233592 |
| TAZ | 0,155878 | 0,184933 | 1,815747 | 0,096962 | 0,1425579 |
| GCLM | 0,150735 | -0,011929 | 3,187728 | 0,008726 | 0,0194173 |
| EGFR | 0,147047 | -0,047453 | 1,591412 | 0,140054 | 0,1943441 |
| NAPSIN-A | 0,146958 | -0,080924 | 1,935652 | 0,079242 | 0,1186224 |
| UBAC1 | 0,143835 | -0,04358 | 1,511115 | 0,159169 | 0,2172084 |
| NF-kB-p65_ | 0,138268 | -0,007839 | 0,931245 | 0,371876 | 0,4480651 |
| eIF4G | 0,133138 | 0,068523 | 1,787907 | 0,101565 | 0,1484409 |
| PDK1 | 0,131864 | -0,009512 | 2,086559 | 0,061208 | 0,0959359 |
| YB1_pS102 | 0,130538 | -0,081959 | 1,322557 | 0,213036 | 0,2813903 |
| MSH6 | 0,117776 | 0,164421 | 0,748335 | 0,470083 | 0,5425722 |
| AMPK-a2_ | 0,116284 | -0,086014 | 1,096294 | 0,296559 | 0,3737254 |
| Notch3 | 0,116038 | 0,05209 | 2,258598 | 0,04539 | 0,0767905 |
| eIF4E_pS2 | 0,113283 | 0,00323 | 1,683749 | 0,12059 | 0,1708614 |
| RIP | 0,100239 | -0,088672 | 0,438797 | 0,669372 | 0,7219861 |
| p16_INK4a | 0,100033 | 0,299067 | 0,373217 | 0,716137 | 0,7657393 |
| PI3K-p110- | 0,09905 | -0,062117 | 2,304663 | 0,04187 | 0,0718194 |
| Pdcd4 | 0,093523 | -0,283527 | 0,544331 | 0,597163 | 0,658033 |
| p27-Kip-1 | 0,092997 | -0,05325 | 1,861886 | 0,089752 | 0,1327472 |
| Tuberin | 0,092488 | 0,051353 | 0,816781 | 0,431528 | 0,5075587 |
| XPF | 0,08906 | -0,131189 | 1,587525 | 0,14093 | 0,1944682 |
| Aurora-B | 0,087973 | -0,024936 | 0,885671 | 0,394897 | 0,4689401 |
| Rab11 | 0,085186 | -0,054335 | 0,738243 | 0,475947 | 0,5467861 |
| FOXO3a_p | 0,083044 | -0,073689 | 1,614393 | 0,134967 | 0,1883432 |
| YAP | 0,081174 | -0,057171 | 1,089494 | 0,299416 | 0,3754096 |
| IGFBP2 | 0,079857 | -0,061101 | 0,593631 | 0,564863 | 0,6341875 |
| PEA15_pS1 | 0,070285 | -0,002652 | 0,382432 | 0,709486 | 0,7619261 |
| Ets-1 | 0,06179 | 0,040532 | 1,220786 | 0,247895 | 0,3172537 |
| p53 | 0,06094 | 0,003464 | 0,455481 | 0,657695 | 0,7156414 |
| MDM2_pS | 0,05173 | -0,168872 | 0,477976 | 0,642101 | 0,7017648 |
| 4E-BP1 | 0,050315 | 0,164505 | 0,250601 | 0,80678 | 0,8408209 |
| Cyclin_D1 | 0,045221 | 0,016667 | 0,563676 | 0,584375 | 0,6472668 |
| TIGAR | 0,039477 | 0,060019 | 0,334432 | 0,744394 | 0,7824057 |
| D-a-Tubuli | 0,037758 | -0,056284 | 0,627801 | 0,543051 | 0,6152922 |
| AR | 0,018574 | -0,040568 | 0,197774 | 0,846858 | 0,8727302 |
| JNK2 | 0,011897 | -0,004291 | 0,121746 | 0,905314 | 0,9240183 |
| cIAP | 0,002476 | -0,046568 | 0,050399 | 0,960715 | 0,96587 |
| b-Catenin_ | 2,89E-05 | 0,070485 | 3,62E-04 | 0,999718 | 0,9997176 |
| Akt | -0,00926 | -0,07518 | -0,0488 | 0,96196 | 0,96587 |
| mTOR_pS2 | -0,01366 | -0,141743 | -0,08319 | 0,935205 | 0,9467033 |
| PKA-a | -0,0203 | -0,164673 | -0,16907 | 0,868835 | 0,8904658 |
| PLC-gamm | -0,02153 | 0,030147 | -0,33716 | 0,742393 | 0,7824057 |
| Slfn11 | -0,02255 | -0,09016 | -0,25396 | 0,804249 | 0,8408209 |
| c-Abl | -0,03094 | -0,081375 | -0,58467 | 0,570663 | 0,6377998 |
| U-Histone- | -0,03409 | -0,00702 | -0,3345 | 0,744342 | 0,7824057 |
| MSI2 | -0,03581 | -0,080021 | -0,57022 | 0,580082 | 0,645407 |

| | | | | | |
|---|---|---|---|---|---|
| Tyro3 | 0,216841 | -0,021857 | 4,213207 | 0,00499 | 0,0102712 |
| Lck | 0,214116 | -0,057782 | 2,749367 | 0,031598 | 0,0478822 |
| PI3K-p110- | 0,206945 | -0,018991 | 4,176665 | 0,005205 | 0,0105379 |
| XPF | 0,206908 | -0,079911 | 4,263259 | 0,004712 | 0,0100325 |
| eEF2 | 0,20647 | -0,256304 | 2,235929 | 0,064486 | 0,0899894 |
| MSI2 | 0,206411 | 0,073123 | 3,944605 | 0,006835 | 0,0130871 |
| IGF1R_pY1 | 0,203077 | 0,268704 | 3,15669 | 0,018323 | 0,0301385 |
| p21 | 0,196038 | -0,030453 | 3,725141 | 0,008911 | 0,0159489 |
| UBAC1 | 0,191785 | -0,046192 | 3,92717 | 0,006979 | 0,0132594 |
| EGFR_pY11 | 0,189005 | -0,087553 | 3,705302 | 0,00913 | 0,0162243 |
| mTOR_pS2 | 0,186438 | -0,018703 | 2,700916 | 0,033761 | 0,0502354 |
| Histone-H3 | 0,184841 | -0,131616 | 1,711879 | 0,135249 | 0,1695762 |
| AR | 0,174091 | 0,04869 | 3,060825 | 0,020789 | 0,0333433 |
| ATR_pS42 | 0,169321 | -0,048818 | 2,39691 | 0,051438 | 0,0734402 |
| Chk1_pS29 | 0,166521 | -0,116373 | 2,190569 | 0,068747 | 0,0953957 |
| Elk1_pS38 | 0,164145 | -0,100894 | 3,258162 | 0,016054 | 0,0267929 |
| Wee1_pS6 | 0,163351 | -0,088256 | 1,790287 | 0,121102 | 0,1557927 |
| SLC1A5 | 0,161474 | 0,51657 | 2,657443 | 0,035836 | 0,0526878 |
| MAPK_pT2 | 0,155805 | 0,083148 | 1,900907 | 0,103572 | 0,1372836 |
| c-Kit | 0,15214 | -0,047281 | 1,584196 | 0,161743 | 0,1997521 |
| GATA6 | 0,150994 | 0,039006 | 3,044622 | 0,02124 | 0,0338469 |
| FOXO3a_p | 0,140839 | -0,05713 | 2,660797 | 0,035671 | 0,0526878 |
| S6_pS240_ | 0,12355 | 0,445644 | 1,144789 | 0,293792 | 0,3406887 |
| YAP | 0,122936 | -0,049824 | 2,363145 | 0,053928 | 0,0765528 |
| Bax | 0,122129 | -0,076393 | 2,416638 | 0,050038 | 0,0718576 |
| p27_pT19 | 0,121086 | -0,04478 | 2,287947 | 0,059933 | 0,0841099 |
| c-Abl | 0,118939 | 0,015301 | 2,172045 | 0,070568 | 0,0973764 |
| Smad3 | 0,11821 | -0,139148 | 0,958144 | 0,373195 | 0,4267554 |
| A-Raf | 0,108797 | -0,10837 | 1,961889 | 0,095005 | 0,1275342 |
| c-Jun_pS7 | 0,106923 | -0,34266 | 1,761632 | 0,126097 | 0,1605466 |
| GCLM | 0,094828 | -0,078361 | 1,893753 | 0,104626 | 0,1374601 |
| PLC-gamm | 0,093728 | 0,104 | 1,902321 | 0,103365 | 0,1372836 |
| C-Raf_pS3 | 0,08842 | -0,395141 | 1,749967 | 0,128188 | 0,1623709 |
| RPA32_pS | 0,08104 | -0,492589 | 0,878326 | 0,411911 | 0,4667061 |
| MEK1_pS2 | 0,071031 | -0,091517 | 1,206356 | 0,27089 | 0,3171079 |
| DJ-1 | 0,060863 | -0,074525 | 0,935965 | 0,383663 | 0,4367036 |
| Bcl-xL | 0,052279 | -0,129883 | 1,046435 | 0,333722 | 0,3833918 |
| U-Histone- | 0,037331 | 0,043273 | 0,397148 | 0,704325 | 0,7531093 |
| Akt | 0,032079 | -0,04836 | 0,486619 | 0,642945 | 0,6965234 |
| IR-b | 0,031783 | 0,316294 | 0,225578 | 0,828649 | 0,8457697 |
| Aurora-B | 0,030001 | -0,078913 | 0,331881 | 0,750711 | 0,7919369 |
| Sox2 | 0,028433 | -0,098909 | 0,367761 | 0,725056 | 0,7686211 |
| Src_pY527 | 0,022202 | -0,353822 | 0,418916 | 0,689144 | 0,7400808 |
| D-a-Tubuli | 0,017926 | -0,076422 | 0,328062 | 0,753462 | 0,7919369 |
| cIAP | 0,014827 | -0,039698 | 0,238545 | 0,819006 | 0,8393962 |
| MDM2_pS | 0,009568 | -0,205456 | 0,088181 | 0,932461 | 0,9400729 |
| PKC-delta_ | 0,009284 | -0,030647 | 0,158939 | 0,878675 | 0,8894786 |
| NAPSIN-A | 0,003427 | -0,199106 | 0,033002 | 0,974691 | 0,9746906 |
| eIF4E_pS2 | 0,002538 | -0,087934 | 0,0336 | 0,974232 | 0,9746906 |
| Ets-1 | -0,00842 | -0,015073 | -0,16254 | 0,875957 | 0,8894786 |
| Rictor | -0,01333 | -0,193772 | -0,24154 | 0,816783 | 0,8393962 |
| JNK2 | -0,01507 | -0,023068 | -0,25977 | 0,803298 | 0,8336747 |
| DUSP4 | -0,01817 | -0,174693 | -0,30705 | 0,768672 | 0,8044994 |
| p38_pT18 | -0,01984 | -0,059012 | -0,276 | 0,791352 | 0,824742 |
| PAICS | -0,02576 | 0,095129 | -0,24407 | 0,814904 | 0,8393962 |
| Slfn11 | -0,03327 | -0,091667 | -0,36891 | 0,72424 | 0,7686211 |
| 14-3-3_bet | -0,03672 | -0,03757 | -0,60925 | 0,563605 | 0,6159749 |
| p53 | -0,04621 | -0,07412 | -0,70626 | 0,505217 | 0,5595896 |
| PDK1 | -0,05047 | -0,14768 | -0,74929 | 0,480636 | 0,5371812 |
| TUFM | -0,05267 | -0,066776 | -0,69002 | 0,514708 | 0,5675577 |
| ER-alpha_ | -0,05461 | -0,008113 | -0,71014 | 0,502968 | 0,5595896 |
| c-Myc | -0,06045 | -0,404563 | -0,68072 | 0,520194 | 0,5710574 |
| MERIT40_ | -0,07311 | 0,13586 | -0,5356 | 0,610552 | 0,6643448 |
| FoxM1 | -0,07797 | 0,059247 | -0,85536 | 0,42359 | 0,4777478 |
| Claudin-7 | -0,07815 | -0,34963 | -1,25117 | 0,255196 | 0,3001594 |
| Notch3 | -0,08199 | -0,092046 | -1,52618 | 0,175339 | 0,2133429 |
| CD20 | -0,08368 | 0,172599 | -1,58464 | 0,161643 | 0,1997521 |
| RBM15 | -0,08377 | 0,02746 | -1,53398 | 0,173451 | 0,2120923 |
| p27-Kip-1 | -0,08378 | -0,179609 | -1,35737 | 0,221134 | 0,2625964 |

| | | | | | |
|---|---|---|---|---|---|
| 14-3-3_bet | -0,0366 | -0,045484 | -0,61735 | 0,549669 | 0,6199463 |
| Smad1 | -0,0399 | 0,026621 | -0,35583 | 0,728754 | 0,775872 |
| Jagged1 | -0,04015 | 0,160316 | -0,19628 | 0,847997 | 0,8727302 |
| PKC-delta_ | -0,04355 | -0,07185 | -0,89381 | 0,390713 | 0,4684759 |
| Heregulin | -0,04479 | -0,211387 | -0,24282 | 0,812649 | 0,8433794 |
| MMP2 | -0,0474 | 0,052953 | -0,98348 | 0,346679 | 0,426019 |
| IRF-1 | -0,05491 | 0,04805 | -0,81101 | 0,434694 | 0,5088597 |
| STING | -0,06749 | 0,211912 | -0,10141 | 0,921061 | 0,9362228 |
| LRP6_pS14 | -0,0724 | -0,15063 | -0,94618 | 0,36454 | 0,4441738 |
| MAPK_pT2 | -0,1042 | -0,095589 | -0,54094 | 0,599423 | 0,658033 |
| Rad51 | -0,12134 | 0,229754 | -0,88946 | 0,392944 | 0,4688757 |
| PDK1_pS24 | -0,12291 | 0,169696 | -1,70236 | 0,116972 | 0,1679774 |
| p70-S6K1 | -0,12351 | 0,129352 | -1,00516 | 0,336594 | 0,4156942 |
| p21 | -0,12913 | -0,253731 | -1,41581 | 0,18474 | 0,246653 |
| Creb | -0,13563 | 0,050198 | -2,6463 | 0,022878 | 0,0434674 |
| IGF1R_pY1 | -0,14277 | 0,030049 | -2,1569 | 0,054193 | 0,0874881 |
| IR-b | -0,15029 | 0,174263 | -0,7185 | 0,487553 | 0,5575264 |
| eEF2K | -0,15551 | 0,19831 | -0,71006 | 0,492563 | 0,5606592 |
| PTEN | -0,16174 | -0,237569 | -0,94207 | 0,366548 | 0,4441738 |
| TTF1 | -0,16512 | 0,039481 | -2,9145 | 0,014186 | 0,0284879 |
| OCT4 | -0,17705 | 0,042496 | -3,68896 | 0,003618 | 0,00961 |
| FoxO3a | -0,1888 | -0,009503 | -2,29361 | 0,04269 | 0,0727205 |
| PEA15 | -0,1898 | 0,194478 | -1,43683 | 0,178817 | 0,2400428 |
| GATA6 | -0,18983 | -0,206902 | -2,21413 | 0,049057 | 0,0807804 |
| Jak2 | -0,19383 | 0,01048 | -2,52848 | 0,028205 | 0,0504822 |
| ER-alpha | -0,19453 | 0,052902 | -3,15482 | 0,009251 | 0,0204008 |
| XBP1 | -0,19696 | 0,006864 | -4,28252 | 0,001317 | 0,0040676 |
| SDHA | -0,19726 | -0,186767 | -2,58089 | 0,025699 | 0,0473706 |
| PREX1 | -0,19958 | 0,220191 | -2,42653 | 0,033783 | 0,0600322 |
| Mcl-1 | -0,20119 | -0,117005 | -0,75625 | 0,465519 | 0,5398267 |
| JNK_pT183 | -0,20163 | 0,079326 | -2,08504 | 0,061368 | 0,0959359 |
| FoxM1 | -0,21101 | -0,066612 | -2,36236 | 0,037832 | 0,065806 |
| B-Raf_pS44 | -0,21397 | -0,024011 | -1,6566 | 0,126047 | 0,176895 |
| COG3 | -0,21831 | -0,254581 | -1,31567 | 0,215261 | 0,282817 |
| INPP4b | -0,2184 | -0,019661 | -3,74148 | 0,003304 | 0,0090666 |
| CD134 | -0,23267 | 0,027245 | -3,45868 | 0,005409 | 0,0133595 |
| MERIT40_p | -0,24118 | -0,017598 | -2,18225 | 0,051857 | 0,0842679 |
| p38_MAPK | -0,24121 | 0,080272 | -1,68139 | 0,121056 | 0,1708614 |
| N-Cadherin | -0,25227 | -0,047889 | -1,68977 | 0,119409 | 0,1704862 |
| HER3_pY12 | -0,2527 | 0,035728 | -3,6702 | 0,003738 | 0,0098224 |
| Glutamate- | -0,26454 | 0,049489 | -2,943 | 0,013484 | 0,0272993 |
| CD20 | -0,28029 | -0,006524 | -3,7106 | 0,003485 | 0,0094595 |
| SLC1A5 | -0,28694 | 0,184915 | -1,26037 | 0,233826 | 0,3023821 |
| PD-L1 | -0,29215 | -0,023057 | -5,77988 | 1,27E-04 | 5,80E-04 |
| ZAP-70 | -0,29337 | 0,082267 | -3,03637 | 0,011418 | 0,0239012 |
| Beclin | -0,30749 | 0,030434 | -4,99438 | 4,16E-04 | 0,0015821 |
| P-Cadherin | -0,30802 | 0,109523 | -3,43219 | 0,005666 | 0,0135884 |
| Rab25 | -0,31623 | -0,072747 | -4,07211 | 0,001876 | 0,005516 |
| Mnk1 | -0,31864 | 0,168786 | -2,7471 | 0,019118 | 0,037778 |
| cdc25C | -0,32948 | 0,176967 | -2,05459 | 0,064672 | 0,1004658 |
| ATM_pS19 | -0,33532 | 0,002799 | -7,52054 | 1,23E-05 | 1,12E-04 |
| MIF | -0,34484 | -0,038735 | -4,13008 | 0,001701 | 0,0050618 |
| G6PD | -0,35346 | -0,118073 | -2,2265 | 0,048009 | 0,080123 |
| PR | -0,35651 | 0,049608 | -6,40147 | 5,26E-05 | 3,02E-04 |
| FAK_pY397 | -0,35757 | 0,306504 | -0,85553 | 0,410652 | 0,4853166 |
| DM-Histon | -0,36085 | 0,075023 | -4,6957 | 6,69E-04 | 0,0023286 |
| Gab2 | -0,36871 | 0,167786 | -1,26068 | 0,233719 | 0,3023821 |
| Tuberin_pT | -0,37218 | 0,042294 | -4,68225 | 6,84E-04 | 0,0023466 |
| STAT5-alph | -0,40104 | -0,212767 | -2,65449 | 0,022546 | 0,0431703 |
| CD4 | -0,41325 | 0,069014 | -5,17645 | 3,14E-04 | 0,0012104 |
| ERCC5 | -0,419 | -0,018824 | -3,96449 | 0,002252 | 0,0065448 |
| GSK-3a-b_p | -0,42261 | 0,040381 | -1,99485 | 0,071644 | 0,1085653 |
| VEGFR-2 | -0,42756 | 0,070836 | -2,55996 | 0,026672 | 0,0488002 |
| Fibronectin | -0,4348 | -0,068722 | -0,80195 | 0,439698 | 0,5122902 |
| Rictor_pT1 | -0,44919 | -0,011196 | -2,89756 | 0,014621 | 0,0291246 |
| SOD2 | -0,44982 | 0,008572 | -6,68145 | 3,60E-05 | 2,41E-04 |
| BRD4 | -0,45613 | -0,010498 | -6,96157 | 2,49E-05 | 1,86E-04 |
| PAK4 | -0,46359 | 0,132529 | -4,87975 | 4,99E-04 | 0,0018389 |
| Hexokinase | -0,46452 | 0,025019 | -7,4514 | 1,34E-05 | 1,18E-04 |

| | | | | | |
|---|---|---|---|---|---|
| PR | -0,09048 | 0,287013 | -1,77088 | 0,124464 | 0,1592882 |
| Atg3 | -0,09665 | 0,16516 | -0,84611 | 0,428357 | 0,4809286 |
| CD134 | -0,09835 | 0,158608 | -1,7218 | 0,133374 | 0,1680785 |
| TIGAR | -0,09866 | -0,031474 | -1,83394 | 0,113861 | 0,1472443 |
| Stat3 | -0,10554 | 0,514912 | -2,07753 | 0,080657 | 0,1100685 |
| eIF4G | -0,1067 | -0,10443 | -1,39794 | 0,209231 | 0,2508744 |
| AMPK-a2_p | -0,11848 | -0,252242 | -1,3651 | 0,21882 | 0,2611044 |
| Rab11 | -0,11934 | -0,195639 | -1,11099 | 0,307043 | 0,3543911 |
| XBP1 | -0,12292 | 0,094257 | -2,48741 | 0,045337 | 0,0654866 |
| TAZ | -0,12631 | -0,018387 | -1,53397 | 0,173452 | 0,2120923 |
| ERCC5 | -0,12795 | 0,247226 | -1,15828 | 0,28864 | 0,3362926 |
| b-Catenin_ | -0,12941 | -0,007183 | -1,28495 | 0,243892 | 0,2882362 |
| IRF-1 | -0,13009 | 0,014927 | -2,03833 | 0,085258 | 0,1157078 |
| Heregulin | -0,13374 | -0,254981 | -2,65088 | 0,036161 | 0,0528507 |
| OCT4 | -0,13794 | 0,10459 | -2,72506 | 0,032665 | 0,0488979 |
| FoxO3a | -0,13835 | 0,061959 | -2,61028 | 0,03824 | 0,0555612 |
| Cyclin_D1 | -0,14334 | -0,106339 | -2,13945 | 0,073894 | 0,1013985 |
| E-Cadherin | -0,14595 | -1,573026 | -0,44404 | 0,671813 | 0,7246191 |
| MMP2 | -0,14934 | 0,002127 | -2,8531 | 0,027448 | 0,0426387 |
| B-Raf_pS44 | -0,15844 | 0,055994 | -2,77263 | 0,030613 | 0,0466748 |
| G6PD | -0,16386 | 0,072803 | -1,40337 | 0,207682 | 0,2502312 |
| Creb | -0,16469 | 0,062356 | -3,28144 | 0,015578 | 0,0261749 |
| Smad1 | -0,17317 | -0,044635 | -2,91578 | 0,025225 | 0,0396852 |
| MSH6 | -0,18649 | -0,043836 | -2,31558 | 0,057649 | 0,0813681 |
| TTF1 | -0,20318 | 0,052673 | -4,3103 | 0,004466 | 0,0095913 |
| INPP4b | -0,20844 | 0,033968 | -3,0191 | 0,021972 | 0,0347891 |
| HSP70 | -0,20846 | -0,365392 | -1,98403 | 0,092073 | 0,1242738 |
| Jak2 | -0,21235 | 0,04166 | -3,77507 | 0,008384 | 0,015115 |
| DM-Histon | -0,22316 | 0,236368 | -3,83638 | 0,007783 | 0,0143458 |
| PD-L1 | -0,22352 | 0,081866 | -4,11713 | 0,005577 | 0,0110677 |
| GSK-3a-b_p | -0,22529 | 0,250974 | -4,24644 | 0,004803 | 0,0101314 |
| p44-42-MA | -0,22796 | 0,137085 | -4,1135 | 0,005601 | 0,0110677 |
| N-Cadherin | -0,23219 | 0,019196 | -4,78009 | 0,002661 | 0,0061419 |
| PEA15_pS1 | -0,23367 | -0,200361 | -1,60621 | 0,156849 | 0,1956648 |
| cdc25C | -0,23414 | 0,306053 | -1,84101 | 0,112729 | 0,1465474 |
| Stat3_pY70 | -0,23629 | 0,395886 | -5,00692 | 0,002095 | 0,0051244 |
| ZAP-70 | -0,23925 | 0,178742 | -3,47374 | 0,012184 | 0,0208984 |
| Rictor_pT1 | -0,24618 | 0,208616 | -5,06932 | 0,001965 | 0,0049116 |
| ER-alpha | -0,24776 | 0,063404 | -4,6993 | 0,002902 | 0,0066368 |
| Rab25 | -0,25567 | 0,032587 | -4,47119 | 0,003726 | 0,0081454 |
| Rad51 | -0,26234 | 0,171627 | -3,13177 | 0,018932 | 0,0307647 |
| HER3_pY12 | -0,26425 | 0,083933 | -4,52409 | 0,003514 | 0,0078198 |
| EGFR | -0,2702 | -0,329884 | -1,42435 | 0,201796 | 0,2443318 |
| ATM_pS19 | -0,2758 | 0,111675 | -5,53879 | 0,001229 | 0,0033368 |
| Beclin | -0,28202 | 0,112803 | -5,74327 | 0,001011 | 0,0028377 |
| HER3 | -0,28705 | -0,064588 | -4,18439 | 0,005159 | 0,0105306 |
| PDHK1 | -0,2881 | 0,170248 | -4,58055 | 0,003302 | 0,0074152 |
| NF-kB-p65_ | -0,29675 | -0,299014 | -4,50104 | 0,003605 | 0,0079502 |
| SOD2 | -0,31034 | 0,190406 | -5,81696 | 9,43E-04 | 0,0027094 |
| p38_MAPK | -0,31181 | 0,090537 | -5,08912 | 0,001925 | 0,0049116 |
| JNK_pT183 | -0,3144 | 0,055657 | -6,22359 | 6,51E-04 | 0,0020354 |
| Hexokinase | -0,32089 | 0,212545 | -5,7514 | 0,001003 | 0,0028377 |
| CD4 | -0,3285 | 0,210027 | -3,89766 | 0,007229 | 0,0135278 |
| Axl | -0,34568 | 0,238258 | -6,02639 | 7,77E-04 | 0,0023137 |
| p70-S6K1 | -0,34604 | 0,022775 | -6,00728 | 7,91E-04 | 0,0023264 |
| Tuberin_pT | -0,34867 | 0,137605 | -6,78324 | 4,02E-04 | 0,0013797 |
| B7-H4 | -0,35764 | 0,196086 | -7,28396 | 2,68E-04 | 9,60E-04 |
| MIF | -0,35975 | 0,027556 | -7,53573 | 2,21E-04 | 8,26E-04 |
| 53BP1 | -0,36826 | -0,440424 | -2,78543 | 0,030084 | 0,0461539 |
| Granzyme- | -0,36875 | 0,378485 | -4,9506 | 0,002222 | 0,0053806 |
| PKM2 | -0,36988 | 0,193229 | -5,66983 | 0,001084 | 0,0030082 |
| Collagen_V | -0,38936 | 0,359319 | -3,44023 | 0,012712 | 0,0216539 |
| Mnk1 | -0,39088 | 0,194965 | -5,10519 | 0,001894 | 0,0049116 |
| Merlin | -0,39201 | -0,407113 | -3,48727 | 0,011977 | 0,0206882 |
| PAR | -0,41043 | -0,597385 | -3,94903 | 0,006799 | 0,0130871 |
| b-Actin | -0,41302 | -0,225716 | -5,63988 | 0,001115 | 0,0030611 |
| BRD4 | -0,41397 | 0,114319 | -6,88796 | 3,69E-04 | 0,001283 |
| PDK1_pS24 | -0,42338 | 0,016233 | -6,74715 | 4,14E-04 | 0,0013834 |
| ULK1_pS75 | -0,42631 | 0,268754 | -6,28088 | 6,19E-04 | 0,0019593 |

| | | | | | |
|---|---|---|---|---|---|
| Atg3 | -0,46667 | -0,158672 | -4,70623 | 6,58E-04 | 0,0023222 |
| p44-42-MA | -0,48204 | -0,120539 | -3,22509 | 0,008167 | 0,0186778 |
| B7-H4 | -0,49165 | 0,008408 | -8,41028 | 4,28E-06 | 6,21E-05 |
| PKM2 | -0,5135 | -0,004977 | -2,67252 | 0,021835 | 0,0421338 |
| SHP-2_pY5 | -0,52558 | -0,033828 | -5,68291 | 1,46E-04 | 6,45E-04 |
| PDHK1 | -0,53004 | -0,09056 | -9,83574 | 9,32E-07 | 2,09E-05 |
| ULK1_pS75 | -0,60626 | 0,028512 | -6,83676 | 2,93E-05 | 2,07E-04 |
| B-Raf | -0,62237 | 0,028469 | -5,20918 | 2,98E-04 | 0,001188 |
| TFAM | -0,63463 | 0,208887 | -3,86464 | 0,002672 | 0,0076739 |
| Axl | -0,64741 | -0,084037 | -7,38521 | 1,45E-05 | 1,24E-04 |
| S6_pS240_ | -0,69437 | -0,196608 | -1,51629 | 0,157871 | 0,2166337 |
| WIPI2 | -0,71436 | 0,135735 | -3,69479 | 0,003582 | 0,00961 |
| Collagen_V | -0,71767 | 0,005751 | -7,86435 | 8,08E-06 | 8,92E-05 |
| PKC-alpha | -0,72069 | -0,189548 | -3,85003 | 0,00274 | 0,0077785 |
| MEK1 | -0,73088 | 0,271173 | -5,64225 | 1,55E-04 | 6,73E-04 |
| Stat3_pY70 | -0,7391 | -0,067062 | -7,84177 | 8,30E-06 | 8,92E-05 |
| NDRG1_pT | -0,74879 | 0,569197 | -2,08564 | 0,061304 | 0,0959359 |
| Caveolin-1 | -0,77492 | 0,094879 | -2,19405 | 0,050804 | 0,0831027 |
| HSP27_pS8 | -0,81976 | 0,102724 | -6,4187 | 5,14E-05 | 3,02E-04 |
| PMS2 | -0,83606 | 0,28608 | -7,01512 | 2,32E-05 | 1,79E-04 |
| IGFRb | -0,85389 | -0,226573 | -8,50891 | 3,82E-06 | 5,90E-05 |
| LDHA | -0,86714 | 0,035181 | -6,00222 | 9,21E-05 | 4,46E-04 |
| TSC1 | -0,88574 | -0,037883 | -6,09638 | 8,05E-05 | 4,06E-04 |
| Atg7 | -0,89547 | 0,079798 | -6,81611 | 3,01E-05 | 2,07E-04 |
| Granzyme- | -0,90222 | -0,138443 | -10,3752 | 5,48E-07 | 1,50E-05 |
| AMPK_pT1 | -0,98203 | -0,097621 | -5,5402 | 1,81E-04 | 7,69E-04 |
| Glutaminas | -0,9895 | -0,064521 | -5,18491 | 3,10E-04 | 0,0012104 |
| LC3A-B | -1,00888 | 0,145092 | -9,86349 | 9,06E-07 | 2,09E-05 |
| Gys_pS641 | -1,07199 | -0,079185 | -15,6395 | 8,13E-09 | 5,02E-07 |
| YAP_pS127 | -1,07328 | -0,052866 | -6,0415 | 8,70E-05 | 4,30E-04 |
| TFRC | -1,08901 | 0,387694 | -5,3443 | 2,43E-04 | 0,0010155 |
| Stat3 | -1,17491 | -0,383054 | -4,7085 | 6,56E-04 | 0,0023222 |
| MYH11 | -1,17492 | 0,096037 | -6,28033 | 6,22E-05 | 3,42E-04 |
| p70S6K_pT | -1,23478 | -0,039359 | -6,59755 | 4,03E-05 | 2,62E-04 |
| Cyclin_B1 | -1,30657 | 0,212369 | -12,9819 | 5,63E-08 | 2,32E-06 |
| Gys | -1,34938 | -0,101855 | -18,948 | 1,07E-09 | 1,33E-07 |
| Cox-IV | -1,37704 | 0,154718 | -11,6248 | 1,74E-07 | 6,15E-06 |
| Myosin-IIa_ | -1,42752 | 0,113416 | -7,65622 | 1,04E-05 | 1,03E-04 |
| PLK1 | -1,44787 | 0,096759 | -9,42633 | 1,42E-06 | 2,91E-05 |
| TRIM25 | -1,51165 | 0,044582 | -7,62208 | 1,08E-05 | 1,03E-04 |
| S6_pS235_ | -1,63268 | -0,119137 | -2,68167 | 0,021482 | 0,041779 |
| MCT4 | -2,06196 | -0,369449 | -8,01736 | 6,74E-06 | 8,49E-05 |
| PDGFR-b | -4,12457 | -0,927044 | -45,2463 | 9,13E-14 | 2,25E-11 |

| | | | | | |
|---|---|---|---|---|---|
| SHP-2_pY5 | -0,44676 | 0,128133 | -7,35017 | 2,55E-04 | 9,25E-04 |
| LDHA | -0,44762 | 0,476085 | -8,05559 | 1,50E-04 | 6,38E-04 |
| Glutamate- | -0,45124 | -0,004813 | -8,2162 | 1,34E-04 | 6,00E-04 |
| eEF2K | -0,4514 | 0,054703 | -3,15247 | 0,018425 | 0,0301385 |
| PREX1 | -0,45485 | 0,110573 | -4,87965 | 0,002394 | 0,0056852 |
| 4E-BP1 | -0,47156 | -0,159597 | -7,88824 | 1,69E-04 | 6,86E-04 |
| Cox2 | -0,47292 | -1,051796 | -7,54332 | 2,19E-04 | 8,26E-04 |
| P-Cadherin | -0,4765 | 0,075639 | -5,44259 | 0,00135 | 0,0035975 |
| Jagged1 | -0,48167 | -0,095837 | -9,99258 | 4,19E-05 | 2,20E-04 |
| MEK1 | -0,48188 | 0,58004 | -7,78747 | 1,83E-04 | 7,27E-04 |
| Caveolin-1 | -0,4844 | 0,438262 | -4,86887 | 0,002421 | 0,0056956 |
| MCT4 | -0,49846 | 1,018534 | -6,5153 | 5,04E-04 | 0,0016613 |
| PEA15 | -0,49975 | 0,049919 | -3,19957 | 0,017324 | 0,028719 |
| TFAM | -0,52316 | 0,414235 | -6,14579 | 6,98E-04 | 0,0021397 |
| WIPI2 | -0,52497 | 0,405226 | -2,88814 | 0,026181 | 0,0409279 |
| VASP | -0,5466 | -0,637872 | -6,32922 | 5,93E-04 | 0,001902 |
| YAP_pS127 | -0,59724 | 0,466929 | -11,6211 | 1,69E-05 | 1,16E-04 |
| B-Raf | -0,61625 | 0,167933 | -13,0013 | 8,51E-06 | 6,70E-05 |
| Gys | -0,69854 | 0,583061 | -13,0449 | 8,34E-06 | 6,70E-05 |
| PKC-alpha | -0,70746 | -0,024366 | -13,1365 | 7,99E-06 | 6,70E-05 |
| VEGFR-2 | -0,73937 | -0,022961 | -13,2104 | 7,72E-06 | 6,70E-05 |
| Gys_pS641 | -0,75884 | 0,342594 | -14,9396 | 3,62E-06 | 3,58E-05 |
| PAK4 | -0,77163 | 0,048855 | -14,9624 | 3,59E-06 | 3,58E-05 |
| S6_pS235_ | -0,77487 | 0,751764 | -4,92621 | 0,002279 | 0,0054662 |
| Glutaminas | -0,77798 | 0,278284 | -11,5562 | 1,75E-05 | 1,17E-04 |
| FAK_pY397 | -0,8357 | 0,097646 | -12,5867 | 1,04E-05 | 7,77E-05 |
| PMS2 | -0,87153 | 0,44721 | -10,5314 | 3,06E-05 | 1,80E-04 |
| Atg7 | -0,88232 | 0,28306 | -18,5744 | 9,40E-07 | 1,11E-05 |
| TFRC | -0,89641 | 0,740853 | -5,06731 | 0,001969 | 0,0049116 |
| HSP27_pS8 | -0,91256 | 0,225903 | -11,3629 | 1,93E-05 | 1,22E-04 |
| p70S6K_pT | -0,94837 | 0,401891 | -7,75291 | 1,87E-04 | 7,34E-04 |
| LC3A-B | -1,03592 | 0,348992 | -17,8253 | 1,21E-06 | 1,36E-05 |
| TSC1 | -1,05448 | 0,054131 | -20,1759 | 5,62E-07 | 8,67E-06 |
| MYH11 | -1,14658 | 0,369384 | -21,5179 | 3,76E-07 | 6,94E-06 |
| Cyclin_B1 | -1,16717 | 0,581082 | -23,7859 | 2,01E-07 | 4,51E-06 |
| NDRG1_pT | -1,23107 | 0,443207 | -22,2046 | 3,09E-07 | 6,36E-06 |
| Cox-IV | -1,23267 | 0,541782 | -18,8318 | 8,63E-07 | 1,11E-05 |
| PLK1 | -1,26306 | 0,523547 | -19,7823 | 6,35E-07 | 9,23E-06 |
| Myosin-IIa_ | -1,37016 | 0,459287 | -10,0217 | 4,12E-05 | 2,20E-04 |
| TRIM25 | -1,37016 | 0,459287 | -10,0217 | 4,12E-05 | 2,20E-04 |
| p16_INK4a | -1,65739 | -0,777213 | -27,0609 | 8,95E-08 | 2,76E-06 |
| Gab2 | -1,68271 | -0,540172 | -27,826 | 7,52E-08 | 2,65E-06 |
| PDGFR-b | -2,43191 | 0,988457 | -45,7566 | 3,29E-09 | 2,03E-07 |

**Table S7.** Changes in protein expression and/or phosphorylation levels occurring in *MMTV-R26 [Met]* MGT4 cells when exposed to single or combined treatment targeting BCL-XL (A1155463) and WEE1 (Adavosertib).

### MGT4 + A11 + ADAV vs. MGT4 + A11

| Protein | logFC | AveExpr | t | P.Value | adj.P.Val | B |
|---|---|---|---|---|---|---|
| PAR | 2,628347 | -0,447016 | 37,96932 | 9,93E-14 | 4,28E-11 | 22,00919 |
| Histone-H3 | 1,565677 | -0,22751 | 20,15493 | 1,59E-10 | 2,29E-08 | 14,67206 |
| CDK1_pT14 | 1,240134 | -0,263332 | 23,3577 | 2,90E-11 | 6,24E-09 | 16,42536 |
| Caspase-7-cleaved- | 1,200468 | 0,044123 | 18,16382 | 5,27E-10 | 3,78E-08 | 13,43042 |
| p53 | 1,089415 | 0,265969 | 12,73397 | 2,90E-08 | 1,14E-06 | 9,21298 |
| Caspase-3-cleaved | 1,049189 | -0,157487 | 12,46308 | 3,68E-08 | 1,32E-06 | 8,960888 |
| DM-Histone-H3 | 0,923131 | 0,044083 | 13,51466 | 1,50E-08 | 6,45E-07 | 9,913311 |
| JNK2 | 0,794061 | 0,368288 | 10,1844 | 3,32E-07 | 8,30E-06 | 6,627543 |
| H2AX_pS140 | 0,79091 | -0,061435 | 16,25579 | 1,87E-09 | 1,15E-07 | 12,10571 |
| KAP1 | 0,786688 | -0,155818 | 3,727147 | 0,002957 | 0,0144311 | -2,94911 |
| Vinculin | 0,759792 | -0,081115 | 2,868058 | 0,014318 | 0,0484051 | -4,54142 |
| p38-MAPK-_pT180_Y182 | 0,699031 | 0,014931 | 11,65955 | 7,66E-08 | 2,36E-06 | 8,18345 |
| H2AX_pS139 | 0,627217 | -0,076421 | 10,14326 | 3,47E-07 | 8,30E-06 | 6,581517 |
| RBM15 | 0,586131 | 0,009748 | 6,343481 | 3,95E-05 | 4,86E-04 | 1,557214 |
| Caveolin-1 | 0,497422 | -0,149569 | 3,132231 | 0,008791 | 0,0347618 | -4,05453 |
| S6_pS235_S236 | 0,492271 | 0,525909 | 2,696065 | 0,019659 | 0,0596703 | -4,85442 |
| Cyclophilin-F | 0,461977 | -0,349202 | 3,065413 | 0,009945 | 0,0375981 | -4,17819 |
| SOD1 | 0,44144 | -0,066911 | 7,220316 | 1,14E-05 | 1,97E-04 | 2,86881 |
| ER-a_pS118 | 0,422159 | -0,029565 | 4,553872 | 6,86E-04 | 0,0043459 | -1,43986 |
| CREB_pS133 | 0,416132 | 0,0246 | 5,047487 | 2,98E-04 | 0,0021796 | -0,57152 |
| p90RSK_pT573 | 0,409055 | 0,127706 | 2,299995 | 0,040483 | 0,1020363 | -5,55473 |
| MAPK_pT202_Y204 | 0,372642 | 0,259927 | 3,404308 | 0,005328 | 0,0244283 | -3,54909 |
| AMPKa_pT172 | 0,369015 | 0,069402 | 6,31148 | 4,14E-05 | 4,95E-04 | 1,507402 |
| Bcl2 | 0,36725 | 0,005942 | 3,090251 | 0,0095 | 0,0363294 | -4,13225 |
| S6 | 0,350749 | -0,221281 | 2,27055 | 0,042686 | 0,1063461 | -5,60527 |
| TRAP1 | 0,348811 | -0,205144 | 3,026304 | 0,01069 | 0,0390462 | -4,25044 |
| MLH1 | 0,344489 | -0,022308 | 4,364849 | 9,51E-04 | 0,0056929 | -1,77969 |
| Claudin-7 | 0,334078 | -0,262171 | 3,942495 | 0,002006 | 0,0107897 | -2,55099 |
| Caspase-8 | 0,325171 | -0,049334 | 2,832361 | 0,015293 | 0,0503401 | -4,60671 |
| c-Jun_pS73 | 0,324502 | -0,075111 | 4,255394 | 0,001152 | 0,0067998 | -1,97811 |
| C-Raf_pS338 | 0,313519 | -0,094016 | 3,878419 | 0,002251 | 0,0116171 | -2,66918 |
| ATP5A | 0,288493 | -0,041445 | 3,192491 | 0,007866 | 0,0326002 | -3,94279 |
| Myosin-IIa | 0,276154 | 0,125587 | 1,109796 | 0,289133 | 0,4311983 | -7,28761 |
| MTCO1 | 0,273638 | 0,022592 | 1,95599 | 0,07449 | 0,1583829 | -6,12722 |
| MDM2_pS166 | 0,271517 | 0,002366 | 3,674281 | 0,003254 | 0,0155852 | -3,04718 |
| Tau | 0,268593 | 0,023815 | 3,868508 | 0,002291 | 0,0116171 | -2,68748 |
| GRB7 | 0,257698 | -0,033547 | 3,184705 | 0,00798 | 0,0327566 | -3,95724 |
| MIG6 | 0,254672 | -0,034288 | 4,738494 | 5,00E-04 | 0,0033696 | -1,1117 |
| Bad_pS112 | 0,252335 | 0,10143 | 5,30182 | 1,97E-04 | 0,0016013 | -0,13596 |
| Wee1_pS642 | 0,250104 | 0,077241 | 4,470019 | 7,92E-04 | 0,0049491 | -1,59016 |
| SF2 | 0,248546 | -0,07376 | 2,060932 | 0,061995 | 0,1406317 | -5,95706 |
| Mitofusin-1 | 0,242395 | 0,041273 | 1,385346 | 0,191507 | 0,3275377 | -6,95901 |
| VHL-EPPK1 | 0,24024 | -0,032711 | 1,062236 | 0,309329 | 0,4488914 | -7,33848 |
| IRF-1 | 0,235718 | 0,017763 | 2,498176 | 0,028256 | 0,0787308 | -5,20861 |
| EGFR | 0,232973 | 0,023382 | 3,722864 | 0,00298 | 0,0144311 | -2,95705 |
| Ambra1_pS52 | 0,22697 | 0,096777 | 2,945236 | 0,012417 | 0,0438658 | -4,3998 |
| Annexin-VII | 0,224964 | -0,018589 | 2,973672 | 0,011782 | 0,0423152 | -4,34747 |
| SOD2 | 0,220183 | 0,043969 | 2,537959 | 0,026275 | 0,0749969 | -5,13801 |
| cdc25C | 0,219803 | 0,066872 | 3,056027 | 0,010119 | 0,0375981 | -4,19554 |
| Caspase-8-cleaved | 0,212609 | -0,011594 | 3,900048 | 0,002165 | 0,0113787 | -2,62926 |
| Fibronectin | 0,212218 | 0,223085 | 3,165356 | 0,00827 | 0,0333121 | -3,99313 |
| ADAR1 | 0,207581 | -0,023956 | 2,583851 | 0,024158 | 0,0708299 | -5,05617 |
| PAX8 | 0,206482 | 0,134307 | 2,679777 | 0,020257 | 0,0610549 | -4,88384 |
| eIF4E_pS209 | 0,204506 | 0,048507 | 4,403309 | 8,89E-04 | 0,0053993 | -1,71025 |
| GATA3 | 0,204245 | -0,050683 | 2,249395 | 0,04434 | 0,1098298 | -5,64141 |
| DNA-Ligase-IV | 0,202696 | 0,096328 | 1,72486 | 0,11055 | 0,2127097 | -6,48506 |
| Cyclin-B1 | 0,199964 | -5,99E-04 | 2,47646 | 0,029398 | 0,0801925 | -5,247 |
| EPHA2 | 0,197891 | -0,649423 | 4,186227 | 0,001301 | 0,0075762 | -2,10406 |
| Stathmin-1 | 0,186112 | -0,093131 | 3,249935 | 0,007076 | 0,0307184 | -3,83613 |
| p70-S6K_pT389 | 0,185756 | 0,269091 | 1,486528 | 0,163293 | 0,2849368 | -6,82504 |
| SETD2 | 0,183254 | -0,035564 | 2,919092 | 0,013031 | 0,0456613 | -4,44785 |
| Collagen-VI | 0,18283 | 0,063135 | 2,602209 | 0,023358 | 0,0694306 | -5,02332 |
| PKC-b-II_pS660 | 0,182643 | 0,022917 | 1,629754 | 0,129467 | 0,2374481 | -6,62459 |
| ATM | 0,182023 | 0,084602 | 1,735465 | 0,108601 | 0,2117958 | -6,4692 |
| 14-3-3-beta | 0,177872 | 0,022584 | 2,057712 | 0,062348 | 0,1406908 | -5,96235 |
| DUSP4 | 0,17682 | -0,042366 | 1,63932 | 0,127443 | 0,2347347 | -6,61078 |
| RIP3 | 0,169619 | -0,015111 | 2,76372 | 0,017357 | 0,0548696 | -4,73178 |
| D-a-Tubulin | 0,165056 | 0,01165 | 2,771548 | 0,017108 | 0,0548696 | -4,71755 |
| Aurora-A | 0,164932 | 0,026589 | 2,274496 | 0,042385 | 0,106208 | -5,59851 |
| N-Ras | 0,160593 | -0,002013 | 2,913742 | 0,01316 | 0,0457428 | -4,45767 |

### MGT4 + A11 + ADAV vs. MGT4 + ADAV

| Protein | logFC | AveExpr | t | P.Value | adj.P.Val | B |
|---|---|---|---|---|---|---|
| PAR | 1,77538 | -0,447016 | 25,64729 | 9,77E-12 | 4,21E-09 | 17,41064 |
| Histone-H3 | 1,24939 | -0,22751 | 16,08338 | 2,11E-09 | 4,54E-07 | 12,17029 |
| Caspase-3- | 0,923867 | -0,157487 | 10,97441 | 1,48E-07 | 1,60E-05 | 7,791781 |
| Caspase-7- | 0,852249 | 0,044123 | 12,89506 | 2,52E-08 | 3,63E-06 | 9,630045 |
| DM-Histon | 0,679048 | 0,044083 | 9,941275 | 4,30E-07 | 3,70E-05 | 6,681096 |
| KAP1 | 0,640602 | -0,155818 | 3,035028 | 0,010519 | 0,1237839 | -3,82686 |
| HES1 | 0,614045 | -0,136402 | 2,333775 | 0,038089 | 0,2534579 | -5,08557 |
| Histone-H3 | 0,476422 | 0,032165 | 6,248991 | 4,54E-05 | 0,002023 | 1,790387 |
| H2AX_pS14 | 0,413435 | -0,061435 | 8,497446 | 2,22E-06 | 1,60E-04 | 4,957271 |
| H2AX_pS13 | 0,399667 | -0,076421 | 6,46336 | 3,31E-05 | 0,0017839 | 2,120802 |
| ER-a_pS11 | 0,368771 | -0,029565 | 3,97797 | 0,001883 | 0,0450839 | -2,08416 |
| Aurora-AB( | 0,359293 | 0,130098 | 4,511615 | 7,37E-04 | 0,0244475 | -1,11808 |
| ERRalpha | 0,348421 | -0,060608 | 6,226119 | 4,69E-05 | 0,002023 | 1,75476 |
| CREB_pS13 | 0,316702 | 0,0246 | 3,841448 | 0,002405 | 0,0518363 | -2,33502 |
| Bcl2 | 0,303528 | 0,005942 | 2,554058 | 0,025512 | 0,1982695 | -4,69945 |
| Cdc6 | 0,285005 | 0,183566 | 4,31231 | 0,001042 | 0,0293093 | -1,47578 |
| Caveolin-1 | 0,265426 | -0,149569 | 1,671369 | 0,120864 | 0,4616099 | -6,15086 |
| Stathmin-1 | 0,25082 | -0,093131 | 4,37989 | 9,26E-04 | 0,0285206 | -1,35404 |
| RBM15 | 0,242631 | 0,009748 | 2,625912 | 0,022364 | 0,1939187 | -4,57124 |
| MLH1 | 0,241882 | -0,022308 | 3,064759 | 0,009957 | 0,1226175 | -3,77209 |
| Cyclophilin | 0,227372 | -0,349202 | 1,508709 | 0,157604 | 0,5031641 | -6,38106 |
| Sox2 | 0,223536 | 0,345513 | 3,856503 | 0,002341 | 0,0518363 | -2,3073 |
| SOD1 | 0,222717 | -0,066911 | 3,642829 | 0,003446 | 0,0550062 | -2,70177 |
| Cox2 | 0,219464 | 0,165493 | 1,294413 | 0,220215 | 0,553027 | -6,65926 |
| c-Myc | 0,211255 | -0,068852 | 1,897212 | 0,082467 | 0,3664253 | -5,80799 |
| SOD2 | 0,195935 | 0,043969 | 2,258454 | 0,043624 | 0,2645265 | -5,21476 |
| PARP | 0,194419 | -0,075124 | 3,713172 | 0,003033 | 0,0522848 | -2,57168 |
| Caspase-8 | 0,193593 | -0,049334 | 1,686268 | 0,117908 | 0,4616099 | -6,12903 |
| Fibronectir | 0,193584 | 0,223085 | 2,887415 | 0,013816 | 0,1452348 | -4,0977 |
| S6 | 0,190408 | -0,221281 | 1,232593 | 0,241658 | 0,5808818 | -6,73365 |
| PAX8 | 0,182963 | 0,134307 | 2,374533 | 0,035381 | 0,2450007 | -5,01501 |
| TRAP1 | 0,178947 | -0,205144 | 1,552559 | 0,146854 | 0,4944847 | -6,32052 |
| ATP5A | 0,175825 | -0,041445 | 1,945699 | 0,075833 | 0,3514424 | -5,73128 |
| MITF | 0,175276 | 0,104632 | 3,294253 | 0,006522 | 0,0878406 | -3,34778 |
| E-Cadherin | 0,172151 | -0,718665 | 0,390062 | 0,703421 | 0,9060647 | -7,42456 |
| Claudin-7 | 0,166088 | -0,262171 | 1,960017 | 0,07397 | 0,3465357 | -5,70844 |
| MTCO1 | 0,161766 | 0,022592 | 1,156321 | 0,270364 | 0,6298742 | -6,82152 |
| GRB7 | 0,15997 | -0,033547 | 1,97695 | 0,071822 | 0,3439474 | -5,68132 |
| Caspase-8- | 0,159073 | -0,011594 | 2,918004 | 0,013057 | 0,1442984 | -4,04174 |
| S100A4 | 0,15705 | 0,323293 | 2,0956 | 0,058317 | 0,3105905 | -5,48809 |
| Mcl-1 | 0,151739 | 0,126503 | 1,999326 | 0,069071 | 0,3382926 | -5,6453 |
| eEF2K | 0,149796 | 0,045657 | 1,455442 | 0,171561 | 0,5186339 | -6,45301 |
| IRF-1 | 0,149114 | 0,017763 | 1,580331 | 0,140378 | 0,4944847 | -6,28158 |
| DNA_POLG | 0,141829 | 0,048448 | 1,331618 | 0,208071 | 0,5484457 | -6,61318 |
| C-Raf_pS3 | 0,141795 | -0,094016 | 1,754088 | 0,105252 | 0,4161802 | -6,0282 |
| EGFR | 0,141214 | 0,023382 | 2,256576 | 0,043772 | 0,2645265 | -5,21796 |
| Bcl-xL | 0,139267 | -0,052409 | 3,339261 | 0,006004 | 0,0834711 | -3,26437 |
| GATA3 | 0,138063 | -0,050683 | 1,520525 | 0,154643 | 0,502496 | -6,36486 |
| Mitofusin-: | 0,136821 | 0,041273 | 0,781969 | 0,449599 | 0,8074041 | -7,18398 |
| SDHA | 0,132624 | -0,103483 | 1,987231 | 0,070546 | 0,3416321 | -5,66479 |
| Bid | 0,127081 | 0,009543 | 1,777647 | 0,101149 | 0,4074308 | -5,99263 |
| b-Actin | 0,124978 | -0,123301 | 1,32571 | 0,209962 | 0,5484457 | -6,62056 |
| Annexin-I | 0,12349 | -0,035592 | 1,300454 | 0,218205 | 0,553027 | -6,65185 |
| ACC_pS79 | 0,119903 | -0,237276 | 1,313618 | 0,213876 | 0,5519799 | -6,6356 |
| Cox-IV | 0,119143 | -0,046332 | 0,595235 | 0,562898 | 0,8406864 | -7,31756 |
| Annexin-VI | 0,118431 | -0,018589 | 1,565464 | 0,143813 | 0,4944847 | -6,30248 |
| SF2 | 0,115018 | -0,07376 | 0,953722 | 0,359303 | 0,7221004 | -7,03256 |
| IGFBP2 | 0,10919 | -0,058578 | 1,465885 | 0,168745 | 0,5158084 | -6,43905 |
| Transglutai | 0,108853 | 0,012303 | 0,825141 | 0,425615 | 0,787296 | -7,14842 |
| Tyro3 | 0,108323 | 0,007668 | 1,881758 | 0,084691 | 0,3724655 | -5,83223 |
| Heregulin | 0,107308 | 0,529459 | 1,337036 | 0,206349 | 0,5484457 | -6,60639 |
| DM-K9-His | 0,105642 | 0,012197 | 1,29297 | 0,220698 | 0,553027 | -6,66103 |
| Cyclin-B1 | 0,103239 | -5,99E-04 | 1,278566 | 0,22556 | 0,5561111 | -6,67859 |
| Coup-TFII | 0,102543 | 0,178384 | 1,444853 | 0,174457 | 0,5186339 | -6,46711 |
| PARG | 0,101702 | 2,89E-04 | 1,46844 | 0,168062 | 0,5158084 | -6,43562 |
| Collagen-V | 0,100183 | 0,063135 | 1,425896 | 0,179744 | 0,5199318 | -6,49215 |
| YAP | 0,098425 | 0,05181 | 1,853687 | 0,08887 | 0,3783138 | -5,87597 |
| XRCC1 | 0,097395 | -0,039169 | 1,241531 | 0,238459 | 0,5773912 | -6,72306 |
| Aurora-A | 0,097341 | 0,026589 | 1,342384 | 0,20466 | 0,5484457 | -6,59966 |
| TFAM | 0,096845 | 0,020213 | 1,434267 | 0,177393 | 0,5186339 | -6,48112 |

| | | | | | | |
|---|---|---|---|---|---|---|
| Notch1-cleaved | 0,158484 | -0,006006 | 3,237015 | 0,007247 | 0,0307184 | -3,86013 |
| Bid | 0,156599 | 0,009543 | 2,19056 | 0,049264 | 0,1186182 | -5,74123 |
| DNMT1 | 0,154231 | -0,244014 | 2,403719 | 0,033557 | 0,0886222 | -5,3748 |
| Notch3 | 0,153731 | -0,012206 | 3,150047 | 0,008507 | 0,0339491 | -4,02151 |
| PARG | 0,149382 | 2,89E-04 | 2,156876 | 0,052311 | 0,1232036 | -5,79787 |
| CDT1 | 0,148304 | 0,052388 | 2,370184 | 0,035661 | 0,0920362 | -5,43329 |
| c-Myc | 0,147521 | -0,068852 | 1,324838 | 0,210242 | 0,3481555 | -7,03587 |
| YB1_pS102 | 0,145909 | -0,035683 | 2,504083 | 0,027953 | 0,0787308 | -5,19815 |
| Src | 0,145128 | -0,207681 | 2,113457 | 0,056503 | 0,1309293 | -5,87032 |
| YAP | 0,144603 | 0,05181 | 2,723384 | 0,018695 | 0,0575552 | -4,80498 |
| Cdc42 | 0,143783 | 0,025328 | 3,121586 | 0,008966 | 0,0351291 | -4,07425 |
| Transglutaminase | 0,143116 | 0,012303 | 1,084873 | 0,299588 | 0,4421999 | -7,3145 |
| CD49b | 0,14114 | -0,01632 | 3,030203 | 0,010613 | 0,0390462 | -4,24324 |
| COG3 | 0,140287 | 0,05772 | 2,201553 | 0,048306 | 0,1169655 | -5,72266 |
| ACC_pS79 | 0,137803 | -0,237276 | 1,509724 | 0,157347 | 0,2756776 | -6,79341 |
| ACVRL1 | 0,136731 | -2,20E-04 | 2,577992 | 0,024419 | 0,0710245 | -5,06665 |
| Syk | 0,135082 | 0,002397 | 2,203825 | 0,04811 | 0,1169655 | -5,71882 |
| Annexin-I | 0,130742 | -0,035592 | 1,37682 | 0,19406 | 0,3305924 | -6,96999 |
| MSI2 | 0,130695 | -0,049706 | 1,753761 | 0,10531 | 0,2072541 | -6,44171 |
| Enolase-1 | 0,130538 | -0,337237 | 1,671938 | 0,12075 | 0,2272622 | -6,5633 |
| Rb | 0,129048 | 0,015017 | 2,10733 | 0,057119 | 0,1316497 | -5,88049 |
| MERIT40_pS29 | 0,127443 | -0,058898 | 1,981759 | 0,071222 | 0,1543987 | -6,08584 |
| PR | 0,124794 | 0,032851 | 1,604207 | 0,135011 | 0,2455262 | -6,66122 |
| FOXM1 | 0,120926 | 0,001348 | 2,420301 | 0,032562 | 0,0871685 | -5,34578 |
| S6_pS240_S244 | 0,116998 | 0,524502 | 0,984701 | 0,344479 | 0,4821139 | -7,41742 |
| PYGM | 0,115132 | -0,031874 | 1,512618 | 0,156619 | 0,2755208 | -6,78944 |
| p21 | 0,115076 | 0,016374 | 1,64396 | 0,126471 | 0,233945 | -6,60406 |
| ATR_pS428 | 0,113369 | -0,007961 | 1,726828 | 0,110186 | 0,2127097 | -6,48212 |
| CA9 | 0,112877 | 0,007759 | 1,93114 | 0,077772 | 0,1643119 | -6,16685 |
| CD4 | 0,11198 | 0,001958 | 0,807017 | 0,435579 | 0,5758734 | -7,57877 |
| CtIP | 0,111624 | -0,049514 | 2,168387 | 0,05125 | 0,1220383 | -5,77856 |
| XBP-1 | 0,111304 | 0,073656 | 2,002886 | 0,068643 | 0,1517183 | -6,05172 |
| GATA6 | 0,11099 | 0,007955 | 1,729934 | 0,109613 | 0,2127097 | -6,47748 |
| MRAP | 0,11001 | 0,030253 | 1,90027 | 0,082033 | 0,171633 | -6,21572 |
| VHL | 0,10966 | 0,077473 | 1,514189 | 0,156224 | 0,2755208 | -6,78729 |
| DAPK1_pS308 | 0,108951 | 0,013646 | 1,995388 | 0,069548 | 0,1521588 | -6,06385 |
| BiP-GRP78 | 0,106921 | 0,01833 | 1,916699 | 0,079739 | 0,1676473 | -6,18977 |
| Rheb | 0,106119 | -0,006062 | 2,004555 | 0,068443 | 0,1517183 | -6,04902 |
| 4E-BP1 | 0,105809 | -0,121582 | 1,78429 | 0,100018 | 0,2014382 | -6,39547 |
| Chk1_pS296 | 0,104451 | 0,030022 | 1,066203 | 0,307605 | 0,4488914 | -7,33431 |
| HER3 | 0,099278 | -0,121268 | 2,08075 | 0,059867 | 0,1365218 | -5,92445 |
| PRAS40 | 0,096575 | -0,009817 | 1,849889 | 0,08945 | 0,1844629 | -6,29457 |
| Complex-II-Subunit | 0,093413 | -0,044895 | 1,761166 | 0,104004 | 0,206595 | -6,43054 |
| Rad50 | 0,09257 | -0,199697 | 1,149163 | 0,273189 | 0,4160578 | -7,24415 |
| DNA_POLG | 0,091061 | 0,048448 | 0,854964 | 0,409549 | 0,553341 | -7,538 |
| IGFBP2 | 0,090832 | -0,058578 | 1,219425 | 0,246434 | 0,3849555 | -7,16359 |
| PD-L1 | 0,089645 | 0,04206 | 0,910622 | 0,380672 | 0,5241836 | -7,48808 |
| NAPSIN-A | 0,089171 | -0,101967 | 1,385731 | 0,191392 | 0,3275377 | -6,95851 |
| Chk2_pT68 | 0,087271 | -0,002757 | 1,324465 | 0,210362 | 0,3481555 | -7,03633 |
| 14-3-3-zeta | 0,086245 | -0,152677 | 1,296713 | 0,219448 | 0,3583302 | -7,07073 |
| Aurora-B | 0,085383 | 0,031936 | 1,371811 | 0,195573 | 0,3318584 | -6,97642 |
| CIITA | 0,084159 | -0,028549 | 1,312669 | 0,214186 | 0,3523437 | -7,05102 |
| BAP1 | 0,08283 | 0,003482 | 1,65086 | 0,125039 | 0,233297 | -6,59405 |
| Cyclin-D1 | 0,081793 | 0,034803 | 1,015438 | 0,330212 | 0,4728286 | -7,38673 |
| VEGF-164 | 0,081308 | 0,03077 | 1,646642 | 0,125913 | 0,2339157 | -6,60017 |
| IL-6 | 0,077952 | -0,01241 | 1,323006 | 0,210832 | 0,3481555 | -7,03815 |
| CD31 | 0,077369 | 0,01401 | 1,611638 | 0,133377 | 0,2435831 | -6,65061 |
| PDH | 0,074612 | 0,038497 | 1,3871 | 0,190985 | 0,3275377 | -6,95674 |
| HSP27 | 0,07056 | -0,024798 | 1,145432 | 0,27467 | 0,4168409 | -7,24832 |
| MIF | 0,06894 | -0,020019 | 1,455836 | 0,171454 | 0,2967739 | -6,86638 |
| Bim | 0,06879 | -0,479038 | 0,546765 | 0,594702 | 0,7184027 | -7,76222 |
| DM-K9-Histone-H3 | 0,065462 | 0,012197 | 0,801204 | 0,438807 | 0,5783662 | -7,58357 |
| PKC-a-b-II_pT638_T641 | 0,06133 | 0,193816 | 1,012725 | 0,331454 | 0,473035 | -7,38947 |
| MEK2 | 0,061318 | -0,028089 | 1,201854 | 0,252923 | 0,3893209 | -7,18409 |
| HSP70 | 0,061097 | -0,104212 | 1,115365 | 0,286835 | 0,4307522 | -7,28154 |
| Calnexin | 0,060677 | -0,064072 | 1,069006 | 0,306391 | 0,4488914 | -7,33135 |
| SOX17 | 0,060094 | -0,001645 | 1,23338 | 0,241375 | 0,382473 | -7,14715 |
| GCLM | 0,058653 | 0,022262 | 1,009472 | 0,332947 | 0,4735979 | -7,39275 |
| Bak | 0,058551 | -0,029684 | 1,035187 | 0,321276 | 0,4646641 | -7,36659 |
| FASN | 0,055383 | -0,105082 | 0,906937 | 0,382539 | 0,5250773 | -7,49147 |
| Hexokinase-I | 0,055313 | 0,008558 | 1,064861 | 0,308187 | 0,4488914 | -7,33572 |
| PAI-1 | 0,055135 | -0,036353 | 0,850796 | 0,41177 | 0,5546027 | -7,54163 |
| Patched | 0,053392 | -0,044418 | 0,814183 | 0,431622 | 0,5723966 | -7,57281 |
| TFAM | 0,053257 | 0,020213 | 0,788736 | 0,445783 | 0,5857693 | -7,59376 |
| Smad4 | 0,052687 | -0,014953 | 0,936875 | 0,367553 | 0,5081363 | -7,46358 |
| CDKN2A | 0,051102 | 0,017704 | 0,593121 | 0,564265 | 0,6988459 | -7,73437 |
| AceCS1 | 0,05108 | 0,175013 | 0,574185 | 0,576596 | 0,7040022 | -7,746 |

| | | | | | | |
|---|---|---|---|---|---|---|
| Lck | 0,095801 | -0,0272 | 1,382813 | 0,192263 | 0,5355361 | -6,5482 |
| CD44 | 0,091911 | -0,048103 | 1,116863 | 0,28622 | 0,6492664 | -6,86522 |
| DUSP4 | 0,090852 | -0,042366 | 0,8423 | 0,416321 | 0,7788793 | -7,13381 |
| GATA6 | 0,08737 | 0,007955 | 1,361785 | 0,198631 | 0,5484457 | -6,5751 |
| Vinculin | 0,081391 | -0,081115 | 0,307234 | 0,764006 | 0,9216856 | -7,45546 |
| COG3 | 0,079435 | 0,05772 | 1,246599 | 0,236659 | 0,5762717 | -6,71704 |
| Chk1_pS29 | 0,079142 | 0,030022 | 0,807856 | 0,435114 | 0,7946369 | -7,16286 |
| Hexokinase | 0,077651 | -0,321777 | 0,604989 | 0,556611 | 0,8406864 | -7,31142 |
| Ambra1_p! | 0,076344 | 0,096777 | 0,990665 | 0,341677 | 0,7012504 | -6,99661 |
| Cdc42 | 0,075523 | 0,025328 | 1,639634 | 0,127377 | 0,4628801 | -6,19695 |
| CD171 | 0,074913 | -0,179758 | 0,339746 | 0,740003 | 0,9216856 | -7,44419 |
| CD86 | 0,074342 | -0,059701 | 1,297426 | 0,219211 | 0,553027 | -6,65557 |
| Enolase-1 | 0,074317 | -0,337237 | 0,951851 | 0,360212 | 0,7221004 | -7,03435 |
| Notch1-cle | 0,074168 | -0,006006 | 1,514867 | 0,156054 | 0,502496 | -6,37263 |
| YAP_pS127 | 0,073161 | 0,902732 | 0,610637 | 0,552989 | 0,8406864 | -7,30782 |
| RPA32 | 0,071498 | -0,242082 | 1,134492 | 0,27905 | 0,6431574 | -6,84585 |
| DNA-Ligase | 0,071404 | 0,096328 | 0,607623 | 0,554921 | 0,8406864 | -7,30974 |
| c-Jun_pS73 | 0,070647 | -0,075111 | 0,926444 | 0,372726 | 0,7369042 | -7,05836 |
| D-a-Tubulir | 0,070607 | 0,01165 | 1,185596 | 0,259046 | 0,6101032 | -6,78831 |
| 4E-BP1 | 0,06702 | -0,121582 | 1,130174 | 0,280793 | 0,643733 | -6,85062 |
| FN14 | 0,067017 | 0,021789 | 0,89254 | 0,389895 | 0,7549078 | -7,08951 |
| U-Histone- | 0,066739 | 0,011354 | 0,599676 | 0,560031 | 0,8406864 | -7,31477 |
| SETD2 | 0,065918 | -0,035564 | 1,050022 | 0,314682 | 0,676246 | -6,93644 |
| Grp75 | 0,064968 | -0,171403 | 1,010149 | 0,332636 | 0,6925892 | -6,97718 |
| SOX17 | 0,064347 | -0,001645 | 1,320654 | 0,211591 | 0,5493724 | -6,62686 |
| RIP3 | 0,063802 | -0,015111 | 1,039567 | 0,319318 | 0,676246 | -6,94725 |
| Snail | 0,06212 | 0,389254 | 0,840201 | 0,41745 | 0,7788793 | -7,13561 |
| CDKN2A | 0,061682 | 0,017704 | 0,715931 | 0,487923 | 0,838916 | -7,23503 |
| Tuberin_p1 | 0,061148 | 0,059566 | 0,778142 | 0,451765 | 0,8079288 | -7,18705 |
| MERIT40_p | 0,05818 | -0,058898 | 0,904712 | 0,383669 | 0,7529167 | -7,07844 |
| Aurora-B | 0,057969 | 0,031936 | 0,931361 | 0,370281 | 0,7354429 | -7,05376 |
| p27-Kip1 | 0,057885 | 0,01314 | 0,643284 | 0,53231 | 0,8406864 | -7,2864 |
| PREX1 | 0,05325 | 0,008897 | 0,796223 | 0,441585 | 0,7988712 | -7,17243 |
| VHL-EPPK1 | 0,052152 | -0,032711 | 0,230596 | 0,821568 | 0,9334929 | -7,47761 |
| ADAR1 | 0,050936 | -0,023956 | 0,634024 | 0,53813 | 0,8406864 | -7,29258 |
| AR | 0,050638 | 0,027125 | 0,903435 | 0,384319 | 0,7529167 | -7,07961 |
| EPHA2 | 0,049966 | -0,649423 | 1,05699 | 0,31162 | 0,6761695 | -6,92918 |
| Rad50 | 0,049587 | -0,199697 | 0,615577 | 0,549831 | 0,8406864 | -7,30464 |
| N-Ras | 0,048613 | -0,002013 | 0,88201 | 0,395336 | 0,7549078 | -7,09898 |
| Wee1_pS6 | 0,048258 | 0,077241 | 0,862498 | 0,405556 | 0,7632947 | -7,11627 |
| cdc25C | 0,047852 | 0,066872 | 0,665306 | 0,518614 | 0,8406864 | -7,27136 |
| IR-b | 0,047754 | -0,023481 | 0,437321 | 0,669754 | 0,8984516 | -7,40372 |
| G6PD | 0,046291 | -0,105384 | 0,479332 | 0,64044 | 0,8984516 | -7,38327 |
| Hexokinase | 0,046108 | 0,008558 | 0,887644 | 0,392419 | 0,7549078 | -7,09393 |
| BiP-GRP78 | 0,045527 | 0,01833 | 0,816135 | 0,430547 | 0,7896418 | -7,15598 |
| SLC1A5 | 0,044073 | -0,020076 | 0,337309 | 0,741792 | 0,9216856 | -7,44507 |
| Tau | 0,043839 | 0,023815 | 0,631406 | 0,539782 | 0,8406864 | -7,29431 |
| SCD | 0,042921 | 0,046091 | 0,293097 | 0,774525 | 0,9216856 | -7,46002 |
| CD4 | 0,04268 | 0,001958 | 0,307589 | 0,763742 | 0,9216856 | -7,45534 |
| Hif-1-alpha | 0,041821 | 0,203091 | 0,392344 | 0,70178 | 0,9060647 | -7,42361 |
| HSP70 | 0,041687 | -0,104212 | 0,761019 | 0,461544 | 0,8134175 | -7,20062 |
| PRAS40 | 0,041436 | -0,009817 | 0,793706 | 0,442994 | 0,7988712 | -7,17448 |
| Bim | 0,040894 | -0,479038 | 0,325037 | 0,750828 | 0,9216856 | -7,44943 |
| Rb | 0,039722 | 0,015017 | 0,648649 | 0,528954 | 0,8406864 | -7,28278 |
| PD-1 | 0,038567 | 0,012291 | 0,678756 | 0,510351 | 0,8406864 | -7,26195 |
| Complex-II | 0,038169 | -0,044895 | 0,719624 | 0,485728 | 0,838916 | -7,23228 |
| GCLM | 0,037951 | 0,022262 | 0,653179 | 0,52613 | 0,8406864 | -7,2797 |
| Syk | 0,03754 | 0,002397 | 0,612455 | 0,551825 | 0,8406864 | -7,30665 |
| P-Cadherin | 0,037431 | 0,004036 | 0,400759 | 0,69574 | 0,9060647 | -7,42005 |
| CD49b | 0,037159 | -0,01632 | 0,797779 | 0,440716 | 0,7988712 | -7,17116 |
| HSP27 | 0,036715 | -0,024798 | 0,596016 | 0,562393 | 0,8406864 | -7,31707 |
| Cyclin-E1 | 0,036164 | 0,131132 | 0,413665 | 0,686518 | 0,9058509 | -7,41444 |
| Myosin-IIa | 0,034994 | 0,125587 | 0,140632 | 0,890526 | 0,9453616 | -7,49561 |
| STING | 0,034502 | 0,244125 | 0,152961 | 0,881006 | 0,9446507 | -7,49366 |
| Cyclin-D3 | 0,034441 | 0,043783 | 0,46426 | 0,650887 | 0,8984516 | -7,39081 |
| FOXM1 | 0,033707 | 0,001348 | 0,674636 | 0,512873 | 0,8406864 | -7,26485 |
| Merlin | 0,032685 | -0,057375 | 0,380642 | 0,710215 | 0,9083162 | -7,42844 |
| Notch3 | 0,030738 | -0,012206 | 0,62983 | 0,540778 | 0,8406864 | -7,29535 |
| MEK1 | 0,030651 | -0,124464 | 0,435224 | 0,671233 | 0,8984516 | -7,4047 |
| RRM1 | 0,030246 | -0,036564 | 0,552337 | 0,590999 | 0,8634592 | -7,34345 |
| BAP1 | 0,029835 | 0,003482 | 0,594628 | 0,56329 | 0,8406864 | -7,31794 |
| PD-L1 | 0,029369 | 0,04206 | 0,298333 | 0,770623 | 0,9216856 | -7,45835 |
| CDT1 | 0,028754 | 0,052388 | 0,459549 | 0,654169 | 0,8984516 | -7,39312 |
| CD29 | 0,028668 | 0,008186 | 0,539145 | 0,599785 | 0,8733357 | -7,35104 |
| LRP6_pS14 | 0,027337 | 0,099524 | 0,369061 | 0,718602 | 0,9115925 | -7,43307 |
| CA9 | 0,027105 | 0,007759 | 0,463726 | 0,651259 | 0,8984516 | -7,39108 |
| ERCC5 | 0,026896 | 0,04876 | 0,306631 | 0,764453 | 0,9216856 | -7,45566 |

| | | | | | | |
|---|---|---|---|---|---|---|
| c-Abl | 0,048813 | 0,06195 | 0,666521 | 0,517864 | 0,6584053 | -7,68593 |
| AR | 0,047109 | 0,027125 | 0,840475 | 0,417303 | 0,5603042 | -7,55054 |
| PI3K-p110-b | 0,044352 | -0,043716 | 0,696791 | 0,499395 | 0,640593 | -7,66442 |
| DAPK2 | 0,041487 | 0,006769 | 0,828256 | 0,423917 | 0,5639149 | -7,56097 |
| MelanA | 0,040197 | -0,07913 | 0,498708 | 0,627127 | 0,7364894 | -7,78881 |
| E2F1 | 0,039477 | -0,015219 | 0,67728 | 0,511254 | 0,6519245 | -7,67838 |
| N-Cadherin | 0,039105 | 0,036274 | 0,749226 | 0,468355 | 0,6080152 | -7,6251 |
| SDHA | 0,037689 | -0,103483 | 0,564726 | 0,582808 | 0,709577 | -7,75168 |
| UQCRC2 | 0,036529 | -0,059451 | 0,777295 | 0,452246 | 0,5906607 | -7,60299 |
| WIPI2 | 0,03641 | -0,02897 | 0,276899 | 0,786636 | 0,857131 | -7,88071 |
| HLA-DR-DP-DQ-DX | 0,035343 | -0,01581 | 0,62175 | 0,5459 | 0,6870819 | -7,7161 |
| PD-1 | 0,034964 | 0,012291 | 0,615337 | 0,549984 | 0,6870819 | -7,72026 |
| IRS2 | 0,03452 | 0,497198 | 0,366643 | 0,720358 | 0,8191933 | -7,84971 |
| Jagged1 | 0,034389 | -0,002552 | 0,762569 | 0,460653 | 0,5998229 | -7,61468 |
| BRD4 | 0,034241 | -0,067661 | 0,390973 | 0,702766 | 0,8034269 | -7,83985 |
| 14-3-3-epsilon | 0,033517 | 0,009189 | 0,537422 | 0,600937 | 0,7234746 | -7,76757 |
| YAP_pS127 | 0,03327 | 0,902732 | 0,277689 | 0,786044 | 0,857131 | -7,88048 |
| Cox-IV | 0,03326 | -0,046332 | 0,166166 | 0,870832 | 0,9148983 | -7,90717 |
| XPA | 0,03231 | 0,037377 | 0,531151 | 0,605142 | 0,7256483 | -7,77111 |
| Bim_1 | 0,031572 | -0,20752 | 0,385155 | 0,706957 | 0,8060805 | -7,84226 |
| CD134 | 0,030915 | -5,21E-04 | 0,462164 | 0,652347 | 0,7598955 | -7,80746 |
| Cdc6 | 0,029744 | 0,183566 | 0,450054 | 0,660806 | 0,7676748 | -7,81334 |
| Gli1 | 0,026904 | 0,045725 | 0,472737 | 0,645001 | 0,7533757 | -7,8022 |
| PDHK1 | 0,026736 | -0,003486 | 0,43086 | 0,674315 | 0,7812626 | -7,82235 |
| Histone-H3_pS10 | 0,025865 | 0,032165 | 0,339264 | 0,740356 | 0,8288143 | -7,86007 |
| XRCC1 | 0,025248 | -0,039169 | 0,321849 | 0,753182 | 0,8388149 | -7,86625 |
| CD38 | 0,025135 | -0,02424 | 0,30849 | 0,763073 | 0,8454618 | -7,87077 |
| MEK1_p_S217_S221 | 0,022659 | 0,022895 | 0,349353 | 0,732963 | 0,8248228 | -7,85634 |
| Grp75 | 0,019198 | -0,171403 | 0,298493 | 0,770504 | 0,8515054 | -7,87403 |
| TFRC | 0,018874 | -0,023369 | 0,199007 | 0,845636 | 0,90023 | -7,90069 |
| Enolase-2 | 0,018453 | -7,87E-04 | 0,278176 | 0,785679 | 0,857131 | -7,88033 |
| SFRP1 | 0,018111 | 0,054441 | 0,343537 | 0,737222 | 0,8274549 | -7,85851 |
| S100A4 | 0,017364 | 0,323293 | 0,231702 | 0,820728 | 0,8821289 | -7,89309 |
| Myt1 | 0,01455 | -0,055936 | 0,242365 | 0,812647 | 0,8756272 | -7,89037 |
| AMPK-a2_pS345 | 0,012675 | -0,111954 | 0,111822 | 0,912839 | 0,9367472 | -7,91533 |
| Ets-1 | 0,010683 | -0,022047 | 0,226562 | 0,824631 | 0,8841192 | -7,89436 |
| P-Cadherin | 0,008282 | 0,004036 | 0,088668 | 0,930829 | 0,9484329 | -7,91785 |
| TAZ | 0,004935 | 0,101754 | 0,093992 | 0,926689 | 0,946452 | -7,91732 |
| UGT1A | 0,004202 | 0,268485 | 0,061073 | 0,952321 | 0,963498 | -7,92008 |
| TUFM | 0,002065 | 0,039828 | 0,025695 | 0,979929 | 0,9844974 | -7,92174 |
| CD29 | 0,001455 | 0,008186 | 0,027363 | 0,978626 | 0,9844974 | -7,9217 |
| Granzyme-B | -1,43E-05 | 0,194614 | -1,67E-04 | 0,99987 | 0,9998699 | -7,9221 |
| CD44 | -0,00108 | -0,048103 | -0,01308 | 0,989782 | 0,9920842 | -7,92201 |
| PREX1 | -0,00274 | 0,008897 | -0,04091 | 0,968053 | 0,9771211 | -7,9212 |
| CD20 | -0,00443 | -0,016963 | -0,06167 | 0,951859 | 0,963498 | -7,92004 |
| Creb | -0,00689 | -0,033283 | -0,11874 | 0,907476 | 0,9366229 | -7,91447 |
| Rab11 | -0,00727 | -0,004335 | -0,11758 | 0,908372 | 0,9366229 | -7,91462 |
| NDUFB4 | -0,00742 | -0,047676 | -0,0856 | 0,933217 | 0,9486234 | -7,91813 |
| Glutaminase | -0,00773 | 0,32211 | -0,10419 | 0,918765 | 0,9405882 | -7,91623 |
| LC3A-B | -0,00793 | -0,040287 | -0,15307 | 0,880923 | 0,9215483 | -7,90943 |
| RIP | -0,00839 | 0,030057 | -0,12257 | 0,904506 | 0,9366229 | -7,91397 |
| p27_pT157 | -0,00864 | 0,027337 | -0,16407 | 0,872444 | 0,9148983 | -7,90754 |
| Puma | -0,01034 | -0,083658 | -0,12816 | 0,900173 | 0,9348782 | -7,91321 |
| MMP2 | -0,01059 | -0,02074 | -0,18605 | 0,855555 | 0,9050349 | -7,90338 |
| CD45 | -0,01091 | -0,03593 | -0,13195 | 0,897239 | 0,934082 | -7,91268 |
| Notch1 | -0,01174 | -0,054826 | -0,1945 | 0,849086 | 0,9013696 | -7,90165 |
| HSP60 | -0,01259 | -0,120894 | -0,19863 | 0,845924 | 0,90023 | -7,90077 |
| NQO1 | -0,01405 | -0,013101 | -0,11318 | 0,911783 | 0,9367472 | -7,91517 |
| Glutamate-D1-2 | -0,01573 | 0,097722 | -0,357 | 0,727381 | 0,8206834 | -7,85345 |
| U-Histone-H2B | -0,01654 | 0,011354 | -0,14862 | 0,884354 | 0,9228969 | -7,91015 |
| PAK1 | -0,01795 | 0,121987 | -0,31199 | 0,760475 | 0,8447543 | -7,8696 |
| Lck | -0,0183 | -0,0272 | -0,26408 | 0,796261 | 0,8622829 | -7,88444 |
| TRIP13 | -0,01846 | -0,047949 | -0,27571 | 0,787526 | 0,857131 | -7,88107 |
| ERCC5 | -0,01861 | 0,04876 | -0,21214 | 0,835612 | 0,8936689 | -7,89778 |
| FOXO3 | -0,01893 | 0,039835 | -0,18357 | 0,857459 | 0,9050349 | -7,90388 |
| LRP6_pS1490 | -0,02046 | 0,099524 | -0,27617 | 0,787181 | 0,857131 | -7,88093 |
| ATM_pS1981 | -0,022 | -0,044366 | -0,26952 | 0,792169 | 0,860012 | -7,88288 |
| c-Kit | -0,02353 | -0,017567 | -0,3334 | 0,744666 | 0,8314791 | -7,86219 |
| SCD | -0,02662 | 0,046091 | -0,18177 | 0,858838 | 0,9050349 | -7,90423 |
| Porin | -0,02718 | -0,05308 | -0,583 | 0,570837 | 0,7018063 | -7,74063 |
| RPA32_pS4_S8 | -0,02793 | 0,003 | -0,42086 | 0,6814 | 0,7857182 | -7,82689 |
| INPP4b | -0,02835 | -0,042678 | -0,57718 | 0,574635 | 0,7036018 | -7,74418 |
| VASP | -0,02857 | 0,087681 | -0,42029 | 0,681807 | 0,7857182 | -7,82715 |
| Mnk1 | -0,02874 | 0,189059 | -0,35786 | 0,726748 | 0,8206834 | -7,85312 |
| p38-a | -0,03069 | -0,037713 | -0,59052 | 0,565952 | 0,6989258 | -7,73599 |
| ENY2 | -0,03274 | -0,11689 | -0,54623 | 0,595057 | 0,7184027 | -7,76252 |
| Beclin | -0,03315 | 0,065552 | -0,72247 | 0,484039 | 0,6246131 | -7,64549 |

| | | | | | | |
|---|---|---|---|---|---|---|
| Mnk1 | 0,026279 | 0,189059 | 0,327209 | 0,749226 | 0,9216856 | -7,44867 |
| PR | 0,024936 | 0,032851 | 0,320548 | 0,754143 | 0,9216856 | -7,45098 |
| XBP-1 | 0,024663 | 0,073656 | 0,4438 | 0,665194 | 0,8984516 | -7,40069 |
| HER3 | 0,024184 | -0,121268 | 0,506872 | 0,621558 | 0,8900047 | -7,36888 |
| MEK1_p_S | 0,024074 | 0,022895 | 0,371172 | 0,71707 | 0,9115925 | -7,43224 |
| PDH | 0,023612 | 0,038497 | 0,438967 | 0,668594 | 0,8984516 | -7,40295 |
| Chk1 | 0,022605 | -0,197309 | 0,395465 | 0,699537 | 0,9060647 | -7,42229 |
| ATR_pS428 | 0,022486 | -0,007961 | 0,342503 | 0,73798 | 0,9216856 | -7,44318 |
| NAPSIN-A | 0,021669 | -0,101967 | 0,336743 | 0,742208 | 0,9216856 | -7,44528 |
| MRAP | 0,019173 | 0,030253 | 0,33119 | 0,746293 | 0,9216856 | -7,44726 |
| A-Raf_pS29 | 0,018922 | 0,023098 | 0,284755 | 0,780755 | 0,9216856 | -7,4626 |
| NRF2 | 0,018762 | -0,109994 | 0,230723 | 0,821471 | 0,9334929 | -7,47758 |
| eIF4E | 0,017942 | 0,084361 | 0,279318 | 0,784823 | 0,9216856 | -7,46425 |
| 14-3-3-beta | 0,01776 | 0,022584 | 0,205453 | 0,840711 | 0,9334929 | -7,48351 |
| PDHK1 | 0,017523 | -0,003486 | 0,282392 | 0,782522 | 0,9216856 | -7,46332 |
| PYGM | 0,015961 | -0,031874 | 0,209693 | 0,837474 | 0,9334929 | -7,48256 |
| BRD4 | 0,015721 | -0,067661 | 0,179509 | 0,860576 | 0,9390078 | -7,48889 |
| N-Cadherin | 0,013995 | 0,036274 | 0,268145 | 0,793206 | 0,9236947 | -7,46754 |
| MIG6 | 0,013683 | -0,034288 | 0,254587 | 0,803413 | 0,9283403 | -7,47135 |
| c-Abl | 0,013308 | 0,06195 | 0,181709 | 0,858887 | 0,9390078 | -7,48847 |
| Src | 0,012614 | -0,207681 | 0,183699 | 0,85736 | 0,9390078 | -7,48807 |
| VHL | 0,012525 | 0,077473 | 0,172944 | 0,865619 | 0,9407674 | -7,49014 |
| DAPK1_pS3 | 0,012352 | 0,013646 | 0,226228 | 0,824885 | 0,9334929 | -7,47868 |
| INPP4b | 0,012192 | -0,042678 | 0,248189 | 0,808243 | 0,9314245 | -7,47308 |
| IRS1 | 0,011322 | 0,130579 | 0,149376 | 0,883773 | 0,9446507 | -7,49424 |
| MIF | 0,010669 | -0,020019 | 0,225312 | 0,825581 | 0,9334929 | -7,4789 |
| Chk2 | 0,009345 | 0,017318 | 0,079352 | 0,938079 | 0,9695736 | -7,50289 |
| Jagged1 | 0,009333 | -0,002552 | 0,206953 | 0,839566 | 0,9334929 | -7,48318 |
| Bak | 0,008164 | -0,029684 | 0,144345 | 0,887658 | 0,9446507 | -7,49504 |
| UQCRC2 | 0,007903 | -0,059451 | 0,168173 | 0,869287 | 0,9407674 | -7,49102 |
| MelanA | 0,007046 | -0,07913 | 0,087413 | 0,931806 | 0,9677306 | -7,50216 |
| Calnexin | 0,006476 | -0,064072 | 0,114087 | 0,911082 | 0,9634726 | -7,49926 |
| Rab11 | 0,006038 | -0,004335 | 0,097628 | 0,923862 | 0,9664673 | -7,50114 |
| IL-6 | 0,002941 | -0,01241 | 0,04991 | 0,961027 | 0,9815226 | -7,50495 |
| 4E-BP1_pS | 0,002867 | 0,087254 | 0,06086 | 0,952486 | 0,9797651 | -7,50429 |
| Ets-1 | 0,002202 | -0,022047 | 0,0467 | 0,963531 | 0,9817537 | -7,50511 |
| HLA-DR-DP | 0,002019 | -0,01581 | 0,035511 | 0,972264 | 0,98338 | -7,50561 |
| Glutaminas | 0,001874 | 0,32211 | 0,02525 | 0,980277 | 0,9848467 | -7,50595 |
| CtIP | 0,001697 | -0,049514 | 0,032963 | 0,974253 | 0,98338 | -7,5057 |
| c-Abl_pY41 | 0,001207 | 0,027149 | 0,015088 | 0,988214 | 0,9905119 | -7,50617 |
| PEA-15 | -9,00E-04 | 0,046253 | -0,01204 | 0,990591 | 0,9905914 | -7,50621 |
| TUFM | -0,00207 | 0,039828 | -0,02577 | 0,979869 | 0,9848467 | -7,50593 |
| MMP14 | -0,00221 | -0,048973 | -0,03419 | 0,973293 | 0,98338 | -7,50566 |
| Bad_pS112 | -0,00291 | 0,10143 | -0,06122 | 0,952205 | 0,9797651 | -7,50427 |
| Bim_1 | -0,00343 | -0,20752 | -0,04185 | 0,967317 | 0,983287 | -7,50535 |
| p21 | -0,00365 | 0,016374 | -0,05213 | 0,959296 | 0,9815226 | -7,50482 |
| PEA-15_pS | -0,00498 | -0,061818 | -0,05254 | 0,958974 | 0,9815226 | -7,5048 |
| MMP2 | -0,00527 | -0,02074 | -0,09262 | 0,927756 | 0,9677306 | -7,50166 |
| MTSS1 | -0,0055 | 0,031705 | -0,08813 | 0,931247 | 0,9677306 | -7,50209 |
| GCN5L2 | -0,00582 | -0,152105 | -0,11148 | 0,913107 | 0,9634726 | -7,49958 |
| Notch1 | -0,006 | -0,054826 | -0,09932 | 0,922543 | 0,9664673 | -7,50096 |
| CD134 | -0,00669 | -5,21E-04 | -0,10006 | 0,921969 | 0,9664673 | -7,50088 |
| DAPK2 | -0,00723 | 0,006769 | -0,14434 | 0,887665 | 0,9446507 | -7,49504 |
| Atg3 | -0,00855 | 0,143336 | -0,18255 | 0,858238 | 0,9390078 | -7,4883 |
| VAV1 | -0,00942 | -0,053778 | -0,08274 | 0,935443 | 0,9691734 | -7,50259 |
| TRIP13 | -0,01005 | -0,047949 | -0,15009 | 0,883224 | 0,9446507 | -7,49413 |
| UVRAG | -0,01046 | 0,098398 | -0,16605 | 0,870919 | 0,9407674 | -7,4914 |
| TAZ | -0,01086 | 0,101754 | -0,20688 | 0,83962 | 0,9334929 | -7,48319 |
| Connexin-4 | -0,01109 | -0,052007 | -0,20308 | 0,842526 | 0,9334929 | -7,48403 |
| Chk2_pT68 | -0,01126 | -0,002757 | -0,17091 | 0,86718 | 0,9407674 | -7,49052 |
| PCNA | -0,01142 | -0,015273 | -0,22479 | 0,825977 | 0,9334929 | -7,47903 |
| HSP60 | -0,01235 | -0,120894 | -0,1948 | 0,848858 | 0,9380966 | -7,48581 |
| 14-3-3-eps | -0,01304 | 0,009189 | -0,20904 | 0,83797 | 0,9334929 | -7,48271 |
| PLK1 | -0,01322 | -0,086862 | -0,10995 | 0,914293 | 0,9634726 | -7,49976 |
| cdc2_pY15 | -0,01332 | -0,23913 | -0,18235 | 0,858397 | 0,9390078 | -7,48834 |
| TTF1 | -0,01375 | -0,024739 | -0,21028 | 0,83703 | 0,9334929 | -7,48243 |
| LDHA | -0,01514 | -0,094454 | -0,15648 | 0,878295 | 0,9446507 | -7,49307 |
| PAI-1 | -0,01557 | -0,036353 | -0,24027 | 0,814234 | 0,9334929 | -7,47516 |
| PI3K-p85 | -0,01773 | 0,139813 | -0,29183 | 0,775467 | 0,9216856 | -7,46041 |
| Smad1 | -0,01782 | 0,042839 | -0,29273 | 0,7748 | 0,9216856 | -7,46013 |
| MSH6 | -0,018 | -0,01407 | -0,26778 | 0,793476 | 0,9236947 | -7,46764 |
| LAD1 | -0,01827 | -0,060637 | -0,28425 | 0,781131 | 0,9216856 | -7,46276 |
| ENY2 | -0,01914 | -0,11689 | -0,31929 | 0,755076 | 0,9216856 | -7,45141 |
| Rheb | -0,0195 | -0,006062 | -0,36834 | 0,719122 | 0,9115925 | -7,43336 |
| E2F1 | -0,02116 | -0,015219 | -0,3631 | 0,722936 | 0,9137405 | -7,43541 |
| Axl | -0,02181 | -0,091902 | -0,20745 | 0,839182 | 0,9334929 | -7,48307 |
| Smac | -0,02201 | -0,021718 | -0,26562 | 0,795106 | 0,9236947 | -7,46826 |

| | | | | | | |
|---|---|---|---|---|---|---|
| ERCC1 | -0,03329 | -0,05099 | -0,60719 | 0,555201 | 0,6906695 | -7,72549 |
| MSH6 | -0,03372 | -0,01407 | -0,5017 | 0,62508 | 0,7364894 | -7,78722 |
| XPF | -0,03464 | 0,014147 | -0,65202 | 0,52685 | 0,6659012 | -7,69591 |
| PEA-15 | -0,03548 | 0,046253 | -0,47475 | 0,643608 | 0,7533757 | -7,80119 |
| MEK1 | -0,03586 | -0,124464 | -0,50914 | 0,620013 | 0,7361582 | -7,78323 |
| A-Raf_pS299 | -0,03724 | 0,023098 | -0,56041 | 0,585652 | 0,711031 | -7,75424 |
| TRIM25 | -0,03884 | -0,075383 | -0,29147 | 0,775736 | 0,8550956 | -7,87626 |
| eIF4E | -0,03966 | 0,084361 | -0,61738 | 0,548679 | 0,6870819 | -7,71894 |
| 4E-BP1_pS65 | -0,0405 | 0,087254 | -0,85972 | 0,407028 | 0,5516632 | -7,53385 |
| NRF2 | -0,04068 | -0,109994 | -0,50021 | 0,626098 | 0,7364894 | -7,78801 |
| PDHA1 | -0,04123 | -0,038326 | -0,58192 | 0,571541 | 0,7018063 | -7,74129 |
| FAK | -0,046 | -0,154643 | -0,60585 | 0,556061 | 0,6906695 | -7,72635 |
| Atg3 | -0,04635 | 0,143336 | -0,98955 | 0,3422 | 0,4821139 | -7,41263 |
| Chk2 | -0,04795 | 0,017318 | -0,40719 | 0,691138 | 0,7922348 | -7,83293 |
| G6PD | -0,0489 | -0,105384 | -0,50634 | 0,62192 | 0,7363942 | -7,78474 |
| Bax | -0,04902 | -0,033866 | -0,94164 | 0,365208 | 0,5077567 | -7,45908 |
| VEGFR-2 | -0,05228 | 0,135944 | -0,51965 | 0,612894 | 0,729716 | -7,77751 |
| CD86 | -0,05365 | -0,059701 | -0,93629 | 0,367839 | 0,5081363 | -7,46413 |
| UVRAG | -0,05463 | 0,098398 | -0,86756 | 0,402887 | 0,547774 | -7,52695 |
| C-Raf | -0,05464 | -0,023119 | -0,9846 | 0,344527 | 0,4821139 | -7,41752 |
| Gli3 | -0,05536 | 2,76E-05 | -0,89343 | 0,389435 | 0,5328465 | -7,50379 |
| Axl | -0,05542 | -0,091902 | -0,52721 | 0,607794 | 0,7256483 | -7,77332 |
| FoxO3a_pS318_S321 | -0,05552 | 0,075277 | -0,87614 | 0,398394 | 0,5433789 | -7,51934 |
| p27-Kip1 | -0,05562 | 0,01314 | -0,61814 | 0,548194 | 0,6870819 | -7,71845 |
| b-Catenin_pT41_S45 | -0,05894 | -0,033569 | -1,02025 | 0,328017 | 0,4712516 | -7,38185 |
| FN14 | -0,06271 | 0,021789 | -0,83517 | 0,420168 | 0,5623988 | -7,55509 |
| Smac | -0,06479 | -0,021718 | -0,78191 | 0,449629 | 0,5890278 | -7,59928 |
| PKC-delta_pS664 | -0,06491 | -0,045451 | -1,1406 | 0,276599 | 0,4182959 | -7,25371 |
| MTSS1 | -0,06642 | 0,031705 | -1,06452 | 0,308337 | 0,4488914 | -7,33608 |
| RRM1 | -0,06676 | -0,036564 | -1,2192 | 0,246514 | 0,3849555 | -7,16385 |
| PEA-15_pS116 | -0,06865 | -0,061818 | -0,72444 | 0,482873 | 0,6246131 | -7,64401 |
| oct-04 | -0,06945 | -0,096692 | -0,24856 | 0,807962 | 0,8727614 | -7,88873 |
| HER2 | -0,07422 | -0,099699 | -0,52788 | 0,607338 | 0,7256483 | -7,77294 |
| PI3K-p85 | -0,07433 | 0,139813 | -1,22322 | 0,245051 | 0,3849555 | -7,15914 |
| TTF1 | -0,07948 | -0,024739 | -1,21583 | 0,247749 | 0,3854873 | -7,1678 |
| PLK1 | -0,07955 | -0,086862 | -0,66161 | 0,520897 | 0,6603134 | -7,68933 |
| Rictor_pT1135 | -0,08016 | 0,247853 | -1,47281 | 0,166898 | 0,2900529 | -6,84359 |
| Chk1_pS345 | -0,08126 | 0,094896 | -1,22724 | 0,24359 | 0,3845689 | -7,1544 |
| TIGAR | -0,08135 | -0,036293 | -1,24129 | 0,238546 | 0,3799484 | -7,13777 |
| GSK-3a-b_pS21_S9 | -0,08229 | 0,350958 | -1,03118 | 0,323074 | 0,4657027 | -7,3707 |
| PDK1_pS241 | -0,08381 | 0,104414 | -1,52827 | 0,152728 | 0,2742739 | -6,76789 |
| LAD1 | -0,08522 | -0,060637 | -1,32578 | 0,209938 | 0,3481555 | -7,03468 |
| GCLC | -0,0873 | -0,02283 | -1,21022 | 0,249816 | 0,3873051 | -7,17436 |
| Smad3 | -0,08751 | -0,00225 | -1,26743 | 0,229378 | 0,3702701 | -7,10643 |
| PHGDH | -0,08832 | -0,245847 | -0,68661 | 0,505559 | 0,6465758 | -7,67175 |
| CD171 | -0,09058 | -0,179758 | -0,41079 | 0,688567 | 0,7913926 | -7,83136 |
| PCNA | -0,09148 | -0,015273 | -1,80117 | 0,097195 | 0,1970097 | -6,3697 |
| Elk1_pS383 | -0,09276 | 0,002093 | -1,08983 | 0,297484 | 0,4406034 | -7,30918 |
| UBQLN4 | -0,09283 | -0,037809 | -1,09481 | 0,295384 | 0,4390016 | -7,30383 |
| B-Raf_pS445 | -0,09538 | 0,108475 | -2,15781 | 0,052225 | 0,1232036 | -5,79631 |
| TSC1 | -0,09703 | 0,030918 | -1,76109 | 0,104017 | 0,206595 | -6,43065 |
| Tuberin_pT1462 | -0,1018 | 0,059566 | -1,29542 | 0,21988 | 0,3583302 | -7,07232 |
| c-Abl_pY412 | -0,10324 | 0,027149 | -1,29055 | 0,221507 | 0,3589076 | -7,07828 |
| Lasu1 | -0,10349 | -0,040635 | -1,97293 | 0,072326 | 0,1558632 | -6,10004 |
| Rab25 | -0,10376 | 0,029024 | -1,16243 | 0,267973 | 0,4095612 | -7,22923 |
| BMK1-Erk5_pT218_Y220 | -0,10552 | 0,018014 | -1,80016 | 0,097362 | 0,1970097 | -6,37124 |
| B-Raf | -0,10806 | 0,001943 | -1,25705 | 0,232982 | 0,3746839 | -7,11892 |
| ATRX | -0,10841 | 0,050761 | -0,98604 | 0,34385 | 0,4821139 | -7,41611 |
| Cyclin-E1 | -0,10864 | 0,131132 | -1,24274 | 0,238029 | 0,3799484 | -7,13604 |
| PRAS40_pT246 | -0,10991 | 0,125751 | -2,13437 | 0,054446 | 0,1273271 | -5,8355 |
| MR1 | -0,11097 | 0,031104 | -0,82882 | 0,423612 | 0,5639149 | -7,5605 |
| CSK | -0,11147 | 0,00617 | -1,82065 | 0,094028 | 0,1929812 | -6,33979 |
| Myosin-IIa_pS1943 | -0,11326 | -0,007141 | -0,96548 | 0,353625 | 0,4932439 | -7,43621 |
| DRP1 | -0,11511 | 0,073839 | -2,38655 | 0,034619 | 0,0904295 | -5,40478 |
| LDHA | -0,11512 | -0,094454 | -1,18991 | 0,257411 | 0,3948196 | -7,19789 |
| B7-H3 | -0,11647 | -0,017587 | -1,66719 | 0,121704 | 0,2280618 | -6,57025 |
| DJ1 | -0,11723 | -0,237303 | -1,20642 | 0,251223 | 0,38809 | -7,17878 |
| BCL2A1 | -0,11938 | -0,035601 | -1,24029 | 0,2389 | 0,3799484 | -7,13895 |
| Pdcd4 | -0,12498 | 0,463311 | -1,2941 | 0,220319 | 0,3583302 | -7,07393 |
| ULK1_pS757 | -0,12592 | 0,319938 | -2,34976 | 0,037004 | 0,0942433 | -5,46876 |
| Hexokinase-II | -0,1285 | -0,321777 | -1,00113 | 0,3368 | 0,4775027 | -7,40112 |
| PMS2 | -0,1285 | -0,057549 | -1,52496 | 0,153543 | 0,2745932 | -6,77246 |
| eEF2 | -0,12857 | 0,086276 | -1,98123 | 0,071288 | 0,1543987 | -6,0867 |
| p44-42-MAPK | -0,12865 | -0,015364 | -1,11057 | 0,288815 | 0,4311983 | -7,28678 |
| PKCa | -0,12908 | 0,324233 | -2,46911 | 0,029794 | 0,0805153 | -5,25997 |
| MERIT40 | -0,12921 | -0,004042 | -1,35436 | 0,200922 | 0,3382707 | -6,99868 |
| PKM2 | -0,1303 | -0,316375 | -2,0262 | 0,065897 | 0,1479243 | -6,01385 |

| | | | | | | |
|---|---|---|---|---|---|---|
| VEGF-164 | -0,0225 | 0,03077 | -0,45574 | 0,656828 | 0,8984516 | -7,39498 |
| PKC-delta_ | -0,0231 | -0,045451 | -0,40587 | 0,692084 | 0,9060647 | -7,41785 |
| CD38 | -0,02375 | -0,02424 | -0,29152 | 0,775699 | 0,9216856 | -7,46051 |
| Elk1_pS38: | -0,02471 | 0,002093 | -0,29035 | 0,776576 | 0,9216856 | -7,46088 |
| Myt1 | -0,02563 | -0,055936 | -0,42693 | 0,677093 | 0,8993053 | -7,4085 |
| YB1_pS102 | -0,02588 | -0,035683 | -0,44413 | 0,664964 | 0,8984516 | -7,40053 |
| ARID1A | -0,02634 | -0,041475 | -0,23253 | 0,820103 | 0,9334929 | -7,47713 |
| Glutamate- | -0,02647 | 0,097722 | -0,60081 | 0,559302 | 0,8406864 | -7,31406 |
| IGFRb | -0,02756 | 0,161739 | -0,38599 | 0,706352 | 0,9060647 | -7,42624 |
| CD20 | -0,02824 | -0,016963 | -0,39313 | 0,701211 | 0,9060647 | -7,42327 |
| CD31 | -0,02923 | 0,01401 | -0,60879 | 0,554171 | 0,8406864 | -7,30899 |
| AMPK-a2_ | -0,02925 | -0,111954 | -0,25807 | 0,80079 | 0,9277965 | -7,47039 |
| Stat3 | -0,0301 | 0,455546 | -0,26584 | 0,79494 | 0,9236947 | -7,4682 |
| A-Raf | -0,03046 | -0,001133 | -0,48284 | 0,63802 | 0,8984516 | -7,38148 |
| PI3K-p110- | -0,03069 | -0,043716 | -0,48218 | 0,638473 | 0,8984516 | -7,38181 |
| XPA | -0,03157 | 0,037377 | -0,51895 | 0,613365 | 0,8841479 | -7,36233 |
| FASN | -0,03189 | -0,105082 | -0,52224 | 0,611144 | 0,8839029 | -7,36052 |
| c-Kit | -0,0321 | -0,017767 | -0,45476 | 0,65751 | 0,8984516 | -7,39545 |
| DNMT1 | -0,03266 | -0,244014 | -0,50902 | 0,620094 | 0,8900047 | -7,36772 |
| FGF-basic | -0,03273 | 0,070481 | -0,46194 | 0,652506 | 0,8984516 | -7,39196 |
| PDK1_pS24 | -0,03406 | 0,104414 | -0,62099 | 0,546383 | 0,8406864 | -7,30114 |
| b-Catenin_ | -0,03481 | -0,033569 | -0,60254 | 0,558185 | 0,8406864 | -7,31296 |
| Creb | -0,03563 | -0,033283 | -0,61429 | 0,550651 | 0,8406864 | -7,30547 |
| Smad4 | -0,03589 | -0,014953 | -0,6382 | 0,535498 | 0,8406864 | -7,2898 |
| Atg7 | -0,03594 | -0,03646 | -0,41261 | 0,68727 | 0,9058509 | -7,41491 |
| Puma | -0,03601 | -0,083658 | -0,44623 | 0,663488 | 0,8984516 | -7,39954 |
| SFRP1 | -0,03692 | 0,054441 | -0,7004 | 0,497222 | 0,840403 | -7,24644 |
| TRIM25 | -0,03733 | -0,075383 | -0,28018 | 0,784178 | 0,9216856 | -7,46399 |
| PHGDH | -0,03738 | -0,245847 | -0,2906 | 0,776386 | 0,9216856 | -7,4608 |
| XIAP | -0,0377 | 0,151229 | -0,57579 | 0,575542 | 0,8467108 | -7,32952 |
| PKC-a-b-II_ | -0,03815 | 0,193816 | -0,63003 | 0,54065 | 0,8406864 | -7,29522 |
| RRM2 | -0,03856 | 0,068136 | -0,42793 | 0,676386 | 0,8993053 | -7,40805 |
| 14-3-3-zeta | -0,03932 | -0,152677 | -0,59116 | 0,565537 | 0,8406864 | -7,3201 |
| Cyclin-D1 | -0,03934 | 0,034803 | -0,48836 | 0,63422 | 0,8984516 | -7,37864 |
| Melanoma | -0,04041 | -0,02963 | -0,61095 | 0,552792 | 0,8406864 | -7,30762 |
| Jak2 | -0,04052 | 0,117213 | -0,46591 | 0,64974 | 0,8984516 | -7,39 |
| Smad3 | -0,04057 | -0,00225 | -0,58763 | 0,567826 | 0,8410075 | -7,32228 |
| Rad51 | -0,04059 | 0,041121 | -0,38651 | 0,705979 | 0,9060647 | -7,42603 |
| PDHA1 | -0,04079 | -0,038326 | -0,5757 | 0,575606 | 0,8467108 | -7,32958 |
| ATR | -0,04326 | -0,111535 | -0,42547 | 0,67813 | 0,8993053 | -7,40916 |
| Patched | -0,04336 | -0,044418 | -0,66118 | 0,521163 | 0,8406864 | -7,27421 |
| DDB-1 | -0,04551 | -0,14144 | -0,5306 | 0,60551 | 0,8787029 | -7,35587 |
| HLA-DQA1 | -0,04576 | -0,058912 | -0,44999 | 0,660853 | 0,8984516 | -7,39775 |
| DJ1 | -0,04596 | -0,237303 | -0,47295 | 0,644852 | 0,8984516 | -7,38649 |
| CD45 | -0,04603 | -0,03593 | -0,55679 | 0,588051 | 0,8620749 | -7,34085 |
| TIGAR | -0,04622 | -0,036293 | -0,70536 | 0,49424 | 0,838916 | -7,24282 |
| CIITA | -0,04667 | -0,028549 | -0,72799 | 0,480781 | 0,838916 | -7,22602 |
| WIPI1 | -0,04676 | 0,170801 | -0,81812 | 0,429454 | 0,7896418 | -7,15431 |
| NQO1 | -0,04801 | -0,013101 | -0,38672 | 0,705829 | 0,9060647 | -7,42595 |
| MSI2 | -0,0481 | -0,049706 | -0,64548 | 0,530934 | 0,8406864 | -7,28492 |
| HER2 | -0,04935 | -0,099699 | -0,351 | 0,731758 | 0,9194979 | -7,44003 |
| HMHA1 | -0,04968 | -0,095438 | -0,62103 | 0,54636 | 0,8406864 | -7,30111 |
| MSH2 | -0,05021 | -0,305486 | -1,08214 | 0,30075 | 0,6647354 | -6,90266 |
| AMPKa_pT | -0,05151 | 0,069402 | -0,88103 | 0,395845 | 0,7549078 | -7,09986 |
| p38-a | -0,05154 | -0,037713 | -0,99185 | 0,341123 | 0,7012504 | -6,99544 |
| RIP | -0,0529 | 0,030057 | -0,773 | 0,45469 | 0,8097984 | -7,19116 |
| ATM_pS19 | -0,05387 | -0,044366 | -0,65984 | 0,521991 | 0,8406864 | -7,27514 |
| p44-42-MA | -0,05544 | -0,015364 | -0,47859 | 0,640954 | 0,8984516 | -7,38365 |
| CD26 | -0,05677 | -0,017648 | -1,023 | 0,326769 | 0,6836772 | -6,96419 |
| Pdcd4 | -0,05707 | 0,463311 | -0,59097 | 0,565659 | 0,8406864 | -7,32021 |
| C-Raf | -0,05729 | -0,023119 | -1,03231 | 0,322566 | 0,6781759 | -6,9547 |
| EGFR_pY11 | -0,05841 | 0,002371 | -0,86987 | 0,401674 | 0,7595346 | -7,10978 |
| p27_pT157 | -0,05885 | 0,027337 | -1,11707 | 0,286136 | 0,6492664 | -6,865 |
| UBAC1 | -0,06053 | -0,032503 | -1,06014 | 0,310244 | 0,6761695 | -6,92589 |
| BMK1-Erk5 | -0,06084 | 0,018014 | -1,03786 | 0,320079 | 0,676246 | -6,949 |
| DUSP6 | -0,06199 | 0,129979 | -1,041 | 0,31868 | 0,676246 | -6,94577 |
| NDUFB4 | -0,06276 | -0,047676 | -0,72411 | 0,483069 | 0,838916 | -7,22893 |
| Beclin | -0,0634 | 0,065552 | -1,3817 | 0,192594 | 0,5355361 | -6,54963 |
| MR1 | -0,06499 | 0,031104 | -0,48545 | 0,636224 | 0,8984516 | -7,38014 |
| TFRC | -0,06518 | -0,023369 | -0,68724 | 0,505179 | 0,8406864 | -7,25592 |
| eIF4E_pS2( | -0,06587 | 0,048507 | -1,41837 | 0,18188 | 0,5226028 | -6,50203 |
| FAK | -0,06604 | -0,154643 | -0,86964 | 0,401796 | 0,7595346 | -7,10999 |
| UGT1A | -0,067 | 0,268485 | -0,9738 | 0,349646 | 0,7108366 | -7,01317 |
| MERIT40 | -0,06756 | -0,004042 | -0,70813 | 0,492579 | 0,838916 | -7,24079 |
| Stat3_pY7( | -0,06769 | 0,23668 | -0,75956 | 0,462383 | 0,8134175 | -7,20176 |
| AceCS1 | -0,06829 | 0,175013 | -0,76761 | 0,457764 | 0,8119186 | -7,19543 |
| Atg4B | -0,06833 | 0,127254 | -1,27786 | 0,225799 | 0,5561111 | -6,67945 |

| | | | | | | |
|---|---|---|---|---|---|---|
| JAB1 | -0,13096 | 0,007015 | -1,74349 | 0,107147 | 0,20991 | -6,45717 |
| Melanoma-gp100 | -0,13262 | -0,02963 | -2,00519 | 0,068366 | 0,1517183 | -6,04798 |
| UBAC1 | -0,13402 | -0,032503 | -2,34725 | 0,037173 | 0,0942433 | -5,47311 |
| IGFRb | -0,13476 | 0,161739 | -1,88747 | 0,083862 | 0,1746119 | -6,23586 |
| AMPKa | -0,13744 | 0,044799 | -2,49132 | 0,028612 | 0,0790486 | -5,22074 |
| FGF-basic | -0,1388 | 0,070481 | -1,95918 | 0,074079 | 0,1583829 | -6,12212 |
| Bcl-xL | -0,1428 | -0,052409 | -3,42397 | 0,005139 | 0,0238166 | -3,51251 |
| mTOR_pS2448 | -0,14429 | 0,087497 | -1,99768 | 0,06927 | 0,1521588 | -6,06015 |
| EGFR_pY1173 | -0,14614 | 0,002371 | -2,17636 | 0,050528 | 0,1209862 | -5,76516 |
| NDRG1_pT346 | -0,14764 | 0,175484 | -1,68748 | 0,117672 | 0,223421 | -6,54049 |
| Atg7 | -0,14892 | -0,03646 | -1,70957 | 0,113414 | 0,2162903 | -6,50781 |
| IR-b | -0,14912 | -0,023481 | -1,3656 | 0,197461 | 0,3337486 | -6,98436 |
| eIF4G | -0,15161 | 0,075302 | -2,36542 | 0,03597 | 0,0922811 | -5,44157 |
| ACSL1 | -0,15219 | 0,059466 | -2,85931 | 0,014551 | 0,0486177 | -4,55744 |
| PAK4 | -0,15378 | 0,010678 | -3,08881 | 0,009525 | 0,0363294 | -4,13492 |
| GSK-3a-b | -0,15507 | 0,145096 | -2,83211 | 0,015301 | 0,0503401 | -4,60717 |
| HLA-DQA1 | -0,15593 | -0,058912 | -1,53343 | 0,151463 | 0,2731411 | -6,76075 |
| STING | -0,15757 | 0,244125 | -0,69858 | 0,498317 | 0,640593 | -7,66313 |
| E-Cadherin | -0,15879 | -0,718665 | -0,35979 | 0,725345 | 0,8206834 | -7,85238 |
| p27_pT198 | -0,1621 | -0,028398 | -1,80739 | 0,096174 | 0,1964504 | -6,36017 |
| c-IAP2 | -0,16608 | -0,079066 | -2,51557 | 0,027372 | 0,0776151 | -5,17778 |
| b-Catenin | -0,16648 | -0,058081 | -2,96857 | 0,011893 | 0,0423626 | -4,35686 |
| PI3K-p110-a | -0,1675 | 0,200034 | -2,40674 | 0,033374 | 0,0886222 | -5,36952 |
| PTPN12 | -0,16914 | 0,448663 | -2,9857 | 0,011523 | 0,0417335 | -4,32532 |
| HMHA1 | -0,17056 | -0,095438 | -2,13223 | 0,054653 | 0,1273271 | -5,83907 |
| 53BP1 | -0,17105 | 0,410082 | -1,95516 | 0,074598 | 0,1583829 | -6,12855 |
| ARID1A | -0,1716 | -0,041475 | -1,51473 | 0,15609 | 0,2755208 | -6,78655 |
| WIPI1 | -0,17492 | 0,170801 | -3,0604 | 0,010038 | 0,0375981 | -4,18746 |
| Atg5 | -0,17764 | 0,237612 | -2,24504 | 0,044687 | 0,1100586 | -5,64884 |
| Mitofusin-2 | -0,17782 | 0,07772 | -1,8598 | 0,087944 | 0,18223 | -6,27915 |
| Atg4B | -0,18092 | 0,127254 | -3,38343 | 0,005536 | 0,0251152 | -3,58793 |
| CD26 | -0,18096 | -0,017648 | -3,26078 | 0,006936 | 0,0305055 | -3,81598 |
| IDO | -0,18098 | -0,045778 | -1,56092 | 0,144877 | 0,2623608 | -6,72244 |
| Raptor | -0,18612 | 0,029375 | -2,88077 | 0,013986 | 0,0478425 | -4,51815 |
| Paxillin | -0,18614 | -0,099849 | -2,60589 | 0,023201 | 0,0694306 | -5,01673 |
| Merlin | -0,18912 | -0,057375 | -2,20243 | 0,04823 | 0,1169655 | -5,72118 |
| Gys | -0,19124 | 0,329948 | -2,46735 | 0,02989 | 0,0805153 | -5,26306 |
| Erk5 | -0,19318 | -0,046955 | -2,55467 | 0,025484 | 0,0732234 | -5,10826 |
| PYGB | -0,19334 | 0,004676 | -3,3156 | 0,006271 | 0,0281526 | -3,71409 |
| RSK | -0,19347 | -0,552241 | -3,24239 | 0,007175 | 0,0307184 | -3,85016 |
| Slfn11 | -0,1971 | -0,015227 | -1,34841 | 0,202773 | 0,3400593 | -7,00623 |
| A-Raf | -0,20048 | -0,001133 | -3,17752 | 0,008087 | 0,0328805 | -3,97057 |
| SHP2 | -0,2016 | -0,044912 | -2,76312 | 0,017376 | 0,0548696 | -4,73287 |
| HER3_pY1289 | -0,20666 | 3,22E-04 | -3,93653 | 0,002028 | 0,0107897 | -2,56198 |
| HSP27_pS82 | -0,21049 | -0,021664 | -1,51486 | 0,156057 | 0,2755208 | -6,78637 |
| B7-H4 | -0,21369 | -0,039993 | -2,49705 | 0,028314 | 0,0787308 | -5,21059 |
| 4E-BP1_pT37_T46 | -0,2144 | -0,149083 | -2,75225 | 0,017727 | 0,0553663 | -4,75262 |
| NF-kB-p65_pS536 | -0,21507 | 0,118054 | -4,07391 | 0,001587 | 0,0089997 | -2,30944 |
| GSK-3B | -0,21579 | 0,32798 | -2,09856 | 0,058013 | 0,1329982 | -5,89503 |
| Aurora-ABC_pT288_pT23 | -0,21695 | 0,130098 | -2,72428 | 0,018665 | 0,0575552 | -4,80337 |
| IRS1 | -0,21722 | 0,130579 | -2,8659 | 0,014376 | 0,0484051 | -4,54537 |
| MSH2 | -0,21838 | -0,305486 | -4,70709 | 5,28E-04 | 0,0034993 | -1,16725 |
| SLC1A5 | -0,21948 | -0,020076 | -1,67974 | 0,119195 | 0,2253199 | -6,55186 |
| ACC1 | -0,21957 | -0,14479 | -4,01458 | 0,001764 | 0,0097457 | -2,41834 |
| Tuberin | -0,22095 | 0,001667 | -2,78026 | 0,016836 | 0,054558 | -4,7017 |
| ER-a | -0,22121 | -0,077341 | -2,58447 | 0,024131 | 0,0708299 | -5,05508 |
| DDR1 | -0,22555 | -0,018116 | -3,87368 | 0,00227 | 0,0116171 | -2,67794 |
| Gys_pS641 | -0,23075 | 0,273752 | -2,80554 | 0,016069 | 0,052468 | -4,65566 |
| Rb_pS807_S811 | -0,23533 | 0,274772 | -4,59662 | 6,37E-04 | 0,0040988 | -1,36353 |
| IGF1R_pY1135_Y1136 | -0,23661 | -0,141052 | -3,83021 | 0,002455 | 0,0123014 | -2,75826 |
| EMA | -0,24261 | 0,046986 | -1,11883 | 0,285412 | 0,430114 | -7,27775 |
| ATR | -0,24415 | -0,111535 | -2,40103 | 0,033722 | 0,0886222 | -5,3795 |
| Gab2 | -0,24626 | 0,045631 | -2,8928 | 0,013679 | 0,047166 | -4,4961 |
| Akt_pT308 | -0,25003 | 0,127704 | -3,61772 | 0,003607 | 0,0168977 | -3,15222 |
| PARP | -0,25049 | -0,075124 | -4,78407 | 4,63E-04 | 0,0031692 | -1,0313 |
| ASNS | -0,25176 | -0,022748 | -3,78878 | 0,002645 | 0,0131036 | -2,83492 |
| Akt_pS473 | -0,25672 | 0,399518 | -2,57498 | 0,024554 | 0,0710245 | -5,07203 |
| ZAP-70 | -0,26283 | -0,001713 | -2,37703 | 0,035222 | 0,0914502 | -5,42138 |
| MCT4 | -0,27109 | 0,021705 | -4,12357 | 0,001453 | 0,0083507 | -2,21851 |
| DDB-1 | -0,28114 | -0,14144 | -3,27753 | 0,006726 | 0,0298838 | -3,78485 |
| Akt2 | -0,28563 | 0,225659 | -4,60106 | 6,32E-04 | 0,0040988 | -1,35562 |
| XIAP | -0,292 | 0,151229 | -4,4592 | 8,07E-04 | 0,0049705 | -1,6096 |
| PDK1 | -0,29333 | -0,003713 | -3,23528 | 0,00727 | 0,0307184 | -3,86335 |
| Smad1 | -0,29718 | 0,042839 | -4,88157 | 3,93E-04 | 0,0028245 | -0,86011 |
| Cox2 | -0,29779 | 0,165493 | -1,75637 | 0,104848 | 0,2072541 | -6,43778 |
| Cyclin-D3 | -0,30003 | 0,043783 | -4,04432 | 0,001673 | 0,0093629 | -2,36373 |
| Rad23A | -0,30217 | -0,014213 | -1,71281 | 0,112802 | 0,2160781 | -6,503 |

| | | | | | | |
|---|---|---|---|---|---|---|
| Paxillin | -0,07021 | -0,099849 | -0,9829 | 0,345331 | 0,7053908 | -7,00427 |
| JAB1 | -0,07169 | 0,007015 | -0,95434 | 0,359005 | 0,7221004 | -7,03197 |
| MAPK_pT2 | -0,07337 | 0,259927 | -0,67024 | 0,515574 | 0,8406864 | -7,26793 |
| Tuberin | -0,07413 | 0,001667 | -0,93279 | 0,369572 | 0,7354429 | -7,05241 |
| ACC1 | -0,07419 | -0,14479 | -1,35655 | 0,200244 | 0,5484457 | -6,58176 |
| B7-H3 | -0,07435 | -0,017587 | -1,06426 | 0,308448 | 0,6761695 | -6,92157 |
| p27_pT198 | -0,07463 | -0,028398 | -0,8321 | 0,421831 | 0,7836602 | -7,14253 |
| CSK | -0,07506 | 0,00617 | -1,22592 | 0,24407 | 0,5811843 | -6,74151 |
| UBQLN4 | -0,07516 | -0,037809 | -0,8865 | 0,393009 | 0,7549078 | -7,09496 |
| p38-MAPK | -0,07553 | 0,104174 | -1,27125 | 0,228063 | 0,5584958 | -6,68746 |
| eIF4G | -0,07654 | 0,075302 | -1,19416 | 0,255807 | 0,6057853 | -6,77848 |
| PAK1 | -0,07698 | 0,121987 | -1,33781 | 0,206103 | 0,5484457 | -6,60541 |
| Rab25 | -0,0791 | 0,029024 | -0,88615 | 0,393191 | 0,7549078 | -7,09527 |
| PI3K-p110- | -0,07947 | 0,200034 | -1,14186 | 0,276096 | 0,6397701 | -6,83768 |
| CDK1_pT14 | -0,08039 | -0,263332 | -1,51417 | 0,156228 | 0,502496 | -6,37358 |
| b-Catenin | -0,0806 | -0,058081 | -1,43733 | 0,176541 | 0,5186339 | -6,47708 |
| Gli1 | -0,08148 | 0,045725 | -1,43177 | 0,178092 | 0,5186339 | -6,48442 |
| DDR1 | -0,08456 | -0,018116 | -1,45234 | 0,172404 | 0,5186339 | -6,45715 |
| LC3A-B | -0,08521 | -0,040287 | -1,64471 | 0,126315 | 0,4628801 | -6,18961 |
| Enolase-2 | -0,0863 | -7,87E-04 | -1,30098 | 0,218032 | 0,553027 | -6,6512 |
| p38-MAPK- | -0,08633 | 0,014931 | -1,43997 | 0,175807 | 0,5186339 | -6,47358 |
| Porin | -0,08667 | -0,05308 | -1,85912 | 0,088046 | 0,3783138 | -5,86753 |
| TSC1 | -0,09035 | 0,030918 | -1,63988 | 0,127326 | 0,4628801 | -6,1966 |
| eEF2 | -0,09043 | 0,086276 | -1,39348 | 0,189097 | 0,535152 | -6,53444 |
| MEK2 | -0,09143 | -0,028089 | -1,79209 | 0,098705 | 0,4013369 | -5,97068 |
| PMS2 | -0,09192 | -0,057549 | -1,09079 | 0,29708 | 0,6600066 | -6,89343 |
| Gli3 | -0,09223 | 2,76E-05 | -1,48857 | 0,162762 | 0,509804 | -6,40847 |
| PAICS | -0,09402 | -0,15572 | -1,55957 | 0,145196 | 0,4944847 | -6,31074 |
| HSP27_pS8 | -0,09473 | -0,021664 | -0,68177 | 0,508509 | 0,8406864 | -7,25982 |
| Wee1 | -0,09497 | -0,01496 | -0,7051 | 0,494396 | 0,838916 | -7,24301 |
| NDRG1_pT | -0,09594 | 0,175484 | -1,09657 | 0,294646 | 0,6600066 | -6,88722 |
| Lasu1 | -0,09597 | -0,040635 | -1,82963 | 0,0926 | 0,3801027 | -5,91318 |
| PYGB | -0,09771 | 0,004676 | -1,67562 | 0,120014 | 0,4616099 | -6,14464 |
| Bax | -0,09977 | -0,033866 | -1,91636 | 0,079786 | 0,362617 | -5,77782 |
| oct-04 | -0,09997 | -0,096692 | -0,35778 | 0,726812 | 0,9159527 | -7,43746 |
| Granzyme- | -0,10128 | 0,194614 | -1,17624 | 0,262624 | 0,615167 | -6,799 |
| AMPKa | -0,10175 | 0,044799 | -1,84434 | 0,090303 | 0,3783138 | -5,89046 |
| SHP2 | -0,10227 | -0,044912 | -1,40168 | 0,186694 | 0,5328811 | -6,52381 |
| IRS2 | -0,10313 | 0,497198 | -1,09538 | 0,295144 | 0,6600066 | -6,8885 |
| ASNS | -0,10411 | -0,022748 | -1,56679 | 0,143505 | 0,4944847 | -6,30062 |
| Rictor_pT1 | -0,10416 | 0,247853 | -1,91373 | 0,08015 | 0,362617 | -5,78198 |
| FoxO3a_pS | -0,10503 | 0,075277 | -1,65761 | 0,12365 | 0,4628801 | -6,17091 |
| BCL2A1 | -0,10513 | -0,035601 | -1,09228 | 0,296452 | 0,6600066 | -6,89183 |
| VEGFR-2 | -0,10621 | 0,135944 | -1,05567 | 0,312199 | 0,6761695 | -6,93056 |
| XPF | -0,10753 | 0,014147 | -2,02401 | 0,06615 | 0,3277096 | -5,60532 |
| Gys | -0,10778 | 0,329948 | -1,39052 | 0,189973 | 0,535152 | -6,53827 |
| Chk1_pS34 | -0,1098 | 0,094896 | -1,65826 | 0,123518 | 0,4628801 | -6,16997 |
| DRP1 | -0,11062 | 0,073839 | -2,29345 | 0,040963 | 0,2635093 | -5,15494 |
| PKA-a | -0,11139 | -0,147499 | -1,34574 | 0,203606 | 0,5484457 | -6,59543 |
| Myosin-IIa_ | -0,11771 | -0,007141 | -1,00337 | 0,335763 | 0,6957393 | -6,98398 |
| GSK-3a-b | -0,11995 | 0,145096 | -2,19071 | 0,04925 | 0,2756727 | -5,32951 |
| ERCC1 | -0,12051 | -0,05099 | -2,19778 | 0,048632 | 0,2756727 | -5,3176 |
| DVL3 | -0,12073 | 0,097953 | -2,36785 | 0,035812 | 0,2450007 | -5,02661 |
| Atg5 | -0,12288 | 0,237612 | -1,55304 | 0,146739 | 0,4944847 | -6,31984 |
| Akt2 | -0,12584 | 0,225659 | -2,02706 | 0,065797 | 0,3277096 | -5,60036 |
| p90RSK_pT | -0,12803 | 0,127706 | -0,7199 | 0,485567 | 0,838916 | -7,23208 |
| B-Raf_pS4 | -0,12818 | 0,108475 | -2,89989 | 0,013501 | 0,1452348 | -4,07489 |
| MCT4 | -0,12951 | 0,021705 | -1,97009 | 0,072686 | 0,3442579 | -5,69232 |
| Rb_pS807_ | -0,13049 | 0,274772 | -2,54875 | 0,025761 | 0,1982695 | -4,70887 |
| B-Raf | -0,13096 | 0,001943 | -1,52347 | 0,153913 | 0,502496 | -6,36081 |
| PKM2 | -0,1319 | -0,316375 | -2,05104 | 0,063084 | 0,3251399 | -5,56129 |
| Raptor | -0,13285 | 0,029375 | -2,05618 | 0,062517 | 0,3251399 | -5,55289 |
| ACVRL1 | -0,133 | -2,20E-04 | -2,50766 | 0,027771 | 0,2099881 | -4,7817 |
| PKCa | -0,13419 | 0,324233 | -2,56682 | 0,024923 | 0,1982695 | -4,67674 |
| RSK | -0,13516 | -0,552241 | -2,26519 | 0,0431 | 0,2645265 | -5,20328 |
| ZAP-70 | -0,136 | -0,001713 | -1,22999 | 0,242596 | 0,5808818 | -6,73672 |
| ACSL1 | -0,13646 | 0,059466 | -2,56373 | 0,025065 | 0,1982695 | -4,68226 |
| p70-S6K1 | -0,13687 | 0,134162 | -1,90928 | 0,080769 | 0,362617 | -5,789 |
| MDM2_pS | -0,13705 | 0,002366 | -1,85463 | 0,088727 | 0,3783138 | -5,87452 |
| B7-H4 | -0,13796 | -0,039993 | -1,61214 | 0,133268 | 0,4786537 | -6,23644 |
| ER-a | -0,14017 | -0,077341 | -1,63761 | 0,127802 | 0,4628801 | -6,19987 |
| GSK-3a-b_p | -0,14136 | 0,350958 | -1,77147 | 0,10221 | 0,4078921 | -6,00197 |
| HER3_pY12 | -0,14265 | 3,22E-04 | -2,71721 | 0,018909 | 0,1733991 | -4,40703 |
| NF-kB-p65_ | -0,14574 | 0,118054 | -2,76059 | 0,017457 | 0,1656751 | -4,32855 |
| RPA32_pS4 | -0,14623 | 0,003 | -2,2038 | 0,048112 | 0,2756727 | -5,30745 |
| Mitofusin-2 | -0,14717 | 0,07772 | -1,53926 | 0,150045 | 0,5013137 | -6,339 |
| c-IAP2 | -0,14783 | -0,079066 | -2,23909 | 0,045167 | 0,2666706 | -5,24771 |

| | | | | | | |
|---|---|---|---|---|---|---|
| PAICS | -0,30731 | -0,15572 | -5,0976 | 2,75E-04 | 0,0021143 | -0,48504 |
| DVL3 | -0,31665 | 0,097953 | -6,21043 | 4,80E-05 | 5,31E-04 | 1,34919 |
| mTOR | -0,31753 | 0,02475 | -5,68632 | 1,07E-04 | 0,0010014 | 0,506244 |
| Akt1 | -0,32109 | -0,036534 | -1,77286 | 0,101971 | 0,2044161 | -6,41284 |
| Stat3_pY705 | -0,3245 | 0,23668 | -3,64101 | 0,003457 | 0,0163746 | -3,10895 |
| ERRalpha | -0,32454 | -0,060608 | -5,79938 | 8,97E-05 | 8,78E-04 | 0,691262 |
| eEF2K | -0,33215 | 0,045657 | -3,22723 | 0,007378 | 0,0308748 | -3,87829 |
| MLKL | -0,33376 | -0,032043 | -2,70875 | 0,019206 | 0,0587069 | -4,83147 |
| Wee1 | -0,33378 | -0,01496 | -2,47799 | 0,029316 | 0,0801925 | -5,24429 |
| MITF | -0,33833 | 0,104632 | -6,35878 | 3,86E-05 | 4,86E-04 | 1,580981 |
| RPA32 | -0,34532 | -0,242082 | -5,47934 | 1,48E-04 | 0,0012646 | 0,163012 |
| Jak2 | -0,34611 | 0,117213 | -3,97961 | 0,001877 | 0,0102422 | -2,48265 |
| VAV1 | -0,35286 | -0,053778 | -3,09766 | 0,00937 | 0,0363294 | -4,11854 |
| p38-MAPK | -0,35463 | 0,104174 | -5,96883 | 6,91E-05 | 7,09E-04 | 0,965284 |
| PTEN | -0,35736 | 0,327711 | -5,061 | 2,92E-04 | 0,0021683 | -0,54816 |
| GCN5L2 | -0,37165 | -0,152105 | -7,11823 | 1,31E-05 | 2,10E-04 | 2,721295 |
| Tyro3 | -0,38247 | 0,007668 | -6,64415 | 2,55E-05 | 3,55E-04 | 2,018466 |
| p70-S6K1 | -0,39486 | 0,134162 | -5,50817 | 1,41E-04 | 0,0012445 | 0,211165 |
| Src_pY527 | -0,39616 | 0,028145 | -5,38593 | 1,72E-04 | 0,0014253 | 0,006224 |
| Rictor | -0,39956 | 0,284828 | -6,17974 | 5,03E-05 | 5,42E-04 | 1,300876 |
| Heregulin | -0,4105 | 0,529459 | -5,11474 | 2,67E-04 | 0,002093 | -0,45554 |
| PLC-gamma2_pY759 | -0,42675 | -0,048337 | -4,85309 | 4,12E-04 | 0,0029141 | -0,91 |
| Akt | -0,4348 | -0,027398 | -6,82411 | 1,98E-05 | 2,88E-04 | 2,288736 |
| Akt2_pS474 | -0,44435 | 0,456442 | -5,5679 | 1,29E-04 | 0,0011693 | 0,310577 |
| Mcl-1 | -0,44681 | 0,126503 | -5,88719 | 7,83E-05 | 7,85E-04 | 0,833756 |
| PKA-a | -0,45301 | -0,147499 | -5,47293 | 1,50E-04 | 0,0012646 | 0,152283 |
| PHLPP | -0,45585 | 0,138066 | -7,96525 | 4,30E-06 | 8,25E-05 | 3,905764 |
| Sox2 | -0,46061 | 0,345513 | -7,94658 | 4,40E-06 | 8,25E-05 | 3,880591 |
| GAPDH | -0,46402 | -0,093054 | -6,51693 | 3,06E-05 | 4,13E-04 | 1,824783 |
| Akt1_pS473 | -0,47762 | 0,132057 | -7,75171 | 5,65E-06 | 1,02E-04 | 3,615459 |
| Stat5a | -0,50318 | 0,376954 | -6,37435 | 3,77E-05 | 4,86E-04 | 1,605132 |
| RRM2 | -0,52111 | 0,068136 | -5,78366 | 9,19E-05 | 8,80E-04 | 0,665629 |
| b-Actin | -0,5242 | -0,123301 | -5,56055 | 1,30E-04 | 0,0011693 | 0,298364 |
| Shc_pY317 | -0,5344 | 0,048967 | -6,23445 | 4,64E-05 | 5,26E-04 | 1,386934 |
| Stat3 | -0,54545 | 0,455546 | -4,81689 | 4,38E-04 | 0,003047 | -0,97355 |
| JNK_pT183_Y185 | -0,5871 | 0,06726 | -6,10202 | 5,65E-05 | 5,94E-04 | 1,177915 |
| FAK_pY397 | -0,655 | 0,127547 | -5,15556 | 2,50E-04 | 0,0019937 | -0,38541 |
| MMP14 | -0,68302 | -0,048973 | -10,5751 | 2,21E-07 | 5,96E-06 | 7,057217 |
| c-Met_pY1234_Y1235 | -0,6895 | -0,612021 | -6,81436 | 2,00E-05 | 2,88E-04 | 2,274211 |
| Src_pY416 | -0,71654 | 0,071301 | -6,95875 | 1,64E-05 | 2,52E-04 | 2,488164 |
| HER2_pY1248 | -0,72084 | 0,074269 | -5,08225 | 2,82E-04 | 0,0021304 | -0,5115 |
| HES1 | -0,72648 | -0,136402 | -2,76109 | 0,017441 | 0,0548696 | -4,73656 |
| FRS2-a_pY196 | -0,79217 | 0,09159 | -6,24453 | 4,57E-05 | 5,26E-04 | 1,402745 |
| cdc2_pY15 | -0,80946 | -0,23913 | -11,0781 | 1,34E-07 | 3,85E-06 | 7,591198 |
| Coup-TFII | -0,84072 | 0,178384 | -11,8459 | 6,44E-08 | 2,14E-06 | 8,367896 |
| Chk1 | -0,85974 | -0,197309 | -15,041 | 4,50E-09 | 2,16E-07 | 11,18095 |
| DUSP6 | -0,90276 | 0,129979 | -15,1602 | 4,12E-09 | 2,16E-07 | 11,27484 |
| SHP-2_pY542 | -0,91189 | 0,322221 | -7,1277 | 1,30E-05 | 2,10E-04 | 2,735042 |
| Rad51 | -0,92203 | 0,041121 | -8,78008 | 1,59E-06 | 3,26E-05 | 4,965088 |
| Connexin-43 | -1,00379 | -0,052007 | -18,3876 | 4,58E-10 | 3,78E-08 | 13,57667 |
| Pyk2_pY402 | -1,02345 | 0,131592 | -8,81671 | 1,52E-06 | 3,26E-05 | 5,010972 |
| Hif-1-alpha | -1,04734 | 0,203091 | -9,8255 | 4,87E-07 | 1,10E-05 | 6,220818 |
| Snail | -1,38606 | 0,389254 | -18,747 | 3,67E-10 | 3,78E-08 | 13,80784 |

| | | | | | | |
|---|---|---|---|---|---|---|
| PKC-b-II_pS | -0,14893 | 0,022917 | -1,32892 | 0,208933 | 0,5484457 | -6,61656 |
| Rictor | -0,1544 | 0,284828 | -2,38797 | 0,03453 | 0,2450007 | -4,99167 |
| EMA | -0,15474 | 0,046986 | -0,71361 | 0,489307 | 0,838916 | -7,23675 |
| IDO | -0,15487 | -0,045778 | -1,33572 | 0,206767 | 0,5484457 | -6,60804 |
| ATM | -0,15562 | 0,084602 | -1,48376 | 0,164015 | 0,509804 | -6,41498 |
| PHLPP | -0,15905 | 0,138066 | -2,77914 | 0,01687 | 0,1656751 | -4,29492 |
| PAK4 | -0,16293 | 0,010678 | -3,27263 | 0,006787 | 0,0886378 | -3,38785 |
| Akt | -0,16642 | -0,027398 | -2,61191 | 0,022946 | 0,1939187 | -4,5963 |
| ATRX | -0,16836 | 0,050761 | -1,53137 | 0,151966 | 0,502496 | -6,34991 |
| GCLC | -0,1712 | -0,02283 | -2,37336 | 0,035457 | 0,2450007 | -5,01705 |
| FOXO3 | -0,1723 | 0,039835 | -1,67056 | 0,121025 | 0,4616099 | -6,15204 |
| PLC-gamm | -0,17834 | -0,048337 | -2,02811 | 0,065676 | 0,3277096 | -5,59866 |
| Erk5 | -0,18134 | -0,046955 | -2,39804 | 0,033905 | 0,2450007 | -4,97413 |
| VASP | -0,18221 | 0,087681 | -2,68052 | 0,020229 | 0,1816439 | -4,47319 |
| Gab2 | -0,18294 | 0,045631 | -2,14896 | 0,053053 | 0,2931512 | -5,3995 |
| MLKL | -0,18411 | -0,032043 | -1,4942 | 0,161307 | 0,509804 | -6,40084 |
| 53BP1 | -0,18424 | 0,410082 | -2,10585 | 0,057269 | 0,3105905 | -5,47115 |
| Akt1 | -0,18804 | -0,036534 | -1,03822 | 0,319917 | 0,676246 | -6,94863 |
| GSK-3B | -0,18819 | 0,32798 | -1,83018 | 0,092514 | 0,3801027 | -5,91233 |
| Akt1_pS47 | -0,18971 | 0,132057 | -3,07886 | 0,009701 | 0,1226175 | -3,74609 |
| p53 | -0,1926 | 0,265969 | -2,25128 | 0,04419 | 0,2645265 | -5,22698 |
| PRAS40_pT | -0,19377 | 0,125751 | -3,76293 | 0,002772 | 0,0522848 | -2,47979 |
| FRS2-a_pY | -0,19741 | 0,09159 | -1,55616 | 0,146001 | 0,4944847 | -6,3155 |
| p70-S6K_p | -0,19983 | 0,269091 | -1,59917 | 0,136127 | 0,4848836 | -6,25491 |
| GAPDH | -0,20186 | -0,093054 | -2,83502 | 0,015218 | 0,1561705 | -4,19331 |
| PDK1 | -0,20918 | -0,003713 | -2,3071 | 0,039968 | 0,2610029 | -5,13151 |
| PTPN12 | -0,2106 | 0,448663 | -3,71753 | 0,003009 | 0,0522848 | -2,56363 |
| 4E-BP1_pT | -0,21502 | -0,149083 | -2,76025 | 0,017468 | 0,1656751 | -4,32919 |
| Slfn11 | -0,21667 | -0,015227 | -1,48223 | 0,164415 | 0,509804 | -6,41704 |
| Akt2_pS47 | -0,21976 | 0,456442 | -2,75364 | 0,017682 | 0,1656751 | -4,34116 |
| JNK_pT183 | -0,22435 | 0,06726 | -2,33181 | 0,038225 | 0,2534579 | -5,08896 |
| Rad23A | -0,22733 | -0,014213 | -1,28859 | 0,222167 | 0,5534901 | -6,66638 |
| Src_pY416 | -0,22829 | 0,071301 | -2,21701 | 0,046989 | 0,2736773 | -5,28513 |
| Akt_pT308 | -0,23395 | 0,127704 | -3,38497 | 0,00552 | 0,0793074 | -3,17963 |
| FAK_pY397 | -0,23423 | 0,127547 | -1,84365 | 0,090409 | 0,3783138 | -5,89152 |
| Pyk2_pY40 | -0,23779 | 0,131592 | -2,04848 | 0,063368 | 0,3251399 | -5,56547 |
| Akt_pS473 | -0,24707 | 0,399518 | -2,47815 | 0,029308 | 0,2177851 | -4,83377 |
| Gys_pS641 | -0,24799 | 0,273752 | -3,0151 | 0,010914 | 0,1237839 | -3,86353 |
| ULK1_pS75 | -0,25007 | 0,319938 | -4,66656 | 5,65E-04 | 0,0203077 | -0,84296 |
| mTOR | -0,26135 | 0,02475 | -4,68024 | 5,52E-04 | 0,0203077 | -0,8188 |
| Src_pY527 | -0,26204 | 0,028145 | -3,56249 | 0,003989 | 0,0614001 | -2,85053 |
| IGF1R_pY1 | -0,26236 | -0,141052 | -4,24708 | 0,001169 | 0,0296286 | -1,59372 |
| c-Met_pY1 | -0,26245 | -0,612021 | -2,59384 | 0,02372 | 0,1965997 | -4,6286 |
| PTEN | -0,26701 | 0,327711 | -3,78148 | 0,00268 | 0,0522848 | -2,44556 |
| mTOR_pS2 | -0,2714 | 0,087497 | -3,75756 | 0,002799 | 0,0522848 | -2,48969 |
| WIPI2 | -0,27549 | -0,02897 | -2,09508 | 0,058371 | 0,3105905 | -5,48895 |
| SHP-2_pY5 | -0,29038 | 0,322221 | -2,26975 | 0,042748 | 0,2645265 | -5,19549 |
| Shc_pY317 | -0,29926 | 0,048967 | -3,49128 | 0,004543 | 0,0675184 | -2,9825 |
| Stat5a | -0,33847 | 0,376954 | -4,28781 | 0,001088 | 0,0293093 | -1,52003 |
| HER2_pY12 | -0,42823 | 0,074269 | -3,01925 | 0,01083 | 0,1237839 | -3,8559 |
| S6_pS240_ | -0,43847 | 0,524502 | -3,69037 | 0,003161 | 0,0523966 | -2,61383 |
| S6_pS235_ | -0,47719 | 0,525909 | -2,61345 | 0,022882 | 0,1939187 | -4,59355 |
| JNK2 | -0,55213 | 0,368288 | -7,08148 | 1,38E-05 | 8,51E-04 | 3,03862 |

**Table S8:** Cell cycle distribution of the *MMTV-R26^Met^* MGT cell lines - Statistical analysis.

| | | MGT2 | | | | | MGT4 | | | | | MGT9 | | | | | MGT11 | | | | | MGT13 | | | | |
|---|---|---|---|---|---|---|---|---|---|---|---|---|---|---|---|---|---|---|---|---|---|---|---|---|---|---|
| | | G0 | G1 | S | G2 | M | G0 | G1 | S | G2 | M | G0 | G1 | S | G2 | M | G0 | G1 | S | G2 | M | G0 | G1 | S | G2 | M |
| MGT7 | G0 | *** | | | | | *** | | | | | *** | | | | | *** | | | | | *** | | | | |
| | G1 | | *** | | | | | *** | | | | | *** | | | | | *** | | | | | *** | | | |
| | S | | | * | | | | | *** | | | | | *** | | | | | *** | | | | | *** | | |
| | G2 | | | | ns | | | | | *** | | | | | *** | | | | | *** | | | | | *** | |
| | M | | | | | ns | | | | | ns | | | | | ns | | | | | ns | | | | | ns |
| MGT2 | G0 | | | | | | *** | | | | | *** | | | | | *** | | | | | *** | | | | |
| | G1 | | | | | | | * | | | | | *** | | | | | ns | | | | | ns | | | |
| | S | | | | | | | | *** | | | | | *** | | | | | *** | | | | | ns | | |
| | G2 | | | | | | | | | *** | | | | | ns | | | | | *** | | | | | *** | |
| | M | | | | | | | | | | ns | | | | | ns | | | | | ns | | | | | ns |
| MGT4 | G0 | | | | | | | | | | | ns | | | | | ns | | | | | * | | | | |
| | G1 | | | | | | | | | | | | ns | | | | | ns | | | | | *** | | | |
| | S | | | | | | | | | | | | | ns | | | | | * | | | | | ns | | |
| | G2 | | | | | | | | | | | | | | * | | | | | ns | | | | | ** | |
| | M | | | | | | | | | | | | | | | ns | | | | | ns | | | | | ns |
| MGT9 | G0 | | | | | | | | | | | | | | | | ns | | | | | ** | | | | |
| | G1 | | | | | | | | | | | | | | | | | *** | | | | | *** | | | |
| | S | | | | | | | | | | | | | | | | | | ns | | | | | ** | | |
| | G2 | | | | | | | | | | | | | | | | | | | ns | | | | | *** | |
| | M | | | | | | | | | | | | | | | | | | | | ns | | | | | ns |
| MGT11 | G0 | | | | | | | | | | | | | | | | | | | | | *** | | | | |
| | G1 | | | | | | | | | | | | | | | | | | | | | | * | | | |
| | S | | | | | | | | | | | | | | | | | | | | | | | *** | | |
| | G2 | | | | | | | | | | | | | | | | | | | | | | | | *** | |
| | M | | | | | | | | | | | | | | | | | | | | | | | | | ns |
| MGT13 | G0 | | | | | | | | | | | | | | | | | | | | | | | | | |
| | G1 | | | | | | | | | | | | | | | | | | | | | | | | | |
| | S | | | | | | | | | | | | | | | | | | | | | | | | | |
| | G2 | | | | | | | | | | | | | | | | | | | | | | | | | |
| | M | | | | | | | | | | | | | | | | | | | | | | | | | |

**Table S9:** In vitro tumorigenic capacity of the *MMTV-R26^Met^* cell lines, as determined by the tumour sphere assay - Statistical analysis.

| | | Passage 1 | | | | | | | | Passage 3 | | | |
|---|---|---|---|---|---|---|---|---|---|---|---|---|---|
| | | MGT2 | | MGT4 | | MGT11 | | MGT13 | | MGT11 | | MGT13 | |
| | | 50-100µm | >100 µm | 50-100µm | >100 µm | 50-100µm | >100 µm | 50-100µm | >100 µm | 50-100µm | >100 µm | 50-100µm | >100 µm |
| **MGT7** | 50-100µm | 0.034 (*) | | 0.086 (ns) | | 0.005 (**) | | 0.003 (**) | | | | | |
| | >100 µm | | 0.116 (ns) | | 0.116 (ns) | | <0.0001 (***) | | 0.007 (**) | | | | |
| **MGT2** | 50-100µm | | | 0.113 (ns) | | 0.006 (**) | | 0.003 (**) | | | | | |
| | >100 µm | | | | >0.999 (ns) | | <0.0001 (***) | | 0.008 (**) | | | | |
| **MGT4** | 50-100µm | | | | | 0.055 (ns) | | 0.008 (**) | | 0.798 (ns) | | 0.032 (*) | |
| | >100 µm | | | | | | <0.0001 (***) | | 0.008 (**) | | 0.009 (**) | | 0.021 (*) |
| **MGT11** | 50-100µm | | | | | | | 0.039 (*) | | | | 0.019 (*) | |
| | >100 µm | | | | | | | | 0.036 (*) | | | | 0.198 (ns) |

**Table S10 :** Migrating capacity of the *MMTV-R26*$^{Met}$ cell lines - Statistical analysis.

| | MGT7 | MGT2 | MGT4 | MGT9 | MGT11 | MGT13 |
|---|---|---|---|---|---|---|
| **MGT7** | | 0.2249 (ns) | 0.0178 (*) | 0.0002 (***) | <0.0001 (***) | <0.0001 (***) |
| **MGT2** | | | 0.6052 (ns) | 0.0139 (*) | 0.0053 (**) | <0.0001 (***) |
| **MGT4** | | | | 0.4207 (ns) | 0.2363 (ns) | <0.0001 (***) |
| **MGT9** | | | | | 0.9987 (ns) | <0.0001 (***) |
| **MGT11** | | | | | | 0.0002 (***) |
| **MGT13** | | | | | | |

**Table S11:** Cell cycle distribution of MGT11 treated cells - Statistical analysis.

| | | A11 | | | | | Adav | | | | | A11 + Adav | | | | |
|---|---|---|---|---|---|---|---|---|---|---|---|---|---|---|---|---|
| | | G0 | G1 | S | G2 | M | G0 | G1 | S | G2 | M | G0 | G1 | S | G2 | M |
| **no** | G0 | ns | | | | | ns | | | | | ns | | | | |
| | G1 | | ns | | | | | ns | | | | | *** | | | |
| | S | | | ns | | | | | ns | | | | | ns | | |
| | G2 | | | | ns | | | | | *** | | | | | *** | |
| | M | | | | | ns | | | | | ns | | | | | ns |
| **A11** | G0 | | | | | | ns | | | | | ns | | | | |
| | G1 | | | | | | | *** | | | | | * | | | |
| | S | | | | | | | | ns | | | | | ns | | |
| | G2 | | | | | | | | | *** | | | | | *** | |
| | M | | | | | | | | | | ns | | | | | ns |
| **Adav** | G0 | | | | | | | | | | | ns | | | | |
| | G1 | | | | | | | | | | | | ns | | | |
| | S | | | | | | | | | | | | | ns | | |
| | G2 | | | | | | | | | | | | | | ns | |
| | M | | | | | | | | | | | | | | | ns |
| **A11 + Adav** | G0 | | | | | | | | | | | | | | | |
| | G1 | | | | | | | | | | | | | | | |
| | S | | | | | | | | | | | | | | | |
| | G2 | | | | | | | | | | | | | | | |
| | M | | | | | | | | | | | | | | | |

**Table S12:** Antibodies used in the study.

| Antibody | Company | Reference | Dilution | Used for | TritonX-100 (%) for IF |
|---|---|---|---|---|---|
| Estrogen receptor alpha (ERα) | Santa Cruz | sc-8005 | 1:100 | IHC | 0.5 % |
| Progesteron receptor (PR) | Santa Cruz | sc-810 | 1:200 | IHC | 0.5 % |
| Human EGFR2 (HER2) | Cell Signaling | 2165 | 1:200 | IHC | 0.2 % |
| MET | Santa Cruz | sc-10 | 1:50 | IHC | 0.2 % |
| Ki67 | Cell Signaling | 9129 | 1:400 | IHC | 0.5 % |
| Ki67 | Dako | M7249 | 1:100 | IF | 0.1 % |
| Krt14 | Covalab | Mab71720 | 1:500 | IF | 0.5 % |
| Krt18 | Covalab | Mab20042 | 1:500 | IF | 0.5 % |
| human MET | Assay Design | 905-076 | 1:150 | IF | 0.5 % |
| $pY_{1234/35}$MET | Cell Signaling | 3126L | 1:50 | IF | 0.5 % |
| pH3 (S10) | Millipore | 06-570 | 1 :500 | IF | 0.3 % |
| alpha-Tubulin | Sigma | T5168 | 1 :5000 | IF | 0.3 % |
| Vimentin | Abcam | Ab-8979 | 1:100 | IF | 0.5 % |
| $pS_{139}$H2AX (γH2AX) | Cell Signaling | 9718 | 1 : 400 | IF, WB | 0.2 % |
| Actin | Sigma | A3853 | 1:6000 | WB | |
| ATM | Cell Signaling | 2873 | 1:1000 | WB | |
| $pS_{1987}$ATM | Invitrogen | PA5-37346 | 1:1000 | WB | |
| ATR | Cell Signaling | 13934 | 1:1000 | WB | |
| $pS_{428}$ATR | Cell Signaling | 2853 | 1:1000 | WB | |
| $pS_{473}$AKT | Cell Signaling | 9271 | 1:2000 | WB | |
| BCL-XL | Transduction Laboratory | B22620 | 1:500 | WB | |
| BIM | Santa Cruz | sc-11425 | 1:1000 | WB | |
| Cleaved-Caspase 3 | Cell Signaling | 9661 | 1:1000 | WB | |
| CDC2 (CDK1) | Cell Signaling | 28493 | 1:20000 | WB | |
| $pY_{15}$CDC2(CDK1) | Cell Signaling | 4539 | 1:1000 | WB | |
| $pT_{202}/Y_{204}$ERKs | Cell Signaling | 9106 | 1:10000 | WB | |
| $pY_{627}$GAB1 | Cell Signaling | 3231 | 1:2000 | WB | |
| MCL1 | Santa Cruz | sc-819 | 1:1000 | WB | |
| $pS_{217/221}$MEK | Cell Signaling | 9121 | 1:1000 | WB | |
| mouse MET | Santa Cruz | sc-8057 | 1:500 | WB | |
| $MET^{25H2}$ | Cell Signaling | 3127 | 1:1000 | WB | |
| human MET | Santa Cruz | sc-161 | 1:1000 | WB | |

| | | | | |
|---|---|---|---|---|
| $pY_{1234/35}MET$ | Cell Signaling | 3126 | 1:2000 | WB |
| PARP | Cell Signaling | 9546S | 1:2000 | WB |
| P53 | Novocastra | CM5 | 1:1000 | WB |
| $pS_{15}P53$ | Cell Signaling | 9284 | 1:1000 | WB |
| RB | Cell Signaling | 9313 | 1:1000 | WB |
| $pS_{795}RB$ | Abcam | Ab47474 | 1:1000 | WB |
| RPA32 (B-4) | Santa Cruz | sc-271578 | 1:200 | WB |
| $pS_{33}RPA32$ | Bethyl | A300-246A | 1 :5000 | WB |
| RRM2 | abbexa | Abx004031 | 1 :5000 | WB |
| $pY_{705}STAT3$ | Cell Signaling | 9145S | 1:2000 | WB |
| XIAP | Transduction laboratory | 610716 | 1:3000 | WB |
| Goat anti-rabbit IgG-peroxidase | Jackson Immuno Research | 115-035-144 | 1:4000 | WB |
| Goat anti-mouse IgG-peroxidase | Jackson Immuno Research | 115-035-146 | 1:4000 | WB |
| CD24-BV421 (clone M1/29) | Biolegend | 101826 | 1:400 | FACS |
| CD29-FITC (clone HMb1-1) | eBioscience | 11-0291-82 | 1:400 | FACS |
| CD61-biotin (clone 2C9.G3) | eBioscience | 13-0611-81 | 1:400 | FACS |
| anti-Ki67-APC (clone SolA15) | eBioscience | 17-5698-82 | 1:200 | FACS |
| Streptavidin, APC-AF750 | Invitrogen | SA1027 | 1:200 | FACS |
| Annexin V | eBioscience | 88-8005-72 | 1:20 | FACS |

**Table S13:** Drugs used for cell viability assays, with the indicated targets and the concentrations used.

| Drug | Company | Target | Concentration (µM) |
|---|---|---|---|
| 3-Methyladdenine (3MA) | TargetMol | Autophagy inhibitor | 1mM |
| 5-Fluorouracil (5-FU) | Targetmol | Thymidylate synthase (nucleotide synthesis) | 0.4, 2, 10 |
| A-1155463 | Selleckchem/Targetmol | Bcl-xL | 0.3, 1, 3, 10 |
| ABT-737 | Selleckchem | Bcl-2,Bcl-xL,Bcl-w | 1, 3, 10 |
| ABT-199 | Selleckchem | Bcl-2 | 1, 3, 10 |
| Adavosertib | Selleckchem | Wee1 | 1, 3, 10 |
| Adavosertib(MK-1775) | Targetmol | Wee1 | 1, 3, 10 |
| AZD6738 | Targetmol | ATR | 1, 3, 10 |
| CB-839 | Selleckchem | Glutaminase | 1, 3, 10 |
| Cisplatin | MedChem Express | DNA replication | 1, 3, 10 |
| Docetaxel | Targetmol | Beta-tubulin (microtubule inhibitor) | 0.4, 2, 10 |
| Doxorubicin | Targetmol | intercalating agent | 0.04, 0.2, 1 |
| Erastin | TargetMol | Ferroptosis inducer | 0,5 - 1 |
| Ferrostatin-1 | TargetMol | Ferroptosis inhibitor | 10, 20, 50, 100 |
| Gefitinib | Calbiochem | EGFR | 1, 3, 10 |
| JNJ-7706621 | Selleckchem | pan-Cdk, Aurora A/B | 1, 3, 10 |
| LY294002 | Calbiochem | PI3K | 1, 3, 10 |
| Necrostatin-1 | TargetMol | Necroptosis inhibitor | 10, 20, 50, 100 |
| Olaparib | TargetMol | PARP1/PARP2 | 1, 3, 10 |
| PF-431396 | Selleckchem | Pyk2/Fak | 1, 3, 10 |
| PHA-665752 | Tocris Bioscience | Met | 0.3, 1, 3 |
| R547 | Selleckchem/Sigma | Cdk1/2/4 | 1, 3, 10 |
| SB225022 | TargetMol | CXCR antagonist | 1, 3, 10 |
| Selumetinib | Selleckchem | Mek1/2 | 1, 3, 10 |
| Sorafenib | Selleckchem | Raf-1, B-Raf, Vegfr2, Pdgfrβ, Flt3,Kit | 1, 3, 10 |
| Staurosporine | TargetMol | Apotosis inducer | 150nM |
| WEHI-539 | Apexbio | Bcl-xL | 1, 3, 10 |
| Z-VAD-FMK | TargetMol | Apoptosis inhibitor | 10, 20, 50, 100 |

**Table S14:** Oligonucleotides used for RT-qPCR experiments.

| Oligonucleotide (name) | Sequence (Forward) | Sequence (Reverse) |
|---|---|---|
| *Met* EXT(mouse) | GTTCTGCTTGGCAACGAGAGCT | GGAGAATGCACTGTATTGCGTCG |
| *Met* (mouse) | GAATTGTCTGCCATTACAGG | CAGCTCTTACTGTTATTGGCGC |
| *Met* (human) | CTGAAGCCGTTTTATGCAGC | GCCACAGGAAAAACCCAAGTAG |
| *Hgf* | GTCCTGAAGGCTCAGACTTGGT | CCAGCCGTAAATACTGCAAGTGG |
| *B2M* | ACAGTTCCACCCGCCTCACATT | TAGAAAGACCAGTCCTTGCTGAAG |

**Table S15:** Proliferation capacity (mitotic index) of the *MMTV-R26$^{Met}$* cell lines - Statistical analysis.

| | MGT7 | MGT2 | MGT4 | MGT9 | MGT11 | MGT13 |
|---|---|---|---|---|---|---|
| **MGT7** | | 0.549 (ns) | <0.0001 (***) | <0.0001 (***) | <0.0001 (***) | <0.0001 (***) |
| **MGT2** | | | <0.0001 (***) | <0.0001 (***) | <0.0001 (***) | <0.0001 (***) |
| **MGT4** | | | | 0.8339 (ns) | 0.6485 (ns) | 0.9936 (ns) |
| **MGT9** | | | | | 0.9996 (ns) | 0.9824 (ns) |
| **MGT11** | | | | | | 0.9121 (ns) |
| **MGT13** | | | | | | |
